# A comprehensive neuroanatomical survey of the *Drosophila* Lobula Plate Tangential Neurons with predictions for their optic flow sensitivity

**DOI:** 10.1101/2023.10.16.562634

**Authors:** Arthur Zhao, Aljoscha Nern, Sanna Koskela, Marisa Dreher, Mert Erginkaya, Connor W. Laughland, Henrique Ludwigh, Alex Thomson, Judith Hoeller, Ruchi Parekh, Sandro Romani, Davi D. Bock, Eugenia Chiappe, Michael B. Reiser

## Abstract

Flying insects exhibit remarkable navigational abilities controlled by their compact nervous systems. *Optic flow*, the pattern of changes in the visual scene induced by locomotion, is a crucial sensory cue for robust self-motion estimation, especially during rapid flight. Neurons that respond to specific, large-field optic flow patterns have been studied for decades, primarily in large flies, such as houseflies, blowflies, and hover flies. The best-known optic-flow sensitive neurons are the large tangential cells of the dipteran lobula plate, whose visual-motion responses, and to a lesser extent, their morphology, have been explored using single-neuron neurophysiology. Most of these studies have focused on the large, Horizontal and Vertical System neurons, yet the lobula plate houses a much larger set of ‘optic-flow’ sensitive neurons, many of which have been challenging to unambiguously identify or to reliably target for functional studies. Here we report the comprehensive reconstruction and identification of the Lobula Plate Tangential Neurons in an Electron Microscopy (EM) volume of a whole *Drosophila* brain. This catalog of 58 LPT neurons (per brain hemisphere) contains many neurons that are described here for the first time and provides a basis for systematic investigation of the circuitry linking self-motion to locomotion control. Leveraging computational anatomy methods, we estimated the visual motion receptive fields of these neurons and compared their tuning to the visual consequence of body rotations and translational movements. We also matched these neurons, in most cases on a one-for-one basis, to stochastically labeled cells in genetic driver lines, to the mirror-symmetric neurons in the same EM brain volume, and to neurons in an additional EM data set. Using cell matches across data sets, we analyzed the integration of optic flow patterns by neurons downstream of the LPTs and find that most central brain neurons establish sharper selectivity for global optic flow patterns than their input neurons. Furthermore, we found that self-motion information extracted from optic flow is processed in distinct regions of the central brain, pointing to diverse foci for the generation of visual behaviors.

## Introduction

As an animal moves through the world, relative motion between its eyes and the environment induces measurable patterns of visual motion, termed optic flow. The patterns of optic flow are a direct consequence of the animal’s movement, and are also a reliable cue for estimating the self-motion producing them (Gibson, 1950). Flying insects, especially the Diptera—true flies—carry out impressive feats of navigation with compact nervous systems that have been studied for well over 100 years (Ramón y Cajal and Sánchez, 1915). These amazing flyers provide an ideal opportunity to understand how the nervous system measures optic flow and uses it to control navigation. Several important goals towards this understanding are now achievable: describing the set of neurons that measure optic flow, explaining how they become tuned to different patterns of optic flow, and exploring how the circuitry downstream of these visual neurons integrates these optic flow-sensitive neurons into what are presumably behaviorally relevant combinations.

Early electrophysiological recordings (Bishop et al., 1968; Dvorak et al., 1975) and anatomical studies (Pierantoni, 1976; Strausfeld, 1970) in blowflies and houseflies identified several large visual neurons that respond to global motion cues, notably, the horizontal system (HS) and vertical system (VS) neurons. Anatomical surveys in *Calliphora* blowflies found about 50 individually identifiable neurons (including HS and VS cells) that span large regions of the Lobula Plate (herein abbreviated as LOP), a neuropil in the optic lobes (Hausen, 1987). These neurons have been collectively known as Lobula Plate Tangential Cells, often abbreviated as LPTCs, however, for greater consistency with nomenclature of other optic lobe neurons, we refer to this group as the LPT neurons (‘C’ is commonly used in abbreviations of columnar neurons, and only rarely is ‘Cell’ included in the abbreviations of neuron types). Neurophysiological measurements of local directional responses across large fields of view suggested that many of these neurons are tuned to optic flow fields induced by specific rotations or translations (Karmeier et al., 2003; Krapp and Hengstenberg, 1996; Longden et al., 2017). While most studies have been carried out in *Calliphora*, comparative studies found many homologous cells in other fly species (Buschbeck and Strausfeld, 1997; Hengstenberg et al., 1982; Nordström et al., 2008). Most of these efforts relied on electrophysiology to characterize the neuron responses, sometimes supported by histology of filled cells for morphological cell type identification. Correlates between neurons’ shapes and their directional responses had been occasionally noted. The better-studied LPT neurons are supported by decades of research, but due to substantial challenges of routinely targeting individual, identified neurons, only limited data is available for the properties of the majority of LPTs. Further, the morphological and physiological diversity of LPT types could not be completely surveyed using these methods.

Over the last two decades, the fruit fly, *Drosophila melanogaster,* with its many experimental advantages, has become an excellent system to investigate the visual motion circuitry (Currier et al., 2023). The availability of cell-type-specific genetic driver lines and updated methods for neuroanatomy have enabled studies describing the diversity of cells and provided experimental “handles” for functional studies. These tools have enabled the identification of the major directionally selective neurons, the T4 and T5 cells, and powered experiments to uncover many details about their function. The T4 neurons receive retinotopically arranged columnar inputs in the medulla from which they compute local preferred directions of moving bright edges (Behnia et al., 2014; Gruntman et al., 2018; Maisak et al., 2013; Strother et al., 2017). Their preferred directions are shaped by the arrangement of synaptic inputs, that are strongly correlated with the principal orientation of their dendritic arbor (Shinomiya et al., 2019; Takemura et al., 2017, 2013). T5 neurons comprise an analogous motion pathway sensitive to dark edge motion (Fisher et al., 2015; Gruntman et al., 2019; Haag et al., 2017). There are four subtypes of T4 (and T5), each with a distinct dendritic orientation, sensitive to a specific direction of motion (but see (Henning et al., 2022; Zhao et al., 2022)), and each projecting an axon terminal to one of the 4 LOP layers (Shinomiya et al., 2022, 2019). Because of this understanding of the organization of the LOP, the shapes and layer-specific-arborization of LPT neurons can now be used as a basis for predicting their motion responses. Using Golgi staining (Heisenberg et al., 1978; Rajashekhar and Shamprasad, 2004) and genetic labeling (Scott et al., 2002; Wasserman et al., 2015; Wei et al., 2020) several LPT neurons closely resembling those in blowflies have been identified, notably, the HS and VS neurons. By using genetic lines to target recordings to these neurons, the HS and VS cells of *Drosophila* were demonstrated to share many of the functional properties of the homologous neurons recorded in *Calliphora* (Joesch et al., 2008; Schnell et al., 2010).

Despite all of this recent progress, it has been challenging to identify the full diversity of LPT neurons in *Drosophila*, and it remains unclear whether the smaller fly has many of the cells described in *Calliphora*. While stochastic labeling (either Golgi or using genetic methods such as (Nern et al., 2015)) provides an effective way to explore the morphology of single cells, a fundamental limitation of applying these methods to large, uniquely identifiable neurons is the uncertainty of whether observed morphological variability reflects animal-to-animal differences of cells of the same type, or indicates the existence of distinct cell types. In addition, it can be challenging to judge when the sampling of cell types in a group of interest by such methods has reached completeness. One remedy to both issues would be to catalog all the LPT neurons in a single animal. Moreover, a comprehensive inventory of the LPTs, together with a complete anatomical reference for the lobula plate, would provide the necessary foundation for predicting the optic flow sensitivity of these neurons.

Recent EM studies have provided important data on some LPTs in *Drosophila*, emphasizing the HS/VS cells (Boergens et al., 2018; Shinomiya et al., 2022), but neither study used a data set suited for surveying the diversity of these neurons. In this study, we present a multi-year effort to manually identify and reconstruct all the LPT neurons in the right-side lobe of a female *Drosophila melanogaster* (example neuron in Figure 1A), in the Full Adult Female Brain (FAFB) EM volume (Zheng et al., 2018), using the CATMAID environment (Saalfeld et al., 2009). The methods and clarifying details of the FAFB brain hemispheres are provided in Methods: Manual reconstruction in CATMAID. The complete catalog of neurons is reviewed in Figure 3 and 4, many of which are identified in *Drosophila* for the first time (with light microcopy matches for many, see Figure 1B). We match these cells, one-for-one, whenever possible, to their contralateral counterparts in the FAFB volume using the FlyWire consortium’s reconstructions (Dorkenwald et al., 2023) to confirm their identities and to report their predicted neurotransmitters (Eckstein et al., 2020). We then use computational anatomy methods to estimate the receptive fields of these neurons based on detailed knowledge of their dendrites’ morphology and predict their optimal motion sensitivity. The overall methodology is explained in Figure 2. Finally, we match the LPTs to neurons in the Hemibrain connectome (Scheffer et al., 2020) to explore how their central brain connectivity could inform their functional role in decoding body movements (Figures 5 and 6). Connecting motion sensing neurons to motor circuit is an important step towards understanding how optic flow is used to regulate animals’ movement and navigation. This study provides an essential resource for exploring these connections in the increasingly well understood *Drosophila* nervous system.

**Figure 1:**
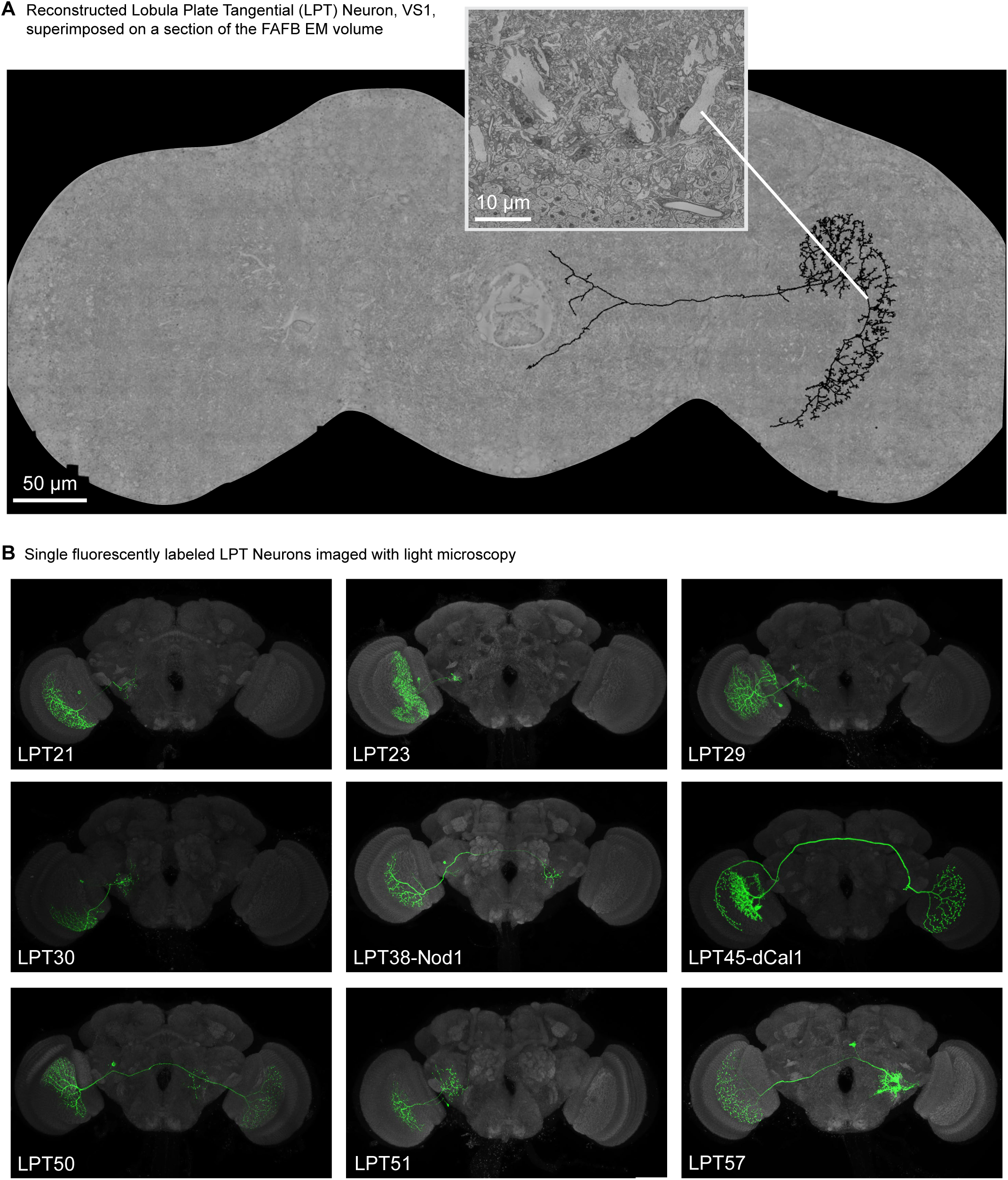
Identification and reconstruction of *Drosophila* LPT neurons in an Electron Microscopy dataset. **A.** The Lobula Plate Tangential Neurons were manually reconstructed on one side of Full Adult Female Brain (FAFB) EM volume (Zheng et al., 2018). The skeleton of one such LPT, the Vertical System number 1, or VS1 neuron, is shown (in black) superimposed on a single slice of the EM volume (Zheng et al., 2018). The data and this image were generated using the CATMAID environment (Saalfeld et al., 2009). The inset shows the very large caliber processes of several VS neurons as well as the much smaller processes of nearby cells, and the somata of neurons in the Lobula Plate cell body rind (bottom of inset). 58 LPT neurons were found on one side of the FAFB brain. The complete inventory of LPT Neurons is detailed Supplementary File 1, and a gallery of their morphologies is in Supplementary File 2. **B.** Examples of light microscopy images of LPT neurons that were used to guide the search, and as a comparison for, reconstructed neurons in the EM volume. Images show manually segmented MulticolorFlpOut (MCFO) labeled LPT neurons (green) together with a neuropile marker (anti-Brp, gray). The names of each cell, listed in the lower left corner, are based on our standardized nomenclature, summarized in the text, for the matching EM-reconstructed LPT neurons.

**Figure 2:**
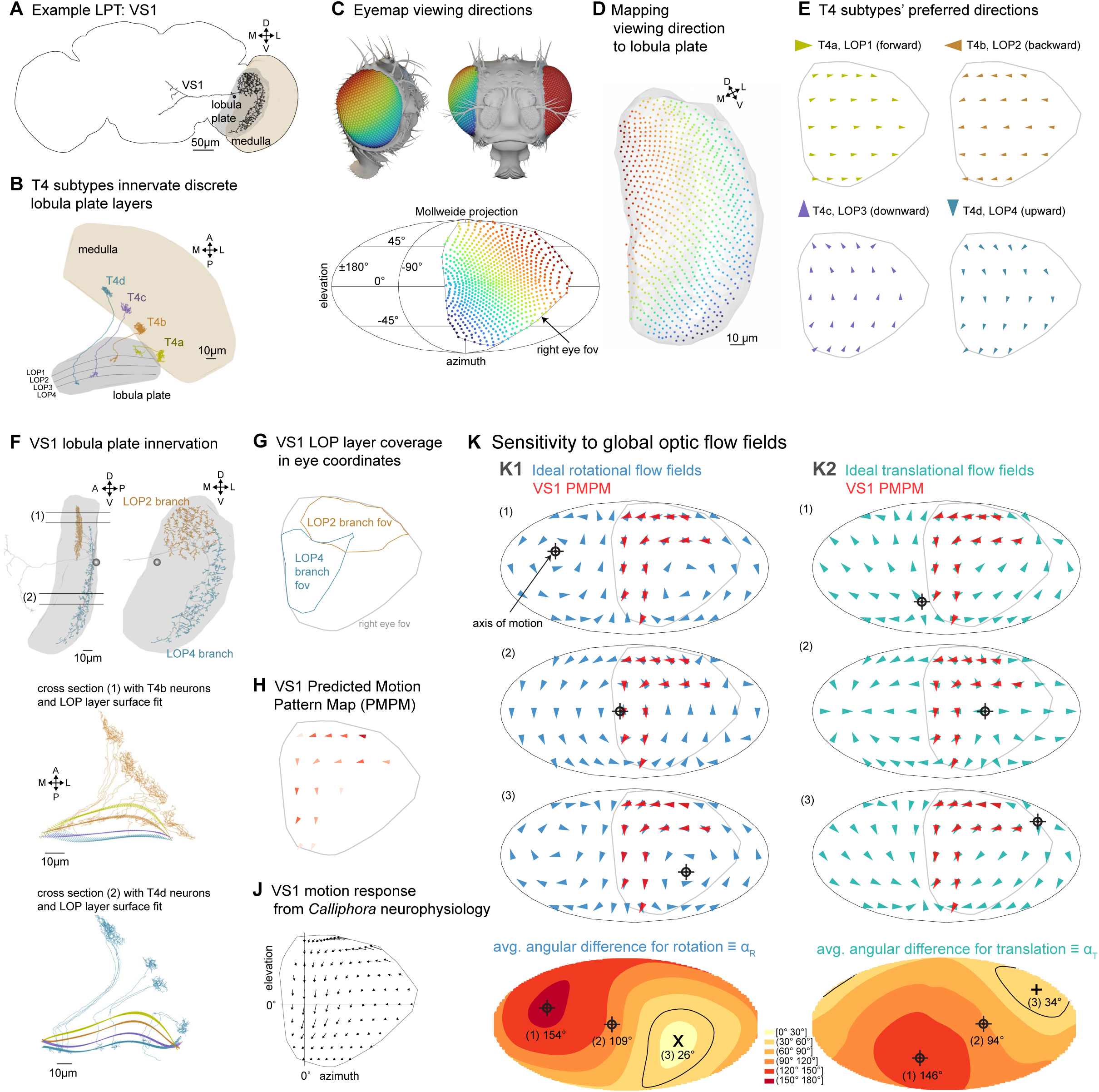
Predicting visual motion response from anatomy. **A.** An EM-reconstructed lobula plate tangential (LPT) neuron, VS1, shown with the outline of the FAFB brain volume and the indicated optic lobe neuropils. **B.** Four subtypes of directionally selective T4 neurons receive input from the proximal layer of the medulla and each project to one of four layers in Lobula Plate (LOP). **C.** Top: a fly head model with lenses color-coded to illustrate retina positions. Bottom: viewing directions of all ommatidia in our reference compound eye (Zhao et al., 2022), presented as a Mollweide 2D projection. **D.** Anatomical positions in LOP mapped to ommatidia directions in (**C**) via set of reconstructed T4 neurons (Figure 2—figure supplement 1A). **E.** Local preferred directions of motion for four T4 subtypes mapped to the eye coordinates, as a Mollweide projection, down-sampled by a factor of 9 (Figure 2—figure supplement 1B,C). **F.** Top: 2 views of the VS1 neuron, whose dendritic branches are colored by the 2 innervated LOP layers, using the same color code as in (**E**). Bellow: two cross sections, corresponding to slices (1) and (2) above, showing several reconstructed T4b and T4d neurons and the 4 LOP layers. **G.** Layer coverage by the VS1 dendrite, displayed in eye coordinates. **H**. Predicated Motion Pattern Map (PMPM) estimated from the dendritic layer coverage and T4 neurons’ preferred directions in (**E**). **J.** Motion response of a *Calliphora* VS1 neuron, measured with electrophysiology (replotted in eye coordinates from (Krapp and Hengstenberg, 1996)). **K1**. Top 3 plots: comparisons of the PMPM to ideal optic flow fields induced by rotations along the 3 indicated axes. Bottom: heat map shows the average angular difference (α_R_) between the PMPM and all sampled ideal flow fields. **K2.** Similar to K1, but for translation (α_T_). The directions with minimal average angular difference for rotational and translational flow fields are indicated by ‘×’ and ‘+’. The black line shows the contour that contains the directions of tested ideal flow fields with the 10% smallest differences.

**Figure 3:**
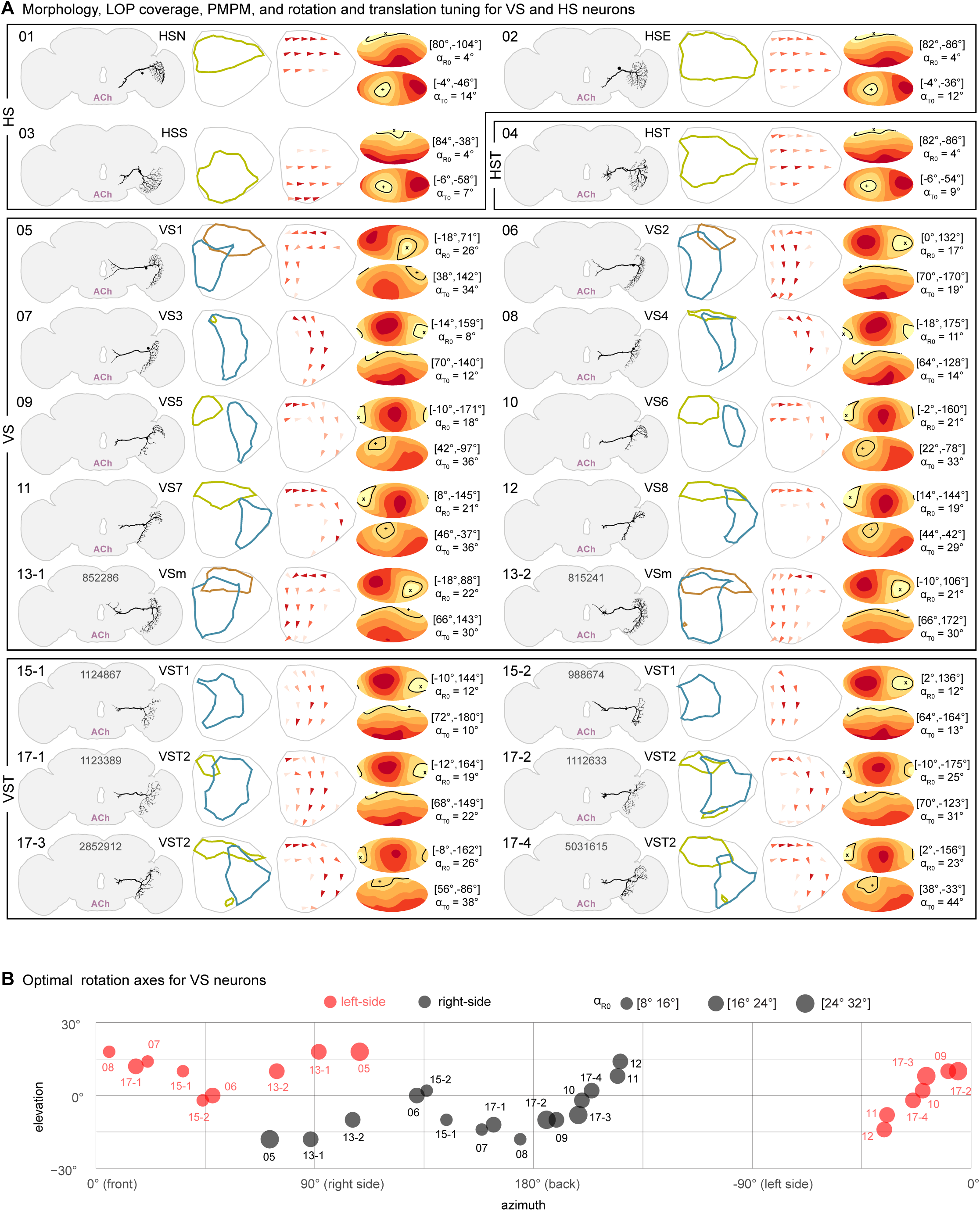
The Horizontal and Vertical System Neurons—morphology, LOP coverage, PMPMs, and optic flow tuning. **A.** The HS and VS neurons reconstructed on one side of the FAFB brain. From left to right: EM reconstruction, LOP layer coverage, color-coded to match Figure 2, PMPM, optic flow tuning as heatmaps of average angular difference (α_R_ for rotation; α_T_ for translation; coordinates [elevation, azimuth] and error value: α_R_ ≡ α_R0_ and α_T_ ≡ α_T0_ for minimal difference). The heatmap coordinate system has positive elevation above the equator, positive azimuth in the right visual field, and [0°,0°] is the front. **B.** Optimal rotation axes for all VS neurons in the right-side optic lobe (in black) and assuming symmetry also for the left-side (in red), shown with a Mercator projection.

**Figure 4:**
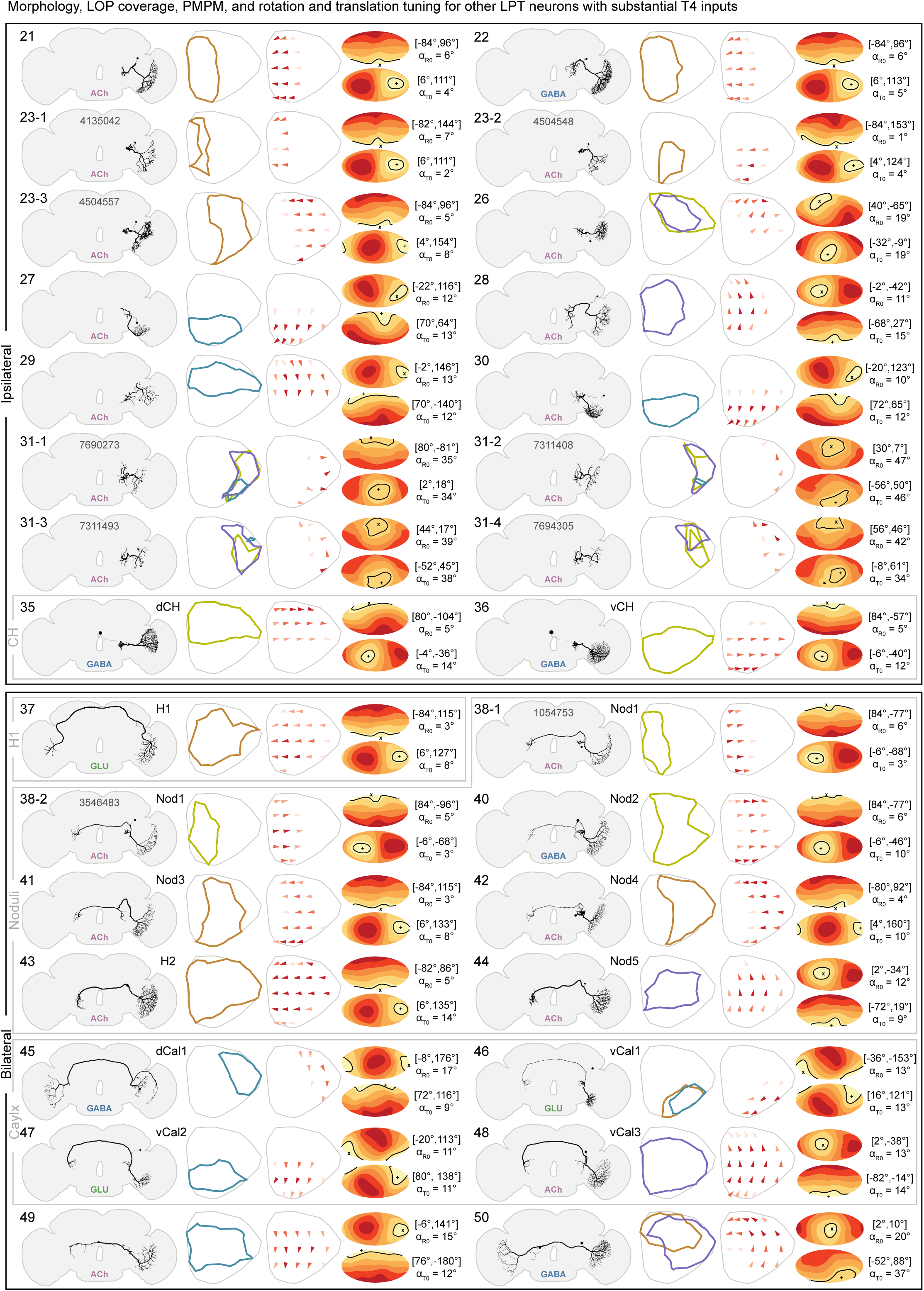
The other LPT neurons with T4 inputs—morphology, LOP coverage, PMPMs, and optic flow tuning. Following the convention of Figure 3, the other LPT neurons are presented, grouped by morphological categories, described in the text. From left to right: EM reconstruction, LOP layer coverage, color-coded to match Figure 2, PMPM, optic flow tuning as heatmaps of average angular difference (α_R_ for rotation; α_T_ for translation; coordinates [elevation, azimuth] and error value: α_R_ ≡ α_R0_ and α_T_ ≡ α_T0_ for minimal difference).

**Figure 5:**
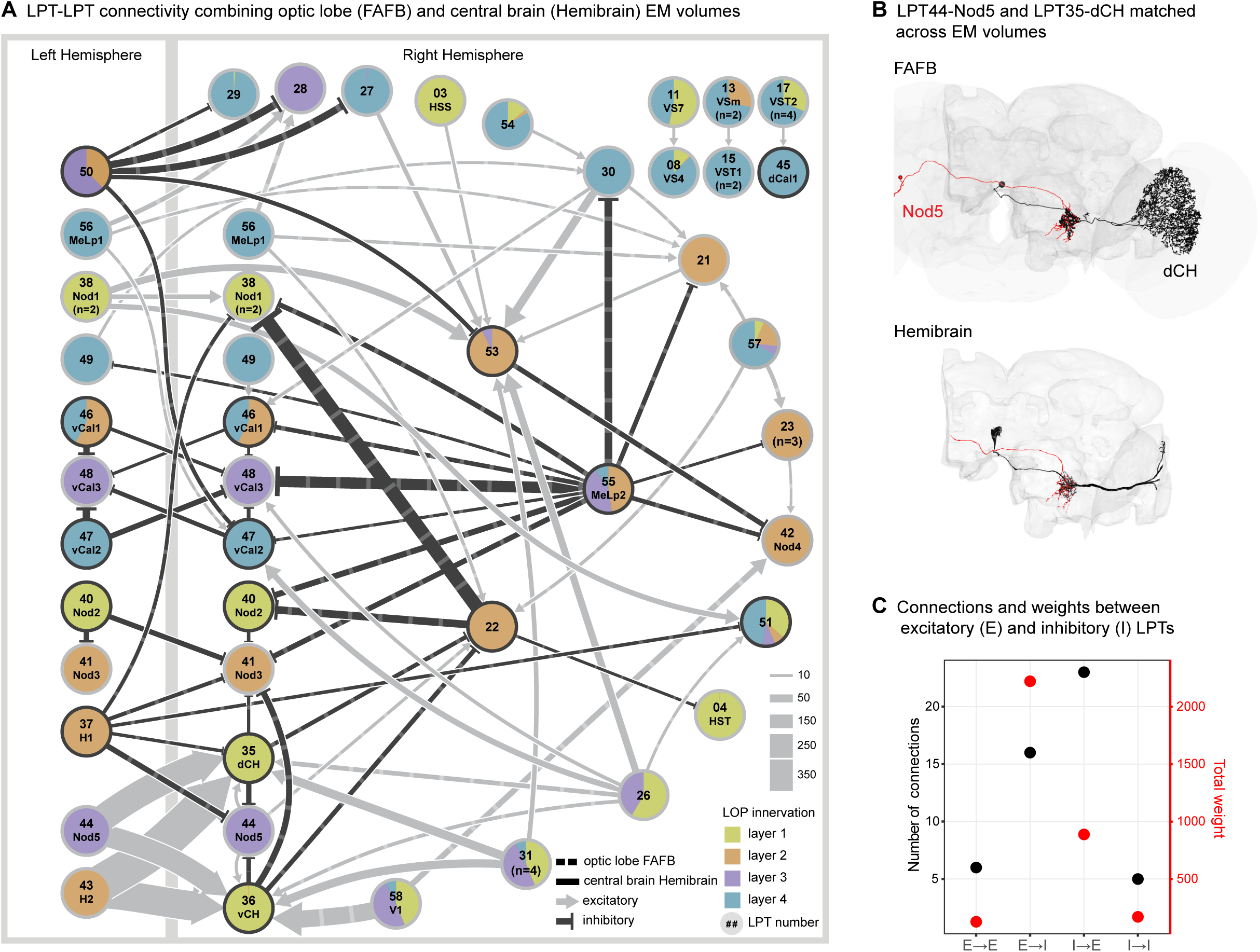
Brain-spanning network connectivity between LPT neurons. **A.** LPT-LPT network graph based on connectivity from the optic lobe in FAFB (reported here) and the central brain in the Hemibrain connectome (Scheffer et al., 2020). All connections ≥ 10 synapses between the LPT neurons are included, with the edges indicating where the connections are made, the strength of the connections, and whether they are expected to be excitatory (pre-synaptic cell predicted to be cholinergic) or inhibitory (pre-synaptic cell is either GABAergic or glutamatergic) based on Figure 3—figure supplement 2 (and Methods). Each LPT neuron is shown as a pie chart representing the layers of LOP innervation. **B.** Examples of two connected LPT neurons, LPT44-Nod5 and LPT35-dCH, with their independent reconstructions in the FAFB data and Hemibrain. **C**. Number of connections and connection weights between LPT neurons grouped by the predicted polarity of their output synapses.

**Figure 6:**
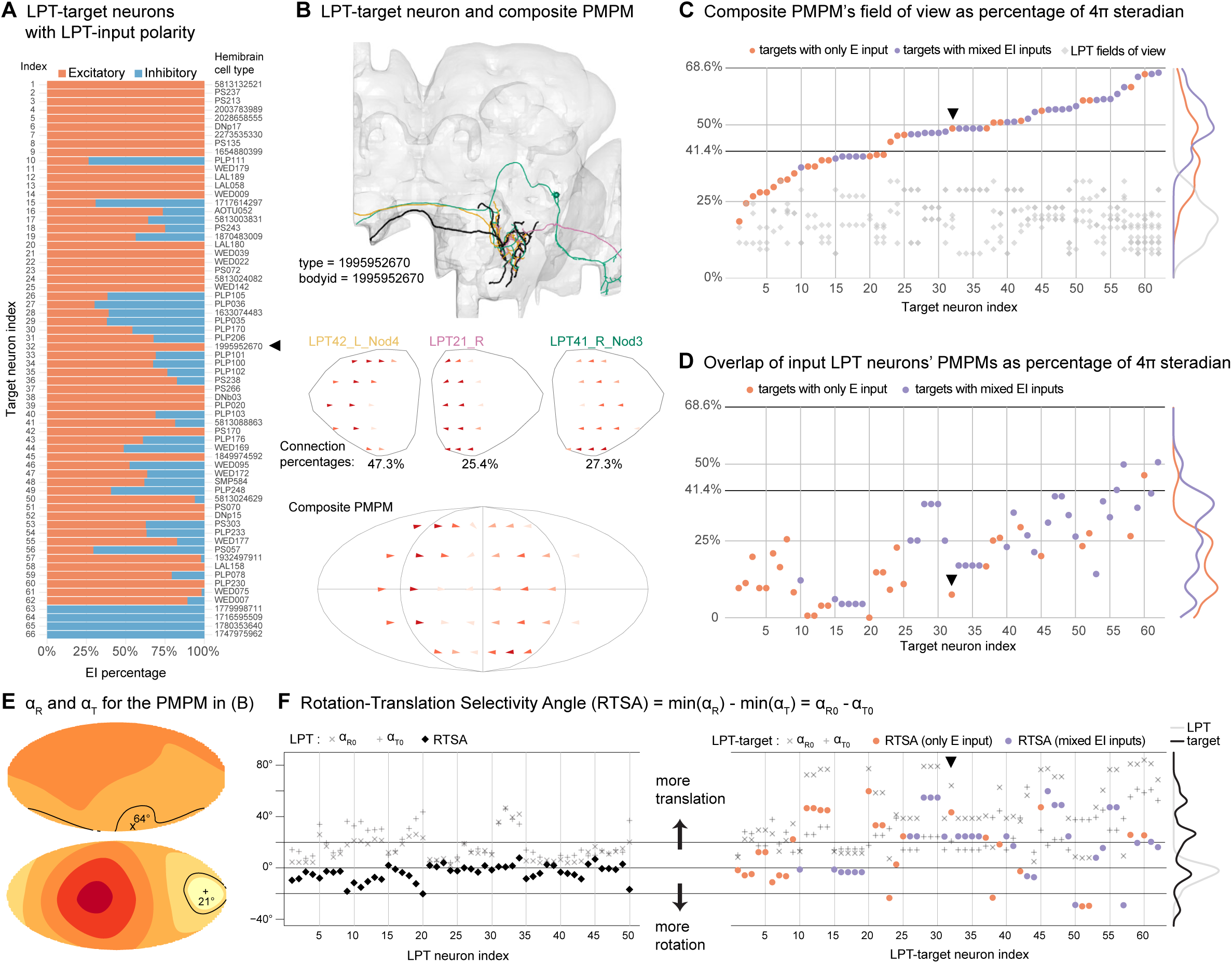
Predicted motion pattern maps for central brain LPT-target neurons. **A.** Excitatory and inhibitory input breakdown of 66 LPT-target cell types, indexed by their PMPM-based field of view (summing up the fields of view of input LPT neurons without double-counting overlaps). The 4 out of 66 cell types that only receive inhibitory inputs are excluded in the following analysis. **B**. An example of an LPT-target neuron in Hemibrain that receives inputs from three LPT types. Its PMPM is constructed from those of the input LPT neurons, weighted by connection weights (given as percentages in this example). We exclude inhibitory inputs in computing PMPMs. **C.** PMPM-based field of view (including inhibitory inputs) as a fraction of 4π solid angle. Also shown (in gray) are the fields of view of input LPT neurons. The field of view of one eye is 41.4%, and of both eyes is 68.6% (based on the eyemap used throughout, (Zhao et al., 2022)). **D**. Overlap of input LPT neurons’ fields of view. **E.** The PMPM for the example target neuron together with optimal rotation and translation axes. **F.** Rotation-translation Selectivity Angle, RTSA, for both LPT neurons and their central brain targets. Large positive values imply a better fit of the PMPM to an optic flow field induced by some translational movement. Large negative values for some rotational movement. Values near zero indicate that the PMPMs are ambiguous, not clearly selective for either translation or rotation.

## Results

### Comprehensive identification and reconstruction of *Drosophila* LPT neurons in an Electron Microscopy dataset

The group of Lobula Plate Tangential neurons have primarily been defined and identified by their anatomy, as large cells that span a substantial portion of the LOP and project into the central brain. Because of their importance for motion vision, as well as their striking neuroanatomy, we caried out a survey of the LPTs using manual neuron reconstruction in the FAFB (Zheng et al., 2018) EM volume. We initially searched for all large neurons in the right lobula plate, and traced their processes to confirm they are indeed projection neurons (we also encountered several large optic lobe intrinsic neurons, such a Am1 (Shinomiya et al., 2022), that we do not consider to be LPTs as they do no extend into the central brain, and were excluded from our survey). As the morphologies of these cells were completed, we began matching them to known LPTs. The Vertical System (VS) cells, many with massive primary dendrites in the 4^th^ layer of the LOP—at the most posterior part of the brain—were the very first cells to be identified (Figure 1A) with the Horizontal System (HS) neurons following shortly thereafter. In the EM data, these neurons resemble the images of the homologous neurons in *Musca domestica* (Pierantoni, 1976), with primary dendritic segments that are substantially larger than the finer processes of the columnar neurons passing nearby (Figure 1A, inset). To complete our survey of the LPTs, we focused on reconstructing all neurons with large axons emanating from the medial edge of the LOP, but this proved insufficient. In a parallel effort, we screened confocal images of stochastically labeled neurons expressed by GAL4 driver lines for candidate LPT neurons (see Methods). We then used these confocal images, with abundant anatomical landmarks, to target our reconstruction effort to find all LPT neurons identified during the Light Microscopy (LM) survey. Examples of cells of nine neuron types that we matched between Light Microscopy and EM reconstructions, are provided as Figure 1B. Most of these cells have not been previously described in *Drosophila*.

From our comprehensive reconstruction of all the large-field cells in the LOP of *Drosophila*, we present here, to the best of our knowledge, the most complete account of the LPT neurons in any insect. In total we find 58 LPT neurons. The set of neurons is listed with supporting reference identifications in Supplementary File 1. Further morphological details of these 58 neurons are provided in a gallery of all the reconstructed cells (Supplementary File 2). We sorted and named all the LPTs as follows: first the 20 total HS and VS neurons (summarized in Figure 3), then 16 ipsilateral neurons (axons that only project to the same side of the brain) and 14 bilateral neurons (with midline-crossing axons), summarized in Figure 4. Finally, we include reconstructions of 8 neurons that resemble LPTs, but these neurons receive a very small number of T4/T5 inputs and are therefore presumed to be feedback neurons, primarily sending information back into the LOP (Figure 4—figure supplement 1). These neurons are included since homologous neurons have been identified as LPTs in prior work (Hausen, 1987). Our uniform naming scheme assigns a number to each cell, incrementing by one each time, except in cases where multiple cells comprise a type that we could not distinguish based on morphology (between LM and EM and between different EM data sets), where we append a numerical identifier for each unique reconstructed cell. Wherever the match to a homologous, named neuron in blowflies is unambiguous, we appended the conventional names following Hausen (1987) to our numeric naming scheme. We named LPT55_MeLp2 and LPT56_MeLp1 to indicate that these cells project from the medulla to the contralateral LOP.

To confirm the identifiability (as unique cells) of all reconstructed LPTs and to gauge whether we may have missed large neurons in our manual search and reconstruction, we took advantage of two recent auto-segmentations of the same FAFB EM volume, FAFB-FFN1 (Li et al., 2020) and FlyWire (Dorkenwald et al., 2023). After an exhaustive search, with substantial editing in FlyWire, we found morphological matches for all tangential cells in the left side of the brain (gallery of left-right matches provided as Supplementary File 3, identifying details also listed in Supplementary File 1). Having these cells in FlyWire allowed us to use automatic synapse detection (Buhmann et al., 2021) and neurotransmitter prediction (Eckstein et al., 2020). LPT-LPT connectivity comparison provided additional confirmation for our cell typing and the left-right matches (Figure 3—figure supplement 1C). We assembled summary figures with many details about each LPT (Figure 3A and Figure 4), showing a brain-scale projection of the reconstructed neuron, the predicted neurotransmitter of the cell, the layer innervation within the LOP, as well as predictions for their sensitivity to optic flow fields.

### Predicting visual motion sensitivity from anatomy

It is now well established that most LPTs integrate local motion signals from directionally selective T4 and T5 neurons (Shinomiya et al., 2022). In this study we use detailed reconstructions of each LPT’s anatomy to estimate their motion sensitivity. We use the well-studied Vertical System neuron VS1 (Figure 1A, 2A) to illustrate our approach. The visual motion responses of VS1 have been measured in *Calliphora* (Krapp and Hengstenberg, 1996) and *Drosophila* (Joesch et al., 2008; Kim et al., 2017). Since the fly visual system maintains a retinotopic organization of columnar neurons across its neuropils, the LOP inherits the retinotopic representation of visual space from the medulla. The four subtypes of the directionally selective T4 neurons, each with a distinct directional tuning, project from the medulla to one of the 4 LOP layers (Figure 2B) (Shinomiya et al., 2022, 2019). In a recent study, we developed an *eyemap* of a female *Drosophila*, that maps the visual space sampled by the retina, onto the columnar organization of the medulla (Figure 2C) (Zhao et al., 2022). We reconstructed several hundred T4 neurons across the right-side medulla (Figure 2—figure supplement 1A) in the FAFB volume (Zheng et al., 2018), which we used to compute the preferred directions of these T4 neurons based on their dendritic orientation (Zhao et al., 2022). In the present study we use these reconstructed T4 neurons to extend the eyemap from the medulla to the LOP (Figure 2C,D) and to define LOP layers using the T4 subtypes’ axonal arbors (see Methods). Taken together, we now can assign retinotopic (visual) coordinates, as well as motion-direction sensitivity, to anatomical positions in the LOP, essentially creating a set of four maps for the local sensitivity to visual space and motion direction. Figure 2E shows these maps, down-sampled and in eye coordinates (further details in Figure 2—figure supplement 1B,C and Methods).

With a few additional steps, we use these maps to estimate the motion pattern sensitivity of LOP neurons. Our manual reconstruction of the VS1 neuron (Figure 1A) matches the morphology of previous descriptions of this cell, that noted a bi-stratified arbor in *Calliphora* (Hausen, 1987; Hengstenberg et al., 1982) and in *Drosophila* (Boergens et al., 2018; Scott et al., 2002; Shinomiya et al., 2022). Indeed, we find that VS1 dendrites innervate LOP2 and LOP4 (second and forth layers; Figure 2F). We found many connections between T4 neurons and the LPTs, but these large neurons receive thousands of synaptic connections from T4/T5s, and so a complete description of this connectivity is too large a task using manual reconstructions. Therefore, after confirming that subtype-specific connectivity indeed exists in accordance with our defined LOP layers, we used the spatial profile of each LPT’s skeletons to determine the retinotopic contribution of each LOP layer (Figure 2G; see Methods: Lobula plate layers; Predicted Motion Pattern Map). Combining the layer contributions and T4 preferred direction maps, we then estimate the visual motion pattern each LPT should be most sensitive to (Figure 2H), based solely on their presumed T4/T5 inputs. We refer to each of these estimates as the Predicted Motion Pattern Map, PMPM (Figure 2H). The assumption that T4/T5 neurons are the primary determinants of motion sensitivity is well-supported by recent connectivity data (Shinomiya et al., 2022), but of course this analysis cannot account for flow-field elaborations through network connections of other LPTs (Farrow et al., 2006) or gap junction coupling between cells (Ammer et al., 2022; Farrow et al., 2005; Haag and Borst, 2005).

The PMPM represents our best estimate for what motion patterns each LPT neuron should be sensitive to based on its LOP morphology. The PMPM for the VS1 neuron shows downward motion sensitivity in the frontal eye (established by the LOP4 dendritic innervation) and a back-to-front motion sensitivity in the dorsal eye (from LOP2 innervation). This response pattern is in excellent agreement with the VS1 response patterns measured in blowflies (Figure 2J, modified from Krapp and Hengstenberg, 1996) using a pioneering method for mapping the flow field sensitivity of LPT neurons (Krapp and Hengstenberg, 1997). As noted in this foundational characterization of these neurons, the VS cells appear to encode the visual motion consequence of body and/or head rotations around specific axes. To determine the tuning of each neuron to global optic flow fields, we compute, within the field of view of the neuron, the average angular difference between the PMPM and idealized optic flow fields induced by either a single rotation or translation, denoted by α_R_ and α_T_, respectively. We employed a ‘brute-force’ search where we densely sampled rotation or translation axes on a globe (Figure 2K). The top panels in Figure 2K show examples of idealized flow fields due to rotations or translations along 6 different axes with VS1’s PMPM overlayed. The distribution of angular differences, for all sampled axes across the globe (4π) are represented as heat maps (bottom row, Figure 2K, see Methods: Optimal motion axes). We found that VS1’s optimal sensitivity is to a rotation along an axis close the equator near the middle of right eye’s field of view (indicated with ‘X’), with a minimal α_R_ value, α_R0_ = 26°. Following the hypothesis that LPTs act as matched filters for optic flow (Franz and Krapp, 2000), the value of α_R0_ can be seen as a proxy for the maximal sensitivity of a neuron to any rotational motion. Small α_R0_ values imply that the neuron is well tuned for some rotational optic flow field. In comparison, the α_T_ heatmap (Figure 2K, bottom right) shows a larger minimal value of α_T0_ = 34°, consistent with prior interpretation of VS cells being more sensitive to rotational optic flow. However, this enhanced sensitivity for rotation does not preclude the possibility that VS1 could be used to estimate translational self-motion since optimal flow estimation benefits from motion detection over the largest field of view possible (Koenderink and van Doorn, 1987).

### The horizontal and vertical system neurons and their predicted motion patterns

During manual reconstruction of the large tangential neurons, we readily identified the HS and VS cells, and found additional neurons that had not been previously described in *Drosophila*, that we group together with these cells (Figure 3). We found 3 Horizontal neurons (following the widely used names of Pierantoni (1976), we also refer to them as North (HSN), Equatorial (HSE), and South (HSS), as well as an additional, quite similar cell type, which we call Horizontal twin (HST), following the suggestion of Pierantoni (1976). We did not find one twin cell per HS cell as was speculated in earlier work, but as LOP1 houses several other large neurons (see Figure 4), we do not think this is a likely difference between *Drosophila* and larger flies. While the classic LM description of the VS neurons in *Drosophila*, either using silver staining (Heisenberg et al., 1978) or genetic labeling (Scott et al., 2002), identified 6 large Vertical System cells, recent EM studies in *Drosophila* found additional neurons that resemble the VS cells, but have smaller profiles (Boergens et al., 2018; Shinomiya et al., 2022), consistent with previous reconstructions in larger flies (Bishop and Bishop, 1981; Pierantoni, 1976). We endeavored to complete the inventory of these cells, and indeed, can confirm these previous findings. We found 6 large VS cells, but then found 10 additional cells, for a total of 16 neurons. Several of these VS cells have medium size profiles, while 6 of them are quite small (Figure 3— figure supplement 1A). We’ve extended the nomenclature of previous studies by using the LOP layer innervations, primary 4^th^ layer dendritic location, and axon diameters to determine the numbering. We arrive at 8 VS cells, 2 VSm cells that have similar dendritic arbors as VS1/2 but smaller axon diameters and axon terminals somewhat different from VS1/2, and 6 VS twin (VST) cells with smaller axon profiles. The 6 twin cells are further divided into 2 subtypes, VST1 and VST2, whose members we cannot further distinguish by morphology. Using FlyWire we also found morphological matches for the 4 HS and 16 VS neurons on the left side of the brain (Figure 3—figure supplement 1B; gallery of left-right matches provided as Supplementary File 3).

By comparing the PMPMs to ideal optic flow fields and computing α_R0_ and α_T0_, we find additional support for the prior claims that VS cells are more selective for rotation, however, in the cases of the neurons with small fields of view, the preference for rotational optic flow fields over translation can be quite small. Furthermore, we see that the set of VS cells from one side of the brain can be used to measure a range of rotation axes around the equator, covering an azimuthal range of about 135° (Figure 3B). Based on the neurotransmitter predictions, all HS and VS neurons are predicted to be cholinergic (Figure 3—figure supplement 2A, 3), and thus presumably excitatory. The implications of these predictions are treated in the Discussion.

### Other large-field tangential neurons and their predicted motion patterns

While decades of attention have focused on the HS and VS cells, it has been clear, at least since Hausen’s pioneering work in *Calliphora* (1987), that the LOP houses dozens of other large-field tangential neurons. However, due to their thinner axons, and variety of ‘exit’ points from the LOP, these cells have been challenging targets for electrophysiology, and most have not been previously described in *Drosophila* using genetic driver lines (with the noteworthy exception of a recent GAL4 driver screen (Wei et al., 2020)). Figure 4 summarizes the (“LOP output”) LPT neurons apart from the HS and VS cells. The table is organized by listing the ipsilateral neurons first, and then the bilateral cell types. For most of the ipsilateral group neurons, we could not confidently match them to blowfly LPTs and we therefore sorted them based on their LOP innervations. While most ipsilateral neurons are individually identifiable, the LPT23 and LPT31 cells are not. While individual LPT23 and LPT31 cells look distinguishable, these differences are not sufficiently stereotyped to enable matching individual cells across brain hemispheres or to LM data.

Wherever we could confidently match Hausen’s descriptions of LPTs in blowflies (1987), we append the names of the homologous cells. Following Hausen’s classifications, we found 3 more neurons in the posterior slope group (together with HS and VS neurons): DCH, VCH, and V1. All three were considered feedback (or “centrifugal”) neurons. However, since we found a reasonably high number of connections from T4s to DCH/VCH, we listed them in Figure 4, while listing V1 in Figure 4—figure supplement 1 along with 7 other ‘feedback’ neurons. We also found a group of LPT neurons whose axons pass through a tract frontal to the noduli of the central complex, just as they do in the blowfly brain. We therefore termed them the noduli group, which included the well-known H2 neuron (reconstruction reported in Zhao, et al. (2022)), and 6 other cells, that we call Nod1-5 (2 of these cells have extremely similar morphology and connectivity, and so both are named as the type Nod1). We found all 4 neurons (vCal1-3, and dCal1) that Hausen (1987) described in the Calyx group, cells whose axons pass near the calyces of the mushroom bodies. Finally we found the H1 neuron of Hausen’s (1987) dorsofrontal group (the only member). All of these neurons could be readily matched, one-for-one, between the *Calliphora* reconstructions (Hausen, 1987) and our *Drosophila* data, despite considerable differences in scale and neuropil shape between these rather different Diptera.

As with the HS/VS neurons, we report the summary predicted neurotransmitter for each reconstructed neuron (Figure 4), which agree for the independent right and left reconstructed neurons in all cases except for 2 (Figure 3—figure supplement 2). We observed two ambiguous predictions with, in both cases, similar scores for predicted GABAergic and glutamatergic phenotypes. In one case, LPT37_H1, recent experimental data (Wei et al., 2020) suggest a glutamatergic phenotype and we assign H1 accordingly. For LPT_55_MeLp2, a neuron that projects from the contralateral medulla to the LOP, with a similar inconclusive prediction, we used a new, specific split-GAL4 line for this cell type together with a recently developed method, Expansion-Assisted Iterative Fluorescence *In Situ* Hybridization (EASI-FISH) (Eddison and Ihrke, 2022; Wang et al., 2021), to assess neurotransmitter marker expression. These experiments indicated that this cell type is likely glutamatergic (Figure 3—figure supplement 2B,C).

We also applied our optic-flow-field “dense-sampling” method to compute the PMPMs for each LPT neuron other than the feedback neurons (Figure 4). Each PMPM highlights the optimal rotation and translation axes, which we assemble for comparison in Figure 4—figure supplement 2, with the diameter of each point indicating the size of the error. This comparison shows that the optimal axes cover much of the sphere, however, noteworthy clusters are seen around the 3 principal axes derived from the morphology of the compound eye (Zhao et al., 2022), an expected result since many neurons primarily receive their inputs from a single LOP layer (Figures 3, 4).

### Brain-spanning network connectivity between LPTs

It has long been appreciated that LPT neurons are not simply outputs of the visual system, and indeed connections between them refine the visual patterns they encode (Farrow et al., 2006, 2005; Haag and Borst, 2003, 2001). With our inventory of LPT neurons we can now more systematically examine the connections between LPTs, summarized in Figure 5, as well as the connections from LPTs to other targets in the central brain (Figure 6). The FAFB data set was ideal for surveying the LPTs and allowed us to manually reconstruct and identify them, however, those methods were impractical for exhaustively sampling their connectivity into the central brain. Fortunately, the Hemibrain connectome (Scheffer et al., 2020), focused on most of the central brain of a female fly, became available during our project. To enable the analysis of circuits downstream of the LPTs, we endeavored to match the cells, one-for-one, between FAFB and Hemibrain, making productive use of both connectome data sets together. Since the Hemibrain contains only a sliver of the LOP, matching neurons relied on their central brain morphology in the Hemibrain and their connectivity (see Methods), in an iterative matching process (coordinated with Erginkaya, et al. (2023)). We noted that the LPT axons were in brain regions that were not as thoroughly proofread as others, and so we initiated additional proofreading in the Hemibrain, targeted to our matched LPTs. In most cases we substantially improved the completeness of the connectivity data (Figure 5—figure supplement 1A), which will be included in an upcoming Hemibrain data update. In total, out of the 58 LPT neurons in FAFB, we found 57 right-side matches and 17 left-side (all bilateral neuron) matches (Hemibrain bodyids for the matched cells listed Supplementary File 1). Figure 5B shows an example of two strongly connected neurons, dCH and Nod5, with our FAFB reconstructions and the corresponding matched cells in Hemibrain. The full set of FAFB-Hemibrain matched LPT neurons, are superimposed and displayed in Supplementary File 4. To confirm the matches between these different brains, we compared the connectivity between the LPT neurons in each data set (Figure 5—figure supplement 1B, and Methods). We note that only the synaptic connections between LPT neurons in the central brain were used, since other target neurons were mostly unidentified in FAFB, and the Hemibrain lacks most of the LOP. The connections are remarkably consistent between the data sets, with the identical set of central brain connections present in both (Figure 5—figure supplement 1B), validating the matches for these cells.

Figure 5A summarizes the synaptic connections (≥ 10 synapses) between LPT neurons, incorporating data from the right-side optic lobe in FAFB and central brain in Hemibrain. Each LPT neuron is shown as a pie chart, representing the layers of LOP innervation. In this diagram, the edges indicate where the connections are made (central brain or optic lobe), the strength of the connections, and whether these connections are likely excitatory (the pre-synaptic cell is predicted to be cholinergic) or inhibitory (pre-synaptic cell is either GABAergic or glutamatergic) based on the neurotransmitter predictions of Figure 3—figure supplement 2A. We’d like to make a few simple observations here: (1) The majority of the connections and the connection weights (equivalently, number of synapses) occur between excitatory and inhibitory neurons (Figure 5C), suggesting that regulation/suppression of responses in the LP network is a major feature, likely more so than broadening of tuning (as would be expected for E→E connections). As we will explore in the next section, there is abundant integration of excitatory cells by downstream neurons, but we only find a few examples of this in the LOP network, e.g. LPTC53. (2) VS and HS neurons are mostly absent from this graph, or form isolated sub-graphs (upper right corner). The VS cells are missing because we did not find strong connections to or from them in the optic lobe, and their axons are mainly cut-off in the Hemibrain volume. It is known that the VS cells are coupled via gap junctions (Ammer et al., 2022; Farrow et al., 2005; Joesch et al., 2008), which are not detectable in our EM data sets. The lack of connectivity to HS cells does not result from any technical limitation, and instead appears to reflect how specialized they are, not sharing direct connections with other LPT neurons (Erginkaya et al., 2023). (3) Many LPT-like LOP input neurons play prominent roles in this network. In our naming, we identify these neurons with a number ≥ 51. LPT35-dCH and LPT36-vCH are also major LOP inputs, as these neurons receive substantial inputs from left-side neurons, notably LPTC43-H2 and LPTC44-Nod5. Other neurons that come to prominence by virtue of their wide-spread contributions (≥ 4 targets) in the LPT network are LPT26, LPT37-H1, LPT50, LPT55-MeLP2, among others.

There is impressive experimental evidence from a variety of blowfly electrophysiology studies for specific identified connections between LPT neurons (Farrow et al., 2006; Haag and Borst, 2007, 2003, 2001; Kurtz et al., 2001; Warzecha et al., 1993). Many of these connections are found here, for example Haag and Borst (2003) found evidence for V1 input into vCH and speculated that this must occur in the LOP, and indeed we find exactly this connection (at the very bottom of the graph in Figure 5A). However, other prominent connections from prior work are not fully in agreement with what we find in *Drosophila* EM connectivity. In studies of connections between horizontal-motion sensitive neurons across the midline, Haag and Borst (2001) found excitatory connections between both H1 and H2 and HSN/HSE and dCH/vCH, whereas our results show the excitatory connections from H2 to the CH cells, but not to the HS cells, and we expect that H1 will be inhibitory, and only find a weak connection from H1 to one of the CH cells, none to HS neurons, and weak connections from HSE to H2 (Figure 5 and Figure 5—figure supplement 1). More recent work showed the H2-HSE connection to be electrical (Farrow et al., 2006; Pokusaeva et al., 2023), so the primary coupling between these cells may be mediated by gap junctions. Furthermore, our network shows many indirect connections between these well-studied cells, illustrating the challenge of a complete reconciliation with this prior work. It is unclear to what extent species differences, the role of electrical synapses, or indirect connections that were interpreted as direct ones in the prior work, will explain these discrepancies. We think there is a golden opportunity to thoroughly understand this network in *Drosophila*, where this graph, together with our PMPMs for the LPTs, establishes a roadmap for experimental validation—marked differences from the predicted flow field sensitivity may well be explained via these network connections. A detailed exploration of the HS-cell network (Erginkaya et al., 2023), shows the power of combining connectomic predictions with functional studies of these networks.

### Enhanced flow-field selectivity in central brain neurons integrating LPT inputs

As LPTs are tuned to a variety of optic flow fields, depicted by the PMPMs in Figures 3 and 4, they already encode information that can be used for estimating state information and for motor control, but the LPTs are hardly the ‘end of the line’ for optic flow estimation. Now that LPT connections can be tabulated in the central brain, we can examine how the signals from different LPTs are integrated by central neurons, and ask whether these combinations provide clues about representations of self-motion. Perhaps the clearest example involves the monocular signals of horizontal-motion-sensitive cells (like HS, H2, etc.) that could be used to encode either yaw rotations or body translations in the equatorial plane; these different body movements could be robustly distinguished by combining inputs from these cells across both eyes (a major theme explored in Erginkaya et al., 2023). To look for these combinations, we investigated the direct synaptic targets of LPT neurons in the central brain. We excluded the LOP-input LPT neurons (with no PMPMs) and were left with 64 LPT neurons (49 right-side and 15 left-side) out of the 74 Hemibrain matches. One can consider this as a 2-layer network: nodes in the first layer are LPT neurons and each node in the second layer has at least one connection from the first layer. The network has 1246 nodes and 52385 directed connections. For this initial exploration of the network, we focused on the LPT-target neurons that combine inputs from multiple (≥ 2) LPTs with strong inputs from them (each connection ≥ 10 synapses), and receive a substantial fraction of their total inputs from LPTs (≥ 10% of all incoming connection weight). These criteria established a set of 138 neurons (nodes in the second layer) that are strong recipients of inputs from multiple LPT neurons. These neurons can be further grouped into 66 types according to Hemibrain annotation. Some LPT target neurons do not have a hemibrain type annotation. We did not attempt to further group these unnamed neurons into types here, although similarities between some of these cells suggest that some may belong to the same types. Four of the 66 cell types only receive inhibitory LPT inputs (assuming both GABA and glutamate are inhibitory) while the rest receive either only excitatory or mixed LPT inputs (Figure 6A).

We constructed a composite PMPM for each LPT-target neuron (or combined for neurons of the same named type) based on its excitatory LPT inputs. Figure 6B shows an example where the target neuron (named by its Hemibrain bodyid as it does not carry a type label) receives 3 excitatory LPT inputs, 1 from a left-side LPT and 2 from right-side LPTs. To generate the composite PMPM for this target neuron, we sum the PMPMs for each input LPT (and impose left-right symmetry for the left-side LPTs), weighted by the fractional connectivity weights (see Methods for details). The composite PMPM for the example neuron (Figure 6B, bottom), shows an expectedly larger receptive field than any of the input neurons, but also shows the structure of a matched filter for a global flow-field that distinguishes between the body movements in the scenario above—preferring backwards translation by integrating regressive motion sensitivity from LOP2 neurons on both sides. As it is less clear how to account for the contribution of inhibitory inputs into PMPMs, we do not include them in our estimates, but separately depict their potential contribution in the more detailed plots for each LPT-target neuron (Supplementary File 5). We note that the predicted tuning of uLPTCrns (#23 and #51) and DNp15 (#52), featuring a marked preference for yaw rotation, are in excellent agreement with recent physiological measurements of these neurons (Erginkaya et al., 2023).

From the LPT neurons’ PMPMs, we can readily compute each neuron’s field of view (Figure 6—figure supplement 1A,B), the fields of view of LPT-target neurons (Figure 6C), and the overlap of their LPT inputs (Figure 6D). Note that in this case, we treated excitatory and inhibitory LPT neurons equivalently, but we excluded the four inhibitory-only LPT-target neurons (bottom of Figure 6A) in our summary data analysis (Figure 6C,D,F). We then ordered the remaining LPT-target neurons (“Target neuron index”) by their fields of view (Figure 6C). In the marginal histograms, we separated targets with excitatory only LPT inputs, indicated in red) from those with mixed inputs (indicated in blue). The marginal histograms show that most of these neurons receive strong LPT connections covering more than the field of view of a single eye (Figure 6C). We find that LPT-target neurons rarely integrate from LPTs with redundant, heavily overlapping fields of view (Figure 6D), with the noteworthy exception of neurons with mixed EI LPT inputs. In most of these mixed EI LPT input cases, the overlapping LPTs feature locally opposing preferred directions (see gallery in Supplementary File 5), consistent with a central brain implementation of motion opponency (Isaacson et al., 2023; Mauss et al., 2015).

Is enhanced tuning for global optic flow patterns a common feature of the central brain LPT-target neurons? In principle these cells could integrate from either synergistic combinations of LPTs (as in the example of Figure 6B) or from seemingly arbitrary combinations of LPTs that may jointly be maximally sensitive to incoherent flow fields, not obviously useful for detecting body movements. We applied the “brute force” search for the optimal rotation or translation axes for each of the LPT-target neurons. As discussed above, we now see that the example neuron in Figure 6B clearly shows a strong preference for translational motion (Figure 6E), along a direction close to straight backwards. Partial measurements of any global optic flow field will lead to ambiguous estimates of that global pattern, and therefore the associated body movement. We have included the heatmaps of the flow-field errors throughout this study to visually emphasize this ambiguity. The estimates are ambiguous both about whether the measurements best “explain” rotational or translational movements, as well as the exact axis of movement. The smallest errors for rotation, α_R0_, and translation, α_T0_, numerically capture the ambiguity of each neuron’s estimates of a flow field.

This ambiguity could be minimized by “cleverly” sampling the visual space, but detecting coherent motion patterns across larger fields of view is the ultimate resolution to this ambiguity. To capture this difference between the sensitivity to the best-aligned flow field for rotation and translation, we defined the Rotation-Translation Selectivity Angle, RTSA = α_R0_ - α_T0_, as a measure of the amount of intrinsic ambiguity of a PMPM for detecting optic flow. The RTSA indirectly compares the sampling strategy: the more positive the RTSA is, the more selective (less ambiguous) that neuron is for detecting translation. Conversely, larger negative values indicate selectivity for rotational flow fields. When RTSA is close to 0, the neuron is equally good (or bad, depending on the absolute values of α_R0_ and α_T0_) at detecting rotational and translation flow fields. The RTSAs along with α_R0_ and α_T0_ of all LPTs (for which we computed PMPMs) are shown on the left side of Figure 6F, suggesting that most LPT neurons are not selective for either rotational or translational flow. The most selective LPTs are mostly those with RTSA < 0, indicating a larger set of neurons with some selectivity for rotational flow fields, such as the VS cells (LPT neuron index 5 - 20).

The “brute force” method assumes the motion of the observer is either a single rotation or a single translation, while a fly moving through the world will often experience some combination of both at the same time. To solve for the more general case, we applied a non-linear regression method to an optic flow model proposed by Koenderink and van Doorn (1987) to find the observer motion as a combination of rotation and translation that best fits the PMPM (see Methods). The resultant minimal average angular difference (α_0_) represents a best-case scenario for finding a body movement that best matches each LPTs flow field (Figure 6—figure supplement 1C). In the majority of cases, α_0_ is just slightly smaller than the smallest of either α_R0_ and α_T0_. This comparison offers important context for understanding the flow-field selectivity of individual LPTs. If α_0_ ≈ α_R0_ (or α_T0_), it suggests the LPT neuron is better tuned to a rotation (or translation), while α_0_ << α_R0_ and α_0_ << α_T0_ suggests the LPT neuron is better tuned to a combination of rotation and translation (but there are few examples of this in our data).

Returning to the selectivity established by integrating from LPTs, we find that many LPT-targets (right side of Figure 6F) feature larger RTSAs with many cells showing a strong selectivity for translational flow fields. In the case of bilateral integration by the neuron in Figure 6B,E, the inclusion of large portions of both left and right fields of view with mirror-symmetric sensitivity, reduces the matching to rotational flow fields (example neuron highlighted with the arrowhead in Figure 6F). This summary shows that more than half of the LPT-target neurons feature marked selectivity for either rotational or translation flow fields, with much higher selectivity (outside of the horizontal bands in Figure 6F) than estimated for any of the LPTs (preferred axes of motion sensitivity for LPT-targets detailed in Supplemental File 5). This result provides evidence for a key insight about optic flow processing—the central brain combines smaller-field LPT responses to establish robust detection of movement-generated global flow fields.

### Patterns of LPT output integration in the central brain

As an initial step towards analyzing the patterns of LPT integration by their central brain targets, we asked whether specific LPTs combinations are integrated with higher probability. Using the set of neurons considered in Figure 5, that receive variable number of LPT inputs (Figure 6— figure supplement 2A), we constructed a “pairwise collaboration matrix” (Figure 6—figure supplement 2B) that was inspired by a similar analysis of the integration of Lobula Columnar neurons (Klapoetke et al., 2022). Each entry in the collaboration matrix represents all the LPT-target neurons that receive inputs from the LPT neuron in that row and the LPT neuron in that column. The value of the entry, however, represents the total connection weight (across all LPT-target neurons in the entry) from the LPT neuron in the row (hence the asymmetry). This matrix shows clear structure, with some sparseness—many pairs of LPT inputs are not found in the connectome, while some pairs are quite prevalent. To further explore this structure, we used spectral clustering (see Methods) to group the LPT neurons into sets of neurons that are more likely to share common targets (Figure 6—figure supplement 2D).

The principle of wiring economy (Chklovskii and Koulakov, 2004; Klapoetke et al., 2022; Rivera-Alba et al., 2011) suggests one perspective for understanding these clusters, since LPT neurons from each group may be anatomically organized together with their common targets into small brain regions that would reduce the total wiring cost (and thus brain volume). To check this, we examined the anatomical distributions of the synapses in each group and compared them with neuropil regions in the central brain. All LPT output synapses in our data set, taken together, are present in 6 central brain neuropils: lateral lobe (LAL), wedge (WED), posterior lateral protocerebrum (PLP), superior posterior slope (SPS), inferior posterior slope (IPS), and gnathal ganglia (GNG). 4 of 5 groups resulting from the spectral clustering have their outgoing synapses present in small anatomical clusters, suggesting support for the idea that these connectivity clusters also have a compact anatomical basis, while the 5^th^ group expands into all 6 neuropils (Figure 6—figure supplement 2E). A more intriguing possibility is that these clusters represent units of movement detection that are organized for motor control, but a proper investigation of this speculation must be left for future work.

## Discussion

Our survey of the LPT neurons in *Drosophila* began from two distinct starting points: manual reconstruction of large neurons in the lobula plate of the FAFB EM volume (Figure 1A) and stochastically labelled neurons expressed by GAL4 driver lines imaged with light microscopy, LM (Figure 1B). During this process we closely compared our EM reconstructions to the LM images, using the LM images to guide the search for neurons not yet found in the EM. Over time, we were able to find all or nearly all the LPTs in FAFB, and then were able to match this set of EM reconstructed neurons to additional data sets, the Hemibrain connectome of the fly central brain (Scheffer et al., 2020)and most recently the FlyWire connectome of the FAFB brain (Dorkenwald et al., 2023). By integrating our extensive survey of the LPTs with neuroanatomical landmarks (Figure 2), we developed predictions for the optic-flow tuning of all the LPTs (Figures 3 and 4). Using our cell-by-cell matches to the Hemibrain connectome, we described the connections between LPTs (Figure 5) and central neurons (Figure 6), allowing us to make dozens of predictions about optic-flow detection in the central brain, setting up multiple specific hypotheses about the pathways linking motion vision with behavioral control.

### Homology of LPTs across distant species

While we made considerable attempts to match as many LPT neurons as possible to homologous cells previously described in blowflies, we preferred to anchor our cross-species matching in a more conservative approach based primarily on the morphology of neurons. Therefore, we can confidently match many groups of cells, such as the HS and VS neurons (Figure 3), and the members of the Noduli and Calyx groups (Figure 4), and some individual neurons: VCH, DCH, H1, H2, V1. To confidently match neurons beyond this set is challenging. Many recordings exist for neurons given names based on their functional properties ascertained with neurophysiology, but often the detailed morphology is either not available, or insufficient to establish confident matches. For example, several of the “Figure-Detecting” FD cells described by Egelhaaf (1985), are almost certainly homologous to some of the cells in *Drosophila*’s Noduli group, but one-for-one matches were hard to establish. The Vi neuron (Haag and Borst, 2007), was similarly hard to match based on its published morphology, however, as suggested when the cell was first described, it’s morphology is a reasonable match to dCal1. Following Hausen’s naming, we did not match the cells named as FD1/2/3/4, Pos1/3/4, H3/4/5, Cen1/2, Ant, and V2. The cell Hausen calls Amc1, we now refer to as Am1 (Shinomiya et al., 2022), was found in FAFB (see Figure 3—figure supplement 3B), but do not include in our LPT survey since it is an optic lobe intrinsic neuron.

Hausen provided the most complete collection of blowfly LPTs, and listed fewer than 50 comparable cells (1987). The literature often reports that there are ∼60 LPTs in the *Calliphora* LOP, but we are unaware of a definitive, original source enumerating this set of neurons. It is noteworthy that our collection of *Drosophila* LPTs finds 58 per side, demonstrating that the LOP of the smaller fly is no less equipped for detecting motion patterns than the LOP of the larger flies, in contrast to speculation in earlier surveys of e.g. the VS cells (Hengstenberg et al., 1982). Given the remarkable morphological consistency of the matched cells, it is likely that most cells have bona-fide homologues, but future physiological measurements of the unmatched neurons in *Drosophila* may be the best way to confidently establish these matches (de Vries and Clandinin, 2012; Tanaka et al., 2023; Wasserman et al., 2015).

### Using neurotransmitter predictions to assign functional signs to LPT output synapses

It is remarkable that a deep neural network trained on ground-truth data can reliably predict the neurotransmitter released from a pre-synaptic site using only the EM data (Eckstein et al., 2020). Based on the reliability of this method in other data sets, we sought to apply it to our reconstructed LPTs, and used right-left matches (Figure 3—figure supplement 1) as an independent check on each neurons’ predicted neurotransmitter phenotype. Of the 58 LPTs in our catalog, we find 56 with right-left agreement, with only two discrepant cells, both of which show high levels of predictions for GABA and Glutamate on both sides. Based on prior work and our new data, we judge these neurons to be most likely glutamatergic. To summarize these Predictions (Figure 3, Figure 4, Figure 3—figure supplement 2), we find 7 GABAergic LPTs, 5 glutamatergic LPTs, and the remaining 46 LPTs are cholinergic. While exceptions to the rule may exist, the cholinergic neurons are expected to excite their post-synaptic partners, while the GABAergic neurons will inhibit their targets. Glutamatergic neurons could either excite or inhibit, but most examples in *Drosophila* are of glutamatergic neurons driving inhibition via the glutamate-gated chloride channel, GluClα (Liu and Wilson, 2013; Mauss et al., 2015; Strother et al., 2017). This convention is used to produce the signs in the network diagram of Figure 5 and in the analysis of Figure 6.

Comparing these predictions to the blowfly catalog of LPTs is not simple. For example, all 20 HS and VS cells are predicted to be cholinergic. We are unaware of direct evidence for the neurotransmitter released by these neurons in blowflies, however, studies using paired recordings, have shown that HS neurons (Haag and Borst, 2001) and VS neurons (Kurtz et al., 2001) are most likely excitatory, consistent with a cholinergic phenotype. We provide further evidence for this prediction, since our FISH labeling of the cell-body region where HS and VS somata are found, shows most cells, including the majority of very large ones, with clear ChaT labelling (and ∼6 GAD1+ somata as well, far fewer than the number of VS and HS cells; Figure 3—figure supplement 3). There are some neurons in blowfly that have been shown to be inhibitory. The best studied are the CH neurons which have been shown to be GABAergic with immunohistochemistry (Meyer et al., 1986) and functionally with electrophysiology and pharmacology (Warzecha et al., 1993), matching our prediction here. While these EM predictions, without independent experimental confirmation must be taken with some caution, the high degree of left-right agreement, as well as consistency, where possible, with prior work, suggests these predictions should be treated as the best available guess at this time, and any unexpected results (such as the effective sign of the glutamatergic H1 neuron) should be prioritized for future experiments.

### Limitations of anatomical predictions

All methods of characterizing a neurons’ properties rely on many assumptions, and no method is perfect. Even the gold standard, neurophysiological measurements of LPT responses, are limited to the field of view that can be experimentally stimulated (< 4π), and even more limited by the assumption that local preferred directions of these large neurons can be measured closer to the ‘output’ of the neurons. While this prediction has been impressively examined in blowfly VS neurons studies (Hopp et al., 2014), it should be an empirical matter whether these findings should apply to all LPT neurons, and perhaps even harder to assess in *Drosophila* experiments, where the most reliable electrophysiological measurements are made from the soma (via whole cell patch clamp) and are thus heavily filtered. Moreover, it is even less likely that the pioneering method for characterizing LPTs (Krapp and Hengstenberg, 1997) would work as well for their target neurons, since these local stimuli’s effect on neuron responses must pass through another synaptic (threshold) non-linearity and the predicted LPT responses are generally not spatially overlapping (Figure 6).

Considering these challenges, we submit that our anatomical predictions are an essential step towards understanding this network, and more generally for understanding optic flow processing in arthropods. As previously discussed, our estimates do not incorporate any effects of electrical synapses, as they are absent in the EM data we used, and do not incorporate network interactions from other LPTs (Figure 5) or other central brain neurons that may mediate behavioral state dependent modulation and gating (Chiappe et al., 2010; Erginkaya et al., 2023; Fujiwara et al., 2022; Maimon et al., 2010; Strother et al., 2018). Another limitation is that these data are based on only 2 fly brains, both of which lack complete connectivity information, but encouragingly the FlyWire connectome (Dorkenwald et al., 2023; Schlegel et al., 2023) will fill in many of these details, as will additional fly connectomes expected in the next few years. These new data present opportunities to reevaluate our analysis and predictions together with future experimental results. These experiments will be critical, and our estimates can be used to design experiments, prioritize cells to record from and manipulate, and to test hypotheses. Any substantial deviations from our predictions should be explored, with the limiting factors discussed above considered as hypotheses to explain these differences.

### The complete Lobula Plate Tangential Neuron catalog

We believe that the completeness of our catalog is the most important contribution of our study, since exposing these details in the *Drosophila* brain sets up a range of future studies. Firstly, having access to all the major Lobula Plate outputs, from this study plus recent work (Isaacson et al., 2023; Shinomiya et al., 2022) provides a clearer and more comprehensive view of the optic flow information that is available to the central brain. The analysis presented in Figure 6 and Figure 6-supplement 2, are just a first step towards exploring these circuits. Secondly, the catalog provides researchers with specific targets, and some functional predictions about them, to guide search and construction of genetic driver lines enabling future experiments. Importantly, these efforts will open the possibility of studying behavioral roles for neurons other than the well-studied HS/VS cells. And finally, by bringing functional predictions into several connectomic data sets, the analysis of full-brain behavioral circuits with well-defined inputs becomes feasible. There are many unanswered questions about how optic flow connects to navigation control, for example to the increasingly well-studied visual-motion sensitive neurons in the Central Complex (Hulse et al., 2023, 2021; Lyu et al., 2022; Weir and Dickinson, 2015). By having complete brain data sets (Zheng et al., 2018; Dorkenwald et al., 2023), we can pursue these pathways in less explored regions of the brain—following visual information wherever the anatomy reveals it is flowing.

## Methods

### Manual reconstruction in CATMAID

All LPT neurons were manually reconstructed in a serial section transmission electron microscopy (ssTEM) volume of the full adult female brain, FAFB, of *Drosophila melanogaster,* (Zheng et al., 2018), using the CATMAID environment (Saalfeld et al., 2009). We followed established guidelines (Schneider-Mizell et al., 2016) for tracing neuron skeletons and identifying synapses. This project began around 2016, with pre-publication access to the FAFB volume prior to auto-segmentation, and as the project evolved, we incorporated new strategies and methods. The high-quality but laborious manual reconstruction of the LPT neurons in the FAFB volume was mainly focused on identifying all these neurons using their detailed morphology. We note that a recent comparison of EM datasets (Schlegel et al., 2023) found that the FAFB EM volume is right-left inverted. Therefore, all neurons we refer to as ‘right-side’ neurons are from the physical left-side of that fly’s brain. Since fly neurons and their connectivity are highly stereotyped across brain hemispheres and animals, and to maintain consistency in nomenclature and the conventions we (and others tracing in FAFB) have used for years, including our collaborators (Erginkaya et al., 2023), we retain the mirrored names here.

We note that this makes the comparison of our neurons to those in Hemibrain (e.g. Supplementary File 4) more intuitive. We started with right hemisphere LPT neurons. The initial strategy was to examine all neurons with large caliber processes that cross into the central brain around the medial edge of the LOP. This strategy quickly found the HS and VS cells and several others, yielding over half of our catalog. The remaining neurons were found using one of two strategies: 1) we endeavored to find all neurons identified in a parallel effort searching for LPT neurons in stochastically labeled confocal images of neurons expressed by GAL4 driver lines (see below and Figure 1B). These cells were matched by using any useful anatomical references from the confocal stacks, including proximity to already reconstructed neurons, cell body and axon tract locations. (2) we surveyed large axons that were near the axons of already reconstructed LPT neurons. Strategy (1) was critical for finding most of the novel ipsilateral neurons and strategy (2) was useful for completing the inventory of bilateral neurons. While we carried out some manual reconstructions in the left hemisphere using symmetry to guide the search, this survey was not nearly as complete. As the project evolved, we primarily confirmed the existence of symmetric neurons using the FAFB data segmented in the FlyWire environment (described below). We did not identify any new LPT neurons on the left side. In other words, our initial manual reconstruction and matching to light strategy appears to have found all (or very nearly all) the LPT neurons. We spent approximately 6 person years to search for and reconstruct all 116 right and left hemisphere LPT neurons, placing a total of 1,592,248 skeleton nodes (only counting these neurons) in CATMAID.

The morphological completion status of the traced LPTs, and the synapse tagging for these neurons vary. Our initial goal was to completely reconstruct all LPT neurons and tag all their pre- and post-synapses, but this proved to be an unreasonable amount of work for our team. We then focused on the right-side LPT neurons and adopted a set of more practical goals: Central brain axon terminals were traced and all synapses tagged to completion, while dendrites, with expansive and often dense arbors, initially only had their main branches traced. Once the main branches were traced, the thinner and more dense arbors were traced on an ad hoc basis – we required the final branches to be space-filling with respect to the resolution of the LOP columns. In other words, we endeavored to have no “holes” in the LOP layer occupancy estimates used to construct the PMPMs. In our final data set, many neurons do have their entire dendrites traced to completion. Our treatment of synapses will be discussed in the next section. We also used a recent auto-segmentations of the same data set, FAFB-Seg (Li et al., 2020) to quickly examine many auto-segmented fragments for neurons of interest. Once a fragment of interest in FAFB-Seg was found, it was imported to CATMAID, followed by manual tracing and identity confirmation. Instead of extensive manual reconstructions of the left-side LPTs, we matched LPT neurons from FAFB-CATMAID to FlyWire (Dorkenwald et al., 2023)—during these initial matches, many neurons in FlyWire required substantial proofreading. With the help of FlyWire, we also quickly found several left-side neurons that we had not found in our limited search, but again, no new LPT types were found (thus far). The left-right comparison, both morphology and neurotransmitter type, is reported in Figure 3—figure supplement 1 and 2, helped confirm our cell typing and identification.

To measure the diameter of the VS Neuron processes (Figure 3—figure supplement 1A), we made a series of measurements in CATMAID (using the ‘x’ key) to capture the membrane-to-membrane distance from representative z-section. For axons, the first measurement was taken from a z-section in which the soma tract branches off, and 4 additional measurements were approximately evenly spaced until the fifth measurement, which was taken just before axon terminal starts to branch (or exhibits boutons). For the dendrites, we took three measurements from nearby z-section at the thickest point of the main dendritic branch.

Summary of the contributors to LPT reconstruction in CATMAID (>=10000 Nodes):

| CATMAID user name | First Name | Last Name | # Nodes | % Traced |
| --- | --- | --- | --- | --- |
| laughlandc | Connor | Laughland | 410995 | 25.81 |
| ludwigh | Henrique | Ludwig | 197968 | 12.43 |
| thomsona | Alex | Thomson | 151624 | 9.52 |
| gezahegnb | Bruck | Gezahegn | 147141 | 9.24 |
| ghorbanip | Padideh | Ghorbani | 140028 | 8.79 |
| moranc | Cullen | Moran | 116396 | 7.31 |
| flynnm | Miriam | Flynn | 98209 | 6.17 |
| erginkayam | Mert | Erginkaya | 71557 | 4.49 |
| gillisz | Zachary | Gillis | 66589 | 4.18 |
| polskyj | Jason | Polsky | 32017 | 2.01 |
| chois | Soonhong | Choi | 27918 | 1.75 |
| smithn | Aidan | Smith | 26883 | 1.69 |
| allredg | Gabrielle | Allred | 20188 | 1.27 |
| wolskyd | David | Wolsky | 13153 | 0.83 |

We also reconstructed hundreds of T4 neurons, some of which appear in a previous study (Zhao et al., 2022). They can be assigned to two overlapping groups for different purposes and with different reconstruction criteria: (1) morphology-complete and (2) axon-complete. The details for group (1) are described in “Extending the eyemap to the LOP” and for (2) are described in “Lobula Plate layers” sections below.

### Synapse Sampling of Neurons in CATMAID

As many LPTs make upwards of 10k synaptic connections, it was impractical to annotate all of them, so we used sampling methods to discover the strongest connections to and from these cells. Over the course of the project, LPTs had their synapses sampled via both a “passive” method and a more direct method, accounting for why some LPTs have more tagged connections than others. The passive method is that when proofreaders were engaged in exploratory reconstructions and would encounter connections (pre or post synaptic) with traced LPT neurons these would generally be tagged. This meant that the synapses we tagged via this method were biased towards particular regions of the LOP where more exploratory tracing was carried out. The direct method relied on the CATMAID widget ‘reconstruction sampler’, which segments a traced skeleton into intervals of pre-defined length, and then randomly select a portion of these intervals for manual examination. Our protocol was to tag all synapses within a segment presented to us by the sampler widget. We would also complete the tracing of any unfinished fine branches on the sampled segments. We used this method for all right-side LPT neurons, where we tagged synapses in 5% of the intervals for VS and HS neurons, and 10% for the rest. These methods provided an excellent foundation of connectivity to both identify strong connections between LPTs (Figure 5) and to capture the LOP layer occupancy of the dendrites (Figure 2).

### Extending the eyemap to the LOP

Following our prior work (Kind et al., 2021; Zhao et al., 2022) we defined medulla columns using the columnar cell type, Mi1, which spans the entire depth of the medulla, is easy to identify, and is a primary input to T4 cells (Takemura et al., 2013). To extend the eyemap of the medulla, we needed to map visual coordinates from the input space of the medulla to the output space of the LOP, which conveniently are connected by the abundant, small-field T4 neurons. We defined T4 medulla positions as the centers-of-mass of their dendritic arbors, and T4 LOP positions as the centers-of-mass of their axon terminals. For this correspondence, we required T4 neurons to be reconstructed with complete dendrite and axon morphology. Building upon earlier work, we completely reconstructed 171 T4b neurons across the whole medulla, ensuring that there is a neuron per every 2-3 columns. Using these sets of points in both neuropils, we then defined column positions in the LOP using kernel regression, which mapped positions from one region (medulla) to another (LOP) while preserving their spatial relationship.

### Lobula plate layers

We defined LOP layers by fitting polynomial surfaces to the axon terminals of T4 neurons. For this purpose, we reconstructed T4 neurons of all 4 types, covering the entire LOP (112 T4a, 187 T4b, 105 T4c, and 139 T4d) with complete axon morphology but many with incomplete dendrites. We manually labeled a node in the T4 neuron skeleton as “axon start”, the axon segment distal to this point showed clear arborization, with pre-synaptic sites, in the LOP. We then collected all the terminal nodes for each T4 subtype (example in Figure 2—figure supplement 1A) and fit a polynomial surface to each. To avoid over-fitting, we used half the nodes from each subtype to generate surfaces of various polynomial orders and then assigned the other half to these 4 surfaces based on the shortest distance (Figure 2—figure supplement 1D). We chose the 5^th^ order polynomial surface for all subsequent analysis. These two steps (eye map extension and layer assignment) essentially maps all retina positions and the subtype-specific local motion directions to each compartment in the LOP. There are in total 3056 compartments = 764 columns × 4 layers.

### Predicted Motion Pattern Map (PMPM)

To estimate the motion responses of each LPT, we use the detailed morphology of the neuron, together with the T4 mapping for space and motion directions. To assemble these estimates, we needed to assign a subset of the 3056 LOP compartments to each LPT neuron (for which we could confirm connections to at least 5 T4 neurons) based on spatial proximity between the compartments and the dendrites. We noticed that there were generally few synapses on the primary dendrite while many on the smaller and higher-order dendrites. So we employed the Strahler Number (SN) (Strahler, 1957) to classify all dendritic segments, which made up the skeleton, into groups: SN = 1 group contained the terminal segments (leaves in a tree graph) and larger SN generally contained larger segments closer to the primary dendrite. We wanted to know which SN groups can best capture the spatial distribution of synapses. Based on the ∼40000 manually tagged LOP synapses on right-side LPTs, we found that ∼98% of them fell within 1.5 µm of the skeleton (Figure 2—figure supplement 1E), and more than 95% within 1.5 µm of the first three SN (=1,2,3) groups (Figure 2—figure supplement 1F). We chose not to use dendritic segments with higher SN to avoid overestimating the size of dendritic field. Next, for each LPT neuron, we performed the following operations:

1. Collect all dendritic segments within the LOP, with SN ≤ 3.
2. Assign nodes in each segment to the closest LOP layers.
3. If more than 90% of the nodes are assigned to a single layer, then that segment is assigned to that layer. This step resolves the layer assignment inaccuracies that come from “vertical” segments that pass through layers which are usually “bridges” between dendritic fields in different layers.
4. We then have a list of nodes with layer and column assignments for each neuron, and we then simply count the number of nodes in each compartment of the LOP.
5. Since layer 1 and 2 have the opposite motion direction tuning, at any location where the counts in these layers are non-zero, we set the smaller one to be zero. Similarly for layer 3 and 4 node counts.
6. We then apply a spatial median filter to assign each retina position the median of 1 + 6 neighboring positions (itself + nearest neighbors in a hexagonal grid). This is done separately for the compartments in each layer.
7. Finally, we add up the T4 preferred directions weighted by the compartment counts for each retina position to construct the PMPM for each LPT neuron.

The illustrative example is in Figure 2, but this method is used throughout the manuscript.

### Optimal motion axes

Given each LPT’s PMPM, we computed the optimal motion axes in two ways. The first approach is a brute force sampling (already introduced in (Zhao et al., 2022)) where we defined 10356 directions (roughly 2 degrees steps) on a unit sphere. For a given LPT neuron, we generated ideal optic flow vectors induced by rotation along a sampling direction, at all ommatidia directions (right eye only) that fall within the neuron’s field of view. We then compared this optic flow field with the neuron’s PMPM to compute the average angular difference, α_R_ (see Figure 2J). Out of the 10356 sampling directions, the one with the minimal α_R_ (= α_R0_) is taken as the optimal rotation direction for this LPT neuron. Similarly for translation with α_T_. The errors for the entire sphere of sampled axes is represented as Mollweide projection color-maps in Figure 2K and also in Figure 3, 4, and 6.

An alternative approach accounts for the possibility of optic flow generated by observer motion that combines rotation and translation. Every body motion can be decomposed as a rotation and a translation. The optic flow vectors depend on the rotation and translation directions, as well as their relative amplitudes (angular speed and translational speed). Another complication results from the fact that the amplitude of the vectors in a translational motion induced optic flow field depends on the distance to the fiducial markers (the visual features without which there would be no optic flow). We treat ommatidia directions as points on a unit sphere S_1_ (with radius R_1_ = 1).

1. Starting from the 4^th^ equation in (Koenderink and van Doorn, 1987), the optic flow vector 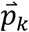 at the k^th^ viewing direction 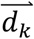 is:

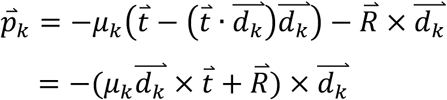

where the second step uses the identity: *a* × (*b* × *c*) = *b*(*c* · *a*) − *c*(*b* · *a*). Here *μ_k_* is the “reduced nearness” which is the ratio of translation speed (*T*) over the distance to the fiducial marker; 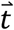 is a unit vector denoting the translation direction; 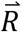 is the rotation vector (both direction and amplitude).
2. Ommatidia viewing direction can be expressed in the spherical coordinates (*ρ* = 1, *θ*, *φ*):

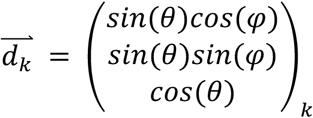
3. 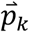 and 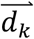 are orthogonal. We can define a 2D orthonormal basis on the 2D unit sphere surface:

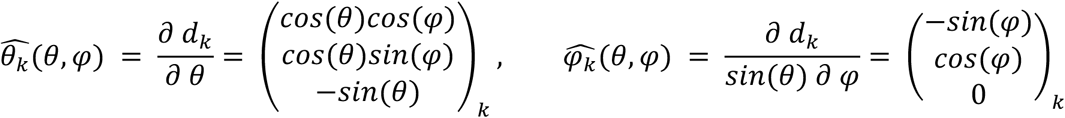
4. Note that 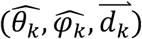 form a right-handed orthonormal basis. The *k^th^* optic flow vector can be written in the 2D orthonormal basis as 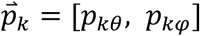:

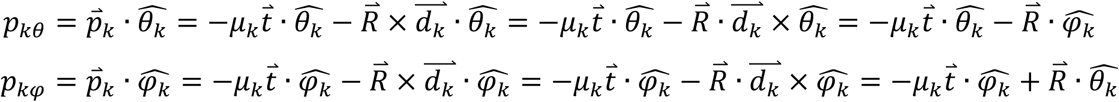
5. We shall make 2 simplifications here: since there is no explicit visual stimulus in our case, we shall assume that all fiducial markers are at equal distance. Indeed, their positions are defined as the intersections between within-field-of-view ommatidia directions and a sphere S_2_ of radius R_2_ that’s concentric with S_1_. This assumption implies *μ_k_* = *μ*. The second simplification is to ignore the amplitude of flow vectors and to only consider their angles, because it’s unknown how local dendritic activity correlates with local motion speed. Now our problem reduces to a non-linear regression problem solving for 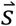:

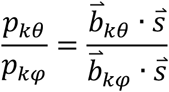

where

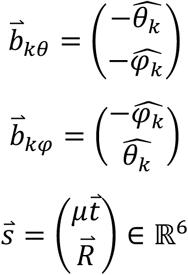
6. A final note on physical constraints. There are 2 free scales in 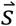: one is an overall scaling factor since we ignore amplitudes of translation and rotation vectors. The other lies in the coupled variables: translation speed *T* and fiducial marker distance R_2_, that only show up as a ratio, 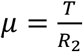. For *Drosophila*, The translation speed can reach up to 1 m s^-1^ in free flight (David, 1982). The fly’s head is about 1 mm in diameter. However, since the fly head is not an exact sphere, the representation of ommatidia directions as points on a unit sphere is valid for distances much larger than the fly head. So we set an ad hoc lower limit, R_2_ ≥ 10 mm. Therefore, *μ* ≤ 100 *s*^−1^. Flies have been observed to make rotational saccade up to 90° within 50 ms (Fry et al., 2003) in free flight, which is about 30 rad s^-1^. The solution 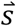 can be scaled within these 2 constraints.

### Flywire Proofreading

Our proofreading of neurons in FlyWire was almost entirely focused on the LPTs we had already identified in CATMAID and a several months search for matching cells on the left side of the brain. Our main contributions were merging and splitting of the auto-segmented fragments. There were several bodies that required splitting from one another, as their large-diameter axons were heavily merged (known by proofreaders as “franken-bodies”). These included more than half of the right and left hemisphere VS cells, as well as the left hemisphere HS cells. No other LPTs required such significant effort.

### Matching LPT neurons Between FAFB and Hemibrain

We mainly relied on central brain morphology to match LPT neurons between FAFB and the Hemibrain data set, since most of the optic lobe wasn’t available in the Hemibrain (Scheffer et al., 2020). We first focused on the large cell bodies near the medial part of the LOP (only a sliver is present). Once this strategy was exhausted, Region of Interest (ROI) queries were made to find most of the remaining LPT neurons. This involved first identifying which brain-region ROIs their axon terminals make synaptic connections, and then querying all neurons in these ROIs in Hemibrain for further comparison. We found 1-to-1 matchings for 56 out of 58 right-side neurons (but note that we matched LPT51 to 2 hemibrain neurons) and 17 left-side neurons (matches shown as Supplementary File 4). We also compared LPT-LPT connections in both data sets to confirm the matching (Figure 5—figure supplement 1B).

### Hemibrain Proofreading

We performed additional proofreading to further improve the connectivity data for the matched LPTs in the Hemibrain data set. Our goal was to achieve a relatively consistent level of completeness for the LPTs. The updated Hemibrain data (also including other proofreading efforts) will be included in a future data release from the FlyEM team. Our proofreading followed a 2-step process:

- Jan 2021: sparsely traced 58 LPT neurons to ensure that there were no major branches missing, with mostly minor fragments added.
- Feb - August 2021: proofreading using FlyEM’s “orphan link” workflow for LPT neurons

- Reached a completeness of 40% for a few “priority neurons”.
- Reached a reduced completeness of 30% for the remaining neurons that required substantial tracing effort. More specifically, if a neuron required >1000 orphan merges to hit 40% completion, we reduced the target completion to 30%

Figure 5—figure supplement 1A shows the improvement of percentage completion between January and August 2021. The completion percentage is calculated as the ratio of the total number of outgoing connections to neurons with the status of “traced” over the total number of output synapses (and reported as a percentage).

### Visualization of LPTs neurons across data sets

For all the renderings of neurons in the supplemental gallery images (Supplementary Files 2, 3, 4), we used the R package “natverse” (Bates et al., 2020), which provides 3D rendering and enables overlaying neurons from different brains. Plotting reconstructed neurons from FAFB and Hemibrain is enabled by the registration of each data set to a common template brain (Bogovic et al., 2020).

### Split-GAL4 lines

Construction of split-GAL4 lines for MeLp2 (SS02082: VT043284-p65ADZp in attP40; VT000926-ZpGDBD in attP2) and LPi2b (SS53141: R91C05-p65ADZp in attP40; VT049130-ZpGDBD in attP2) was as previously described for other optic lobe cell types (Davis et al., 2020; Wu et al., 2016). Split-GAL4 screening and characterization was done with the help of Janelia’s FlyLight Project Team. Driver lines will be made available at https://splitgal4.janelia.org/

### Light microscopy images (Figure 1B)

We used MultiColorFlpOut, MCFO (Nern et al., 2015), to visualize single LPT neuron morphologies. The GAL4 driver lines used for the cells shown in Figure 1B are listed in Table 1. Samples were prepared and imaged by the Janelia FlyLight Project Team following established protocols (https://www.janelia.org/project-team/flylight/protocols). All image stacks used were recently made publicly available as part of a large collection of MCFO images (Meissner et al., 2023). MCFO-labeled LPT cells were displayed using VVD viewer (https://github.com/takashi310/VVD_Viewer). This process involved manual segmentation to remove additional labeled cells and background signal. Original confocal stacks are available online as part of the Janelia FlyLight MCFO collection, with details provided in the caption of Table 1.

**Table 1:** GAL4 driver lines used to produce images of labeled LOP neurons. All images in Figure 1B show manually segmented cells from confocal stacks that are part of the public Janelia MCFO image collection (Meissner et al., 2023) and available online at https://gen1mcfo.janelia.org/cgi-bin/gen1mcfo.cgi. The individual original image stacks can be directly accessed using links formatted as follows: https://gen1mcfo.janelia.org/cgi-bin/view_gen1mcfo_imagery.cgi?slidecode=slidecode. For example, for the first image in the table this would be: https://gen1mcfo.janelia.org/cgi-bin/view_gen1mcfo_imagery.cgi?slidecode=20140313_32_D7.

| Figure Panel | Driver line | slidecode |
| --- | --- | --- |
| Figure 1B - LPT21 | R73C02 | 20140411_21_B6 |
| Figure 1B - LPT23 | VT009671 | 20131017_33_B6 |
| Figure 1B – LPT29 | VT009671 | 20131017_33_A4 |
| Figure 1B - LPT30 | VT004726 | 20140306_31_C5 |
| Figure 1B – LPT38-Nod1 | VT026759 | 20131024_32_H2 |
| Figure 1B - LPT45-dCal1 | VT026927 | 20140313_32_D7 |
| Figure 1B - LPT50 | VT026477 | 20140305_33_F4 |
| Figure 1B – LPT51 | R67D09 | 20130624_2_C4 |
| Figure 1B - LPT57 | VT049340 | 20140307_33_E4 |
| Figure 3-S2B,C - LPT55-MeLp2 | SS02082 | NA |
| Figure 3-S3A – Lpi2b | SS53141 | NA |

### Neurotransmitter Predictions (Figure 3—figure supplement 2)

As described in the “FlyWire Proofreading” section above, we identified and edited the LPT neurons in FlyWire to confirm right-left side matches and also to obtain access to automatic synapse predictions (Buhmann et al., 2021) and the neurotransmitter predictions (Eckstein et al., 2020) for each of these synapses. We use two main components of the synapse predictions: first, a quality score called the cleft score (Buhmann et al., 2021) for each identified synapse, and second, a per-synapse prediction as one of six transmitter types: dopamine, octopamine, serotonin, GABA, glutamate, and acetylcholine (Eckstein et al., 2020). Following a common recommendation from members of the FlyWire community, we used the transmitter predictions for all synapses with a cleft score of 50 or higher. The neurotransmitter predictions for each neuron are determined by simply adding up all the per-synapse predictions and taking the most frequent prediction (Figure 3—figure supplement 2A). Since many LPTs have very few pre-synapses in the optic lobes (Shinomiya et al., 2022), we used the central brain synapse transmitter predictions for most neurons and the optic lobe synapse predictions for the LOP-input LPTs (with a simple spatial cut-off to separate OL from the central brain). We found matching transmitter predictions for the left- and right-side instance of nearly all LPT neurons with 2 exceptions discussed in the text.

### Expansion-assisted iterative fluorescence in situ hybridization (EASI-FISH)

EASI-FISH detection of transcripts for VGlut, ChAT and GAD1, plus a nuclear marker (DAPI), was as described (Eddison and Ihrke, 2022). Members of the Janelia Project Technical Resources team (Mark Eddison, Yisheng He) performed the EASI-FISH experiments. We used Fiji to examine the images, adjust brightness and contrast and select sections for display. Newly generated split-GAL4 driver lines, described above, were used to label MeLp2 (Figure 3—figure supplement 2B) and LPi2b (Figure 3—figure supplement 3) cells in these experiments. Candidate HS and VS cell somata were identified by their size and location in the lobula plate cell body rind and are compared with the somata of the reconstructed counterparts in FAFB (Figure 3—figure supplement 3B).

### Cosine Similarity Analysis (Figure 3—figure supplement 1C)

We compared the LPT-LPT connectivity to verify our left-right pairing of LPT neurons (equivalently, their identification). We first collected all the synapse predictions in FlyWire for connections between left and right LPT neurons, discarding all synapses with a cleft score of <50 (Buhmann et al., 2021). Cells with more than one per type, as well as the HS neurons and 3 subgroups of VS neurons were grouped, and the connection weights for all neurons within each group were averaged. These groups are all indicated with (N = # of grouped cells) in the labels of Figure 3—figure supplement 1C. Each LPT neuron’s connection weights to all the other LPT neurons was ordered into a vector, taking into account the left-right symmetry. The cosine of the angles between these vectors were calculated for all left-right pairs (Figure 3—figure supplement 1C). Here smaller angles have a larger cosine similarity (max of 1) and indicate more similar connectivity patterns.

### LPT-target neuron network

We were able to find 74 neurons in Hemibrain that match our manually reconstructed LPT neurons in FAFB (Supplementary File 4). We queried their direct downstream partner and constructed a connectivity graph of 1246 nodes and 52385 edges. We grouped together LPT-target neurons of the same type given by Hemibrain annotation, with a new edge weight taking the average value. We then removed the LOP-input LPT neurons that receive few T4 inputs, and removed edges that either have a weight of less than 10 or contribute to less than 10% of total incoming connections to a given LPT-target neuron. Additionally, we removed single LPT-target neurons that receive input from a single LPT neuron, such that all remaining LPT-target neurons receive inputs from at least 2 LPTs. The resultant graph has 66 LPT-target nodes (cell types), 4 of which receive only inhibitory LPT inputs and are the basis for the analysis in Figure 5, and are further detailed in Supplementary File 5.

### Pairwise collaboration matrix and spectral clustering (Figure 6—figure supplement 2)

For each pair of LPT neurons, say, A and B, we examine their direct common targets in Hemibrain, using set of LPT-targets described above. The total connection weight from A to these targets is assigned as the value in the collaboration matrix’s row A and column B, and the total weights from B are the entry in row B and column A, resulting in an asymmetric matrix (Figure 6—figure supplement 2B).

Spectral clustering is a popular method that group nodes together based on their “affinity” (Ng et al., 2001). Here we define the affinity between neuron A and B as their average collaboration value, that is, we symmetrize the pairwise collaboration matrix and set its diagonal to 0. The values in the symmetric matrix represent how strongly any pair of LPT neurons are jointly connected to common targets (in the set of neurons considered for this analysis). Following the algorithm of (Ng et al., 2001), we constructed the symmetrized Laplacian matrix whose eigen spectrum (Figure 6—figure supplement 2C) has a “gap” after the 5^th^ largest eigenvalue. This suggests 5 clusters with high within-cluster affinities, which are used to group the cells in Figure 6—figure supplement 2D and shown spatially in Figure 6—figure supplement 2E.

### Code availability

All reconstructed neurons from the FAFB dataset will be made available at https://fafb.catmaid.virtualflybrain.org/. All code to carry out the analysis and generate the figures will be available at https://github.com/reiserlab/LPT-catalog

## Acknowledgments

The authors thank Gerry Rubin for supporting Aljoscha Nern and fly connectomics projects at Janelia, Janelia’s Fly Core for fly care, Mark Edison and Yisheng He of Janelia’s Project Technical Resources team for EASI_FISH experiments, Janelia FlyLight Project Team for help with preparation and imaging of light microscopy samples, Kit Longden and Stephen Huston for extensive discussions about LPTs and all members of the Reiser lab for helpful feedback, and David Wolsky, Gabrielle Allred, Aidan Smith, Soonhong Choi, Jason Polsky, Zachary Gillis, Miriam Flynn, Cullen Moran, Padideh Ghorbani, and Bruck Gezahegn for assistance tracing neurons in CATMAID, and Caroline Mooney, Audrey Francis, and Chelsea Alvarado for proofreading Hemibrain neurons. We thank the FAFB tracing community for supportive and open sharing of methods and data, especially Greg Jefferis, Marta Costa, Philipp Schlegel, and the “FAFB optic lobe working group”, especially the groups of Mathias Wernet, Marion Silies, and Rudy Behnia for collaborations and feedback. We thank Holger Krapp and Klaus Hausen for access to Hausen’s unpublished report on the LPTs of *Calliphora*. Development and administration of the FAFB tracing environment and analysis tools were funded in part by National Institutes of Health BRAIN Initiative grant 1RF1MH120679-01 to Davi Bock and Greg Jefferis, with software development effort and administrative support provided by Tom Kazimiers (Kazmos GmbH) and Eric Perlman (Yikes LLC). We thank Peter Li for sharing his automatic segmentation (https://doi.org/10.1101/605634). We thank the Princeton FlyWire team and members of the Murthy and Seung labs, as well as members of the Allen Institute for Brain Science, for development and maintenance of FlyWire (supported by BRAIN Initiative grants MH117815 and NS126935 to Murthy and Seung). We also acknowledge members of the Princeton FlyWire team and the FlyWire consortium for neuron proofreading and annotation, especially members of the Wernet, Linneweber, Borst, Jefferis, Murthy, and Seung Labs and the Eyewire citizen scientists, who contributed to the LPTs neurons (individual contributions detailed in Supplemental File 6). This work made use of VVDviewer, based on software funded by the NIH: Fluorender: An Imaging Tool for Visualization and Analysis of Confocal Data as Applied to Zebrafish Research, R01-GM098151-01. This work is funded by the Howard Hughes Medical Institute through its support of the Janelia Research Campus. This article is subject to HHMI’s Open Access to Publications policy. HHMI lab heads have previously granted a nonexclusive CC BY 4.0 license to the public and a sublicensable license to HHMI in their research articles. Pursuant to those licenses, the author-accepted manuscript of this article can be made freely available under a CC BY 4.0 license immediately upon publication.

---

**Figure 2—figure supplement 1:**
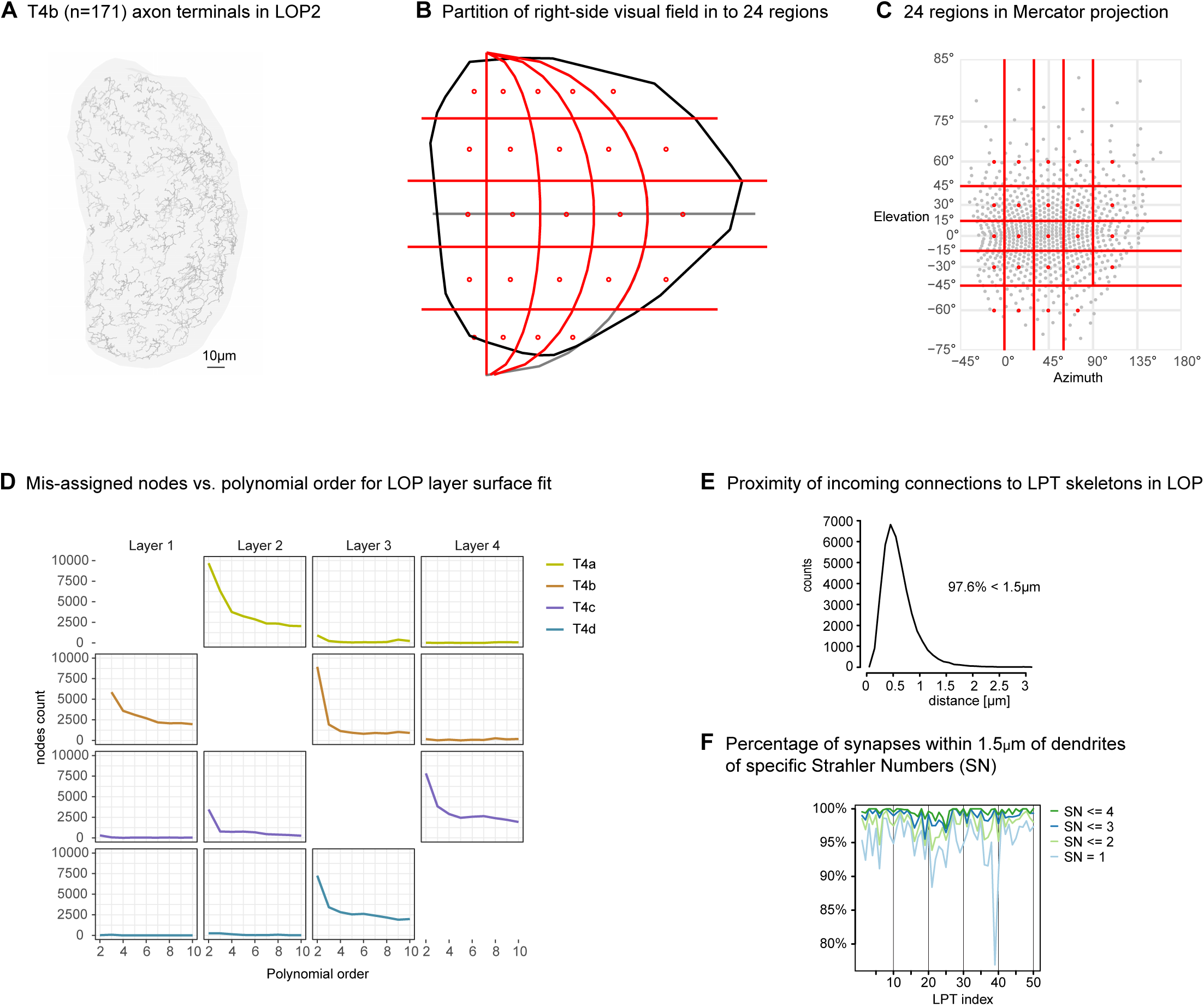
Analysis details supporting the computational predictions of LPT PMPS. **A.** EM reconstructed T4b axon terminals in LOP2. **B** and **C.** To present down-sampled PMPMs, the field of view of the fly’s right eye is divided into 24 regions. The average preferred direction in each region is represented by arrows in the PMPMs, whose locations are denoted by red dots here. This averaging is carried out for all flow field plots in this manuscript. Here we show both Mollweide and Mercator projections. **D.** For the analysis presented in the manuscript, the LOP layers are constructed as a polynomial surface fit to the axon terminals of each T4 subtype. We use 5^th^ order polynomials, and evaluated the quality of assignments of nodes in T4 axon terminals to layers to avoid over-fitting. The mis-assigned nodes are plotted as a function of the order of the polynomial surface fit. **E.** Distance between annotated synaptic connector and the nearest node of the reconstructed neuron skeleton, for all LPT neurons described in this paper. Note that 97.6% of connectors fall within 1.5 µm of the skeleton. **F.** Percentage of synapses captured by including varying levels of Strahler branches of the dendritic tree, within this 1.5 µm distance, for the main LPTs described in this manuscript (except for the feedback neurons that are not major T4/T5 targets). Note that SN <=3 captures most synapses, using SN>=4 may lead to overestimate of the dendritic coverage.

**Figure 3—figure supplement 1:**
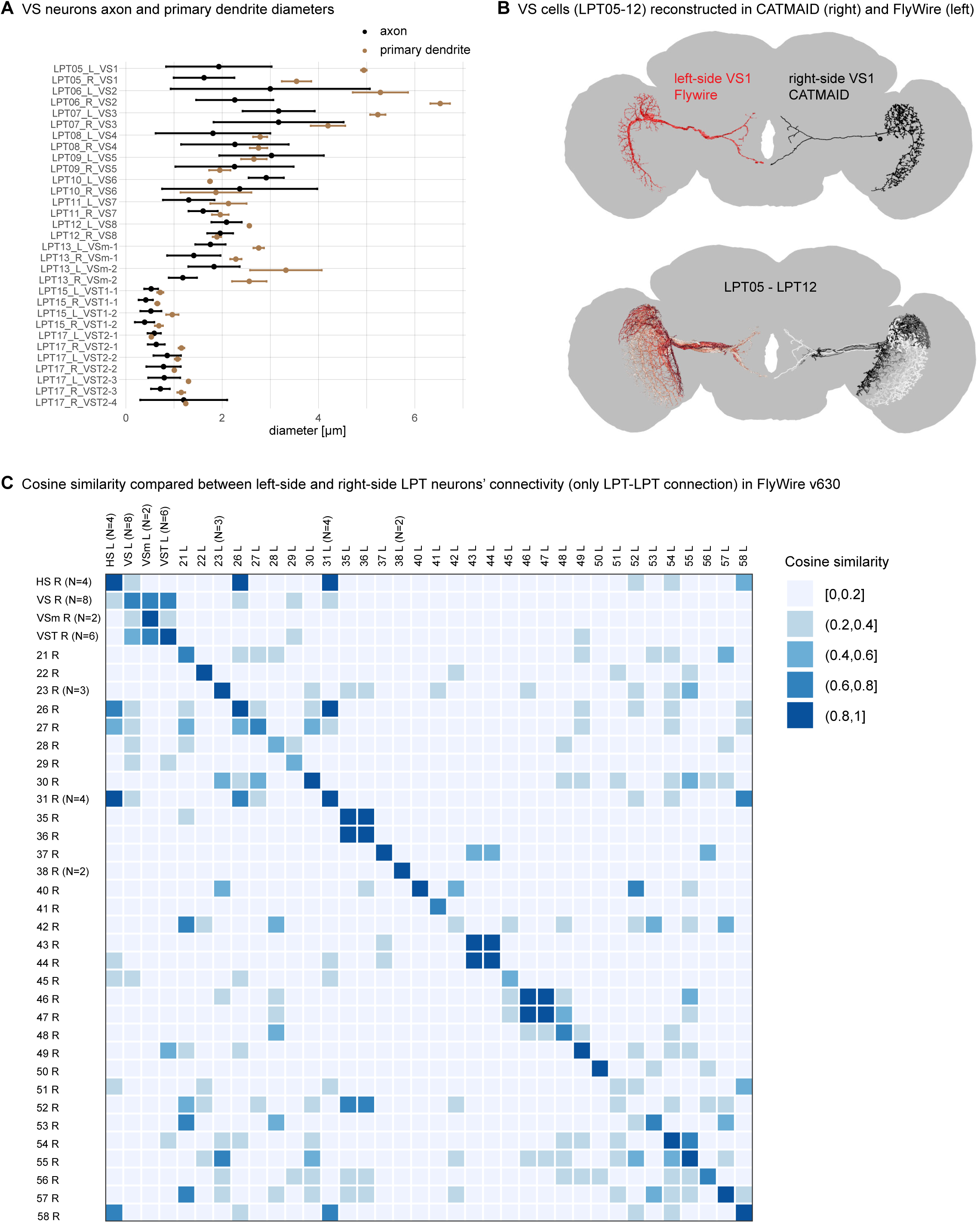
Supporting evidence for identification of LPT neurons. **A.** Axon and primary dendrite diameters for all VS neurons in FAFB. We note that 16 VS cells were eventually found on the right and left sides of the brain, but at the time these measurements were made, one of the LPT17 cells on the left side had not yet been located. **B**. Comparison of EM reconstruction of VS neurons, in the FAFB brain, left-side neurons from FlyWire (in shades of red) and right-side neurons manually reconstructed in CATMAID (in shades of black). **C.** Cosine-similarity comparison of left vs. right-side LPT neurons in Flywire. Some cells are treated as a group, as indicated, and the cosine similarity is computed on the group connectivity average. For this computation, an LPT neuron’s connectivity is characterized by its connection with all other LPT neurons only. All FlyWire data based on the March 2023 public release (version 630).

**Figure 3—figure supplement 2:**
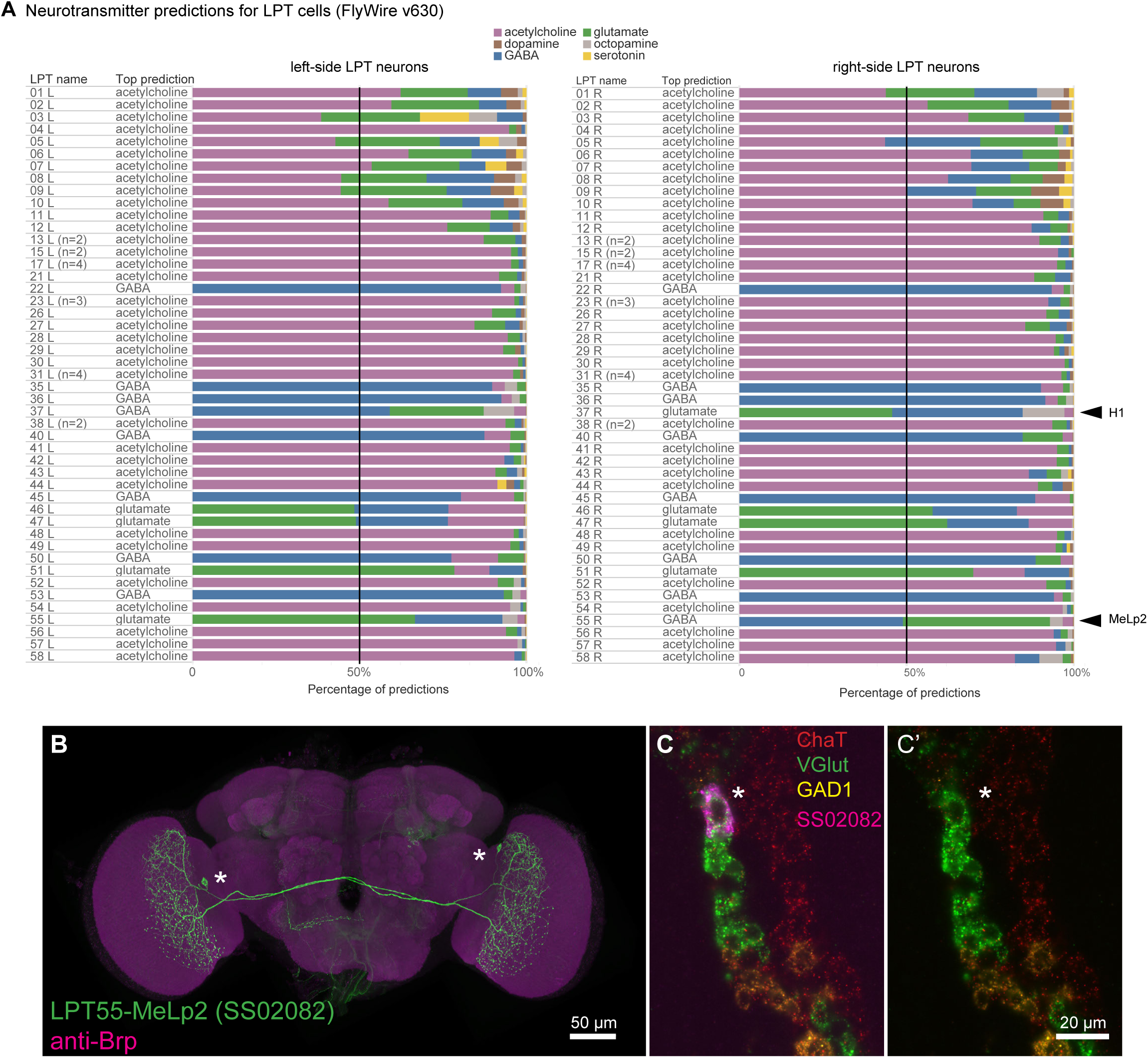
Neurotransmitter predictions for the LPT neurons. **A.** Neurotransmitter prediction for all LPT neurons, based on the method of (Eckstein et al., 2020) for the matched neurons in FlyWire. Only 2 cell types show right/left discrepancy in the top prediction, indicated with arrowheads. **B,C,C’.** Experimental evidence that MeLp2 neurons are glutamatergic. **B.** Expression pattern of a split-GAL4 line (SS02082) labelling LPT55-MeLp2 neurons. GAL4-driven expression of a membrane-targeted GFP is in green, a general neuropile label (anti-Brp) in magenta. The asterisks show the soma locations of the right and left MeLp2 neuron. **C and C’.** EASI-FISH labeling (Eddison and Ihrke, 2022; Wang et al., 2021) of transcripts indicative of cholinergic (ChAT; red), glutamatergic (VGlut; green) or GABAergic (GAD1; yellow) transmitter phenotypes. A MeLp2 cell body (indicated by the adjacent asterisk in both C and C’) is labeled in magenta in C. Images show different channel combinations of a single confocal section.

**Figure 3—figure supplement 3:**
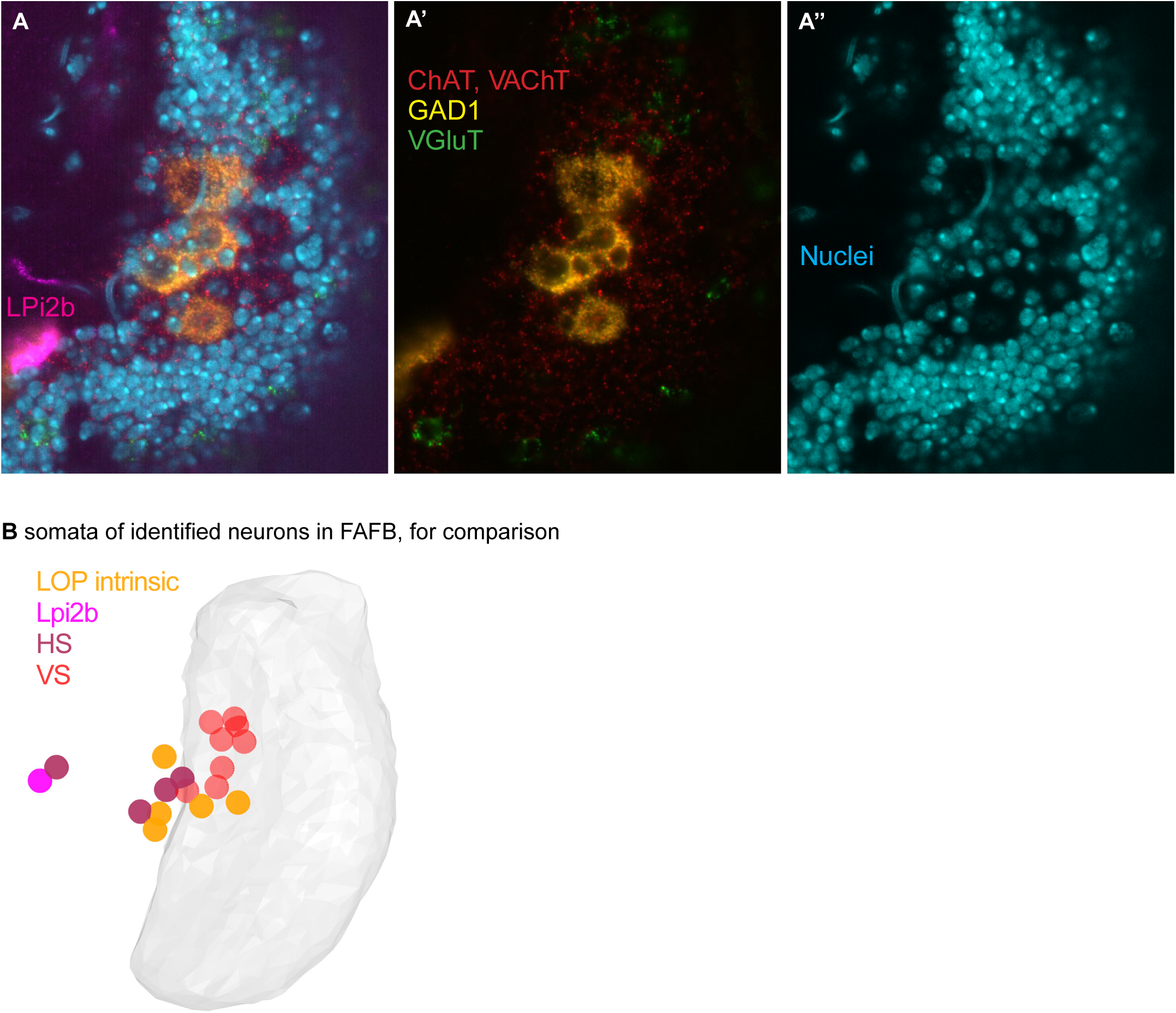
Additional supporting evidence for the neurotransmitter predictions. **A,A’,A’’.** EASI-FISH labeling (Eddison and Ihrke, 2022; Wang et al., 2021) of ChAT (red), VGlut (green) and GAD1 (yellow) in the lobula plate cell body rind, in the region where the HS and VS somata are found. Nuclei are also labeled (blue). An LPi2b neuron labelled by a split-GAL4 driver line (SS 53141) is shown in magenta. Images show different channel combinations of a single confocal section (A, all channels; A’ transmitter indicators only; A’’, nuclei only). Large cells in this region appear to be either GABAergic (GAD1 marker, orange) or cholinergic (ChAT marker, red). **B.** For comparison with (A), we use the soma locations from the FAFB LPT set: LPi2b (magenta), other large optic lobe intrinsic LOP neurons (orange; these include Am1, 2 LPi12, Lpi21, and an undescribed cell that appears to be a large layer four LPi), 4 HS neurons (maroon), and 8 VS neurons (red). The large optic lobe intrinsic neurons are only partially reconstructed and not included in the LPT survey reported in this study, but could be matched to known cell types (Shinomiya et al., 2022). Given that these are all the large cell bodies we found in this region, and several of the LPis are expected to be inhibitory, we suggest that the GABAergic cells in the EASI-FISH images are the large LPi and Am1 neurons and the cholinergic cells are the HS and large VS cells.

**Figure 4—figure supplement 1:**
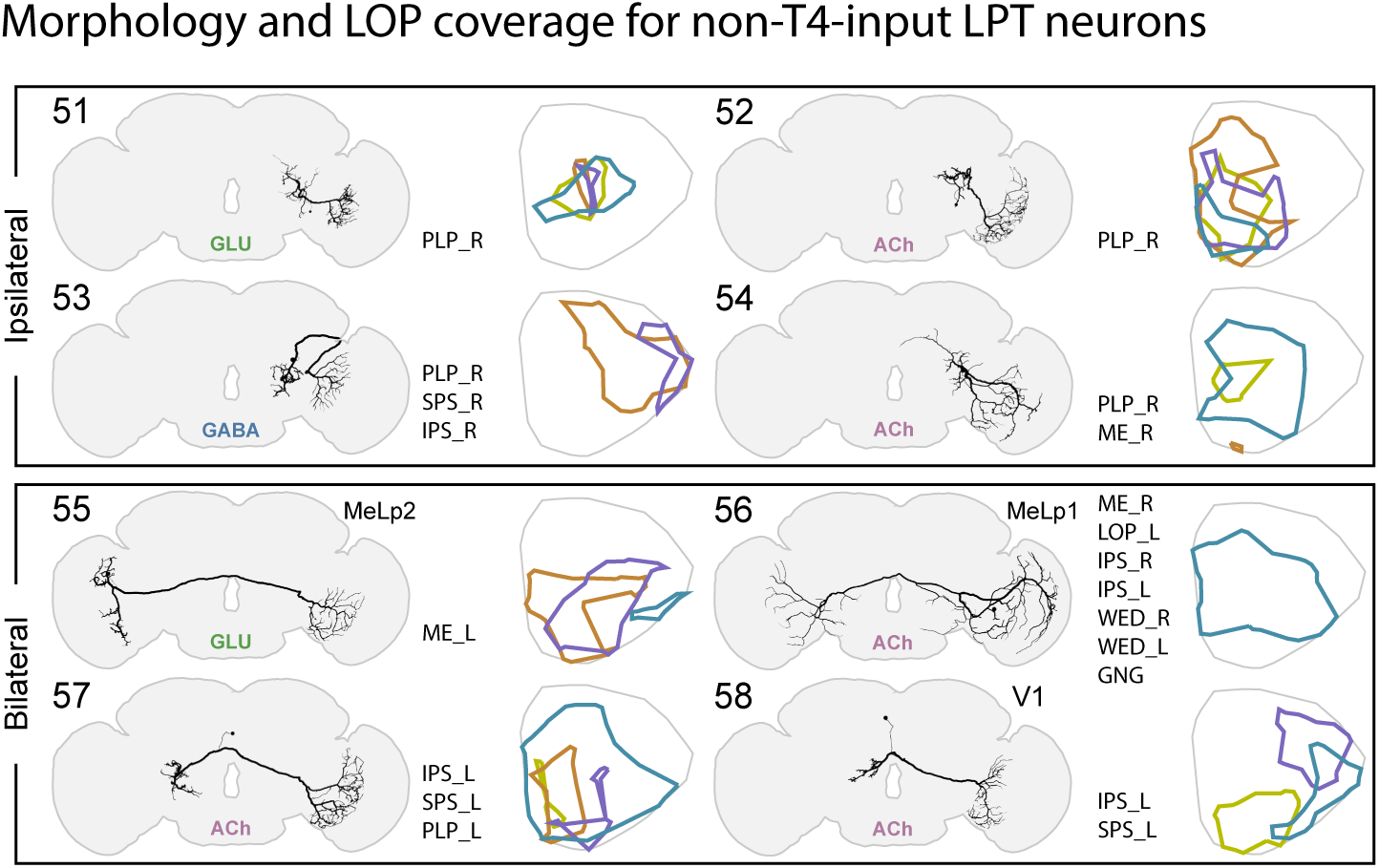
morphology and layer coverage of the LOP-input LPT neurons. The remaining LPT neurons receive little T4 inputs, instead they most likely provide inputs to the LOP, some as feedback from the central brain. As these neurons receive minimal T4 inputs, we do not predict their motion pattern maps.

**Figure 4—figure supplement 2:**
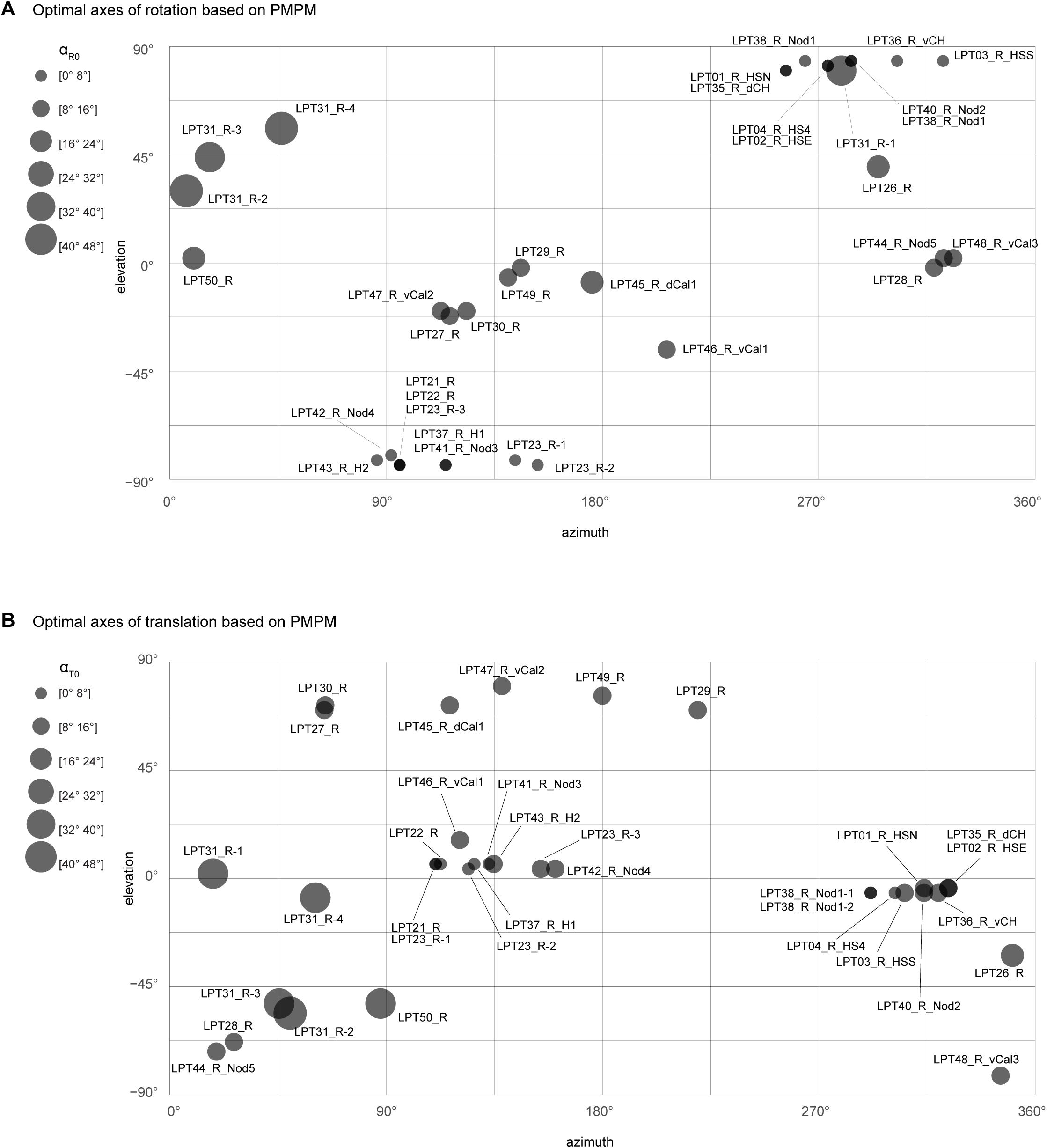
Optimal rotation or translation axes predicted by PMPMs. These two plots summarize the optimal rotation (top) and translation (bottom) axes for the LPTs of Figures 3 and 4, excluding the VS neurons, which are treated in Figure 3B. The size of each neuron’s marker indicates the size of the angular error.

**Figure 5—figure supplement 1:**
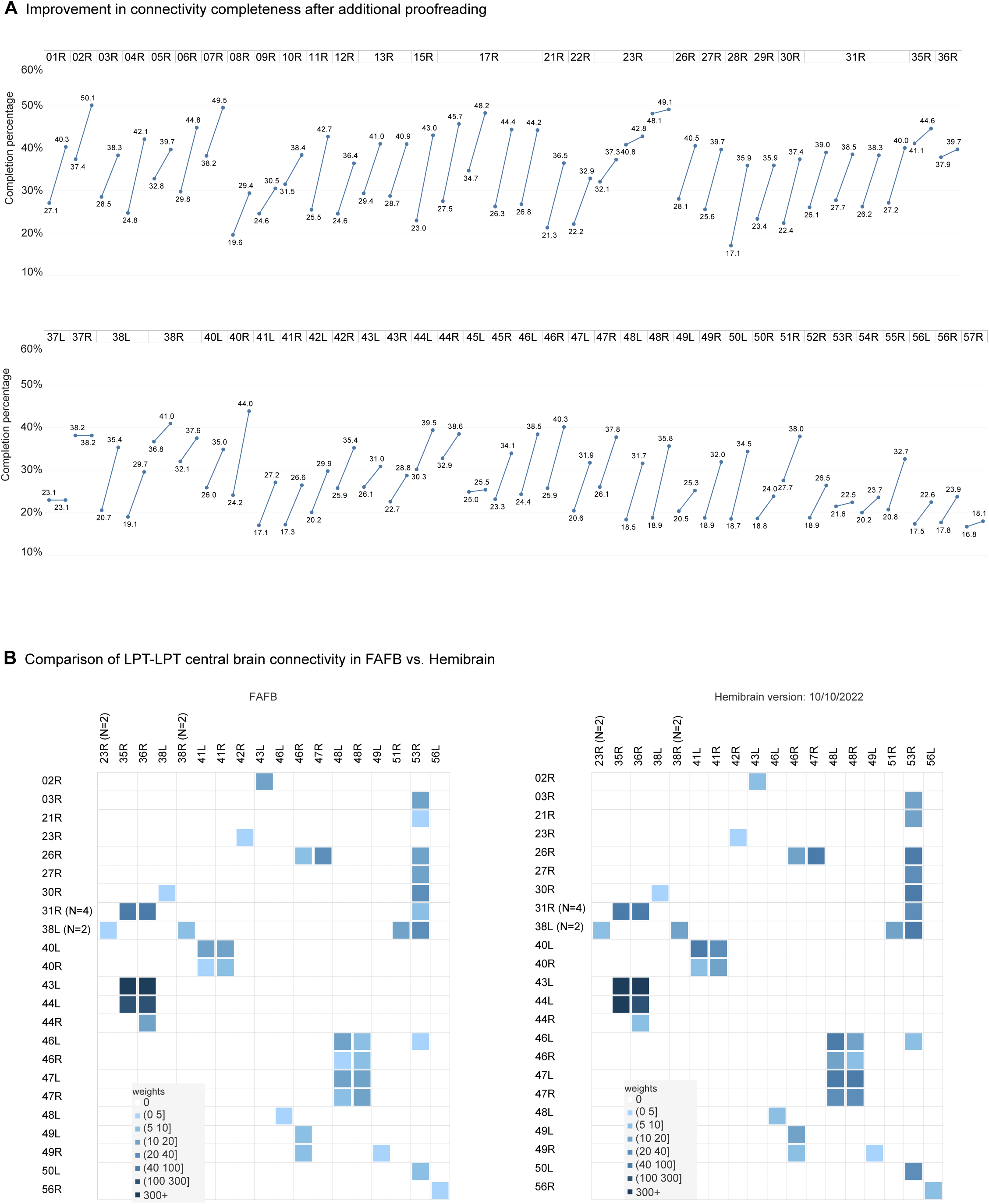
LPT connectivity in the Hemibrain connectome. **A.** Improvement in connectivity completeness for the matched set of LPT neurons, after manual proofreading in Hemibrain. **B.** Comparison of central brain LPT-LPT connectivity in FAFB (left) and Hemibrain (right) data sets.

**Figure 6—figure supplement 1:**
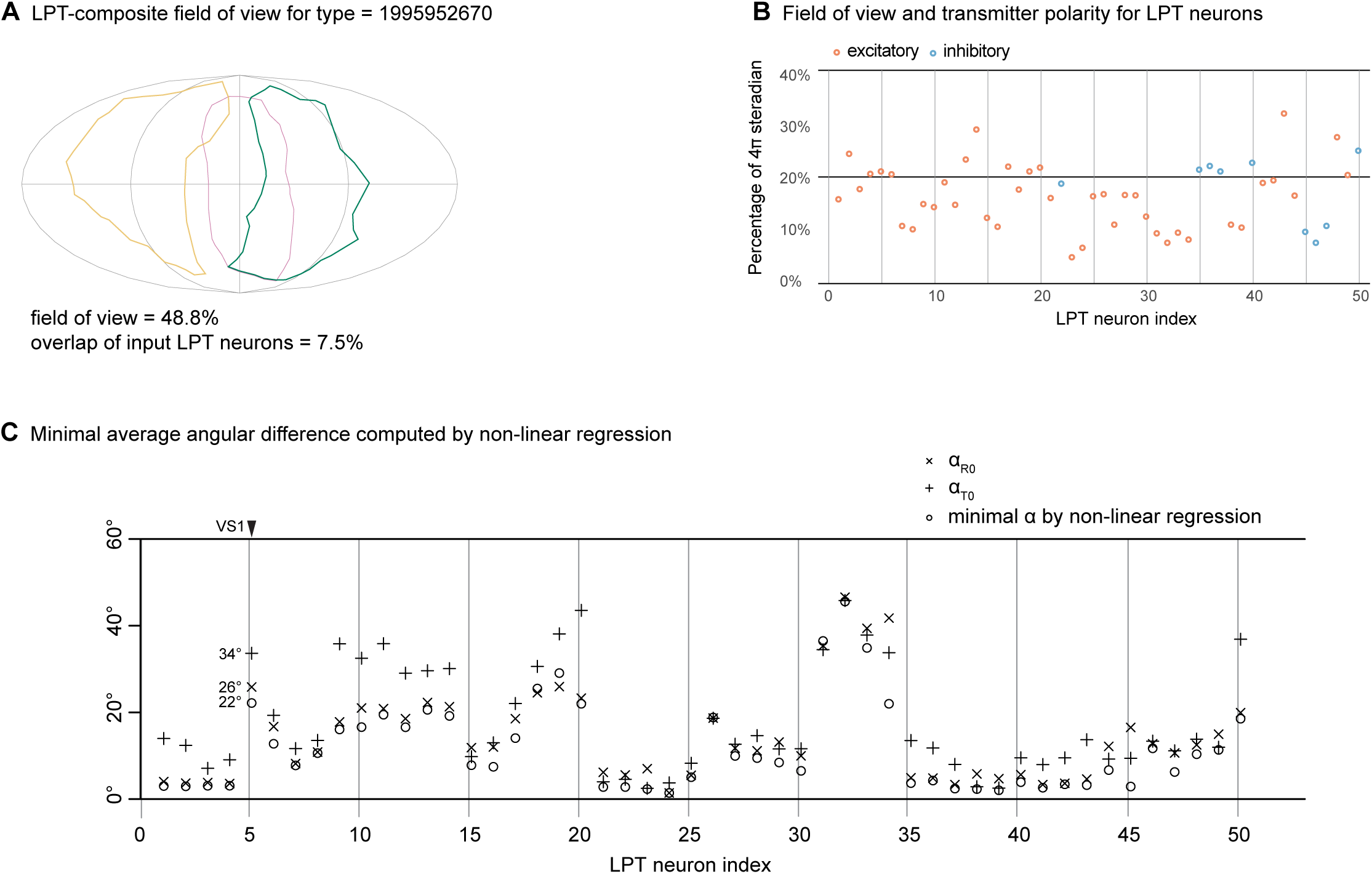
supporting data for LPT integration analysis. **A.** Composite field of view for the example LPT target neuron in Figure 6A. **B**. Fields of view and predicted synapse polarity for the LPT neurons (the indexing used throughout indicates the corresponding LPT neuron). **C**. Comparing the minimal average angular differences computed via the brute force method for rotation and translation separately, and a non-linear regression method treating them jointly (see Methods).

**Figure 6—figure supplement 2:**
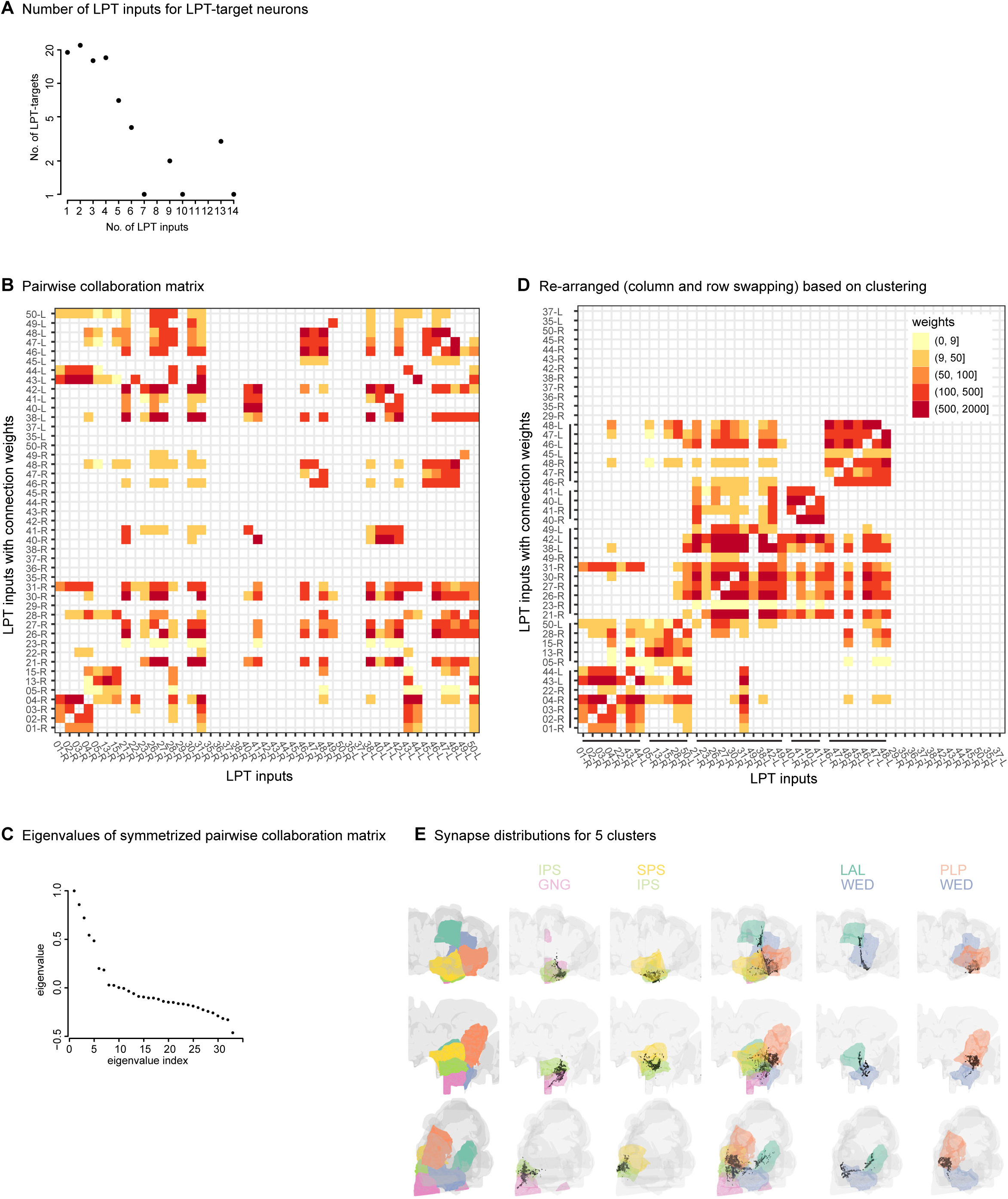
Patterns of LPT output integration in the central brain. **A.** Number of target cells vs. number of their LPT inputs for the central neurons considered in this analysis (based on Hemibrain connectivity). **B**. Pairwise “collaboration matrix” where each entry represents all the LPT-target neurons that receive inputs from the LPT neuron indicated in the row and the LPT neuron in that column. The matrix is asymmetric since the value of each entry is the total connection weight from the LPT neuron in the row. **C.** The eigen spectrum of the Laplacian matrix constructed from the pairwise collaboration matrix, showing a gap after the 5^th^ largest eigenvalue. This was used to select the number of clusters (=5) for the spectral clustering in C. **D.** The pairwise collaboration matrix re-ordered based on spectral clustering. There are 5 clusters within the connected graph. **E.** Synapse distributions and the relevant neuropils capturing the majority of synapses for each cluster in the central brain. The synapses shown are restricted to those between LPT neurons and their targets (i.e. those in the collaboration matrix).

**Supplementary File 1: Table of reconstructed LPTs, with R-L matches, and correspondences to matched neurons in FlyWire and Hemibrain.**

**Supplementary File 2: Gallery of all the right-side LPTs reconstructed in FAFB.**

**Supplementary File 3: Gallery of all right-side LPTs reconstructed in FAFB and all matched left-side LPTs from FlyWire.**

**Supplementary File 4: Gallery of LPT neurons overlayed with putative Hemibrain matches.** The LPT neurons reconstructed in FAFB are shown in red, and the matched Hemibrain neurons are shown in black (56 right side matches, note that LPT51 is matched to 2 Hemibrain neurons, and 17 left side matches).

**Supplementary File 5: Gallery of LPT-target neurons from Hemibrain, combined with estimated optic flow sensitivity.** Blue arrows indicate the PMPM contribution of inhibitory LPTs, while red arrows represent excitatory LPTs contributions.

**Supplemental File 6: Table of FlyWire LPT neuron proofreading contributions.**

## References

Ammer G, Vieira RM, Fendl S, Borst A. 2022. Anatomical distribution and functional roles of electrical synapses in Drosophila. Current Biology 32:2022–2036.e4. doi:10.1016/j.cub.2022.03.040

Bates AS, Manton JD, Jagannathan SR, Costa M, Schlegel P, Rohlfing T, Jefferis GS. 2020. The natverse, a versatile toolbox for combining and analysing neuroanatomical data. eLife 9:e53350. doi:10.7554/eLife.53350

Behnia R, Clark DA, Carter AG, Clandinin TR, Desplan C. 2014. Processing properties of ON and OFF pathways for Drosophila motion detection. Nature 512:427–430. doi:10.1038/nature13427

Bishop CA, Bishop LG. 1981. Vertical motion detectors and their synaptic relations in the third optic lobe of the fly. Journal of Neurobiology 12:281–296. doi:10.1002/neu.480120308

Bishop LG, Keehn DG, McCann GD. 1968. Motion detection by interneurons of optic lobes and brain of the flies Calliphora phaenicia and Musca domestica. Journal of Neurophysiology 31:509–525. doi:10.1152/jn.1968.31.4.509

Boergens KM, Kapfer C, Helmstaedter M, Denk W, Borst A. 2018. Full reconstruction of large lobula plate tangential cells in Drosophila from a 3D EM dataset. PLOS ONE 13:e0207828. doi:10.1371/journal.pone.0207828

Bogovic JA, Otsuna H, Heinrich L, Ito M, Jeter J, Meissner G, Nern A, Colonell J, Malkesman O, Ito K, Saalfeld S. 2020. An unbiased template of the Drosophila brain and ventral nerve cord. PLOS ONE 15:e0236495. doi:10.1371/journal.pone.0236495

Buhmann J, Sheridan A, Malin-Mayor C, Schlegel P, Gerhard S, Kazimiers T, Krause R, Nguyen TM, Heinrich L, Lee W-CA, Wilson R, Saalfeld S, Jefferis GSXE, Bock DD, Turaga SC, Cook M, Funke J. 2021. Automatic detection of synaptic partners in a whole-brain Drosophila electron microscopy data set. Nat Methods 18:771–774. doi:10.1038/s41592-021-01183-7

Buschbeck EK, Strausfeld NJ. 1997. The relevance of neural architecture to visual performance: Phylogenetic conservation and variation in dipteran visual systems. Journal of Comparative Neurology 383:282–304. doi:10.1002/(SICI)1096-9861(19970707)383:3<282::AID-CNE2>3.0.CO;2-#

Chiappe ME, Seelig JD, Reiser MB, Jayaraman V. 2010. Walking modulates speed sensitivity in drosophila motion vision. Current Biology 20. doi:10.1016/j.cub.2010.06.072

Chklovskii DB, Koulakov AA. 2004. Maps in the brain: what can we learn from them? Annu Rev Neurosci 27:369–392. doi:10.1146/annurev.neuro.27.070203.144226

Currier TA, Pang MM, Clandinin TR. 2023. Visual processing in the fly, from photoreceptors to behavior. Genetics iyad064. doi:10.1093/genetics/iyad064

David CT. 1982. Compensation for height in the control of groundspeed byDrosophila in a new, ‘barber’s pole’ wind tunnel. J Comp Physiol 147:485–493. doi:10.1007/BF00612014

Davis FP, Nern A, Picard S, Reiser MB, Rubin GM, Eddy SR, Henry GL. 2020. A genetic, genomic, and computational resource for exploring neural circuit function. eLife. doi:10.7554/elife.50901

de Vries SEJ, Clandinin TR. 2012. Loom-Sensitive Neurons Link Computation to Action in the Drosophila Visual System. Current Biology 22:353–362. doi:10.1016/j.cub.2012.01.007

Dorkenwald S, Matsliah A, Sterling AR, Schlegel P, Yu S, McKellar CE, Lin A, Costa M, Eichler K, Yin Y, Silversmith W, Schneider-Mizell C, Jordan CS, Brittain D, Halageri A, Kuehner K, Ogedengbe O, Morey R, Gager J, Kruk K, Perlman E, Yang R, Deutsch D, Bland D, Sorek M, Lu R, Macrina T, Lee K, Bae JA, Mu S, Nehoran B, Mitchell E, Popovych S, Wu J, Jia Z, Castro M, Kemnitz N, Ih D, Bates AS, Eckstein N, Funke J, Collman F, Bock DD, Jefferis GSXE, Seung HS, Murthy M, Consortium the F. 2023. Neuronal wiring diagram of an adult brain. doi:10.1101/2023.06.27.546656

Dvorak DR, Bishop LG, Eckert HE. 1975. On the identification of movement detectors in the fly optic lobe. J Comp Physiol 100:5–23. doi:10.1007/BF00623928

Eckstein N, Bates AS, Du M, Hartenstein V, Jefferis GS, Funke J. 2020. Neurotransmitter classification from electron microscopy images at synaptic sites in Drosophila. BioRxiv 2020–06.

Eddison M, Ihrke G. 2022. Expansion-assisted iterative fluorescence in situ hybridization (EASI-FISH) in Drosophila CNS.

Egelhaaf M. 1985. On the neuronal basis of figure-ground discrimination by relative motion in the visual system of the fly: II. Biol Cybern 52:195–209. doi:10.1007/BF00339948

Erginkaya M, Cruz T, Brotas M, Steck K, Nern A, Torrão F, Varela N, Bock D, Reiser M, Chiappe ME. 2023. A competitive disinhibitory network for robust optic flow processing in Drosophila. doi:10.1101/2023.08.06.552150

Farrow K, Borst A, Haag J. 2005. Sharing Receptive Fields with Your Neighbors: Tuning the Vertical System Cells to Wide Field Motion. J Neurosci 25:3985–3993. doi:10.1523/JNEUROSCI.0168-05.2005

Farrow K, Haag J, Borst A. 2006. Nonlinear, binocular interactions underlying flow field selectivity of a motion-sensitive neuron. Nat Neurosci 9:1312–1320. doi:10.1038/nn1769

Fisher YE, Silies M, Clandinin TR. 2015. Orientation Selectivity Sharpens Motion Detection in Drosophila. Neuron 88:390–402. doi:10.1016/j.neuron.2015.09.033

Franz MO, Krapp HG. 2000. Wide-field, motion-sensitive neurons and matched filters for optic flow fields. Biol Cybern 83:185–197. doi:10.1007/s004220000163

Fry SN, Sayaman R, Dickinson MH. 2003. The Aerodynamics of Free-Flight Maneuvers in Drosophila. Science 300:495–498. doi:10.1126/science.1081944

Fujiwara T, Brotas M, Chiappe ME. 2022. Walking strides direct rapid and flexible recruitment of visual circuits for course control in Drosophila. Neuron 110:2124–2138.e8. doi:10.1016/j.neuron.2022.04.008

Gibson JJ. 1950. The perception of the visual world., The perception of the visual world. Oxford, England: Houghton Mifflin.

Gruntman E, Romani S, Reiser MB. 2019. The computation of directional selectivity in the Drosophila OFF motion pathway. eLife 8:e50706. doi:10.7554/eLife.50706

Gruntman E, Romani S, Reiser MB. 2018. Simple integration of fast excitation and offset, delayed inhibition computes directional selectivity in Drosophila. Nature Neuroscience 21:250. doi:10.1038/s41593-017-0046-4

Haag J, Borst A. 2007. Reciprocal inhibitory connections within a neural network for rotational optic-flow processing. Frontiers in Neuroscience 1.

Haag J, Borst A. 2005. Dye-coupling visualizes networks of large-field motion-sensitive neurons in the fly. J Comp Physiol A 191:445–454. doi:10.1007/s00359-005-0605-0

Haag J, Borst A. 2003. Orientation tuning of motion-sensitive neurons shaped by vertical-horizontal network interactions. J Comp Physiol A 189:363–370. doi:10.1007/s00359-003-0410-6

Haag J, Borst A. 2001. Recurrent Network Interactions Underlying Flow-Field Selectivity of Visual Interneurons. J Neurosci 21:5685–5692. doi:10.1523/JNEUROSCI.21-15-05685.2001

Haag J, Mishra A, Borst A. 2017. A common directional tuning mechanism of Drosophila motion-sensing neurons in the ON and in the OFF pathway. eLife. doi:10.7554/elife.29044

Hausen K. 1987. The neural architecture of the lobula plate of the blowfly, Calliphora erythrocephala. Unpublished work.

Heisenberg M, Wonneberger R, Wolf R. 1978. Optomotor-blindH31—aDrosophila mutant of the lobula plate giant neurons. J Comp Physiol 124:287–296. doi:10.1007/BF00661379

Hengstenberg R, Hausen K, Hengstenberg B. 1982. The number and structure of giant vertical cells (VS) in the lobula plate of the blowflyCalliphora erythrocephala. J Comp Physiol 149:163–177. doi:10.1007/BF00619211

Henning M, Ramos-Traslosheros G, Gür B, Silies M. 2022. Populations of local direction– selective cells encode global motion patterns generated by self-motion. Science Advances 8:eabi7112. doi:10.1126/sciadv.abi7112

Hopp E, Borst A, Haag J. 2014. Subcellular mapping of dendritic activity in optic flow processing neurons. J Comp Physiol A 200:359–370. doi:10.1007/s00359-014-0893-3

Hulse BK, Haberkern H, Franconville R, Turner-Evans D, Takemura S, Wolff T, Noorman M, Dreher M, Dan C, Parekh R, Hermundstad AM, Rubin GM, Jayaraman V. 2021. A connectome of the Drosophila central complex reveals network motifs suitable for flexible navigation and context-dependent action selection. eLife 10:e66039. doi:10.7554/eLife.66039

Hulse BK, Stanoev A, Turner-Evans DB, Seelig JD, Jayaraman V. 2023. A rotational velocity estimate constructed through visuomotor competition updates the fly’s neural compass. doi:10.1101/2023.09.25.559373

Isaacson MD, Eliason JLM, Nern A, Rogers EM, Lott GK, Tabachnik T, Rowell WJ, Edwards AW, Korff WL, Rubin GM, Branson K, Reiser MB. 2023. Small-field visual projection neurons detect translational optic flow and support walking control. doi:10.1101/2023.06.21.546024

Joesch M, Plett J, Borst A, Reiff DF. 2008. Response Properties of Motion-Sensitive Visual Interneurons in the Lobula Plate of Drosophila melanogaster. Current Biology 18:368–374. doi:10.1016/j.cub.2008.02.022

Karmeier K, Krapp HG, Egelhaaf M. 2003. Robustness of the Tuning of Fly Visual Interneurons to Rotatory Optic Flow. Journal of Neurophysiology 90:1626–1634. doi:10.1152/jn.00234.2003

Kim AJ, Fenk LM, Lyu C, Maimon G. 2017. Quantitative Predictions Orchestrate Visual Signaling in Drosophila. Cell. doi:10.1016/j.cell.2016.12.005

Kind E, Longden KD, Nern A, Zhao A, Sancer G, Flynn MA, Laughland CW, Gezahegn B, Ludwig HD, Thomson AG, Obrusnik T, Alarcón PG, Dionne H, Bock DD, Rubin GM, Reiser MB, Wernet MF. 2021. Synaptic targets of photoreceptors specialized to detect color and skylight polarization in Drosophila. eLife 10:e71858. doi:10.7554/eLife.71858

Klapoetke NC, Nern A, Rogers EM, Rubin GM, Reiser MB, Card GM. 2022. A functionally ordered visual feature map in the Drosophila brain. Neuron 110:1700–1711.e6. doi:10.1016/j.neuron.2022.02.013

Koenderink JJ, van Doorn AJ. 1987. Facts on optic flow. Biological Cybernetics 1987 56:4 56:247–254. doi:10.1007/BF00365219

Krapp HG, Hengstenberg R. 1997. A fast stimulus procedure to determine local receptive field properties of motion-sensitive visual interneurons. Vision research 37:225–34.

Krapp HG, Hengstenberg R. 1996. Estimation of self-motion by optic flow processing in single visual interneurons. Nature 384:463–466. doi:10.1038/384463a0

Kurtz R, Warzecha A-K, Egelhaaf M. 2001. Transfer of Visual Motion Information via Graded Synapses Operates Linearly in the Natural Activity Range. J Neurosci 21:6957–6966. doi:10.1523/JNEUROSCI.21-17-06957.2001

Li PH, Lindsey LF, Januszewski M, Zheng Z, Bates AS, Taisz I, Tyka M, Nichols M, Li F, Perlman E, Maitin-Shepard J, Blakely T, Leavitt L, Jefferis GSXE, Bock D, Jain V. 2020. Automated Reconstruction of a Serial-Section EM Drosophila Brain with Flood-Filling Networks and Local Realignment. doi:10.1101/605634

Liu WW, Wilson RI. 2013. Glutamate is an inhibitory neurotransmitter in the Drosophila olfactory system. Proc Natl Acad Sci U S A 110:10294–10299. doi:10.1073/pnas.1220560110

Longden KD, Wicklein M, Hardcastle BJ, Huston SJ, Krapp HG. 2017. Spike Burst Coding of Translatory Optic Flow and Depth from Motion in the Fly Visual System. Current biology : CB 27:3225–3236.e3. doi:10.1016/j.cub.2017.09.044

Lyu C, Abbott LF, Maimon G. 2022. Building an allocentric travelling direction signal via vector computation. Nature 601:92–97. doi:10.1038/s41586-021-04067-0

Maimon G, Straw AD, Dickinson MH. 2010. Active flight increases the gain of visual motion processing in Drosophila. Nature Neuroscience 2010 13:3 13:393–399. doi:10.1038/nn.2492

Maisak MS, Haag J, Ammer G, Serbe E, Meier M, Leonhardt A, Schilling T, Bahl A, Rubin GM, Nern A, Dickson BJ, Reiff DF, Hopp E, Borst A. 2013. A directional tuning map of Drosophila elementary motion detectors. Nature 500:212–216. doi:10.1038/nature12320

Mauss AS, Pankova K, Arenz A, Nern A, Rubin GM, Borst A. 2015. Neural Circuit to Integrate Opposing Motions in the Visual Field. Cell 162:351–362. doi:10.1016/J.CELL.2015.06.035

Meissner GW, Nern A, Dorman Z, DePasquale GM, Forster K, Gibney T, Hausenfluck JH, He Y, Iyer NA, Jeter J, Johnson L, Johnston RM, Lee K, Melton B, Yarbrough B, Zugates CT, Clements J, Goina C, Otsuna H, Rokicki K, Svirskas RR, Aso Y, Card GM, Dickson BJ, Ehrhardt E, Goldammer J, Ito M, Kainmueller D, Korff W, Mais L, Minegishi R, Namiki S, Rubin GM, Sterne GR, Wolff T, Malkesman O, FlyLight Project Team. 2023. A searchable image resource of Drosophila GAL4 driver expression patterns with single neuron resolution. eLife 12:e80660. doi:10.7554/eLife.80660

Meyer EP, Matute C, Streit P, Nässel DR. 1986. Insect optic lobe neurons identifiable with monoclonal antibodies to GABA. Histochemistry 84:207–216. doi:10.1007/BF00495784

Nern A, Pfeiffer BD, Rubin GM. 2015. Optimized tools for multicolor stochastic labeling reveal diverse stereotyped cell arrangements in the fly visual system. Proceedings of the National Academy of Sciences 112:E2967–E2976. doi:10.1073/pnas.1506763112

Ng A, Jordan M, Weiss Y. 2001. On Spectral Clustering: Analysis and an algorithmAdvances in Neural Information Processing Systems. MIT Press.

Nordström K, Barnett PD, Miguel IMM de, Brinkworth RSA, O’Carroll DC. 2008. Sexual Dimorphism in the Hoverfly Motion Vision Pathway. Current Biology 18:661–667. doi:10.1016/j.cub.2008.03.061

Pierantoni R. 1976. A look into the cock-pit of the fly. Cell Tissue Res 171:101–122. doi:10.1007/BF00219703

Pokusaeva VO, Satapathy R, Symonova O, Jösch M. 2023. Gap junctions arbitrate binocular course control in flies. doi:10.1101/2023.05.31.543181

Rajashekhar KP, Shamprasad VR. 2004. Golgi analysis of tangential neurons in the lobula plate ofDrosophila melanogaster. J Biosci 29:93–104. doi:10.1007/BF02702566

Ramón y Cajal S, Sánchez D. 1915. Contribución al conocimiento de los centros nerviosos de los insectos. Imprenta de Hijos de Nicolás Moya,.

Rivera-Alba M, Vitaladevuni SN, Mishchenko Y, Lu Z, Takemura S, Scheffer L, Meinertzhagen IA, Chklovskii DB, de Polavieja GG. 2011. Wiring Economy and Volume Exclusion Determine Neuronal Placement in the Drosophila Brain. Current Biology 21:2000–2005. doi:10.1016/j.cub.2011.10.022

Saalfeld S, Cardona A, Hartenstein V, Tomančák P. 2009. CATMAID: Collaborative annotation toolkit for massive amounts of image data. Bioinformatics. doi:10.1093/bioinformatics/btp266

Scheffer LK, Xu CS, Januszewski M, Lu Z, Takemura Shin-ya, Hayworth KJ, Huang GB, Shinomiya K, Maitlin-Shepard J, Berg S, Clements J, Hubbard PM, Katz WT, Umayam L, Zhao T, Ackerman D, Blakely T, Bogovic J, Dolafi T, Kainmueller D, Kawase T, Khairy KA, Leavitt L, Li PH, Lindsey L, Neubarth N, Olbris DJ, Otsuna H, Trautman ET, Ito M, Bates AS, Goldammer J, Wolff T, Svirskas R, Schlegel P, Neace E, Knecht CJ, Alvarado CX, Bailey DA, Ballinger S, Borycz JA, Canino BS, Cheatham N, Cook M, Dreher M, Duclos O, Eubanks B, Fairbanks K, Finley S, Forknall N, Francis A, Hopkins GP, Joyce EM, Kim S, Kirk NA, Kovalyak J, Lauchie SA, Lohff A, Maldonado C, Manley EA, McLin S, Mooney C, Ndama M, Ogundeyi O, Okeoma N, Ordish C, Padilla N, Patrick CM, Paterson T, Phillips EE, Phillips EM, Rampally N, Ribeiro C, Robertson MK, Rymer JT, Ryan SM, Sammons M, Scott AK, Scott AL, Shinomiya A, Smith C, Smith K, Smith NL, Sobeski MA, Suleiman A, Swift J, Takemura Satoko, Talebi I, Tarnogorska D, Tenshaw E, Tokhi T, Walsh JJ, Yang T, Horne JA, Li F, Parekh R, Rivlin PK, Jayaraman V, Costa M, Jefferis GS, Ito K, Saalfeld S, George R, Meinertzhagen IA, Rubin GM, Hess HF, Jain V, Plaza SM. 2020. A connectome and analysis of the adult Drosophila central brain. eLife 9:e57443. doi:10.7554/eLife.57443

Schlegel P, Yin Y, Bates AS, Dorkenwald S, Eichler K, Brooks P, Han DS, Gkantia M, Santos M dos, Munnelly EJ, Badalamente G, Capdevila LS, Sane VA, Pleijzier MW, Tamimi IFM, Dunne CR, Salgarella I, Javier A, Fang S, Perlman E, Kazimiers T, Jagannathan SR, Matsliah A, Sterling AR, Yu S, McKellar CE, Consortium F, Costa M, Seung HS, Murthy M, Hartenstein V, Bock DD, Jefferis GSXE. 2023. Whole-brain annotation and multi-connectome cell typing quantifies circuit stereotypy in Drosophila. doi:10.1101/2023.06.27.546055

Schneider-Mizell CM, Gerhard S, Longair M, Kazimiers T, Li F, Zwart MF, Champion A, Midgley FM, Fetter RD, Saalfeld S, Cardona A. 2016. Quantitative neuroanatomy for connectomics in Drosophila. eLife 5:e12059. doi:10.7554/eLife.12059

Schnell B, Joesch M, Forstner F, Raghu SV, Otsuna H, Ito K, Borst A, Reiff DF. 2010. Processing of Horizontal Optic Flow in Three Visual Interneurons of the Drosophila Brain. Journal of Neurophysiology 103:1646–1657. doi:10.1152/jn.00950.2009

Scott EK, Raabe T, Luo L. 2002. Structure of the vertical and horizontal system neurons of the lobula plate in Drosophila. Journal of Comparative Neurology. doi:10.1002/cne.10467

Shinomiya K, Huang G, Lu Z, Parag T, Xu CS, Aniceto R, Ansari N, Cheatham N, Lauchie S, Neace E, Ogundeyi O, Ordish C, Peel D, Shinomiya A, Smith C, Takemura S, Talebi I, Rivlin PK, Nern A, Scheffer LK, Plaza SM, Meinertzhagen IA. 2019. Comparisons between the ON- and OFF-edge motion pathways in the Drosophila brain. eLife. doi:10.7554/eLife.40025

Shinomiya K, Nern A, Meinertzhagen IA, Plaza SM, Reiser MB. 2022. Neuronal circuits integrating visual motion information in Drosophila melanogaster. Current Biology. doi:10.1016/j.cub.2022.06.061

Strahler AN. 1957. Quantitative analysis of watershed geomorphology. *Eos*, Transactions American Geophysical Union 38:913–920. doi:10.1029/TR038i006p00913

Strausfeld NJ. 1970. The Optic Lobes of Diptera. Philosophical Transactions of the Royal Society of London Series B, Biological Sciences 258:135–223.

Strother JA, Wu S-T, Rogers EM, Eliason JLM, Wong AM, Nern A, Reiser MB. 2018. Behavioral state modulates the on visual motion pathway of drosophila. Proceedings of the National Academy of Sciences of the United States of America 115. doi:10.1073/pnas.1703090115

Strother JA, Wu S-T, Wong AM, Nern A, Rogers EM, Le JQ, Rubin GM, Reiser MB. 2017. The Emergence of Directional Selectivity in the Visual Motion Pathway of Drosophila. Neuron 94:168–182.e10. doi:10.1016/j.neuron.2017.03.010

Takemura S, Nern A, Chklovskii DB, Scheffer LK, Rubin GM, Meinertzhagen IA. 2017. The comprehensive connectome of a neural substrate for ‘ON’ motion detection in Drosophila. eLife 6:e24394. doi:10.7554/eLife.24394

Takemura SY, Bharioke A, Lu Z, Nern A, Vitaladevuni S, Rivlin PK, Katz WT, Olbris DJ, Plaza SM, Winston P, Zhao T, Horne JA, Fetter RD, Takemura S, Blazek K, Chang LA, Ogundeyi O, Saunders MA, Shapiro V, Sigmund C, Rubin GM, Scheffer LK, Meinertzhagen IA, Chklovskii DB. 2013. A visual motion detection circuit suggested by Drosophila connectomics. Nature. doi:10.1038/nature12450

Tanaka R, Zhou B, Agrochao M, Badwan BA, Au B, Matos NCB, Clark DA. 2023. Neural mechanisms to incorporate visual counterevidence in self motion estimation. doi:10.1101/2023.01.04.522814

Wang Y, Eddison M, Fleishman G, Weigert M, Xu S, Wang T, Rokicki K, Goina C, Henry FE, Lemire AL, Schmidt U, Yang H, Svoboda K, Myers EW, Saalfeld S, Korff W, Sternson SM, Tillberg PW. 2021. EASI-FISH for thick tissue defines lateral hypothalamus spatio-molecular organization. Cell 184:6361–6377.e24. doi:10.1016/j.cell.2021.11.024

Warzecha AK, Egelhaaf M, Borst A. 1993. Neural circuit tuning fly visual interneurons to motion of small objects. I. Dissection of the circuit by pharmacological and photoinactivation techniques. J Neurophysiol 69:329–339. doi:10.1152/jn.1993.69.2.329

Wasserman SM, Aptekar JW, Lu P, Nguyen J, Wang AL, Keles MF, Grygoruk A, Krantz DE, Larsen C, Frye MA. 2015. Olfactory Neuromodulation of Motion Vision Circuitry in Drosophila. Current Biology 25:467–472. doi:10.1016/j.cub.2014.12.012

Wei H, Kyung HY, Kim PJ, Desplan C. 2020. The diversity of lobula plate tangential cells (LPTCs) in the Drosophila motion vision system. J Comp Physiol A 206:139–148. doi:10.1007/s00359-019-01380-y

Weir PT, Dickinson MH. 2015. Functional divisions for visual processing in the central brain of flying Drosophila. Proceedings of the National Academy of Sciences 112:E5523–E5532. doi:10.1073/pnas.1514415112

Wu M, Nern A, Ryan Williamson W, Morimoto MM, Reiser MB, Card GM, Rubin GM. 2016. Visual projection neurons in the Drosophila lobula link feature detection to distinct behavioral programs. eLife 5. doi:10.7554/eLife.21022

Zhao A, Gruntman E, Nern A, Iyer NA, Rogers EM, Koskela S, Siwanowicz I, Dreher M, Flynn MA, Laughland CW, Ludwig HDF, Thomson AG, Moran CP, Gezahegn B, Bock DD, Reiser MB. 2022. Eye structure shapes neuron function in Drosophila motion vision. doi:10.1101/2022.12.14.520178

Zheng Z, Lauritzen JS, Perlman E, Robinson CG, Nichols M, Milkie D, Torrens O, Price J, Fisher CB, Sharifi N, Calle-Schuler SA, Kmecova L, Ali IJ, Karsh B, Trautman ET, Bogovic JA, Hanslovsky P, Jefferis GSXE, Kazhdan M, Khairy K, Saalfeld S, Fetter RD, Bock DD. 2018. A Complete Electron Microscopy Volume of the Brain of Adult Drosophila melanogaster. Cell 174:730–743.e22. doi:10.1016/J.CELL.2018.06.019

