## Supplemental File 1 for "A comprehensive neuroanatomical survey of the *Drosophila* Lobula Plate Tangential Neurons with predictions for their optic flow sensitivity"

| | Right hemisphere | | | | | | | Left hemisphere | | | | | |
|---|---|---|---|---|---|---|---|---|---|---|---|---|---|
| **group** | **name** | **CATMAID skid** | **Hemibrain bodyid** | **FlyWire ID** | **FlyWire soma xyz** | **transmitter** | | **name** | **CATMAID skid** | **Hemibrain bodyid** | **FlyWire ID** | **FlyWire soma xyz** | **transmitter** |
| HS | LPT01_R_HSN | 830793 | 2211443902 | 720575940628031249 | 92419, 60544, 6514 | acetylcholine | | LPT01_L_HSN | 1088632 | | 720575940615933919 | 168936, 65983, 5843 | acetylcholine |
| HS | LPT02_R_HSE | 827034 | 1807537598 | 720575940642723981 | 98622, 55662, 5601 | acetylcholine | | LPT02_L_HSE | 1088754 | | 720575940629148007 | 165324, 58221, 5440 | acetylcholine |
| HS | LPT03_R_HSS | 4058824 | 2179731270 | 720575940622312965 | 94415, 61273, 6326 | acetylcholine | | LPT03_L_HSS | 3796480 | | 720575940620960347 | 173701, 59833, 5826 | acetylcholine |
| HST | LPT04_R_HST | 985774 | 5813042688 | 720575940612296154 | 90246, 60095, 6567 | acetylcholine | | LPT04_L_HST | 9158488 | | 720575940633182291 | 172785, 57390, 5750 | acetylcholine |
| VS | LPT05_R_VS1 | 982897 | 5813024262 | 720575940626477498 | 86605, 61925, 6776 | acetylcholine | | LPT05_L_VS1 | 1066332 | | 720575940619878961 | 180732, 56951, 5935 | acetylcholine |
| VS | LPT06_R_VS2 | 793032 | 1868619183 | 720575940615269794 | 82996, 59458, 6768 | acetylcholine | | LPT06_L_VS2 | 1027307 | | 720575940625331860 | 175038, 66315, 5995 | acetylcholine |
| VS | LPT07_R_VS3 | 815776 | 5813025091 | 720575940622831740 | 85204, 54954, 6650 | acetylcholine | | LPT07_L_VS3 | 1031609 | | 720575940641812699 | 176523, 67292, 5996 | acetylcholine |
| VS | LPT08_R_VS4 | 17686499 | 1557885051 | 720575940633017939 | 82041, 55308, 6788 | acetylcholine | | LPT08_L_VS4 | 1038415 | | 720575940659799937 | 178209, 66353, 6020 | acetylcholine |
| VS | LPT09_R_VS5 | 807401 | 5813033533 | 720575940626457406 | 85255, 59585, 6818 | acetylcholine | | LPT09_L_VS5 | 1037034 | | 720575940639151694 | 177141, 63573, 6015 | acetylcholine |
| VS | LPT10_R_VS6 | 804539 | 1992161820 | 720575940605688492 | 84161, 56888, 6776 | acetylcholine | | LPT10_L_VS6 | 5053485 | | 720575940626928521 | 176791, 65544, 5982 | acetylcholine |
| VS | LPT11_R_VS7 | 851432 | 1868255513 | 720575940624931564 | 89772, 60887, 6632 | acetylcholine | | LPT11_L_VS7 | 1040470 | | 720575940618681709 | 178365, 61493, 6006 | acetylcholine |
| VS | LPT12_R_VS8 | 804092 | 1558230909 | 720575940633923298 | 80405, 58223, 6579 | acetylcholine | | LPT12_L_VS8 | 1034264 | | 720575940636972400 | 175995, 64465, 5995 | acetylcholine |
| VS | LPT13_R_VSm-1 | 852286 | 1836516710 | 720575940620463307 | 87866, 58912, 6715 | acetylcholine | | LPT13_L_VSm-1 | 1038271 | | 720575940631615545 | 178690, 65008, 6029 | acetylcholine |
| VS | LPT13_R_VSm-2 | 815241 | 1805481901 | 720575940630825527 | 80973, 56494, 6842 | acetylcholine | | LPT13_L_VSm-2 | 1030967 | | 720575940624500412 | 175032, 64338, 5995 | acetylcholine |
| VST | LPT15_R_VST1-1 | 1124867 | | 720575940626947971 | 89787, 63442, 6664 | acetylcholine | | LPT15_L_VST1-1 | 4646358 | | 720575940630700348 | 172987, 65432, 5938 | acetylcholine |
| VST | LPT15_R_VST1-2 | 988674 | 5812993165 | 720575940645317412 | 90491, 63121, 6700 | acetylcholine | | LPT15_L_VST1-2 | 4909997 | | 720575940620885158 | 176705, 64882, 6083 | acetylcholine |
| VST | LPT17_R_VST2-1 | 1123389 | 1837198715 | 720575940607420290 | 88409, 61439, 6677 | acetylcholine | | LPT17_L_VST2-1 | 4930213 | | 720575940626690078 | 175782, 61463, 5953 | acetylcholine |
| VST | LPT17_R_VST2-2 | 1112633 | 5813023313 | 720575940646114926 | 86093, 56825, 6691 | acetylcholine | | LPT17_L_VST2-2 | 1059146 | | 720575940624981908 | 177331, 58779, 5908 | acetylcholine |
| VST | LPT17_R_VST2-3 | 2852912 | 5813049185 | 720575940635615339 | 87772, 60990, 6701 | acetylcholine | | LPT17_L_VST2-3 | 3526291 | | 720575940622939101 | 176251, 59718, 5901 | acetylcholine |
| VST | LPT17_R_VST2-4 | 5031615 | 5813023581 | 720575940628452520 | 84643, 62966, 6845 | acetylcholine | | LPT17_L_VST2-4 | 1058778 | | 720575940624695275 | 174516, 66583, 5944 | acetylcholine |
| Ipsilateral | LPT21_R | 1110765 | 1850310331 | 720575940610781560 | 91734, 46381, 5476 | acetylcholine | | LPT21_L | 7764572 | | 720575940627341736 | 167934, 53014, 4893 | acetylcholine |
| Ipsilateral | LPT22_R | 3510999 | 1501708149 | 720575940610061763 | 97775, 46462, 5376 | gaba | | LPT22_L | 17077102 | | 720575940628685099 | 165247, 52598, 4767 | gaba |
| Ipsilateral | LPT23_R-1 | 4135042 | 1842090544 | 720575940617304949 | 94555, 50477, 5512 | acetylcholine | | LPT23_L-1 | 17053936 | | 720575940621790017 | 167897, 56581, 4842 | acetylcholine |
| Ipsilateral | LPT23_R-2 | 4504548 | 1751364895 | 720575940632545080 | 95706, 51204, 5502 | acetylcholine | | LPT23_L-2 | 17092277 | | 720575940645636887 | 168672, 53867, 4835 | acetylcholine |
| Ipsilateral | LPT23_R-3 | 4504557 | 1751028010 | 720575940619638395 | 93785, 48704, 5586 | acetylcholine | | LPT23_L-3 | 17076395 | | 720575940630171642 | 168507, 55106, 4878 | acetylcholine |
| Ipsilateral | LPT26_R | 1107045 | 1747289181 | 720575940630691895 | 95022, 67847, 4005 | acetylcholine | | LPT26_L | 4702640 | | 720575940625992073 | 162875, 73838, 3644 | acetylcholine |
| Ipsilateral | LPT27_R | 3514698 | 1747608690 | 720575940630990300 | 94413, 68614, 3895 | acetylcholine | | LPT27_L | 7937241 | | 720575940634371996 | 164593, 72304, 3618 | acetylcholine |
| Ipsilateral | LPT28_R | 1071860 | 1407904797 | 720575940620898324 | 87242, 41135, 4875 | acetylcholine | | LPT28_L | 4970951 | | 720575940631468485 | 176196, 44172, 4262 | acetylcholine |
| Ipsilateral | LPT29_R | 7760435 | 5813087982 | 720575940632216899 | 91436, 65820, 4007 | acetylcholine | | LPT29_L | 17096943 | | 720575940633500185 | 163367, 69763, 5142 | acetylcholine |
| Ipsilateral | LPT30_R | 7674107 | 1777301335 | 720575940624047654 | 81985, 59904, 3607 | acetylcholine | | LPT30_L | 7701590 | | 720575940621204673 | 180205, 62796, 2749 | acetylcholine |
| Posterial | LPT31_R-1 | 7690273 | 1622471963 | 720575940634166242 | 89099, 65661, 3909 | acetylcholine | | LPT31_L-1 | 7841427 | | 720575940623263453 | 175392, 68820, 2642 | acetylcholine |
| Posterial | LPT31_R-2 | 7311408 | 1434427171 | 720575940616260146 | 89798, 66530, 3874 | acetylcholine | | LPT31_L-2 | 7841256 | | 720575940622087429 | 172799, 69268, 2782 | acetylcholine |
| Posterial | LPT31_R-3 | 7311493 | 1434427905 | 720575940629577475 | 89987, 68909, 3709 | acetylcholine | | LPT31_L-3 | 8035397 | | 720575940607401138 | 177419, 65843, 2633 | acetylcholine |
| Posterial | LPT31_R-4 | 7694305 | 1344082025 | 720575940629567055 | 88553, 66092, 3832 | acetylcholine | | LPT31_L-4 | 8035389 | | 720575940629012240 | 174034, 68456, 2659 | acetylcholine |
| CH | LPT35_R_dCH | 1077174 | 1466485353 | 720575940636933751 | 131387, 52610, 1235 | gaba | | LPT35_L_dCH | 6243409 | 1545158404 | 720575940628391848 | 125941, 49686, 1261 | gaba |
| CH | LPT36_R_vCH | 1078535 | 5813024201 | 720575940627138562 | 133918, 52245, 838 | gaba | | LPT36_L_vCH | 1078048 | | 720575940634274017 | 129340, 51844, 1347 | gaba |
| H1 | LPT37_R_H1 | 1121730 | 676832896 | 720575940660765569 | 176884, 43788, 2168 | glutamate | | LPT37_L_H1 | 1121335 | 1167783603 | 720575940627348617 | 92329, 43685, 2675 | glutamate |
| Noduli | LPT38_R_Nod1-1 | 1054753 | 5812996970 | 720575940609132043 | 101804, 63992, 5521 | acetylcholine | | LPT38_L_Nod1-1 | 6439204 | 1789306586 | 720575940623997949 | 160483, 67115, 5219 | acetylcholine |
| Noduli | LPT38_R_Nod1-2 | 3546483 | 5812993603 | 720575940628438427 | 94671, 46204, 5314 | acetylcholine | | LPT38_L_Nod1-2 | 6437945 | 1758621675 | 720575940629456860 | 168023, 48186, 4930 | acetylcholine |
| Noduli | LPT40_R_Nod2 | 902072 | 1315529069 | 720575940623235683 | 107626, 47015, 5121 | gaba | | LPT40_L_Nod2 | 16615842 | 1871778911 | 720575940625670949 | 157474, 51216, 4838 | gaba |
| Noduli | LPT41_R_Nod3 | 1058996 | 1352706891 | 720575940623384781 | 100729, 52368, 5430 | acetylcholine | | LPT41_L_Nod3 | 15060018 | 1758617327 | 720575940611869849 | 164666, 53244, 4966 | acetylcholine |
| Noduli | LPT42_R_Nod4 | 1106958 | 1566524544 | 720575940625992781 | 108161, 62214, 5644 | acetylcholine | | LPT42_L_Nod4 | 16615419 | 1725837767 | 720575940619612142 | 159693, 68616, 5133 | acetylcholine |
| Noduli | LPT43_R_H2 | 1088678 | 1534124048 | 720575940627079938 | 104111, 49333, 5272 | acetylcholine | | LPT43_L_H2 | 5232902 | 5813078454 | 720575940632427603 | 164234, 55937, 5175 | acetylcholine |
| Noduli | LPT44_R_Nod5 | 1121795 | 1899956171 | 720575940633685459 | 98853, 47380, 5337 | acetylcholine | | LPT44_L_Nod5 | 7231304 | 5812994338 | 720575940620431808 | 164930, 52607, 5201 | acetylcholine |
| Calyx | LPT45_R_dCal1 | 7204844 | 943813788 | 720575940628350997 | 86606, 64556, 6802 | gaba | | LPT45_L_dCal1 | 3509763 | 5813057267 | 720575940629288271 | 166160, 69437, 5555 | gaba |
| Calyx | LPT46_R_vCal1 | 7510076 | 1005174931 | 720575940641223888 | 92000, 41560, 5217 | glutamate | | LPT46_L_vCal1 | 8747266 | 943468720 | 720575940624943504 | 169463, 47531, 4854 | glutamate |
| Calyx | LPT47_R_vCal2 | 3529071 | 1005174975 | 720575940618653524 | 89976, 42233, 5046 | glutamate | | LPT47_L_vCal2 | 7449616 | 943472755 | 720575940649181817 | 169732, 45848, 4848 | glutamate |
| Calyx | LPT48_R_vCal3 | 1056097 | 974502819 | 720575940626919780 | 98267, 43905, 5284 | acetylcholine | | LPT48_L_vCal3 | 11230125 | 943472763 | 720575940626630693 | 164609, 51684, 4749 | acetylcholine |
| Bilateral | LPT49_R | 1110693 | 1654969939 | 720575940638155998 | 81754, 59034, 3645 | acetylcholine | | LPT49_L | 17072553 | 1685896788 | 720575940629791643 | 180014, 63132, 2609 | acetylcholine |
| Bilateral | LPT50_R | 905761 | 1496497366 | 720575940655599777 | 102903, 50290, 5211 | gaba | | LPT50_L | 3503997 | 5813049974 | 720575940611348834 | 164426, 56537, 5078 | gaba |
| Ipsilateral | LPT51_R | 4224711 | 1313496323 | 720575940608287701 | 96820, 68844, 4150 | glutamate | | LPT51_L | 17083003 | | 720575940621153670 | 165710, 62759, 4543 | glutamate |
| Ipsilateral | LPT52_R | 1107296 | 1469291436 | 720575940640716928 | 102698, 64957, 5710 | acetylcholine | | LPT52_L | 17039407 | | 720575940621164532 | 164302, 60961, 5020 | acetylcholine |
| Ipsilateral | LPT53_R | 1111992 | 1231610379 | 720575940653093110 | 103717, 51483, 5709 | gaba | | LPT53_L | 17072462 | | 720575940629704210 | 162429, 54039, 5238 | gaba |
| Ipsilateral | LPT54_R | 4235388 | 1129033939 | 720575940655602849 | 100006, 56145, 5476 | acetylcholine | | LPT54_L | 17196151 | | 720575940632066578 | 163177, 58561, 5143 | acetylcholine |
| Bilateral | LPT55_R_MeLp2 | 1061368 | 1558226341 | 720575940617422731 | 182002, 56164, 2020 | glutamate | | LPT55_L_MeLo2 | 17652973 | | 720575940622735649 | 79037, 46479, 3736 | glutamate |
| Bilateral | LPT56_R_MeLp1 | 3509520 | 5813068976 | 720575940655602849 | 88337, 67218, 3555 | acetylcholine | | LPT56_L_MeLo1 | 4224594 | 5813063227 | 720575940632061738 | 171781, 75310, 2802 | acetylcholine |
| Bilateral | LPT57_R | 886797 | 5901198180 | 720575940632041746 | 132727, 42772, 4246 | acetylcholine | | LPT57_L | 17059178 | 5813045086 | 720575940619709342 | 128014, 38440, 4550 | acetylcholine |
| Bilateral | LPT58_R_V1 | 1059420 | | 720575940621291873 | 136567, 34979, 4354 | acetylcholine | | LPT58_L_V1 | 9539098 | | 720575940622275199 | 131179, 29675, 3714 | acetylcholine |
