## Supplemental File 2 for "A comprehensive neuroanatomical survey of the *Drosophila* Lobula Plate Tangential Neurons with predictions for their optic flow sensitivity"

LPT01_R_HSN_skid=830793

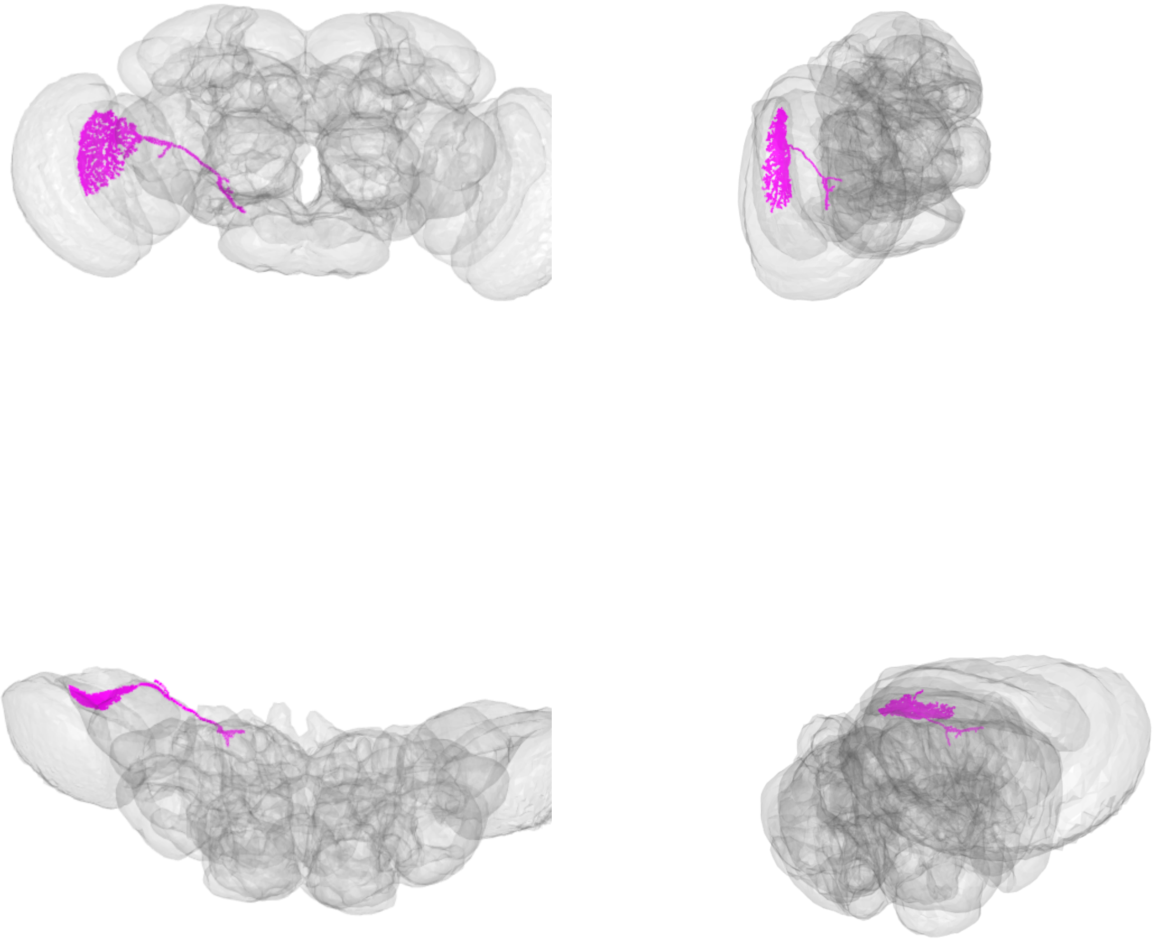

LPT01_R_HSN_skid=830793

LPT02_R_HSE_skid=827034

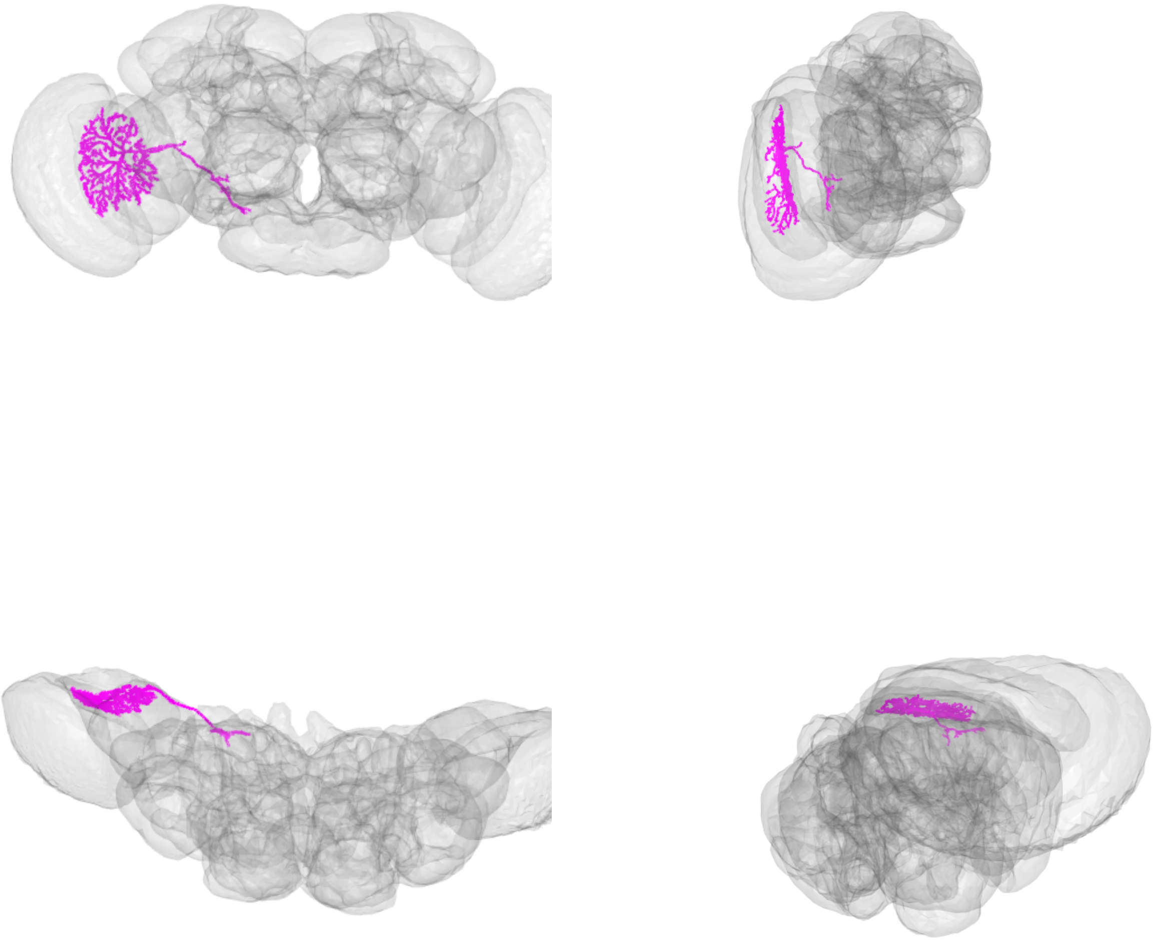

LPT02_R_HSE_skid=827034

LPT03_R_HSS_skid=4058824

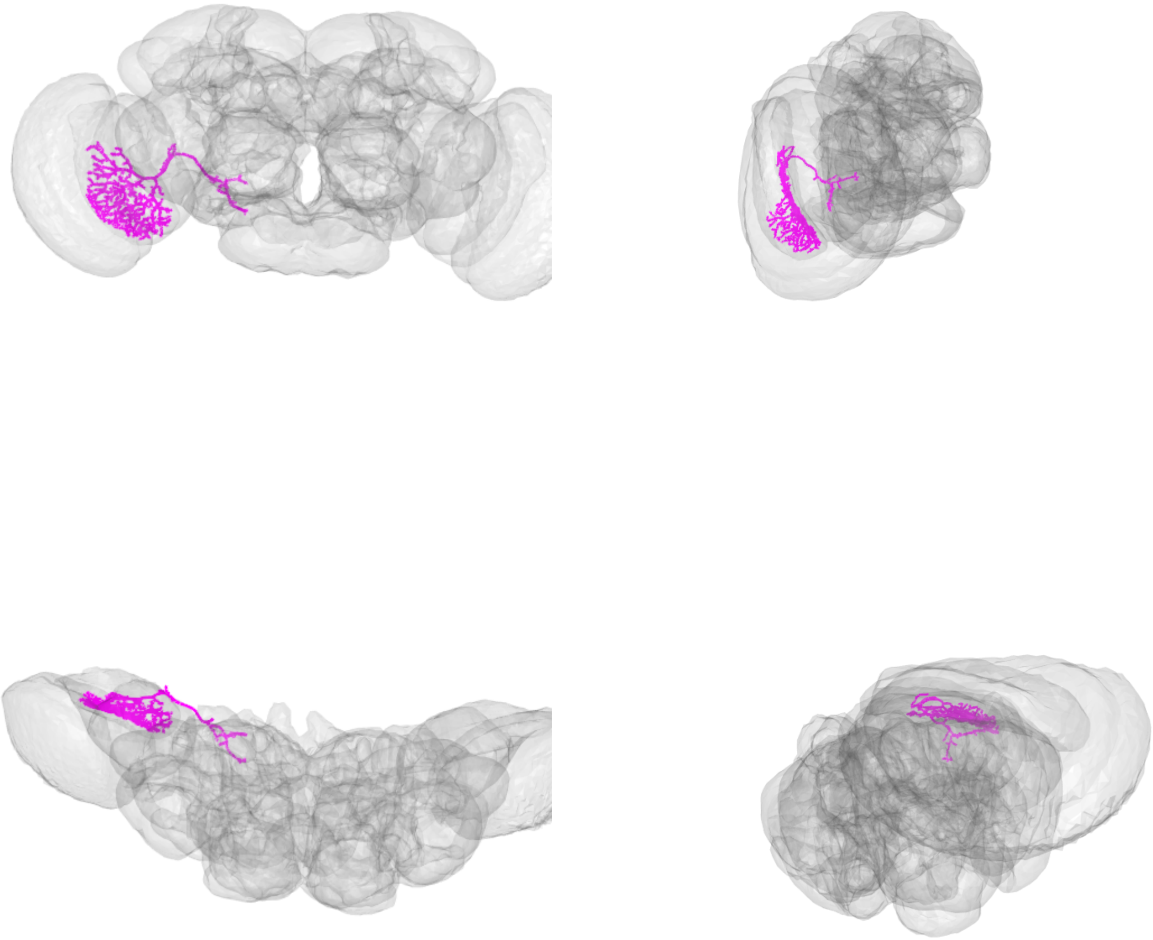

LPT03_R_HSS_skid=4058824

LPT04_R_HS4_skid=985774

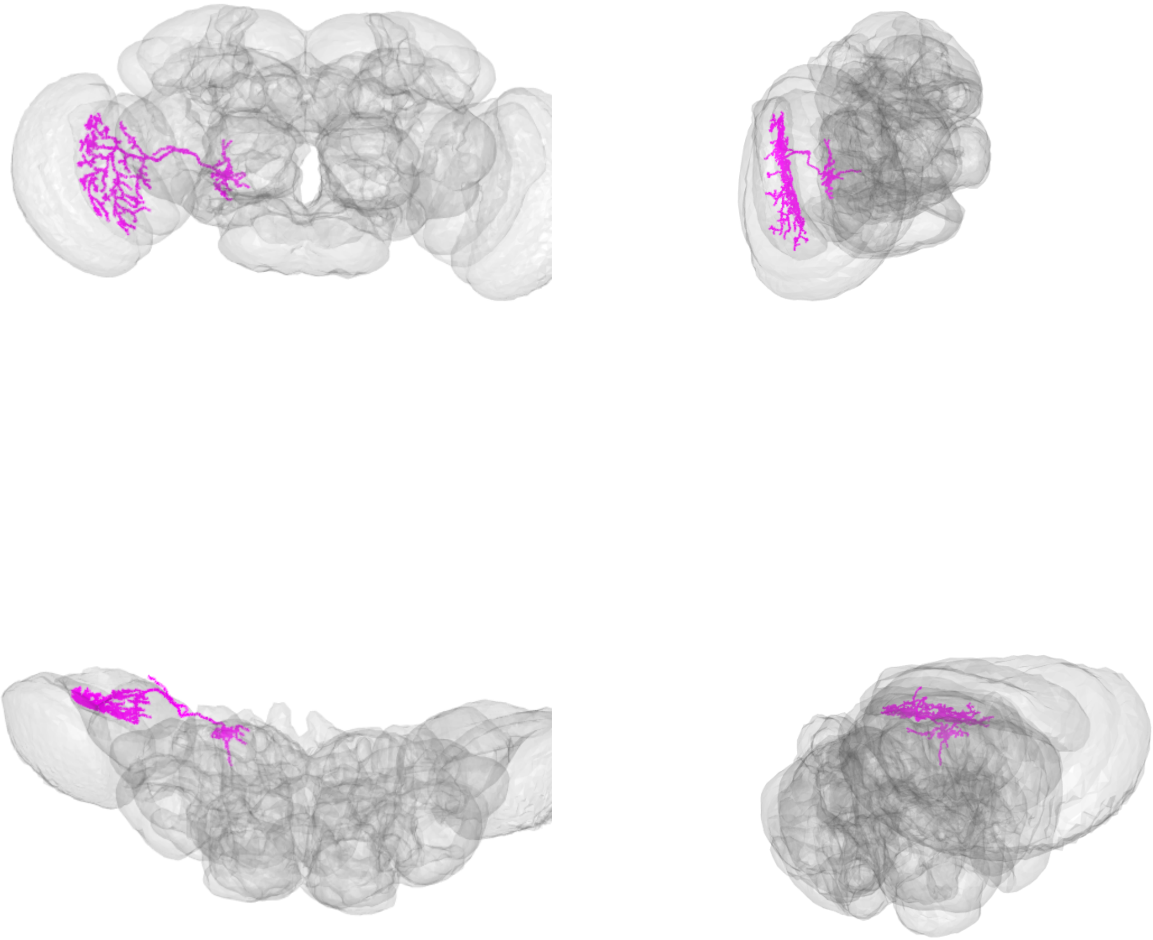

LPT04_R_HS4_skid=985774

LPT05_R_VS1_skid=982897

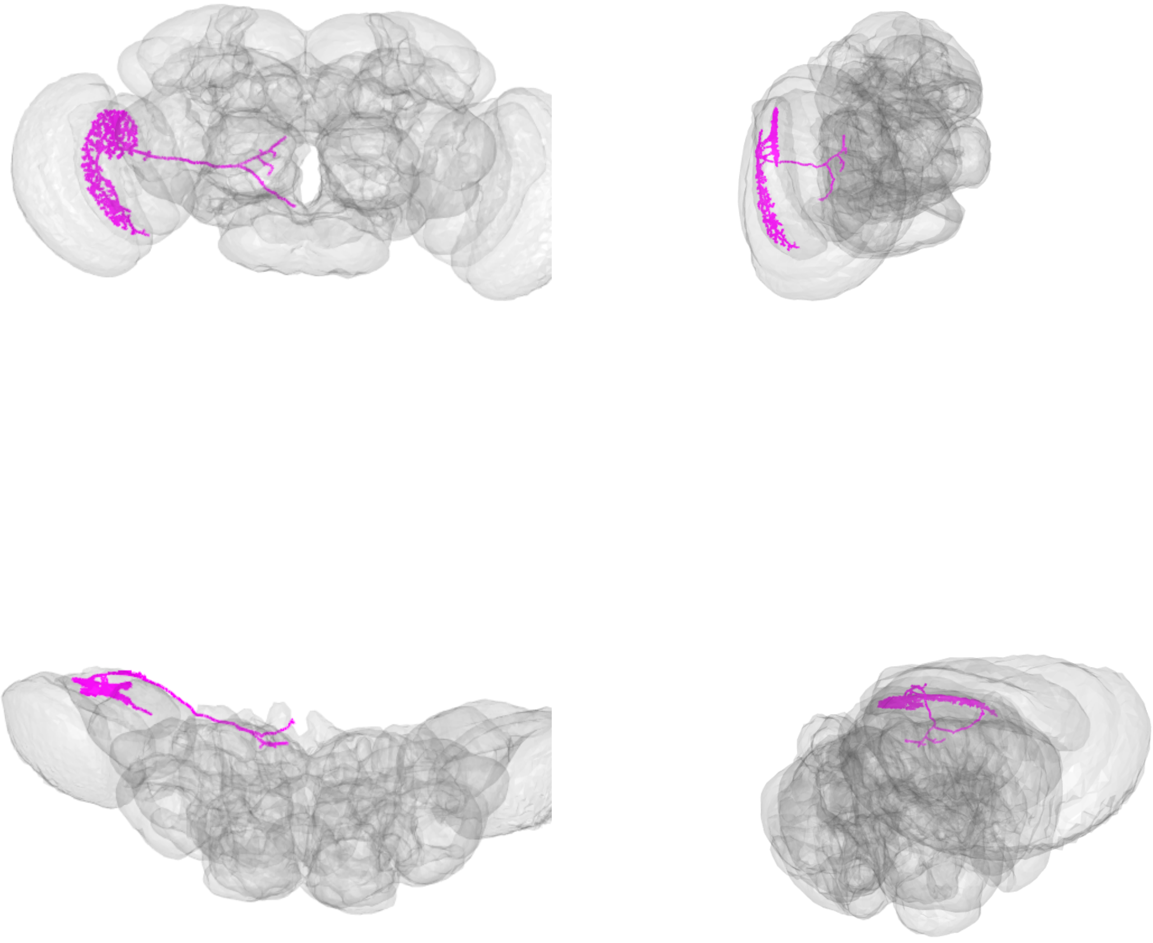

LPT05_R_VS1_skid=982897

LPT06_R_VS2_skid=793032

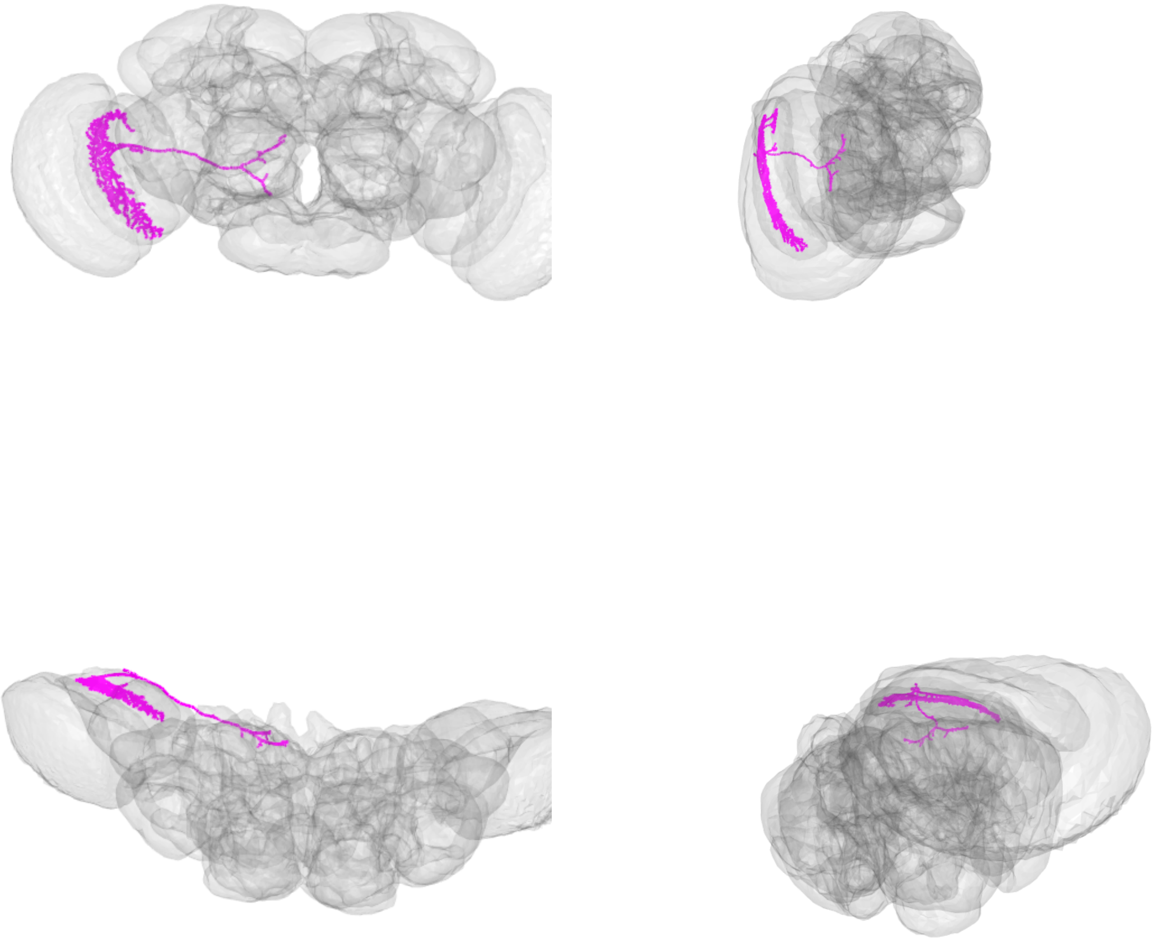

LPT06_R_VS2_skid=793032

LPT07_R_VS3_skid=815776

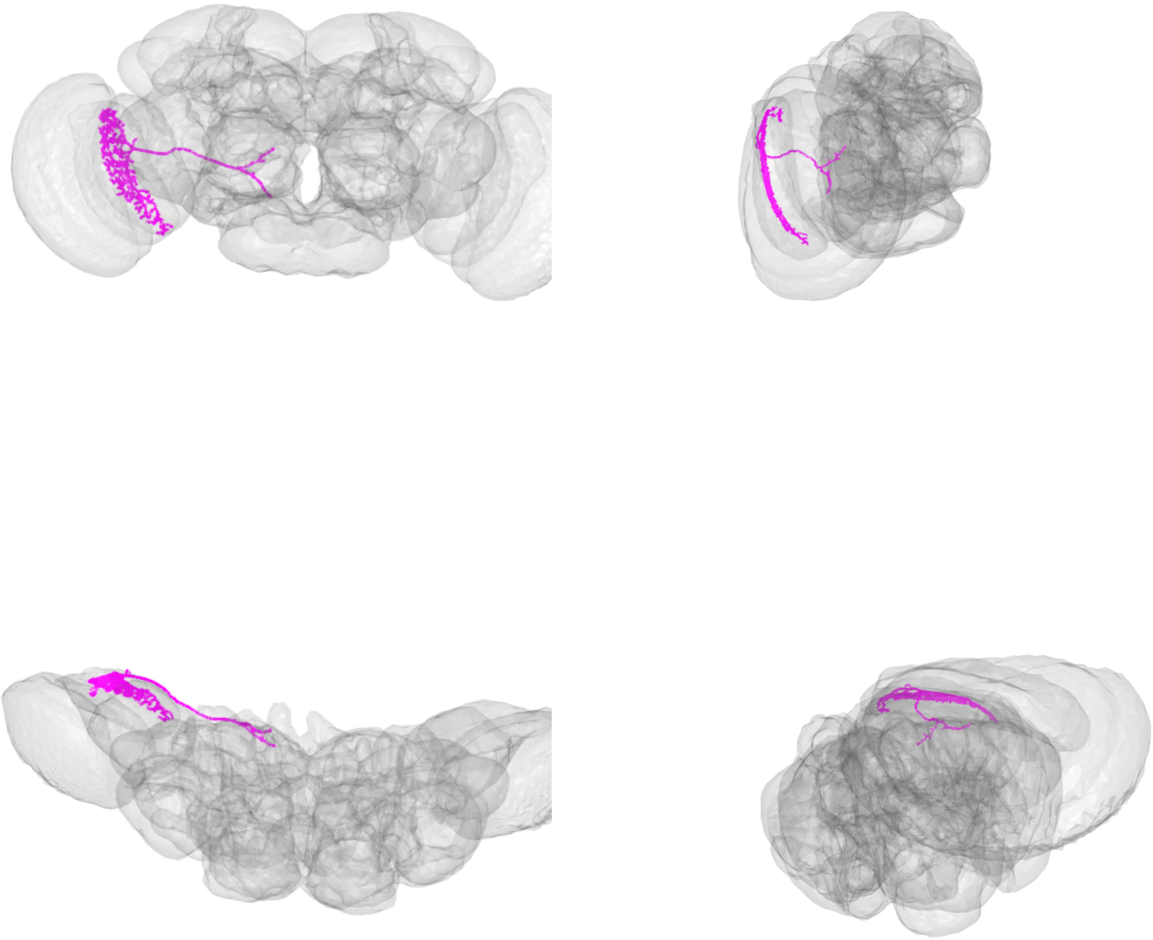

LPT07_R_VS3_skid=815776

LPT08_R_VS4_skid=17686499

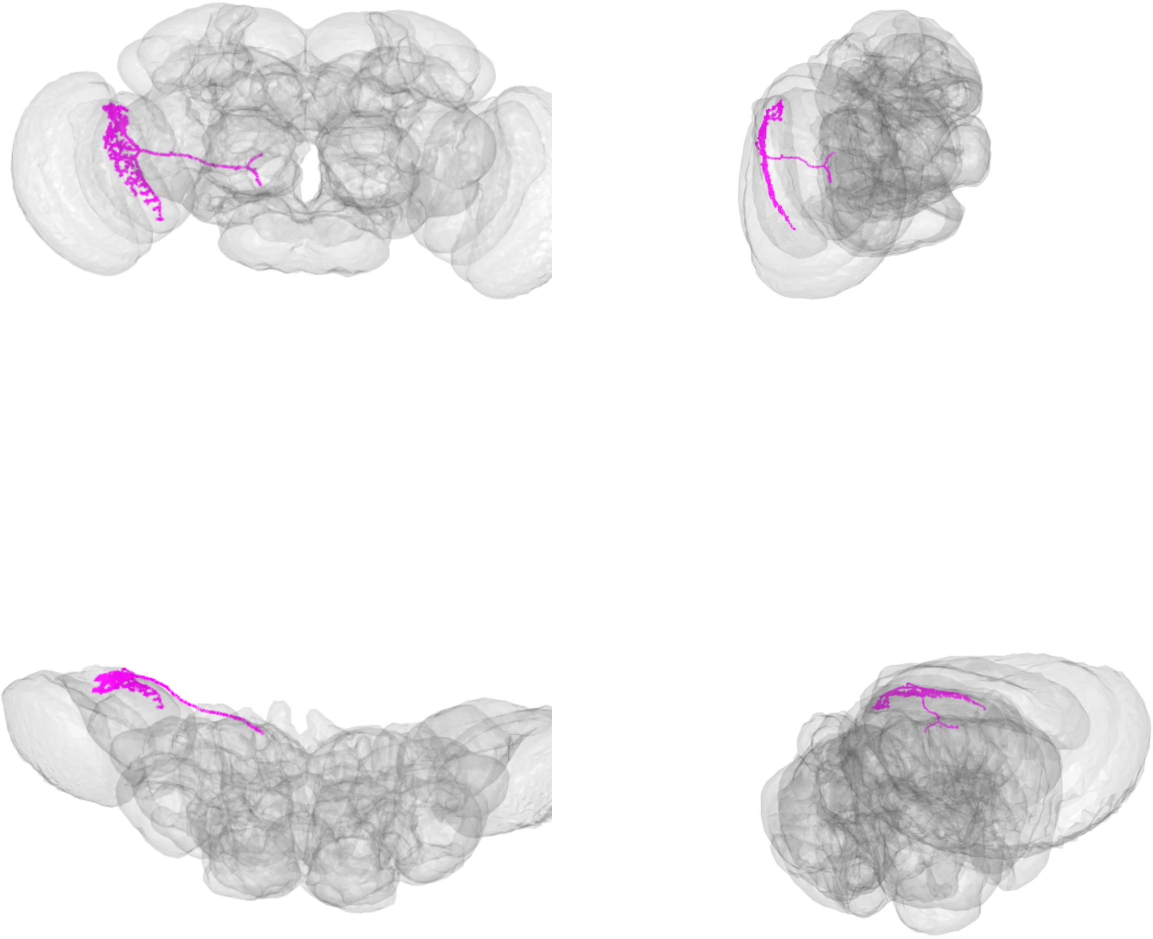

LPT08_R_VS4_skid=17686499

LPT09_R_VS5_skid=807401

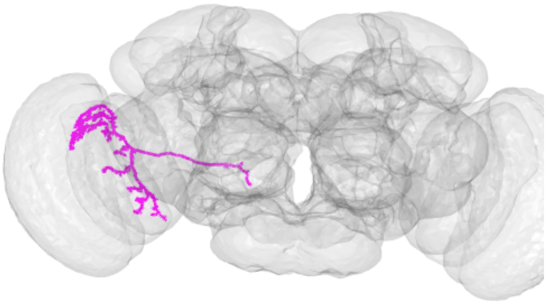

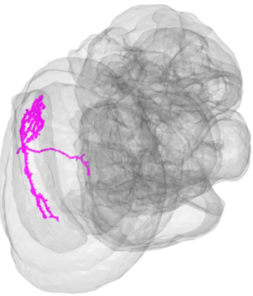

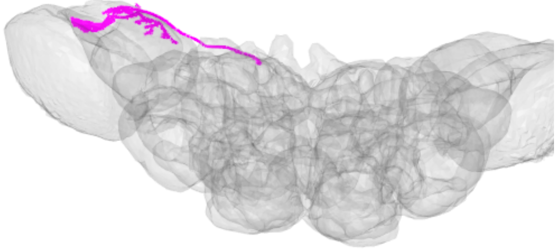

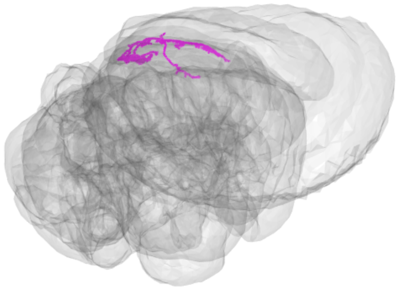

LPT09_R_VS5_skid=807401

LPT10_R_VS6_skid=804539

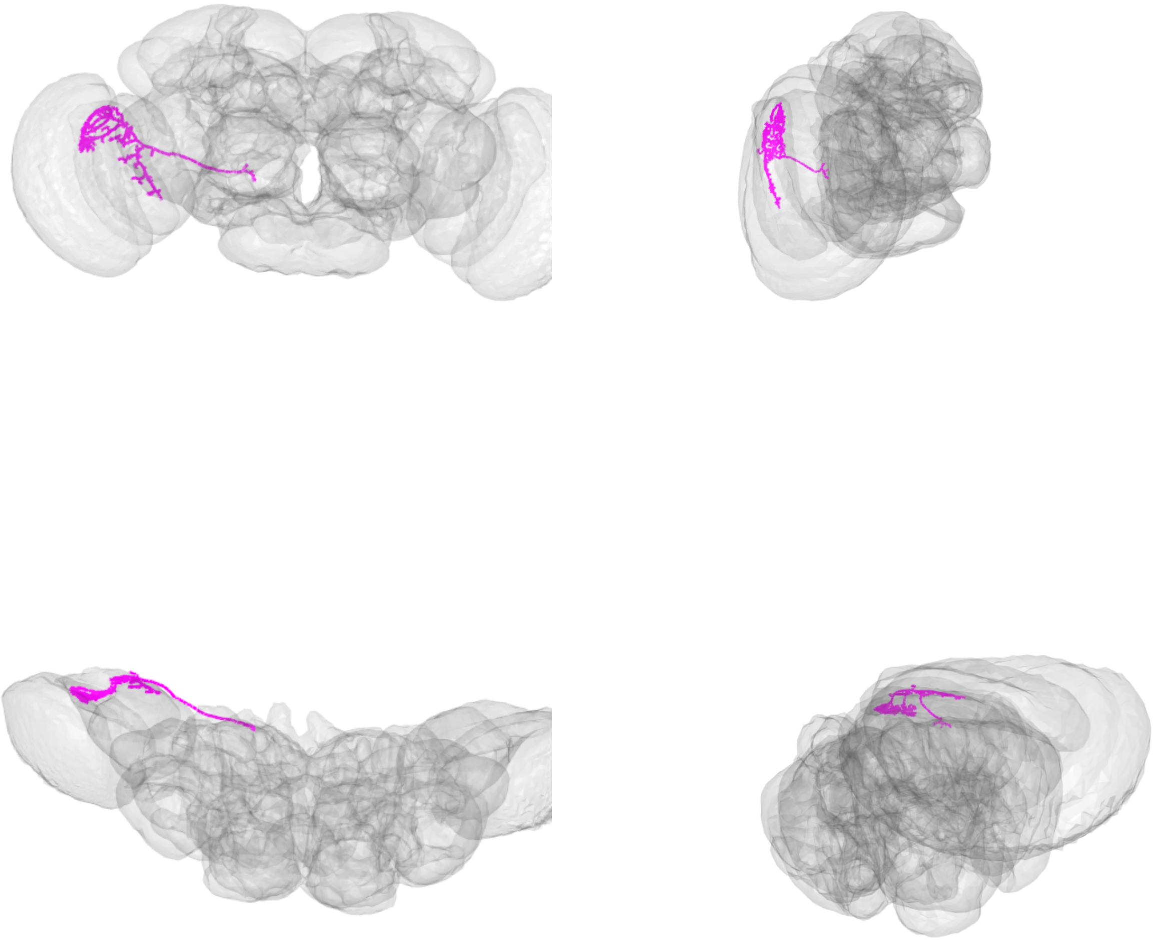

LPT10_R_VS6_skid=804539

LPT11_R_VS7_skid=851432

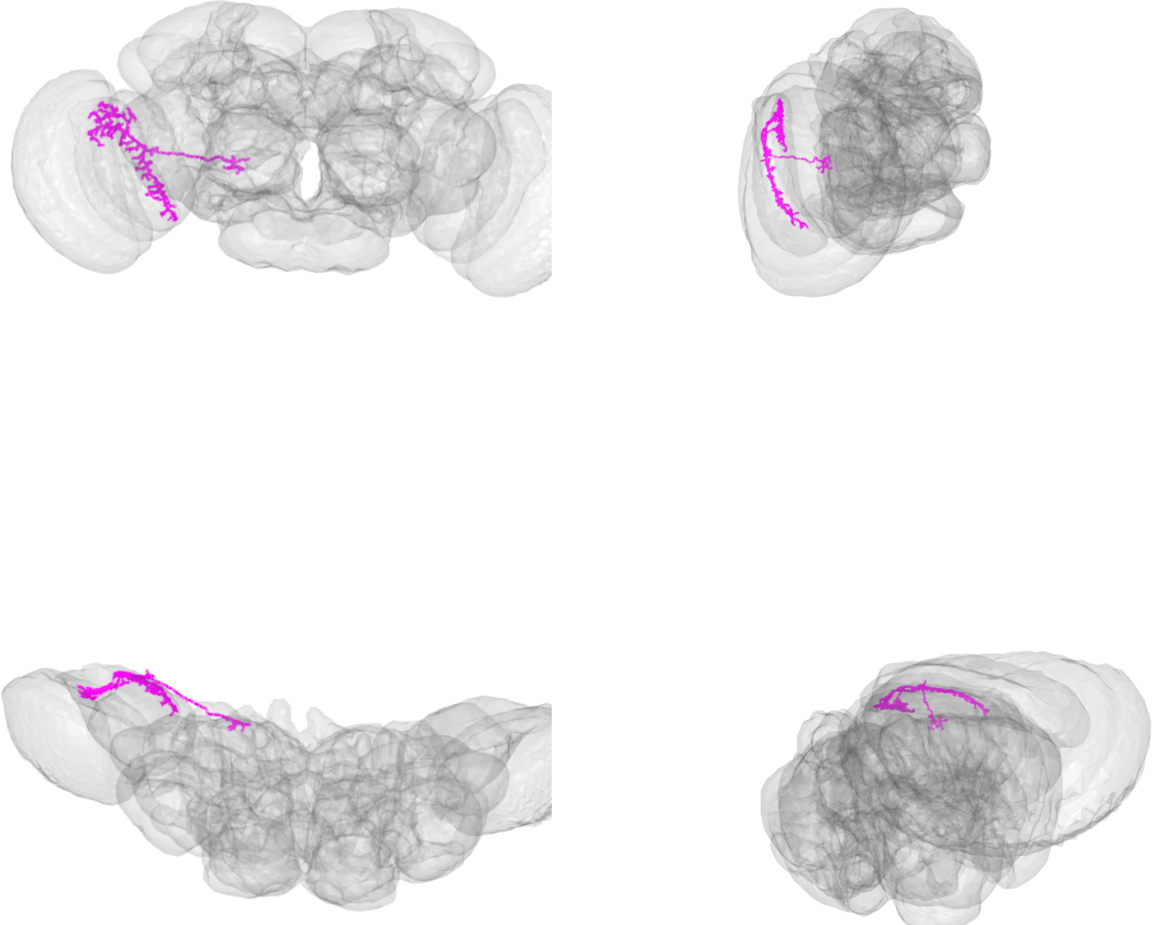

LPT11_R_VS7_skid=851432

LPT12_R_VS8_skid=804092

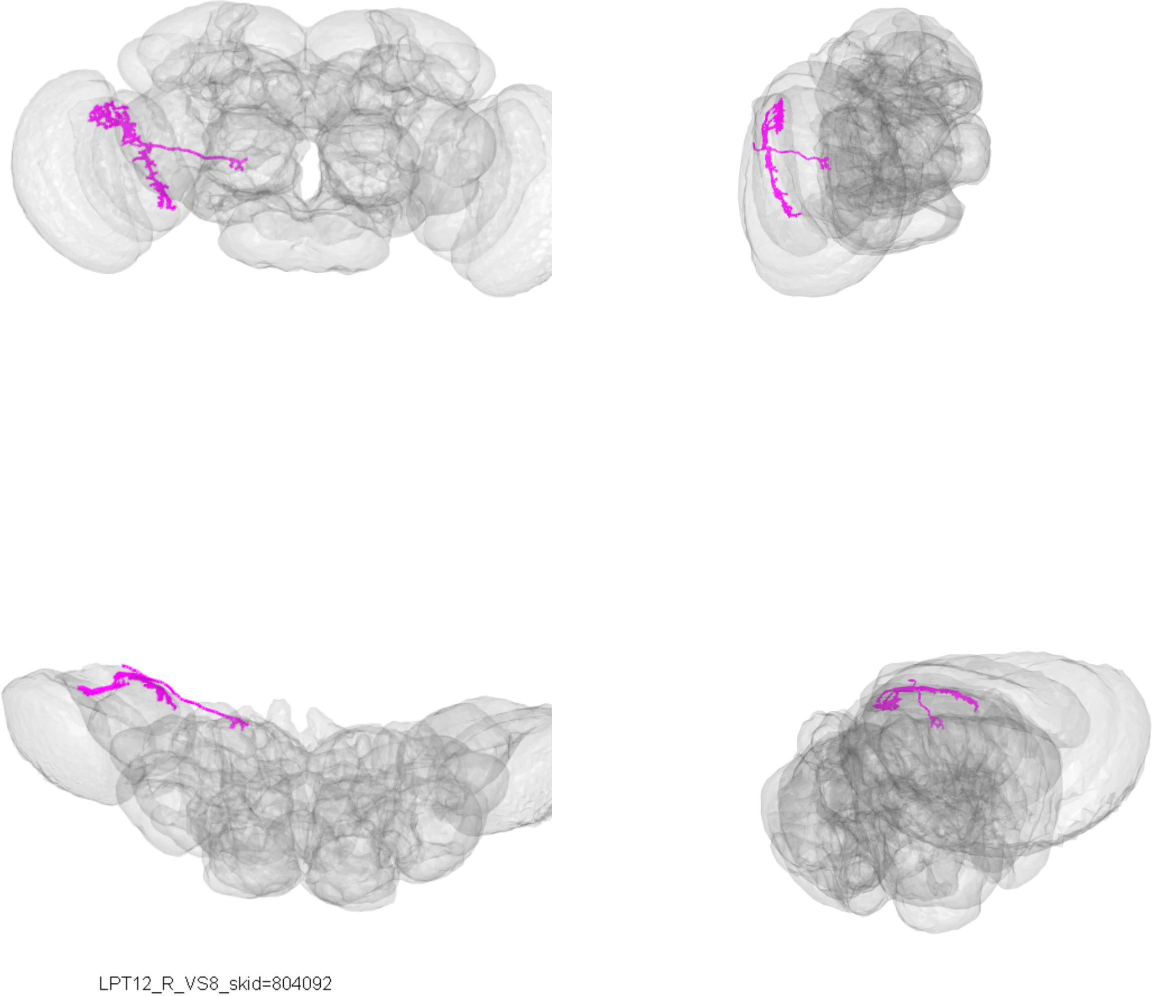

LPT12_R_VS8_skid=804092

LPT13_R_VSm_skid=852286

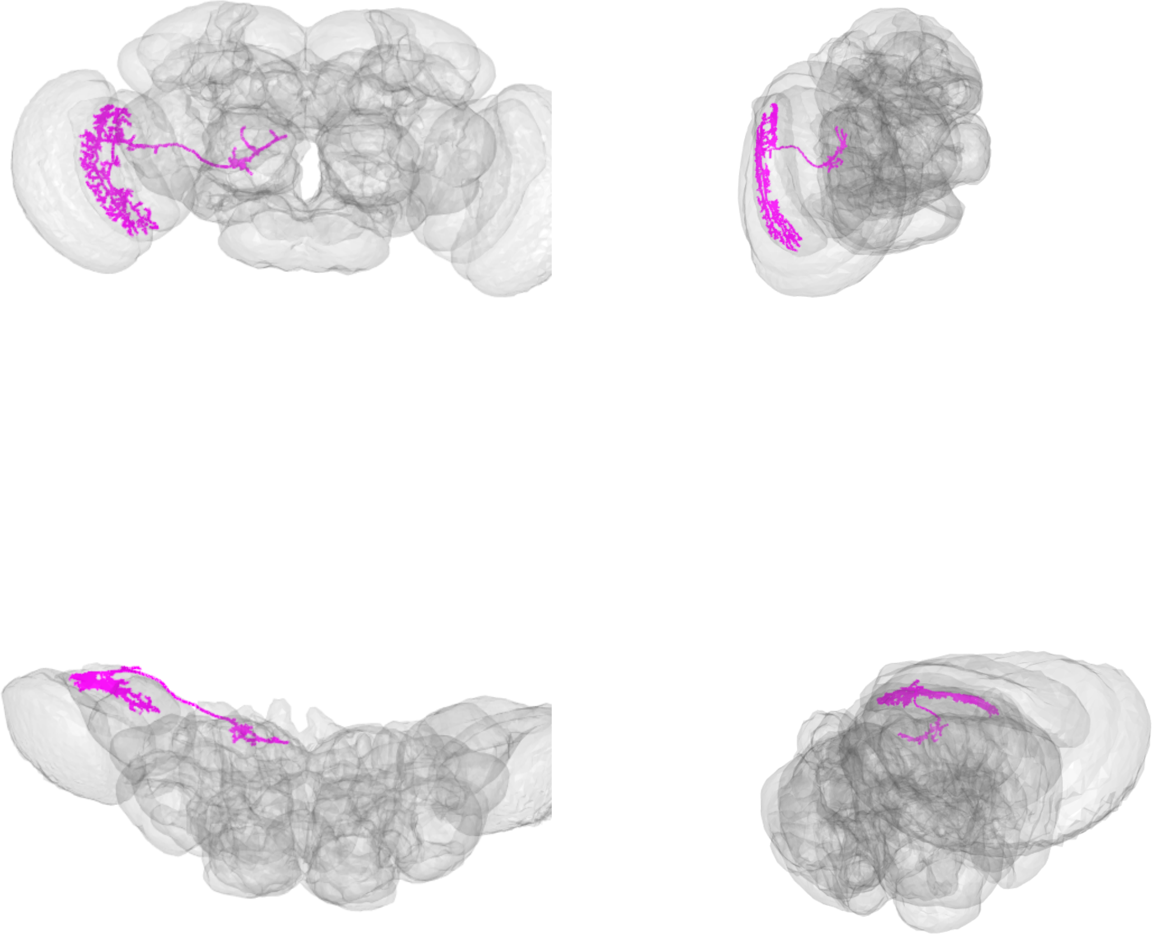

LPT13_R_VSm_skid=852286

LPT13_R_VSm_skid=815241

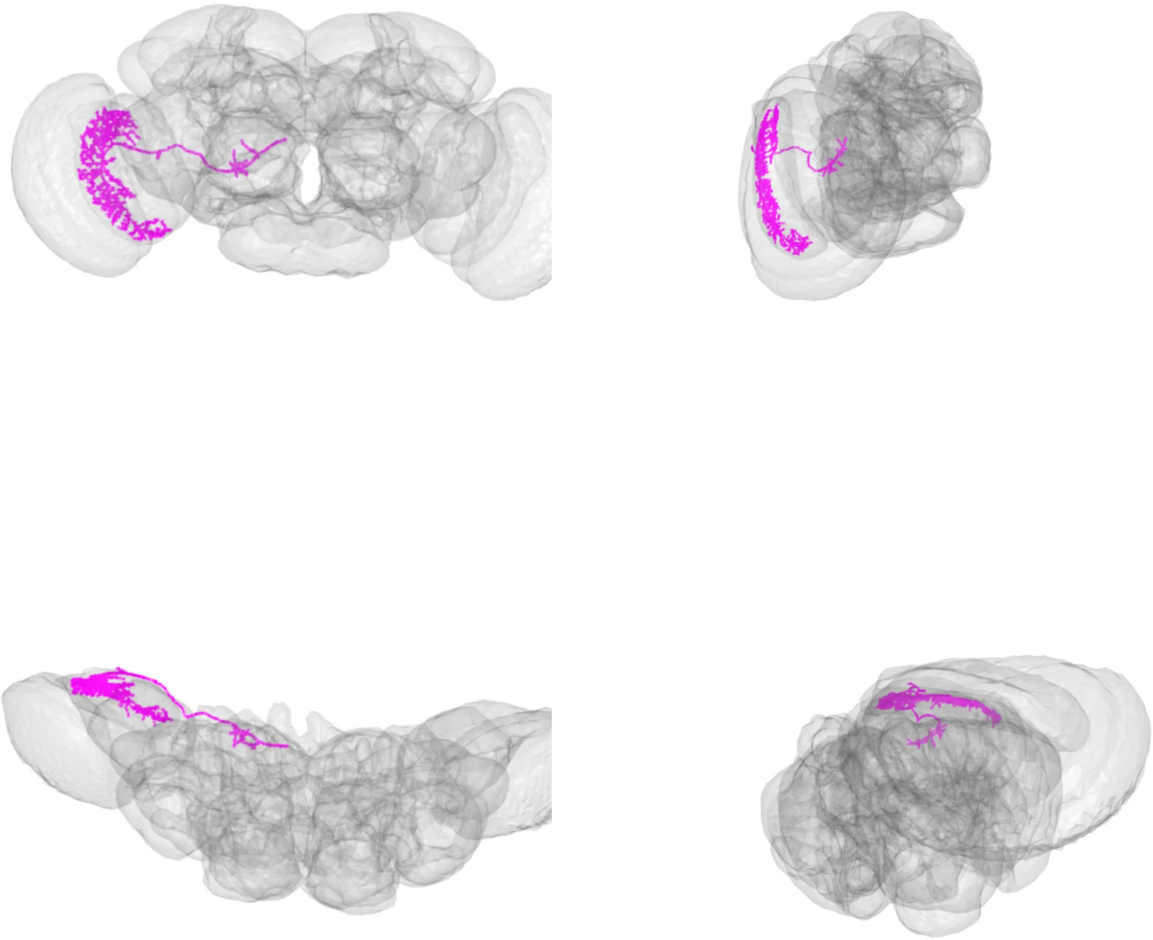

LPT13_R_VSm_skid=815241

LPT15_R_VT1_skid=1124867

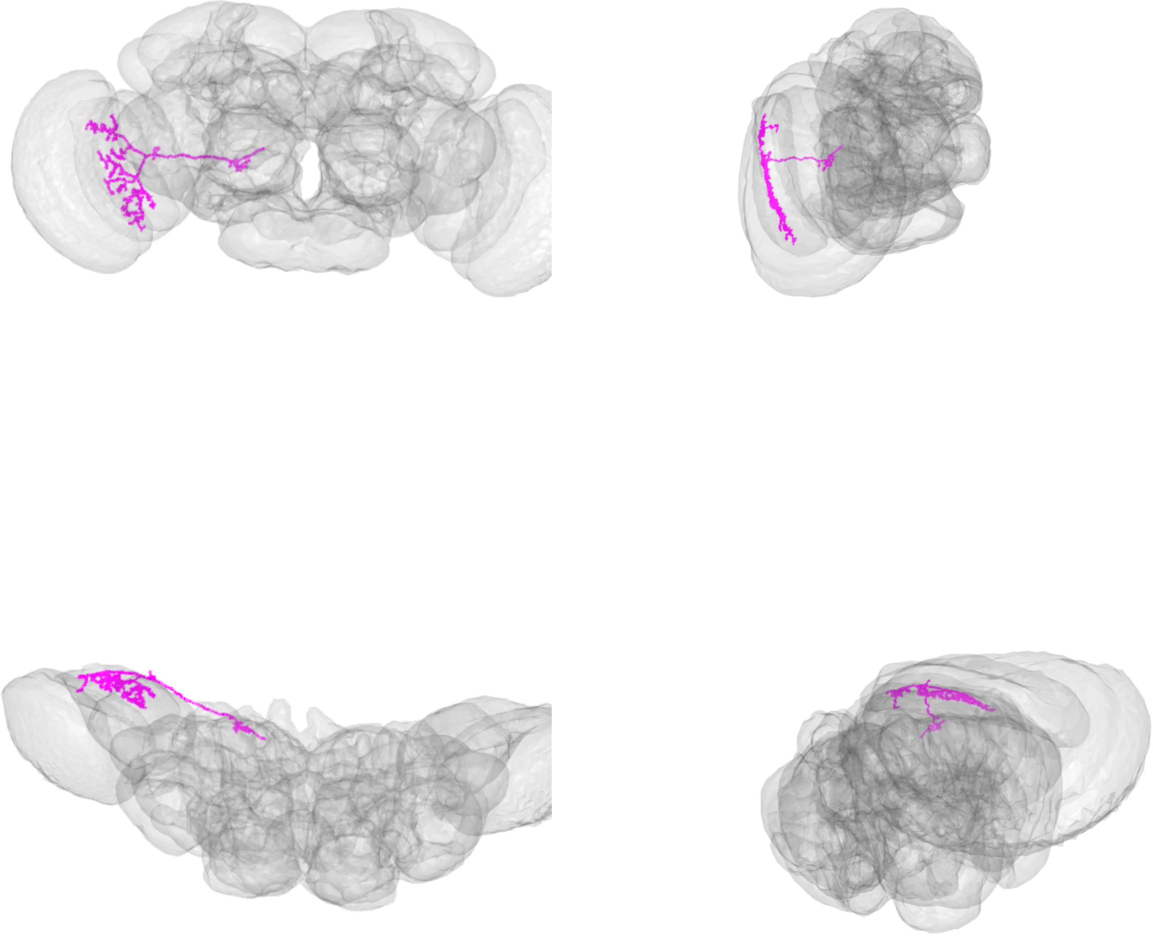

LPT15_R_VT1_skid=1124867

LPT15_R_VT1_skid=988674

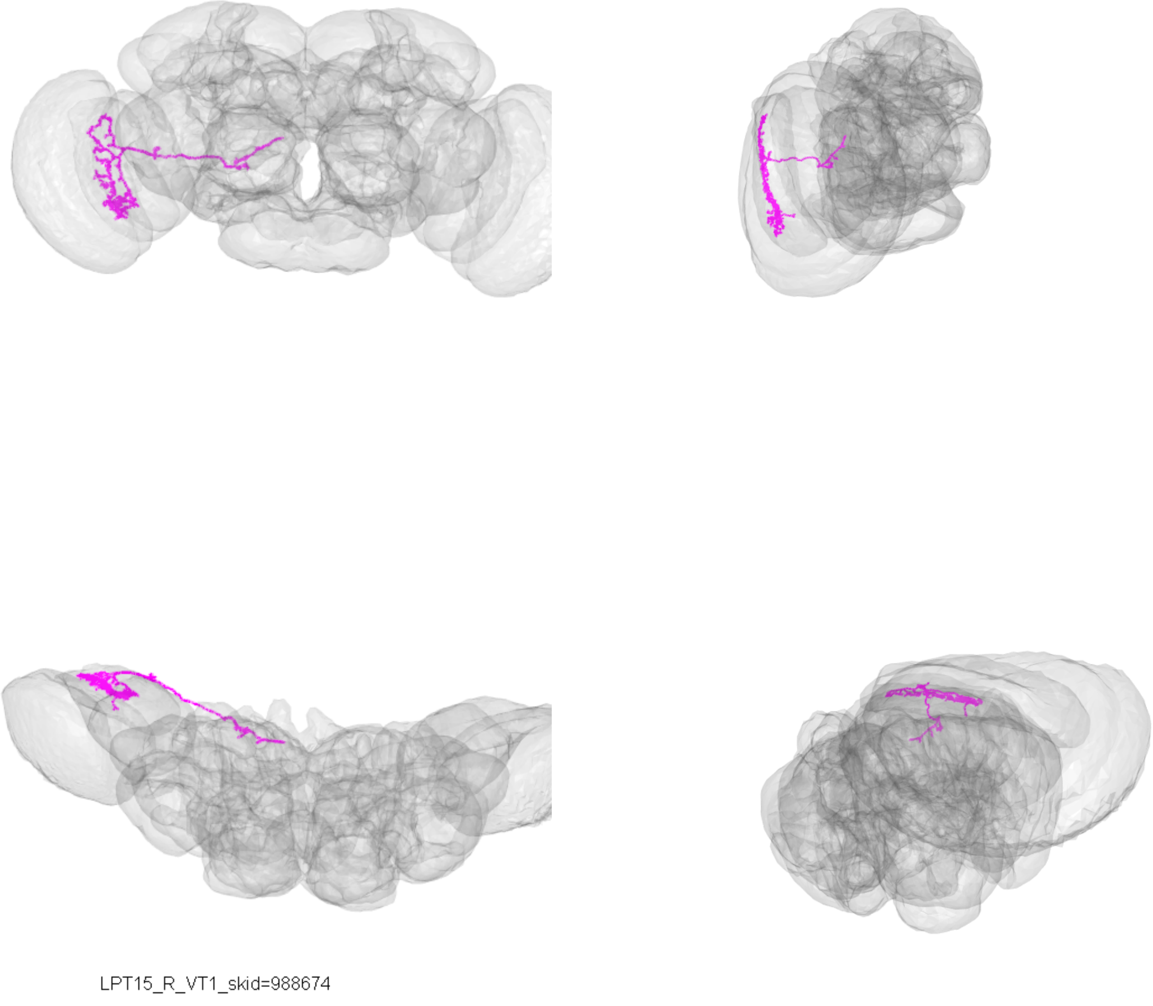

LPT15_R_VT1_skid=988674

LPT17_R_VT2_skid=1123389

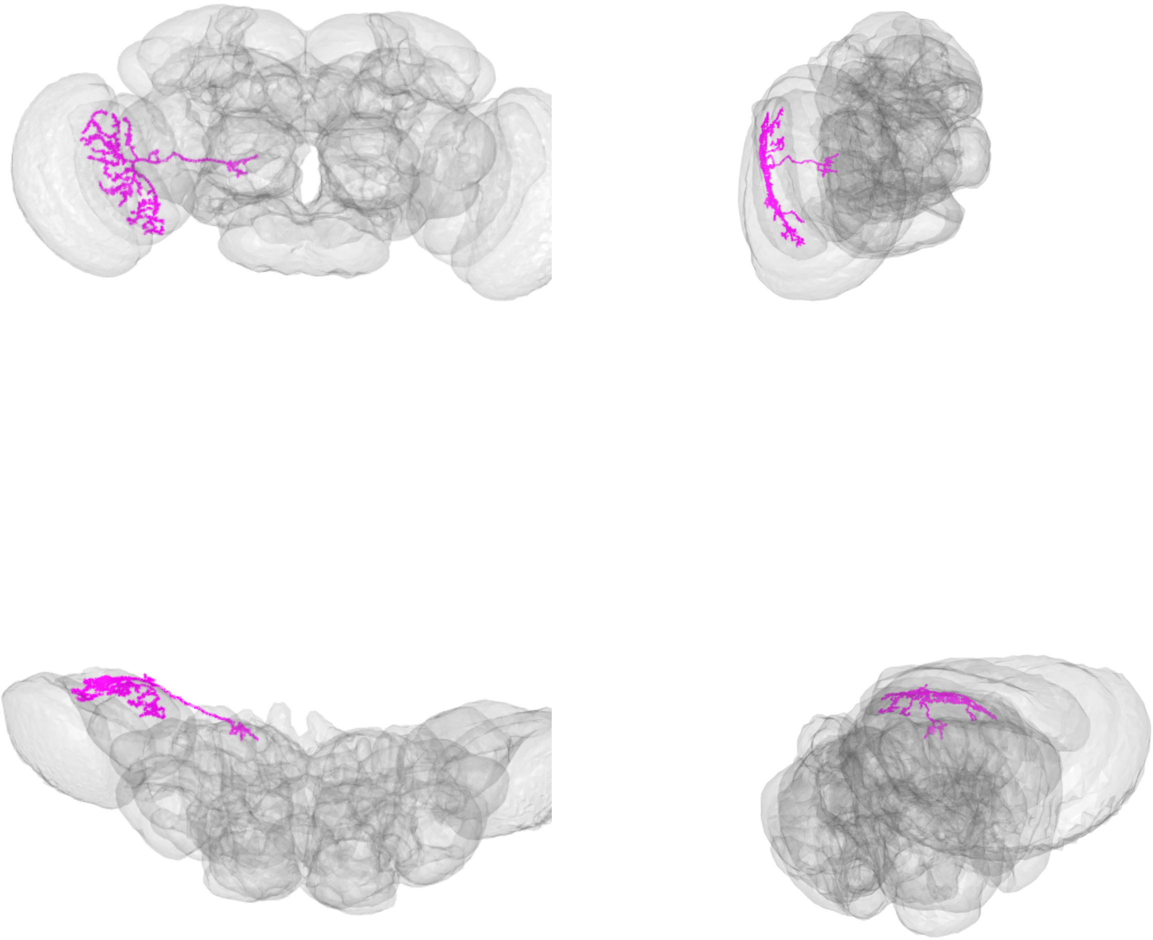

LPT17_R_VT2_skid=1123389

LPT17_R_VT2_skid=1112633

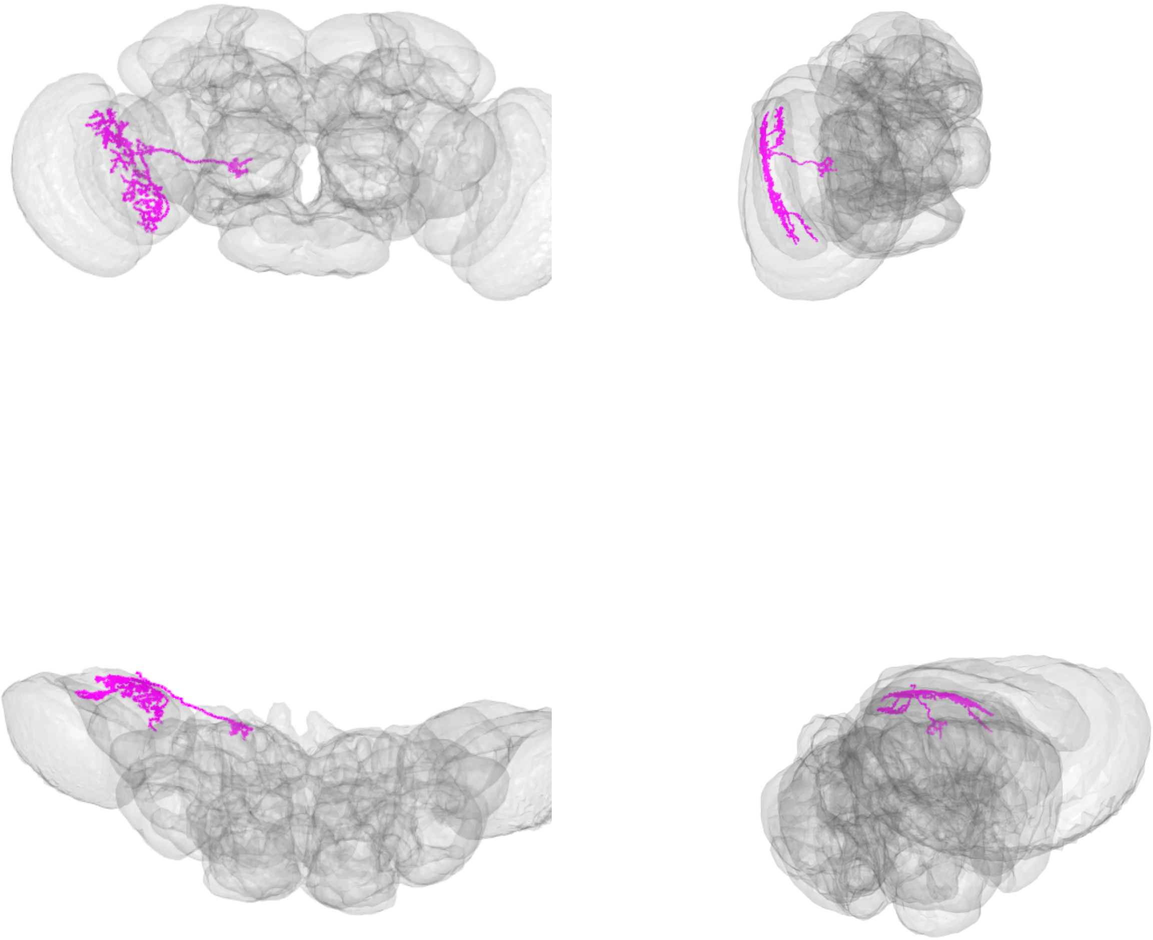

LPT17_R_VT2_skid=1112633

LPT17_R_VT2_skid=2852912

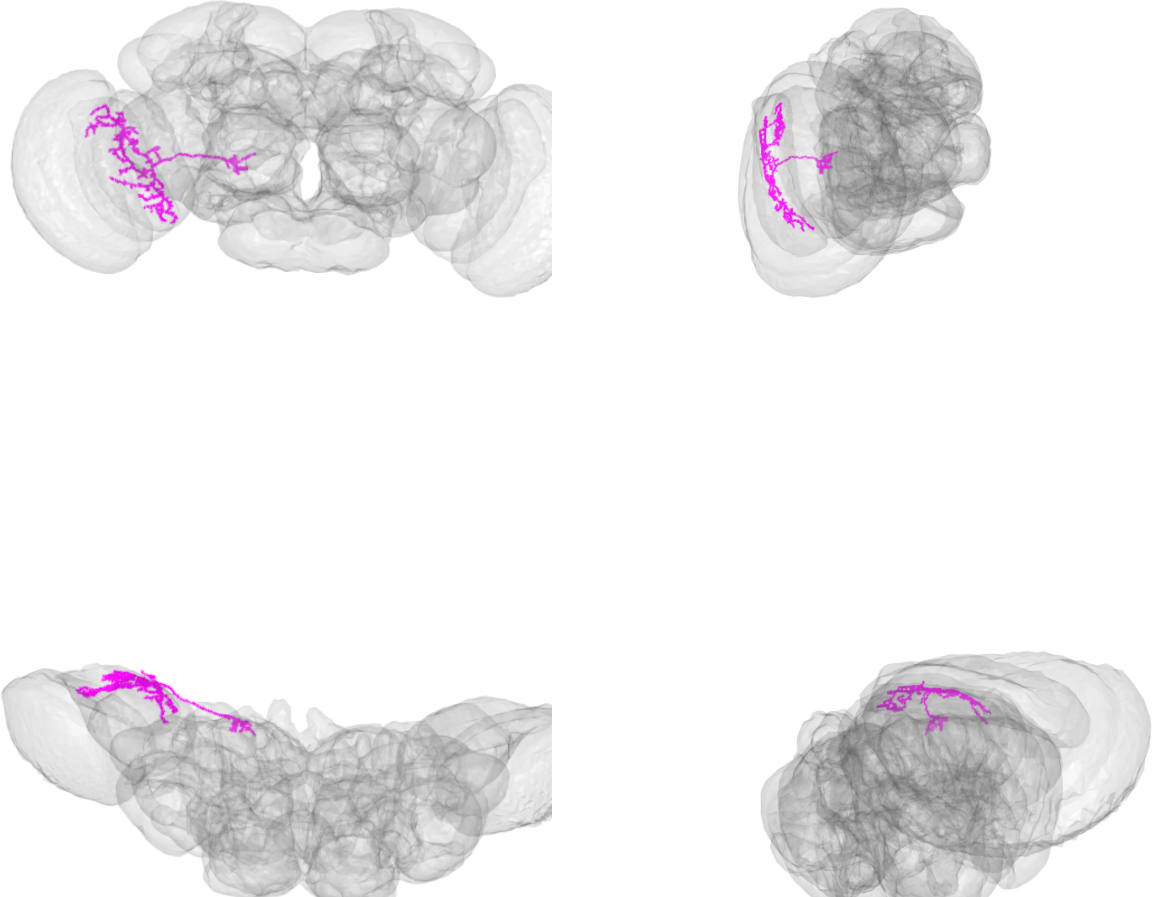

LPT17_R_VT2_skid=2852912

LPT17_R_VT2_skid=5031615

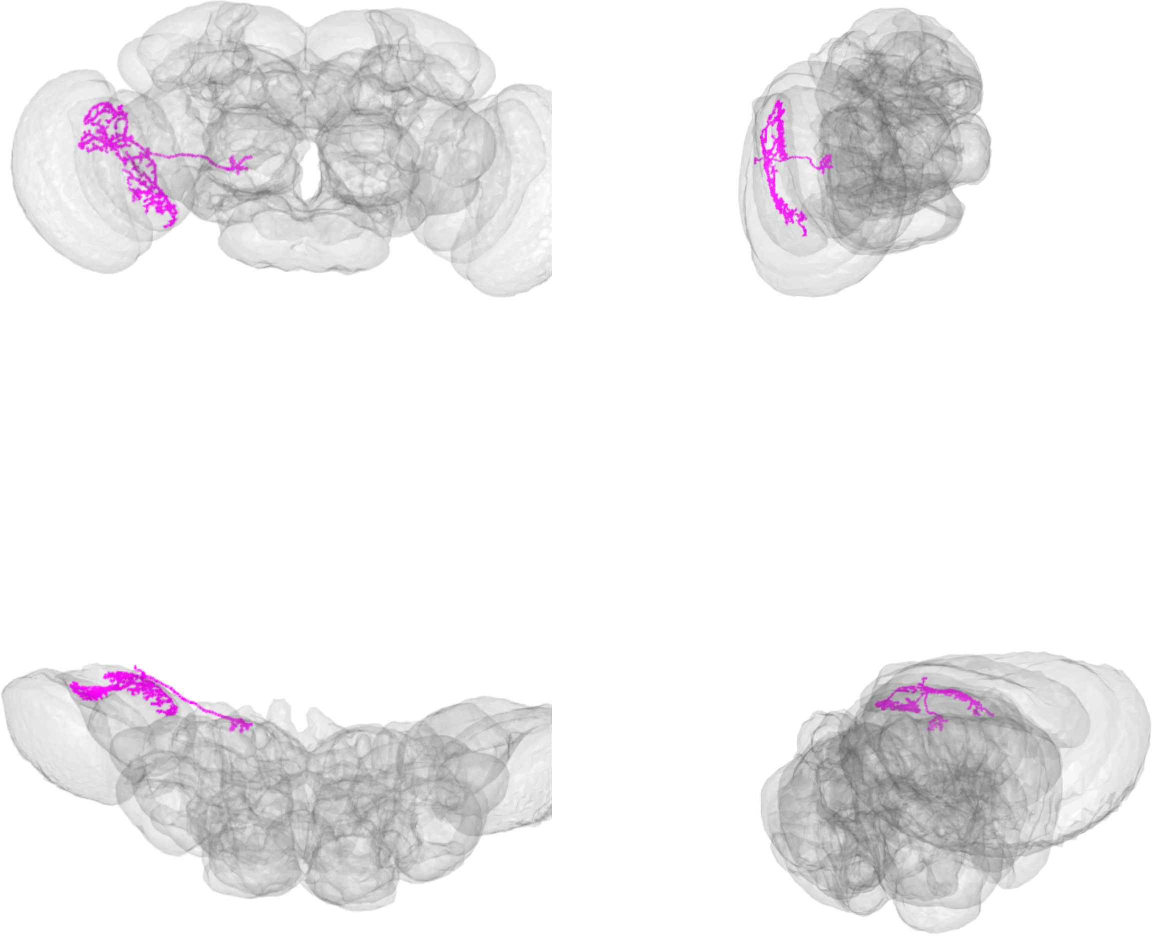

LPT17_R_VT2_skid=5031615

LPT21_R_skid=1110765

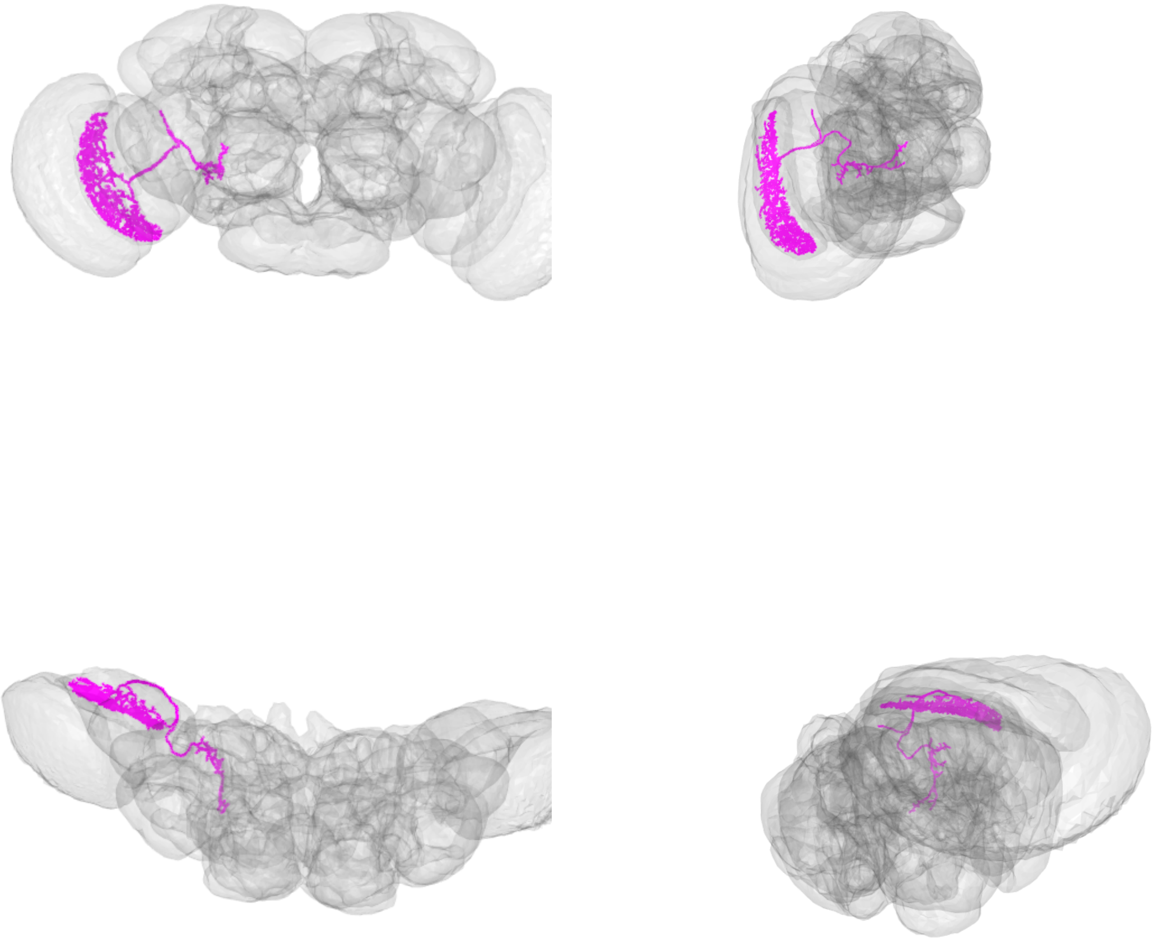

LPT21_R_skid=1110765

LPT22_R_skid=3510999

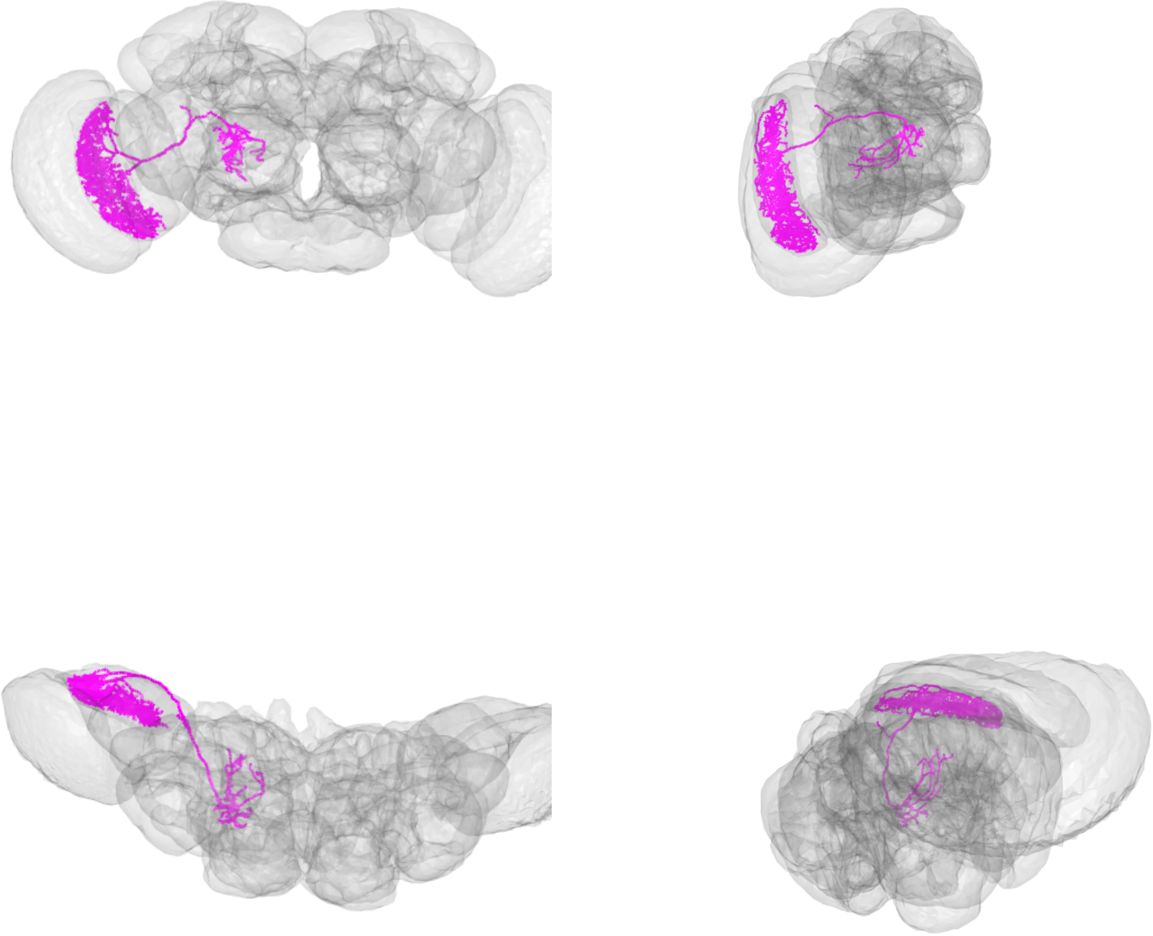

LPT22_R_skid=3510999

LPT23_R_skid=4135042

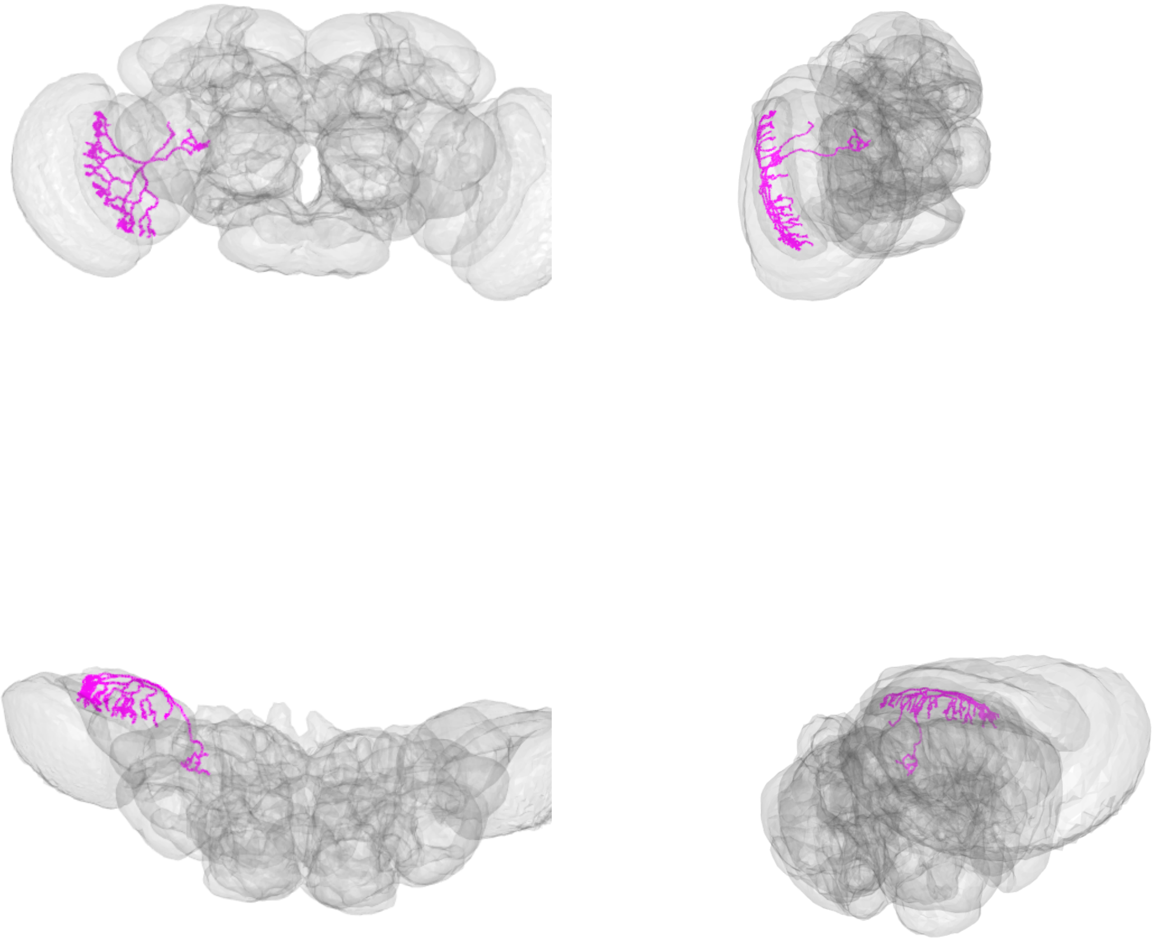

LPT23_R_skid=4135042

LPT23_R_skid=4504548

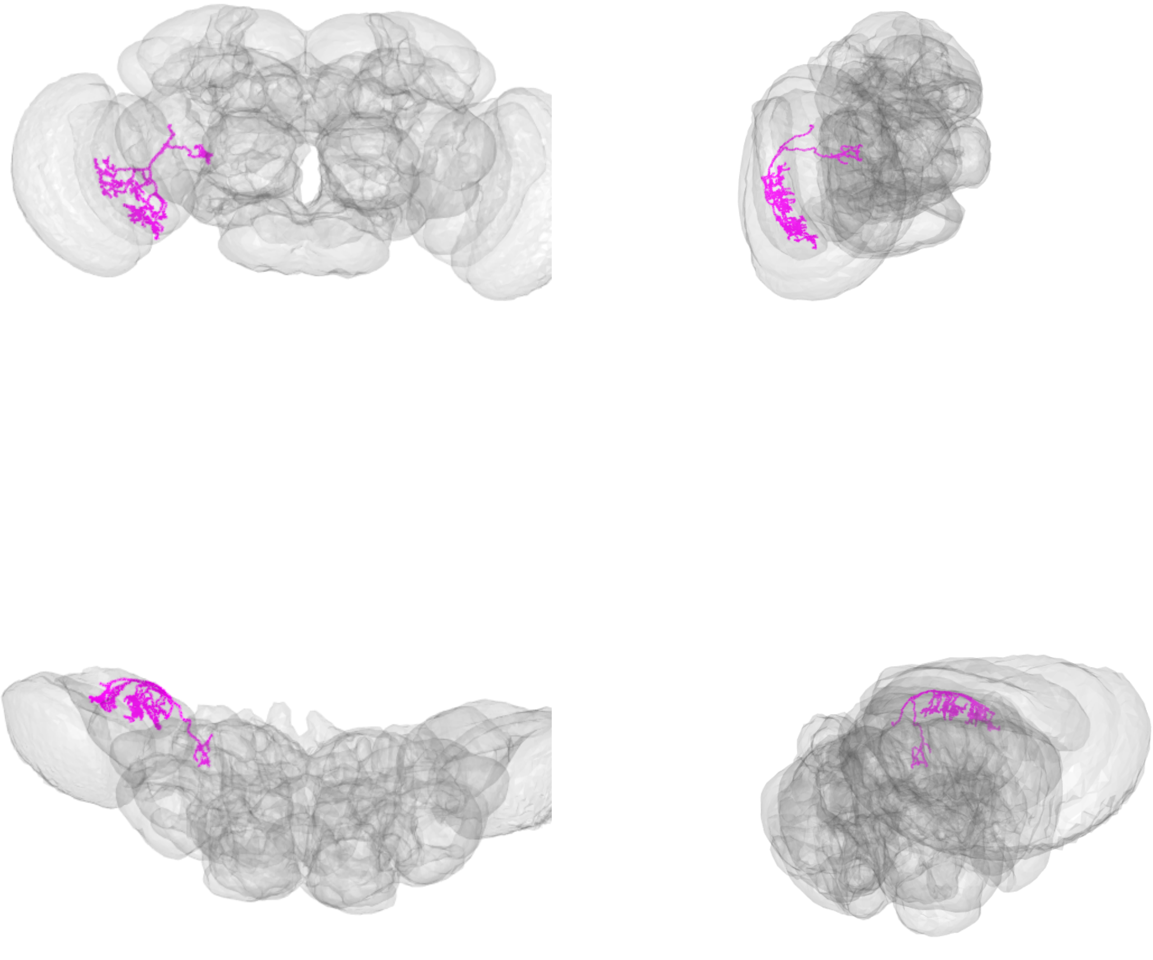

LPT23_R_skid=4504548

LPT23_R_skid=4504557

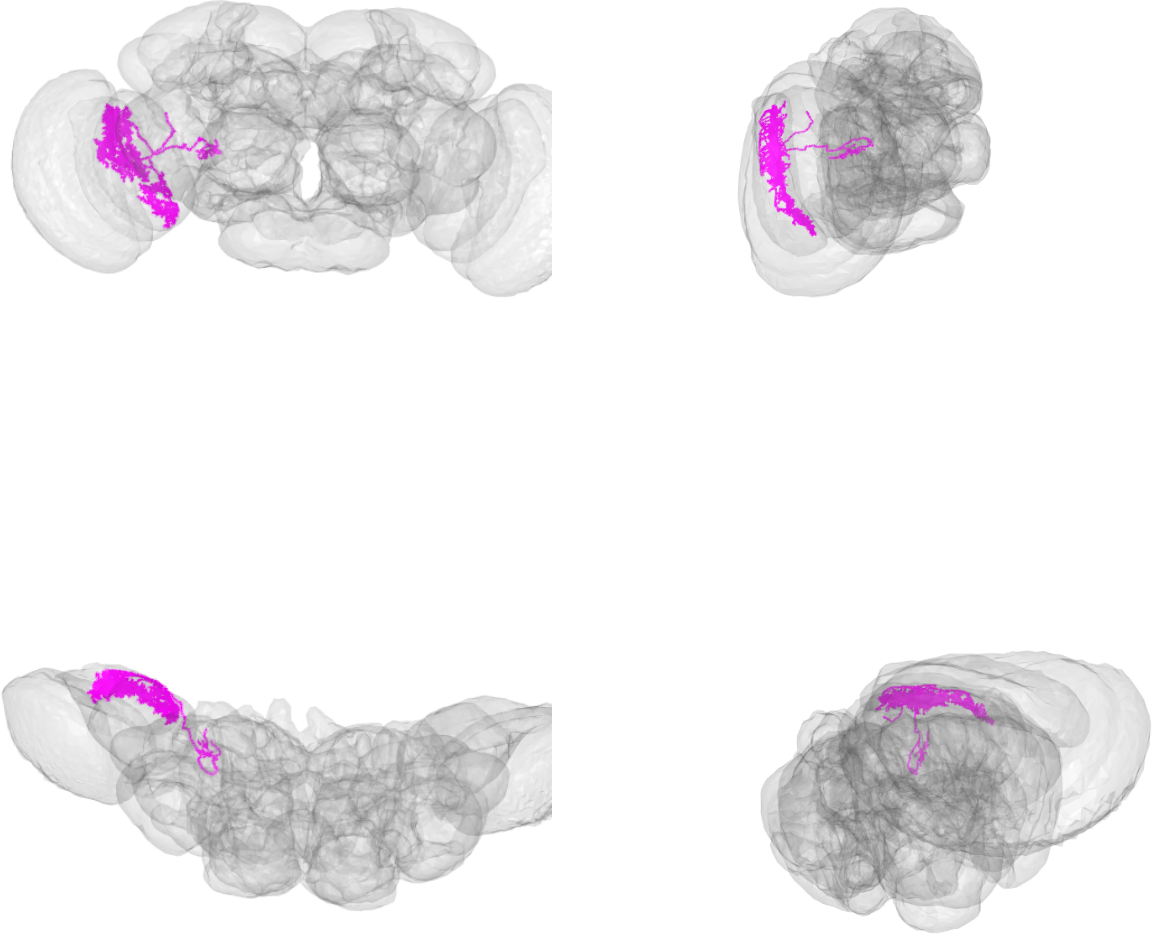

LPT23_R_skid=4504557

LPT26_R_skid=1107045

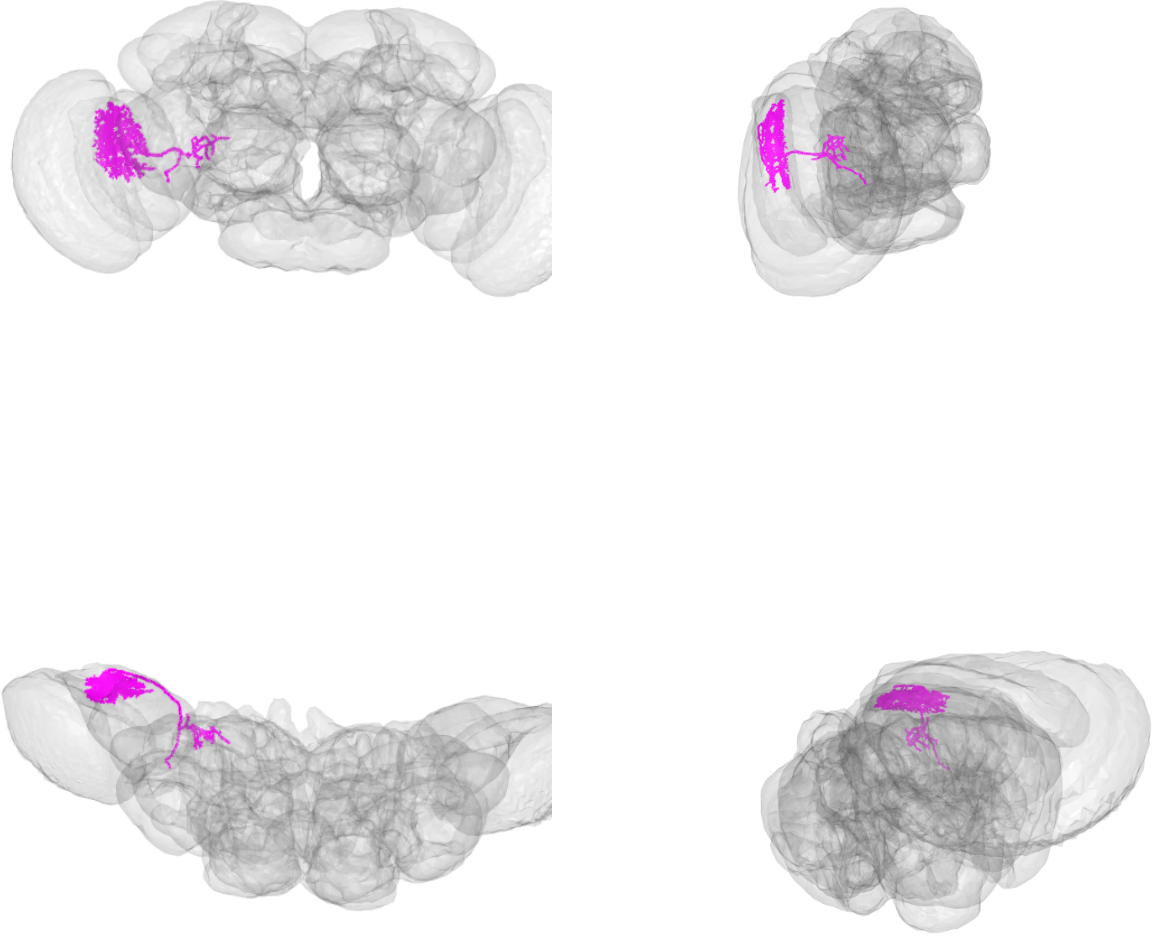

LPT26_R_skid=1107045

LPT27_R_skid=3514698

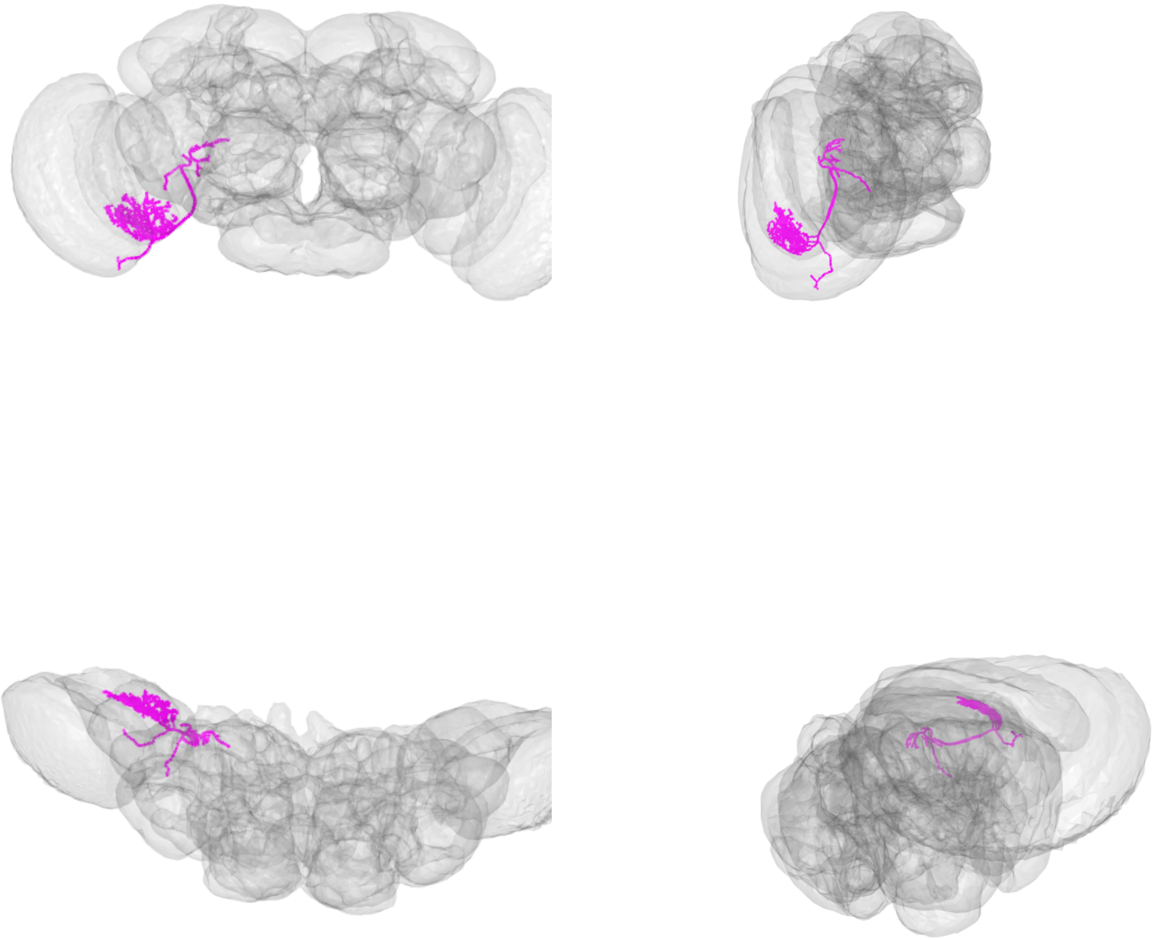

LPT27_R_skid=3514698

LPT28_R_skid=1071860

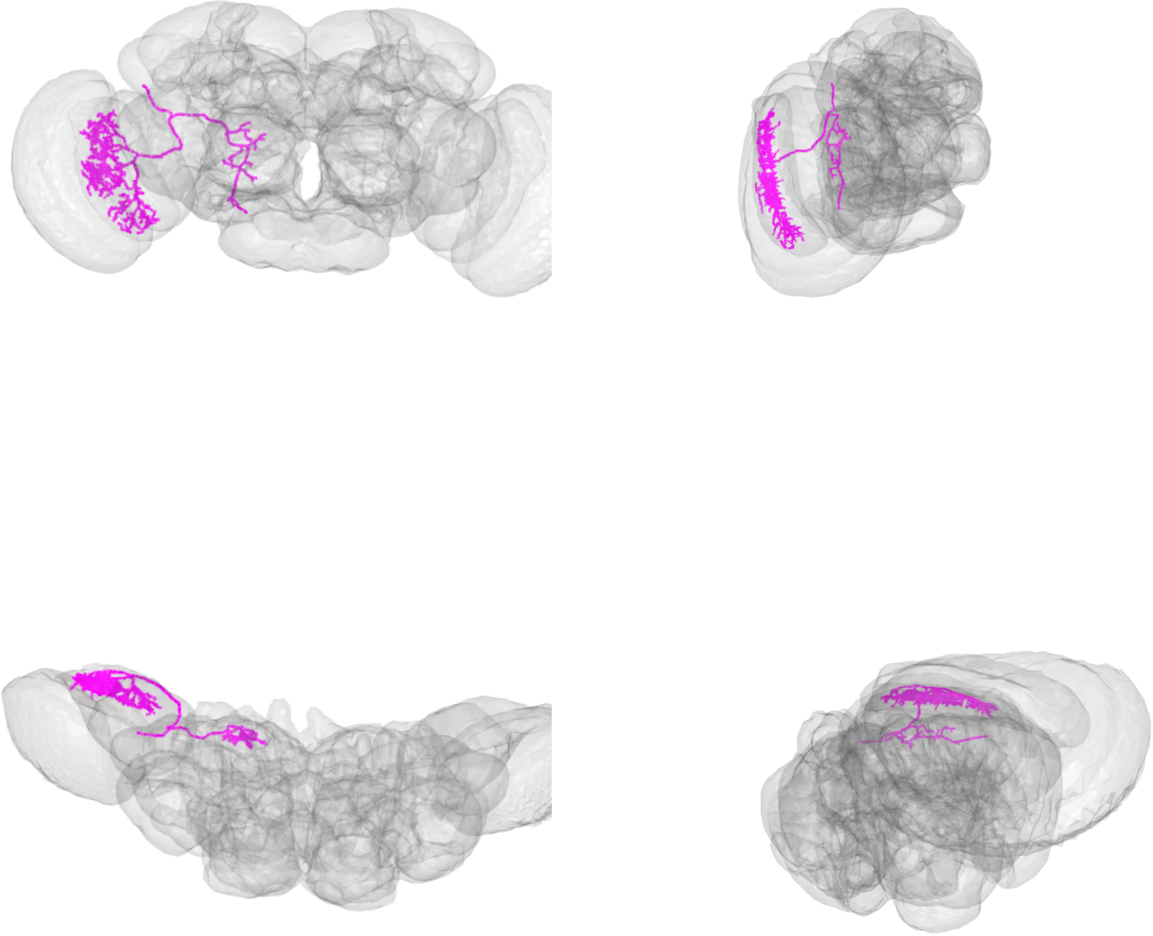

LPT28_R_skid=1071860

LPT29_R_skid=7760435

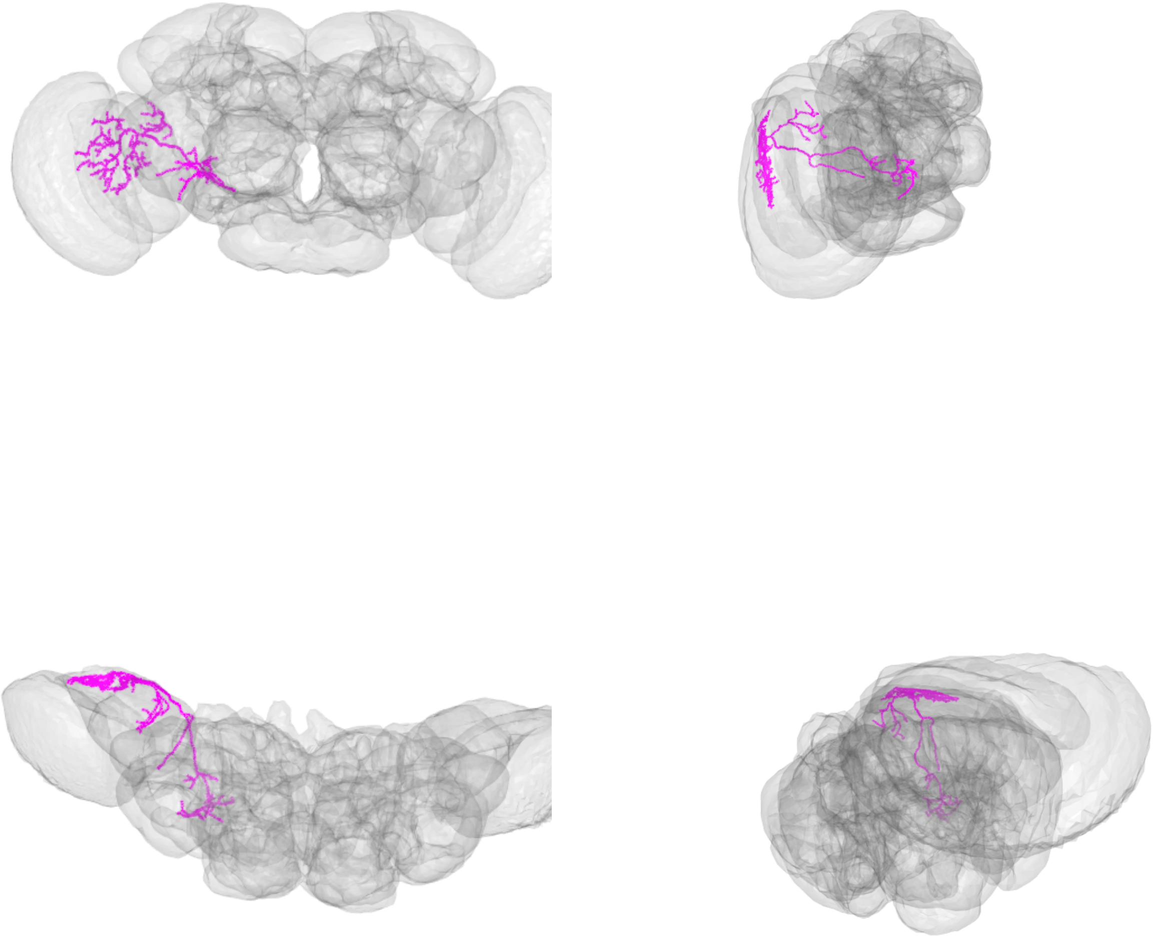

LPT29_R_skid=7760435

LPT30_R_skid=7674107

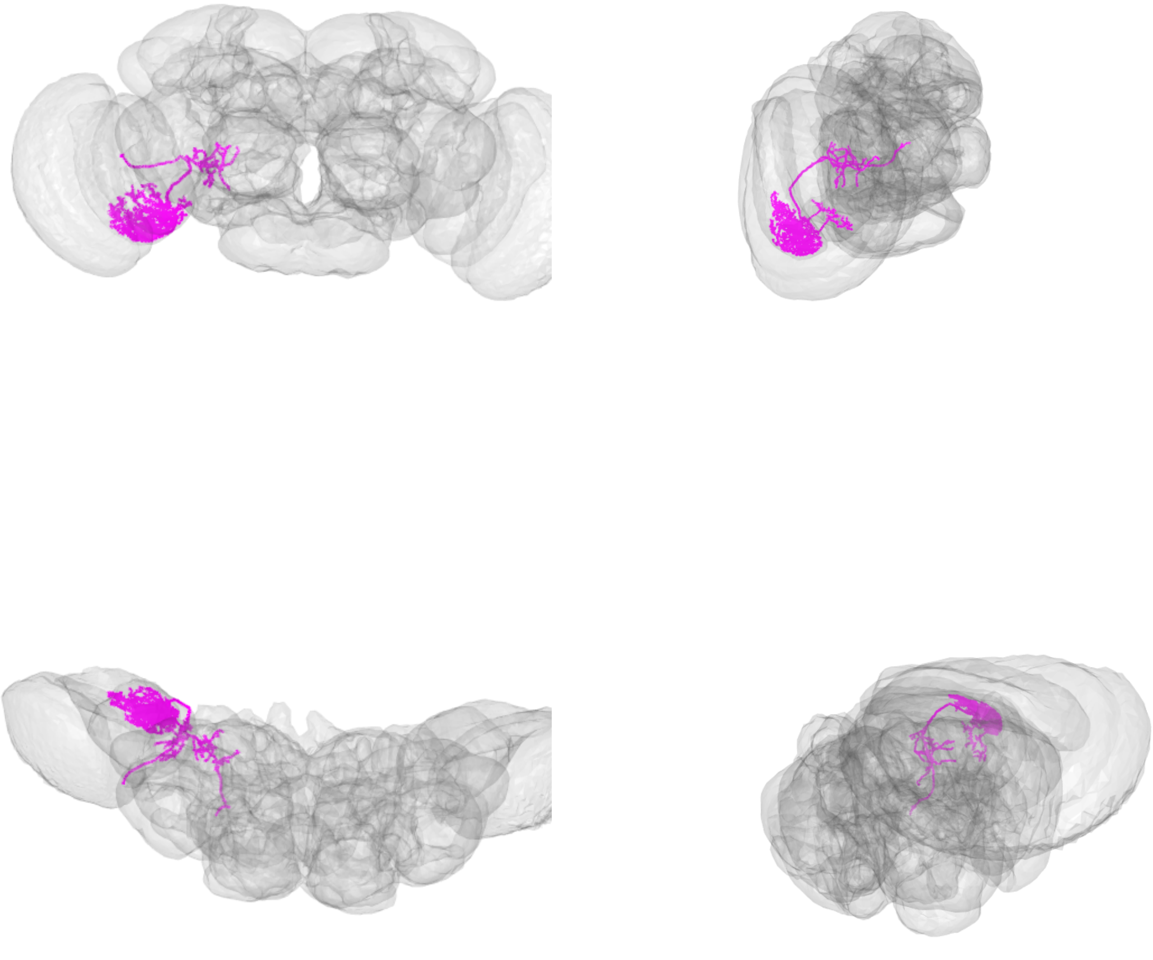

LPT30_R_skid=7674107

LPT31_R_skid=7690273

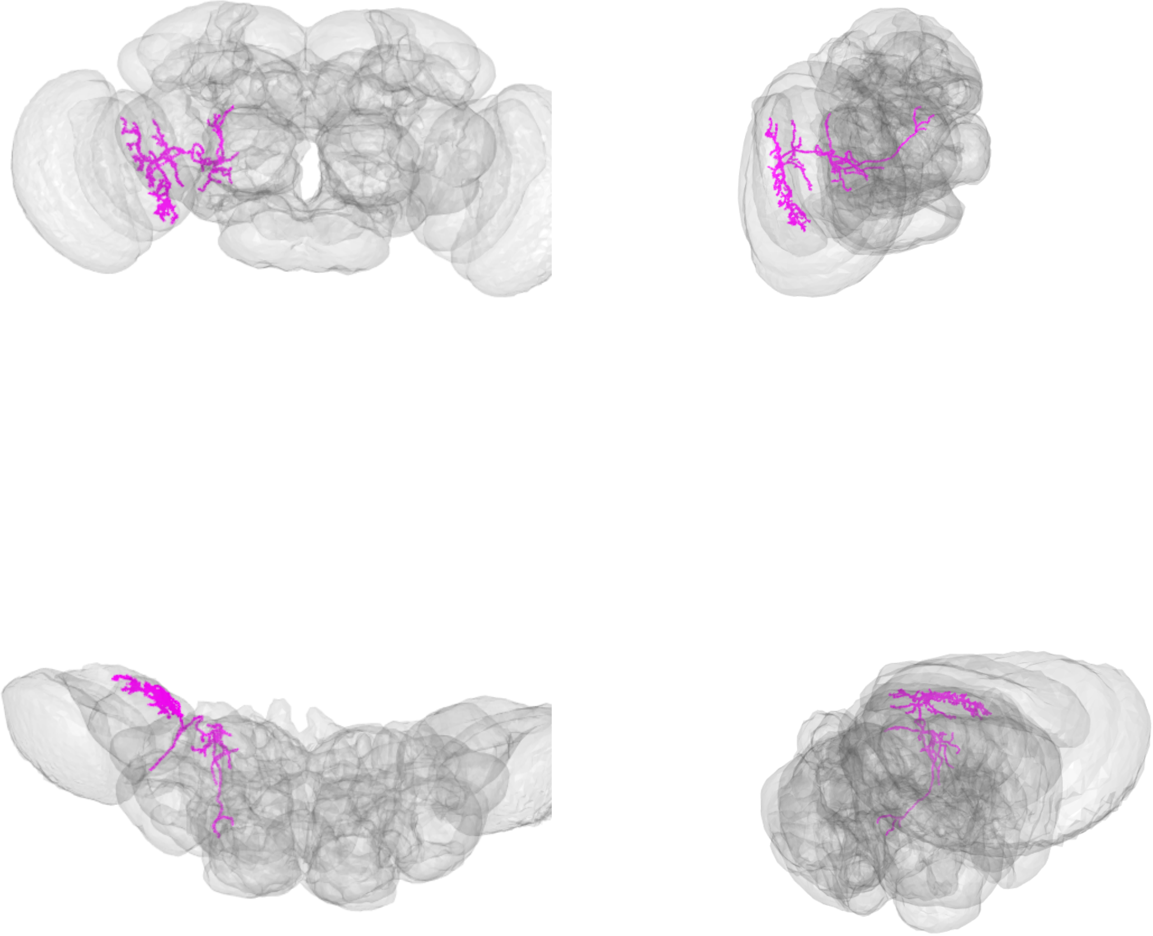

LPT31_R_skid=7690273

LPT31_R_skid=7311408

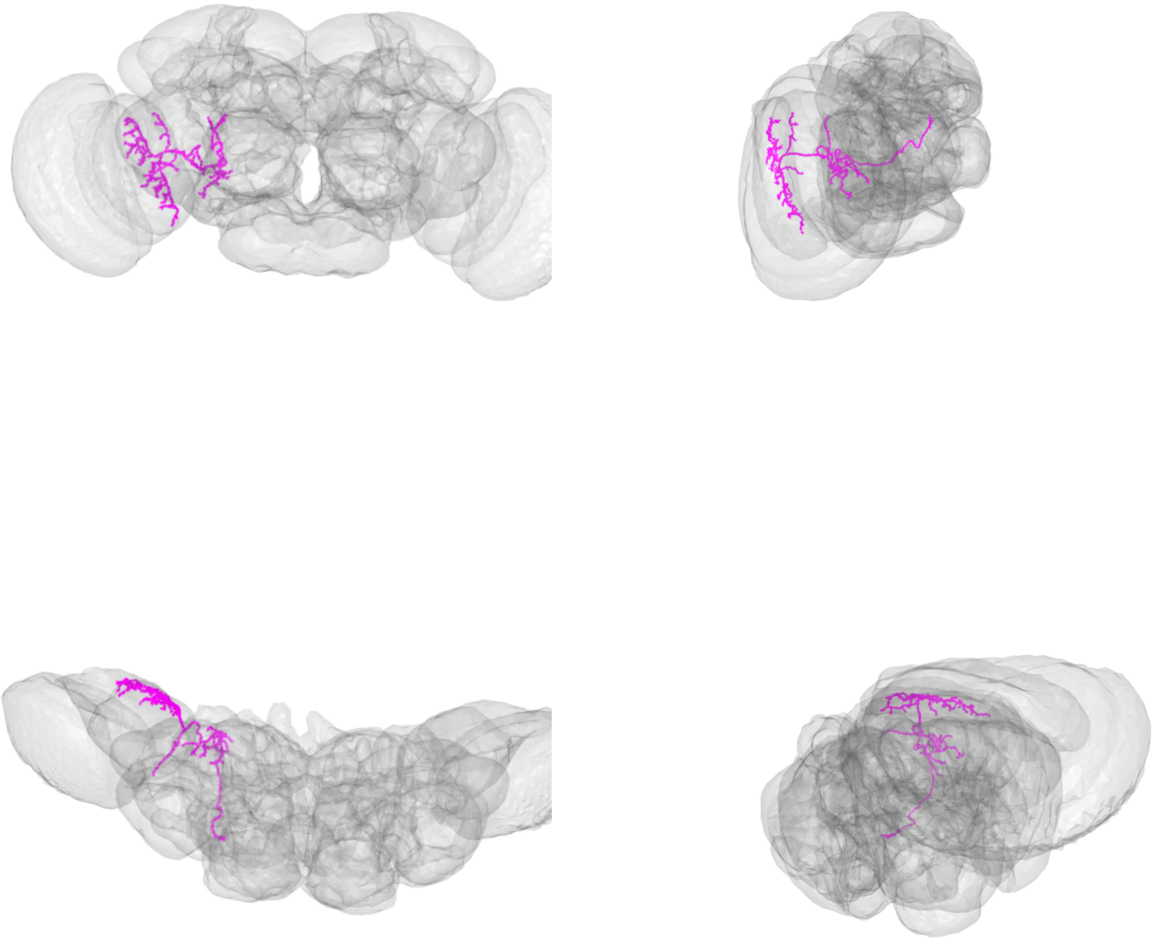

LPT31_R_skid=7311408

LPT31_R_skid=7311493

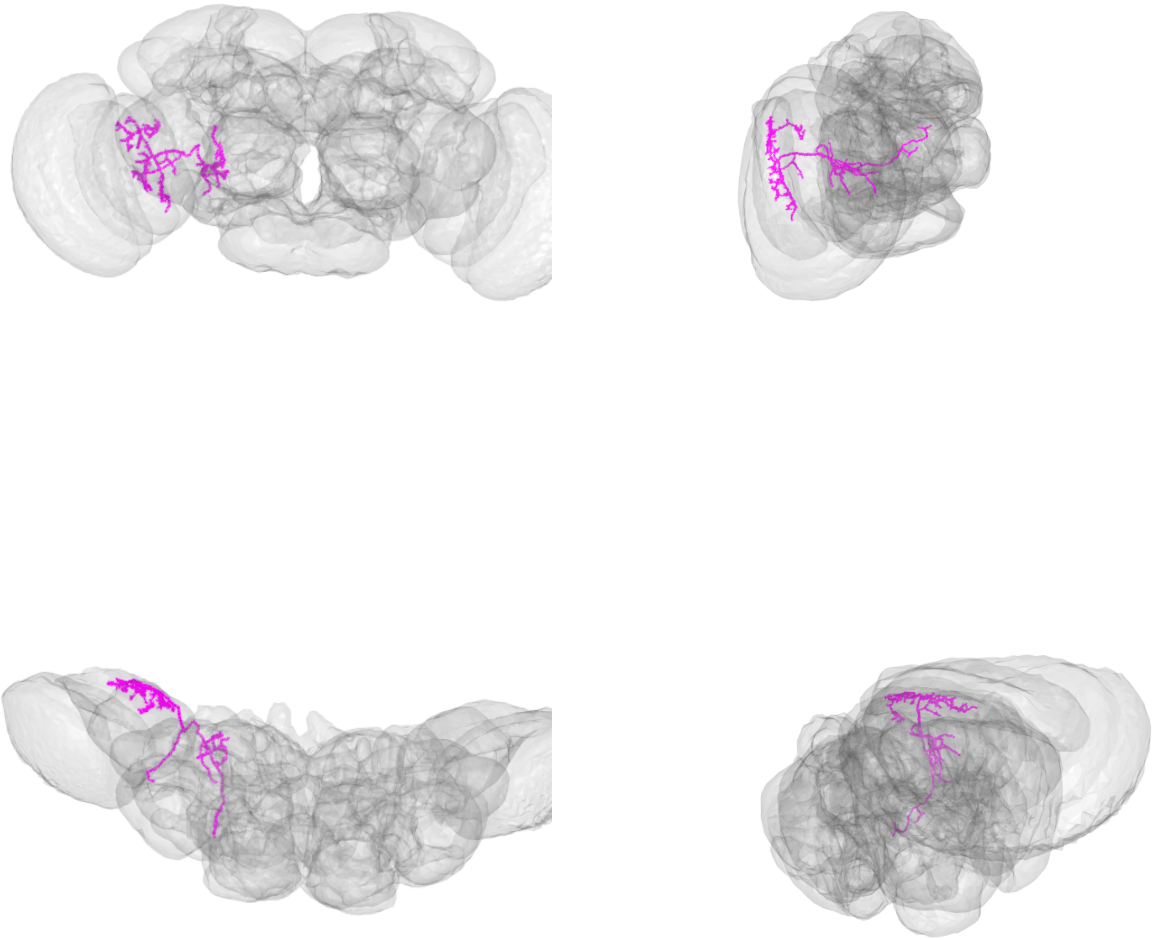

LPT31_R_skid=7311493

LPT31_R_skid=7694305

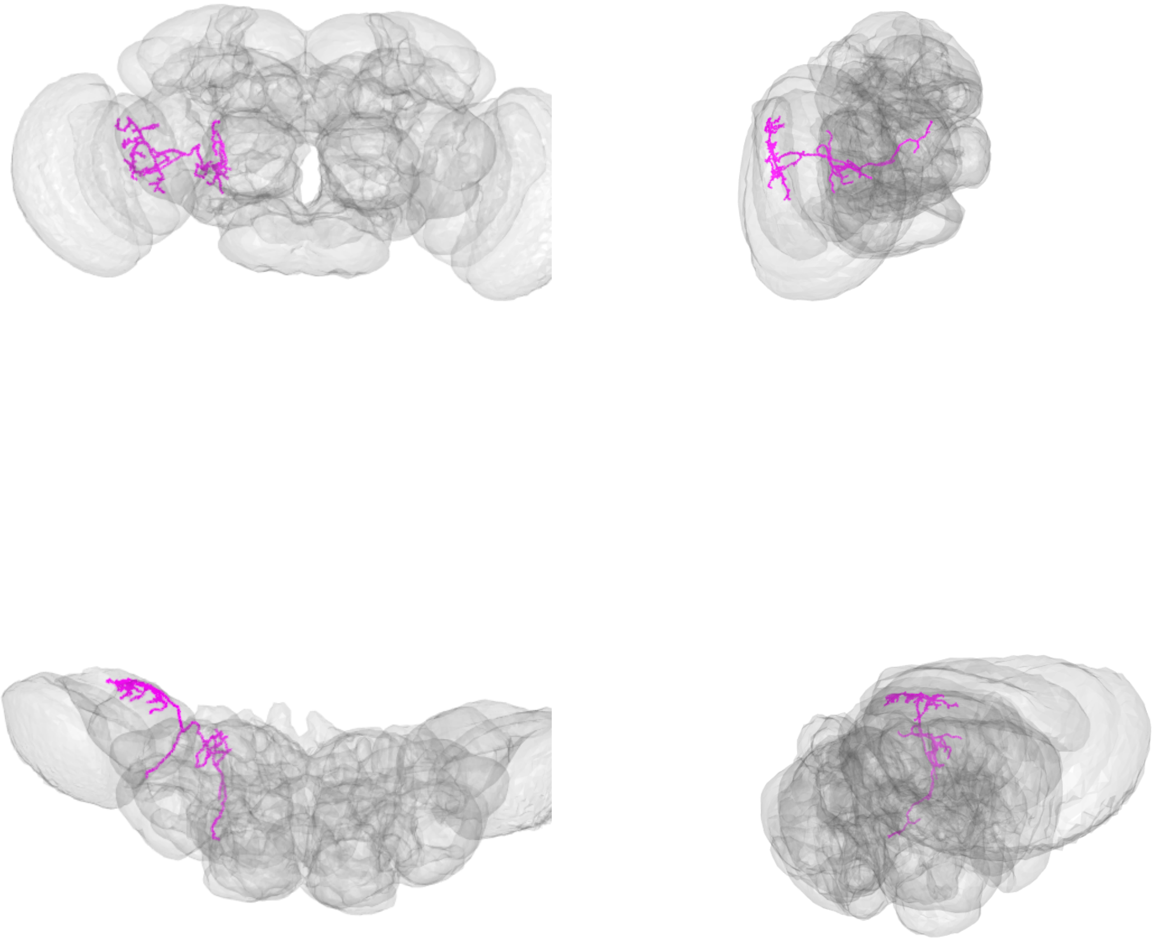

LPT31_R_skid=7694305

LPT35_R_dCH_skid=1077174

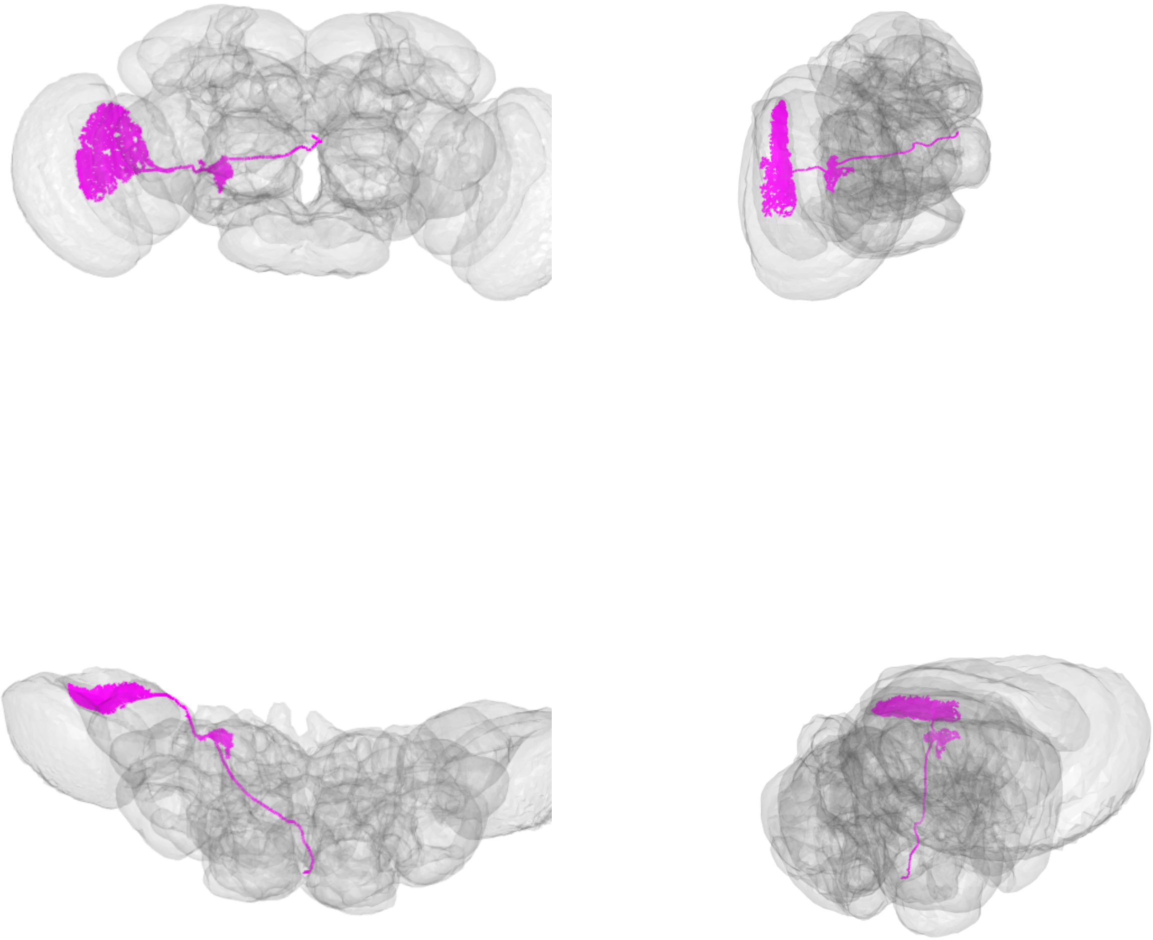

LPT35_R_dCH_skid=1077174

LPT36_R_vCH_skid=1078535

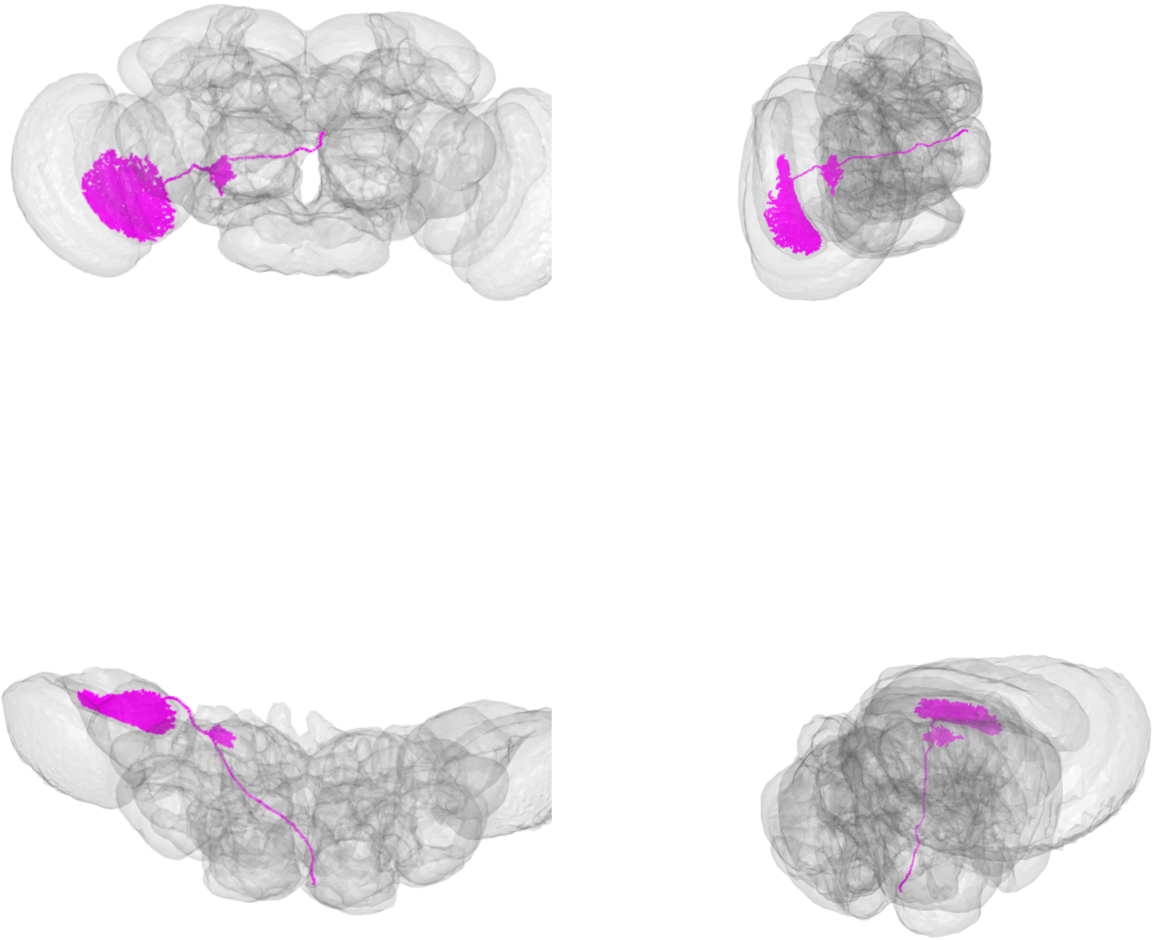

LPT36_R_vCH_skid=1078535

LPT37_R_H1_skid=1121730

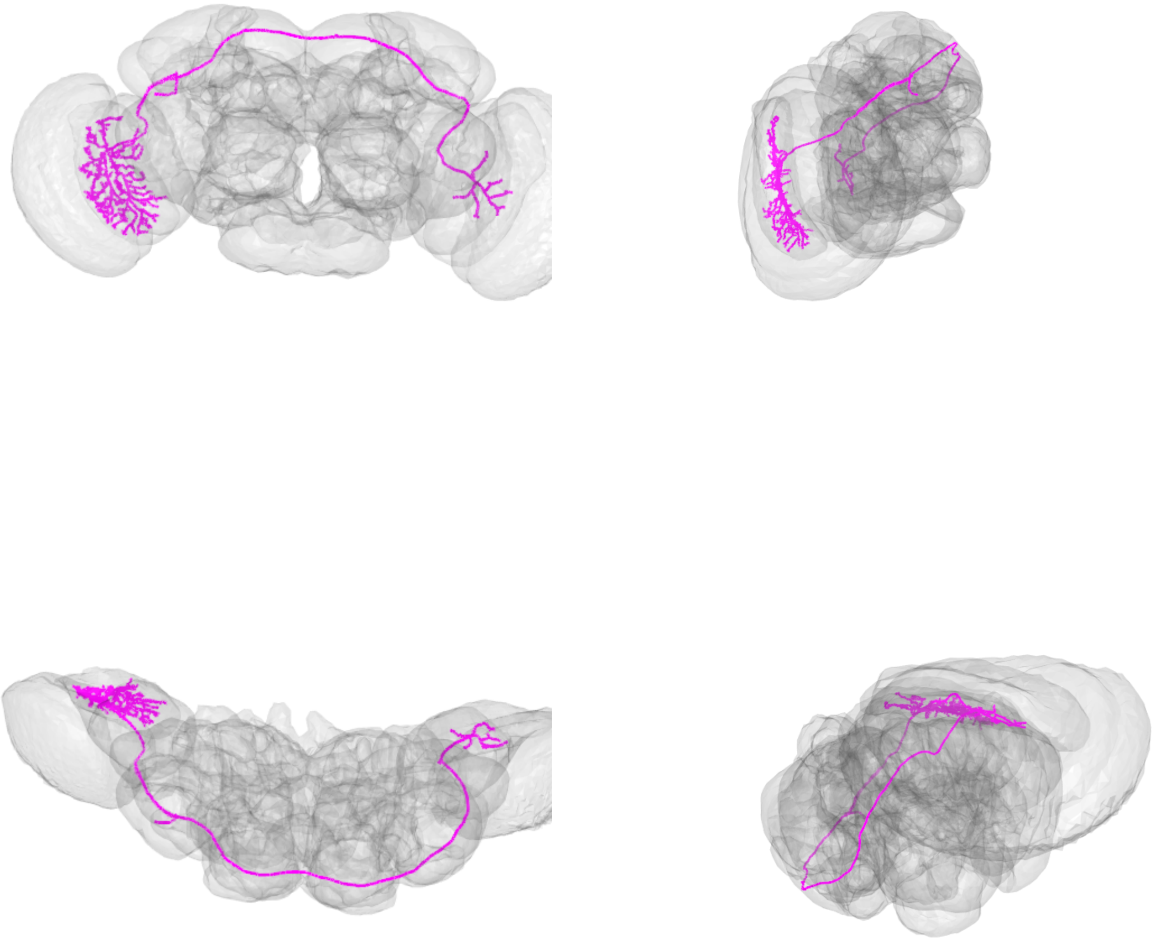

LPT37_R_H1_skid=1121730

LPT38_R_Nod1_skid=1054753

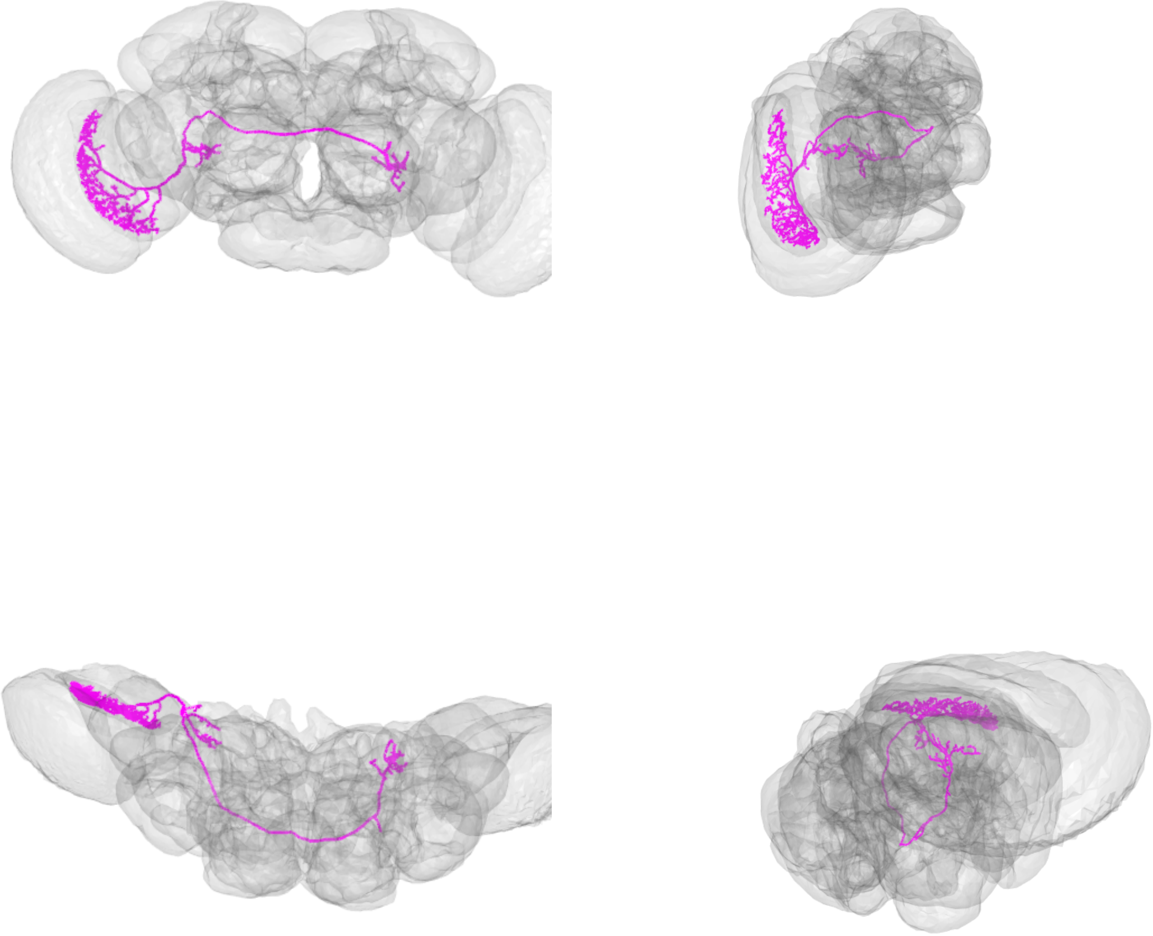

LPT38_R_Nod1_skid=1054753

LPT38_R_Nod1_skid=3546483

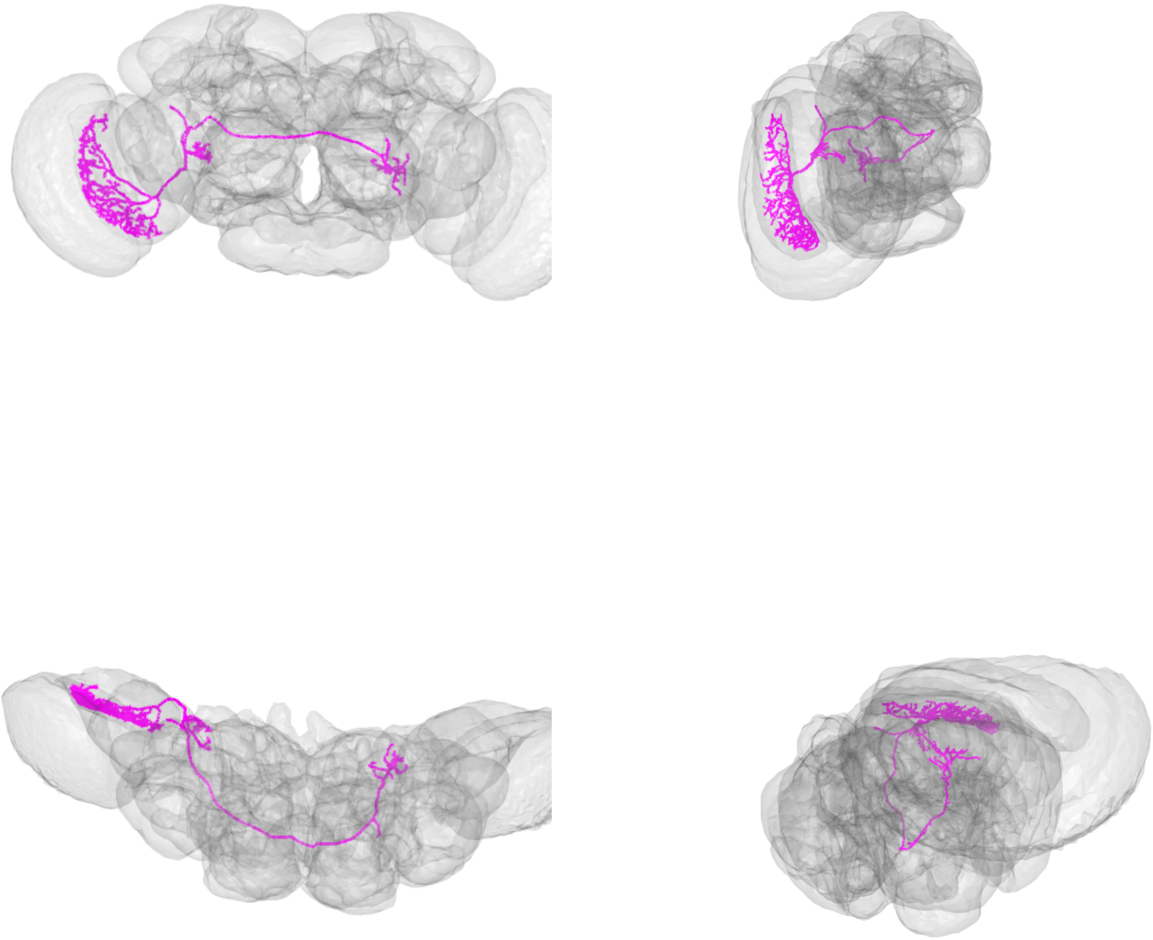

LPT38_R_Nod1_skid=3546483

LPT40_R_Nod2_skid=902072

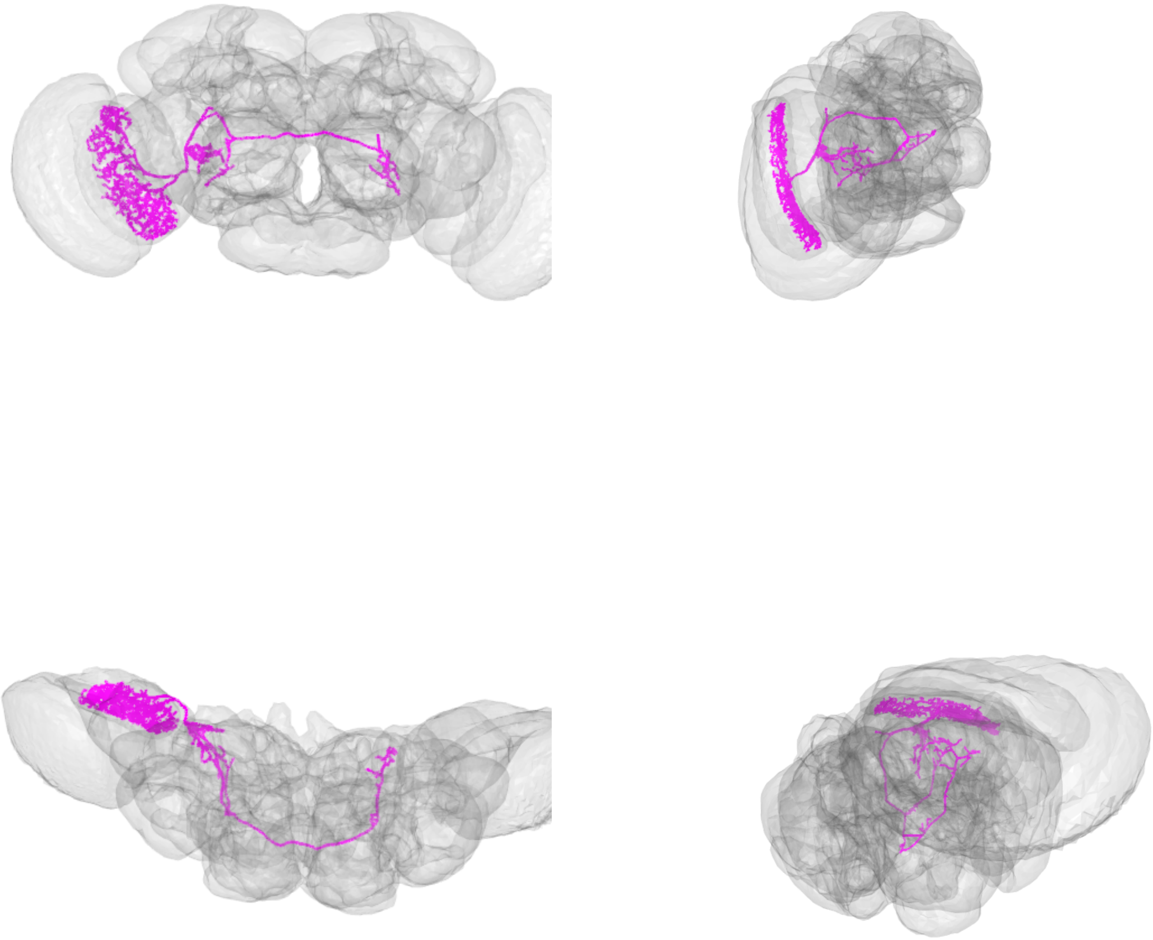

LPT40_R_Nod2_skid=902072

LPT41_R_Nod3_skid=1058996

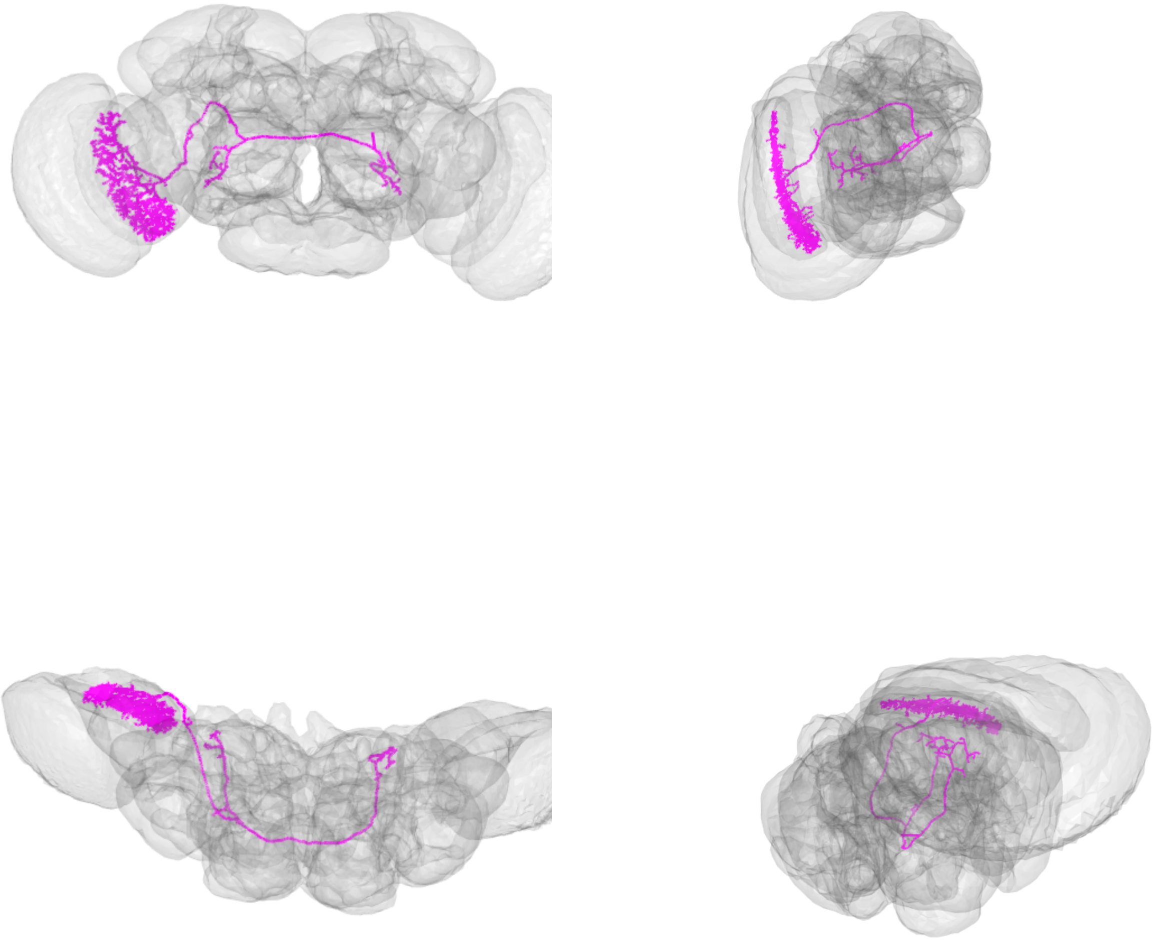

LPT41_R_Nod3_skid=1058996

LPT42_R_Nod4_skid=1106958

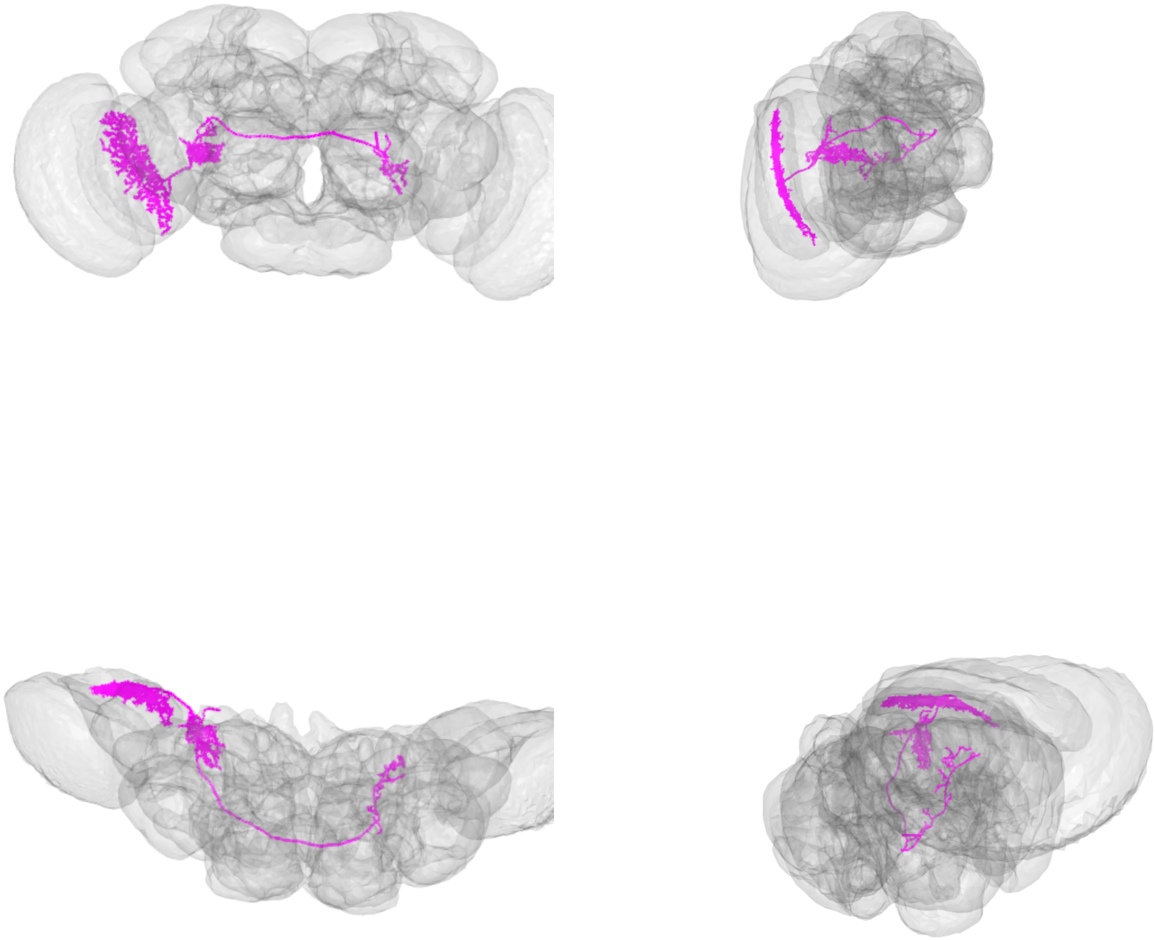

LPT42_R_Nod4_skid=1106958

LPT43_R_H2_skid=1088678

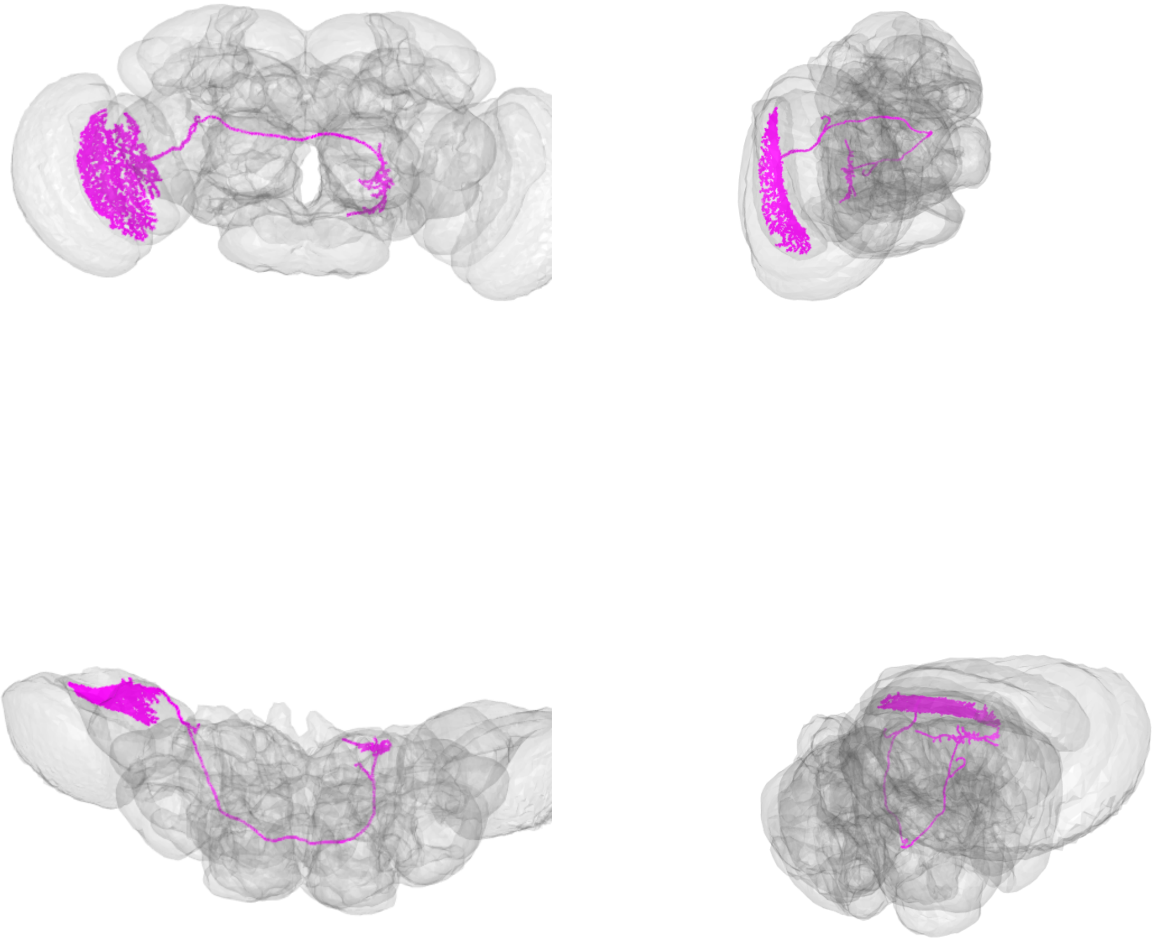

LPT43_R_H2_skid=1088678

LPT44_R_Nod5_skid=1121795

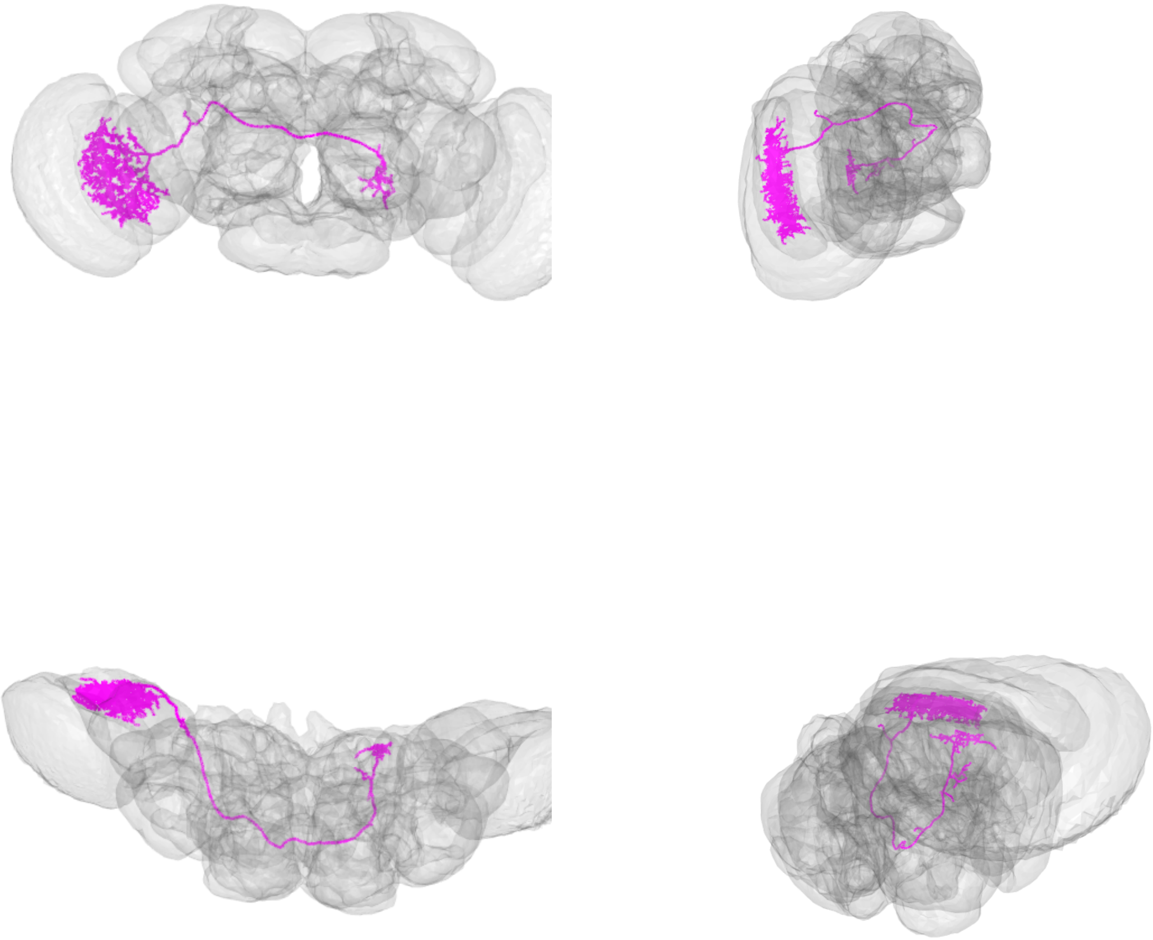

LPT44_R_Nod5_skid=1121795

LPT45_R_dCal1_skid=7204844

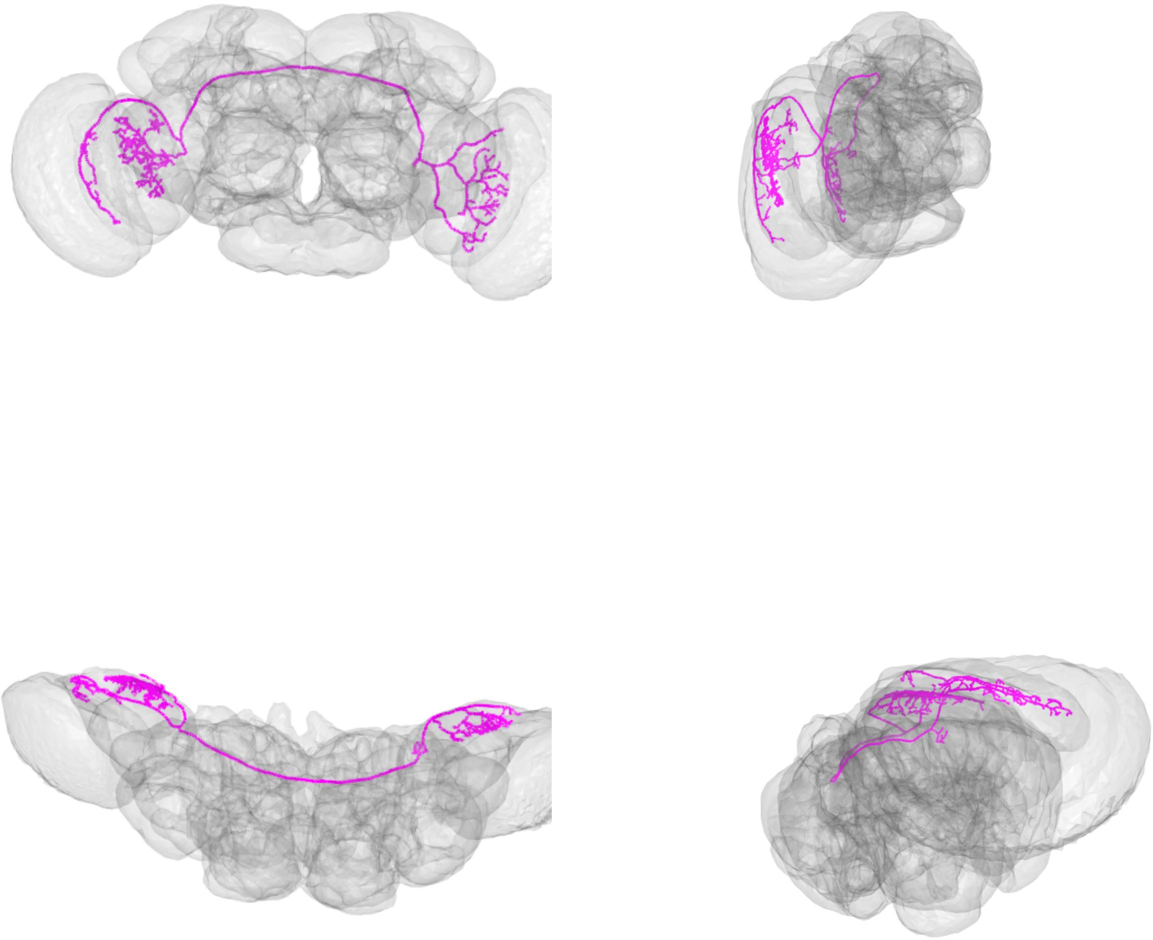

LPT45_R_dCal1_skid=7204844

LPT46_R_vCal1_skid=7510076

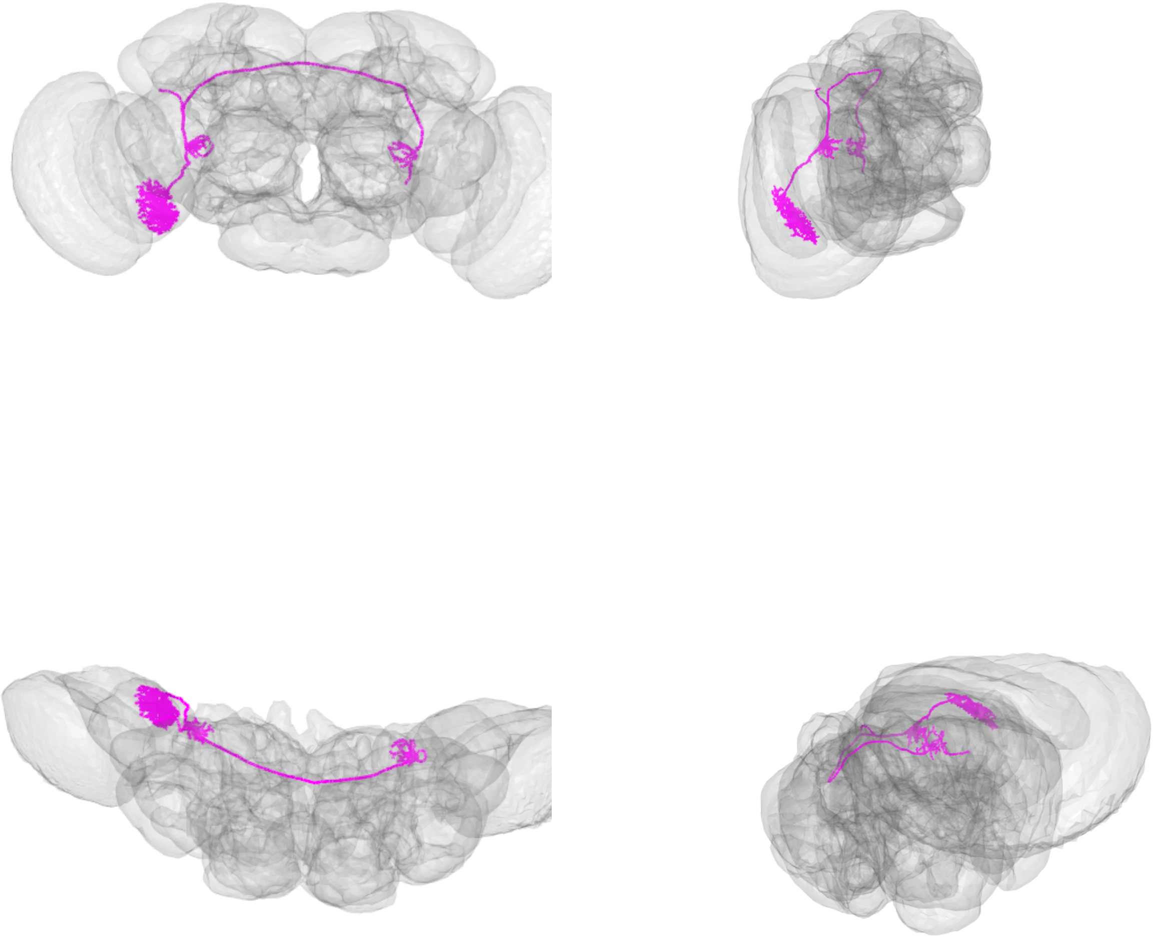

LPT46_R_vCal1_skid=7510076

LPT47_R_vCal2_skid=3529071

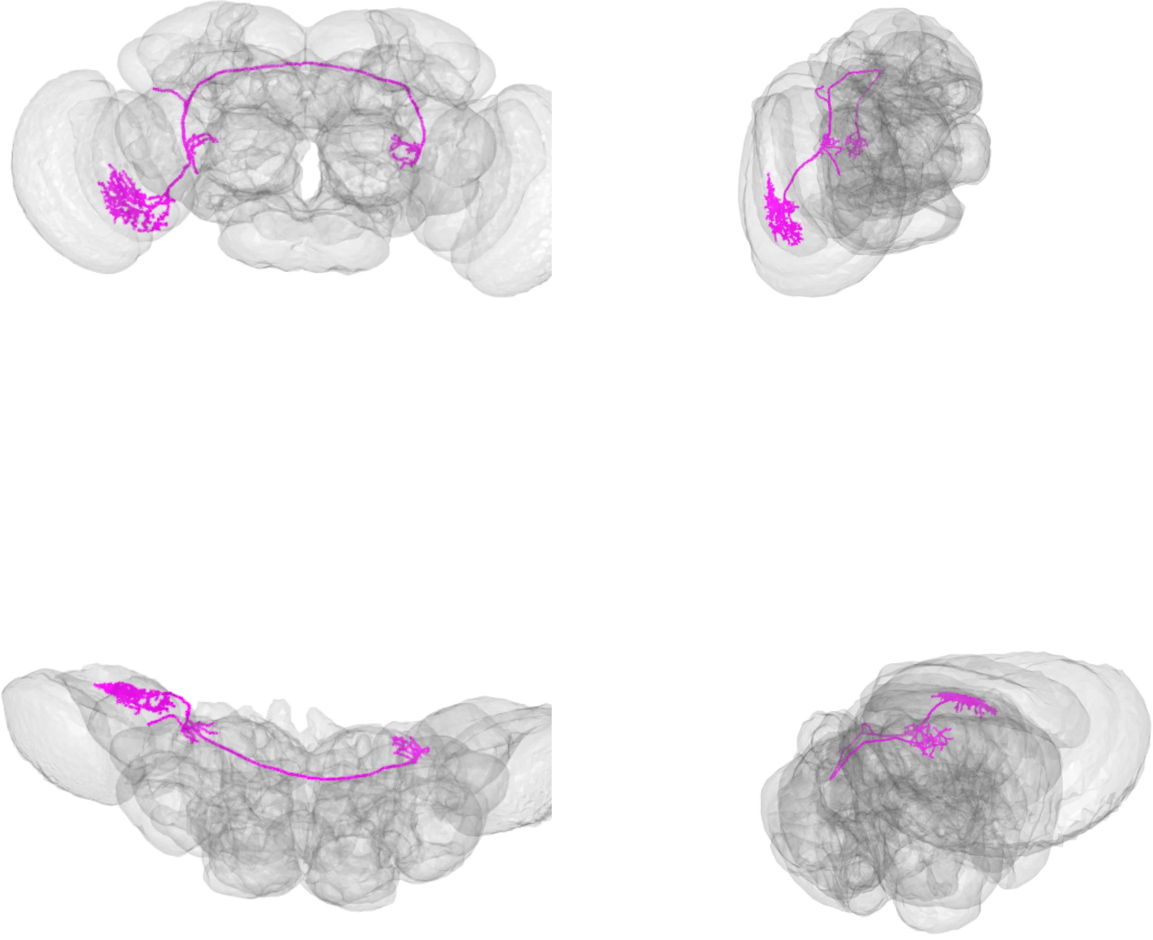

LPT47_R_vCal2_skid=3529071

LPT48_R_vCal3_skid=1056097

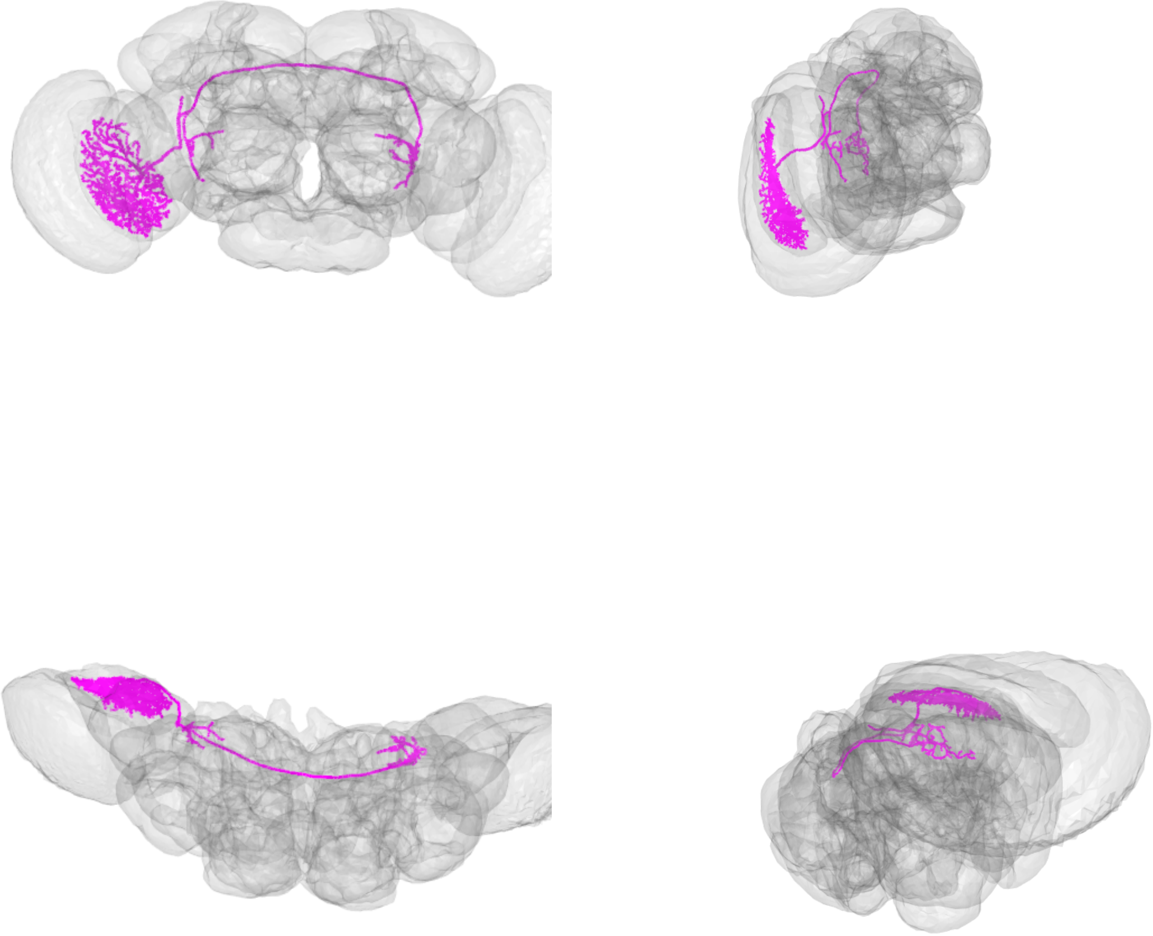

LPT48_R_vCal3_skid=1056097

LPT49_R_skid=1110693

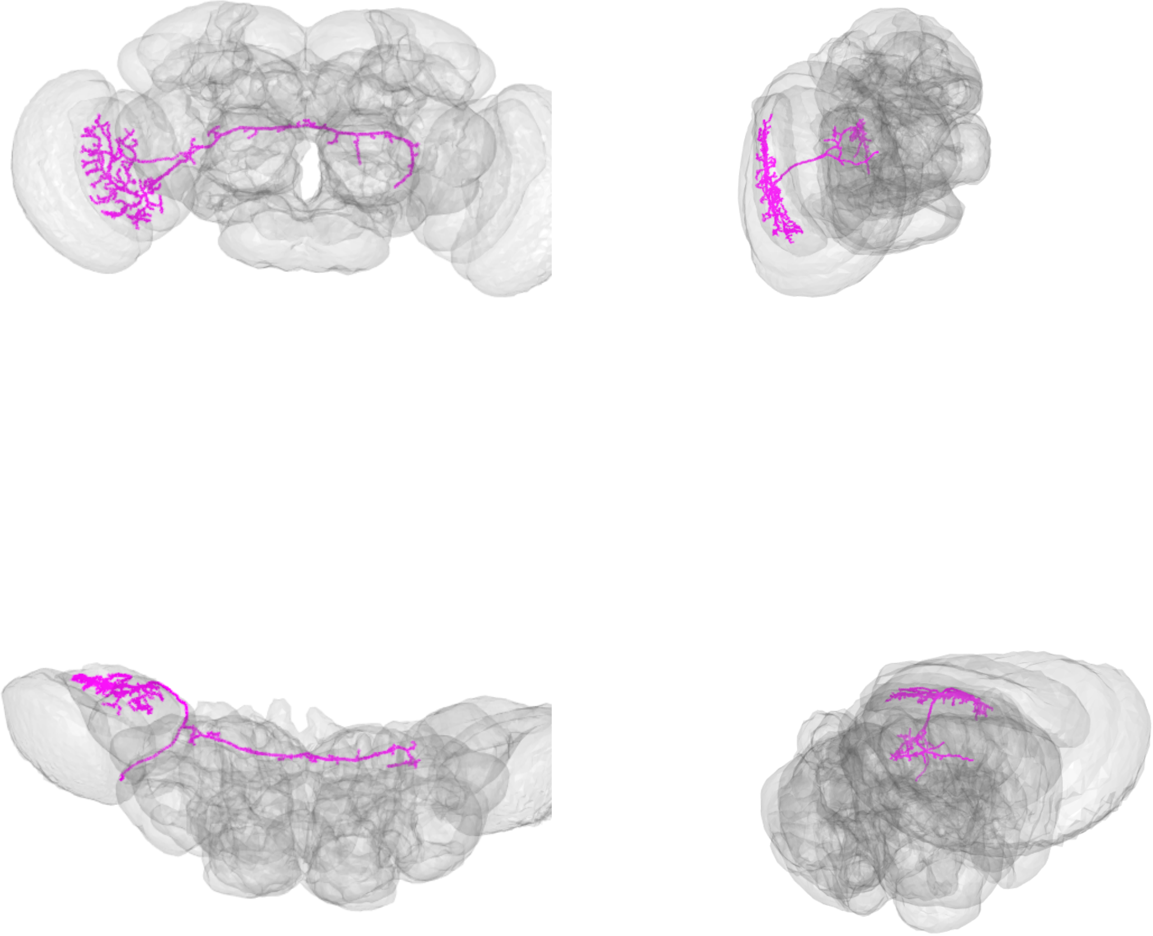

LPT49_R_skid=1110693

LPT50_R_skid=905761

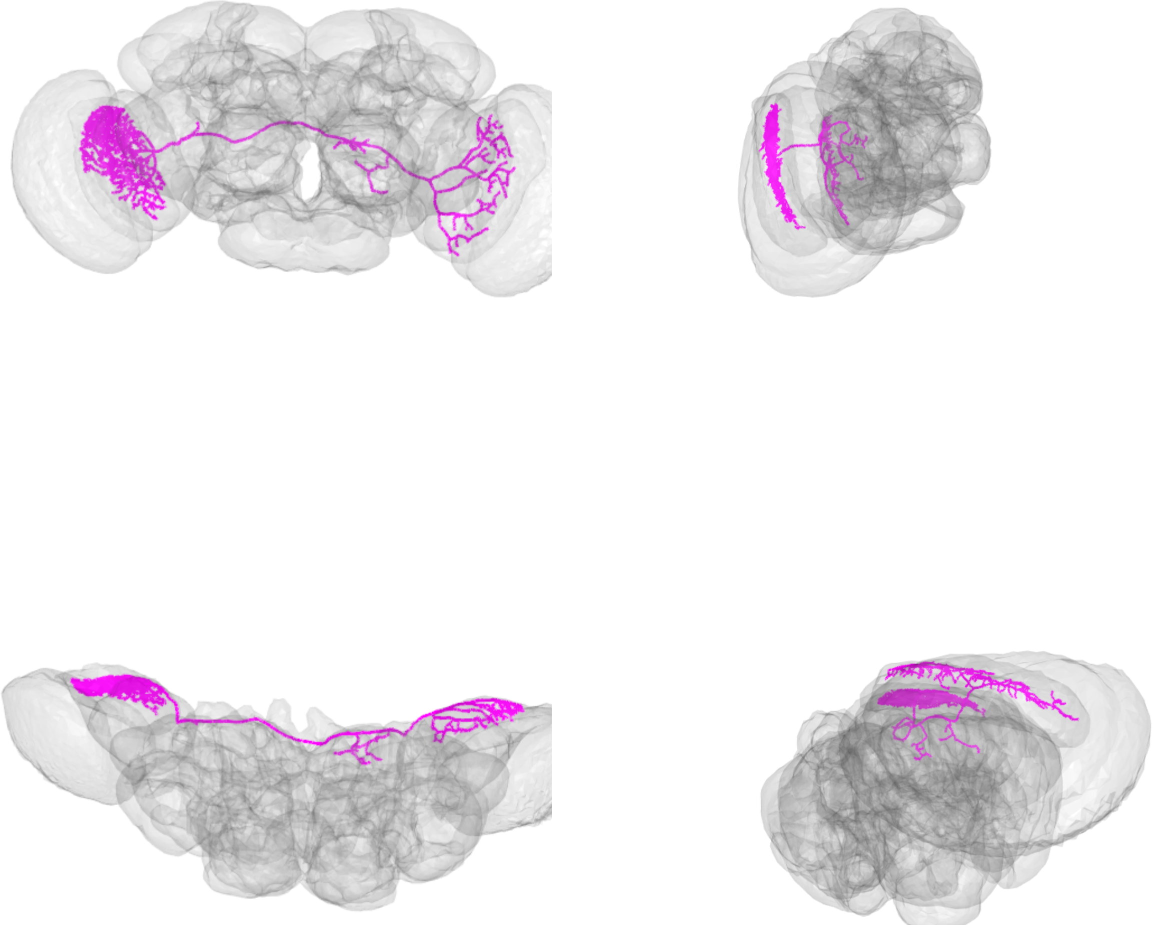

LPT50_R_skid=905761

LPT51_R_skid=4224711

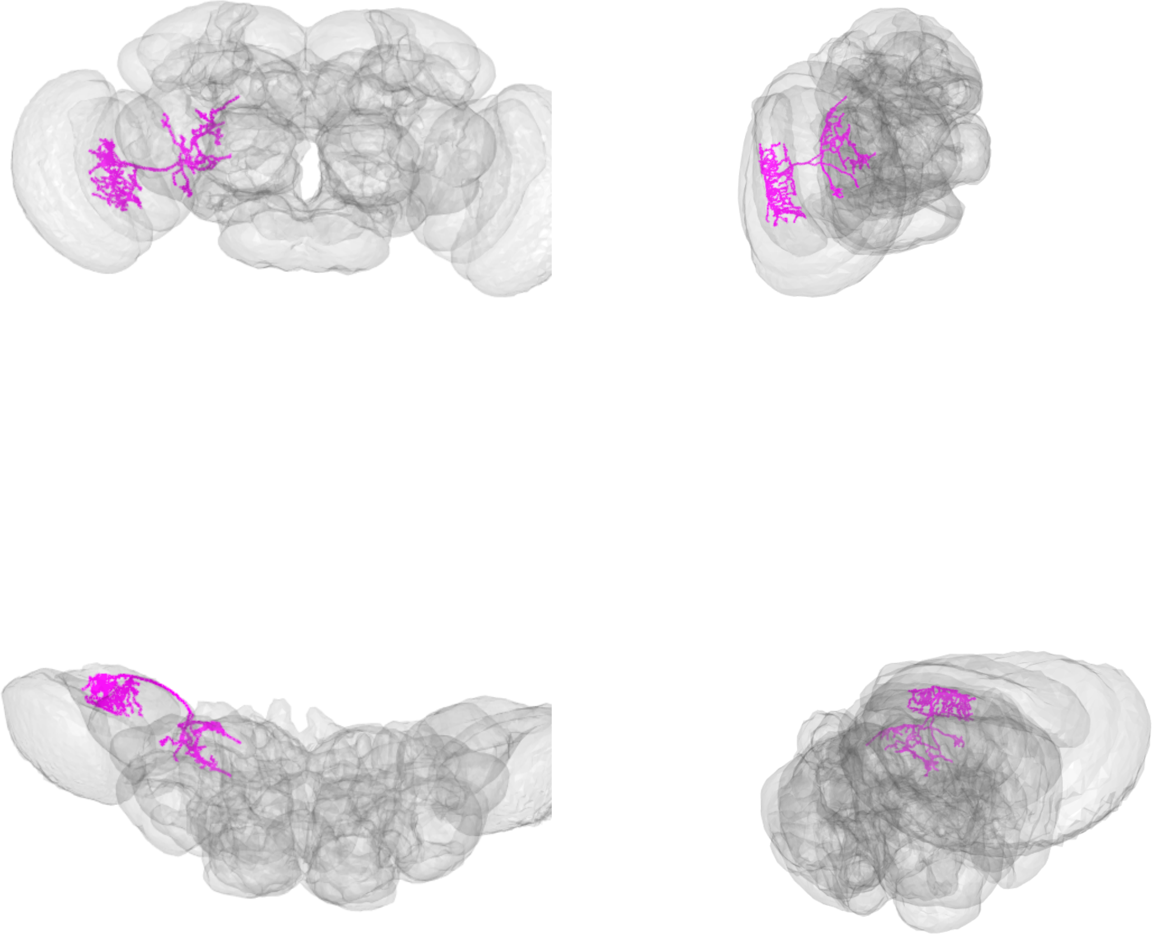

LPT51_R_skid=4224711

LPT52_R_skid=1107296

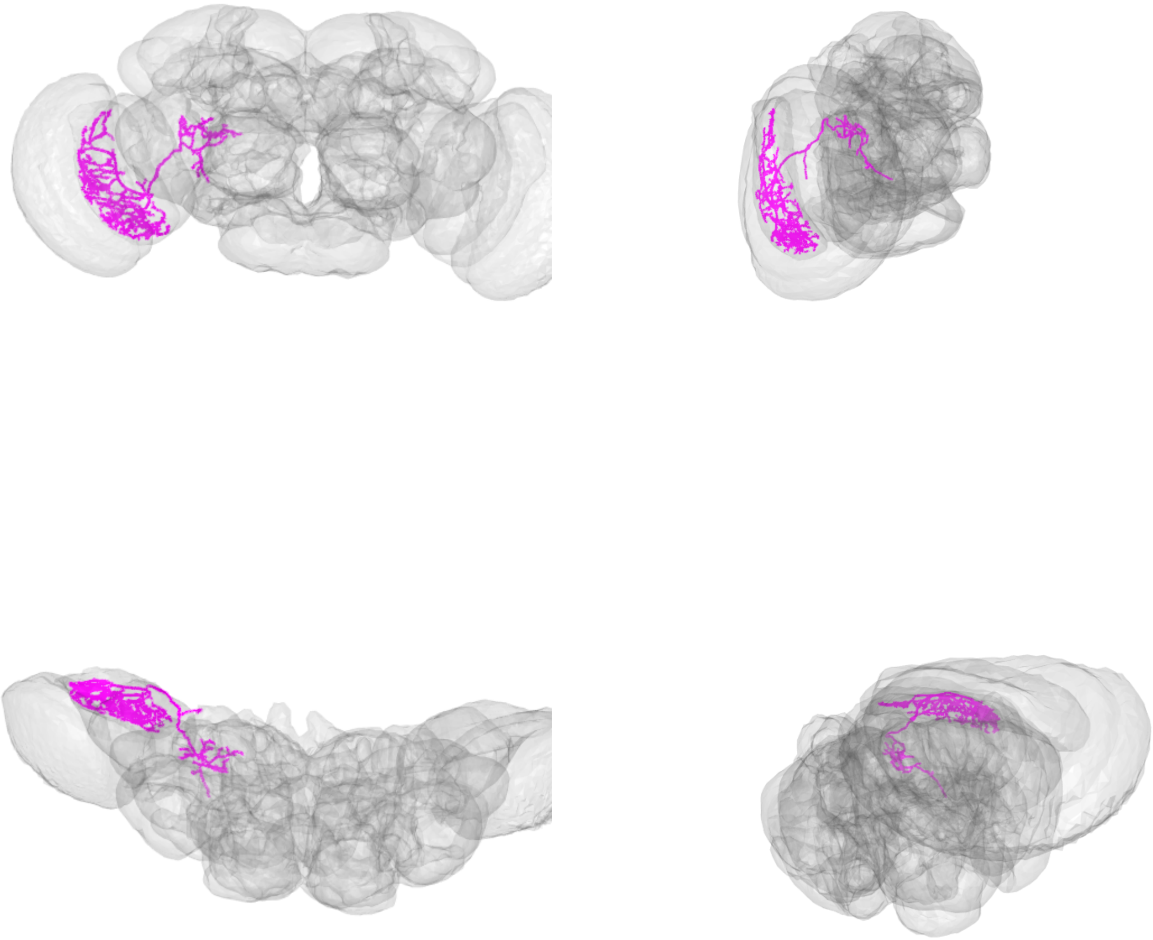

LPT52_R_skid=1107296

LPT53_R_skid=1111992

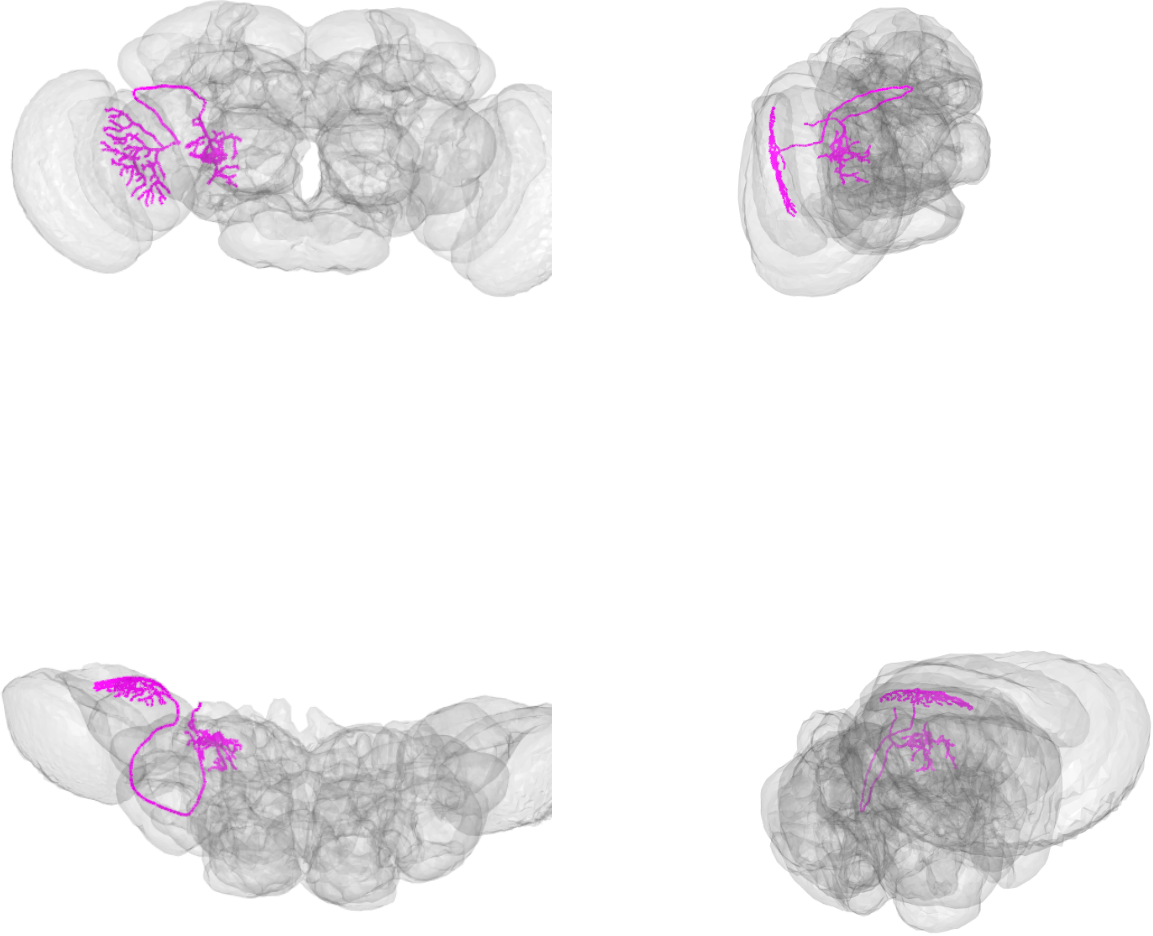

LPT53_R_skid=1111992

LPT54_R_skid=4235388

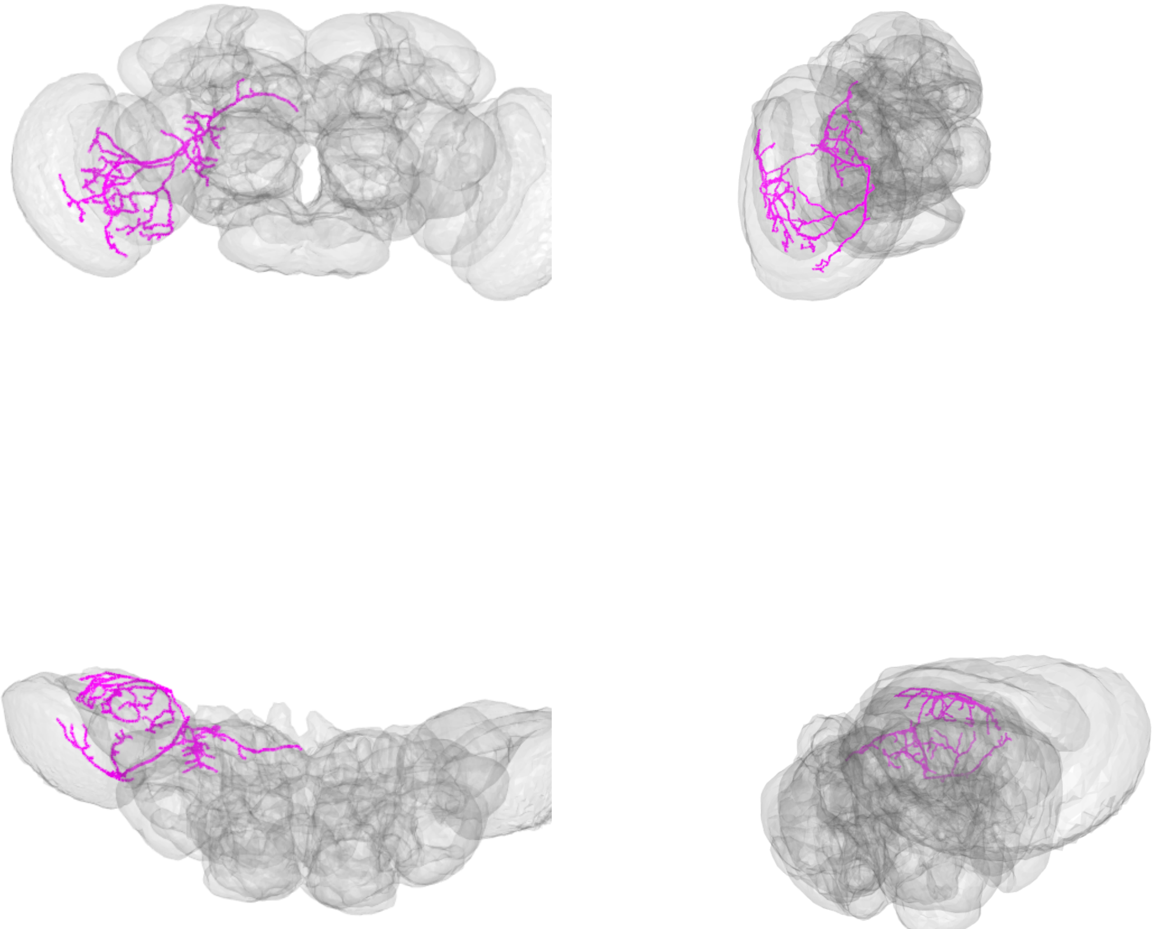

LPT54_R_skid=4235388

LPT55_R_MeLp2_skid=1061368

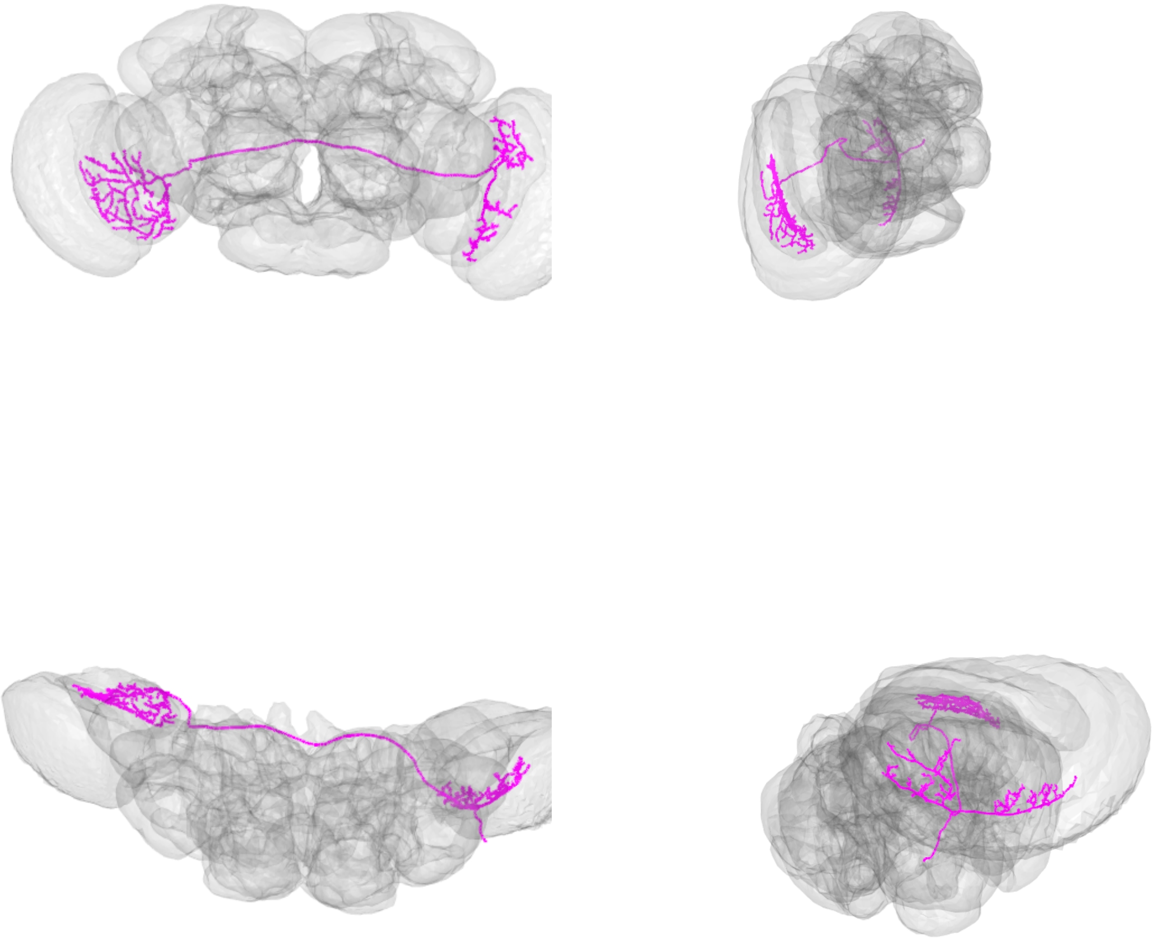

LPT55_R_MeLp2_skid=1061368

LPT56_R_MeLp1_skid=3509520

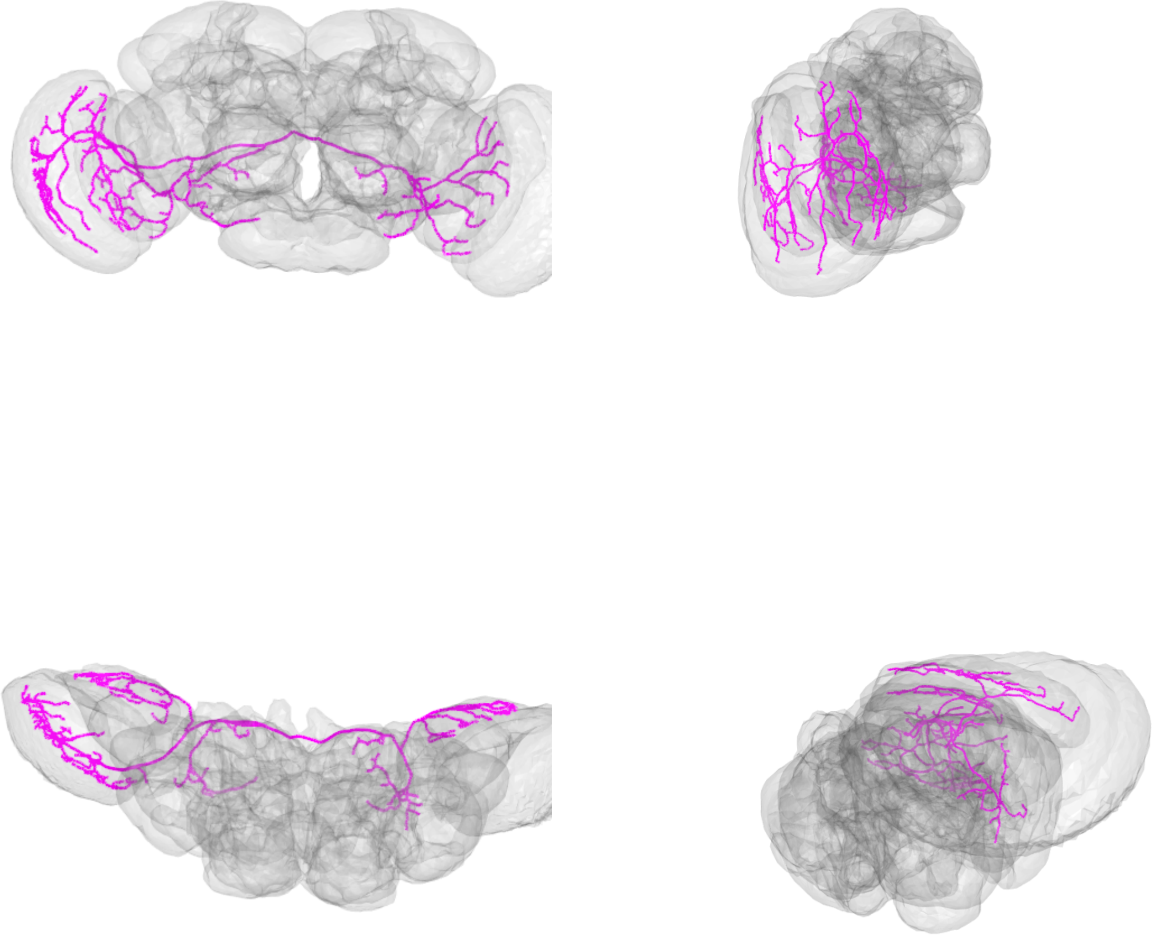

LPT56_R_MeLp1_skid=3509520

LPT57_R_skid=886797

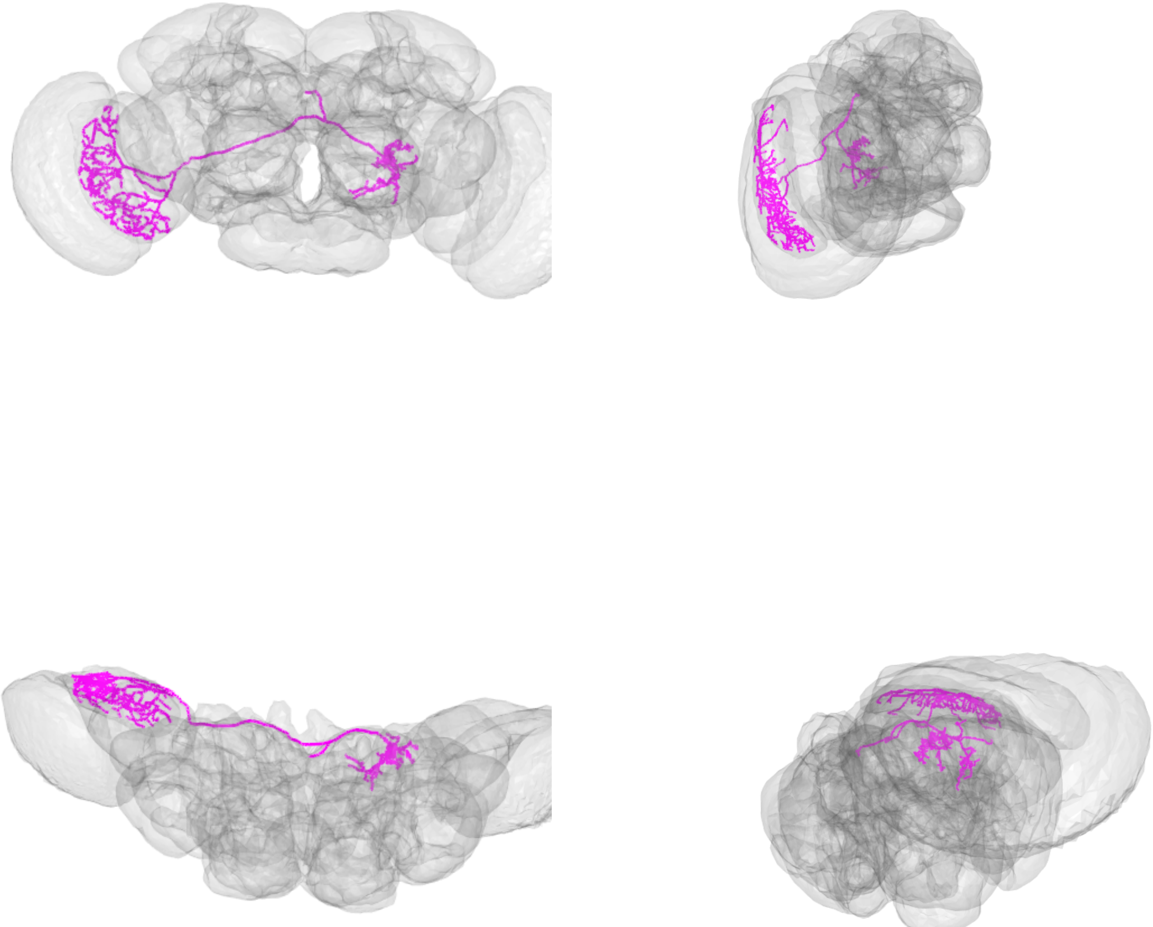

LPT57_R_skid=886797

LPT58_R_V1_skid=1059420

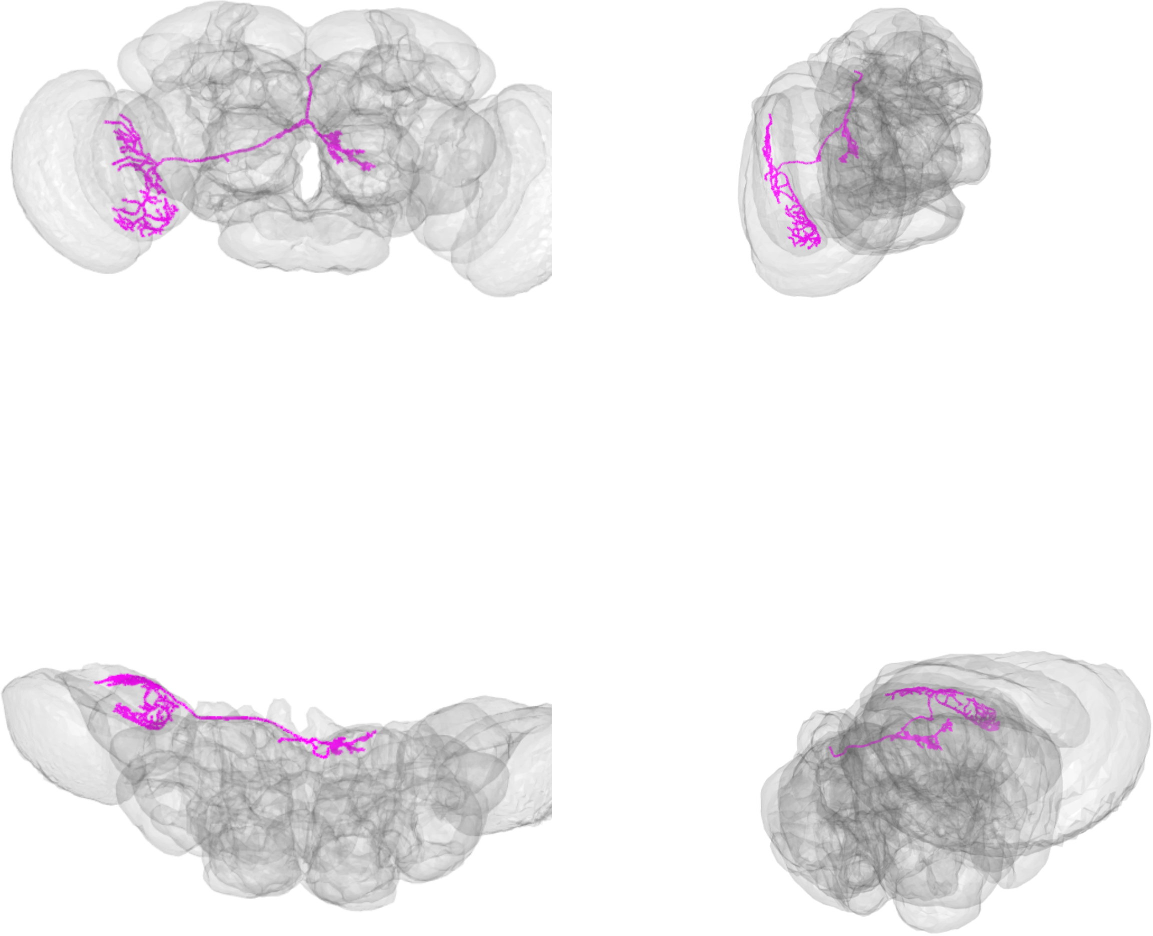

LPT58_R_V1_skid=1059420
