## Supplemental File 3 for "A comprehensive neuroanatomical survey of the *Drosophila* Lobula Plate Tangential Neurons with predictions for their optic flow sensitivity"

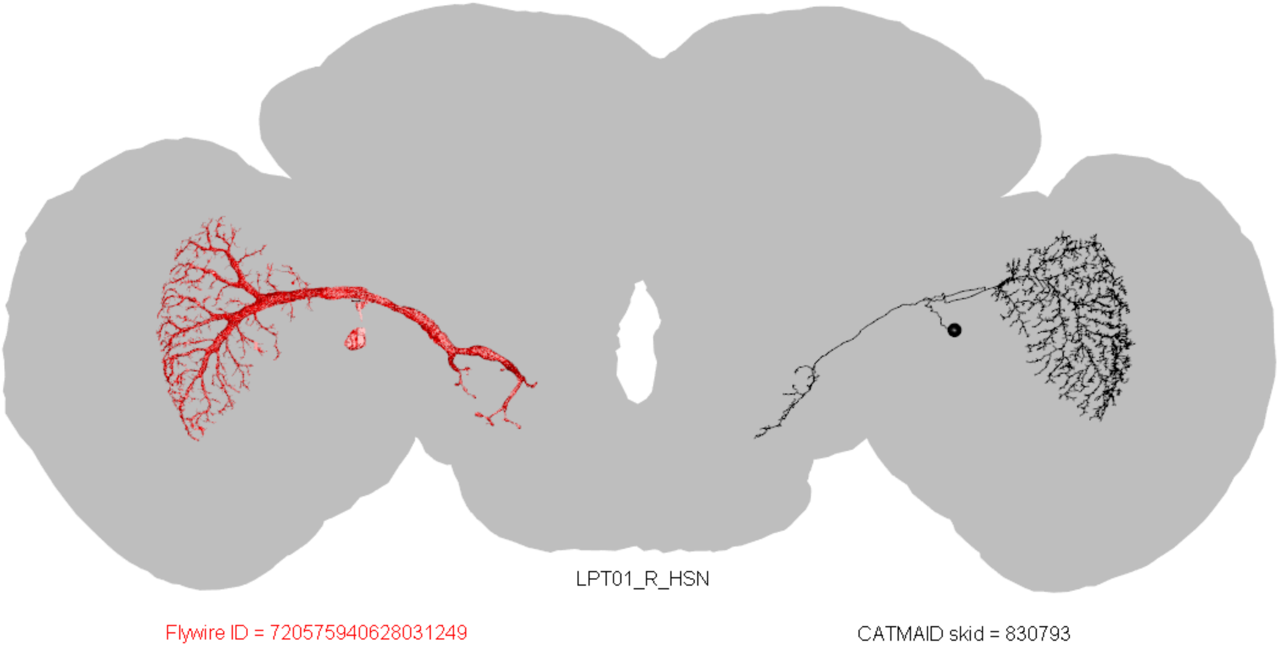
LPT01_R_HSN
Flywire ID = 720575940628031249
CATMAID skid = 830793

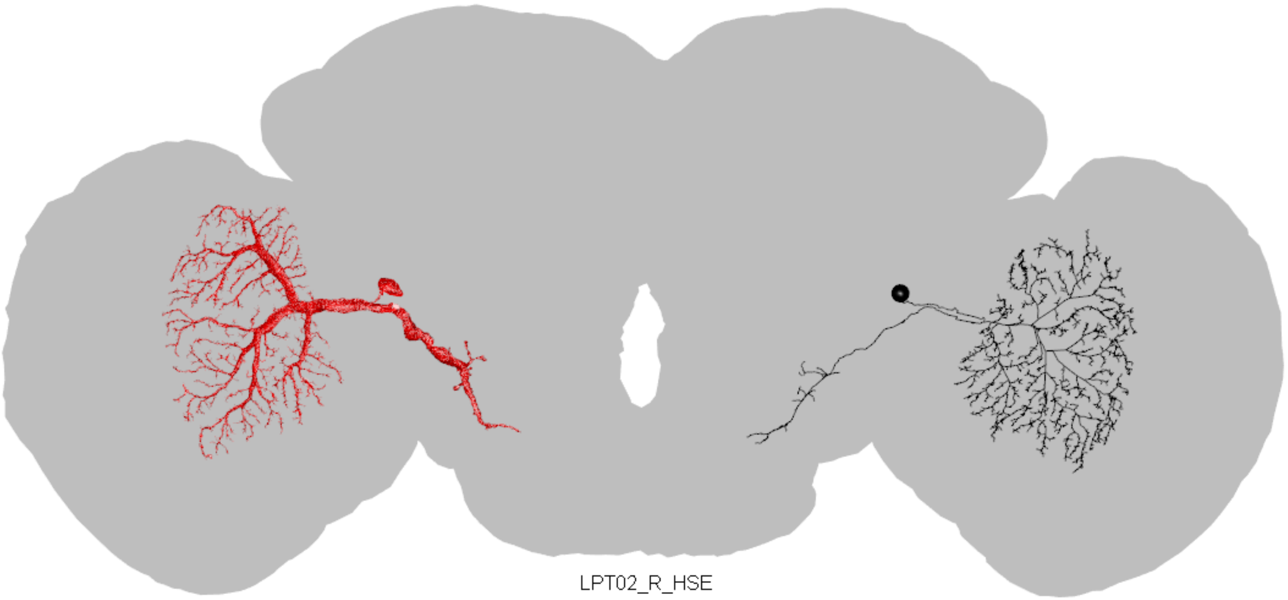

LPT02_R_HSE

Flywire ID = 720575940642723981

CATMAID skid = 827034

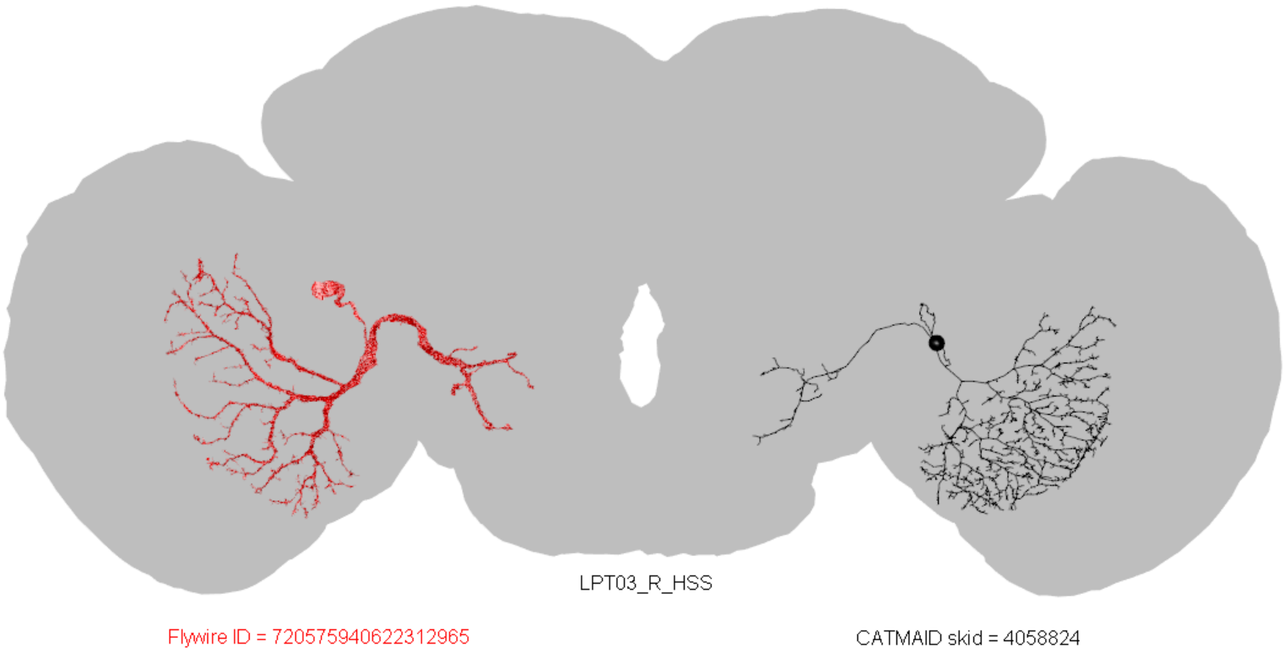
LPT03_R_HSS
Flywire ID = 720575940622312965
CATMAID skid = 4058824

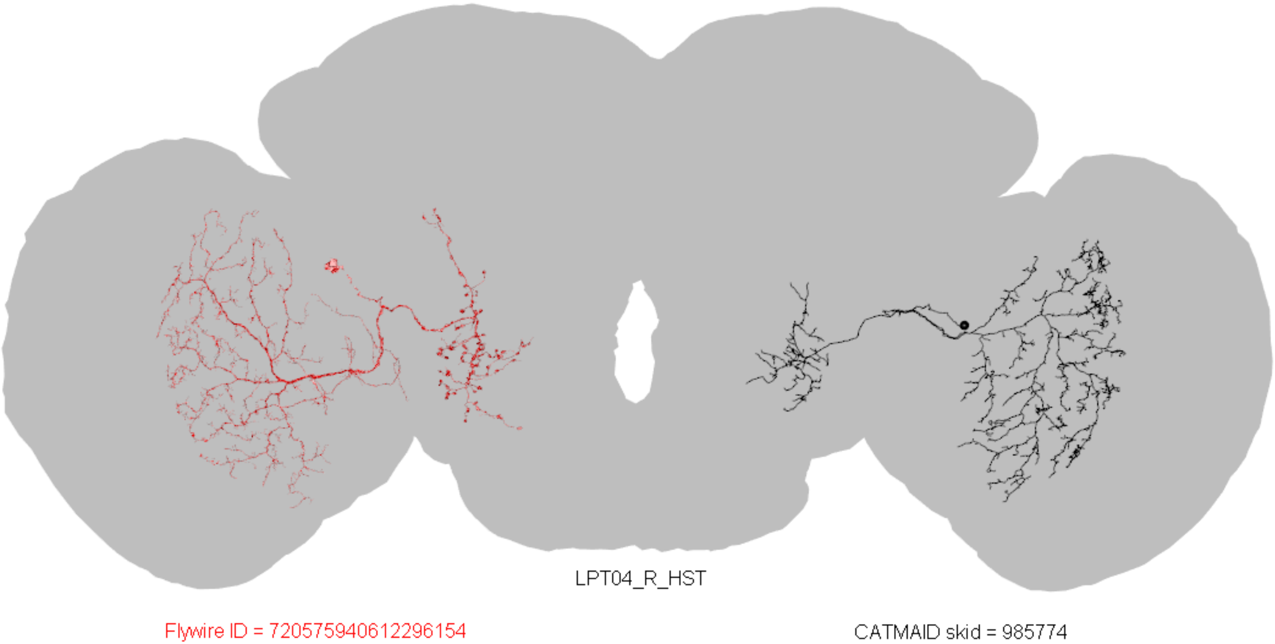
LPT04_R_HST
Flywire ID = 720575940612296154
CATMAID skid = 985774

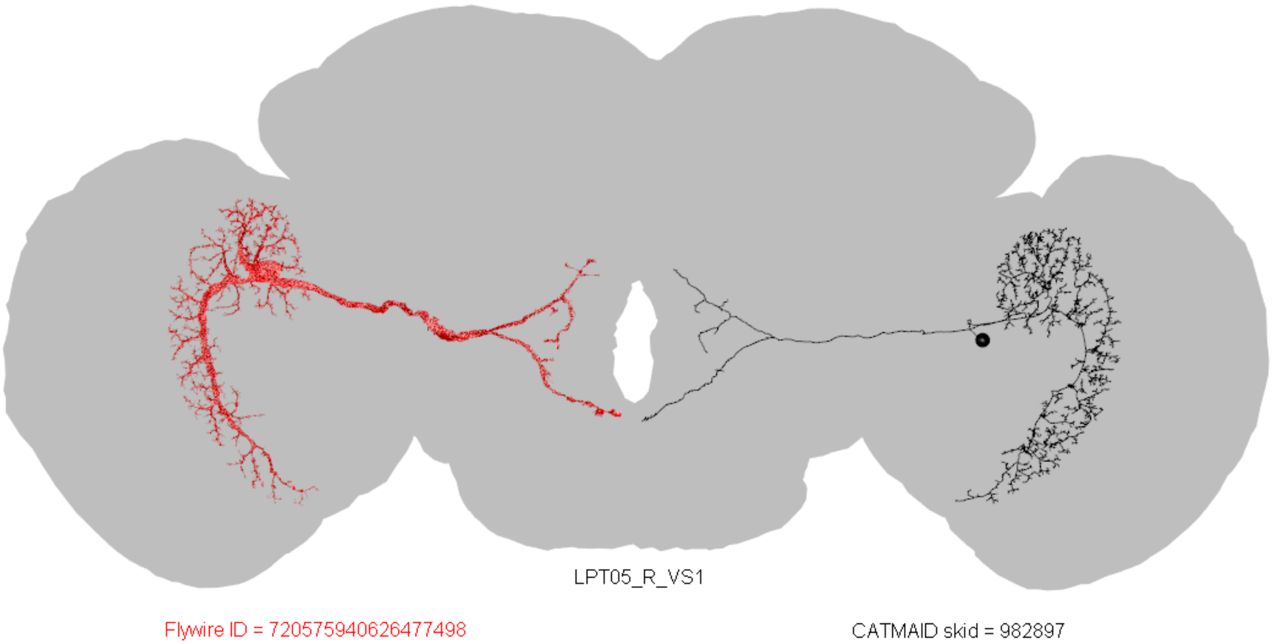
LPT05_R_VS1
Flywire ID = 720575940626477498
CATMAID skid = 982897

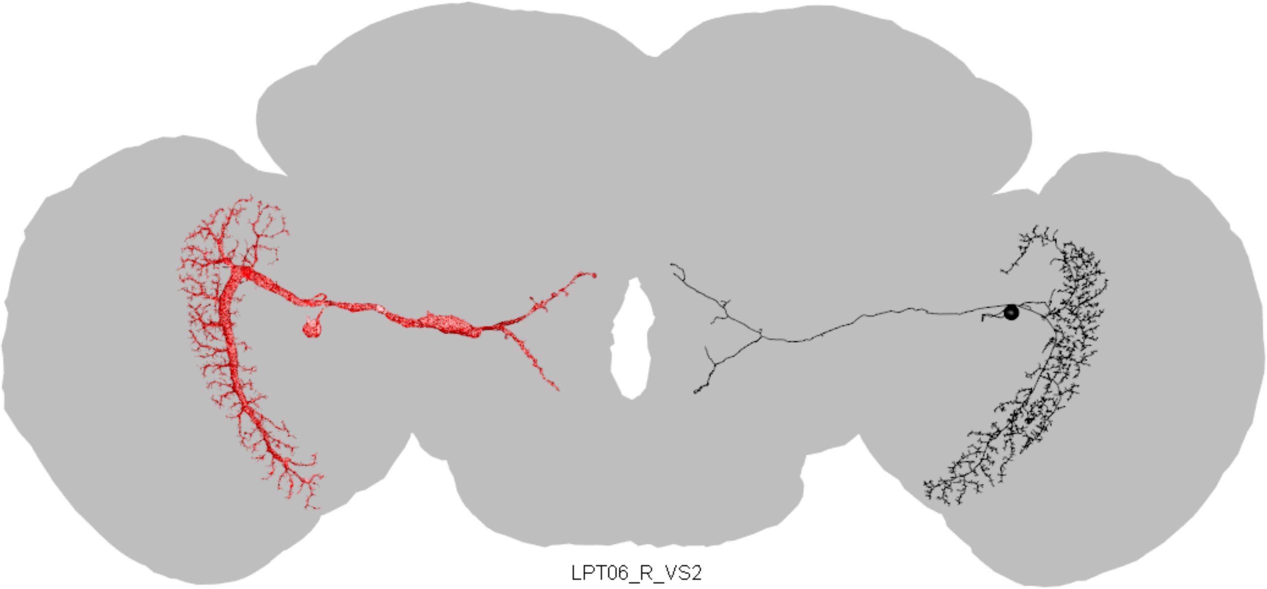

LPT06_R_VS2

Flywire ID = 720575940615269794

CATMAID skid = 793032

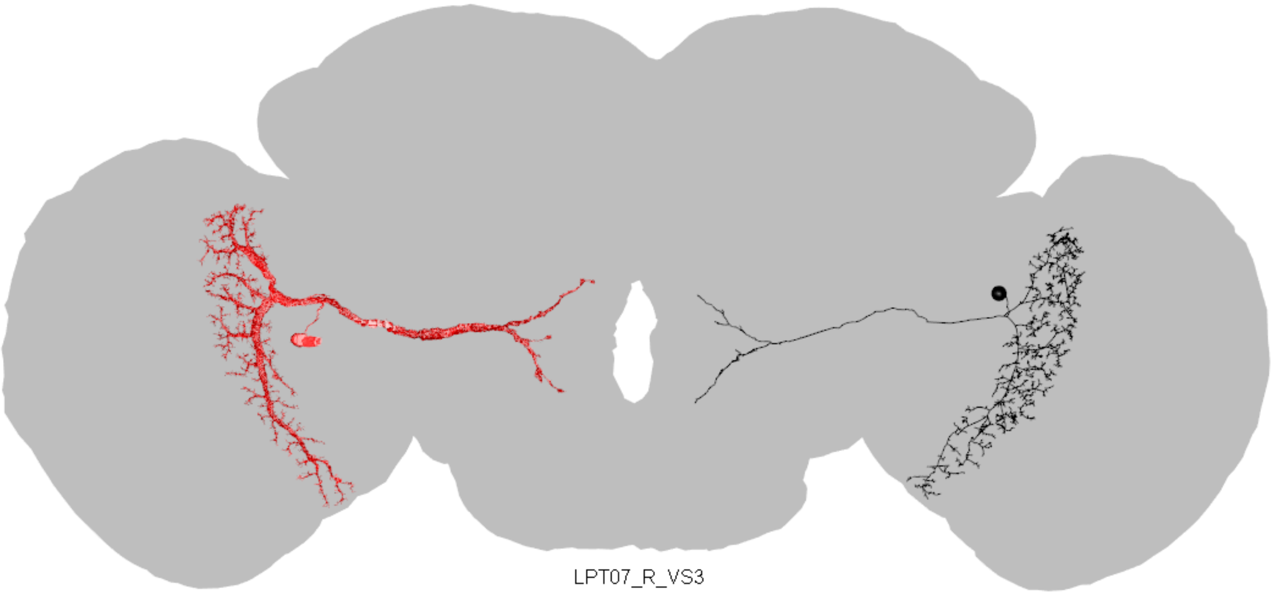

LPT07_R_VS3

Flywire ID = 720575940622831740

CATMAID skid = 815776

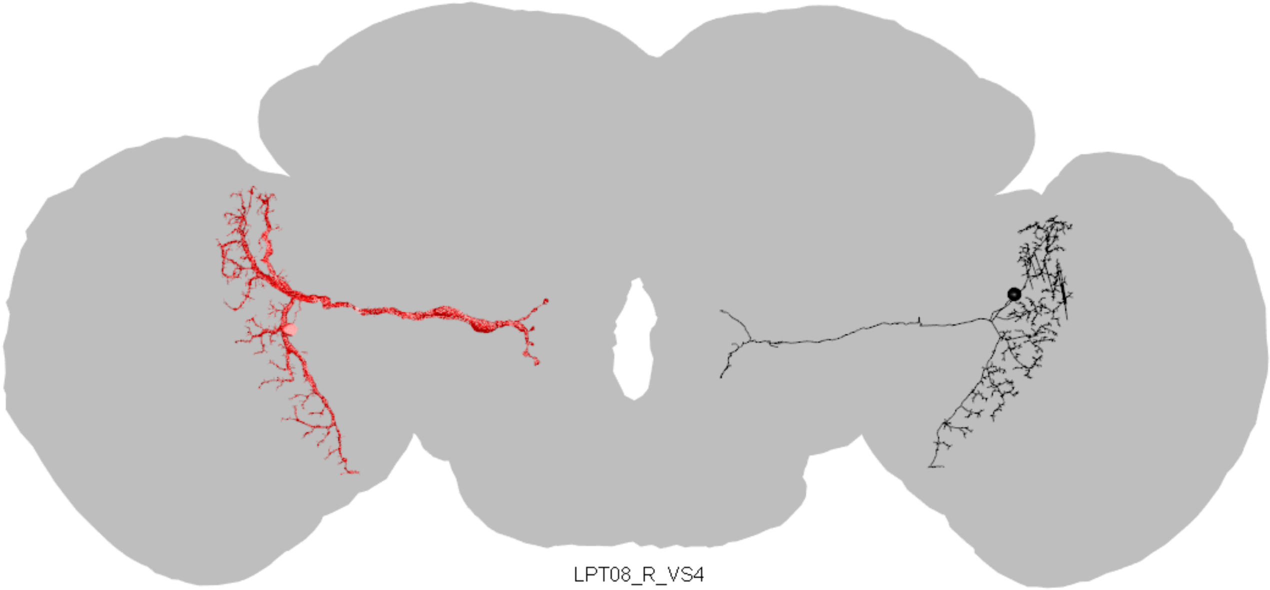

LPT08_R_VS4

Flywire ID = 720575940633017939

CATMAID skid = 17686499

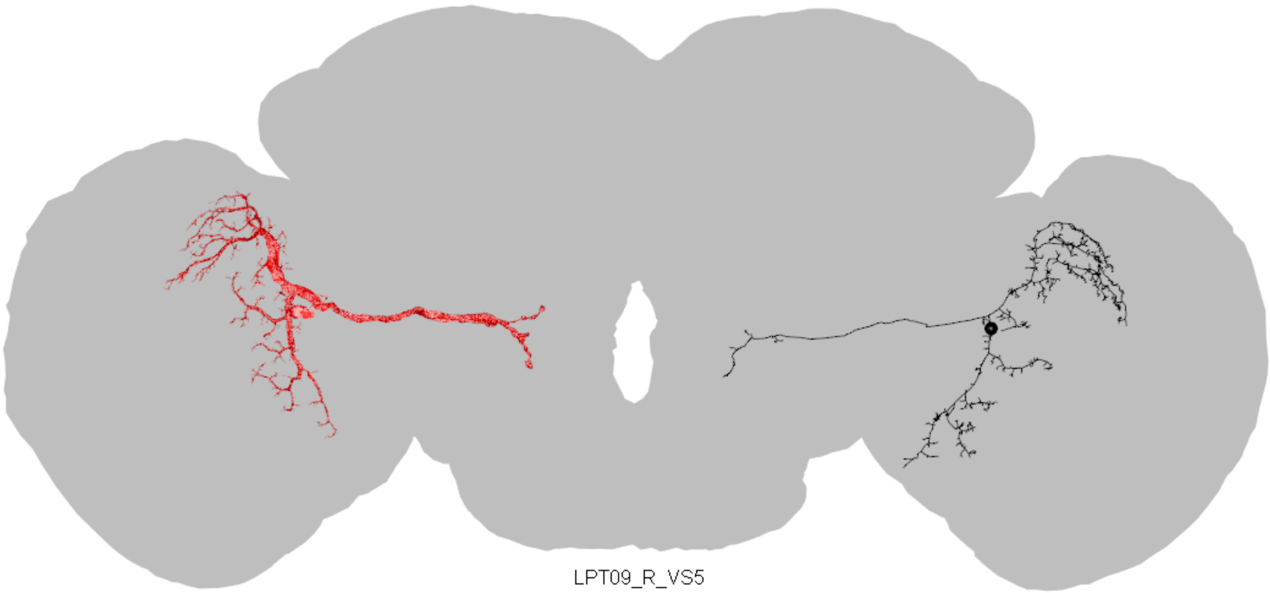

LPT09_R_VS5

Flywire ID = 720575940626457406

CATMAID skid = 807401

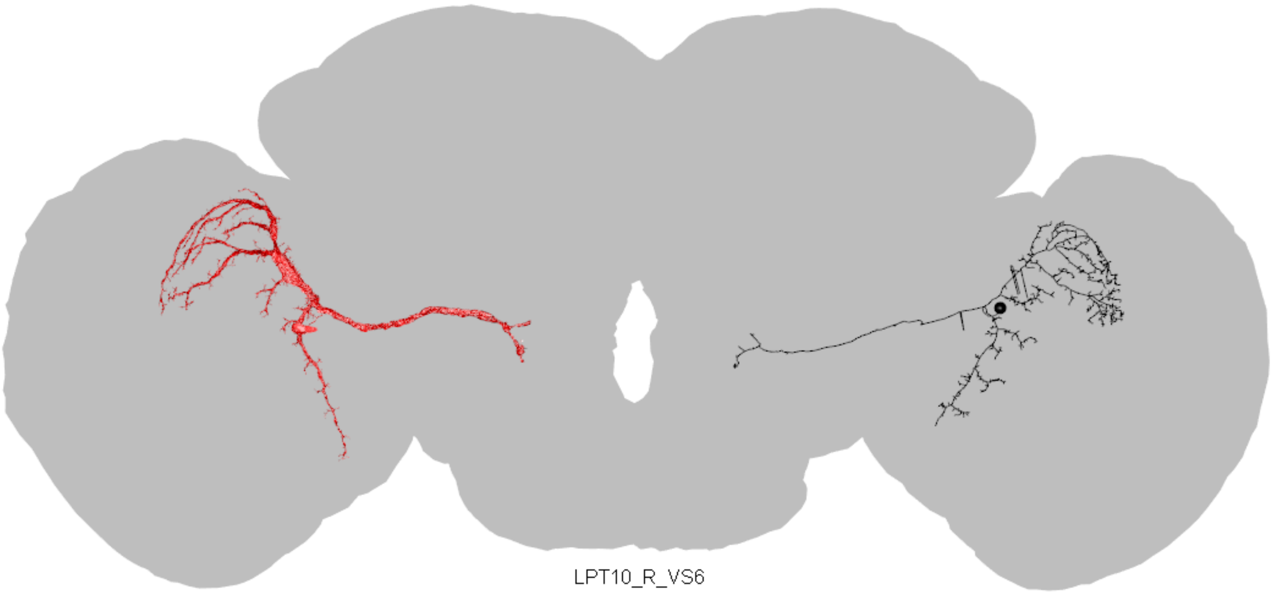

LPT10_R_VS6

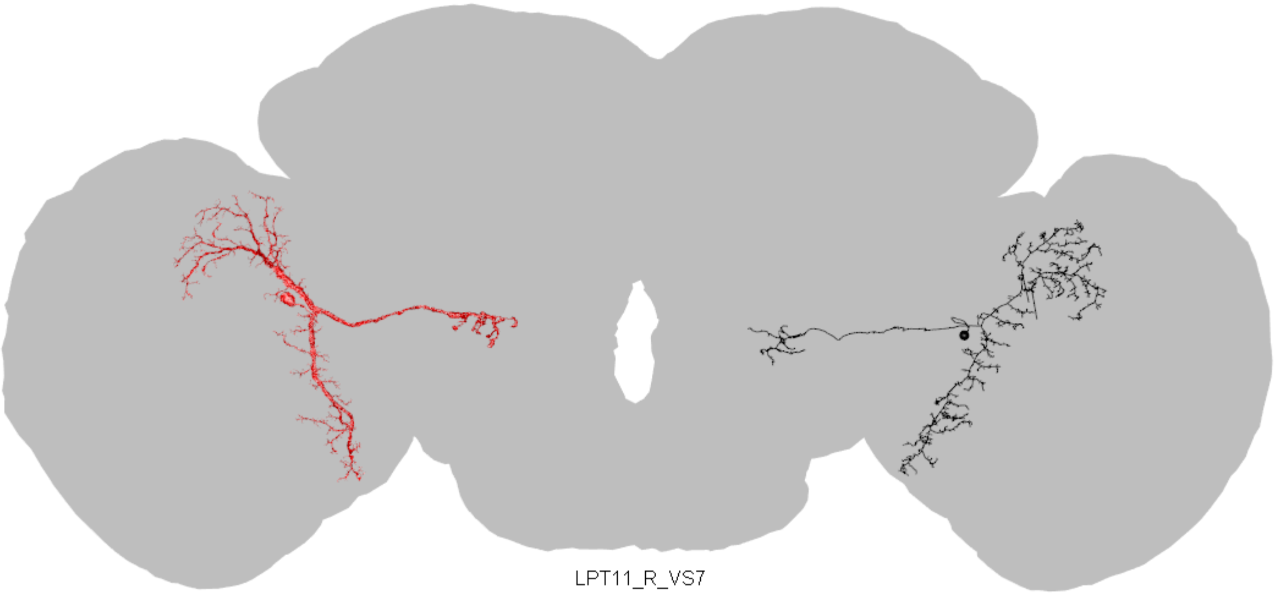

LPT11_R_VS7

Flywire ID = 720575940624931564

CATMAID skid = 851432

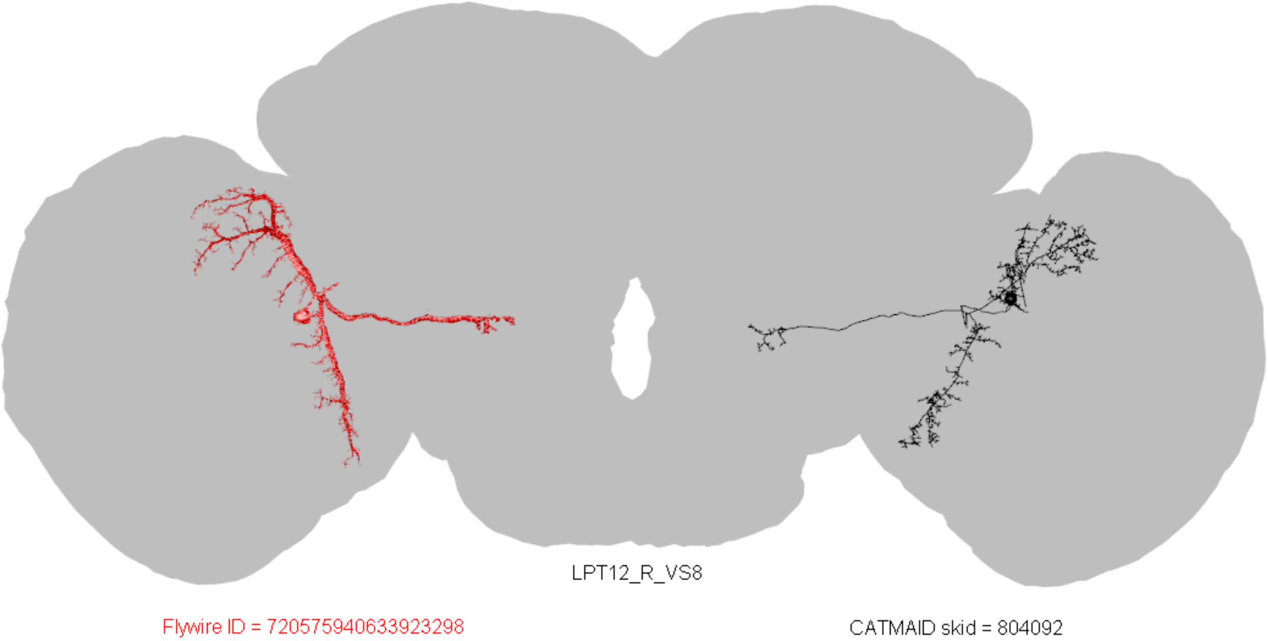
LPT12_R_VS8
Flywire ID = 720575940633923298
CATMAID skid = 804092

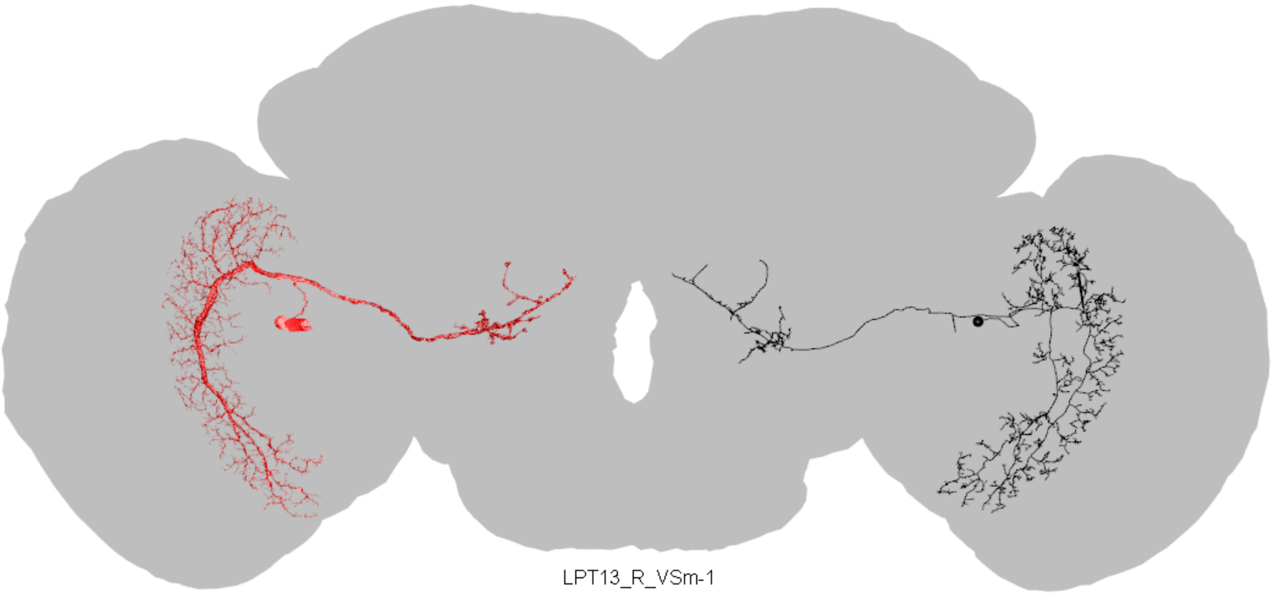

LPT13_R_VSm-1

Flywire ID = 720575940620463307

CATMAID skid = 852286

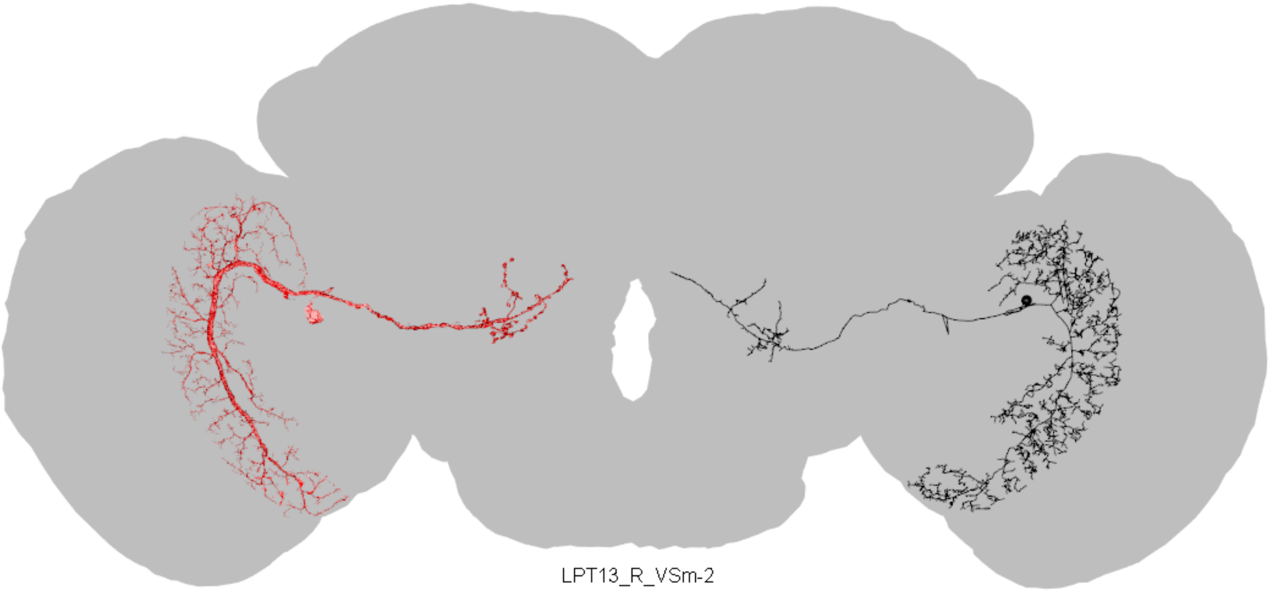

LPT13_R_VSm-2

Flywire ID = 720575940630825527

CATMAID skid = 815241

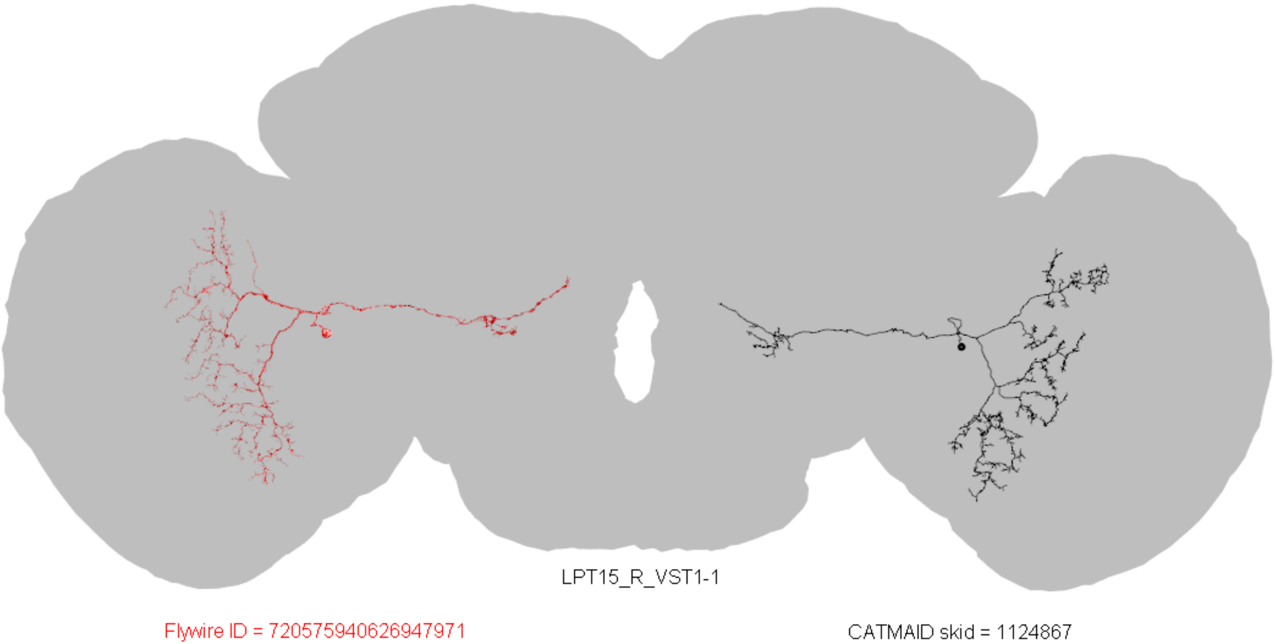
LPT15_R_VST1-1
Flywire ID = 720575940626947971
CATMAID skid = 1124867

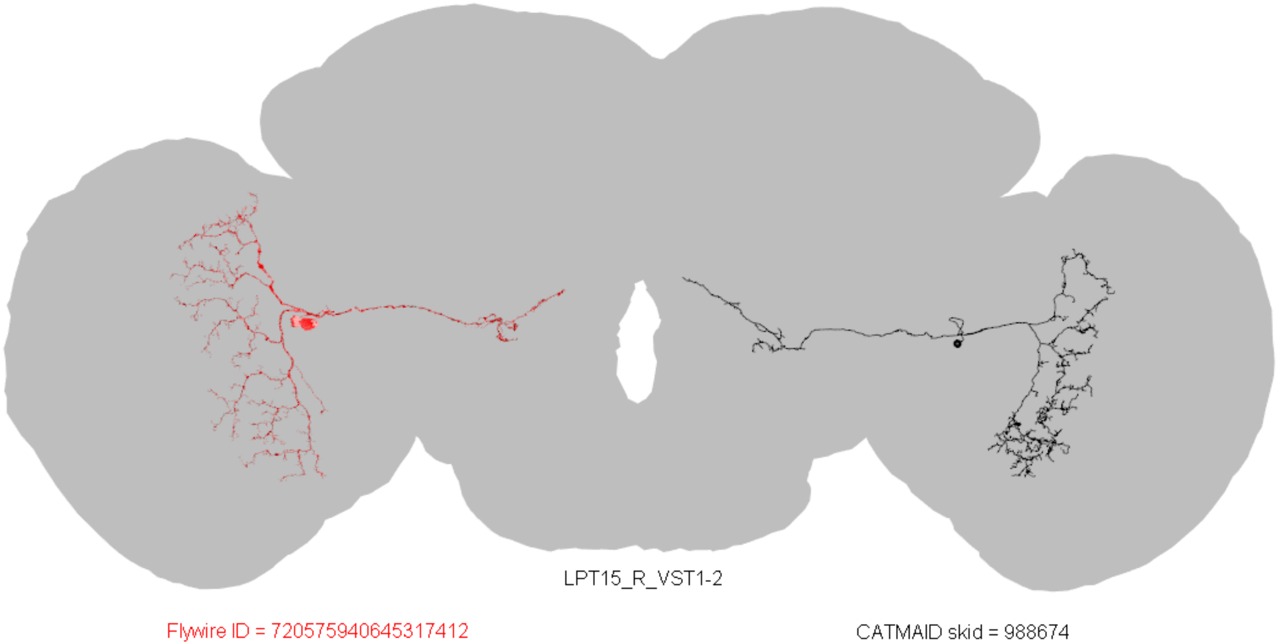
LPT15_R_VST1-2
Flywire ID = 720575940645317412
CATMAID skid = 988674

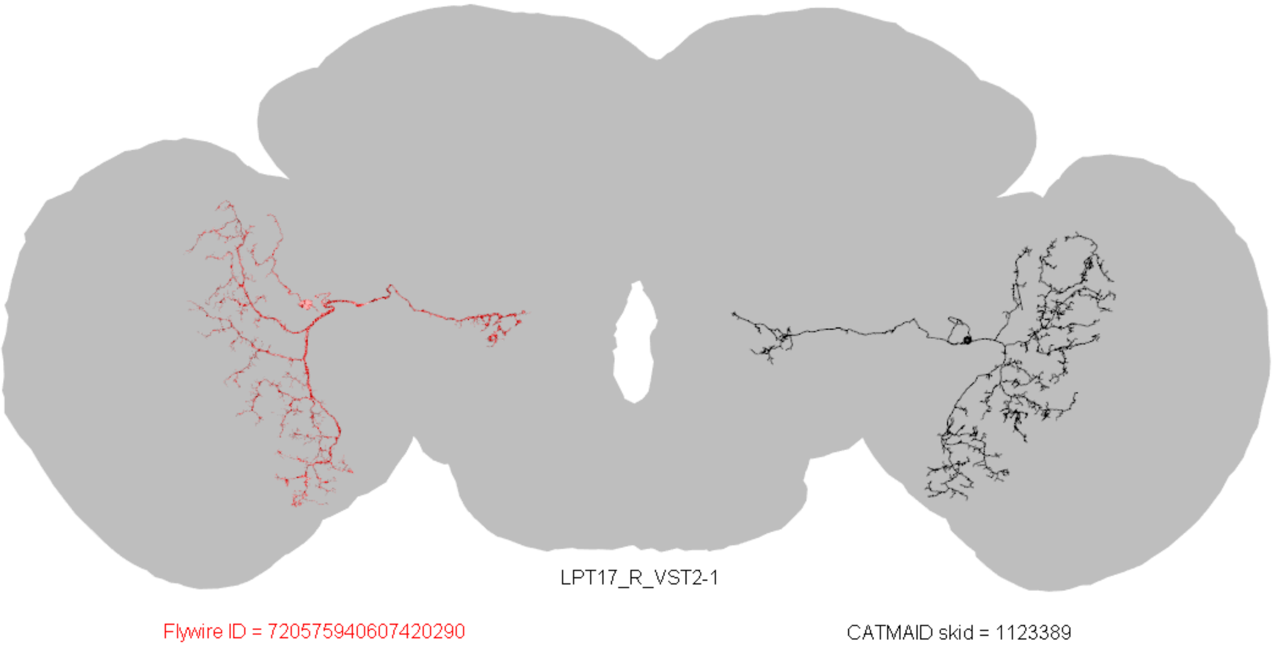
LPT17_R_VST2-1
Flywire ID = 720575940607420290
CATMAID skid = 1123389

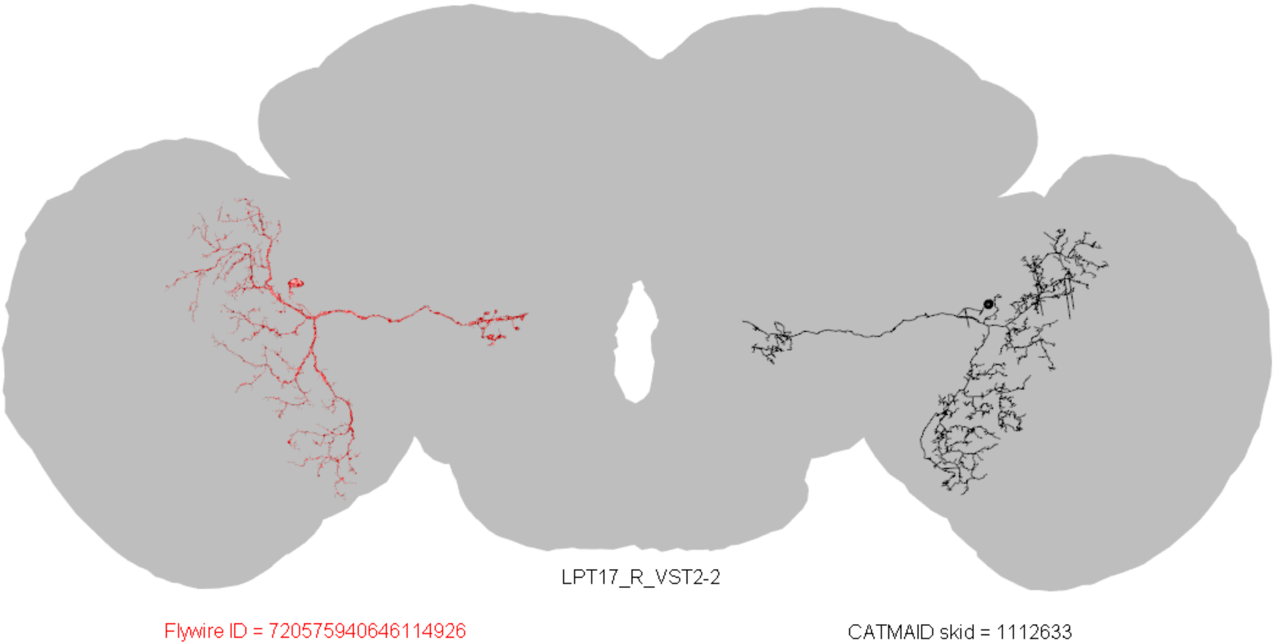
LPT17_R_VST2-2
Flywire ID = 720575940646114926
CATMAID skid = 1112633

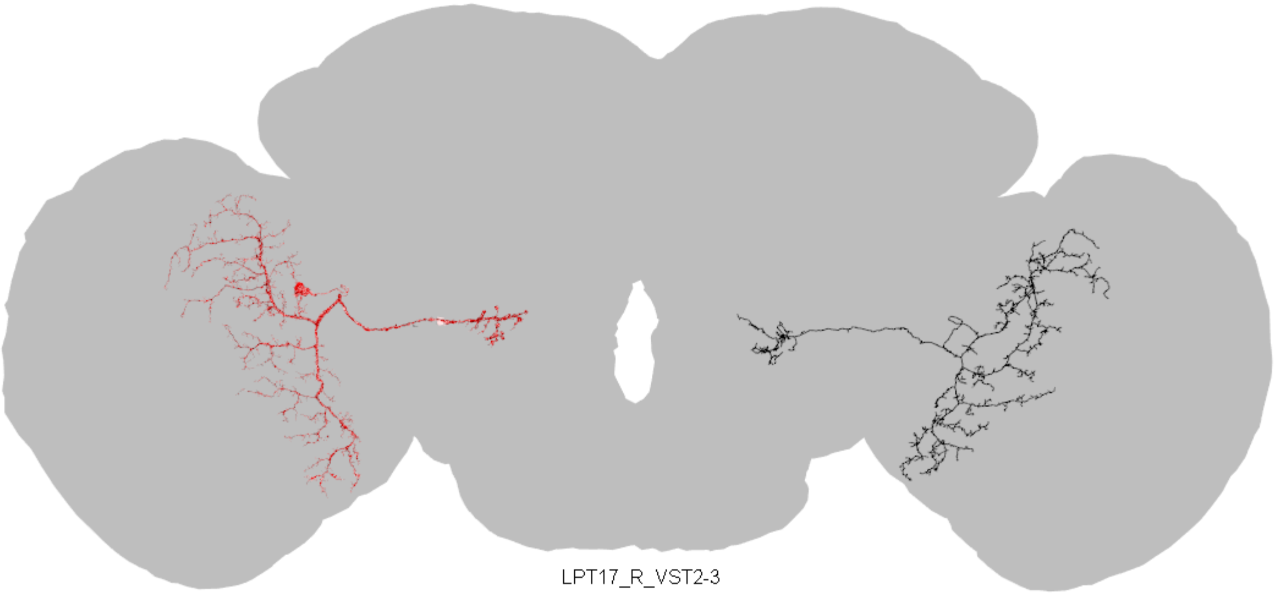

LPT17_R_VST2-3

Flywire ID = 720575940635615339

CATMAID skid = 2852912

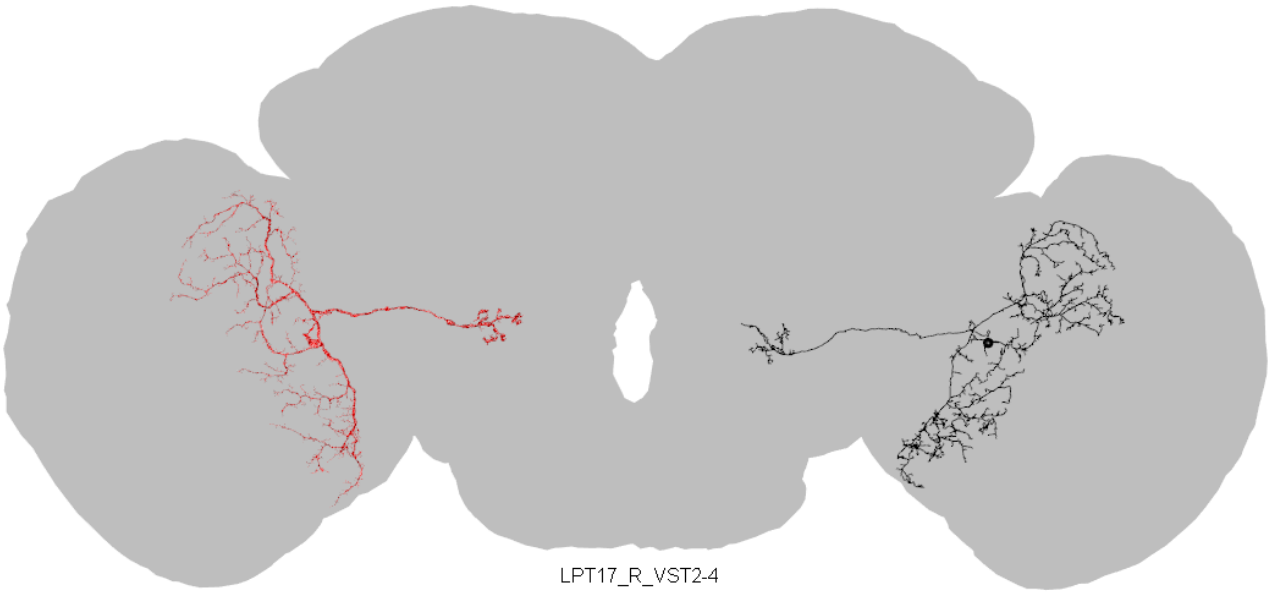

LPT17_R_VST2-4

Flywire ID = 720575940628452520

CATMAID skid = 5031615

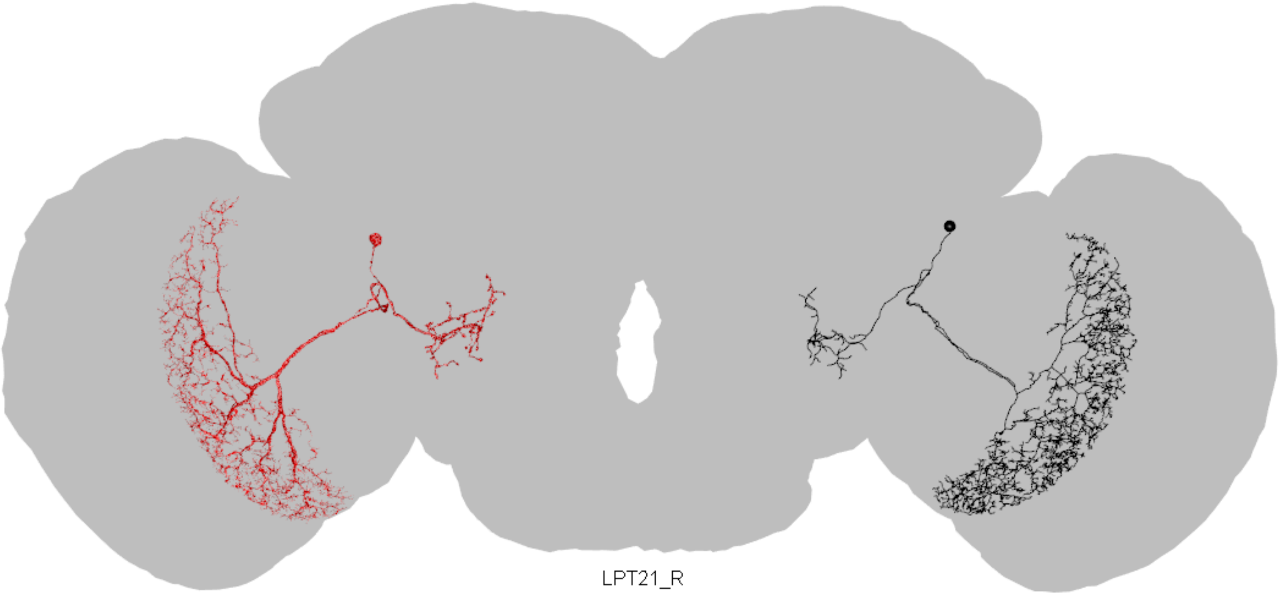

LPT21_R

Flywire ID = 720575940610781560

CATMAID skid = 1110765

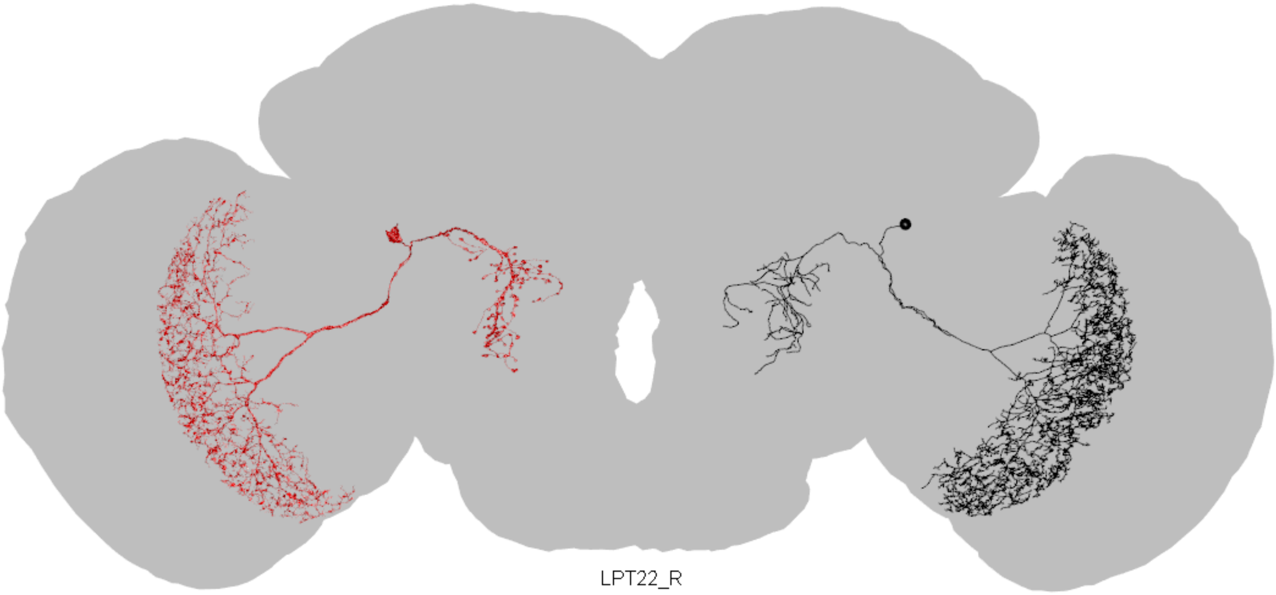


Flywire ID = 720575940610061763

CATMAID skid = 3510999

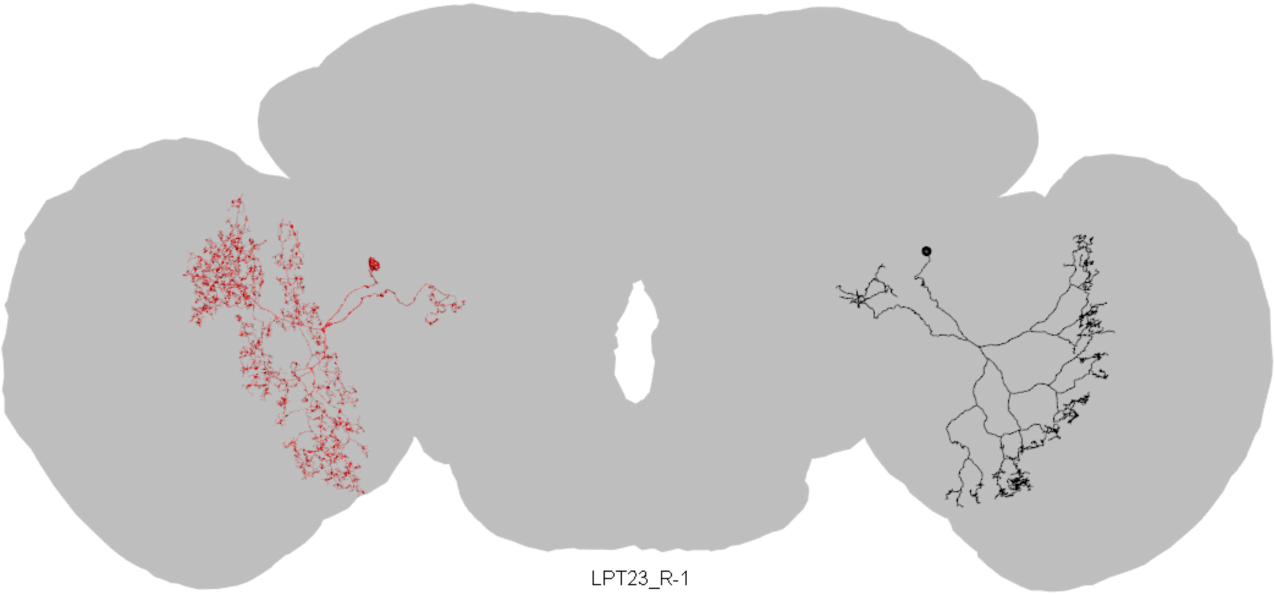

LPT23_R-1

Flywire ID = 720575940617304949

CATMAID skid = 4135042

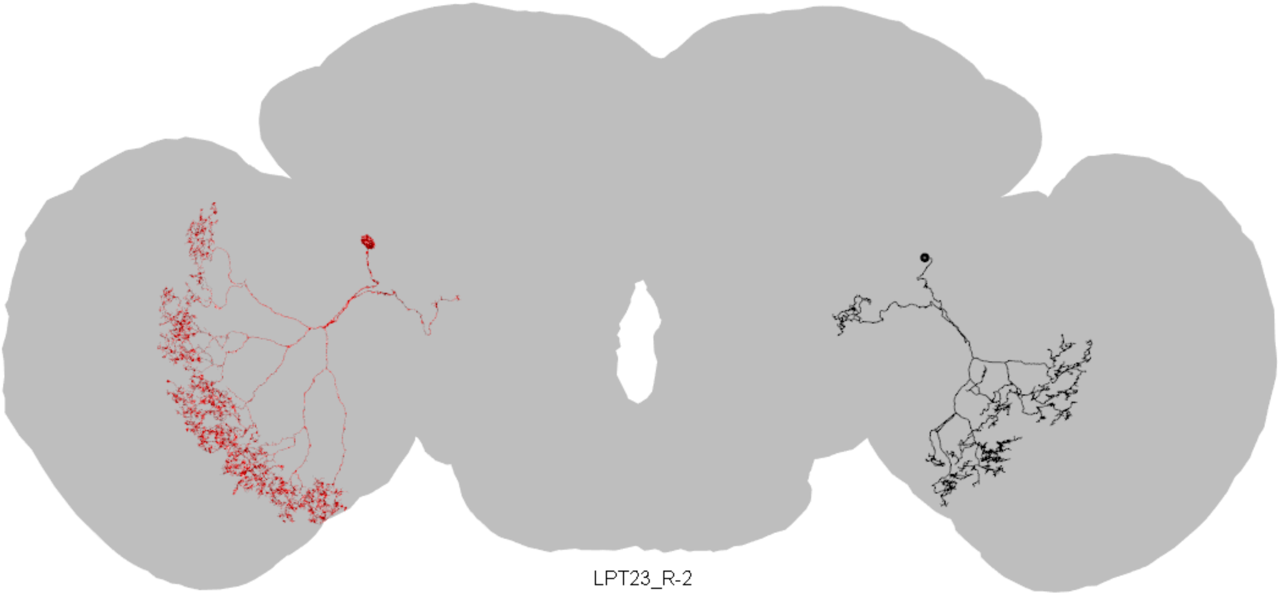

LPT23_R-2

Flywire ID = 720575940632545080

CATMAID skid = 4504548

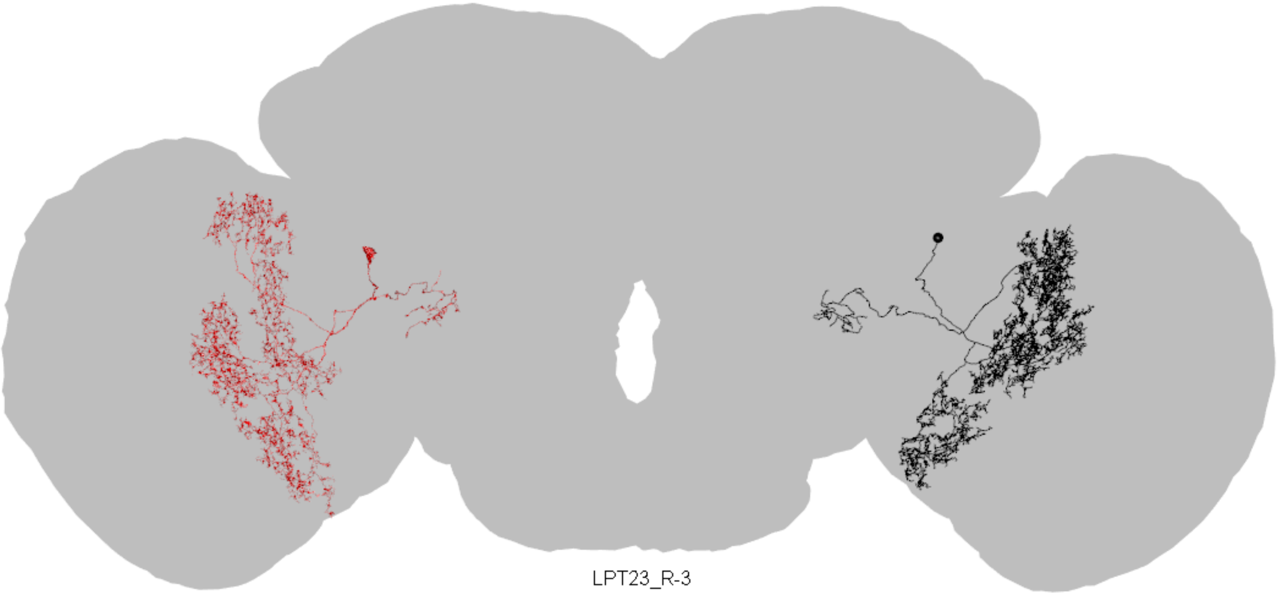

LPT23_R-3

Flywire ID = 720575940619638395

CATMAID skid = 4504557

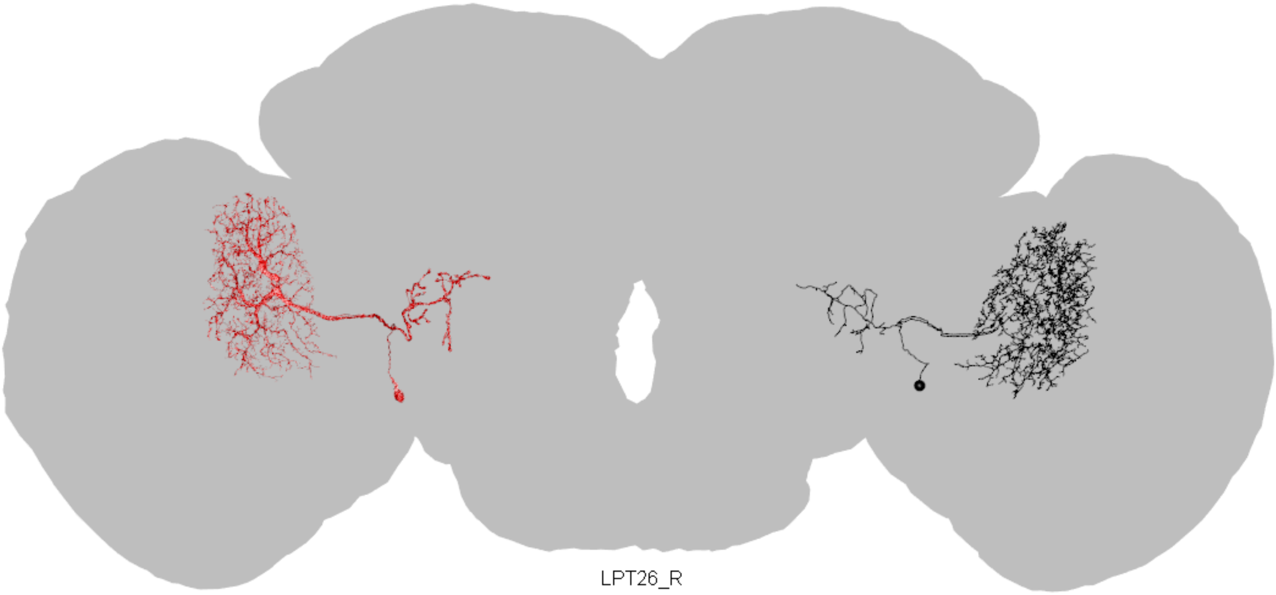

LPT26_R

Flywire ID = 720575940630691895

CATMAID skid = 1107045

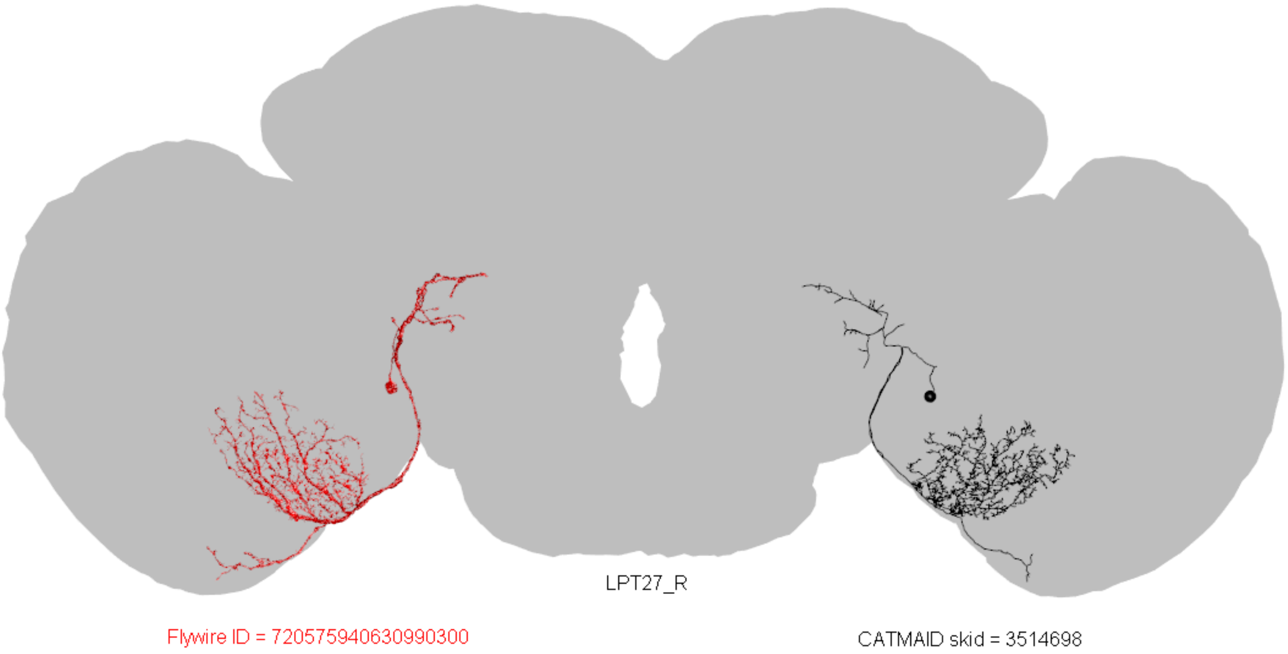
LPT27_R
Flywire ID = 720575940630990300
CATMAID skid = 3514698

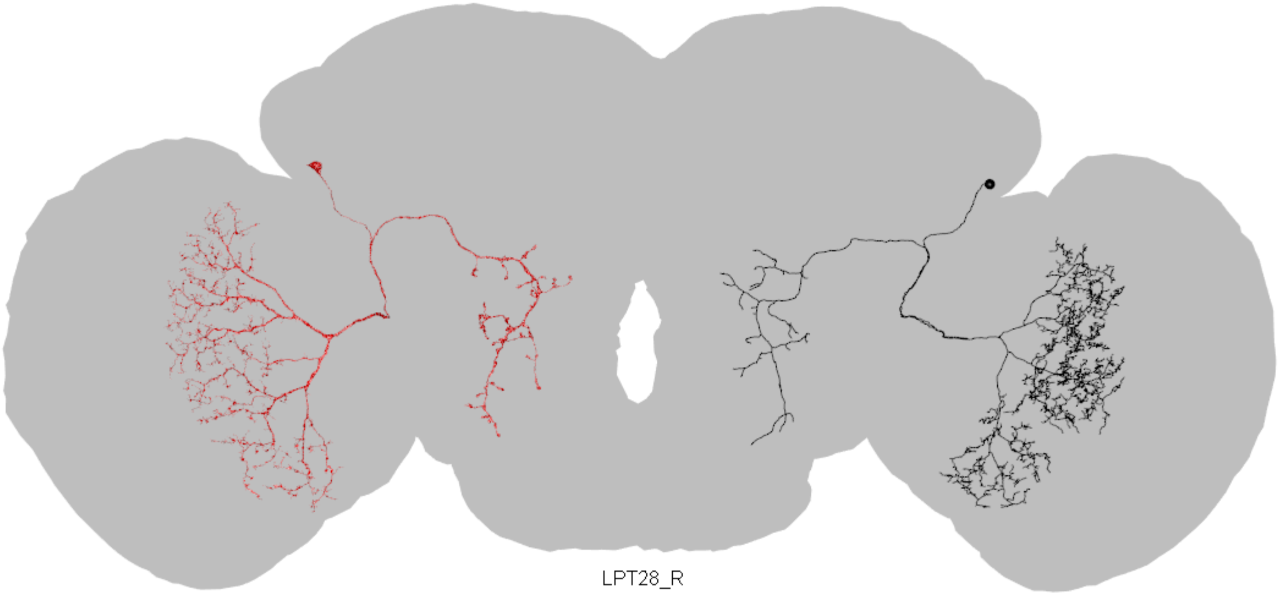

LPT28_R

Flywire ID = 720575940620898324

CATMAID skid = 1071860

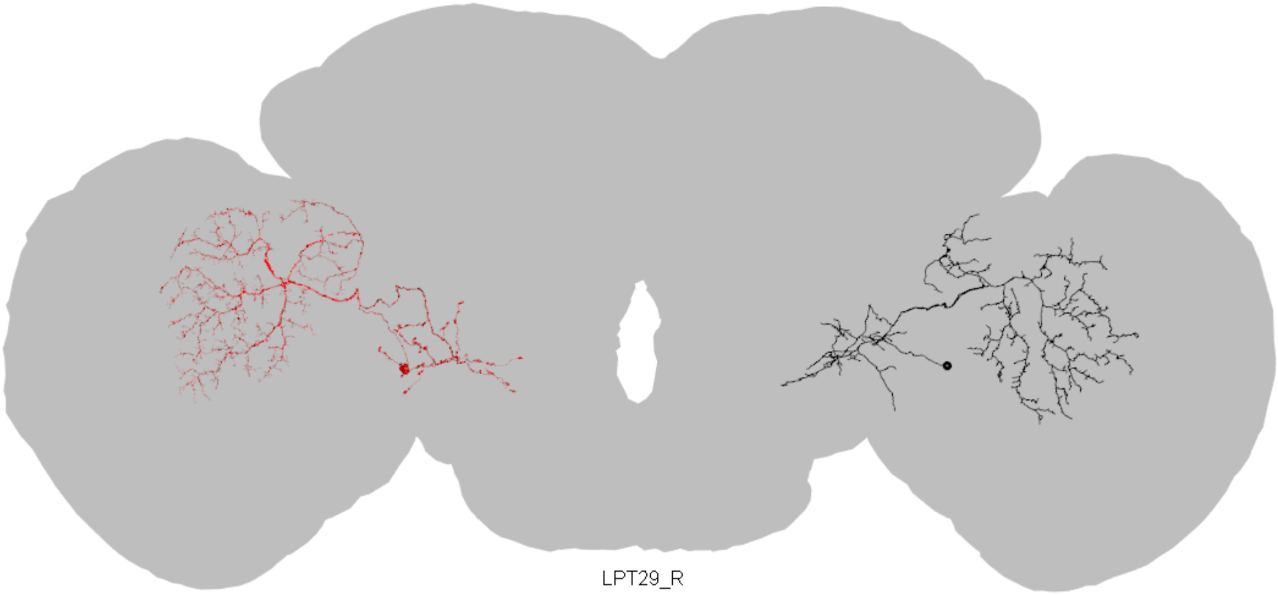
LPT29_R
Flywire ID = 720575940632216899
CATMAID skid = 7760435

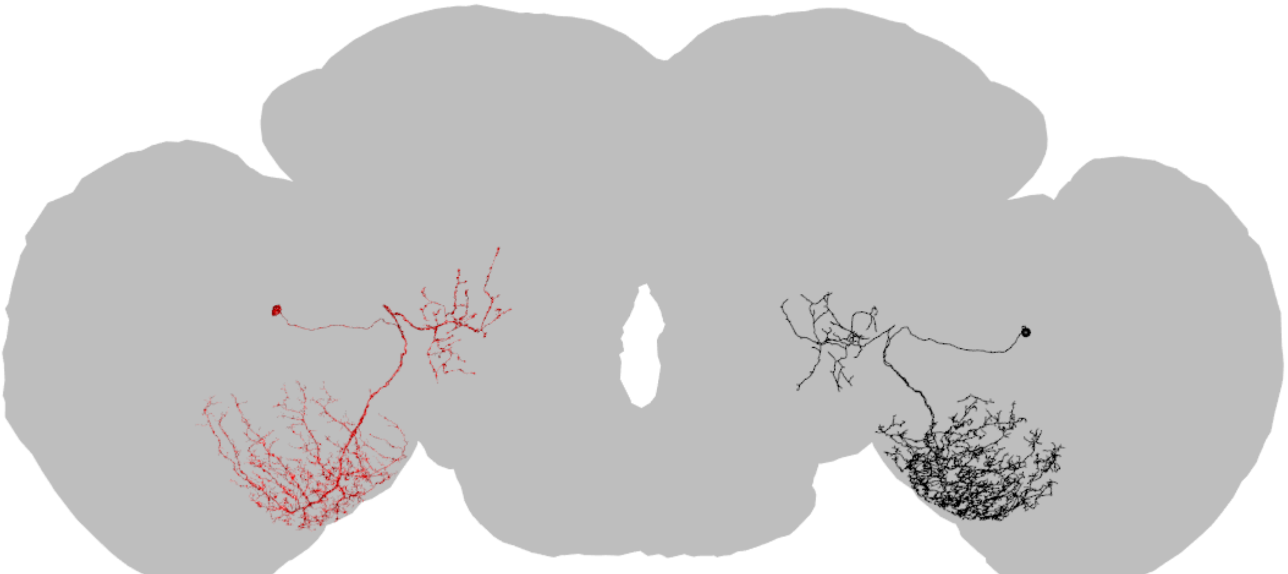

LPT30_R

Flywire ID = 720575940624047654

CATMAID skid = 7674107

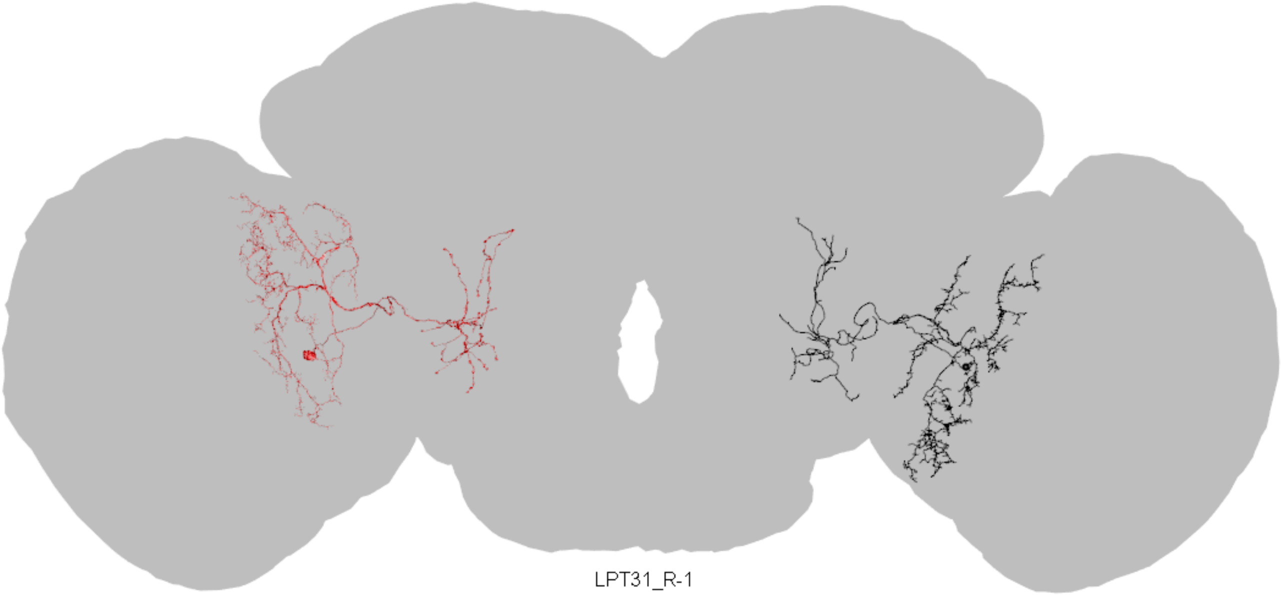


Flywire ID = 720575940634166242

CATMAID skid = 7690273

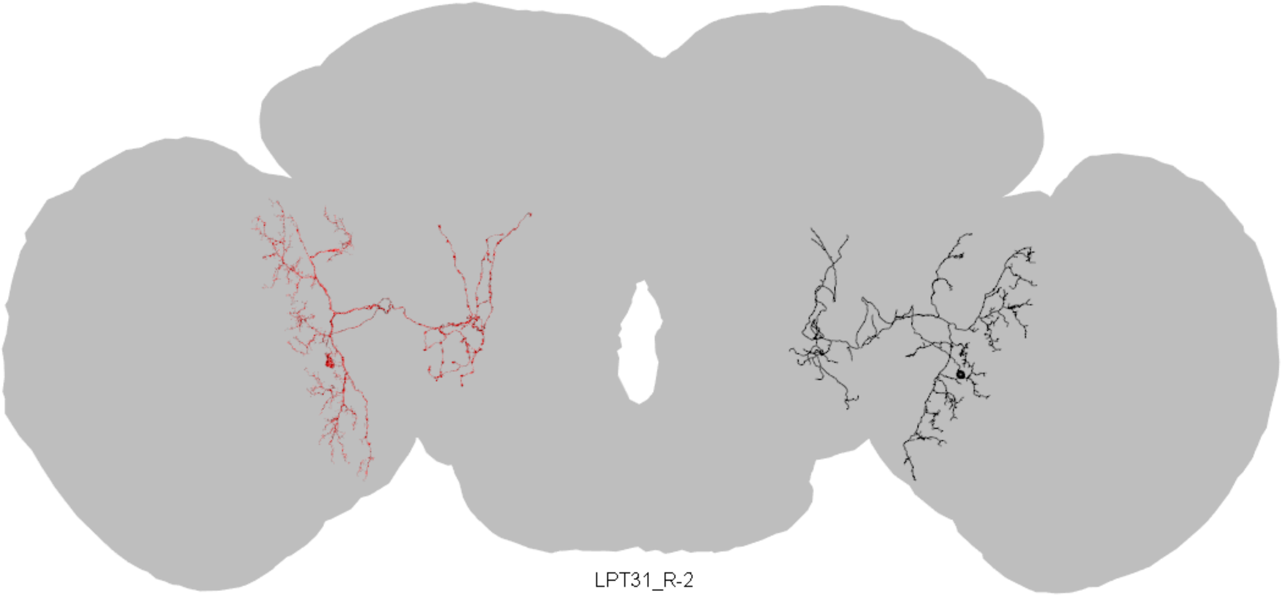

LPT31_R-2

Flywire ID = 720575940616260146

CATMAID skid = 7311408

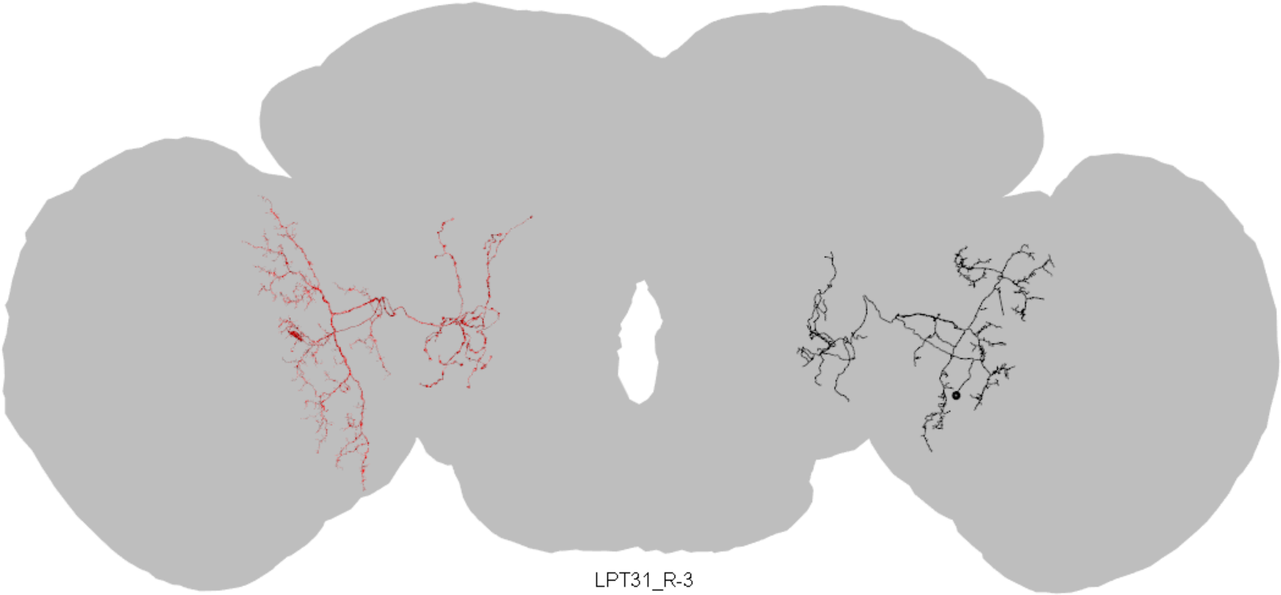

LPT31_R-3

Flywire ID = 720575940629577475

CATMAID skid = 7311493

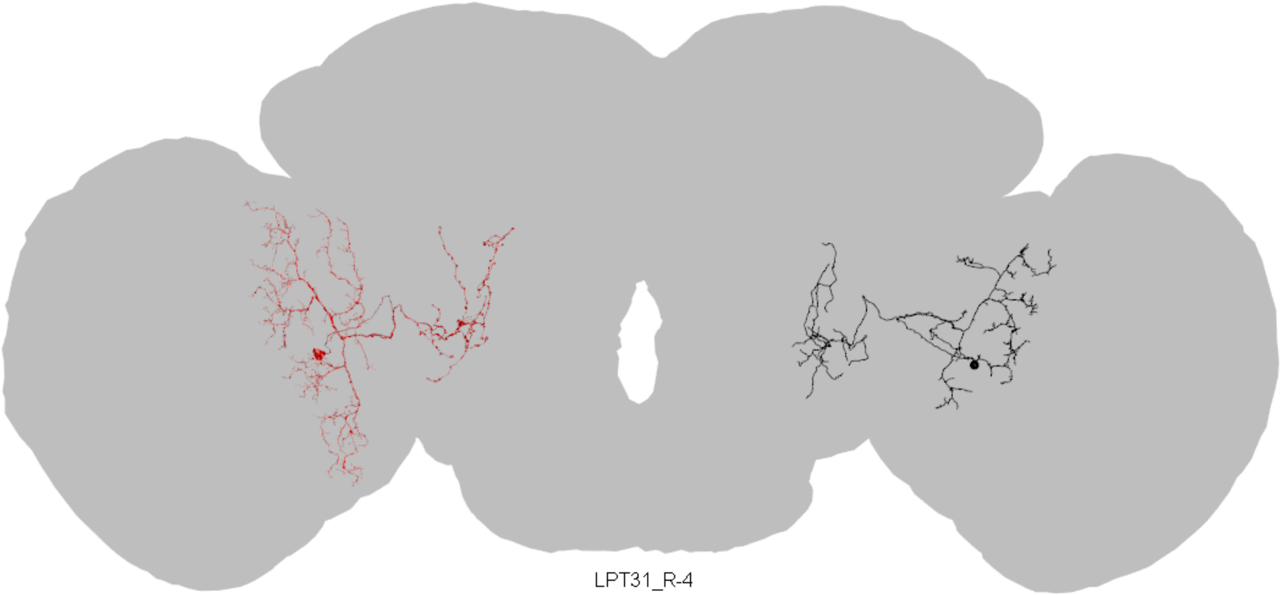

LPT31_R-4

Flywire ID = 720575940629567055

CATMAID skid = 7694305

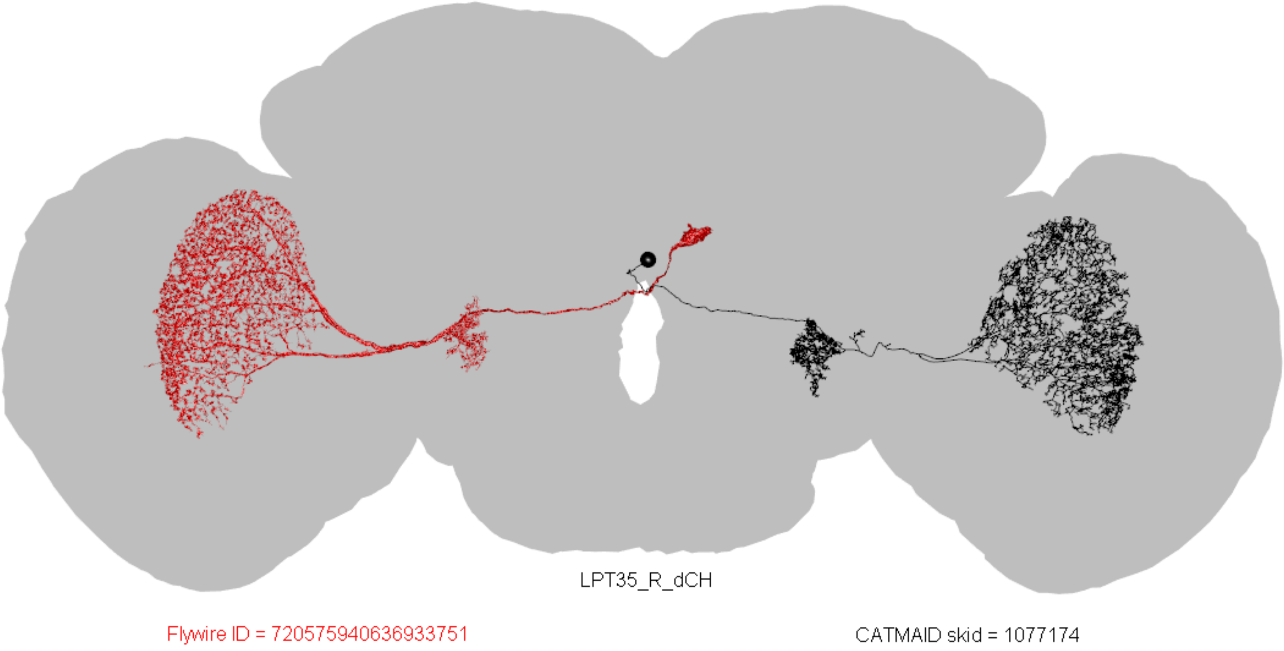
LPT35_R_dCH
Flywire ID = 720575940636933751
CATMAID skid = 1077174

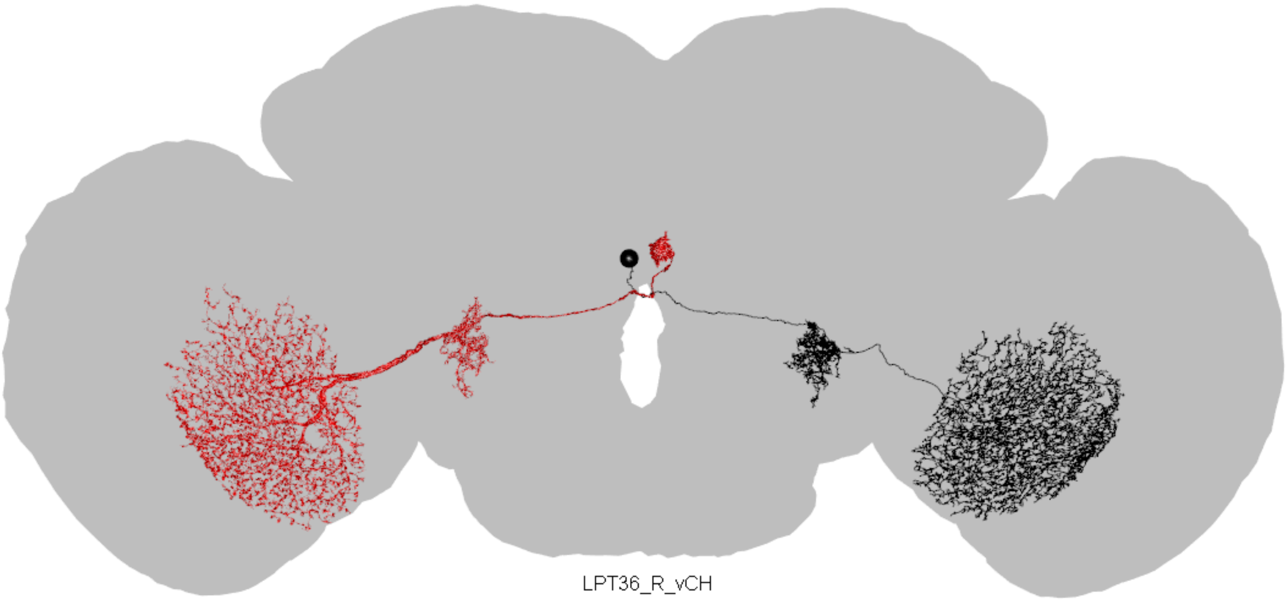

LPT36_R_vCH

Flywire ID = 720575940627138562

CATMAID skid = 1078535

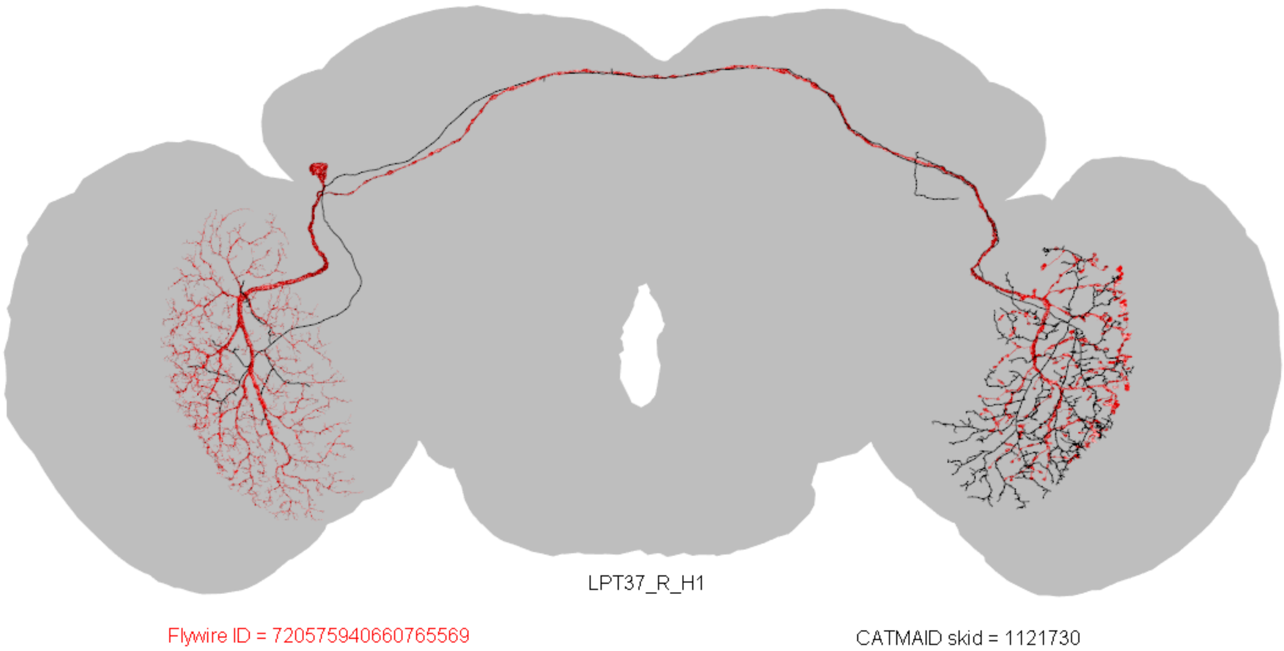
LPT37_R_H1
Flywire ID = 720575940660765569
CATMAID skid = 1121730

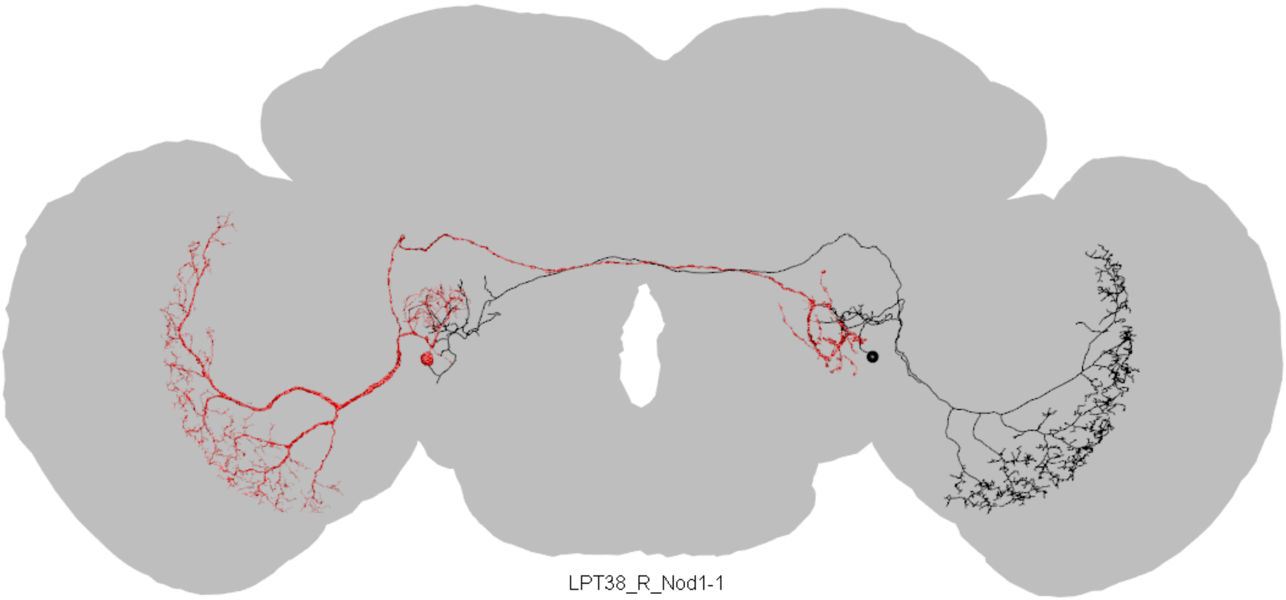

LPT38_R_Nod1-1

Flywire ID = 720575940609132043

CATMAID skid = 1054753

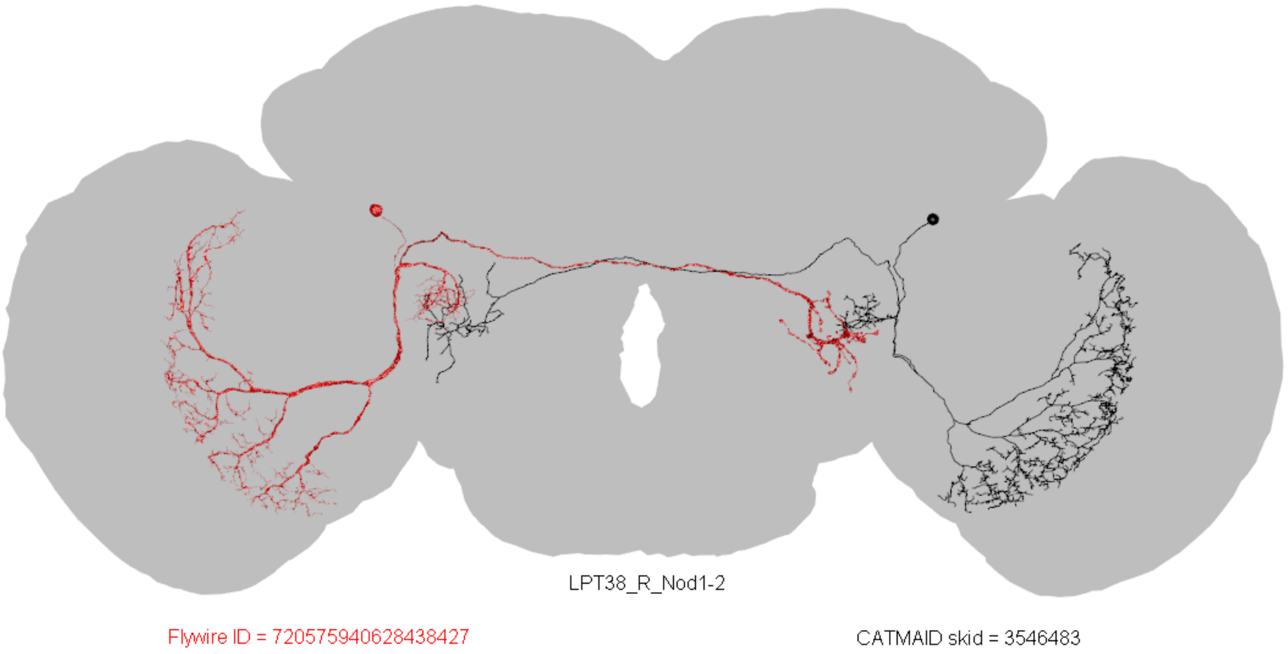
LPT38_R_Nod1-2
Flywire ID = 720575940628438427
CATMAID skid = 3546483

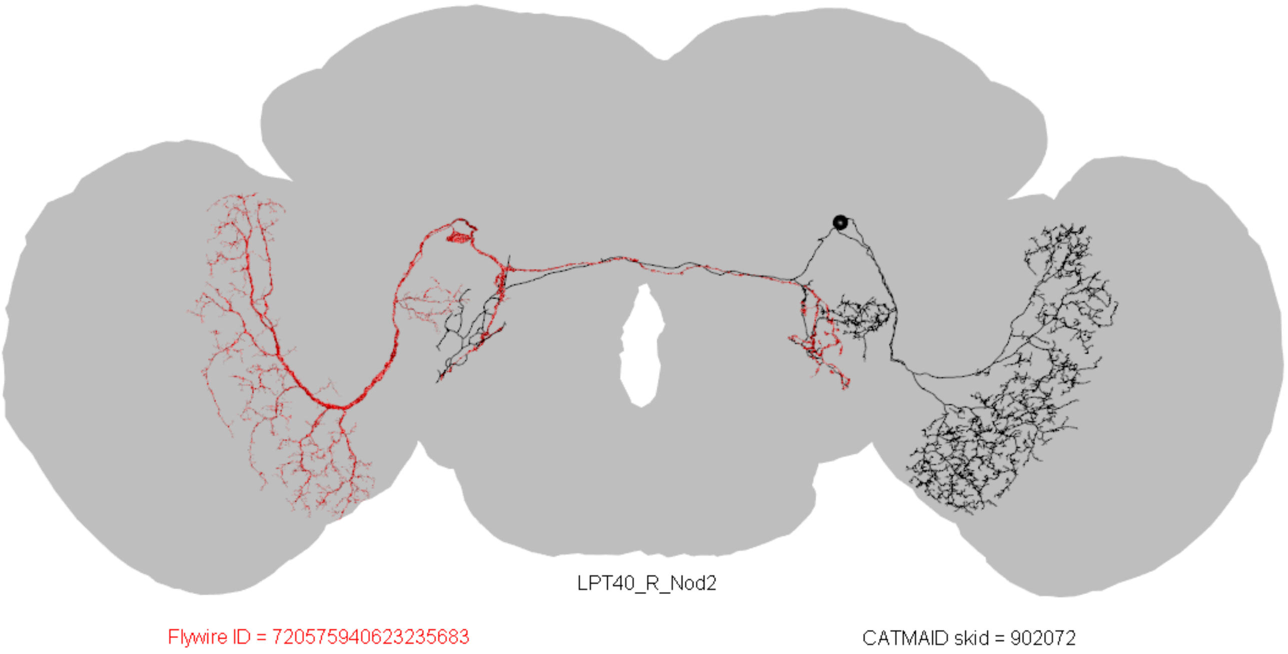
LPT40_R_Nod2
Flywire ID = 720575940623235683
CATMAID skid = 902072

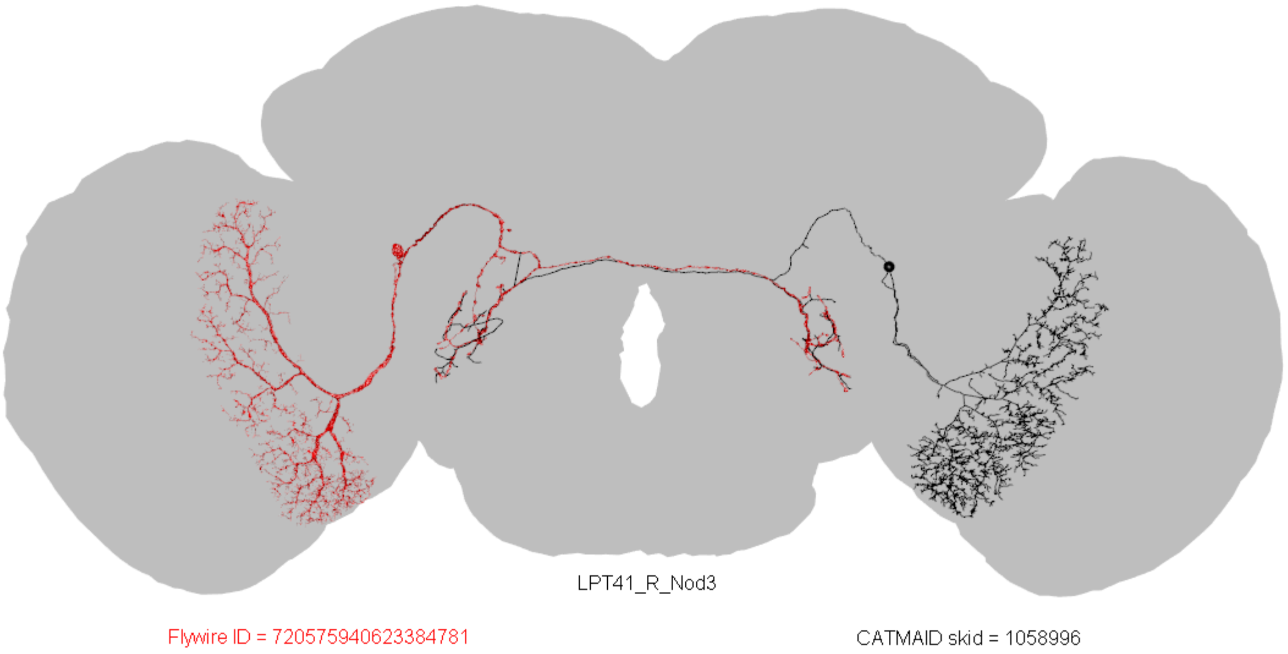
LPT41_R_Nod3
Flywire ID = 720575940623384781
CATMAID skid = 1058996

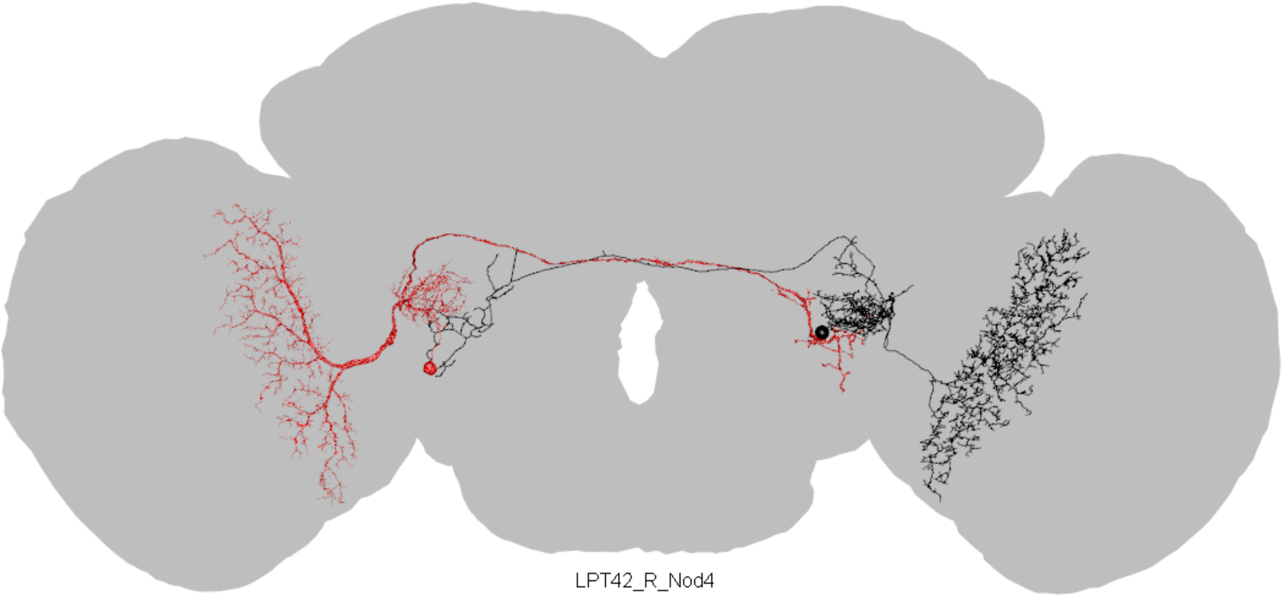

LPT42_R_Nod4

Flywire ID = 720575940625992781

CATMAID skid = 1106958

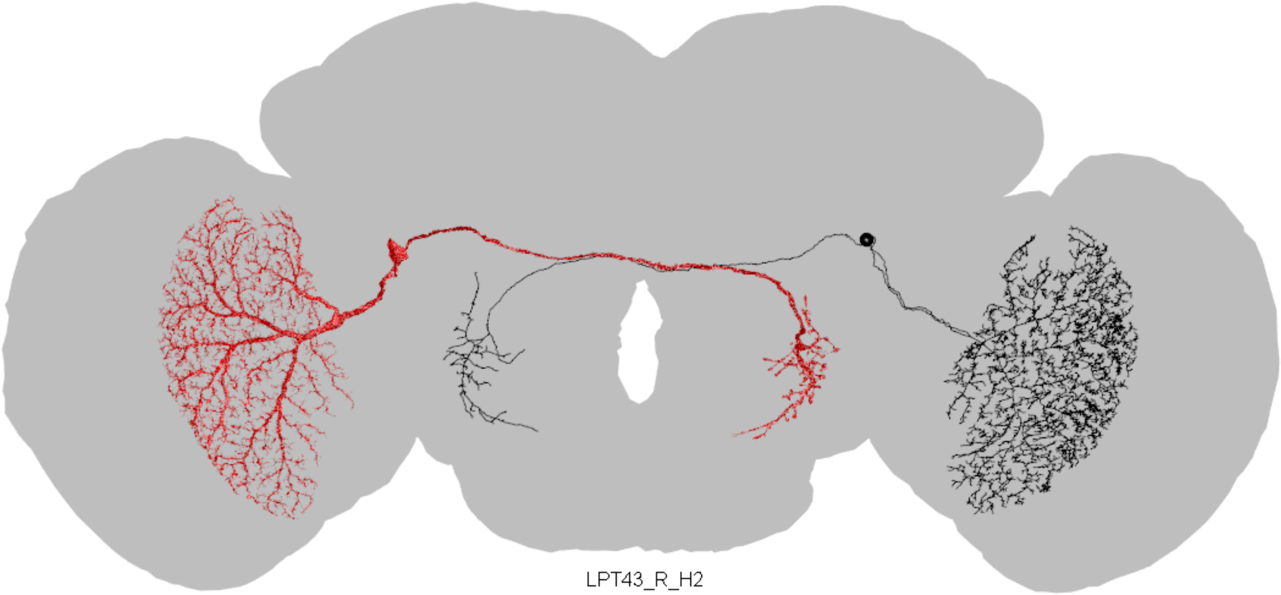

LPT43_R_H2

Flywire ID = 720575940627079938

CATMAID skid = 1088678

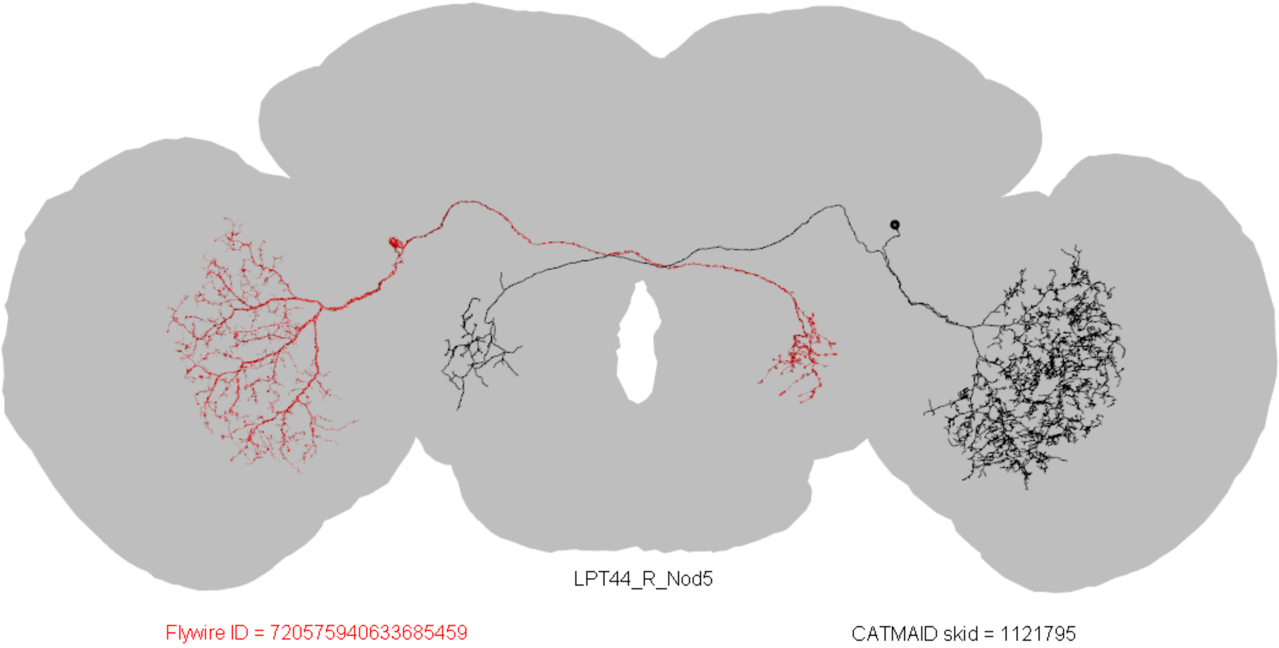
LPT44_R_Nod5
Flywire ID = 720575940633685459
CATMAID skid = 1121795

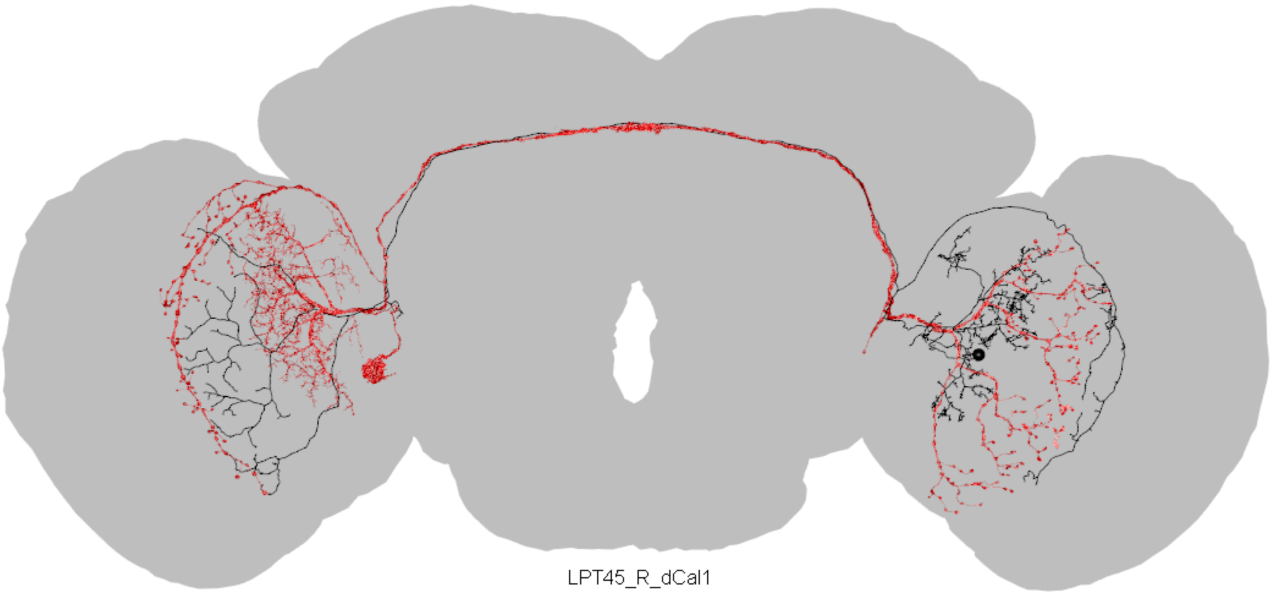

LPT45_R_dCal1

Flywire ID = 720575940628350997

CATMAID skid = 7204844

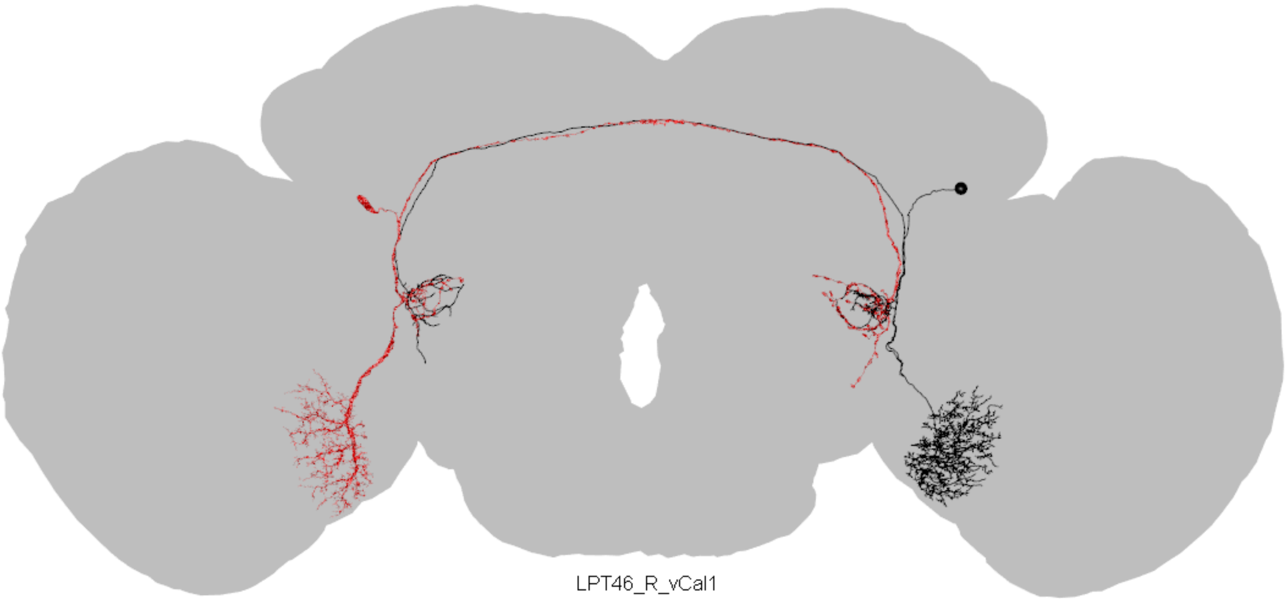

LPT46_R_vCal1

Flywire ID = 720575940641223888

CATMAID skid = 7510076

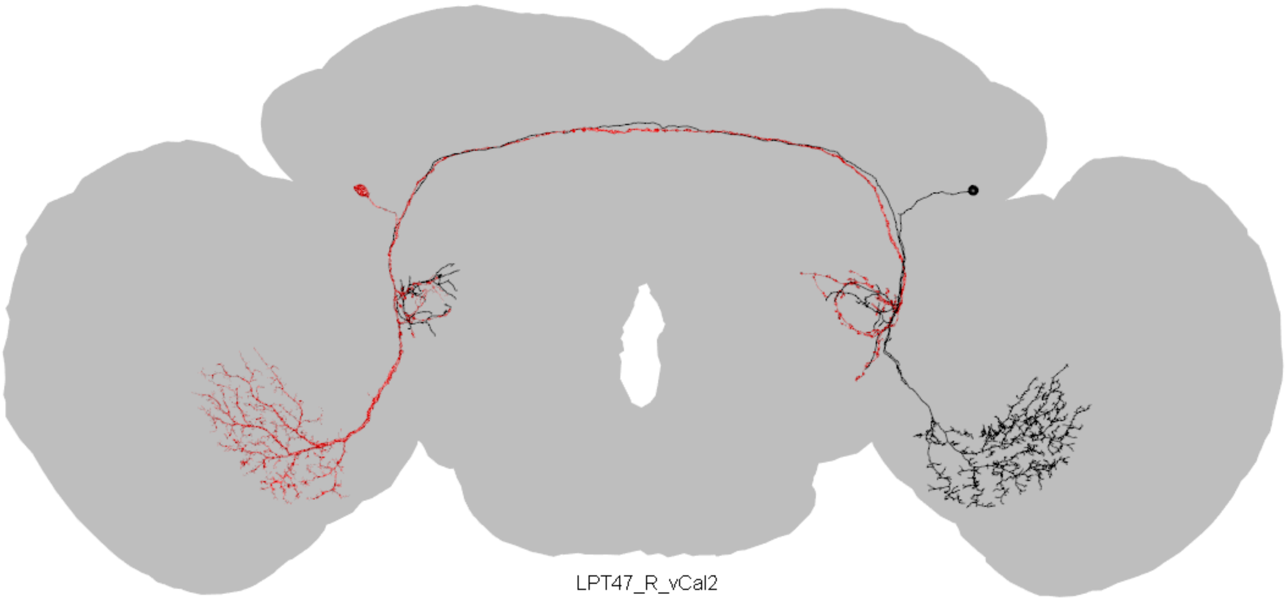

LPT47_R_vCal2

Flywire ID = 720575940618653524

CATMAID skid = 3529071

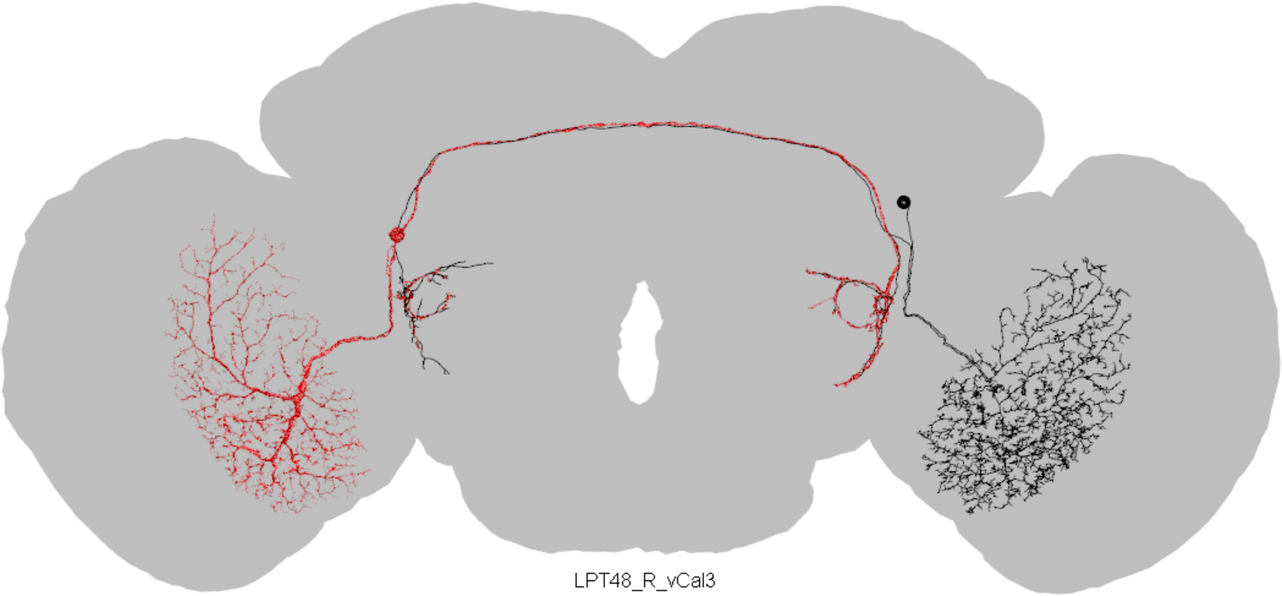

LPT48_R_vCal3

Flywire ID = 720575940626919780

CATMAID skid = 1056097

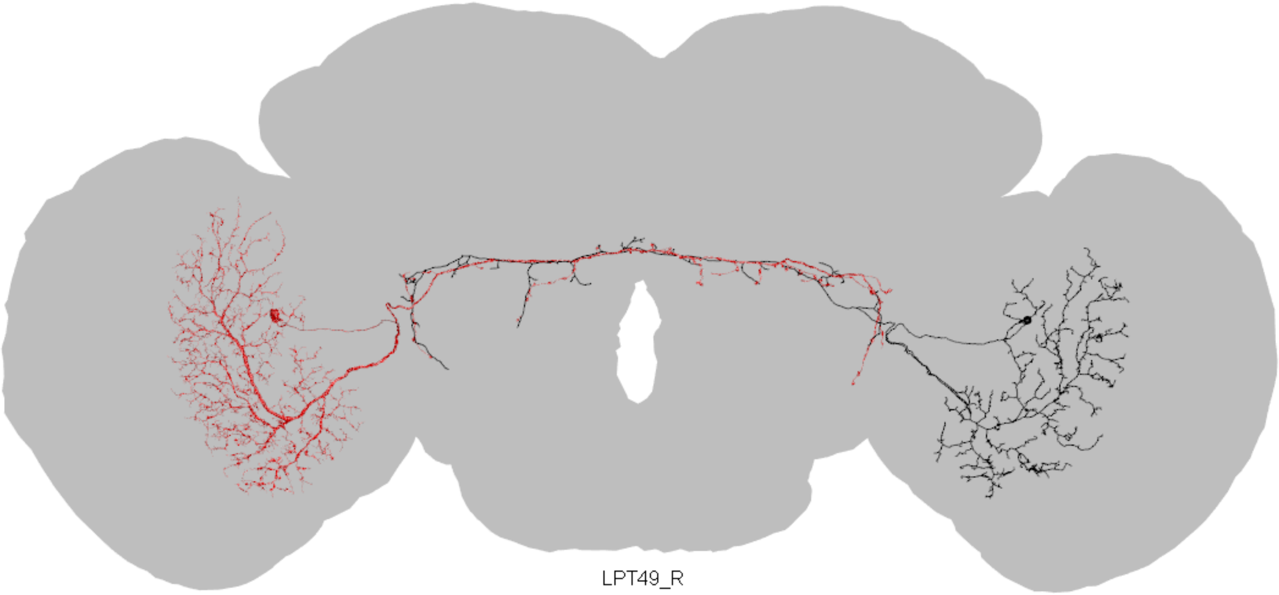

LPT49_R

Flywire ID = 720575940638155998

CATMAID skid = 1110693

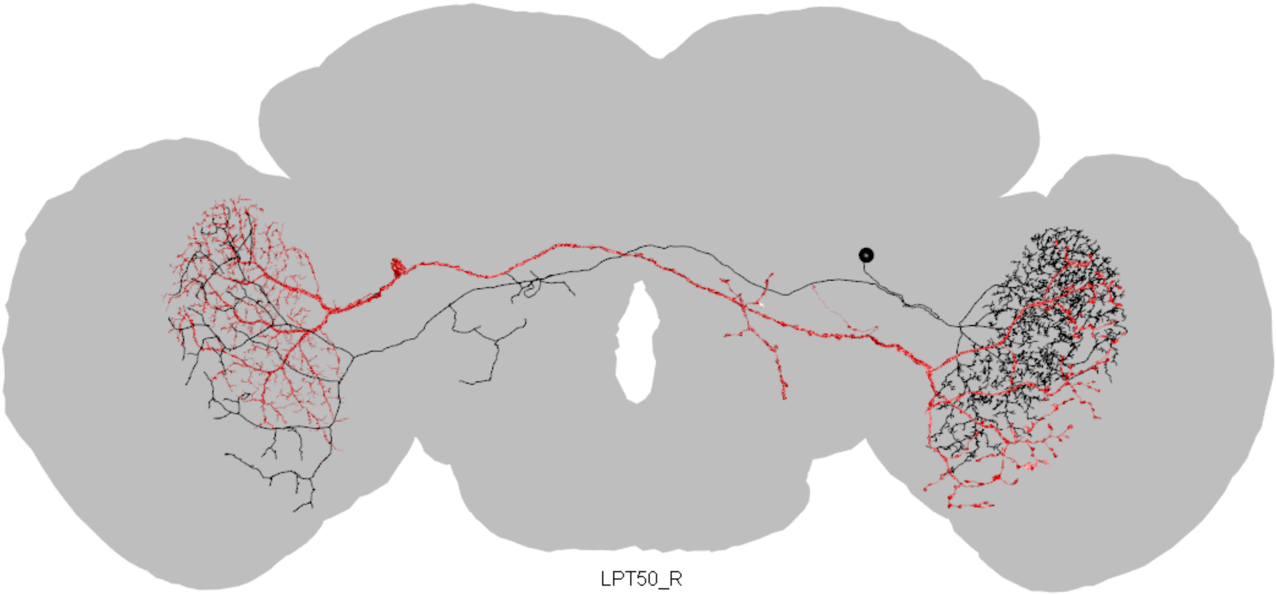

LPT50_R

Flywire ID = 720575940655599777

CATMAID skid = 905761

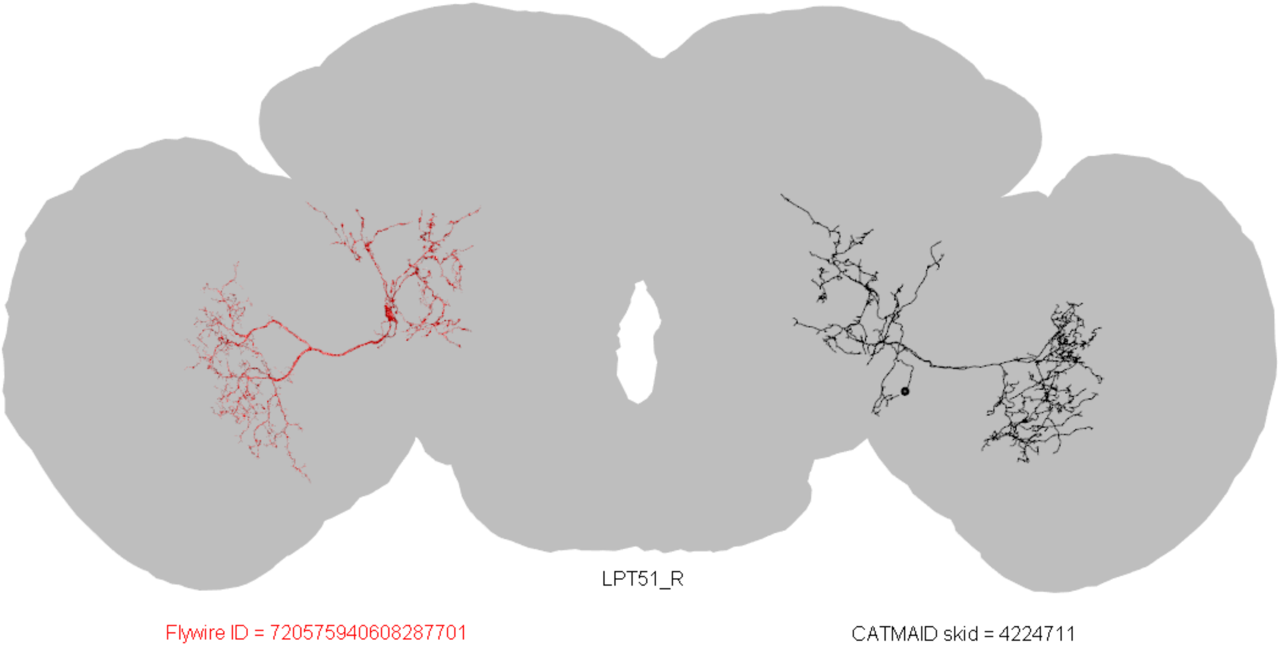

LPT51_R
Flywire ID = 720575940608287701
CATMAID skid = 4224711

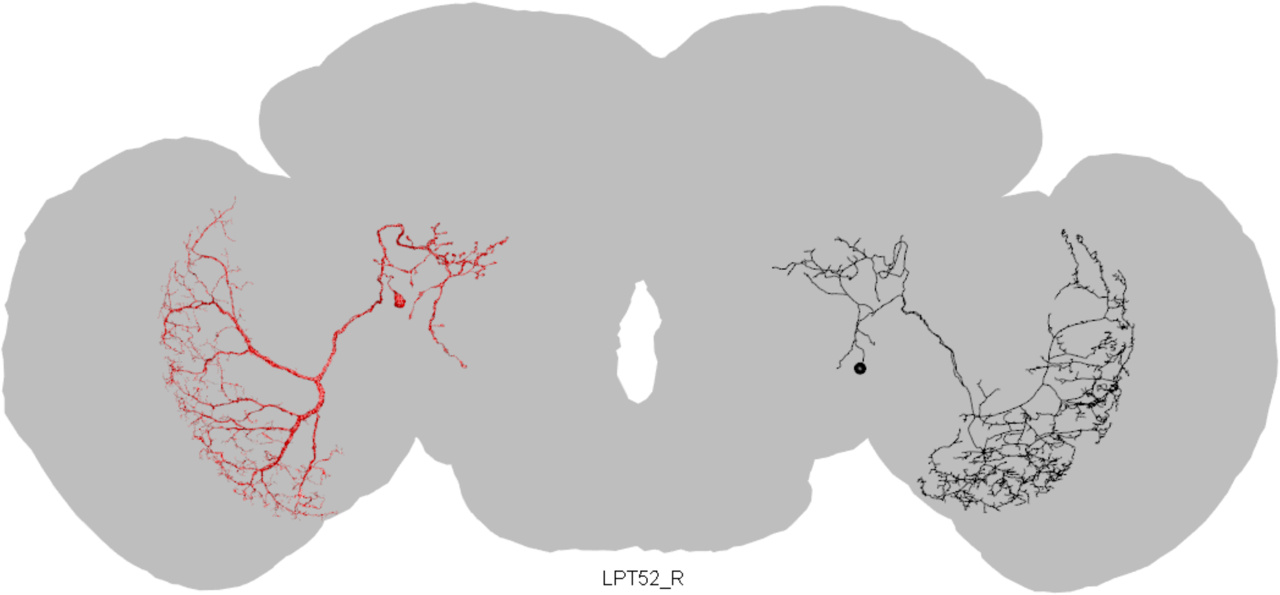

LPT52_R

Flywire ID = 720575940640716928

CATMAID skid = 1107296

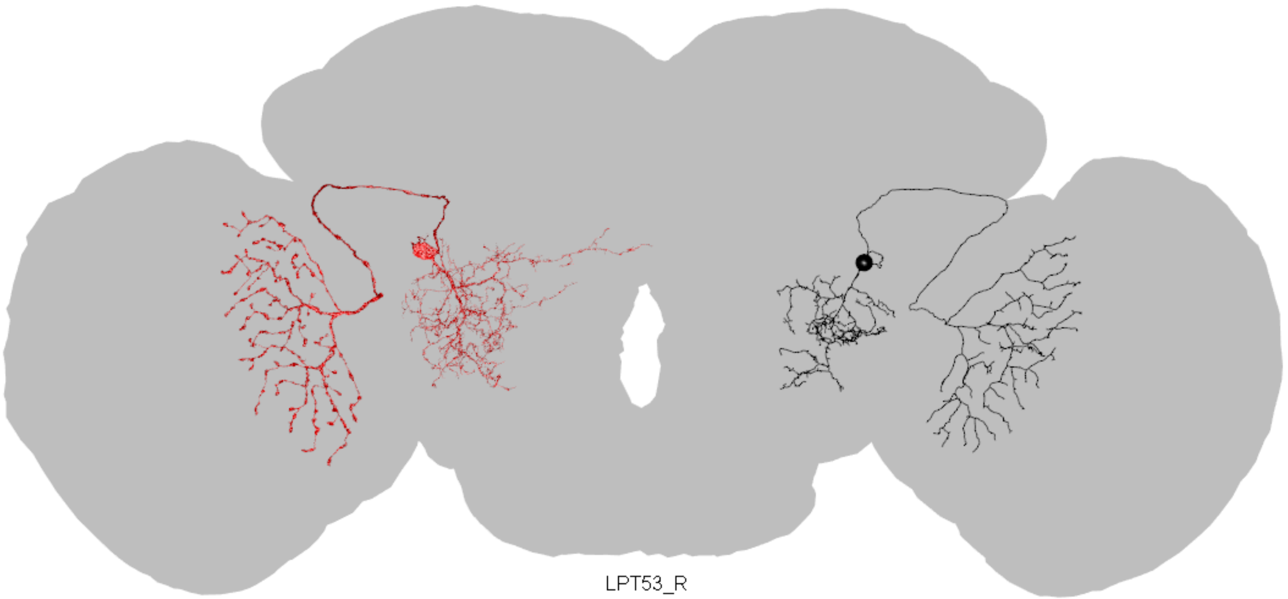

LPT53_R

Flywire ID = 720575940653093110

CATMAID skid = 1111992

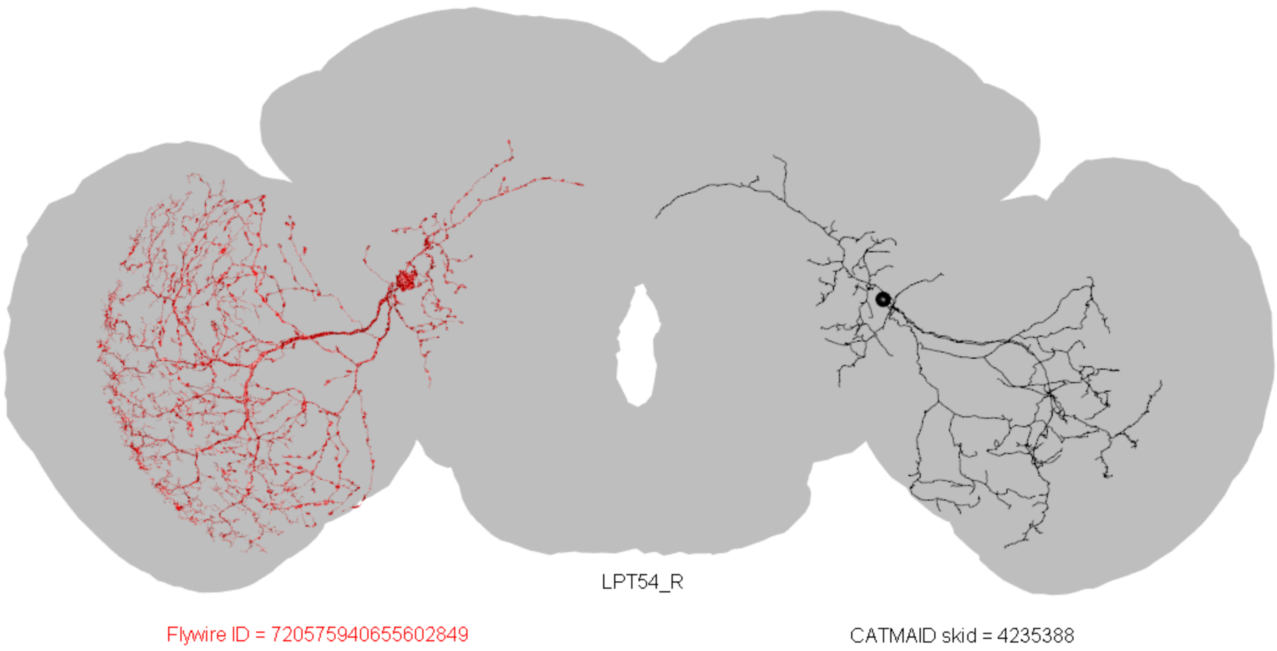
LPT54_R
Flywire ID = 720575940655602849
CATMAID skid = 4235388

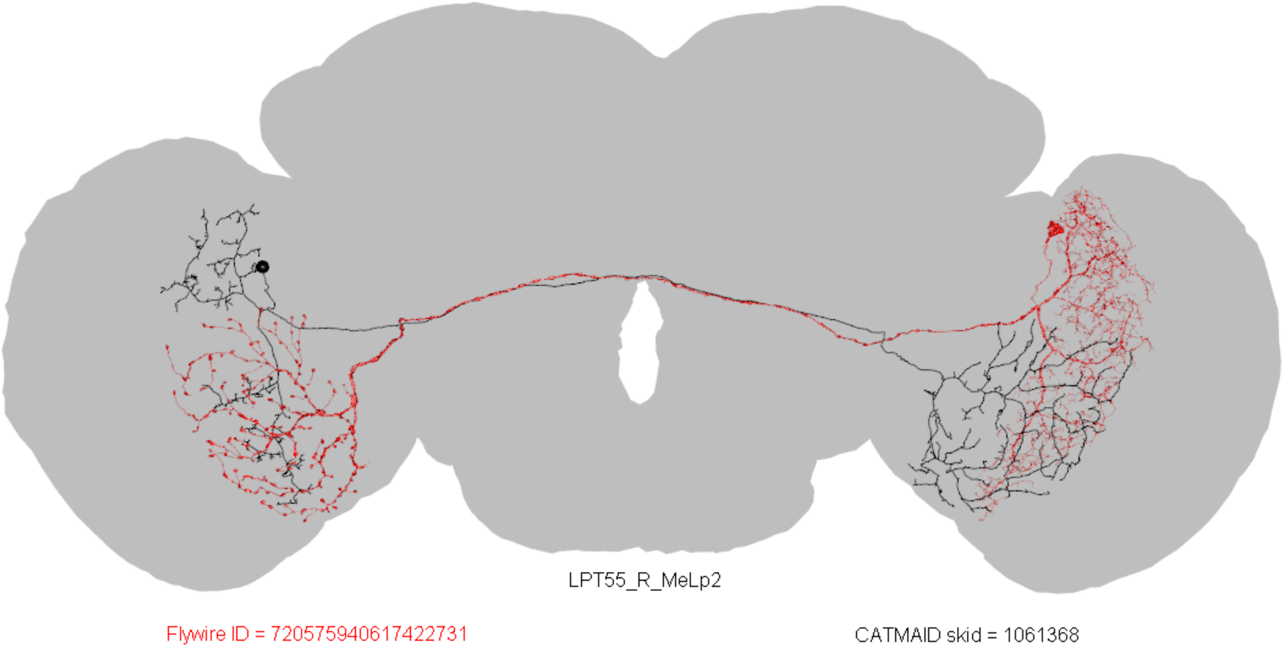
LPT55_R_MeLp2
Flywire ID = 720575940617422731
CATMAID skid = 1061368

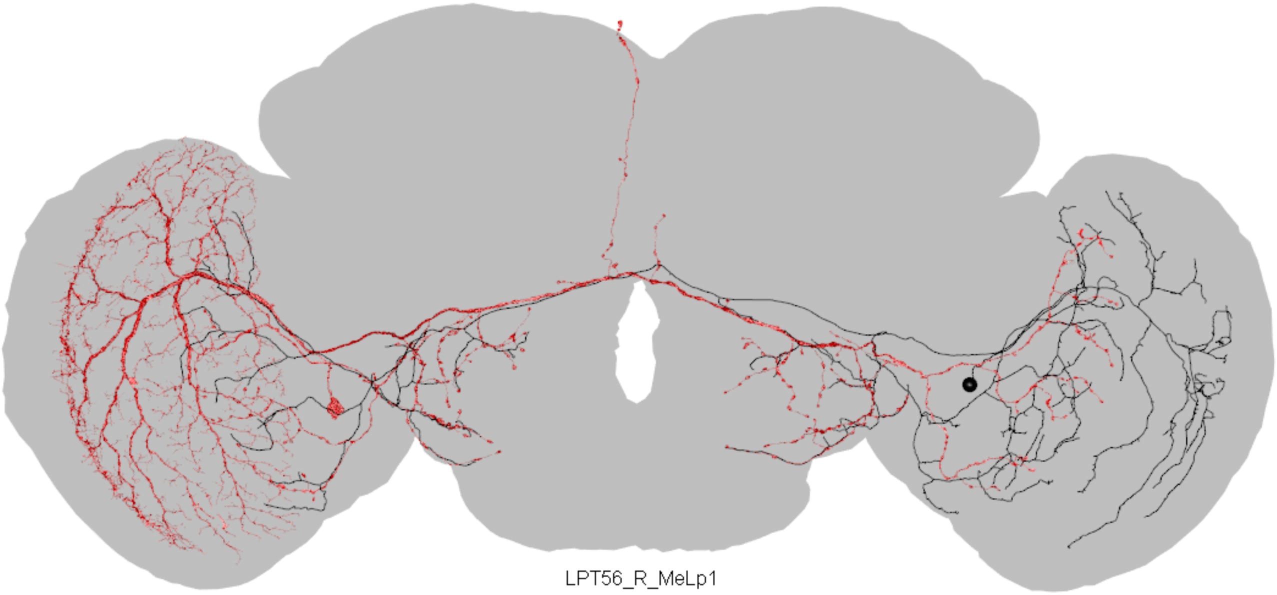
LPT56_R_MeLp1
Flywire ID = 720575940655602849
CATMAID skid = 3509520

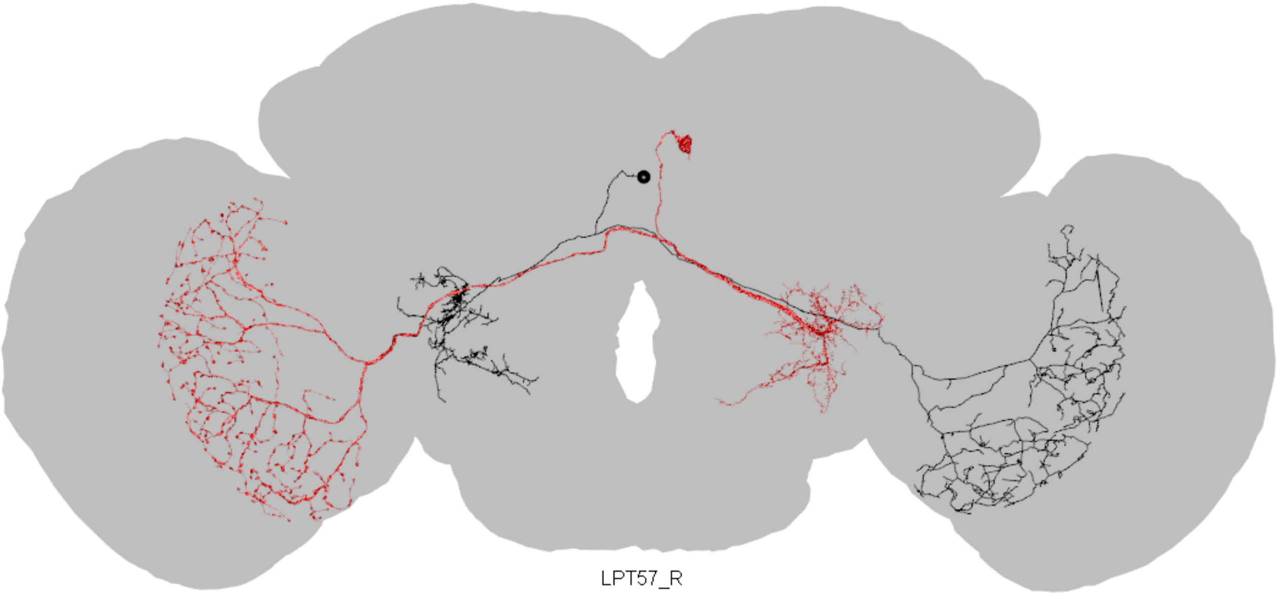

LPT57_R

Flywire ID = 720575940632041746

CATMAID skid = 886797

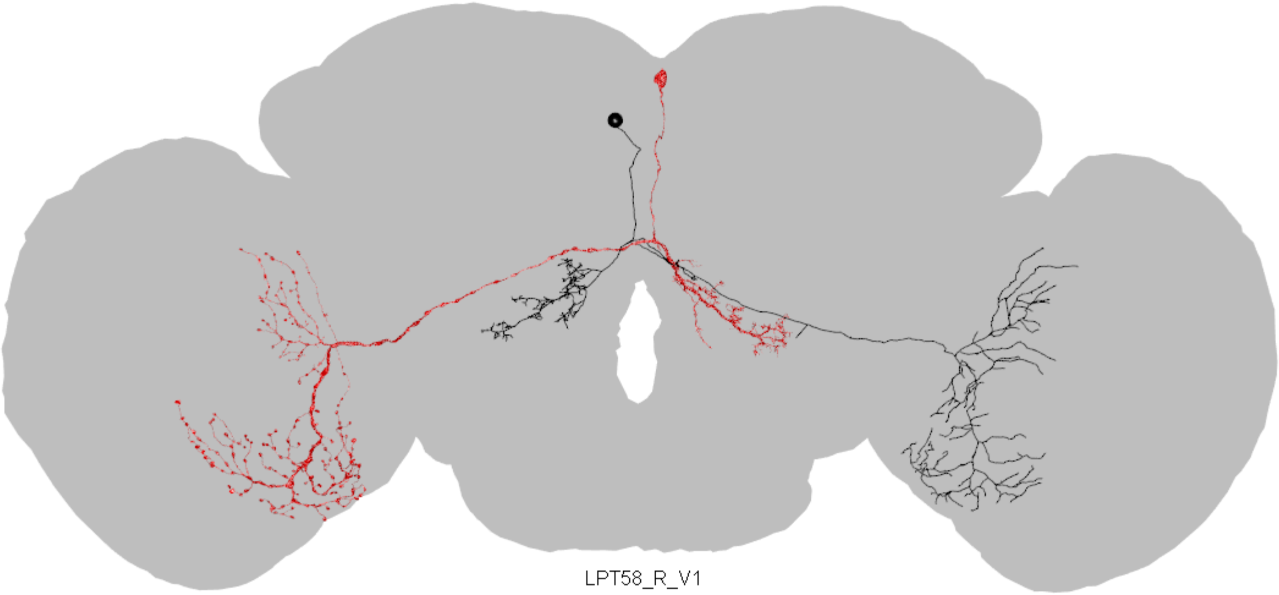

LPT58_R_V1

Flywire ID = 720575940621291873

CATMAID skid = 1059420
