## Supplemental File 4 for "A comprehensive neuroanatomical survey of the *Drosophila* Lobula Plate Tangential Neurons with predictions for their optic flow sensitivity"

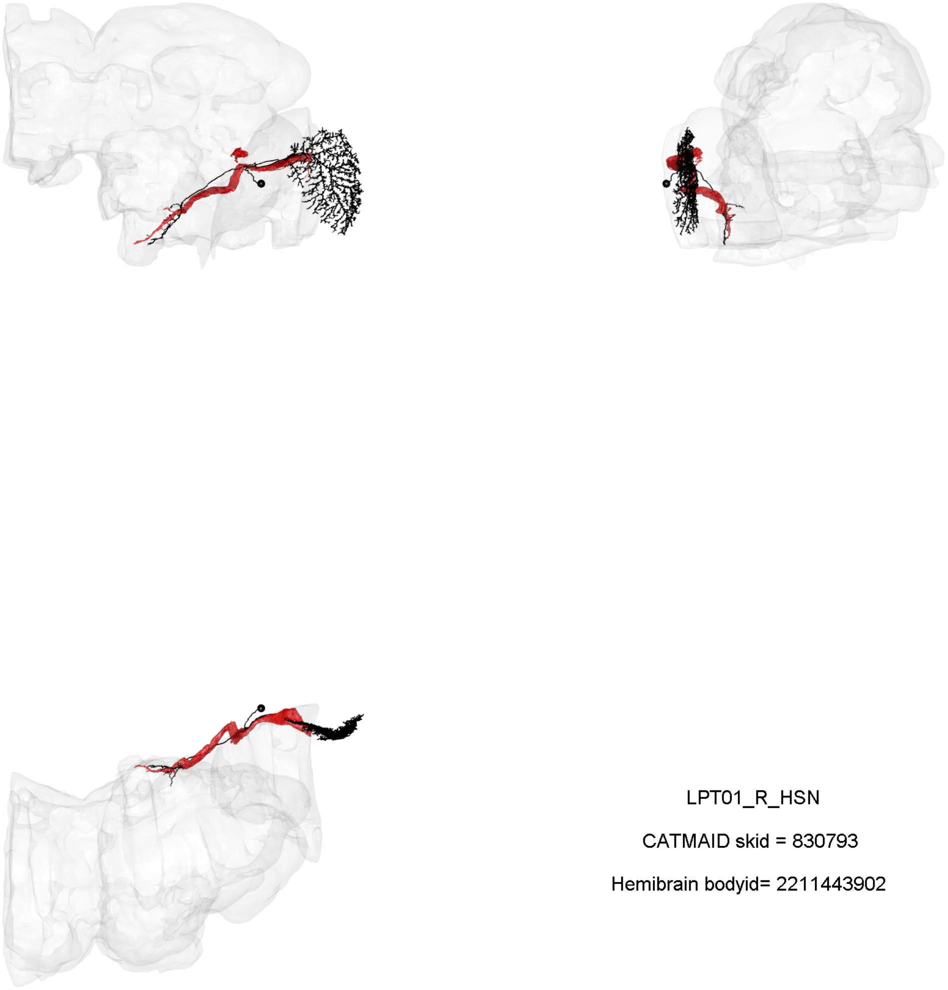

LPT01_R_HSN

CATMAID skid = 830793

Hemibrain bodyid= 2211443902

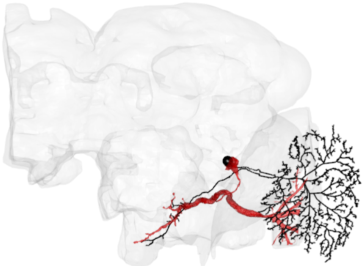

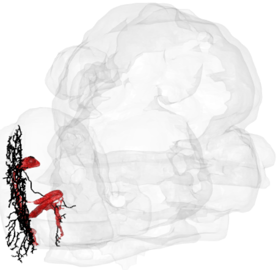

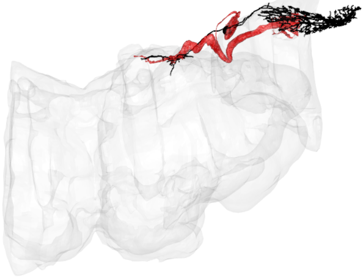

LPT02_R_HSE

CATMAID skid = 827034

Hemibrain bodyid= 1807537598

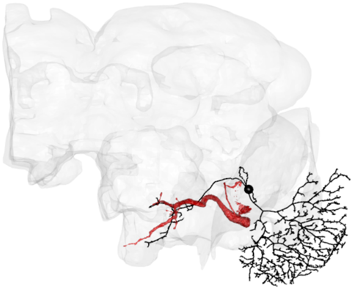

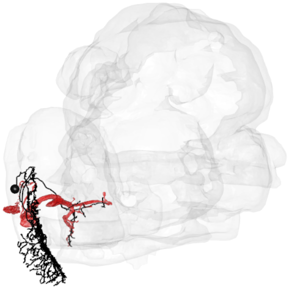

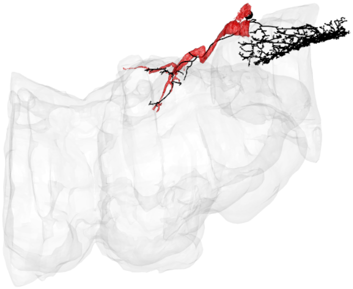

LPT03_R_HSS

CATMAID skid = 4058824

Hemibrain bodyid= 2179731270

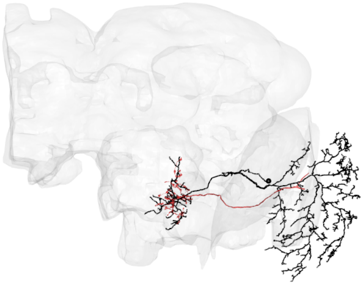

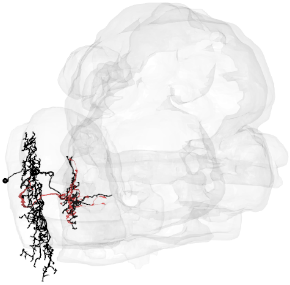

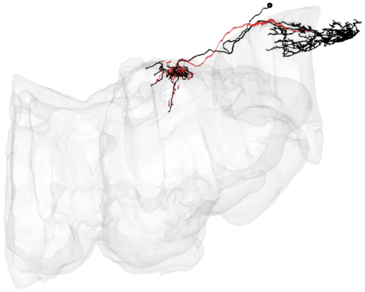

LPT04_R_HST

CATMAID skid = 985774

Hemibrain bodyid= 5813042688

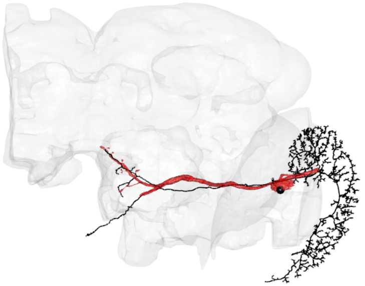

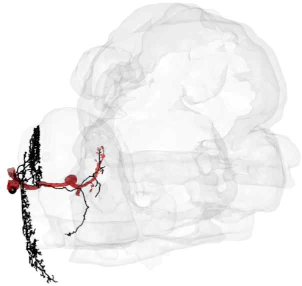

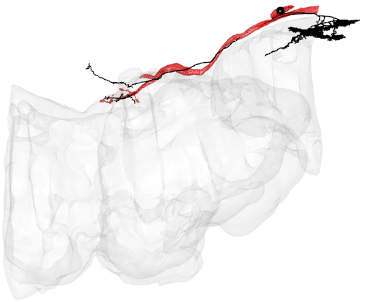

LPT05_R_VS1

CATMAID skid = 982897

Hemibrain bodyid= 5813024262

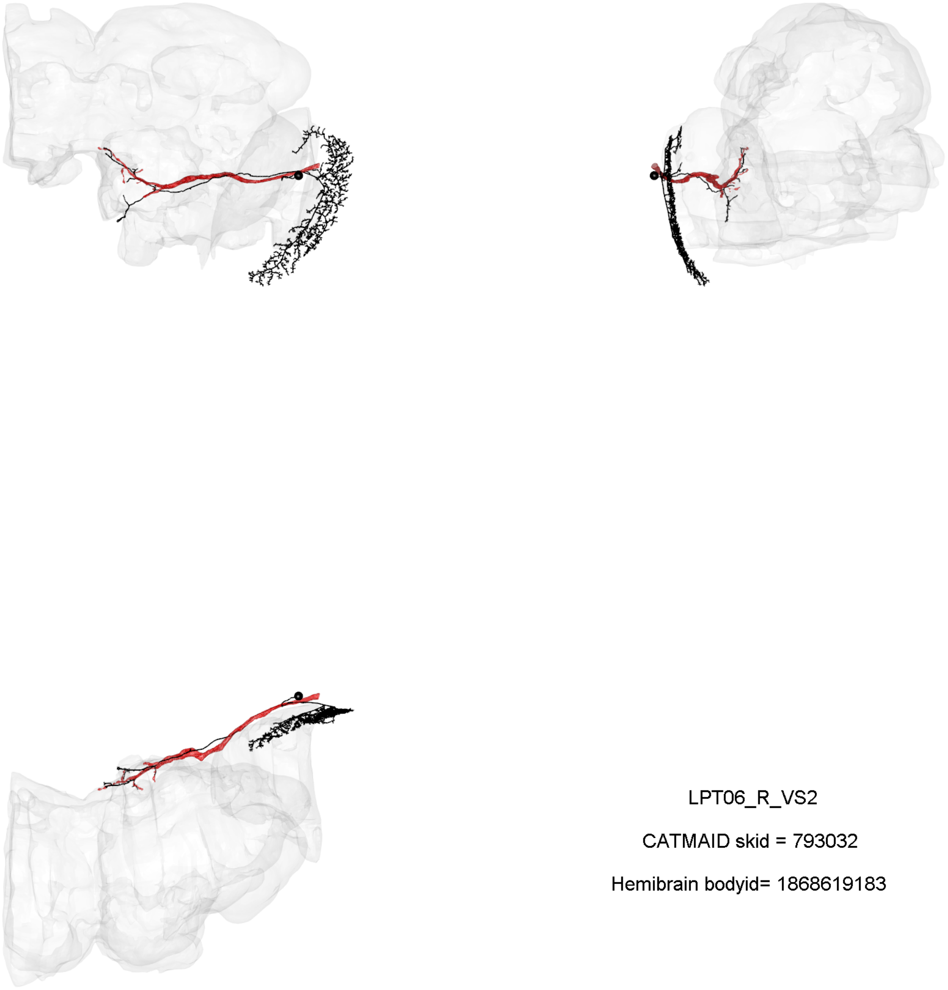

LPT06_R_VS2

CATMAID skid = 793032

Hemibrain bodyid= 1868619183

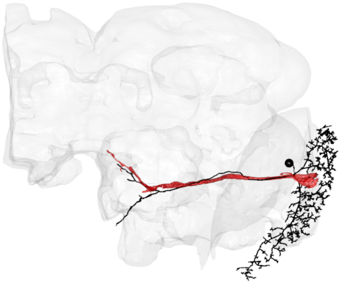
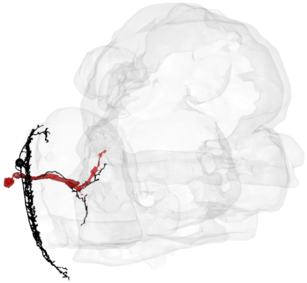
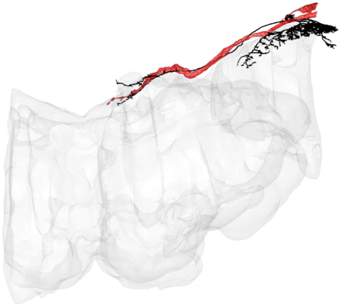

LPT07_R_VS3

CATMAID skid = 815776

Hemibrain bodyid= 5813025091

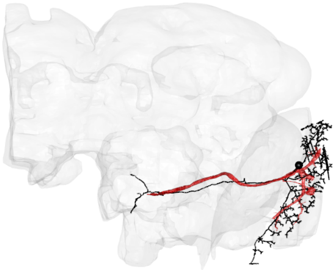

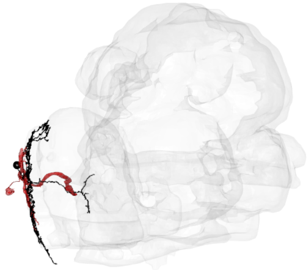

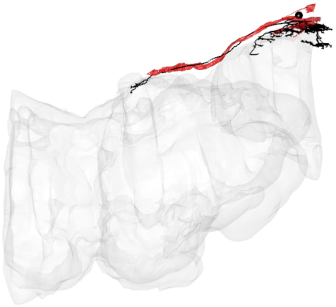

LPT08_R_VS4

CATMAID skid = 17686499

Hemibrain bodyid= 1557885051

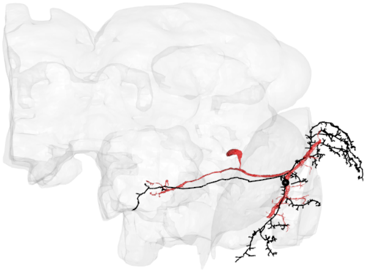

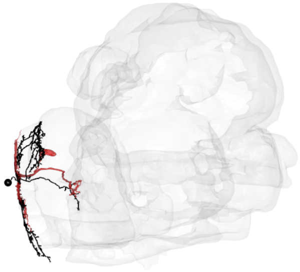

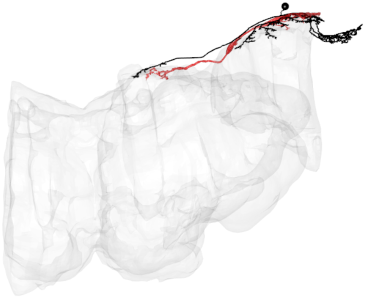

LPT09_R_VS5

CATMAID skid = 807401

Hemibrain bodyid= 5813033533

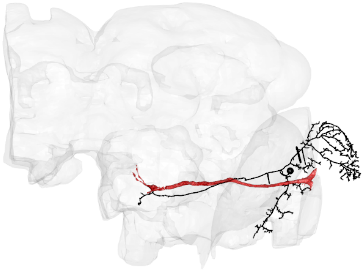

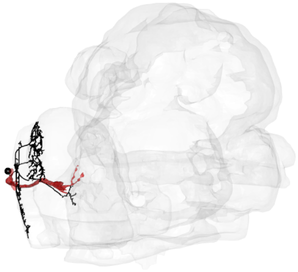

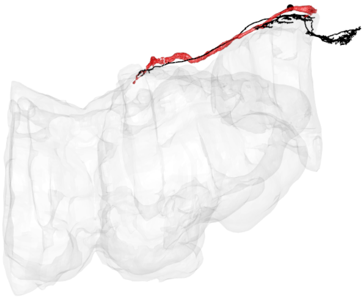

LPT10_R_VS6

CATMAID skid = 804539

Hemibrain bodyid= 1992161820

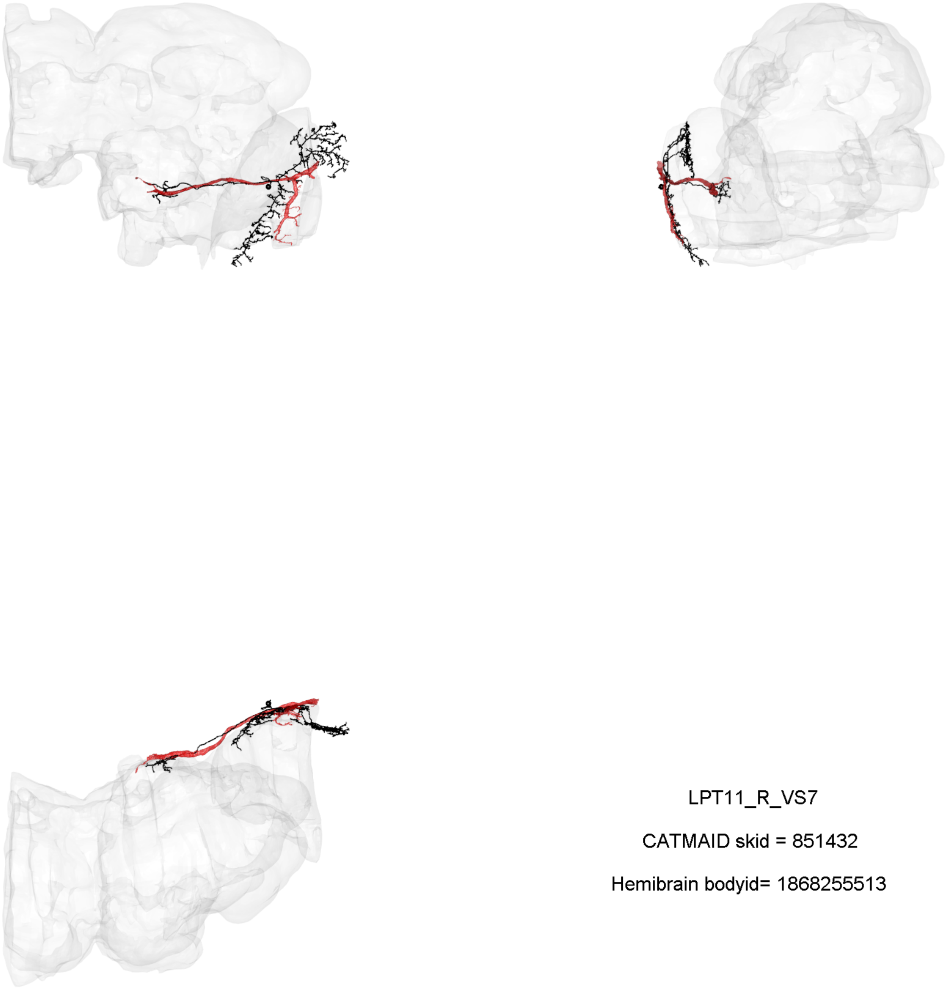

LPT11_R_VS7

CATMAID skid = 851432

Hemibrain bodyid= 1868255513

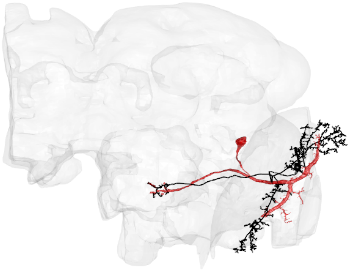

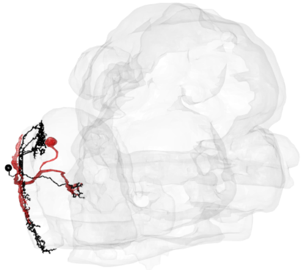

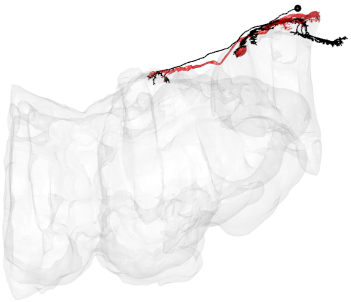

LPT12_R_VS8

CATMAID skid = 804092

Hemibrain bodyid= 1558230909

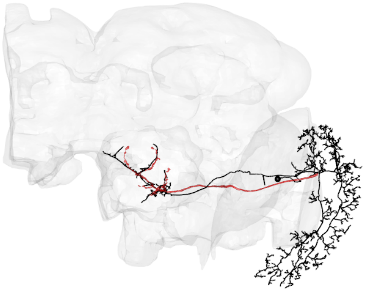

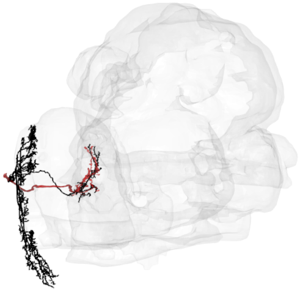

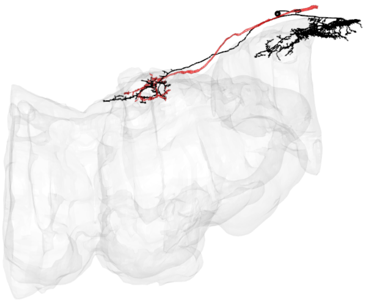

LPT13_R_VSm-1

CATMAID skid = 852286

Hemibrain bodyid= 1836516710

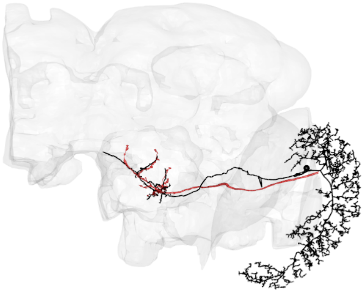

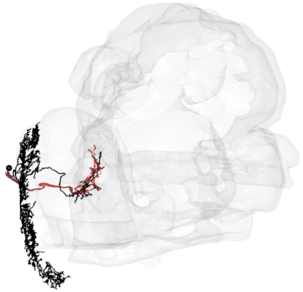

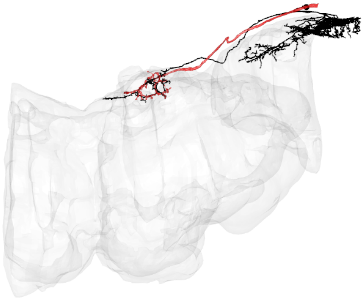

LPT13_R_VSm-2

CATMAID skid = 815241

Hemibrain bodyid= 1805481901

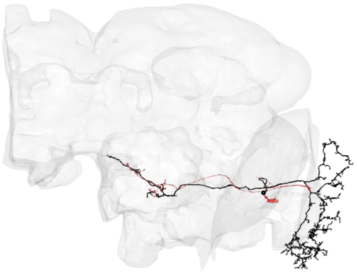

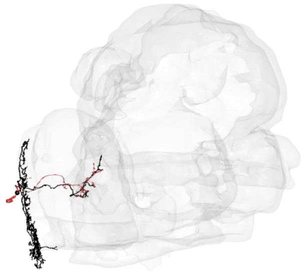

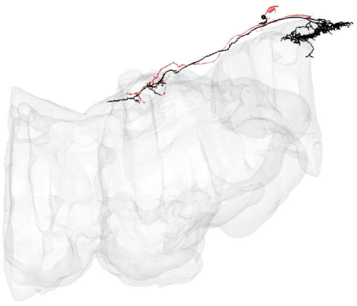

LPT15_R_VST1-2

CATMAID skid = 988674

Hemibrain bodyid= 5812993165

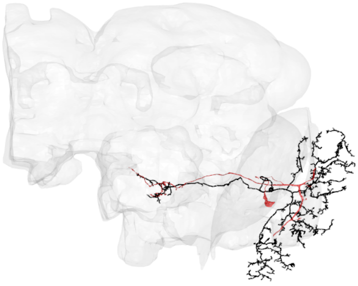

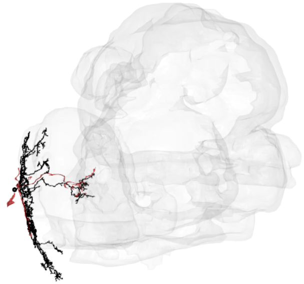

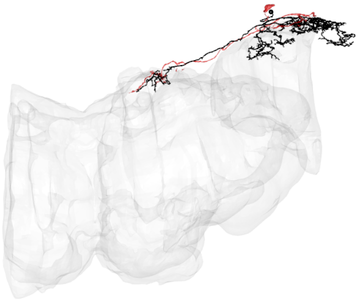

LPT17_R_VST2-1

CATMAID skid = 1123389

Hemibrain bodyid= 1837198715

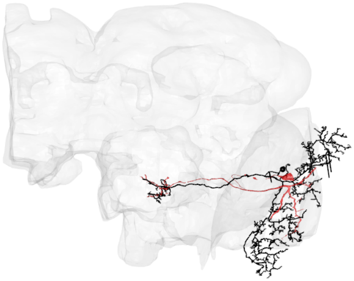
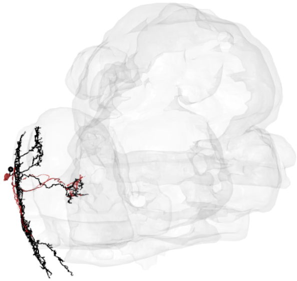
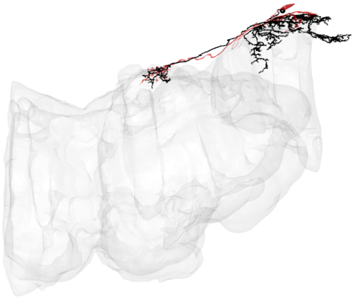

LPT17_R_VST2-2

CATMAID skid = 1112633

Hemibrain bodyid= 5813023313

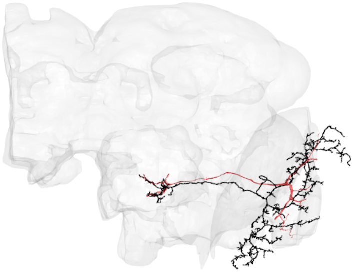

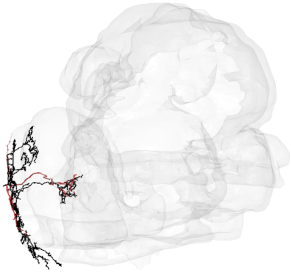

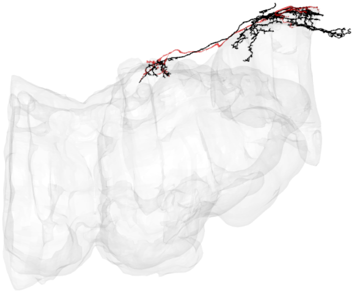

LPT17_R_VST2-3

CATMAID skid = 2852912

Hemibrain bodyid= 5813049185

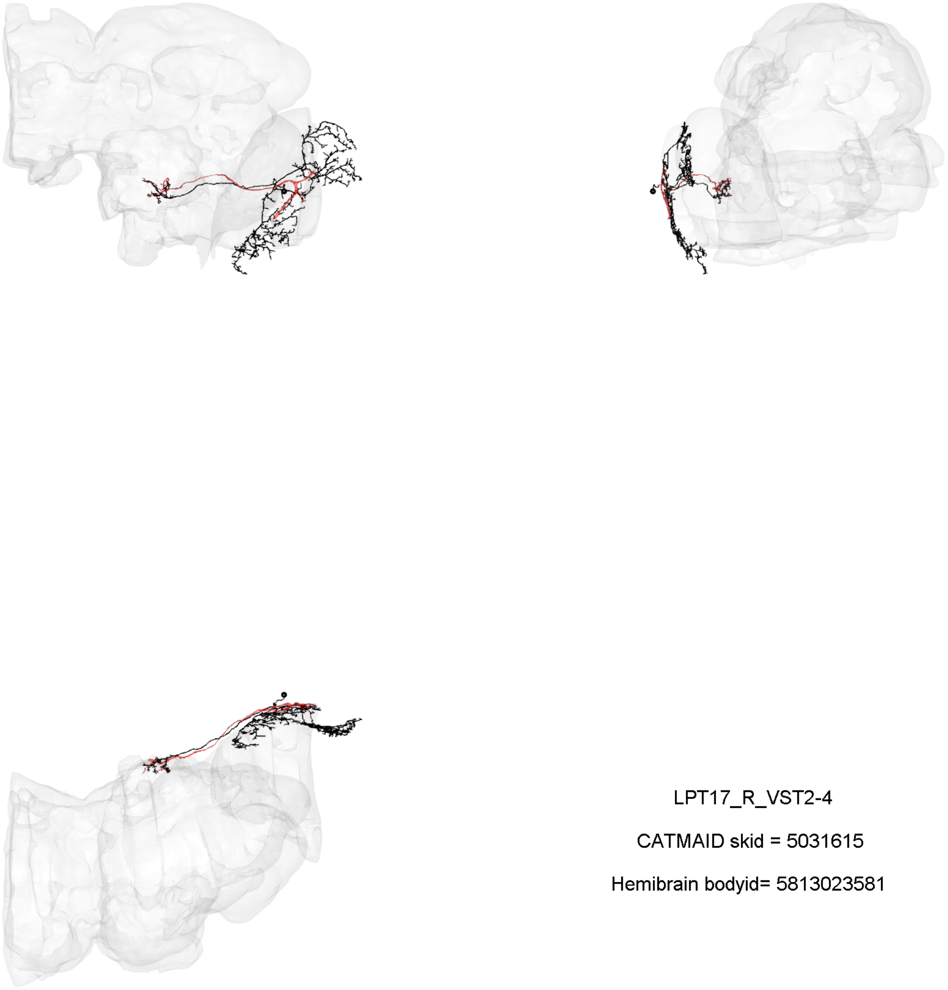

LPT17_R_VST2-4

CATMAID skid = 5031615

Hemibrain bodyid= 5813023581

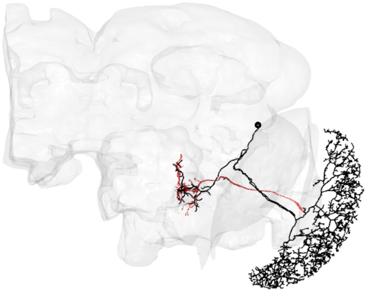

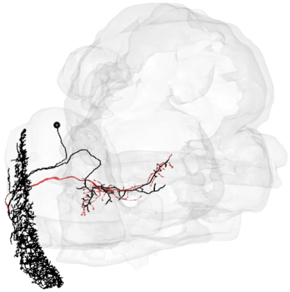

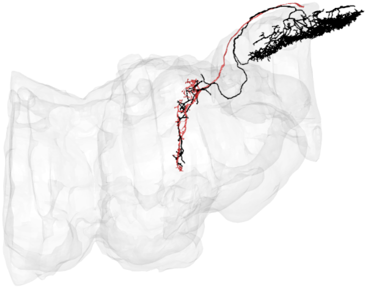

LPT21_R

CATMAID skid = 1110765

Hemibrain bodyid= 1850310331

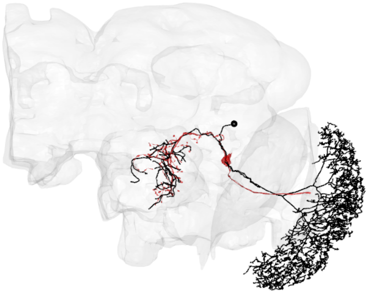

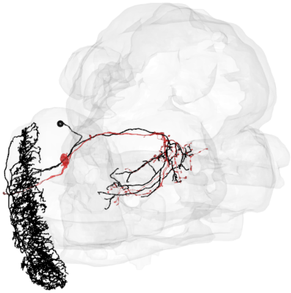

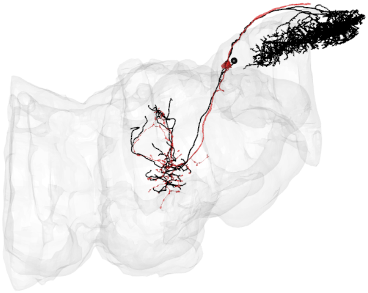

LPT22_R

CATMAID skid = 3510999

Hemibrain bodyid= 1501708149

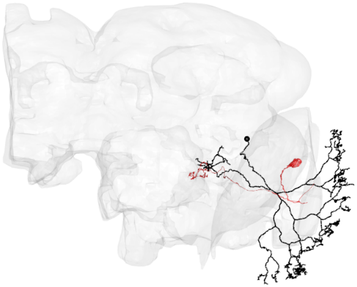

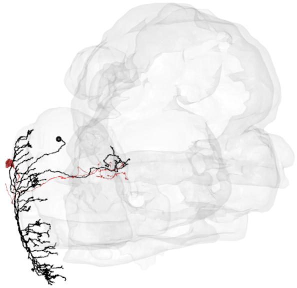

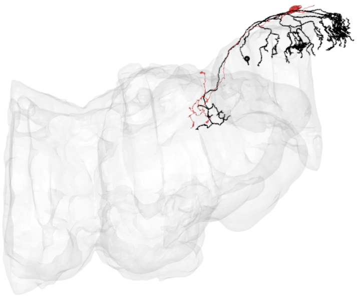

LPT23_R-1

CATMAID skid = 4135042

Hemibrain bodyid= 1842090544

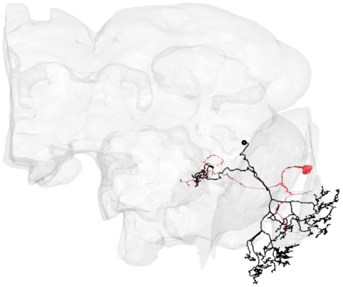

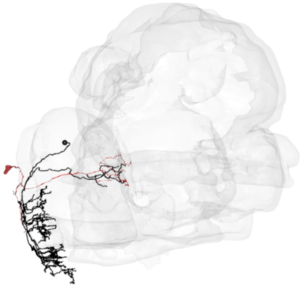

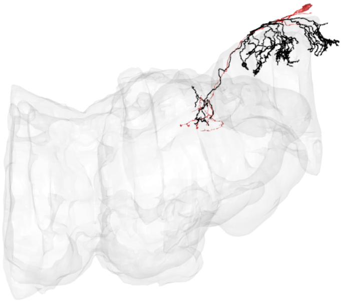

LPT23_R-2

CATMAID skid = 4504548

Hemibrain bodyid= 1751364895

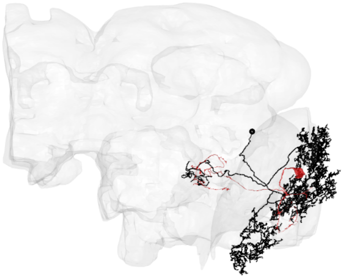

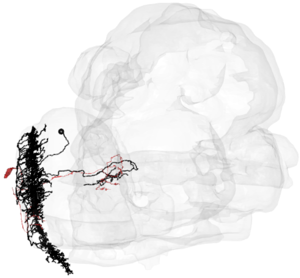

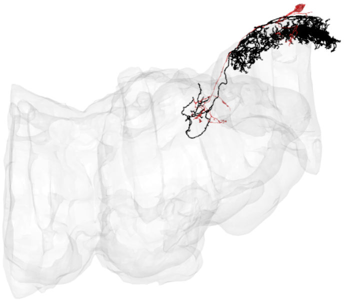

LPT23_R-3

CATMAID skid = 4504557

Hemibrain bodyid= 1751028010

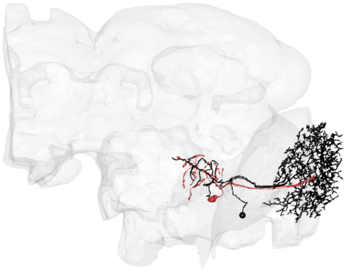

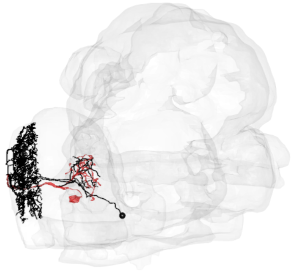

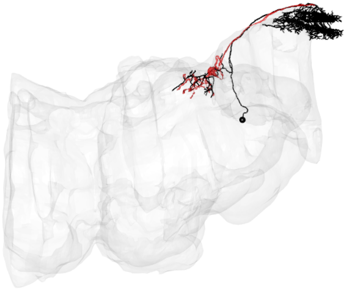

LPT26_R

CATMAID skid = 1107045

Hemibrain bodyid= 1747289181

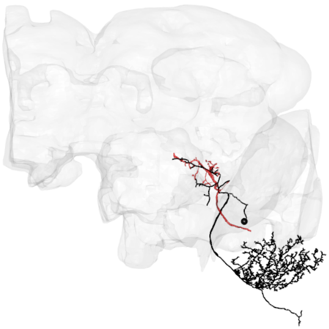

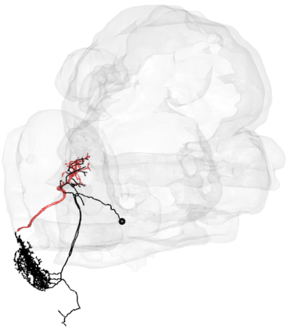

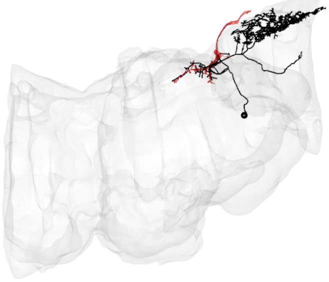

LPT27_R

CATMAID skid = 3514698

Hemibrain bodyid= 1747608690

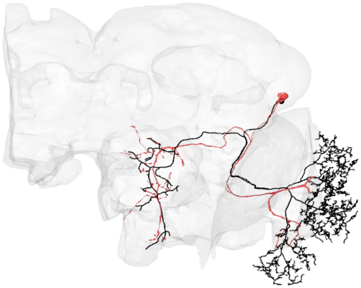

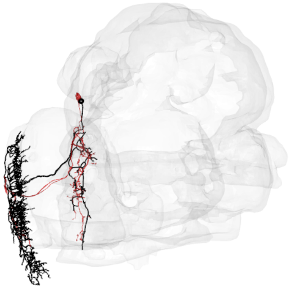

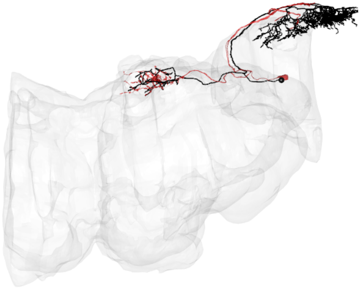

LPT28_R

CATMAID skid = 1071860

Hemibrain bodyid= 1407904797

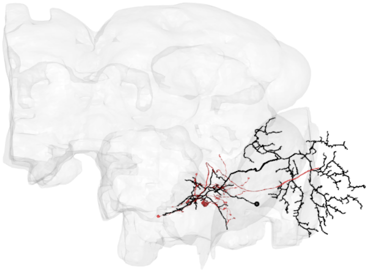

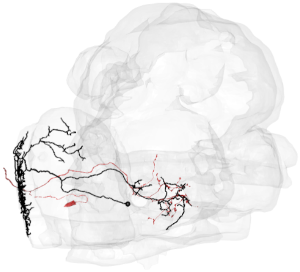

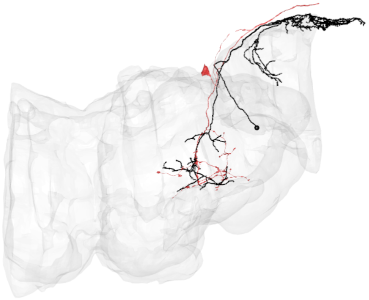

LPT29_R

CATMAID skid = 7760435

Hemibrain bodyid= 5813087982

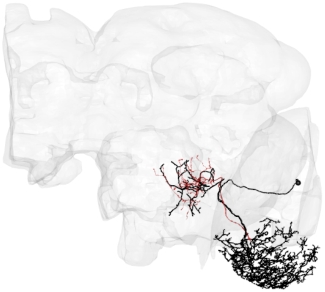

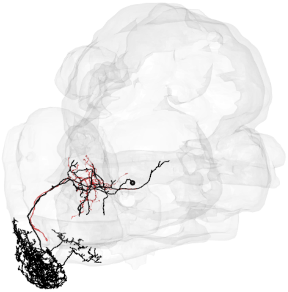

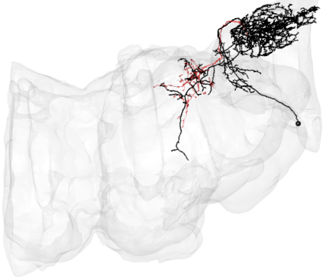

LPT30_R

CATMAID skid = 7674107

Hemibrain bodyid= 1777301335

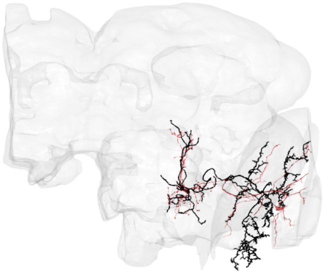

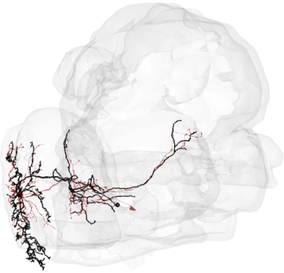

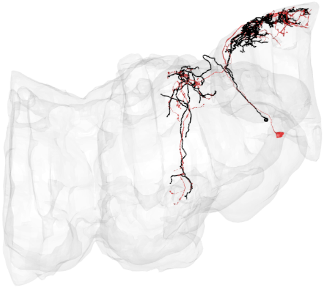

LPT31_R-1

CATMAID skid = 7690273

Hemibrain bodyid= 1622471963

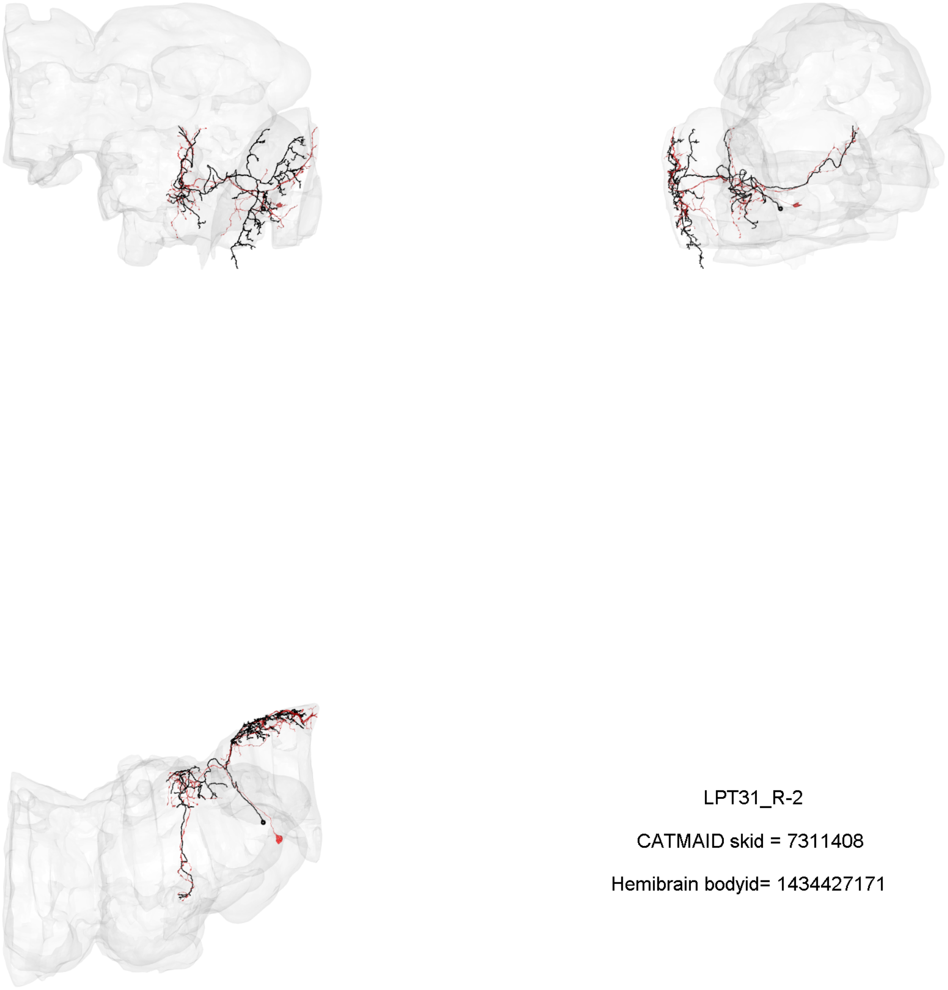

LPT31_R-2

CATMAID skid = 7311408

Hemibrain bodyid= 1434427171

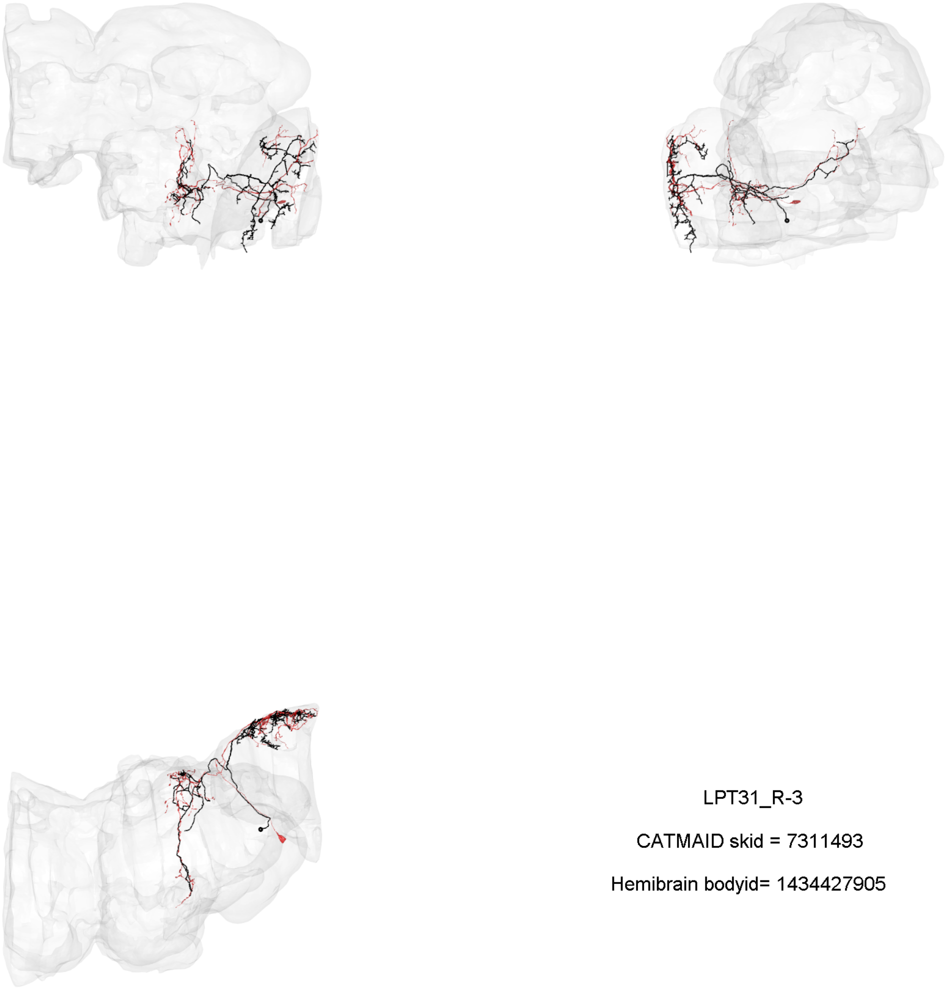

LPT31_R-3

CATMAID skid = 7311493

Hemibrain bodyid= 1434427905

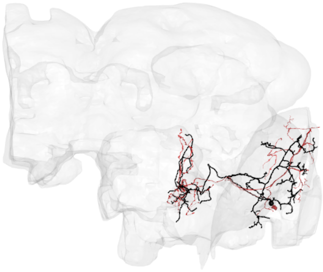

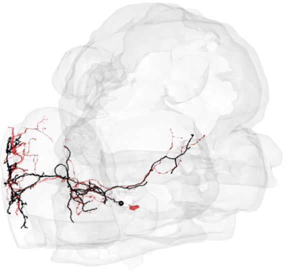

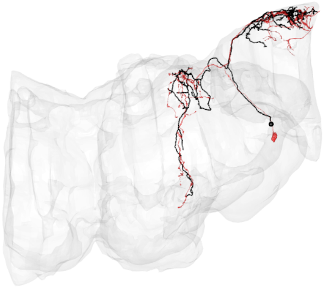

LPT31_R-4

CATMAID skid = 7694305

Hemibrain bodyid= 1344082025

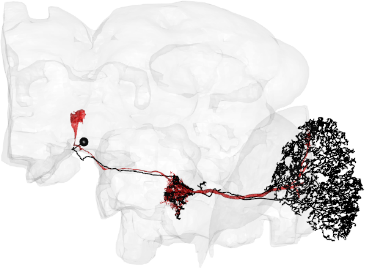

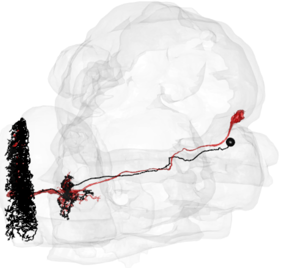

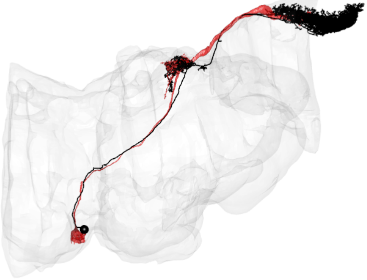

LPT35_R_dCH

CATMAID skid = 1077174

Hemibrain bodyid= 1466485353

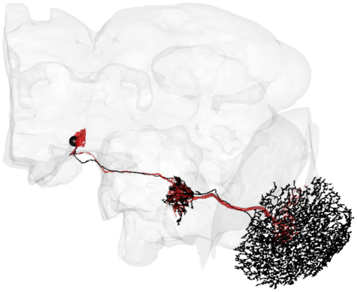
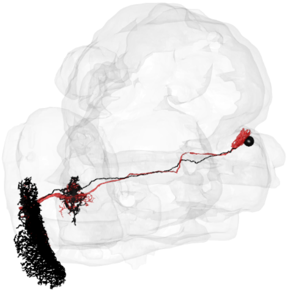
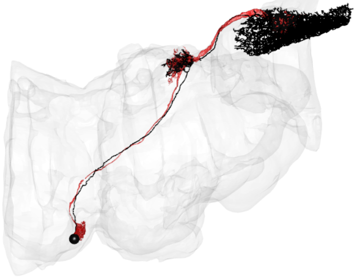

LPT36_R_vCH

CATMAID skid = 1078535

Hemibrain bodyid= 5813024201

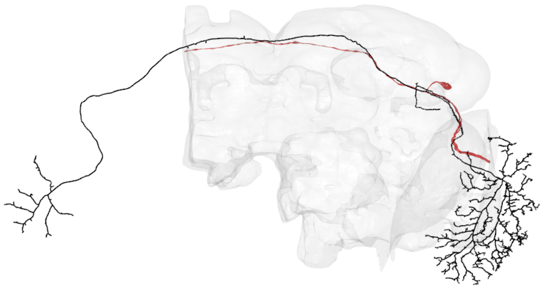

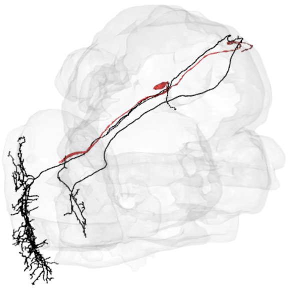

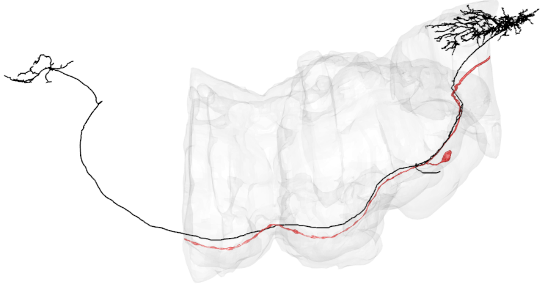

LPT37_R_H1

CATMAID skid = 1121730

Hemibrain bodyid= 676832896

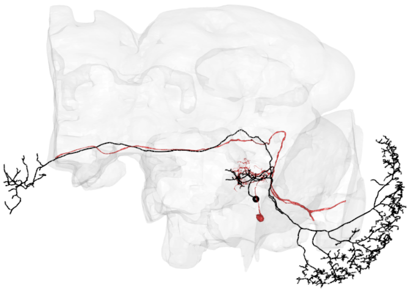

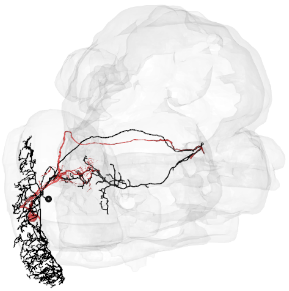

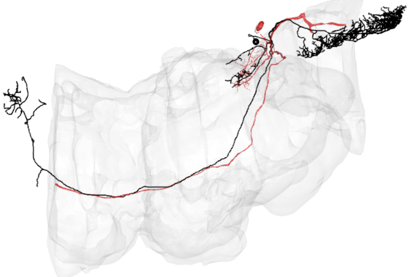

LPT38_R_Nod1-1

CATMAID skid = 1054753

Hemibrain bodyid= 5812996970

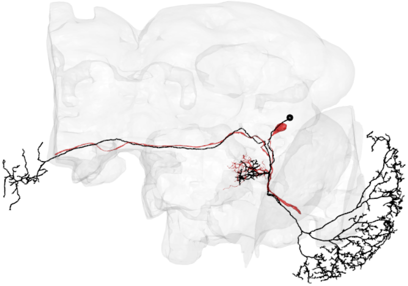

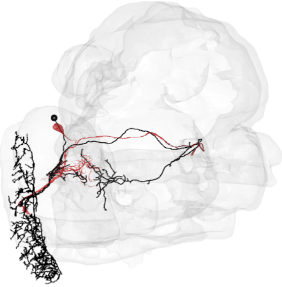

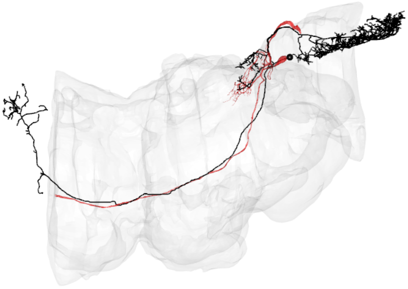

LPT38_R_Nod1-2

CATMAID skid = 3546483

Hemibrain bodyid= 5812993603

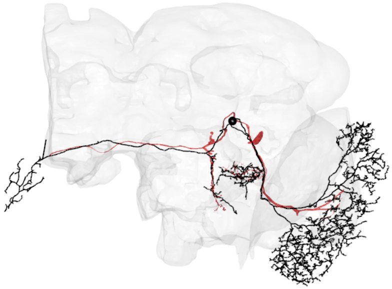

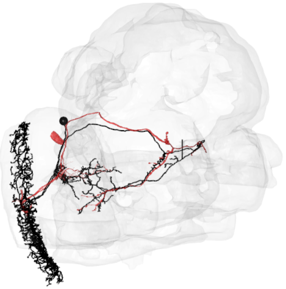

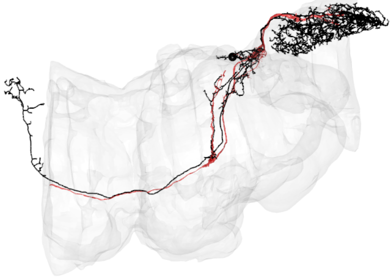

LPT40_R_Nod2

CATMAID skid = 902072

Hemibrain bodyid= 1315529069

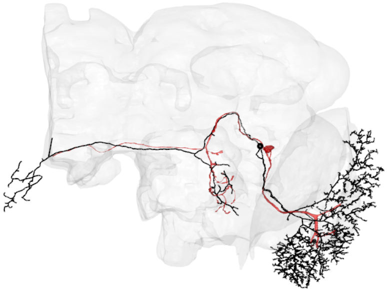

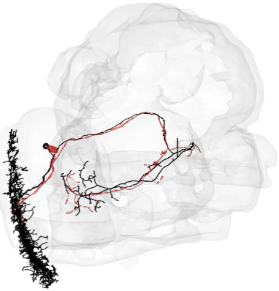

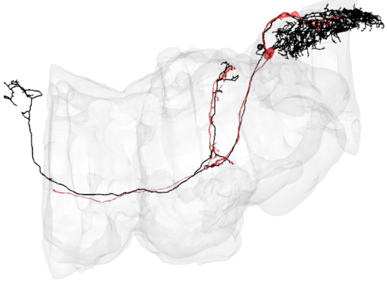

LPT41_R_Nod3

CATMAID skid = 1058996

Hemibrain bodyid= 1352706891

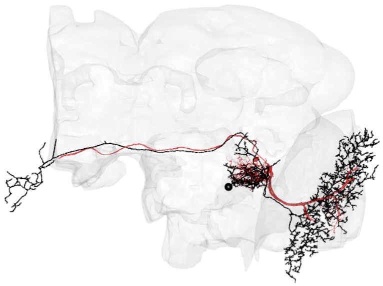

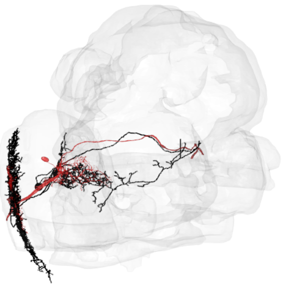

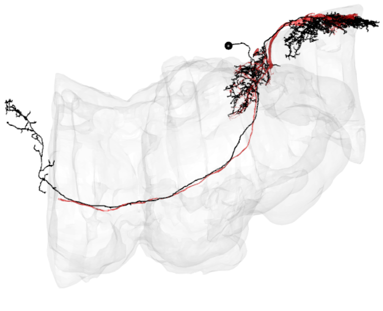

LPT42_R_Nod4

CATMAID skid = 1106958

Hemibrain bodyid= 1566524544

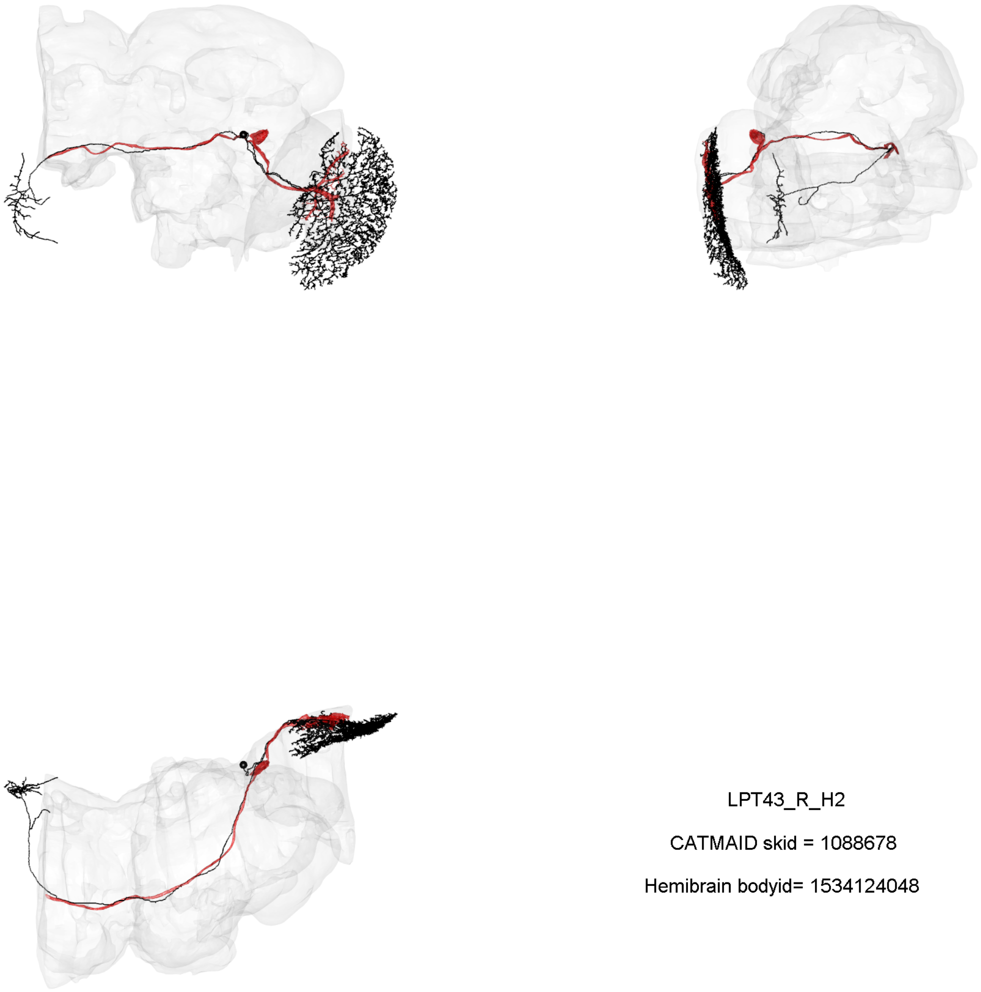

LPT43_R_H2

CATMAID skid = 1088678

Hemibrain bodyid= 1534124048

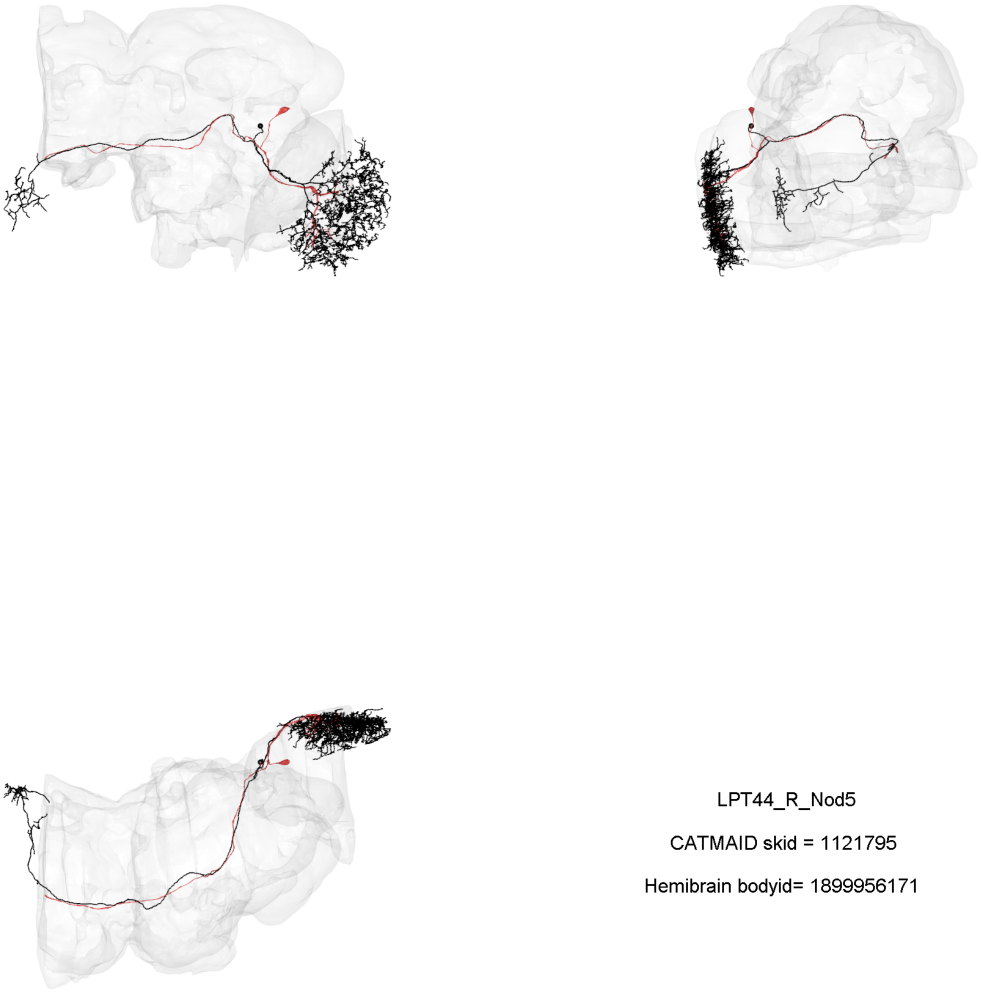

LPT44_R_Nod5

CATMAID skid = 1121795

Hemibrain bodyid= 1899956171

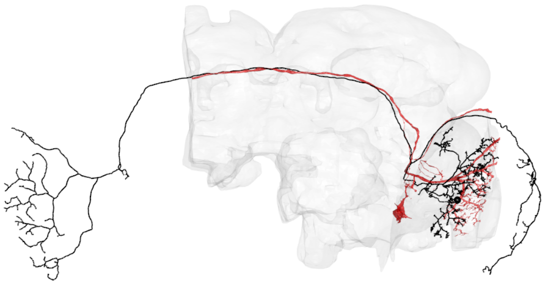

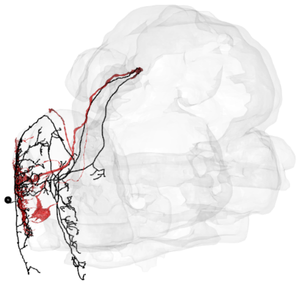

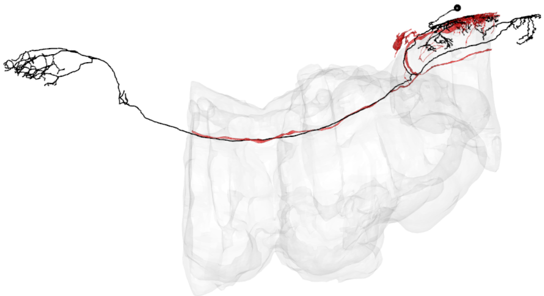

LPT45_R_dCal1

CATMAID skid = 7204844

Hemibrain bodyid= 943813788

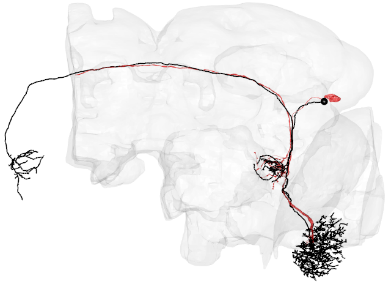

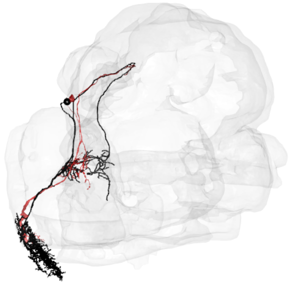

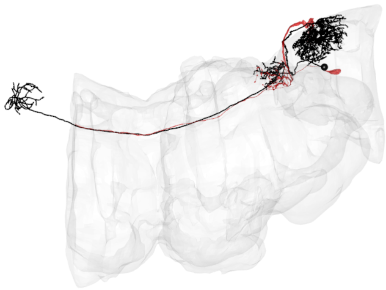

LPT46_R_vCal1

CATMAID skid = 7510076

Hemibrain bodyid= 1005174931

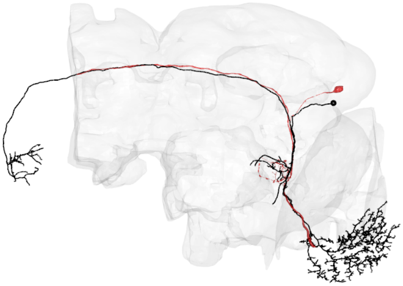

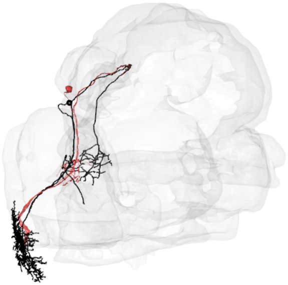

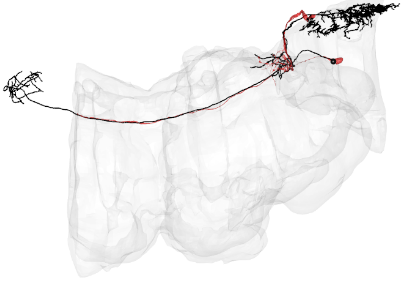

LPT47_R_vCal2

CATMAID skid = 3529071

Hemibrain bodyid= 1005174975

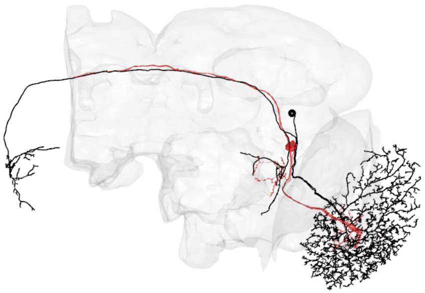

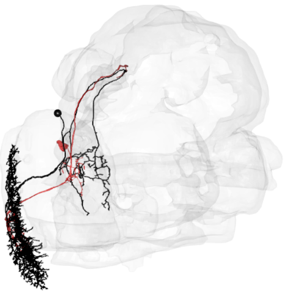

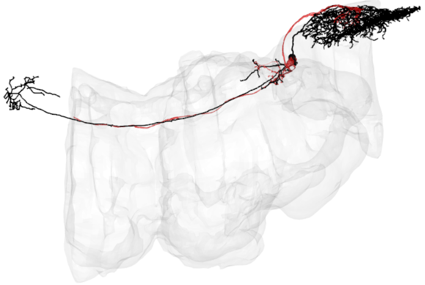

LPT48_R_vCal3

CATMAID skid = 1056097

Hemibrain bodyid= 974502819

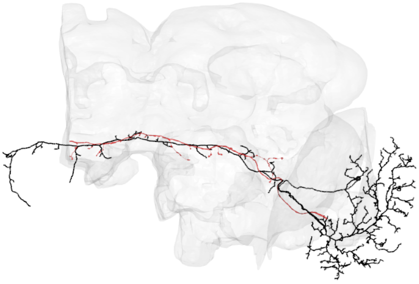

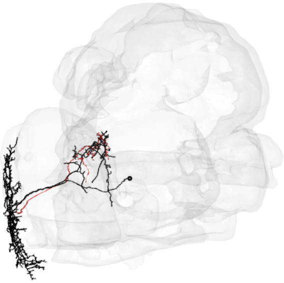

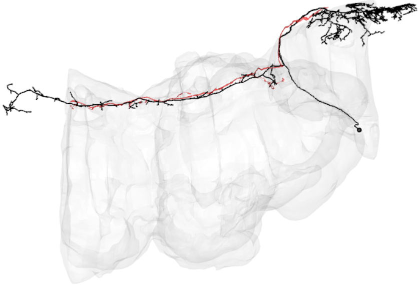

LPT49_R

CATMAID skid = 1110693

Hemibrain bodyid= 1654969939

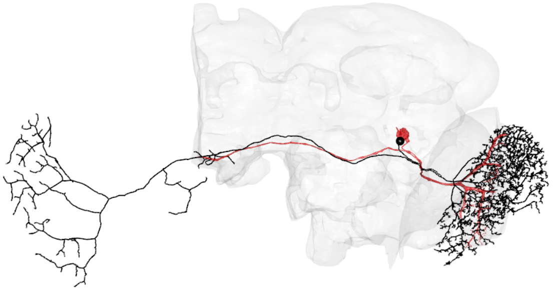

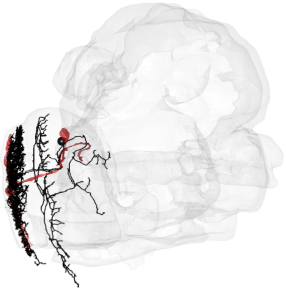

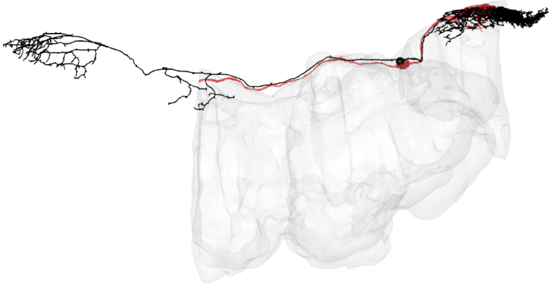

LPT50_R

CATMAID skid = 905761

Hemibrain bodyid= 1496497366

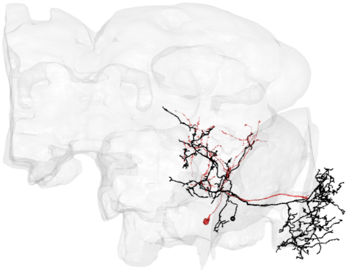

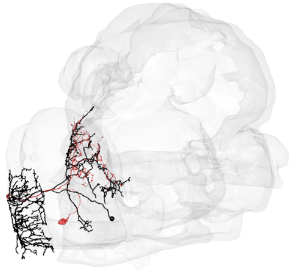

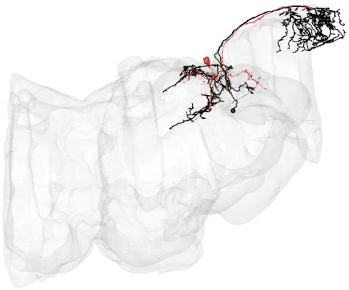

LPT51_R

CATMAID skid = 4224711

Hemibrain bodyid= 1313496323

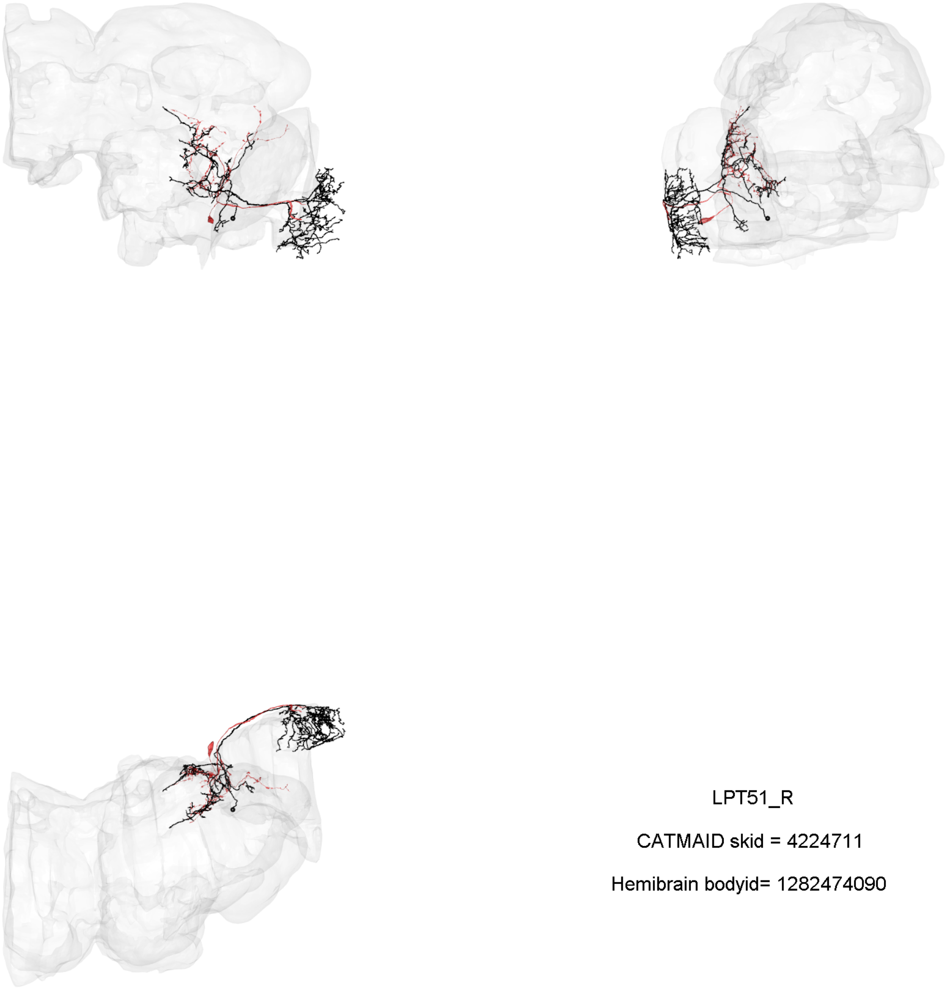

LPT51_R

CATMAID skid = 4224711

Hemibrain bodyid= 1282474090

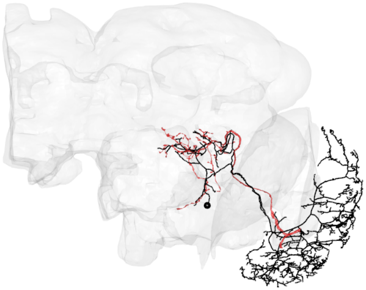

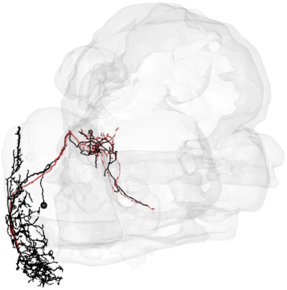

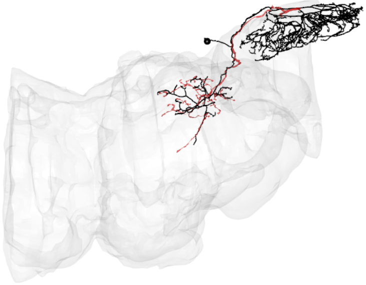

LPT52_R

CATMAID skid = 1107296

Hemibrain bodyid= 1469291436

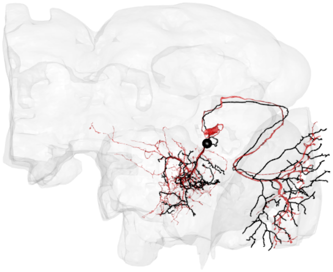

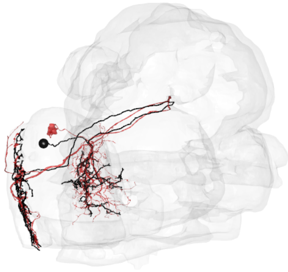

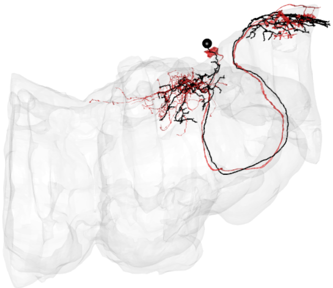

LPT53_R

CATMAID skid = 1111992

Hemibrain bodyid= 1231610379

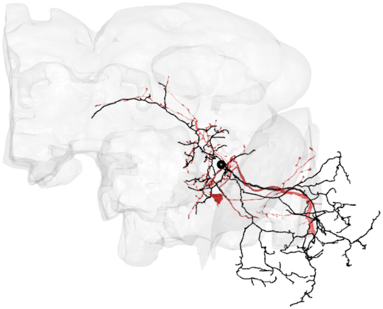

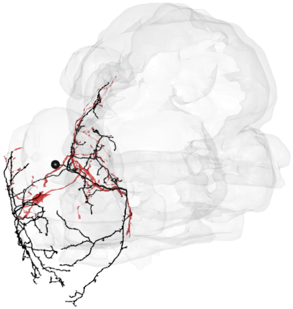

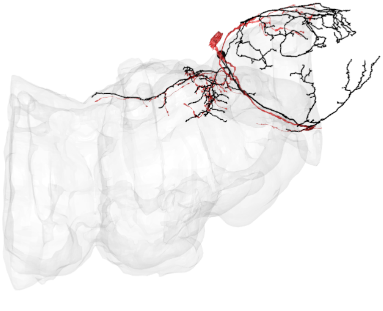

LPT54_R

CATMAID skid = 4235388

Hemibrain bodyid= 1129033939

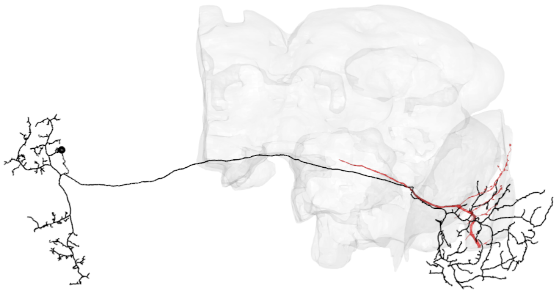

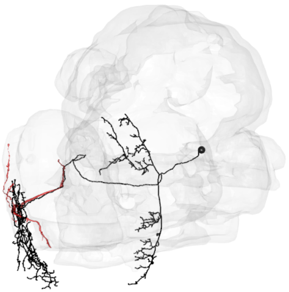

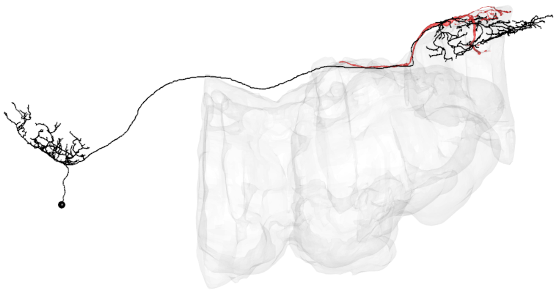

LPT55_R_MeLp2

CATMAID skid = 1061368

Hemibrain bodyid= 1558226341

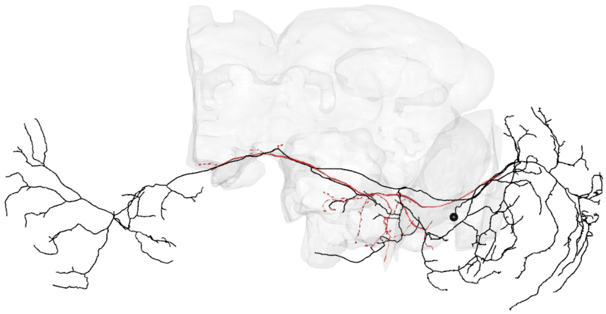

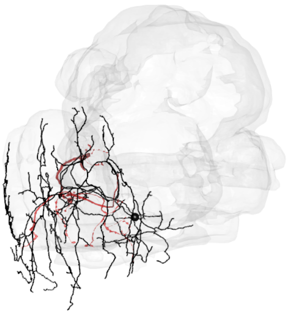

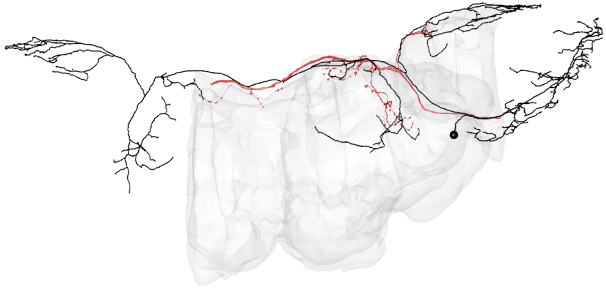

LPT56_R_MeLp1

CATMAID skid = 3509520

Hemibrain bodyid= 5813068976

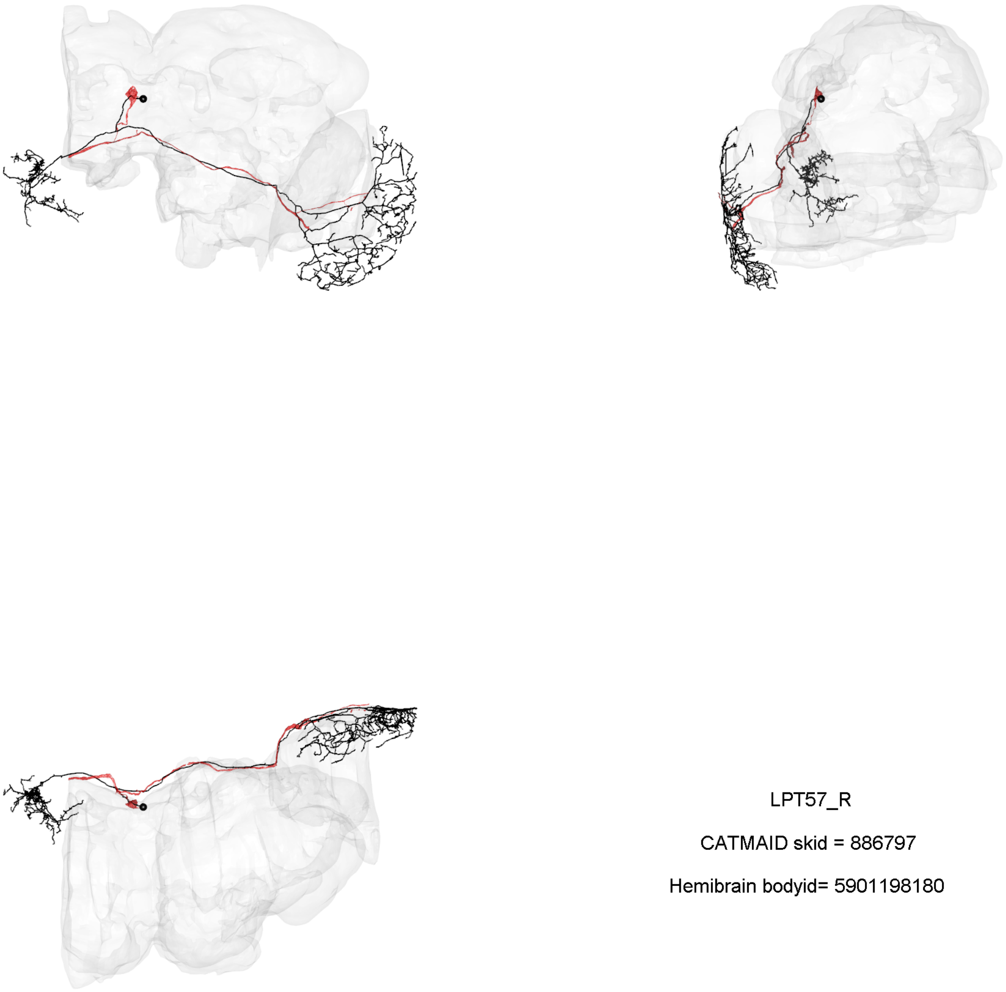

LPT57_R

CATMAID skid = 886797

Hemibrain bodyid= 5901198180

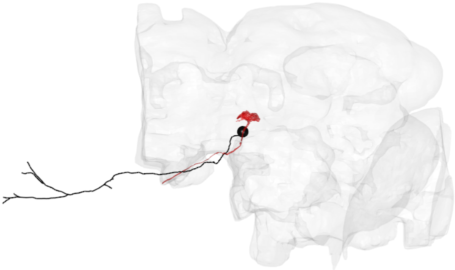

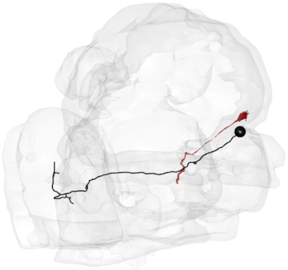

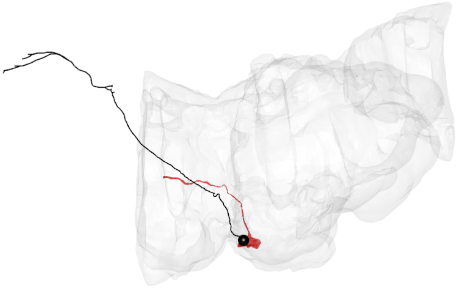

LPT35_L_dCH

CATMAID skid = 6243409

Hemibrain bodyid= 1545158404

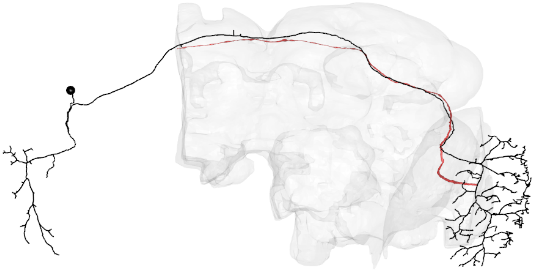

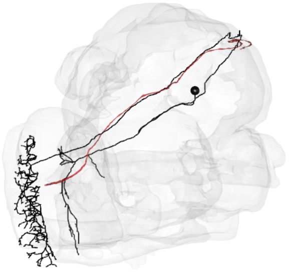

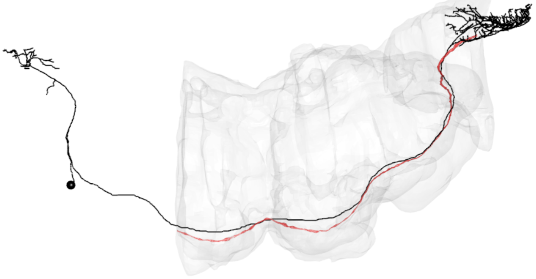

LPT37_L_H1

CATMAID skid = 1121335

Hemibrain bodyid= 1167783603

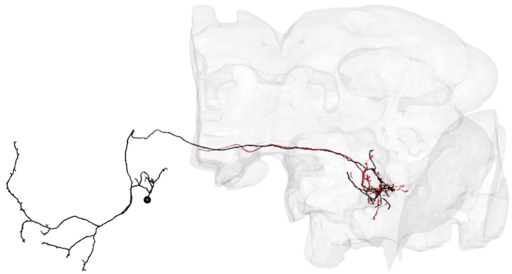

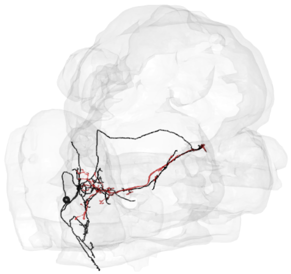

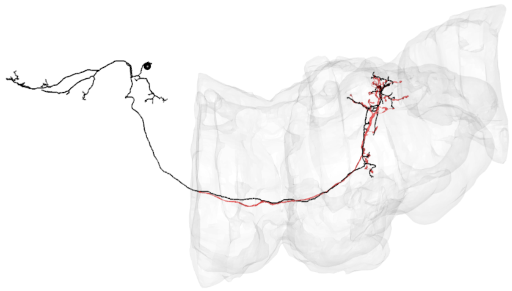

LPT38_L_Nod1-1

CATMAID skid = 6439204

Hemibrain bodyid= 1789306586

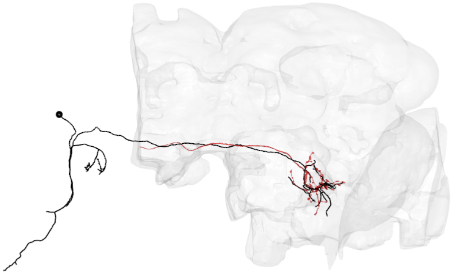

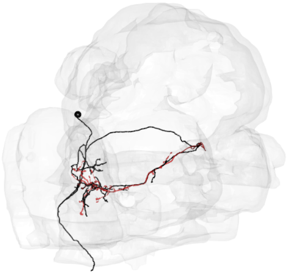

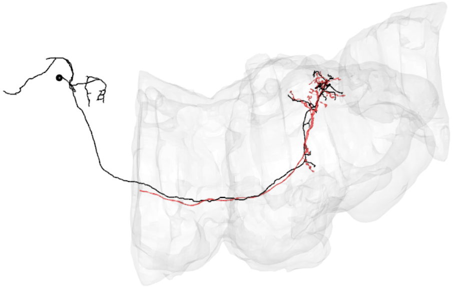

LPT38_L_Nod1-2

CATMAID skid = 6437945

Hemibrain bodyid= 1758621675

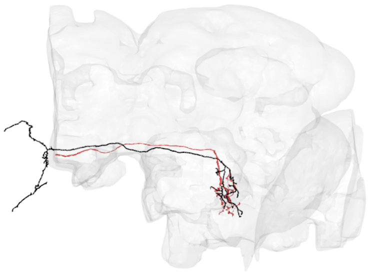

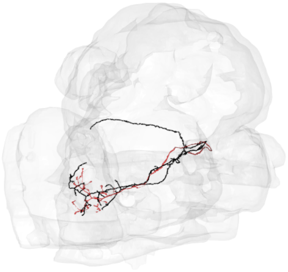

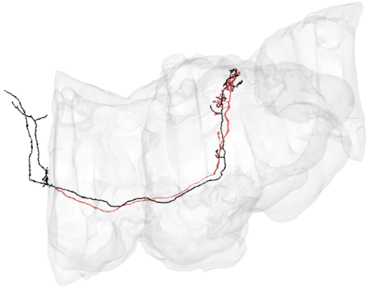

LPT40_L_Nod2

CATMAID skid = 16615842

Hemibrain bodyid= 1871778911

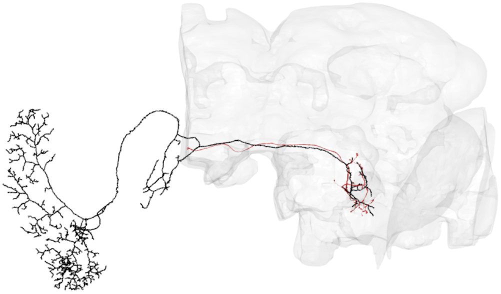

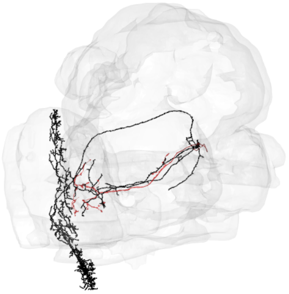

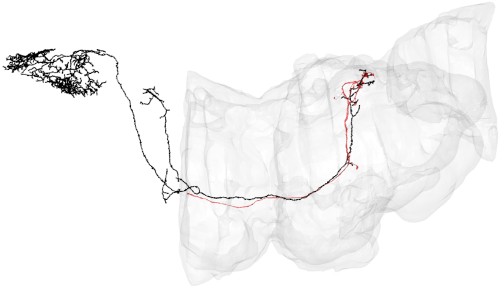

LPT41_L_Nod3

CATMAID skid = 15060018

Hemibrain bodyid= 1758617327

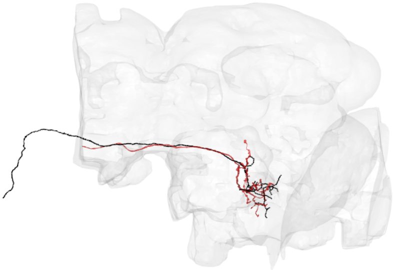

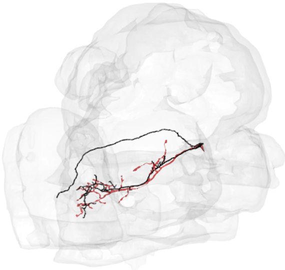

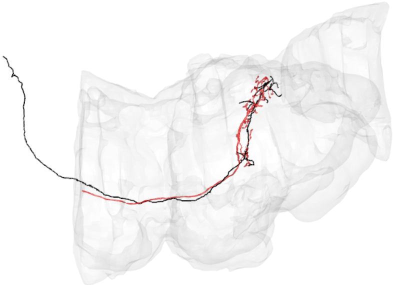

LPT42_L_Nod4

CATMAID skid = 16615419

Hemibrain bodyid= 1725837767

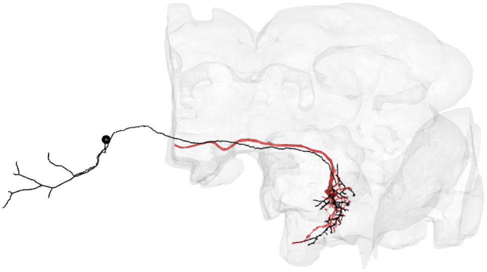

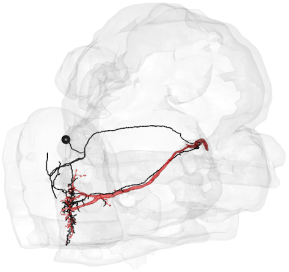

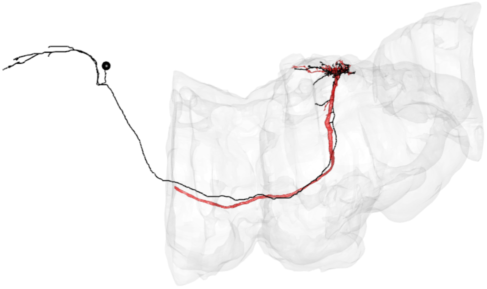

LPT43_L_H2

CATMAID skid = 5232902

Hemibrain bodyid= 5813078454

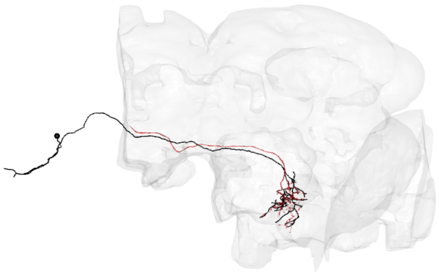

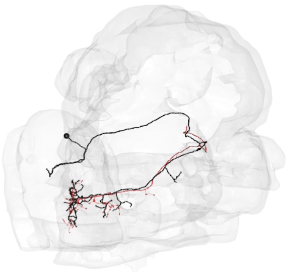

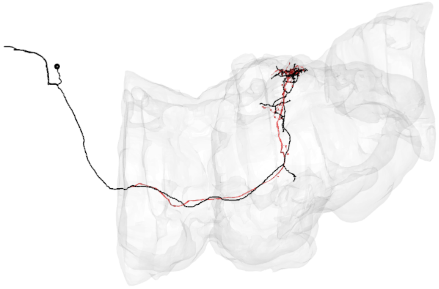

LPT44_L_Nod5

CATMAID skid = 7231304

Hemibrain bodyid= 5812994338

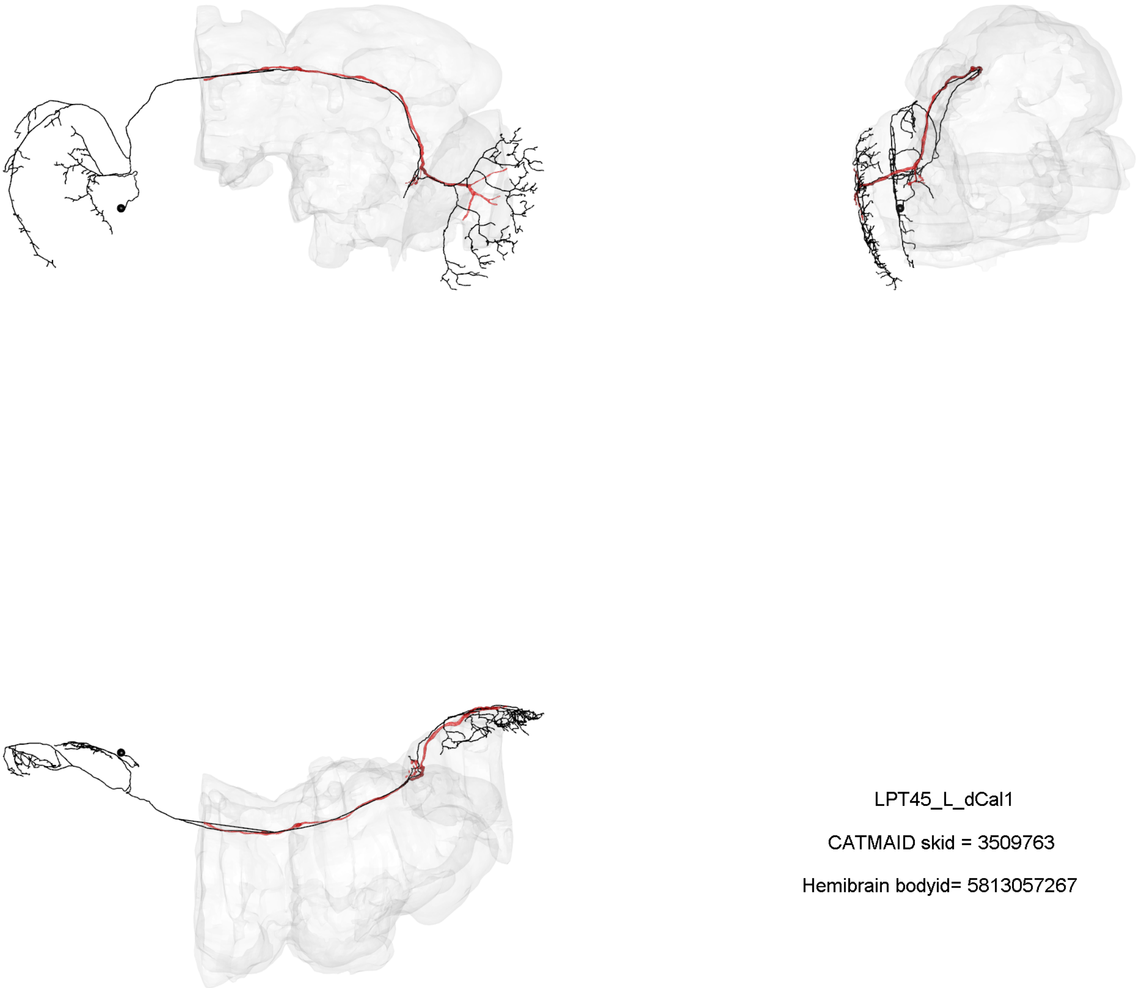

LPT45_L_dCal1

CATMAID skid = 3509763

Hemibrain bodyid= 5813057267

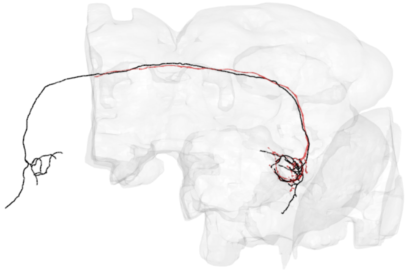

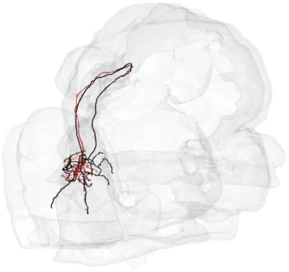

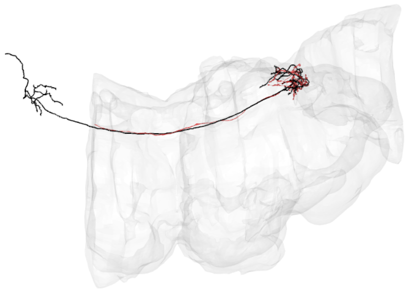

LPT46_L_vCal1

CATMAID skid = 8747266

Hemibrain bodyid= 943468720

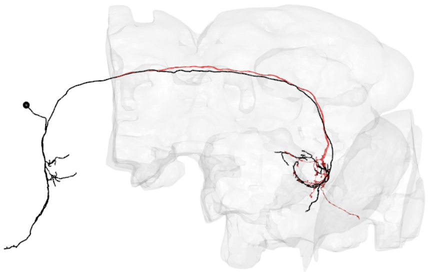

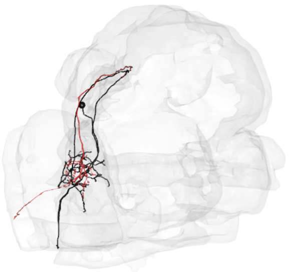

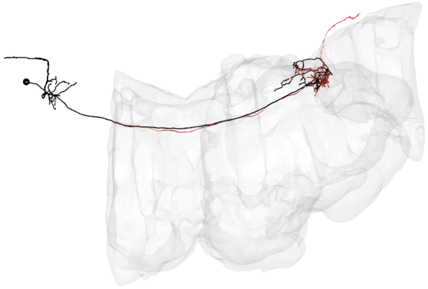

LPT47_L_vCal2

CATMAID skid = 7449616

Hemibrain bodyid= 943472755

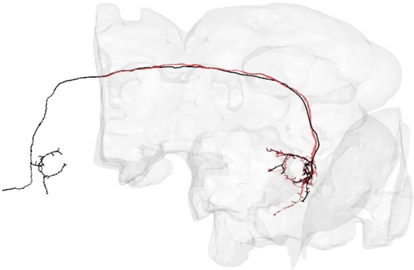

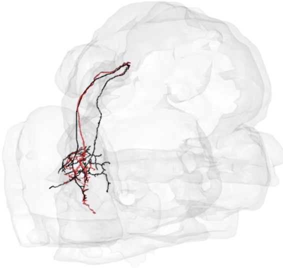

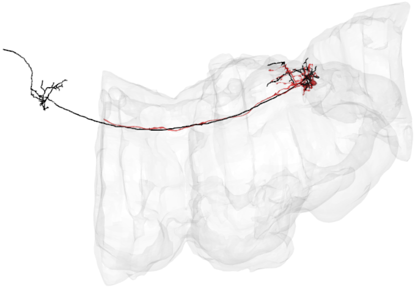

LPT48_L_vCal3

CATMAID skid = 11230125

Hemibrain bodyid= 943472763

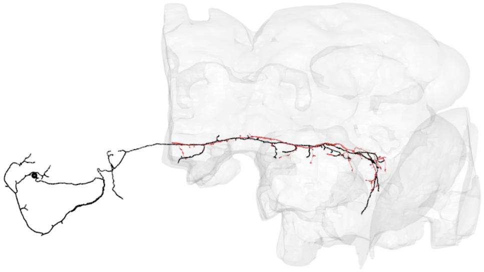

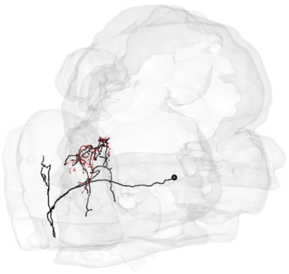

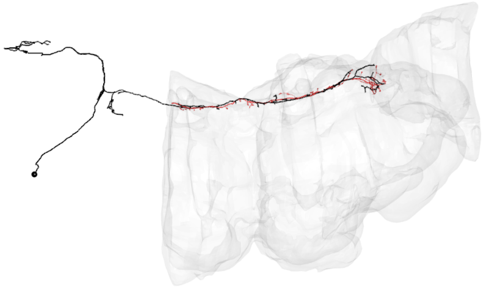

LPT49_L

CATMAID skid = 17072553

Hemibrain bodyid= 1685896788

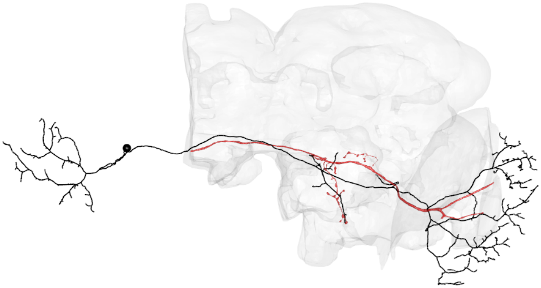

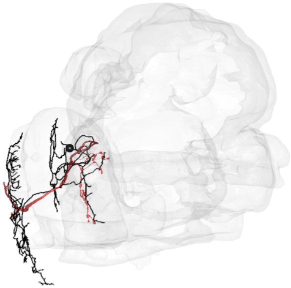

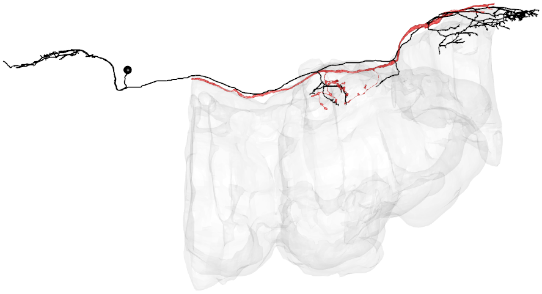

LPT50_L

CATMAID skid = 3503997

Hemibrain bodyid= 5813049974

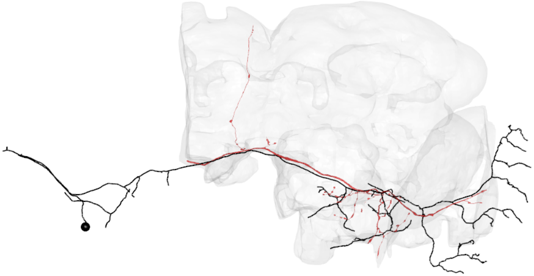

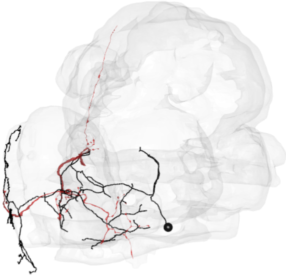

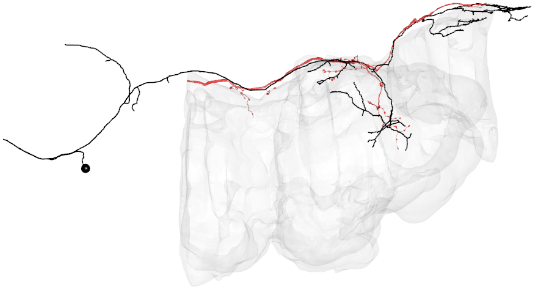

LPT56_L_MeLo1

CATMAID skid = 4224594

Hemibrain bodyid= 5813063227

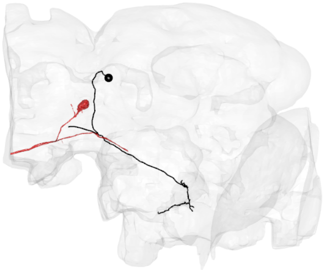

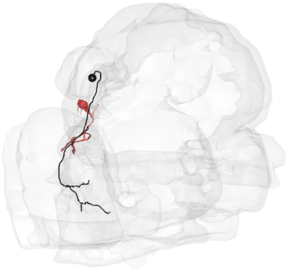

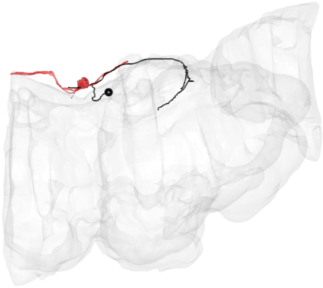

LPT57_L

CATMAID skid = 17059178

Hemibrain bodyid= 5813045086
