## Supplemental File 5 for "A comprehensive neuroanatomical survey of the *Drosophila* Lobula Plate Tangential Neurons with predictions for their optic flow sensitivity"

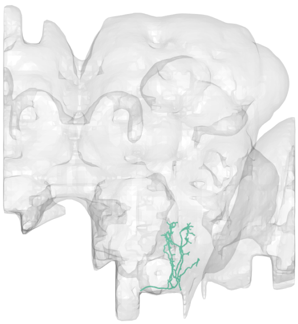

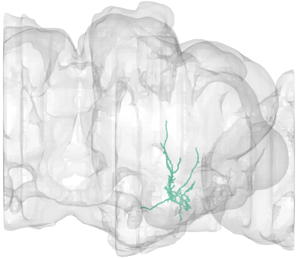

#1
bodyid =
5813132521

FOV = 19%
overlap of input LPTs' FOV = 10%
optimal rotation axis = [−80°,−180°]
$\alpha_{R0} = 8°$
optimal translation axis = [4°,102°]
$\alpha_{T0} = 10°$
RTSA = −2°

input LPT neuons:
LPT21_R
LPT38_L_Nod1

PMPM

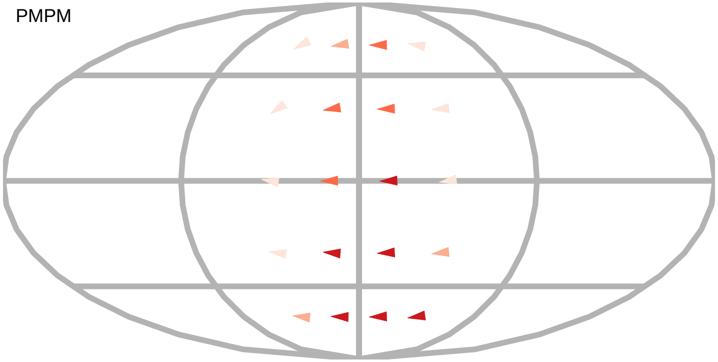

$\alpha_R$

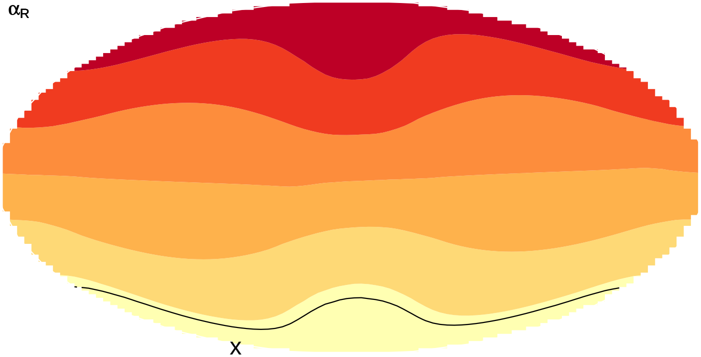


layer innervation

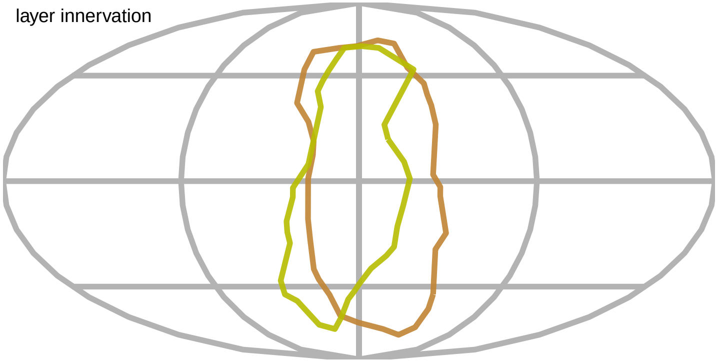

$\alpha_T$

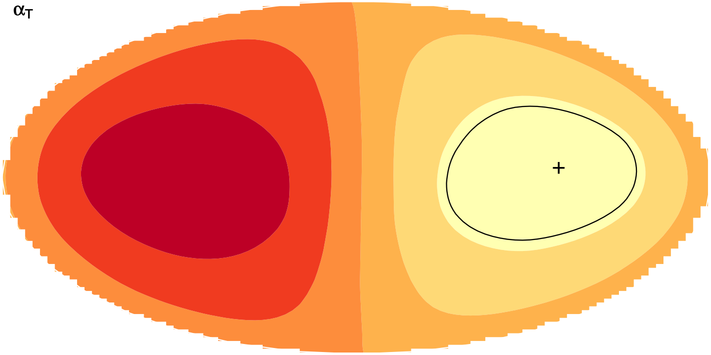

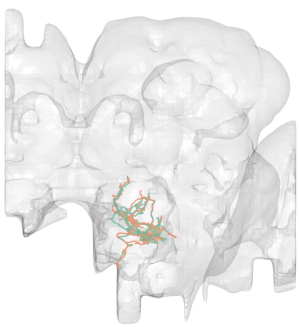

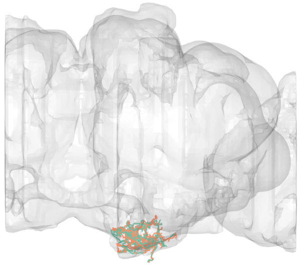

#2
bodyid =
1775638620
5901195929

FOV = 24%
overlap of input LPTs' FOV = 11%
optimal rotation axis = [−10°,97°]
$\alpha_{R0} = 22°$
optimal translation axis = [70°,164°]
$\alpha_{T0} = 28°$
RTSA = −6°

input LPT neuons:
LPT13_R_VSm
LPT15_R_VST1

PMPM

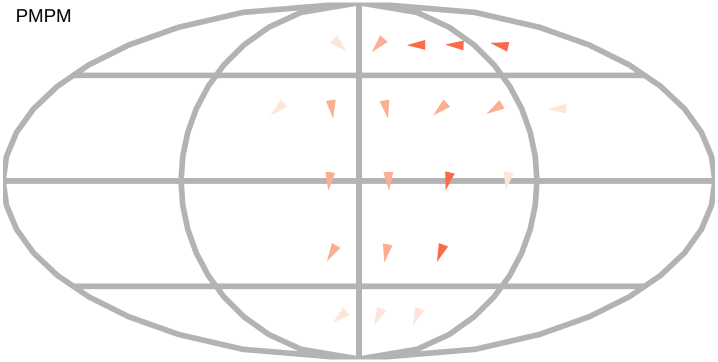

$\alpha_R$

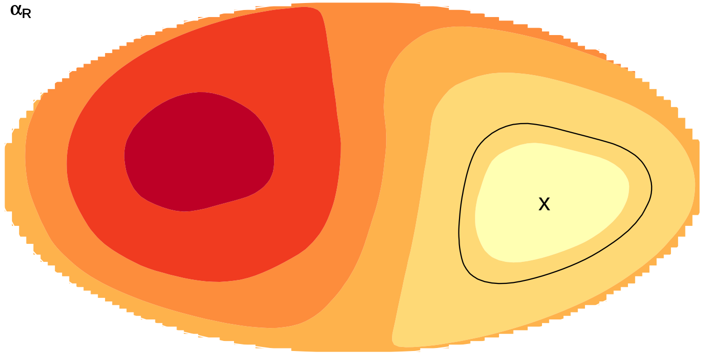


$\alpha_T$

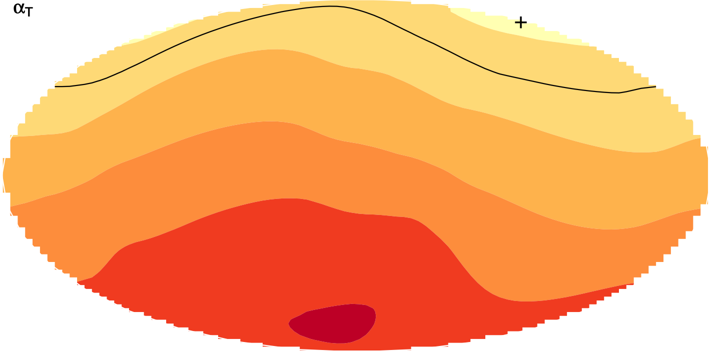


layer innervation

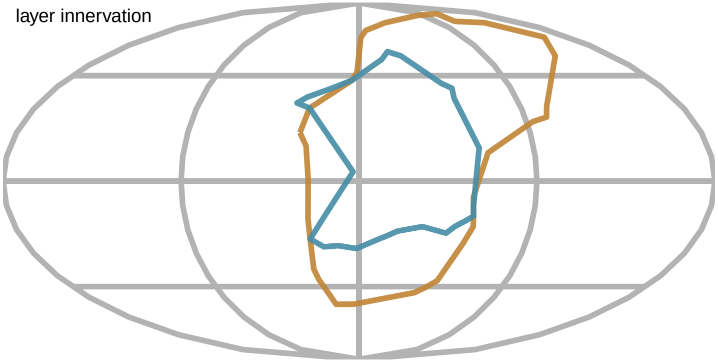

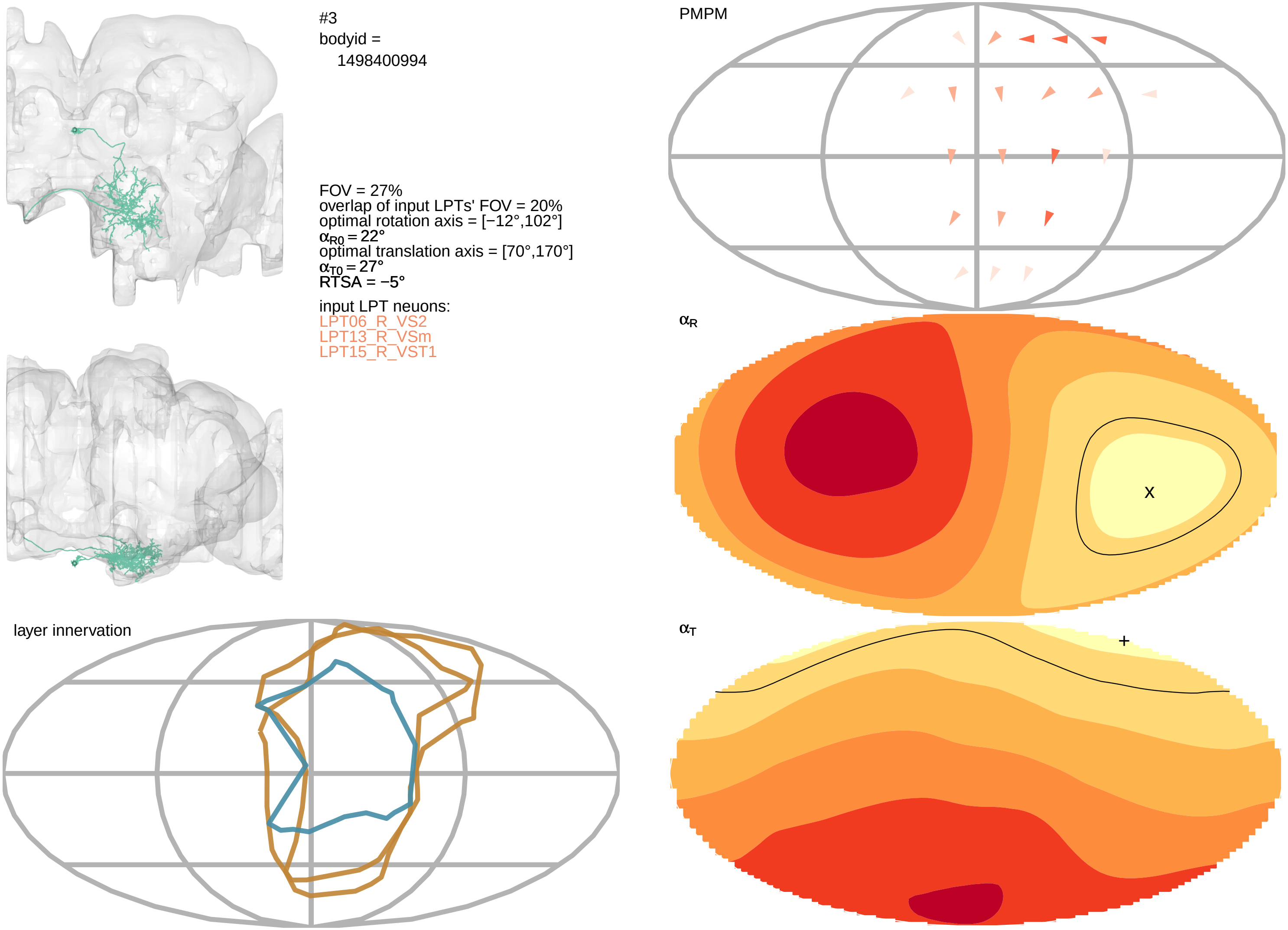

#3
bodyid =
1498400994
FOV = 27%
overlap of input LPTs' FOV = 20%
optimal rotation axis = [−12°,102°]
$\alpha_{R0} = 22°$
optimal translation axis = [70°,170°]
$\alpha_{T0} = 27°$
RTSA = −5°
input LPT neuons:
LPT06_R_VS2
LPT13_R_VSm
LPT15_R_VST1
layer innervation
PMPM
$\alpha_R$
x
$\alpha_T$
+

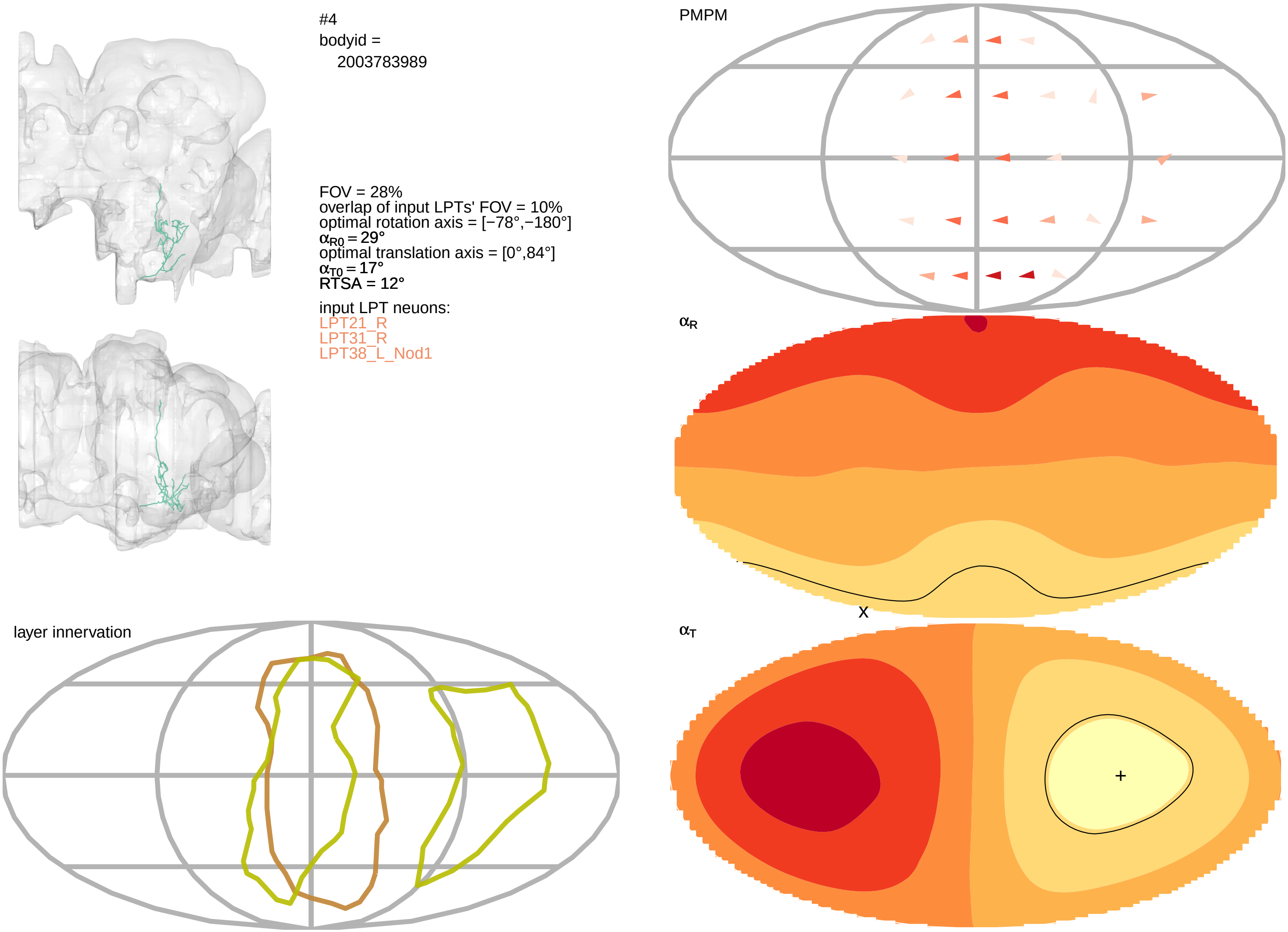
#4
bodyid =
2003783989
FOV = 28%
overlap of input LPTs' FOV = 10%
optimal rotation axis = [−78°,−180°]
αR0 = 29°
optimal translation axis = [0°,84°]
αT0 = 17°
RTSA = 12°
input LPT neuons:
LPT21_R
LPT31_R
LPT38_L_Nod1
PMPM
αR
X
αT
+
layer innervation

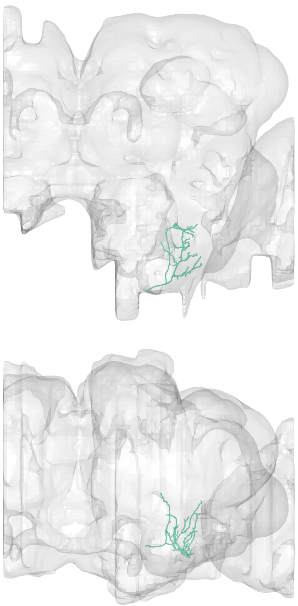

#5
bodyid =
2028658555

FOV = 28%
overlap of input LPTs' FOV = 10%
optimal rotation axis = [−78°,−180°]
$\alpha_{R0} = 29°$
optimal translation axis = [0°,84°]
$\alpha_{T0} = 17°$
RTSA = 12°

input LPT neuons:
LPT21_R
LPT31_R
LPT38_L_Nod1

PMPM

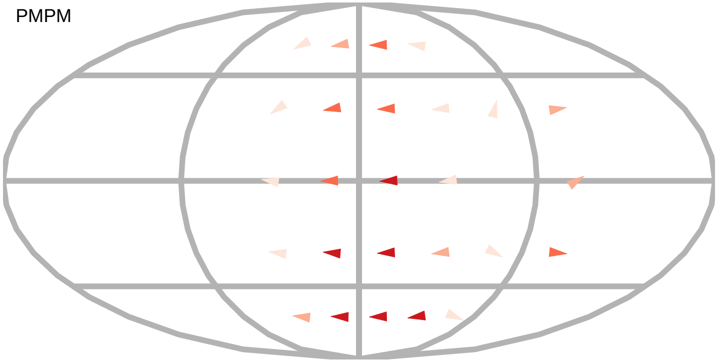

$\alpha_R$

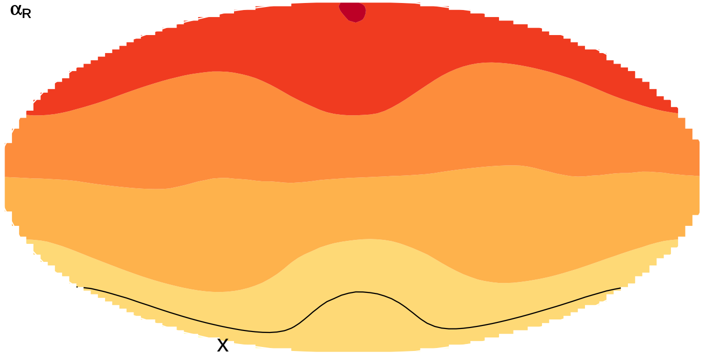


layer innervation

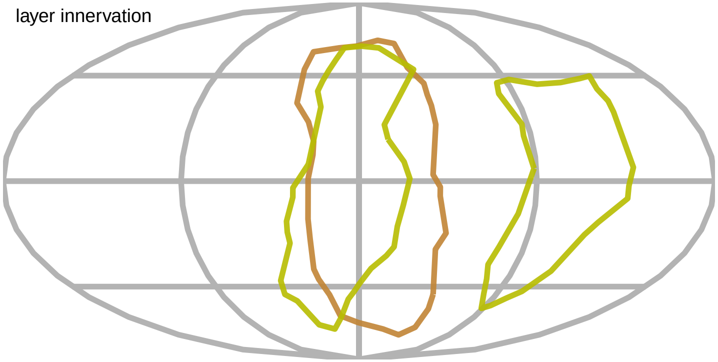

$\alpha_T$

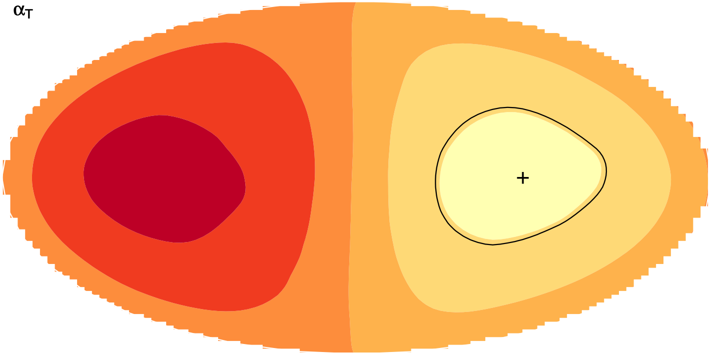

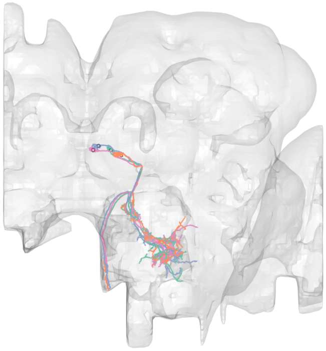

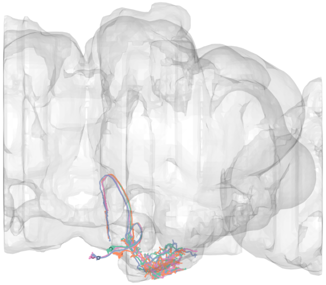

#6
bodyid =
5812993956
1654940174
5813045400
5813054498

FOV = 30%
overlap of input LPTs' FOV = 21%
optimal rotation axis = [38°,−170°]
$\alpha_{R0} = 26°$
optimal translation axis = [38°,−56°]
$\alpha_{T0} = 38°$
RTSA = −12°

input LPT neuons:
LPT04_R_HST
LPT09_R_VS5
LPT17_R_VST2

layer innervation

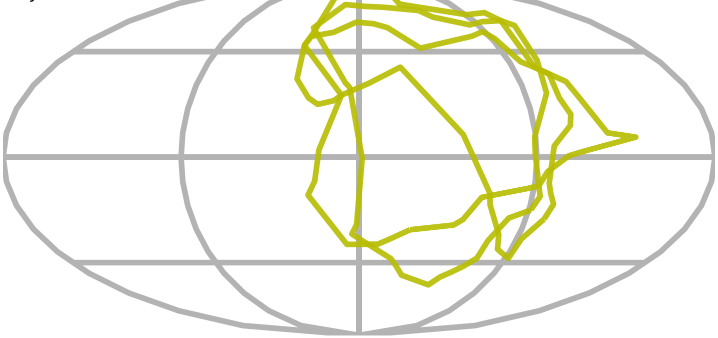

PMPM

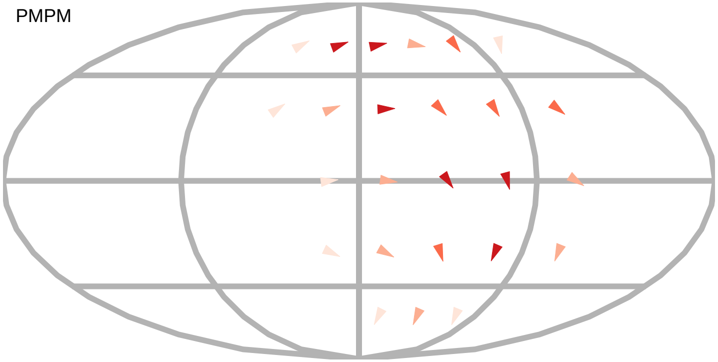

$\alpha_R$

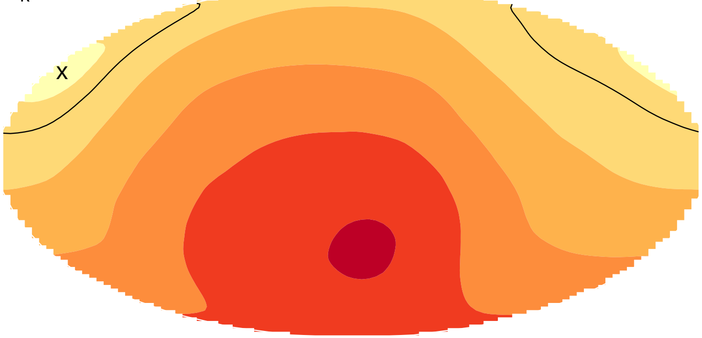


$\alpha_T$

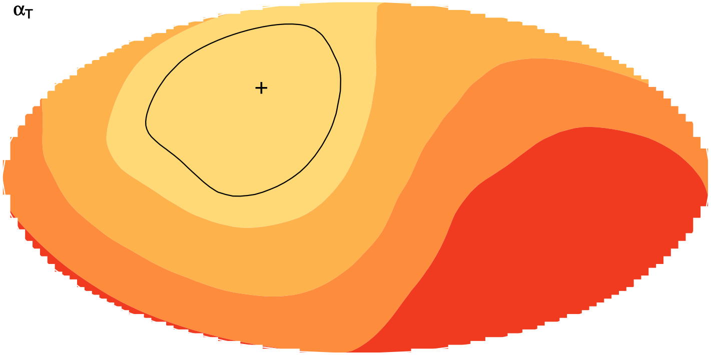

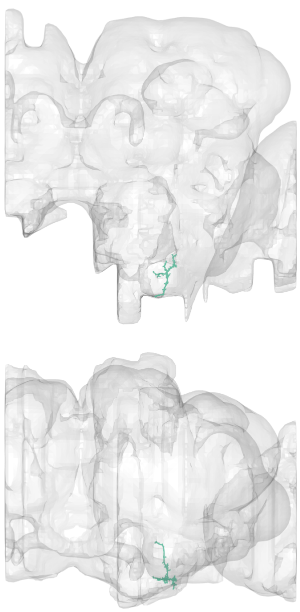

#7
bodyid =
2273535330

FOV = 32%
overlap of input LPTs' FOV = 16%
optimal rotation axis = [68°,−166°]
$\alpha_{R0} = 13°$
optimal translation axis = [−14°,−136°]
$\alpha_{T0} = 19°$
RTSA = −6°

input LPT neuons:
LPT43_L_H2
LPT44_L_Nod5

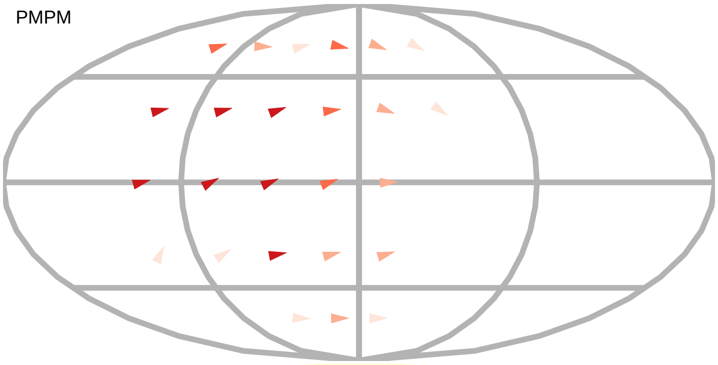


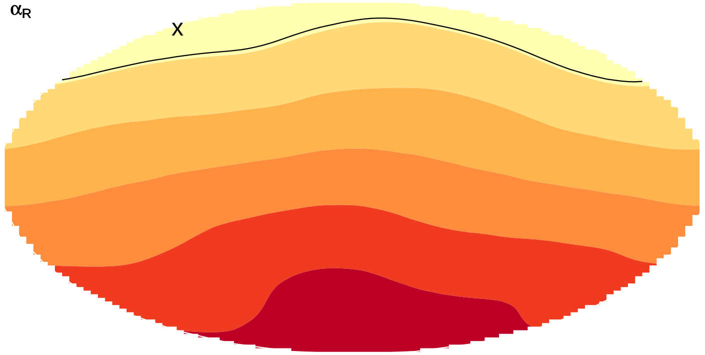


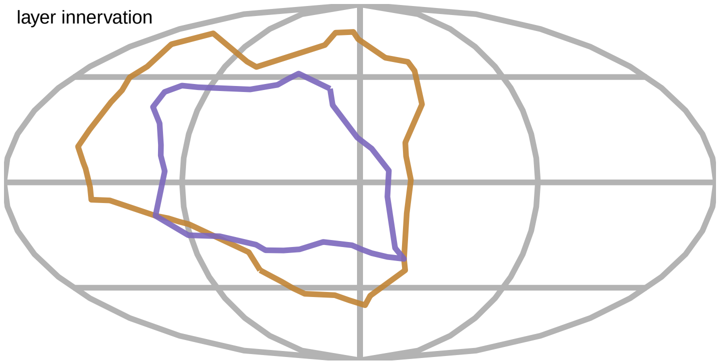


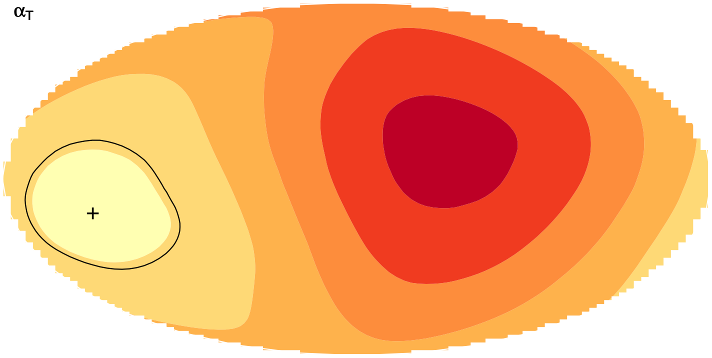

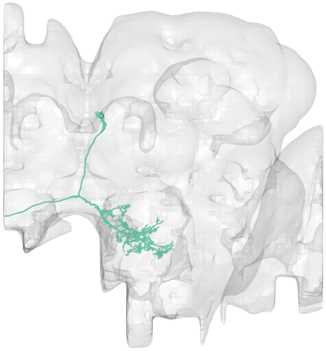

#8
bodyid =
1653295260

FOV = 32%
overlap of input LPTs' FOV = 26%
optimal rotation axis = [−16°,106°]
$\alpha_{R0} = 21°$
optimal translation axis = [68°,166°]
$\alpha_{T0} = 28°$
RTSA = −7°

input LPT neuons:
LPT05_R_VS1
LPT06_R_VS2
LPT13_R_VSm
LPT15_R_VST1
LPT17_R_VST2

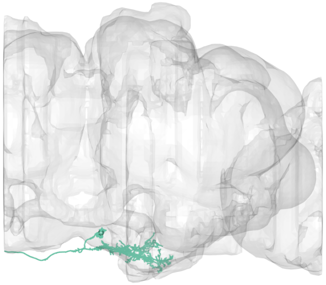

PMPM

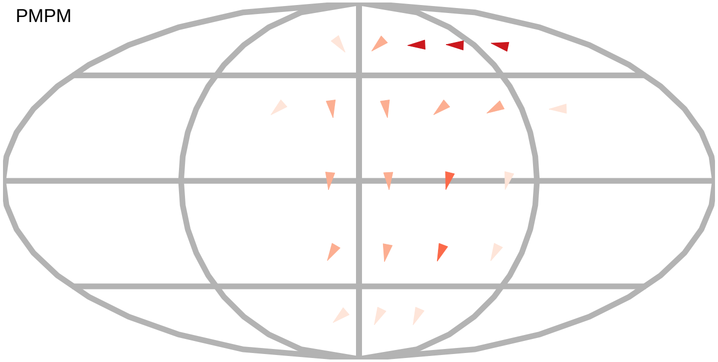

$\alpha_R$

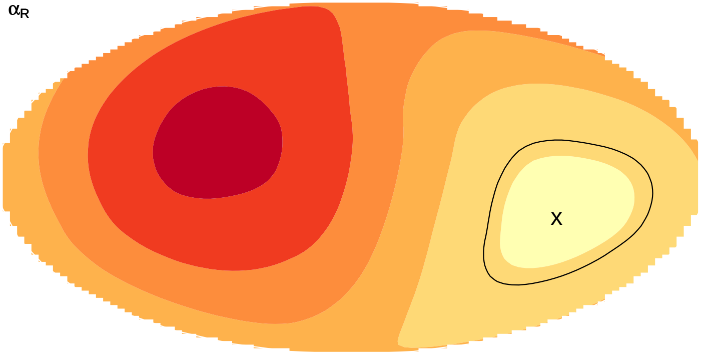


layer innervation

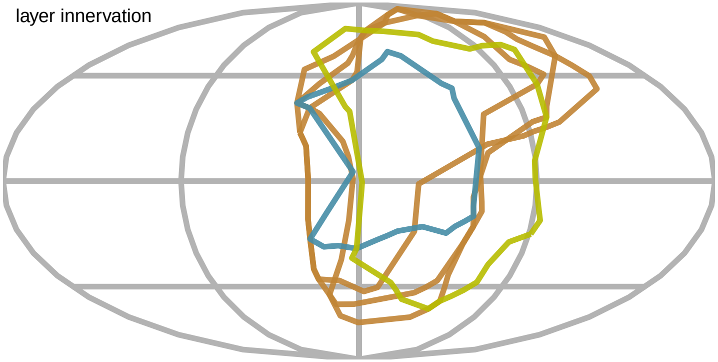

$\alpha_T$

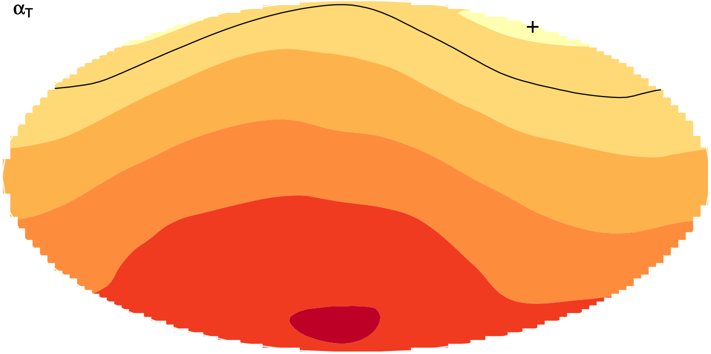

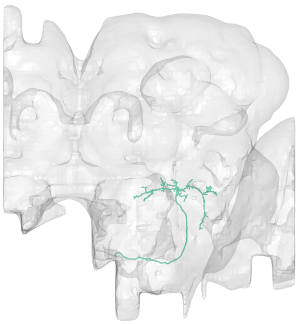

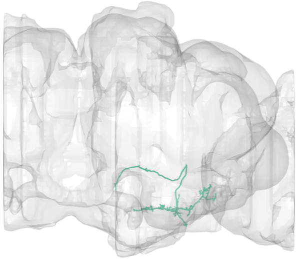

#9
bodyid =
1654880399

FOV = 34%
overlap of input LPTs' FOV = 8%
optimal rotation axis = [0°,90°]
$\alpha_{R0} = 34°$
optimal translation axis = [76°,−180°]
$\alpha_{T0} = 12°$
RTSA = 22°

input LPT neuons:
LPT49_R
LPT49_L

PMPM

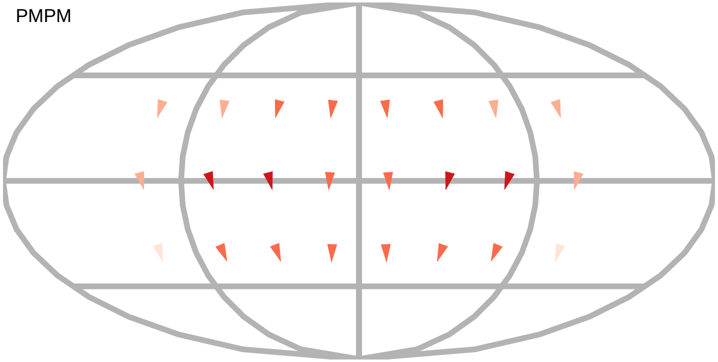

$\alpha_R$

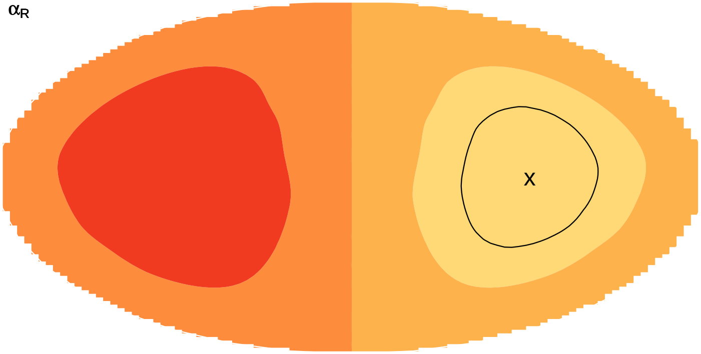

$\alpha_T$

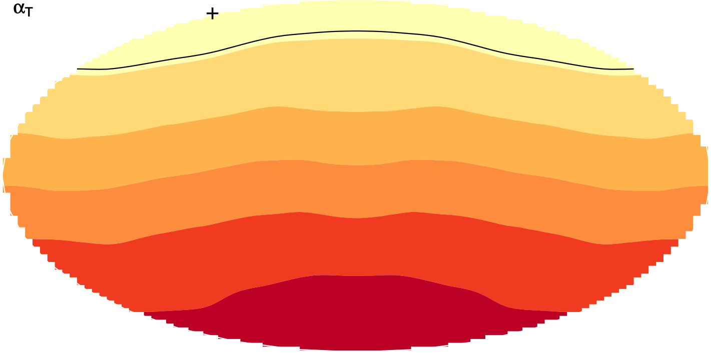

layer innervation

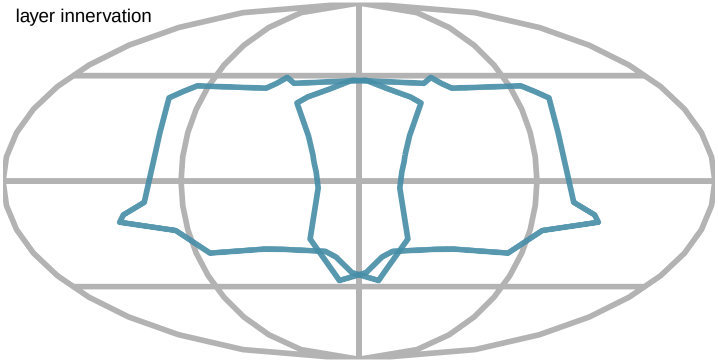

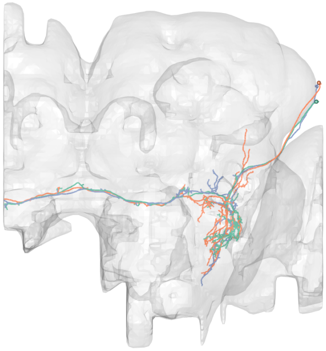

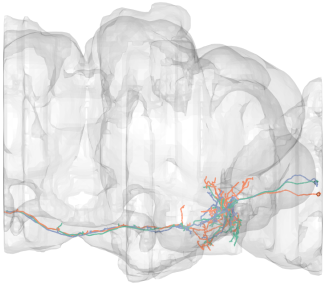

#10
bodyid =
1162305766
1471337986
1471337698

FOV = 36%
overlap of input LPTs' FOV = 12%
optimal rotation axis = [−2°,−142°]
$\alpha_{R0} = 13°$
optimal translation axis = [−82°,14°]
$\alpha_{T0} = 14°$
RTSA = −1°

input LPT neuons:
LPT48_L_vCal3
LPT47_R_vCal2
LPT47_L_vCal2

layer innervation

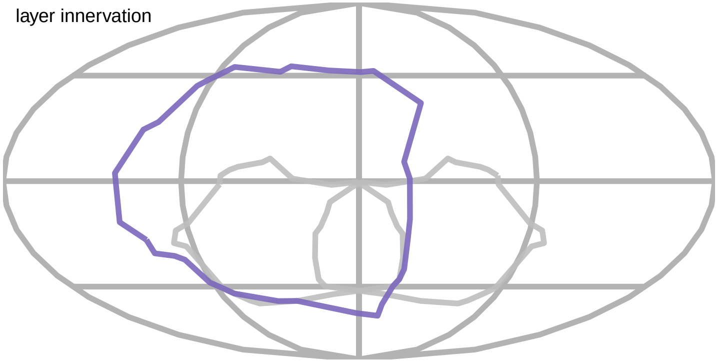

PMPM

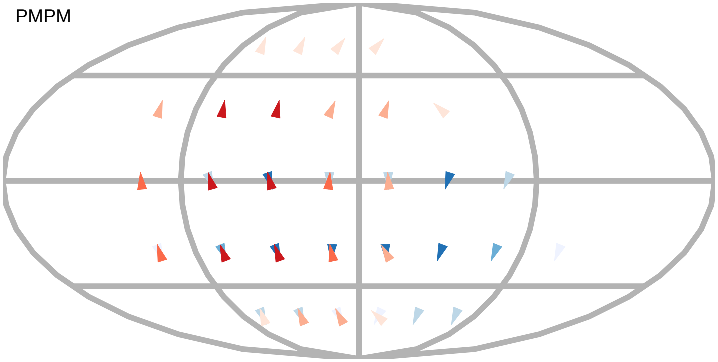

$\alpha_R$

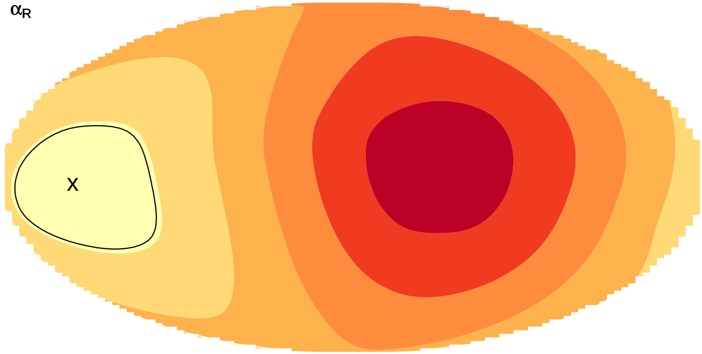


$\alpha_T$

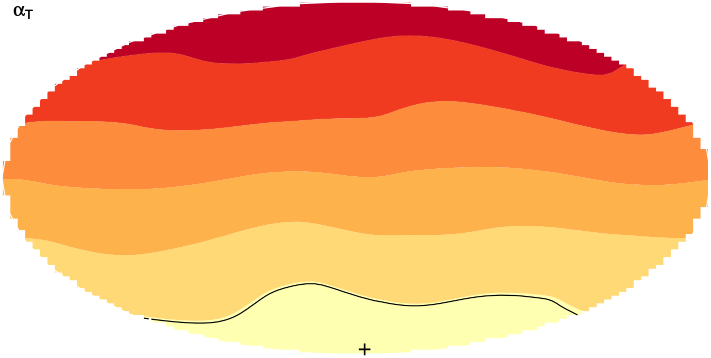

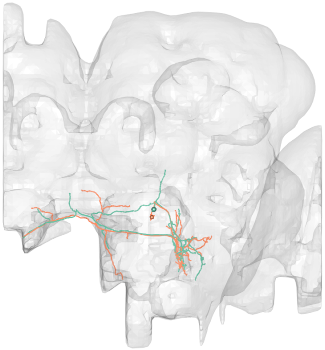

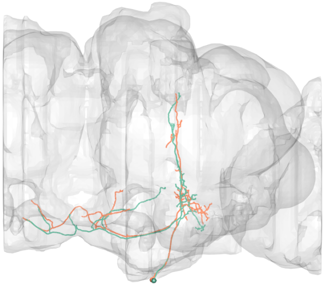

#11
bodyid =
1881013117
1849969844

FOV = 36%
overlap of input LPTs' FOV = 1%
optimal rotation axis = [32°,97°]
$\alpha_{R0} = 72°$
optimal translation axis = [2°,144°]
$\alpha_{T0} = 26°$
RTSA = 46°

input LPT neuons:
LPT21_R
LPT42_L_Nod4

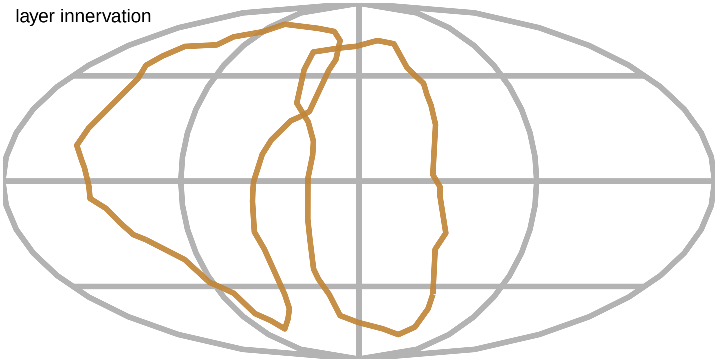


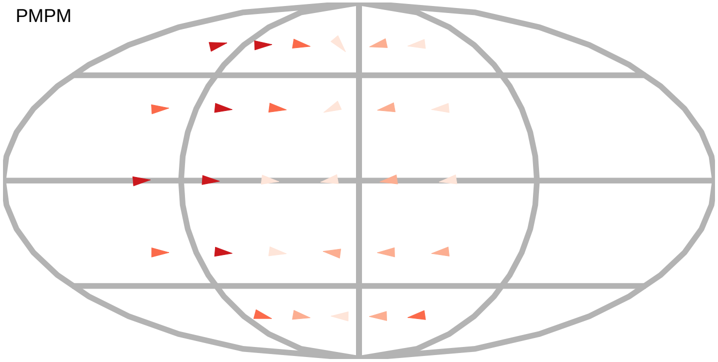


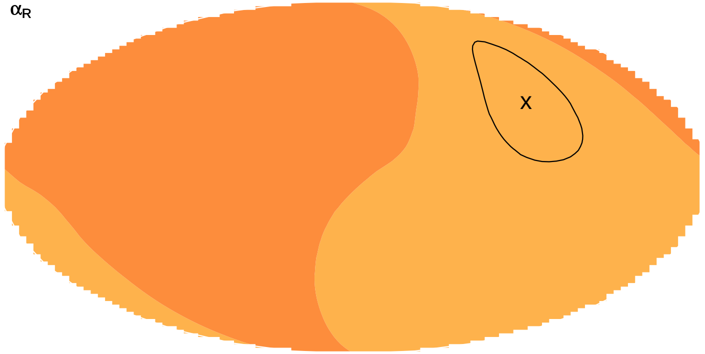


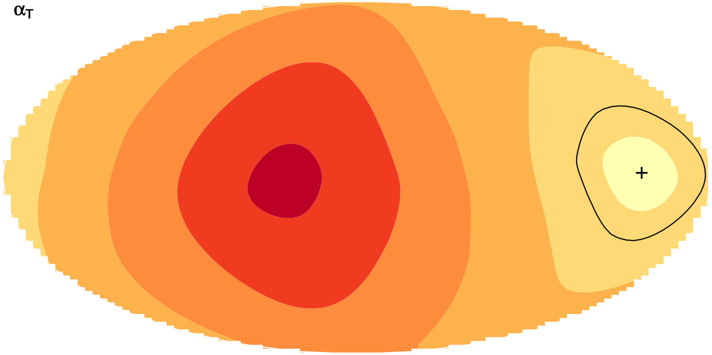

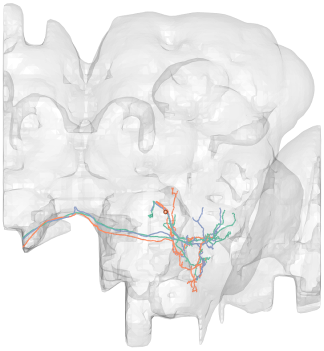

#12
bodyid =
1905170140
5813034397
1744608077

FOV = 36%
overlap of input LPTs' FOV = 1%
optimal rotation axis = [32°,97°]
$\alpha_{R0} = 72°$
optimal translation axis = [2°,144°]
$\alpha_{T0} = 26°$
RTSA = 46°

input LPT neuons:
LPT21_R
LPT42_L_Nod4

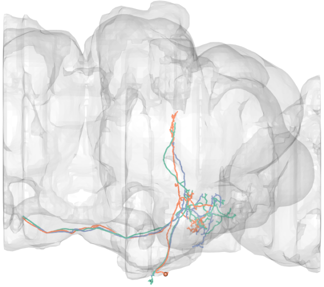

layer innervation

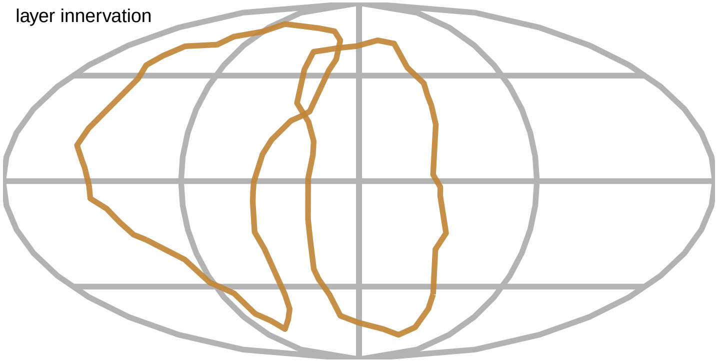

PMPM

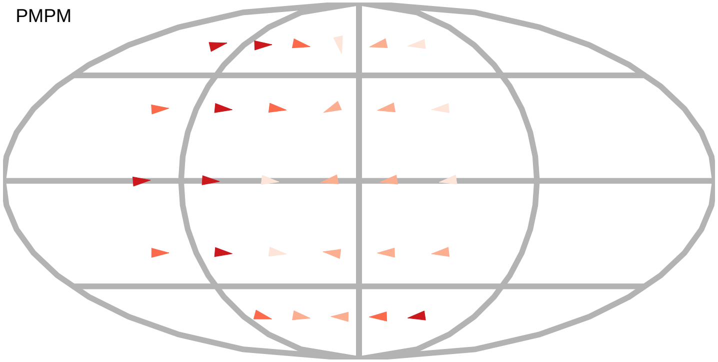

$\alpha_R$

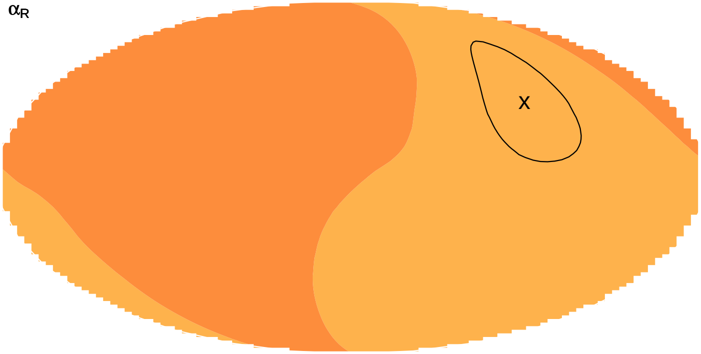


$\alpha_T$

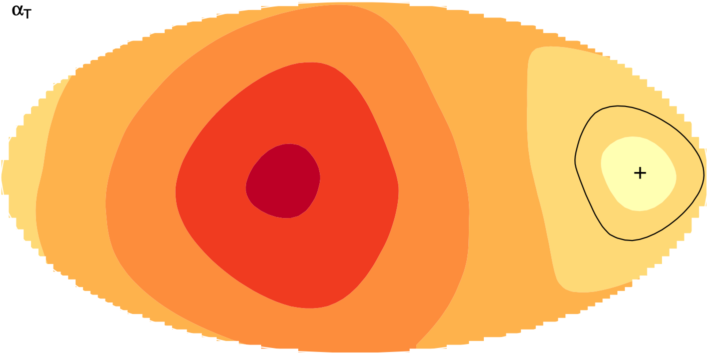

#13
bodyid =
1749313891
5813054847

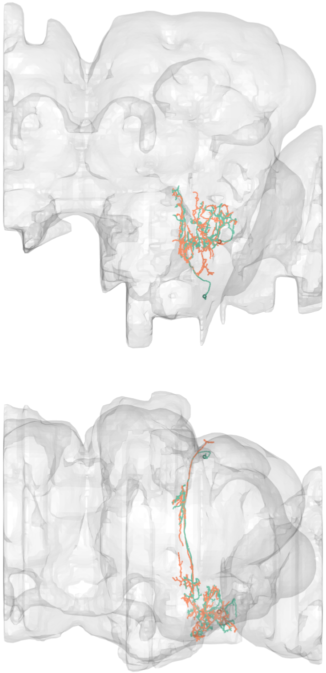

FOV = 38%
overlap of input LPTs' FOV = 4%
optimal rotation axis = [0°,−118°]
$\alpha_{R0} = 77°$
optimal translation axis = [0°,34°]
$\alpha_{T0} = 32°$
RTSA = 45°

input LPT neuons:
LPT26_R
LPT30_R
LPT38_L_Nod1

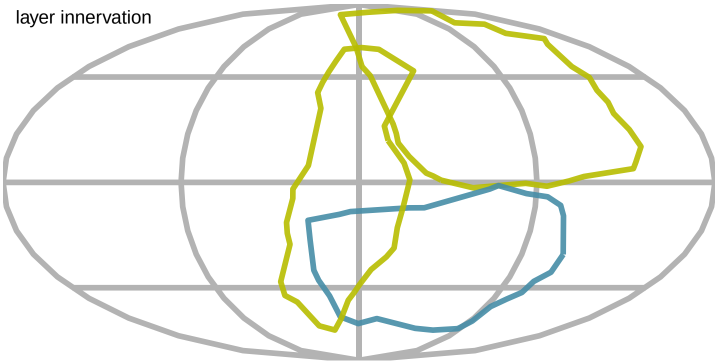


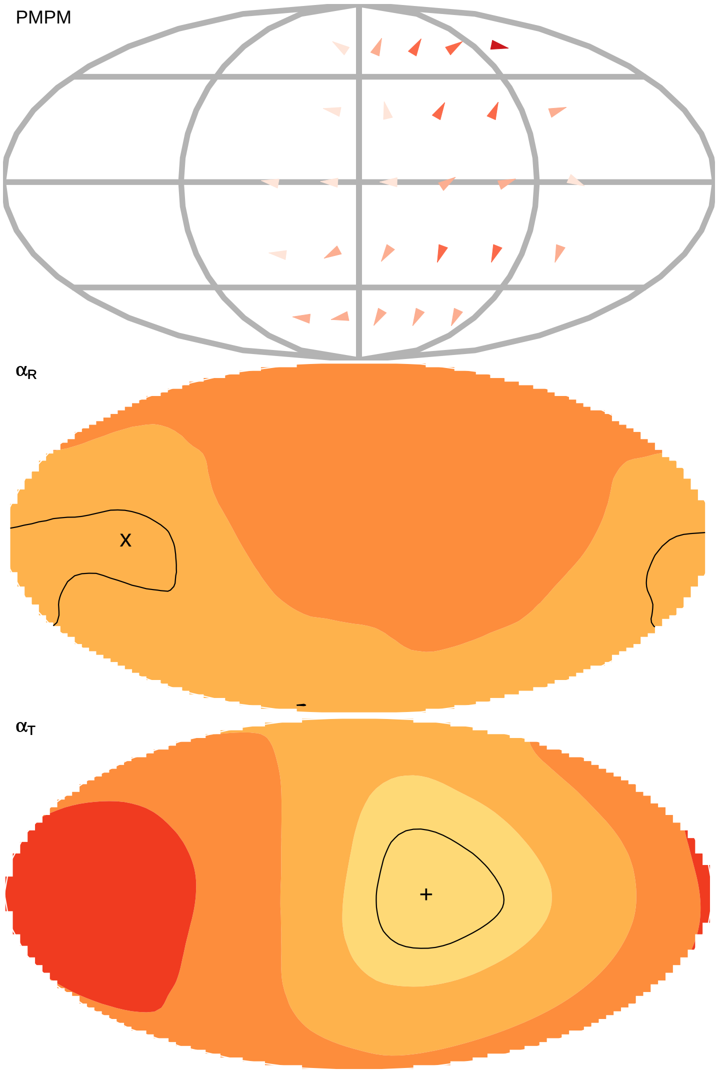

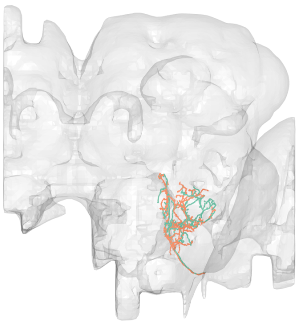

#14
bodyid =
1778311161
1633747719

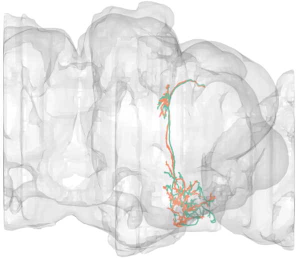

FOV = 38%
overlap of input LPTs' FOV = 4%
optimal rotation axis = [0°,−118°]
$\alpha_{R0} = 77°$
optimal translation axis = [0°,34°]
$\alpha_{T0} = 32°$
RTSA = 45°

input LPT neuons:
LPT26_R
LPT30_R
LPT38_L_Nod1

PMPM

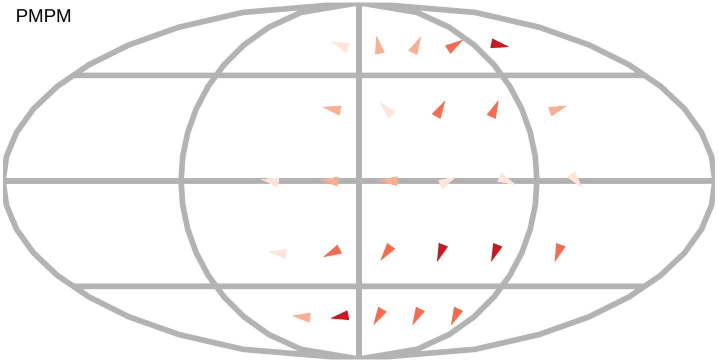

$\alpha_R$

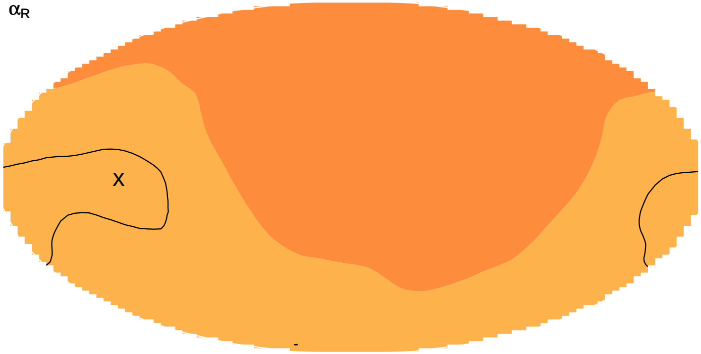


layer innervation

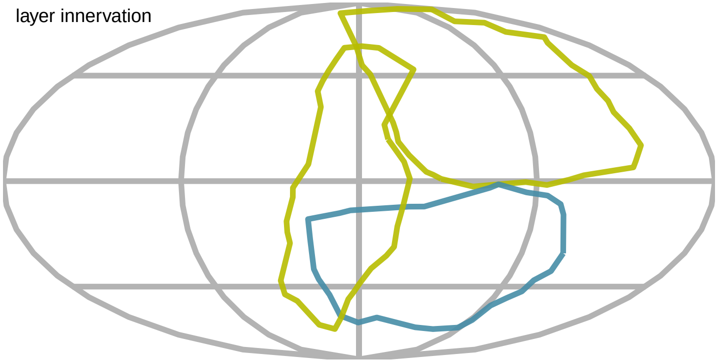

$\alpha_T$

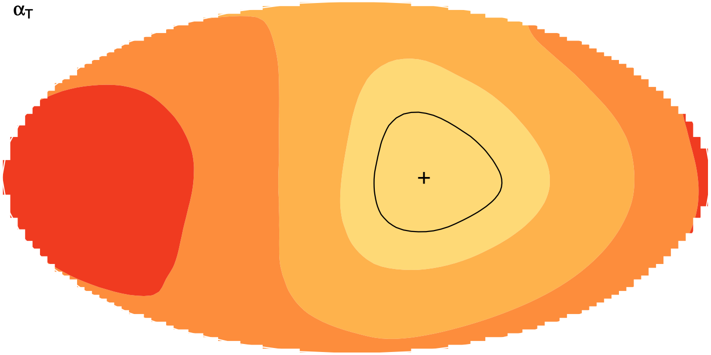

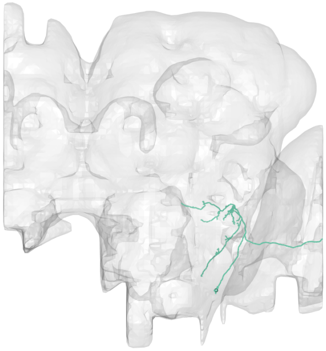

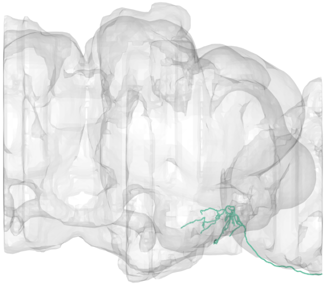

#15
bodyid =
1717614297

FOV = 39%
overlap of input LPTs' FOV = 6%
optimal rotation axis = [−2°,−142°]
$\alpha_{R0} = 13°$
optimal translation axis = [−82°,14°]
$\alpha_{T0} = 14°$
RTSA = −1°

input LPT neuons:
LPT48_L_vCal3
LPT46_R_vCal1
LPT46_L_vCal1

layer innervation

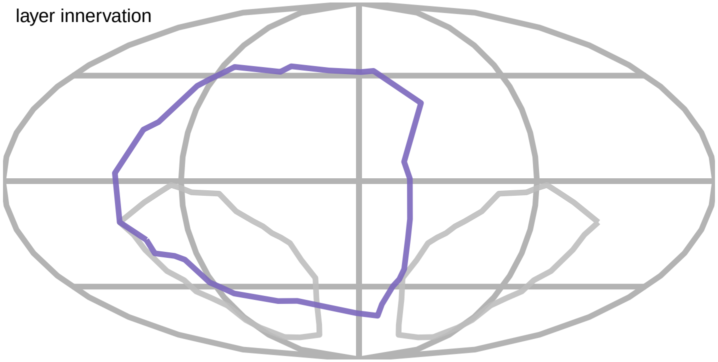

PMPM

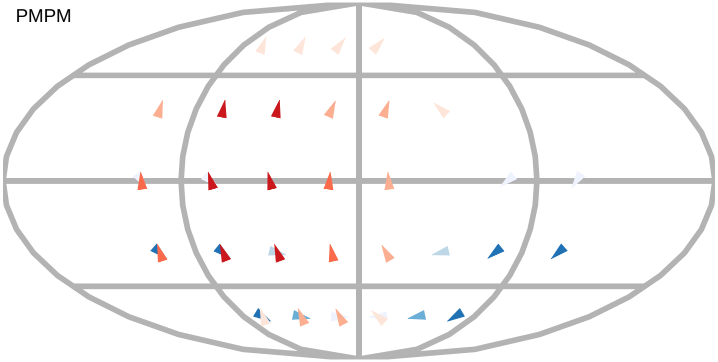

$\alpha_R$

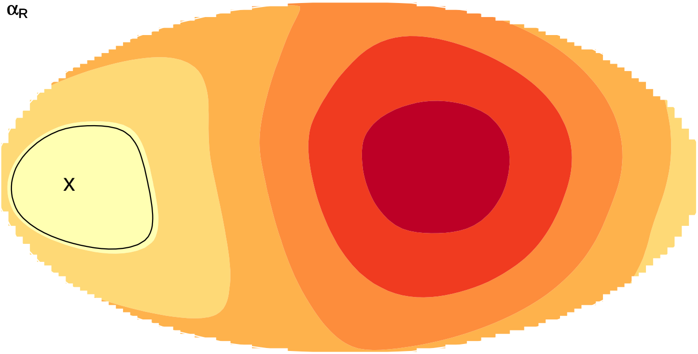


$\alpha_T$

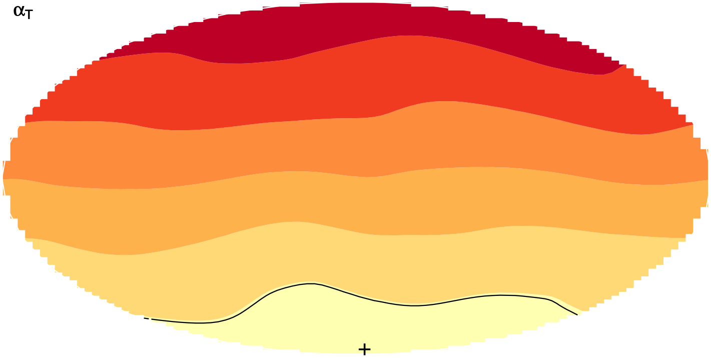

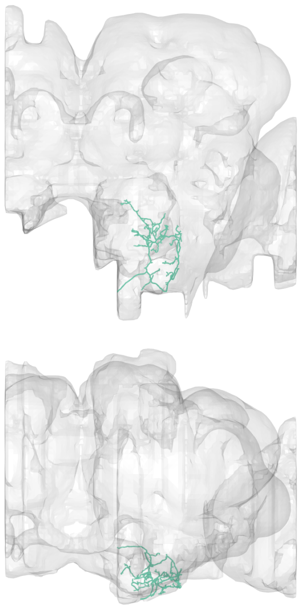

#16
bodyid =
1870483009

FOV = 40%
overlap of input LPTs' FOV = 4%
optimal rotation axis = [−2°,−42°]
$\alpha_{R0} = 11°$
optimal translation axis = [−68°,27°]
$\alpha_{T0} = 15°$
RTSA = −4°

input LPT neuons:
LPT28_R
LPT50_L

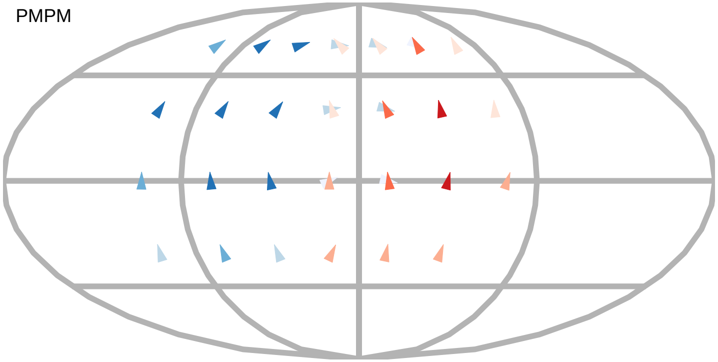


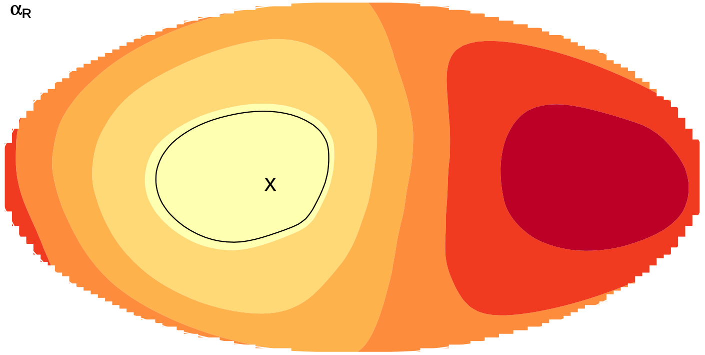


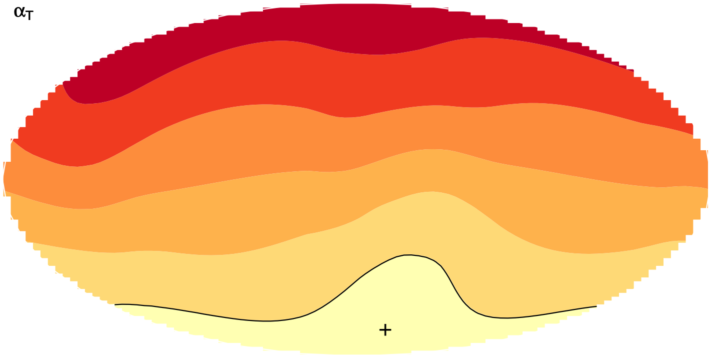


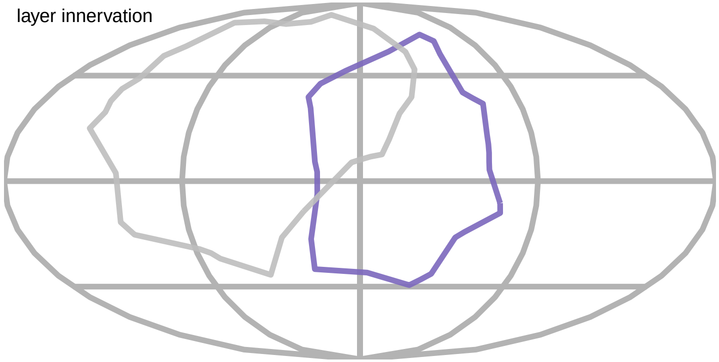

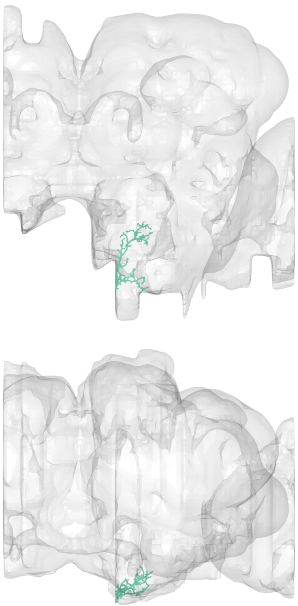

#17
bodyid =
5813003831

FOV = 40%
overlap of input LPTs' FOV = 4%
optimal rotation axis = [−2°,−42°]
$\alpha_{R0} = 11°$
optimal translation axis = [−68°,27°]
$\alpha_{T0} = 15°$
RTSA = −4°

input LPT neuons:
LPT28_R
LPT50_L

layer innervation

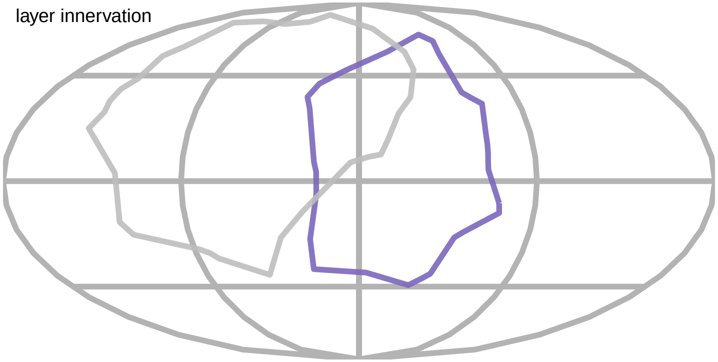

PMPM

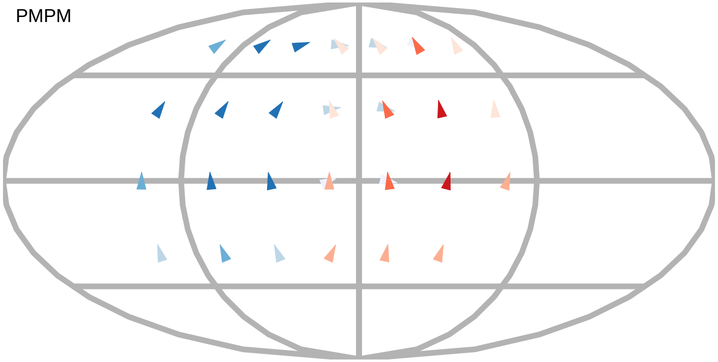

$\alpha_R$

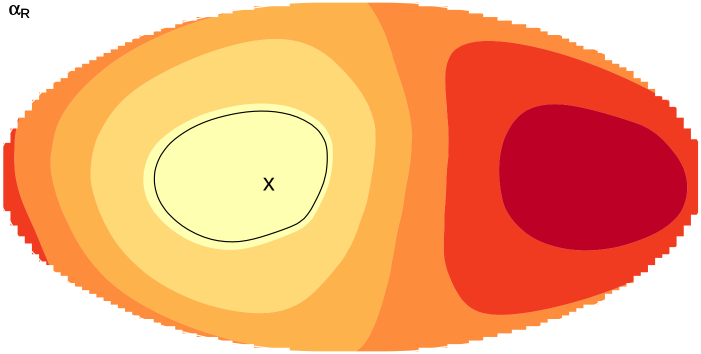


$\alpha_T$

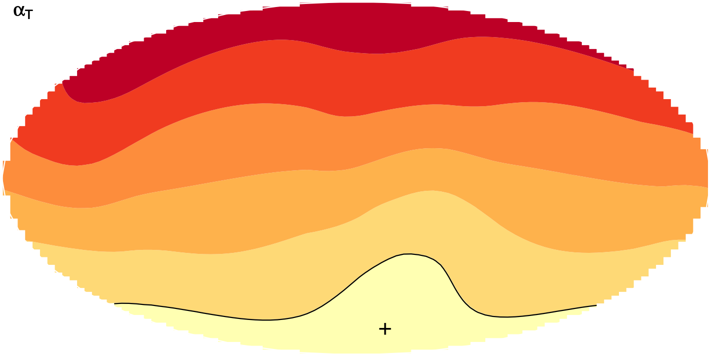

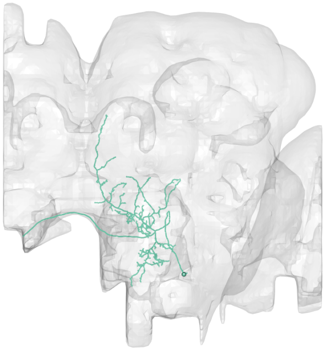

#18
bodyid =
1222195618

FOV = 40%
overlap of input LPTs' FOV = 4%
optimal rotation axis = [−2°,−42°]
$\alpha_{R0} = 11°$
optimal translation axis = [−68°,27°]
$\alpha_{T0} = 15°$
RTSA = −4°

input LPT neuons:
LPT28_R
LPT50_L

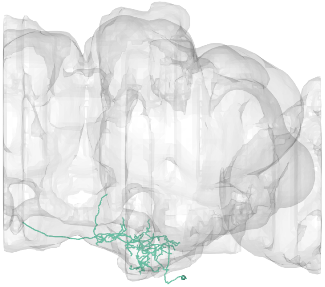

layer innervation

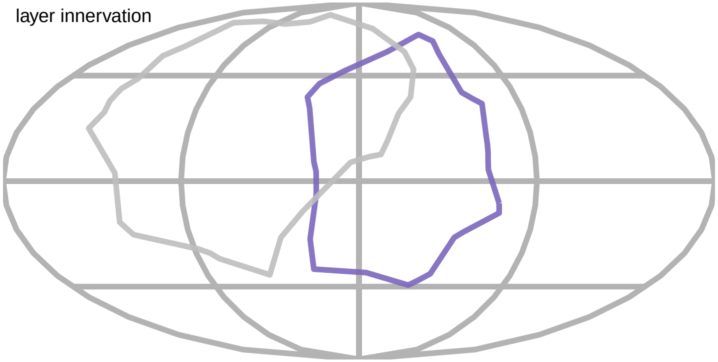

PMPM

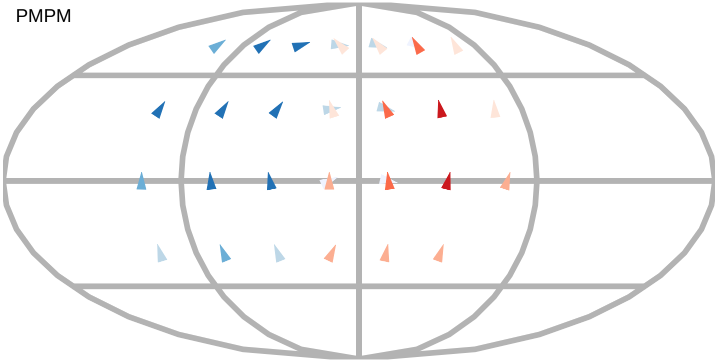

$\alpha_R$

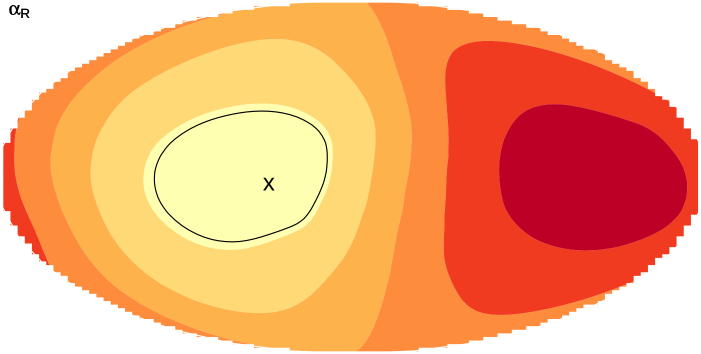


$\alpha_T$

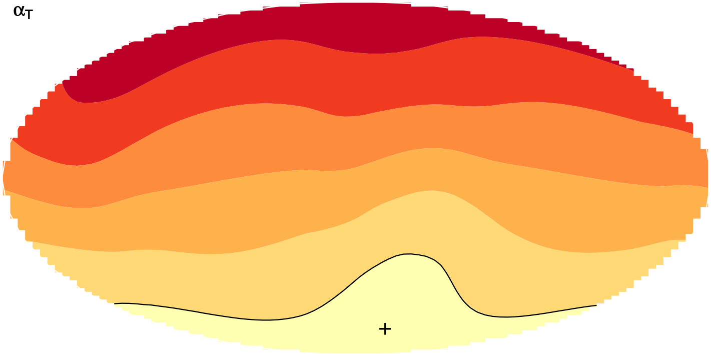

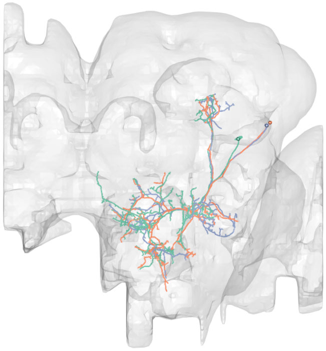

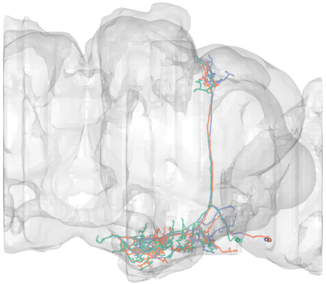

#19
bodyid =
1374839724
1413058496
1500693034

FOV = 40%
overlap of input LPTs' FOV = 4%
optimal rotation axis = [−2°,−42°]
$\alpha_{R0} = 11°$
optimal translation axis = [−68°,27°]
$\alpha_{T0} = 15°$
RTSA = −4°

input LPT neuons:
LPT28_R
LPT50_L

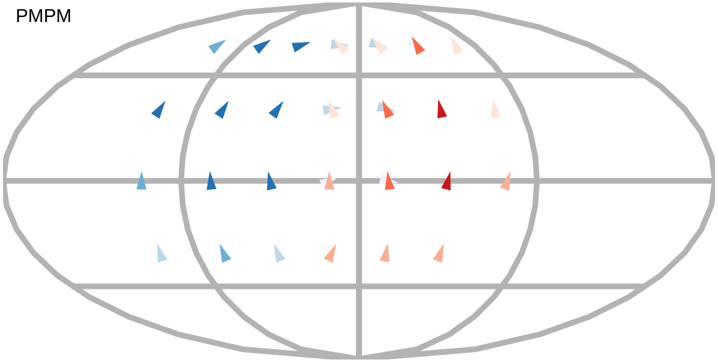


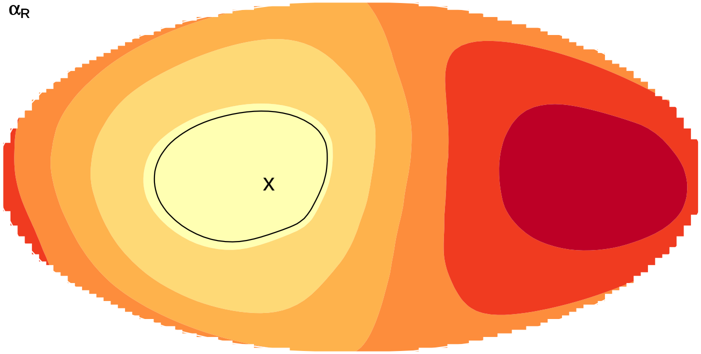


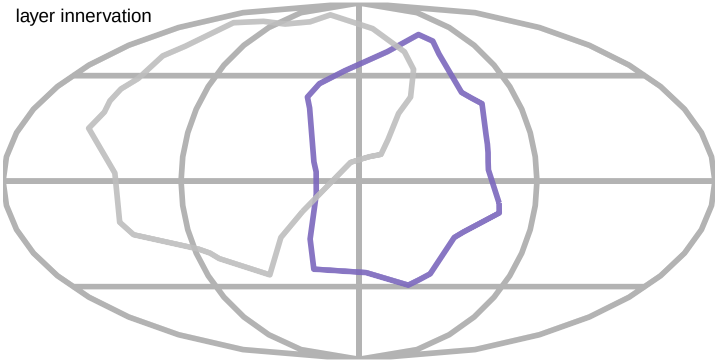


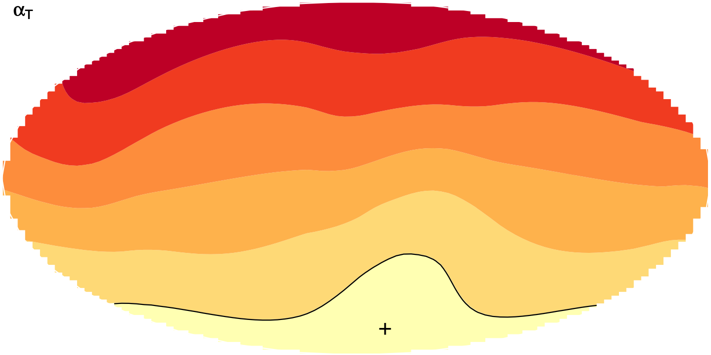

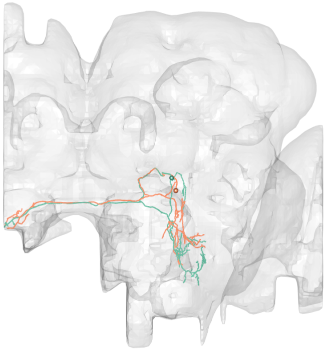

#20
bodyid =
2029345079
1819102805

FOV = 40%
overlap of input LPTs' FOV = 0%
optimal rotation axis = [−58°,−159°]
$\alpha_{R0} = 77°$
optimal translation axis = [2°,166°]
$\alpha_{T0} = 17°$
RTSA = 60°

input LPT neuons:
LPT41_R_Nod3
LPT42_L_Nod4

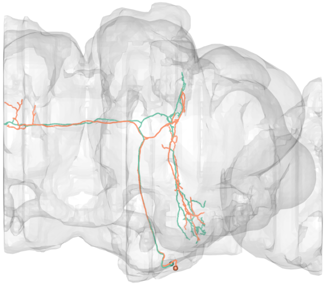

layer innervation

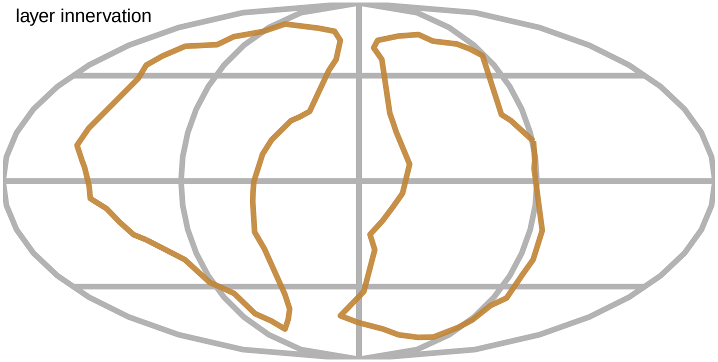

PMPM

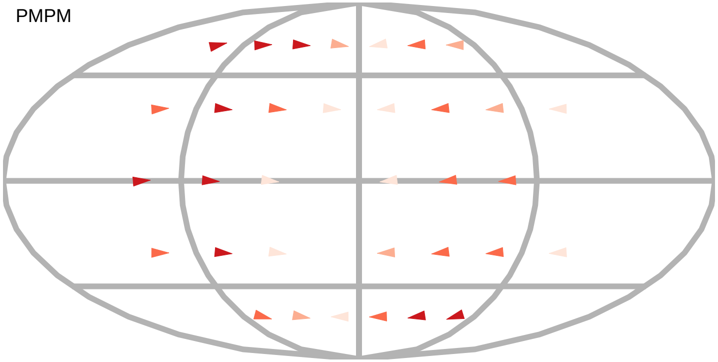

$\alpha_R$

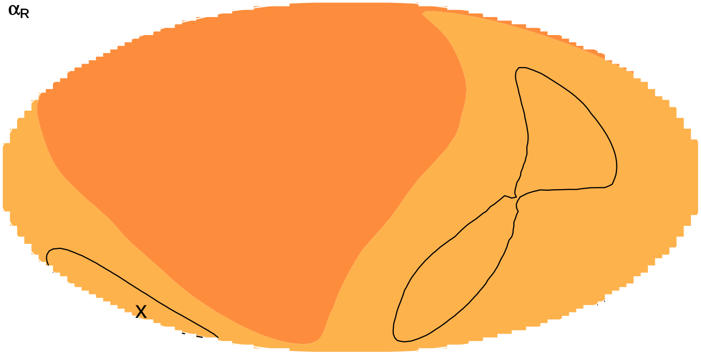


$\alpha_T$

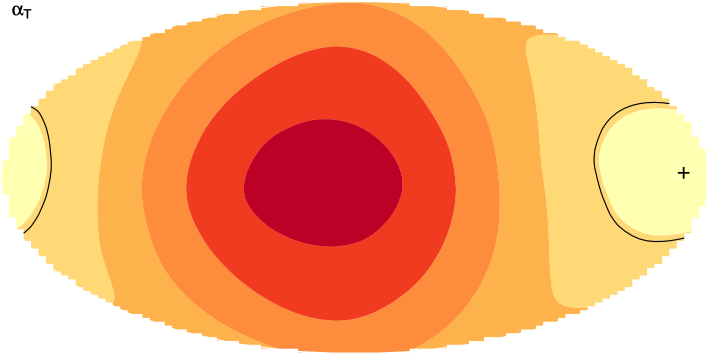

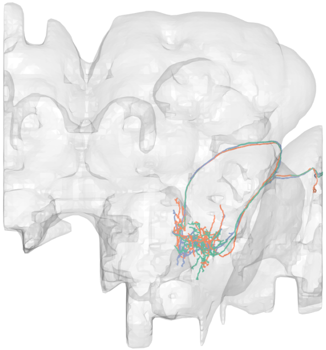

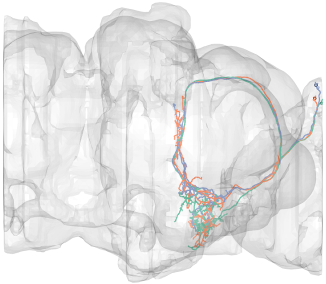

#21
bodyid =
1720300315
1932816628
1913066247

FOV = 40%
overlap of input LPTs' FOV = 15%
optimal rotation axis = [−78°,173°]
$\alpha_{R0} = 58°$
optimal translation axis = [0°,56°]
$\alpha_{T0} = 25°$
RTSA = 33°

input LPT neuons:
LPT21_R
LPT26_R
LPT30_R
LPT38_L_Nod1

PMPM

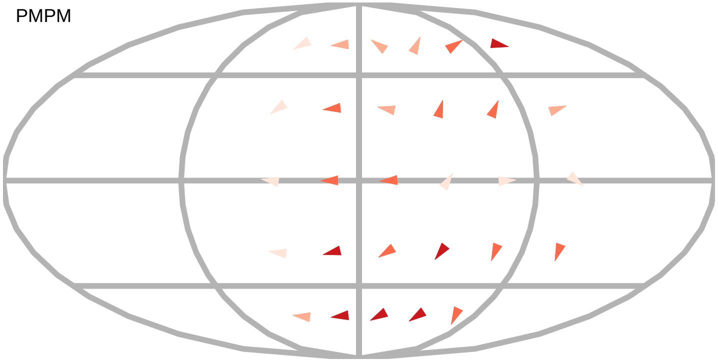

$\alpha_R$

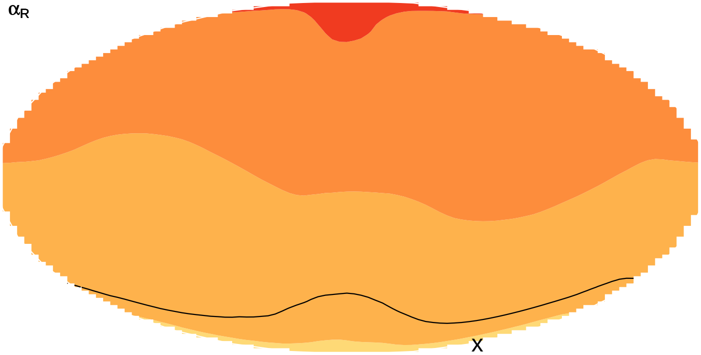


$\alpha_T$

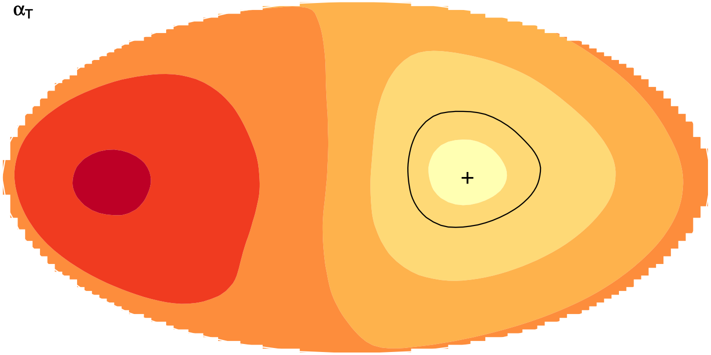


layer innervation

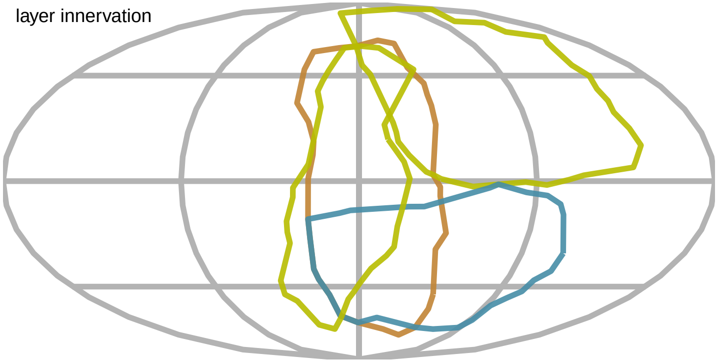

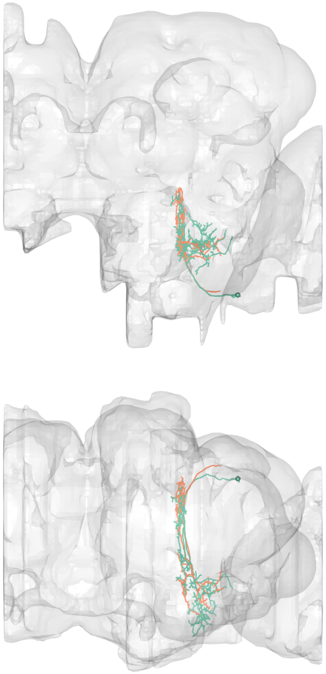

#22
bodyid =
1664782583
1634088865

FOV = 40%
overlap of input LPTs' FOV = 15%
optimal rotation axis = [−78°,173°]
$\alpha_{R0} = 58°$
optimal translation axis = [2°,56°]
$\alpha_{T0} = 25°$
RTSA = 33°

input LPT neuons:
LPT21_R
LPT26_R
LPT30_R
LPT38_L_Nod1

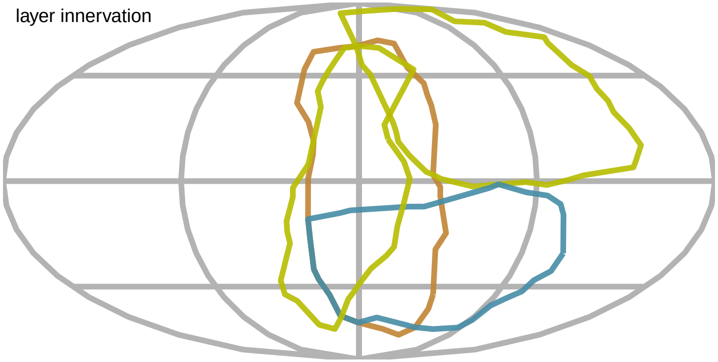


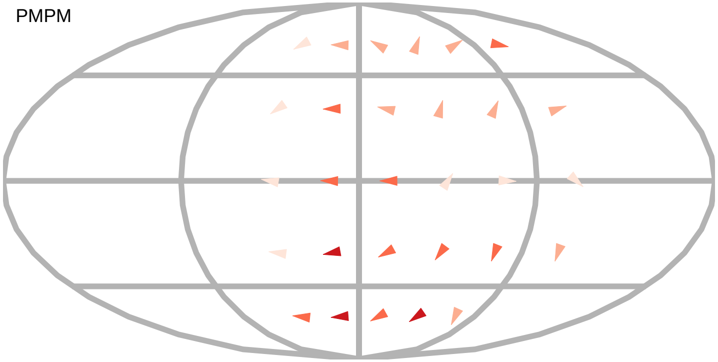


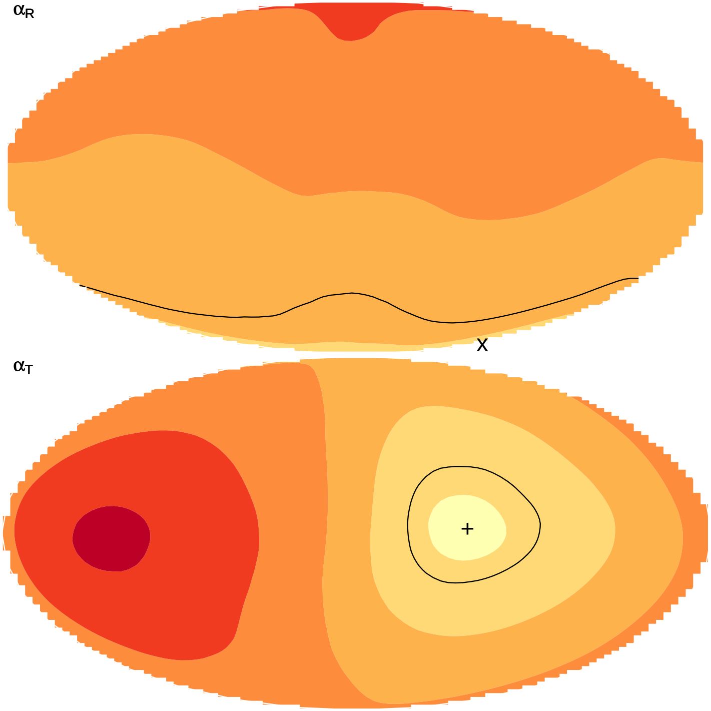

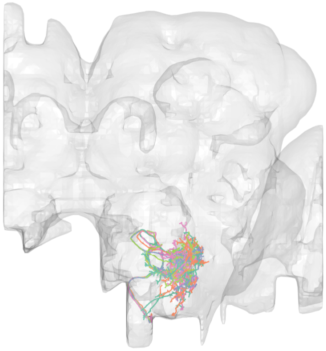

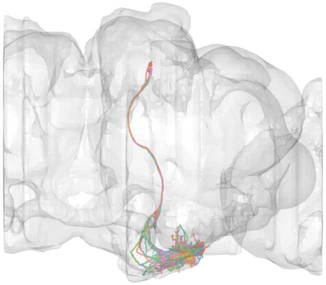

#23
bodyid =
5813034469
5813108981
5813022617
1933563873
1933904658

FOV = 44%
overlap of input LPTs' FOV = 9%
optimal rotation axis = [86°,29°]
$\alpha_{R0} = 7°$
optimal translation axis = [6°,−109°]
$\alpha_{T0} = 30°$
RTSA = −23°

input LPT neuons:
LPT04_R_HST
LPT43_L_H2

layer innervation

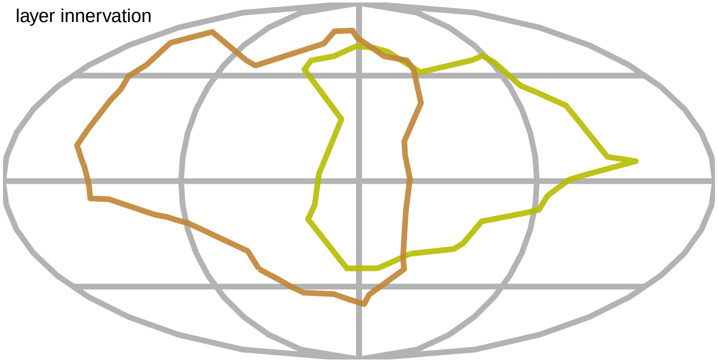

PMPM

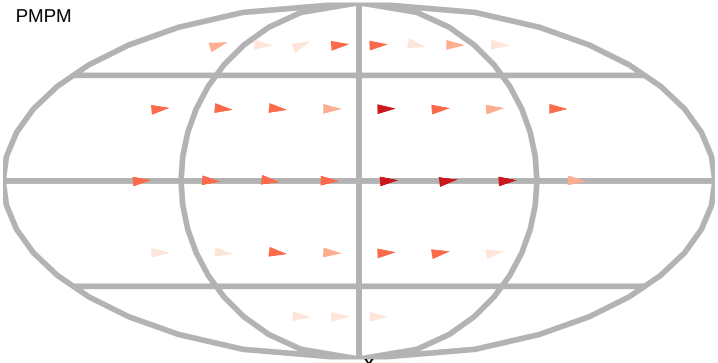

$\alpha_R$

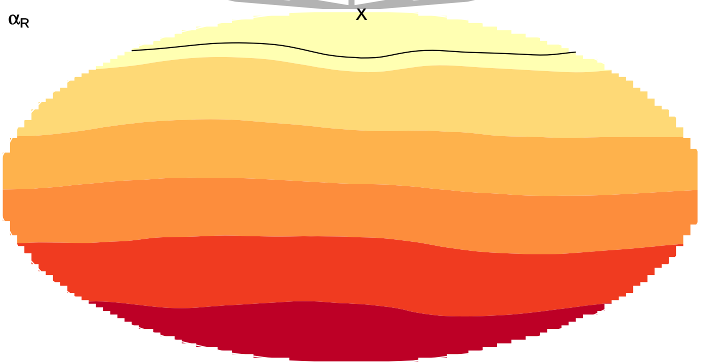

$\alpha_T$

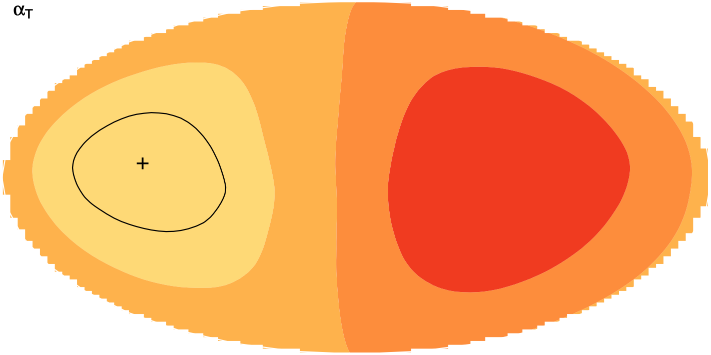

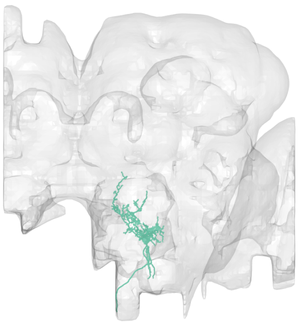

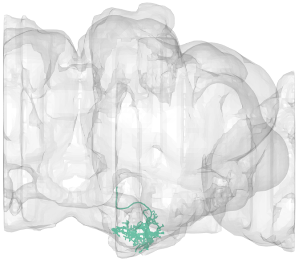

#24
bodyid =
5813024082

FOV = 46%
overlap of input LPTs' FOV = 23%
optimal rotation axis = [36°,109°]
$\alpha_{R0} = 41°$
optimal translation axis = [24°,−149°]
$\alpha_{T0} = 39°$
RTSA = 2°

input LPT neuons:
LPT06_R_VS2
LPT13_R_VSm
LPT15_R_VST1
LPT43_L_H2

PMPM

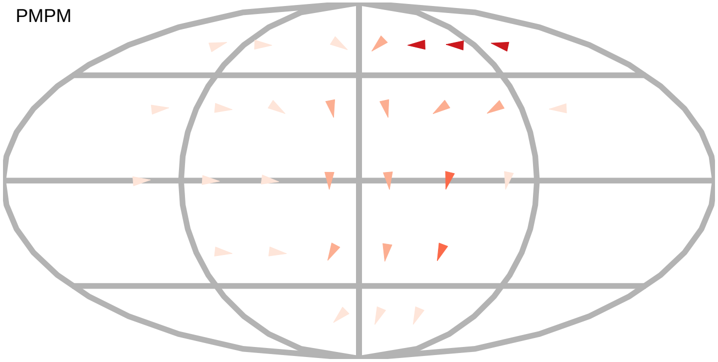

$\alpha_R$

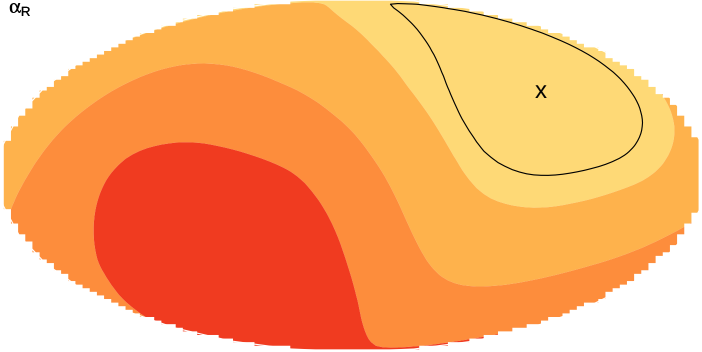


layer innervation

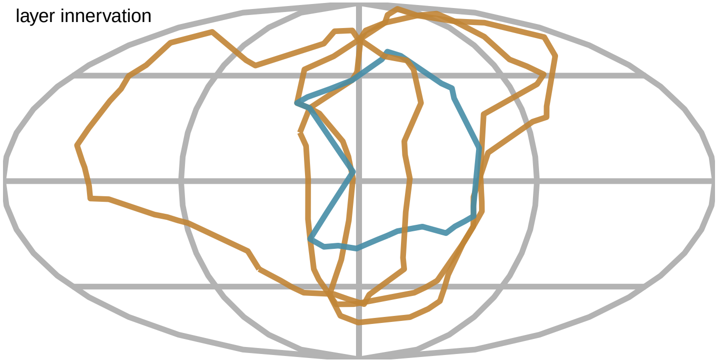

$\alpha_T$

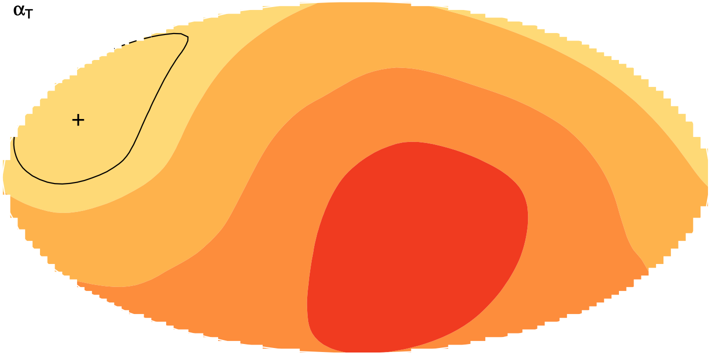

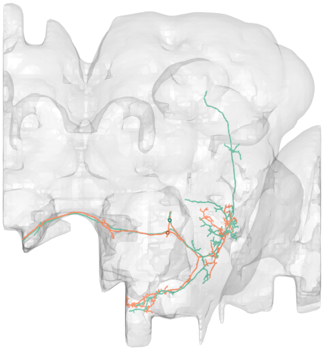

#25
bodyid =
1284468417
1933576833

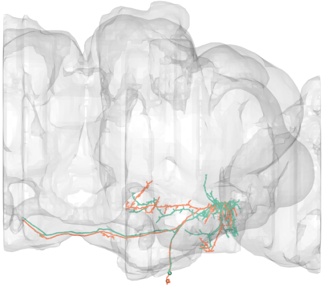

FOV = 47%
overlap of input LPTs' FOV = 11%
optimal rotation axis = [0°,−90°]
$\alpha_{R0} = 39°$
optimal translation axis = [−80°,0°]
$\alpha_{T0} = 14°$
RTSA = 25°

input LPT neuons:
LPT48_R_vCal3
LPT48_L_vCal3

PMPM

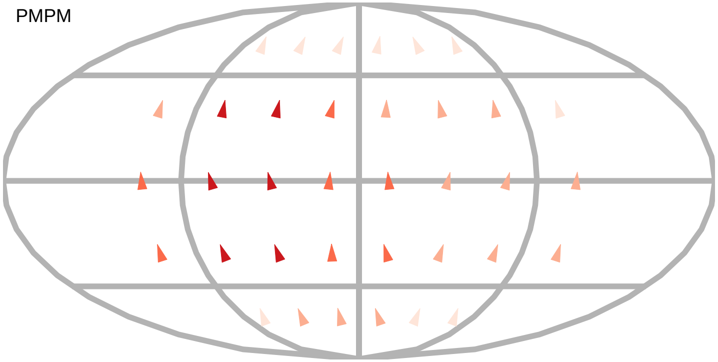

$\alpha_R$

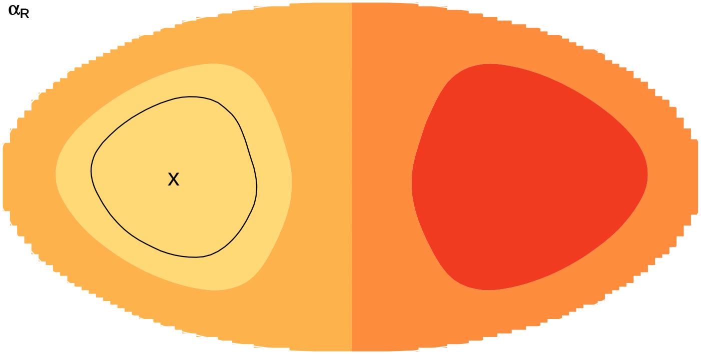

$\alpha_T$

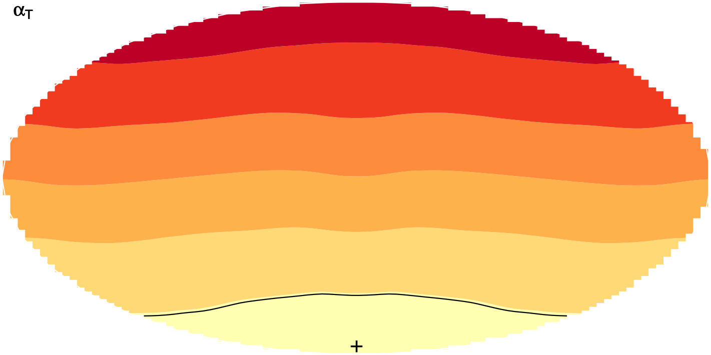

layer innervation

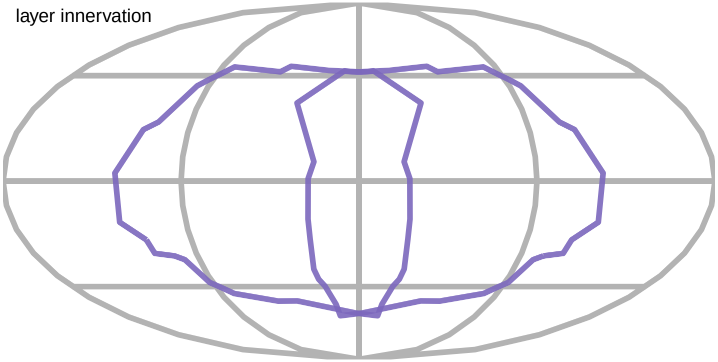

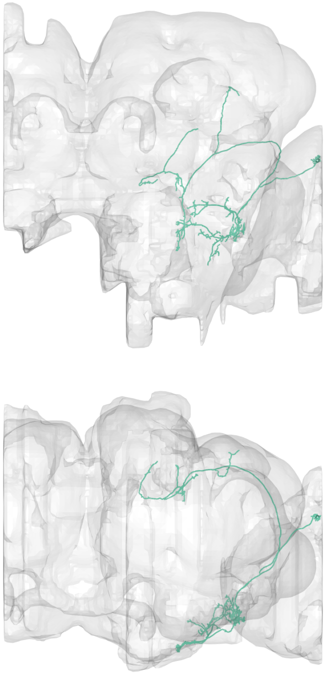

#26
bodyid =
1504052107

FOV = 47%
overlap of input LPTs' FOV = 25%
optimal rotation axis = [0°,−90°]
$\alpha_{R0} = 39°$
optimal translation axis = [−80°,0°]
$\alpha_{T0} = 14°$
RTSA = 25°

input LPT neuons:
LPT48_R_vCal3
LPT48_L_vCal3
LPT47_R_vCal2
LPT47_L_vCal2

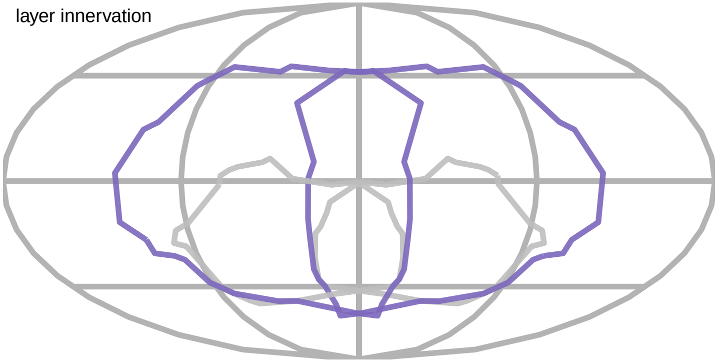


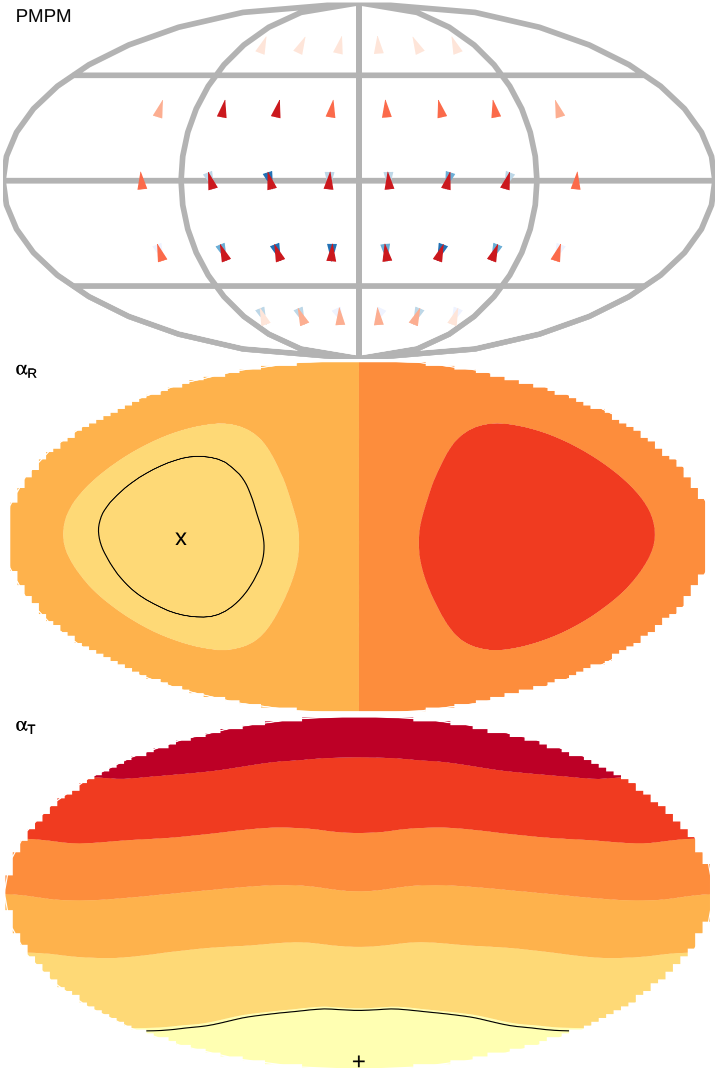

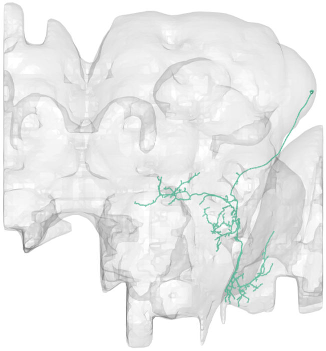

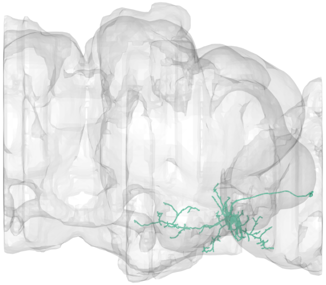

#27
bodyid =
1686229077

FOV = 47%
overlap of input LPTs' FOV = 25%
optimal rotation axis = [0°,−90°]
$\alpha_{R0} = 39°$
optimal translation axis = [−80°,0°]
$\alpha_{T0} = 14°$
RTSA = 25°

input LPT neuons:
LPT48_R_vCal3
LPT48_L_vCal3
LPT47_R_vCal2
LPT47_L_vCal2

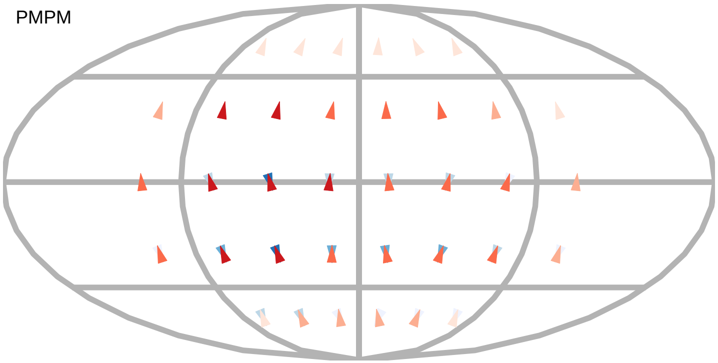


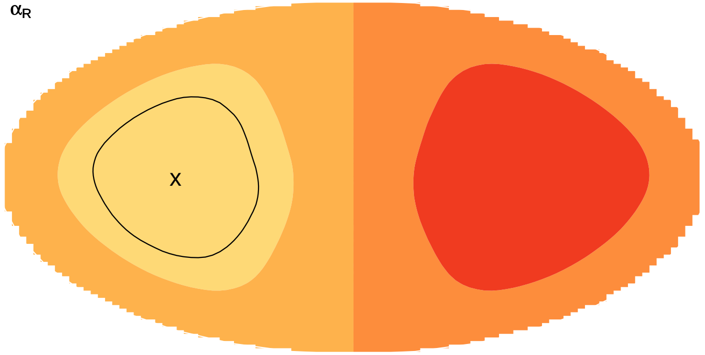


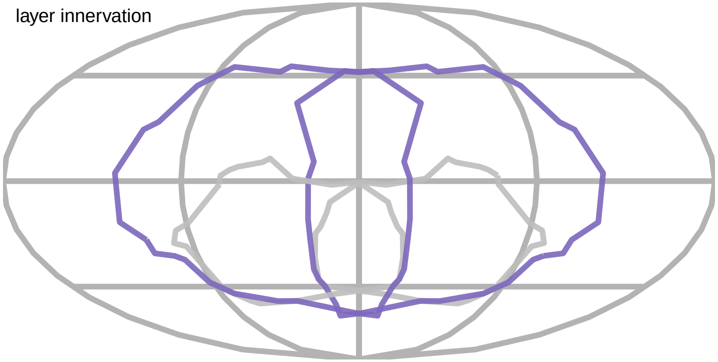


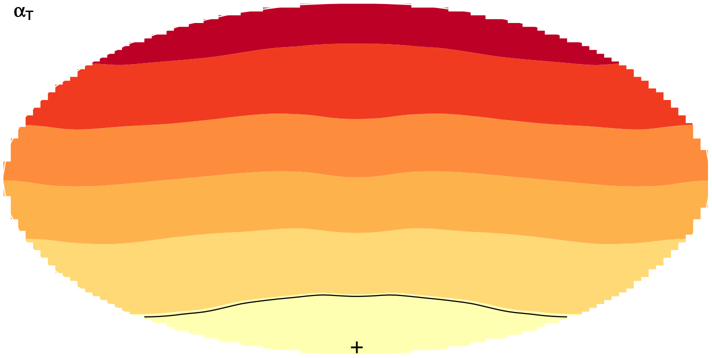

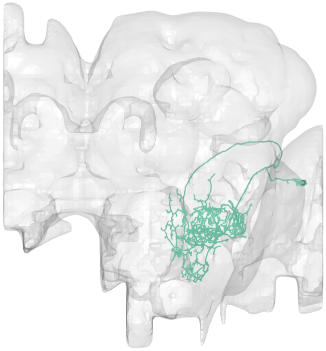

#28
bodyid =
1441978094

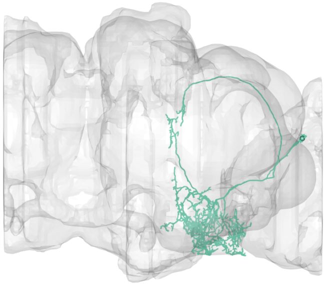

FOV = 47%
overlap of input LPTs' FOV = 37%
optimal rotation axis = [48°,126°]
$\alpha_{R0} = 79°$
optimal translation axis = [4°,−180°]
$\alpha_{T0} = 24°$
RTSA = 55°

input LPT neuons:
LPT41_R_Nod3
LPT41_L_Nod3
LPT40_R_Nod2
LPT40_L_Nod2

PMPM

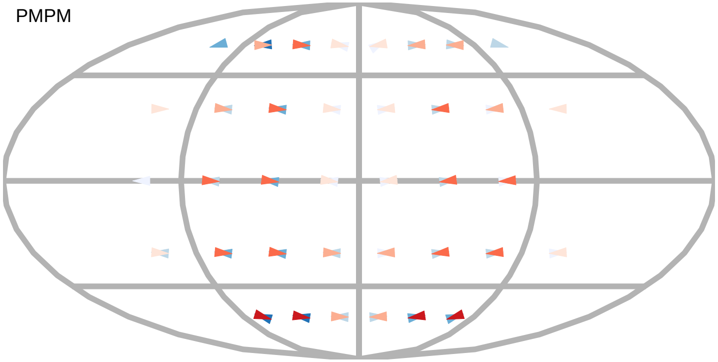

$\alpha_R$

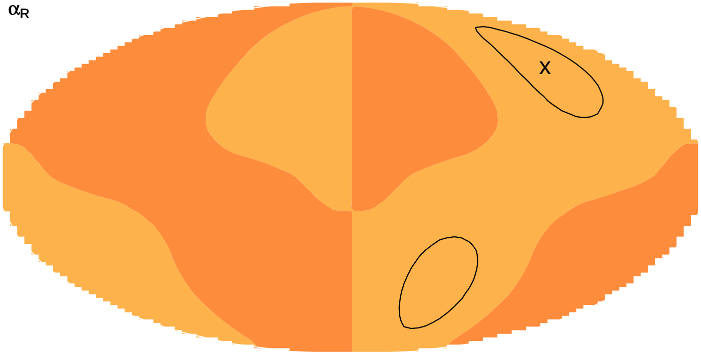



$\alpha_T$

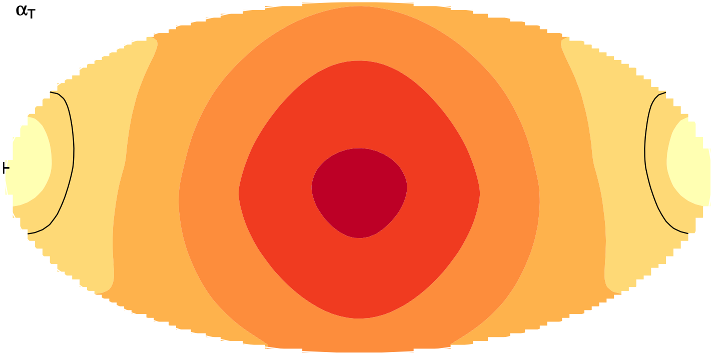

layer innervation

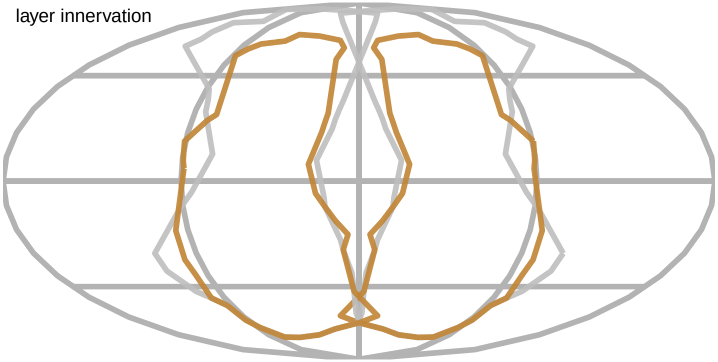

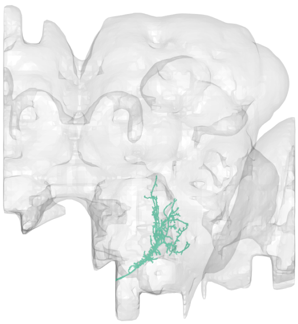

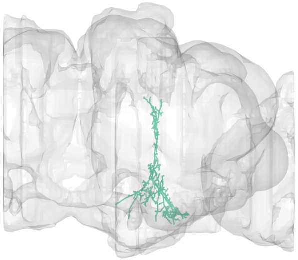

#29
bodyid =
1633074483

FOV = 47%
overlap of input LPTs' FOV = 37%
optimal rotation axis = [48°,126°]
$\alpha_{R0} = 79°$
optimal translation axis = [4°,−180°]
$\alpha_{T0} = 24°$
RTSA = 55°

input LPT neuons:
LPT41_R_Nod3
LPT41_L_Nod3
LPT40_R_Nod2
LPT40_L_Nod2

PMPM

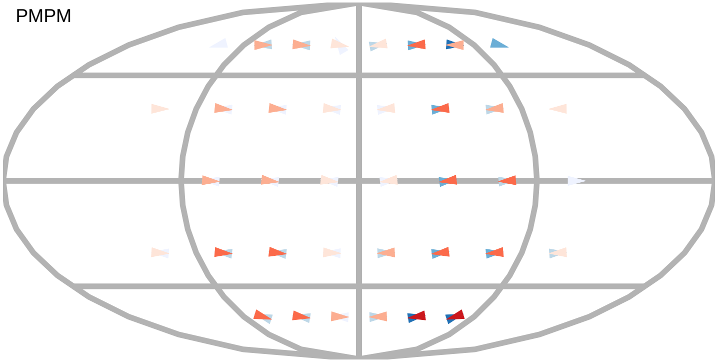

$\alpha_R$

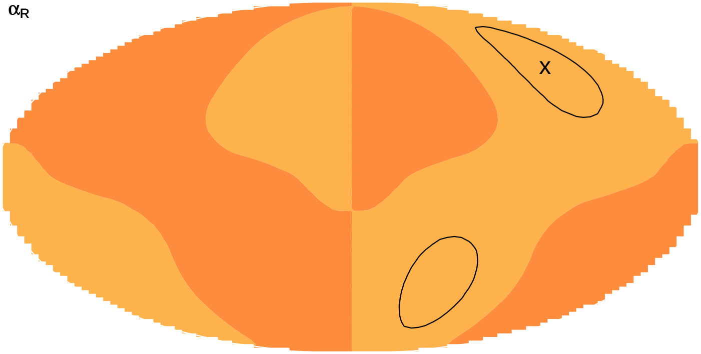


layer innervation

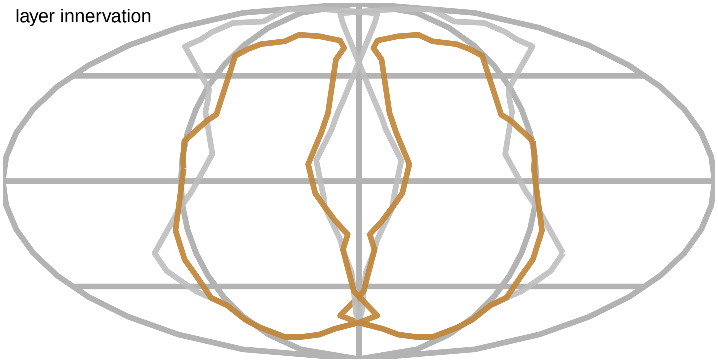

$\alpha_T$

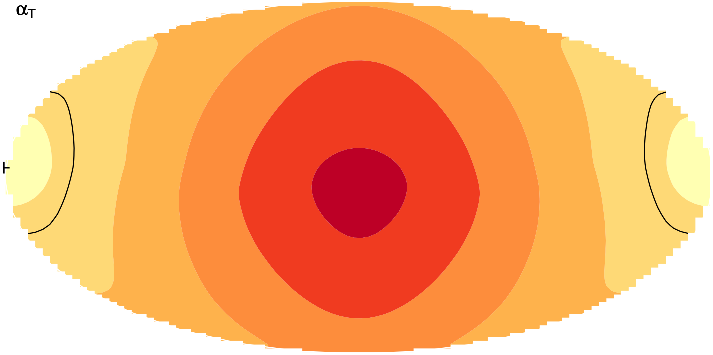

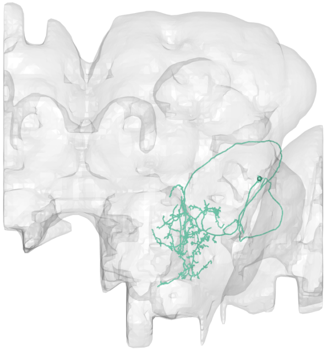

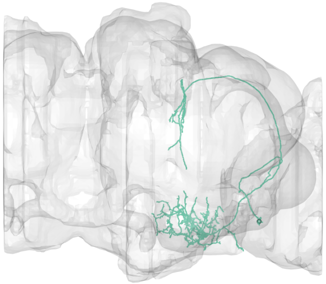

#30
bodyid =
1504388714

FOV = 47%
overlap of input LPTs' FOV = 37%
optimal rotation axis = [48°,126°]
$\alpha_{R0} = 79°$
optimal translation axis = [4°,−180°]
$\alpha_{T0} = 24°$
RTSA = 55°

input LPT neuons:
LPT41_R_Nod3
LPT41_L_Nod3
LPT40_R_Nod2
LPT40_L_Nod2

layer innervation

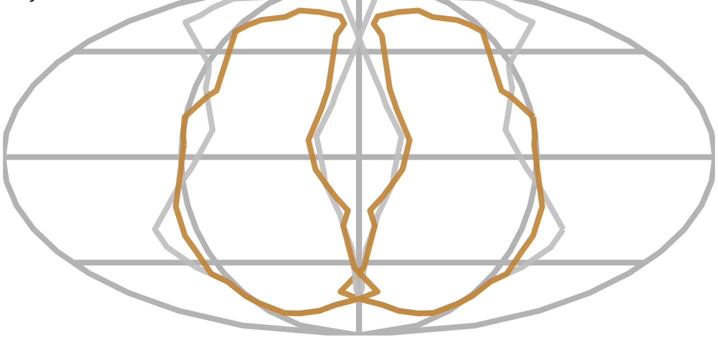

PMPM

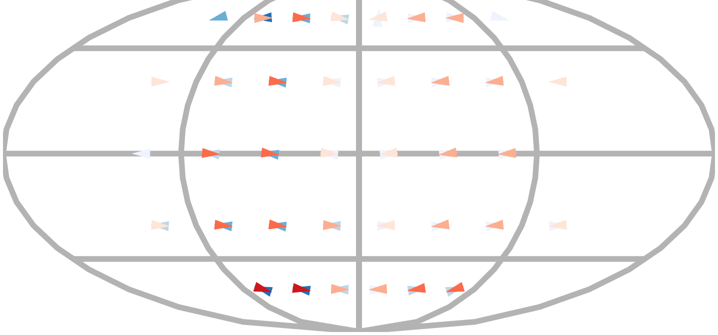

$\alpha_R$

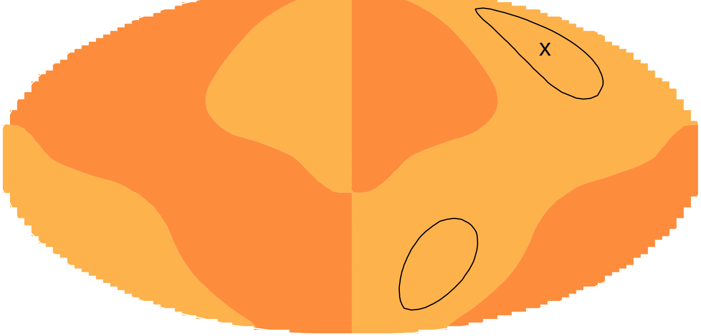


$\alpha_T$

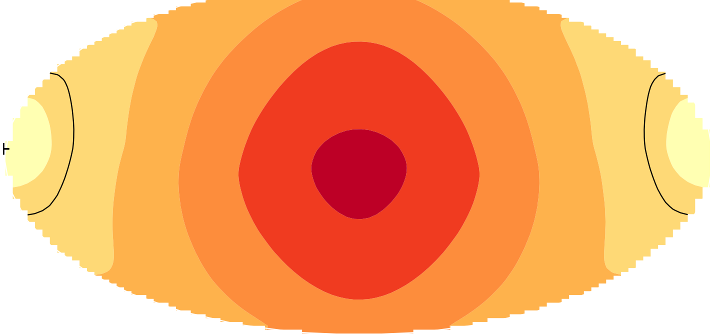

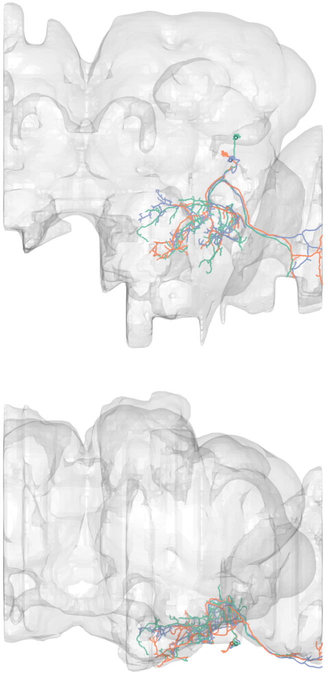

#31

bodyid =

1342777248

1563100015

1467944196

FOV = 48%

overlap of input LPTs' FOV = 25%

optimal rotation axis = [−2°,−90°]

$\alpha_{R0} = 38°$

optimal translation axis = [−82°,0°]

$\alpha_{T0} = 14°$

RTSA = 24°

input LPT neuons:

LPT28_R

LPT48_R_vCal3

LPT48_L_vCal3

LPT47_L_vCal2

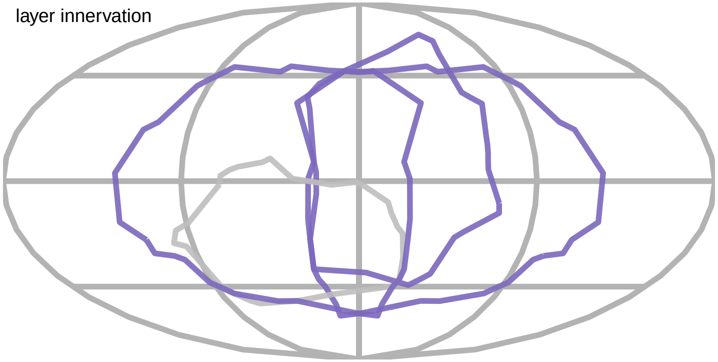


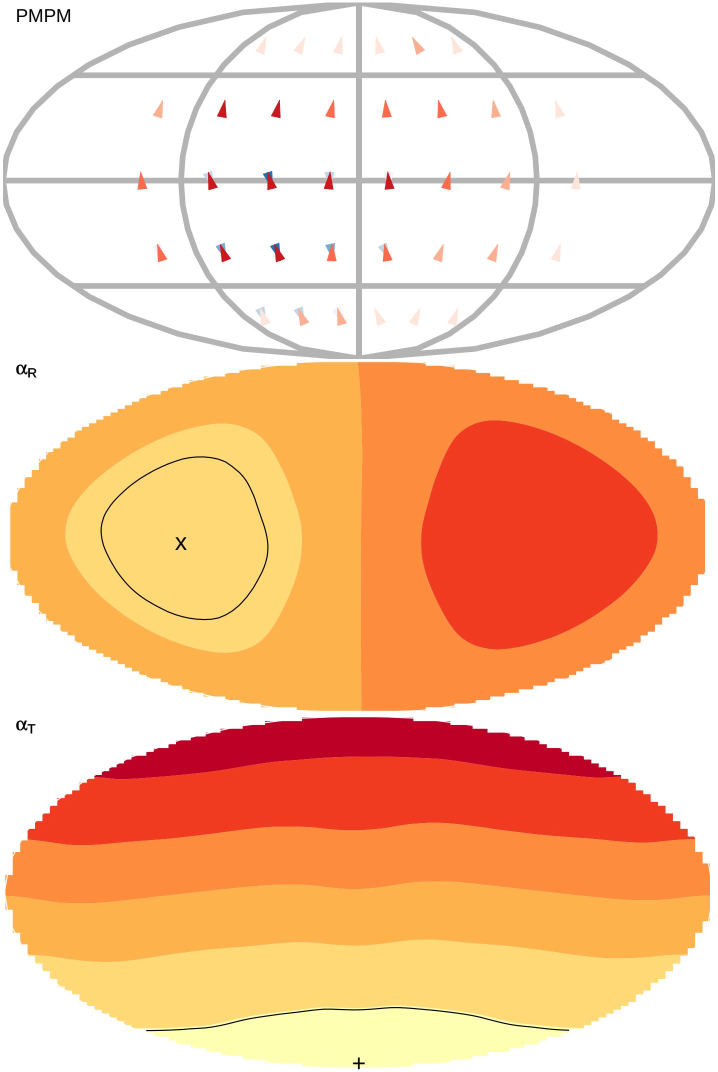

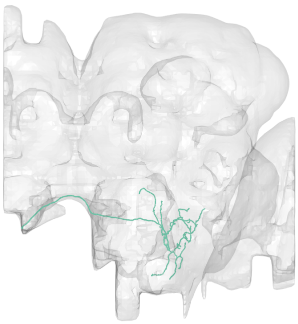

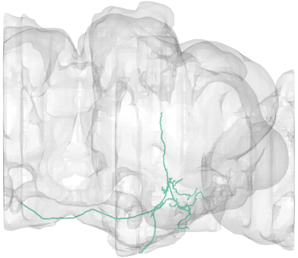

#32
bodyid =
1995952670

FOV = 49%
overlap of input LPTs' FOV = 8%
optimal rotation axis = [−74°,58°]
$\alpha_{R0} = 64°$
optimal translation axis = [4°,144°]
$\alpha_{T0} = 21°$
RTSA = 43°

input LPT neuons:
LPT21_R
LPT41_R_Nod3
LPT42_L_Nod4

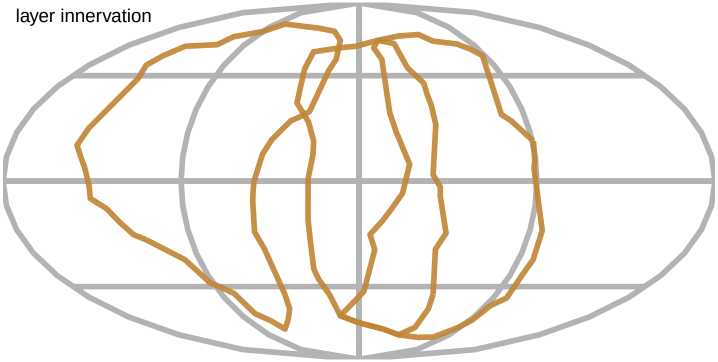


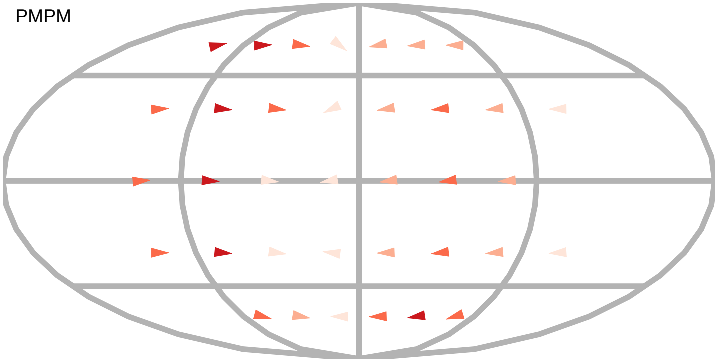


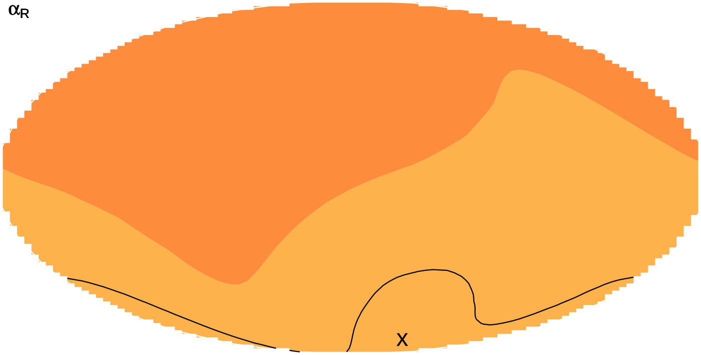


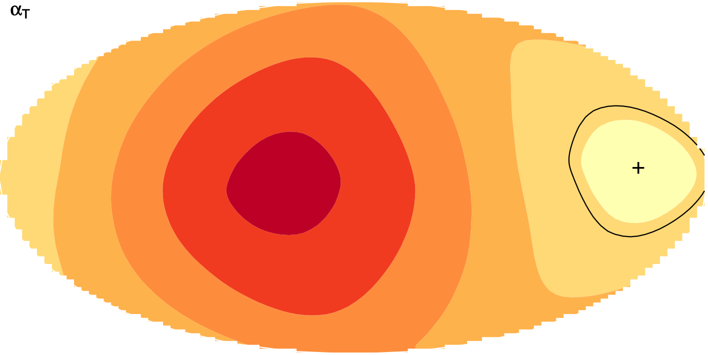

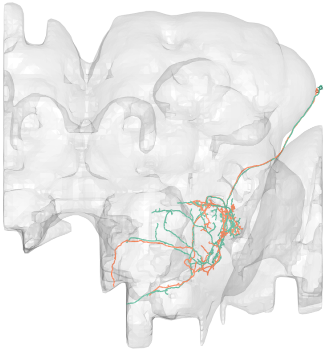

#33
bodyid =
1471337816
1471337463

FOV = 49%
overlap of input LPTs' FOV = 17%
optimal rotation axis = [0°,−90°]
$\alpha_{R0} = 39°$
optimal translation axis = [−80°,0°]
$\alpha_{T0} = 14°$
RTSA = 25°

input LPT neuons:
LPT48_R_vCal3
LPT48_L_vCal3
LPT46_L_vCal1

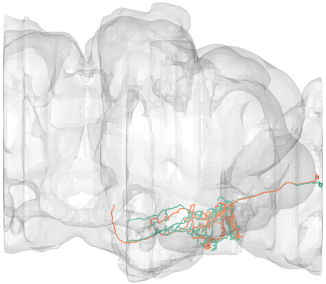

layer innervation

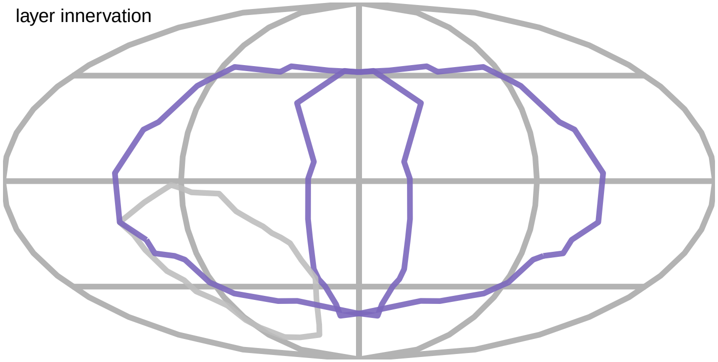

PMPM

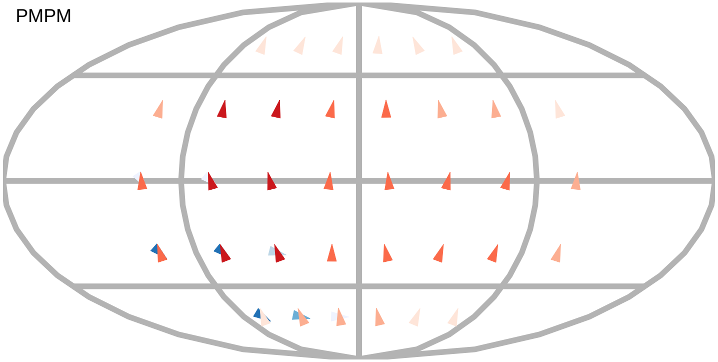

$\alpha_R$

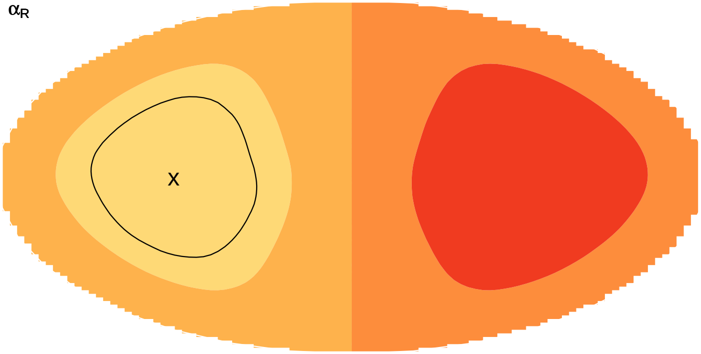


$\alpha_T$

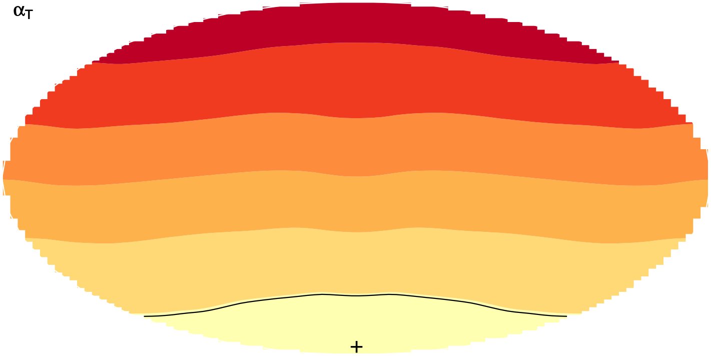

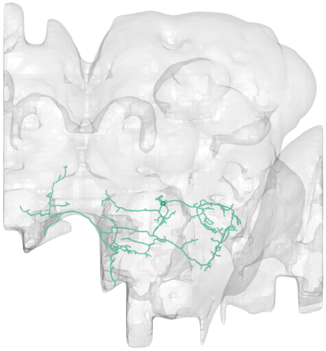

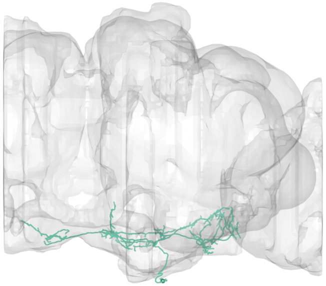

#34
bodyid =
1623253338

FOV = 49%
overlap of input LPTs' FOV = 17%
optimal rotation axis = [0°,−90°]
$\alpha_{R0} = 39°$
optimal translation axis = [−80°,0°]
$\alpha_{T0} = 14°$
RTSA = 25°

input LPT neuons:
LPT48_R_vCal3
LPT48_L_vCal3
LPT46_L_vCal1

layer innervation

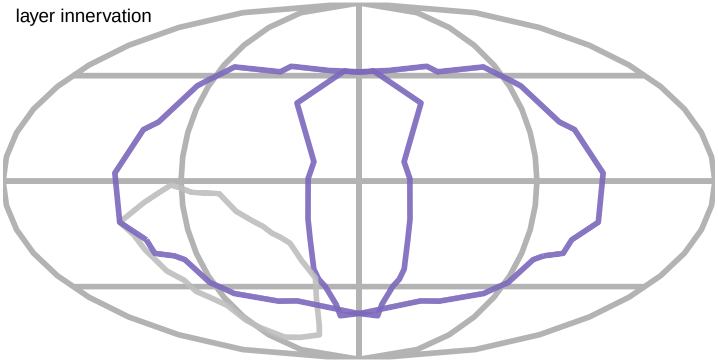

PMPM

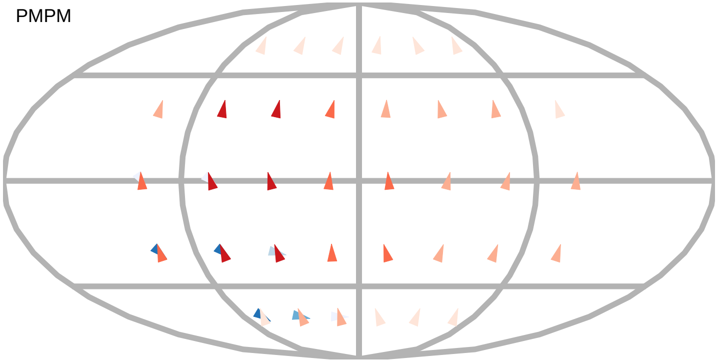

$\alpha_R$

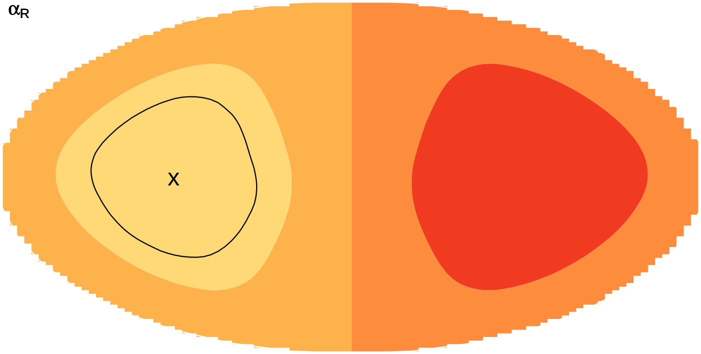


$\alpha_T$

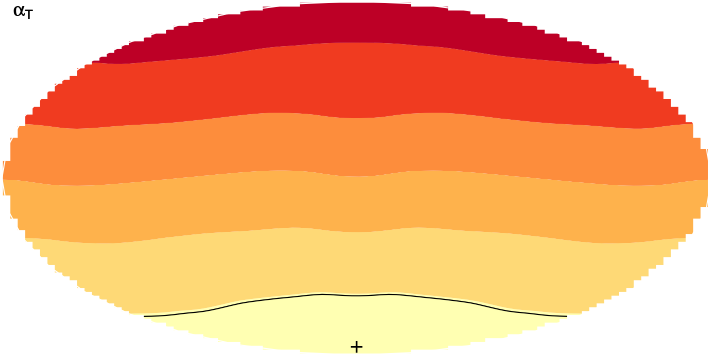

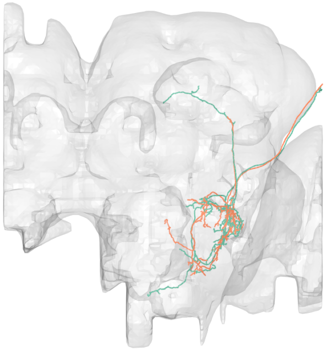

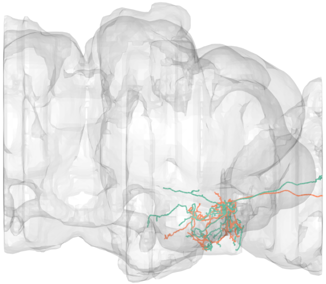

#35
bodyid =
1686233836
1471337908

FOV = 49%
overlap of input LPTs' FOV = 17%
optimal rotation axis = [0°,−90°]
$\alpha_{R0} = 39°$
optimal translation axis = [−80°,0°]
$\alpha_{T0} = 14°$
RTSA = 25°

input LPT neuons:
LPT48_R_vCal3
LPT48_L_vCal3
LPT46_L_vCal1

layer innervation

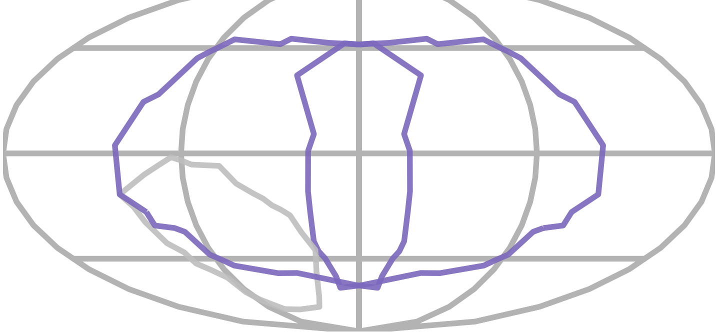

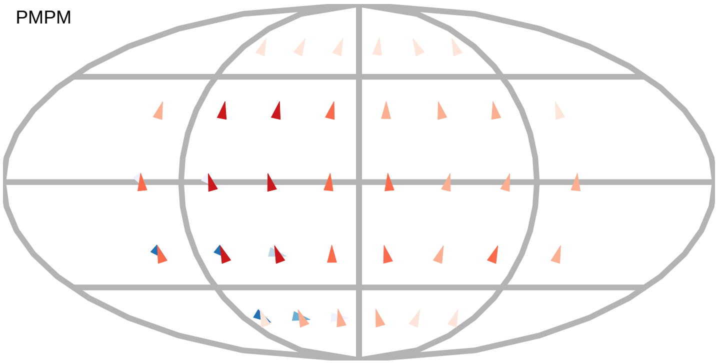


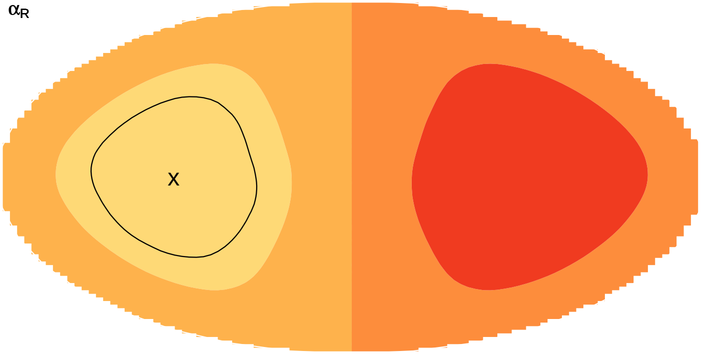


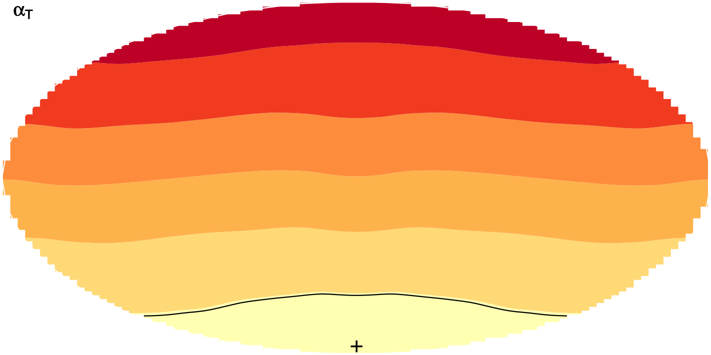

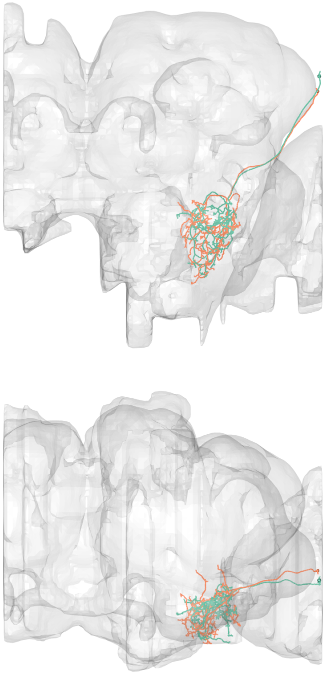

#36
bodyid =
1471337938
1471337364

FOV = 49%
overlap of input LPTs' FOV = 17%
optimal rotation axis = [0°,−90°]
$\alpha_{R0} = 39°$
optimal translation axis = [−80°,0°]
$\alpha_{T0} = 14°$
RTSA = 25°

input LPT neuons:
LPT48_R_vCal3
LPT48_L_vCal3
LPT46_L_vCal1

layer innervation

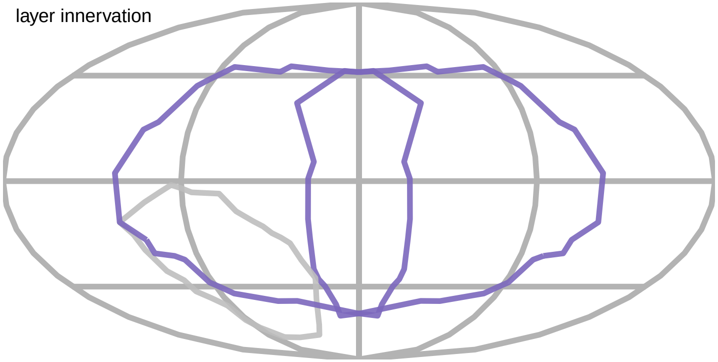

PMPM

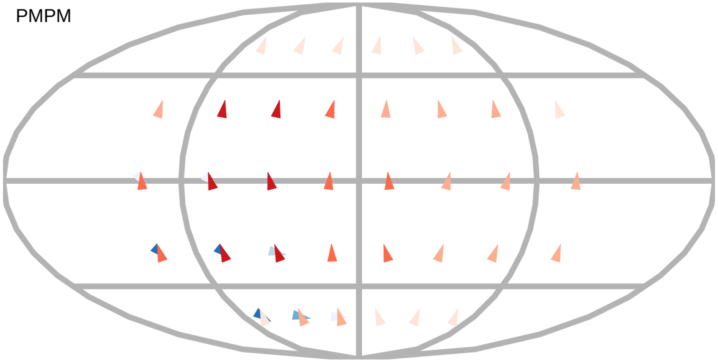

$\alpha_R$

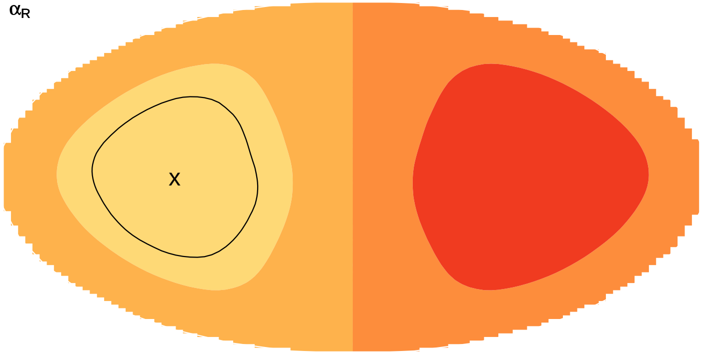


$\alpha_T$

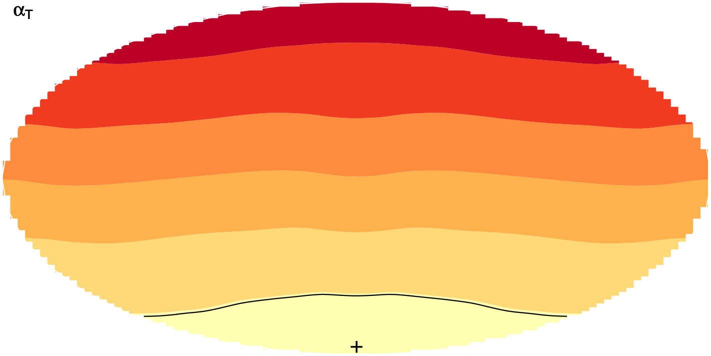

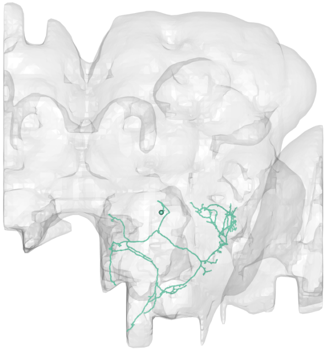

#37

bodyid =

1717290865

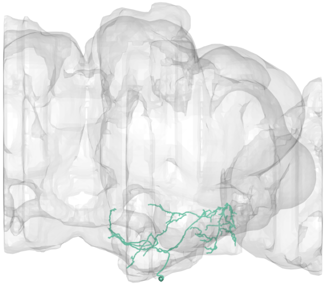

FOV = 49%

overlap of input LPTs' FOV = 17%

optimal rotation axis = [0°,−90°]

$\alpha_{R0} = 40°$

optimal translation axis = [−80°,0°]

$\alpha_{T0} = 17°$

RTSA = 23°

input LPT neuons:

LPT27_R

LPT48_R_vCal3

LPT48_L_vCal3

layer innervation

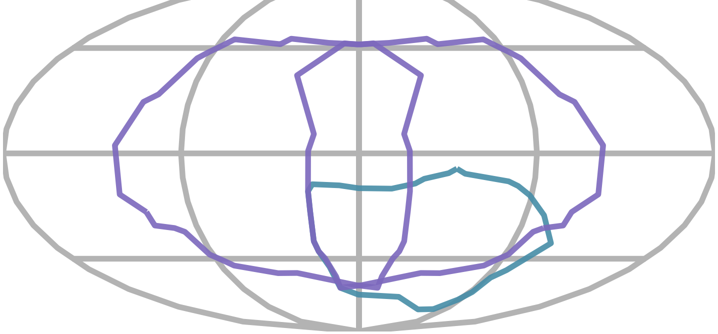

PMPM

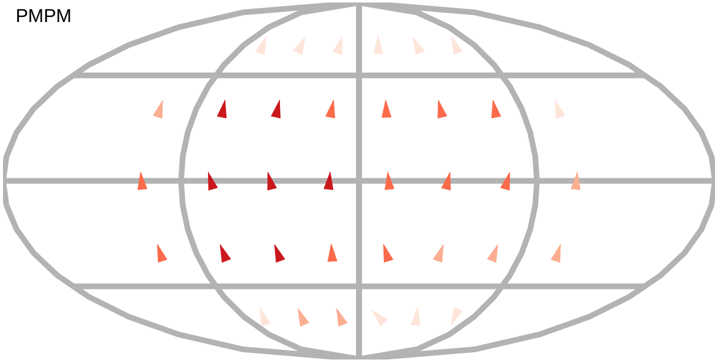

$\alpha_R$

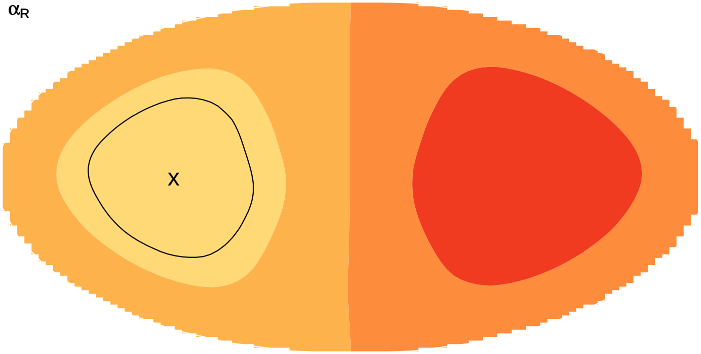

$\alpha_T$

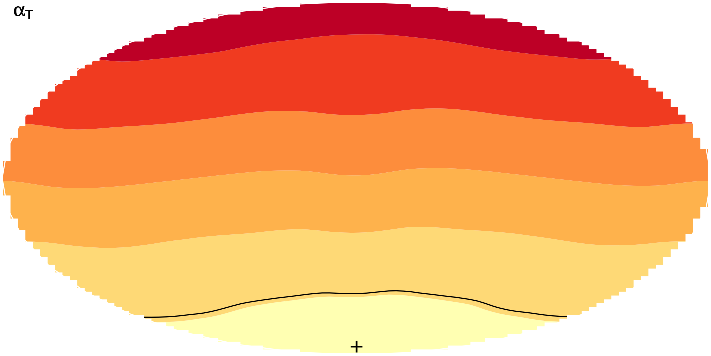

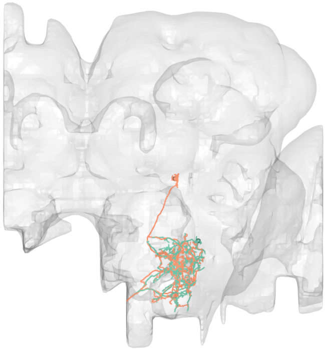

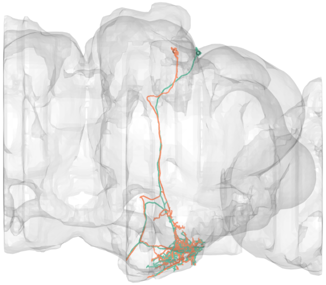

#38
bodyid =
1963869286
5813022608

FOV = 51%
overlap of input LPTs' FOV = 25%
optimal rotation axis = [74°,−131°]
$\alpha_{R0} = 16°$
optimal translation axis = [−14°,−89°]
$\alpha_{T0} = 39°$
RTSA = −23°

input LPT neuons:
LPT04_R_HST
LPT31_R
LPT43_L_H2
LPT44_L_Nod5

layer innervation

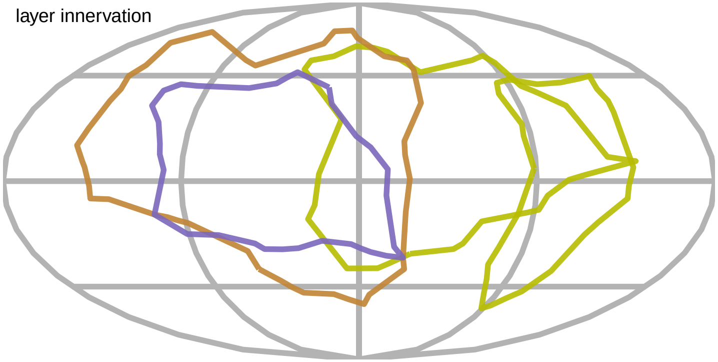

PMPM

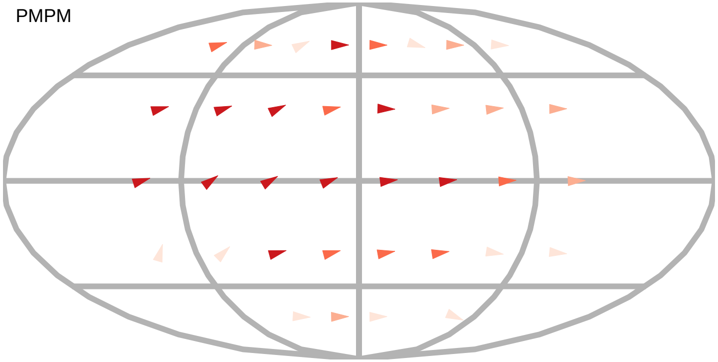

$\alpha_R$

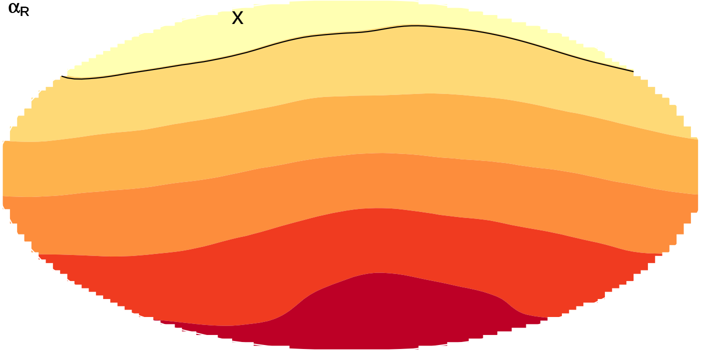

$\alpha_T$

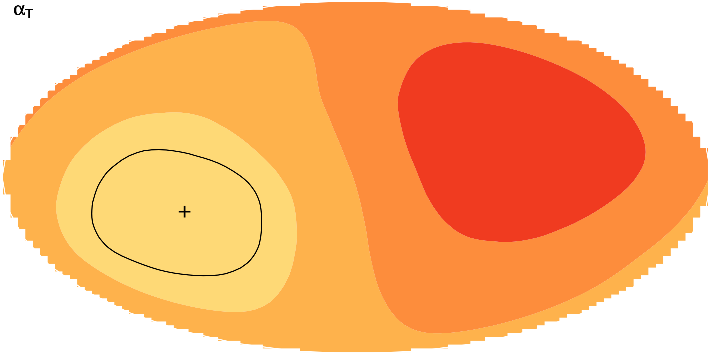

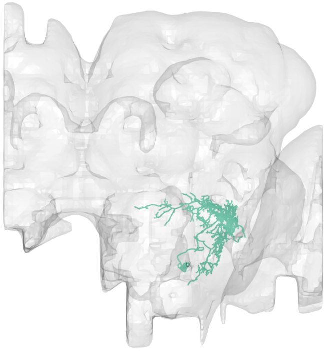

#39
bodyid =
1654879954

FOV = 51%
overlap of input LPTs' FOV = 26%
optimal rotation axis = [36°,54°]
$\alpha_{R0} = 63°$
optimal translation axis = [60°,−4°]
$\alpha_{T0} = 44°$
RTSA = 19°

input LPT neuons:
LPT26_R
LPT27_R
LPT30_R
LPT49_R
LPT49_L

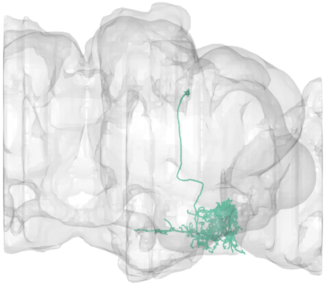

layer innervation

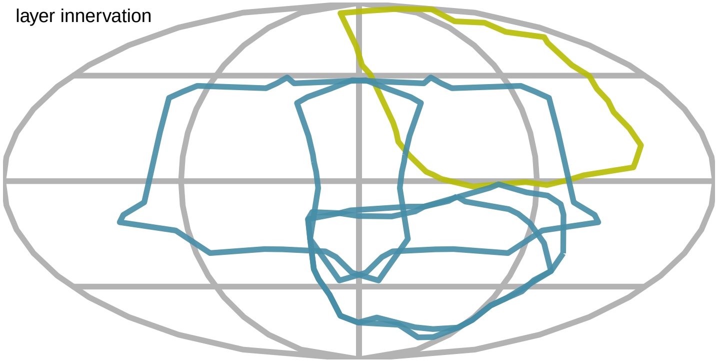

PMPM

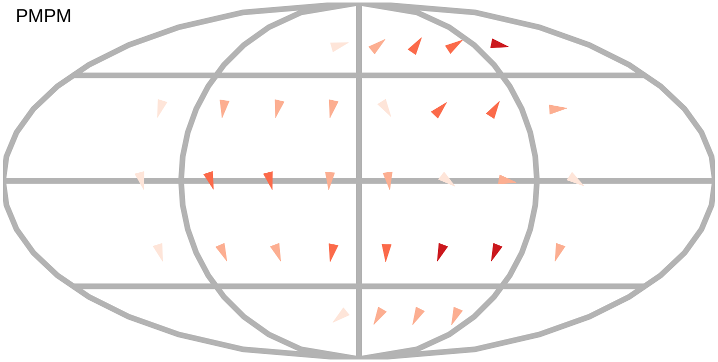

$\alpha_R$

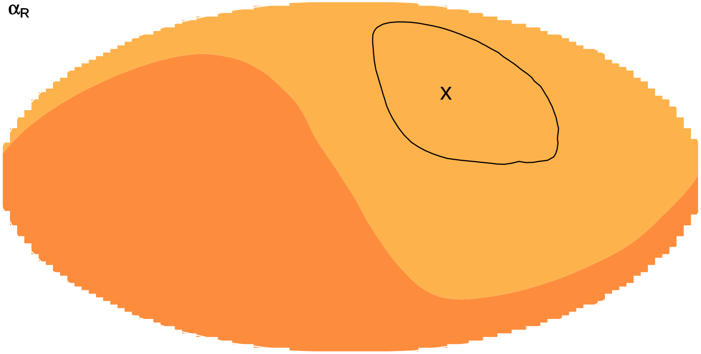


$\alpha_T$

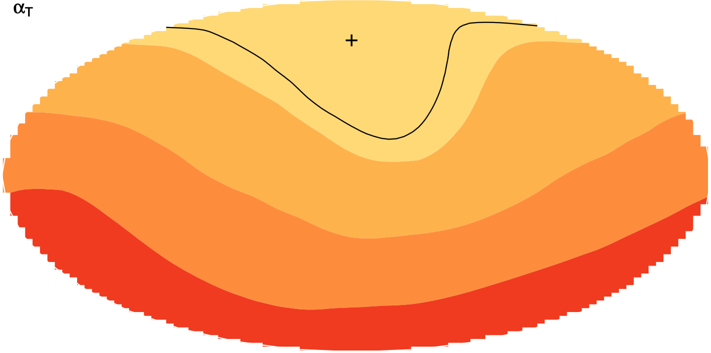

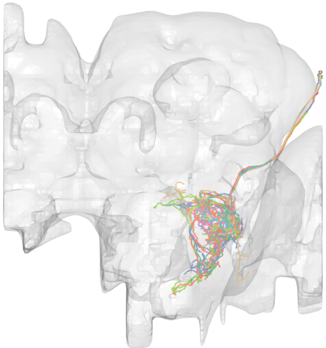

#40

bodyid =

1471337767
1654866496
1471337439
1378544024
1471337887
1687230765

FOV = 51%
overlap of input LPTs' FOV = 23%
optimal rotation axis = [0°,−90°]
$\alpha_{R0} = 39°$
optimal translation axis = [−80°,0°]
$\alpha_{T0} = 14°$
RTSA = 25°

input LPT neuons:
LPT48_R_vCal3
LPT48_L_vCal3
LPT46_R_vCal1
LPT46_L_vCal1

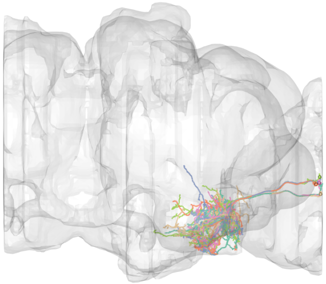

layer innervation

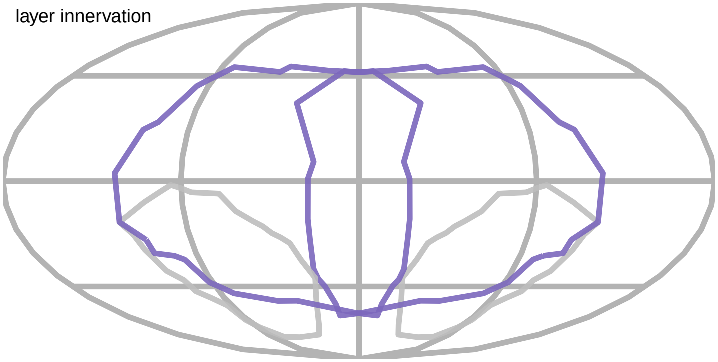

PMPM

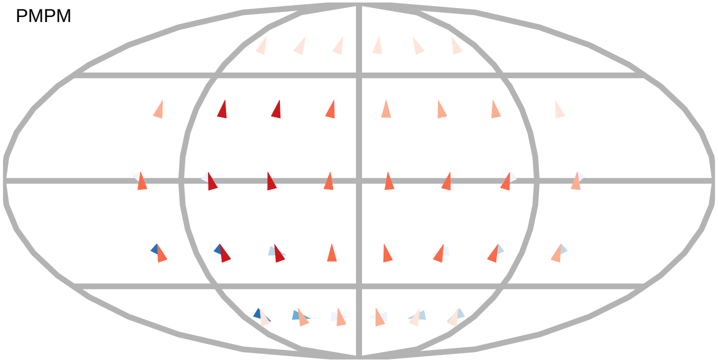

$\alpha_R$

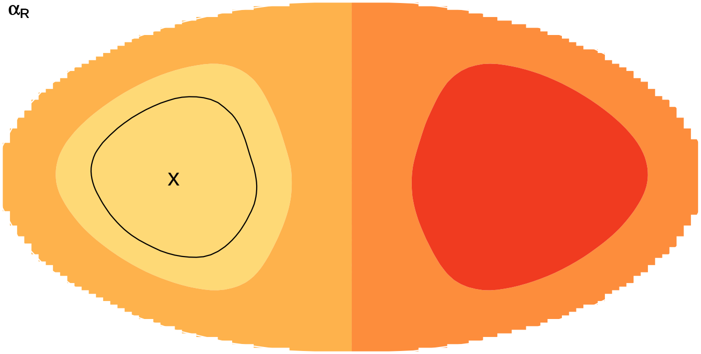


$\alpha_T$

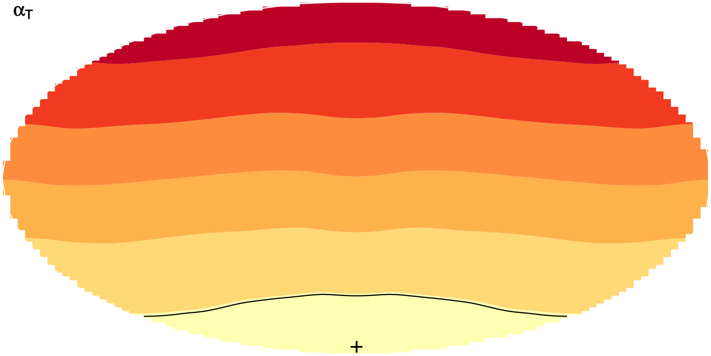

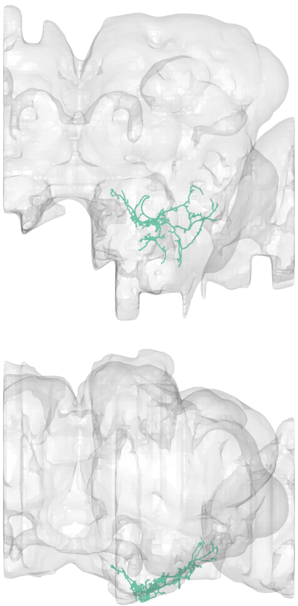

#41
bodyid =
5813088863

FOV = 51%
overlap of input LPTs' FOV = 34%
optimal rotation axis = [16°,−102°]
$\alpha_{R0} = 40°$
optimal translation axis = [−72°,−39°]
$\alpha_{T0} = 23°$
RTSA = 17°

input LPT neuons:
LPT04_R_HST
LPT17_R_VST2
LPT48_R_vCal3
LPT48_L_vCal3
LPT47_L_vCal2

PMPM

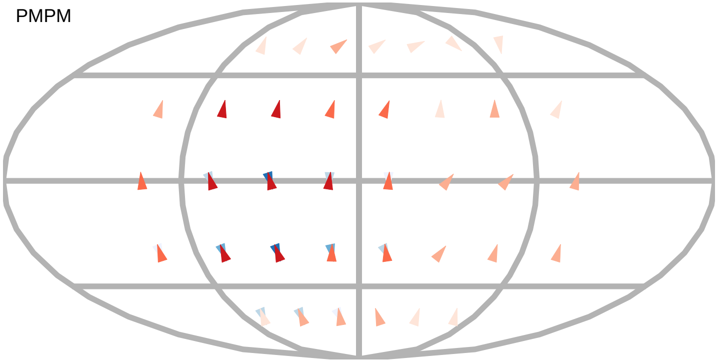

$\alpha_R$

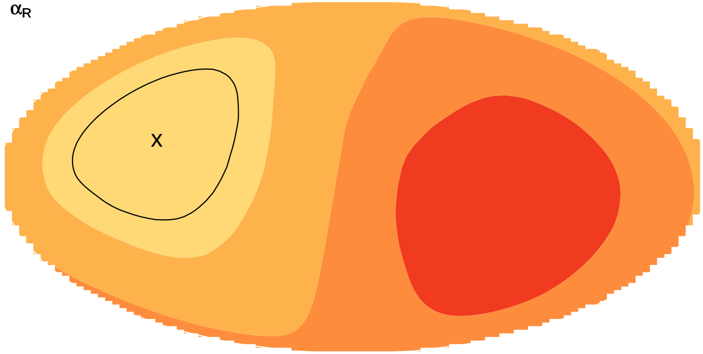


layer innervation

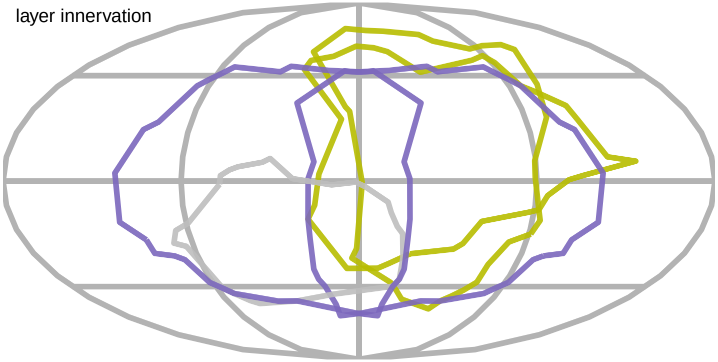

$\alpha_T$

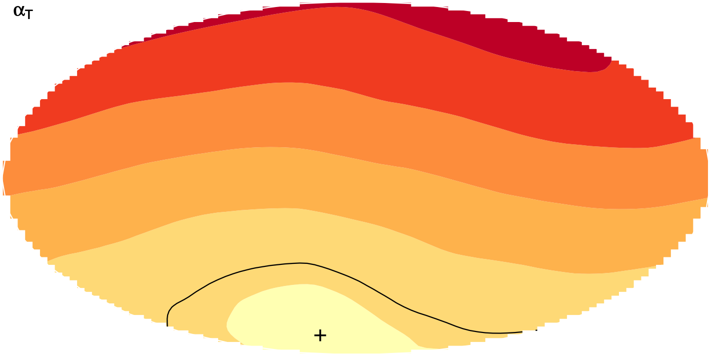

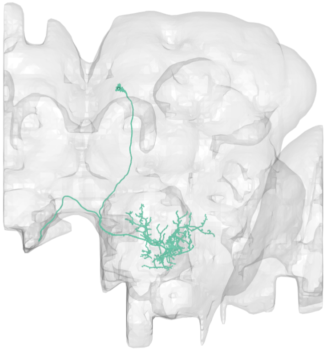

#42
bodyid =
1685020627

FOV = 51%
overlap of input LPTs' FOV = 29%
optimal rotation axis = [−4°,−120°]
$\alpha_{R0} = 36°$
optimal translation axis = [−82°,−14°]
$\alpha_{T0} = 39°$
RTSA = −3°

input LPT neuons:
LPT09_R_VS5
LPT17_R_VST2
LPT28_R
LPT48_R_vCal3
LPT48_L_vCal3

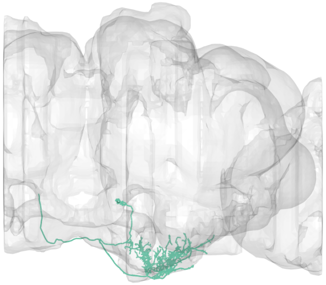

layer innervation

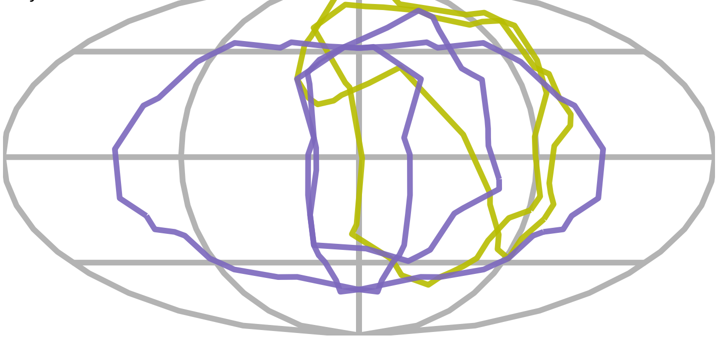

PMPM

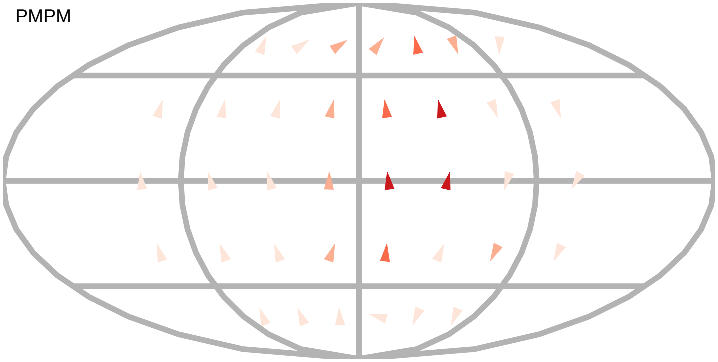

$\alpha_R$

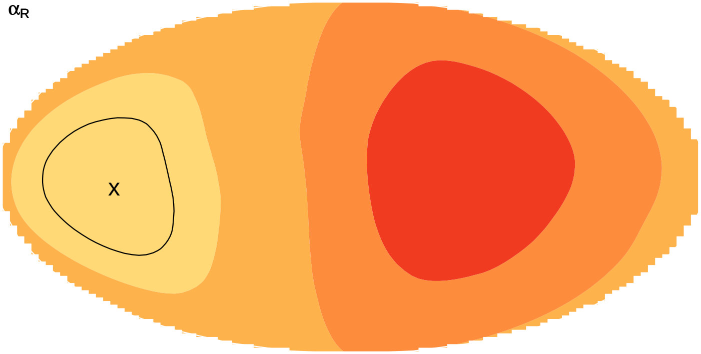


$\alpha_T$

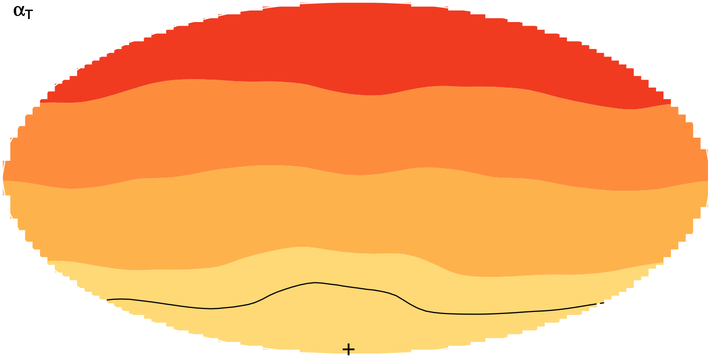

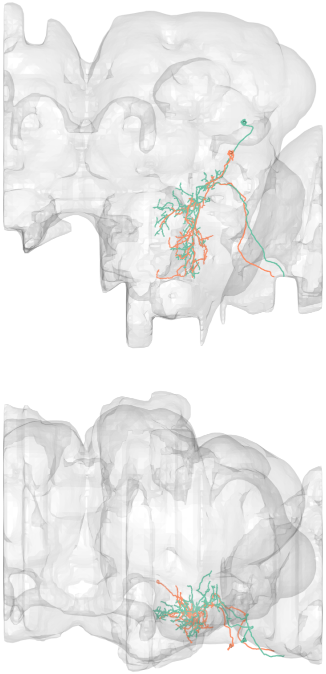

#43
bodyid =
5812993576
1563790913

FOV = 52%
overlap of input LPTs' FOV = 27%
optimal rotation axis = [52°,−42°]
$\alpha_{R0} = 56°$
optimal translation axis = [8°,−176°]
$\alpha_{T0} = 63°$
RTSA = −7°

input LPT neuons:
LPT26_R
LPT41_R_Nod3
LPT41_L_Nod3
LPT40_L_Nod2

layer innervation

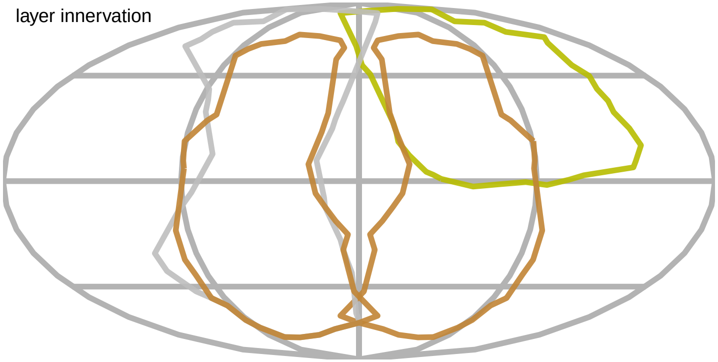

PMPM

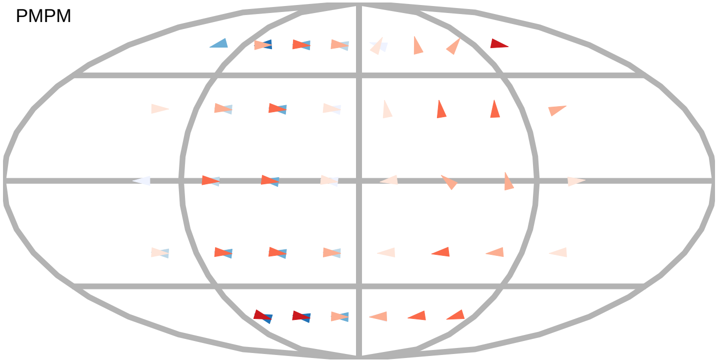

$\alpha_R$

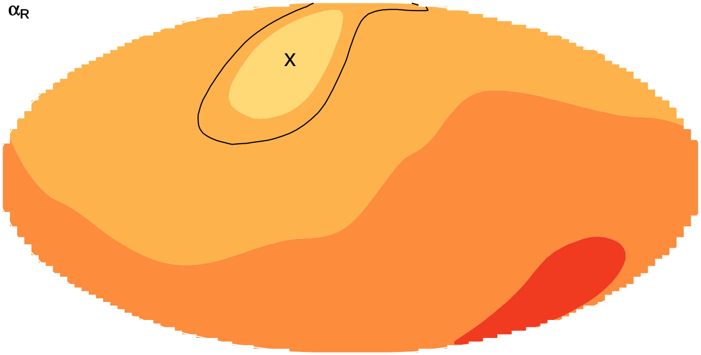


$\alpha_T$

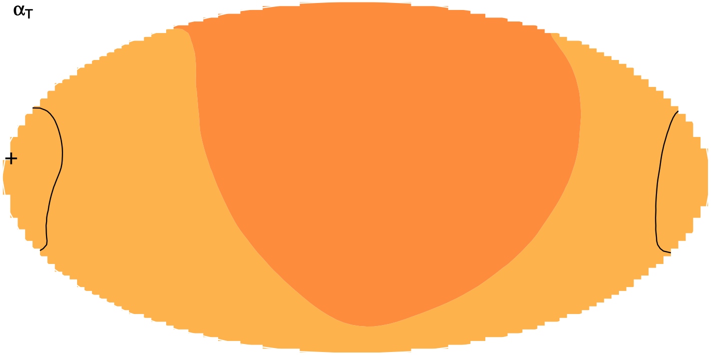

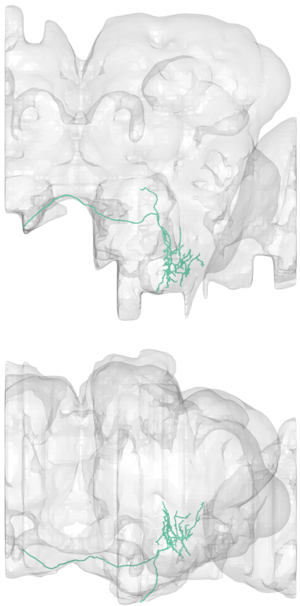

#44
bodyid =
5813133658

FOV = 54%
overlap of input LPTs' FOV = 21%
optimal rotation axis = [84°,77°]
$\alpha_{R0} = 4°$
optimal translation axis = [6°,−149°]
$\alpha_{T0} = 12°$
RTSA = −8°

input LPT neuons:
LPT41_L_Nod3
LPT42_L_Nod4
LPT40_R_Nod2
LPT40_L_Nod2

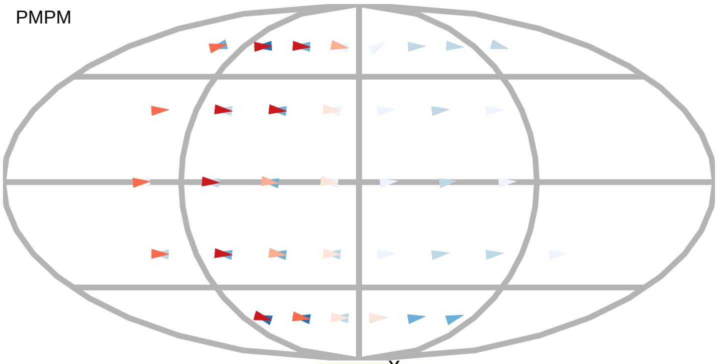


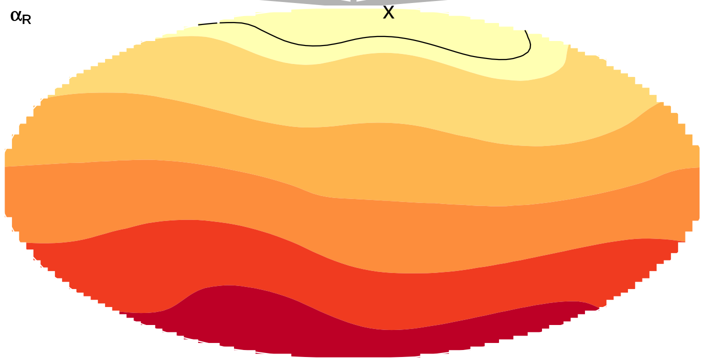


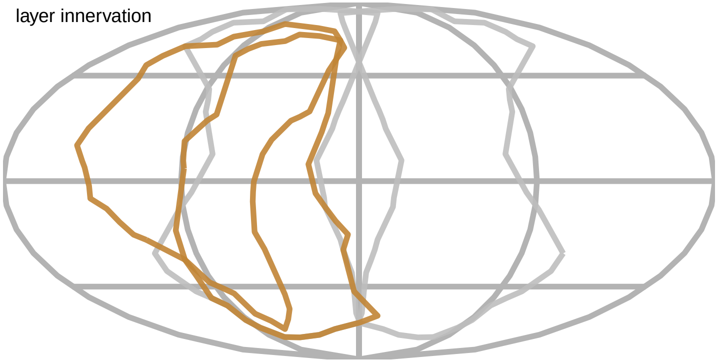


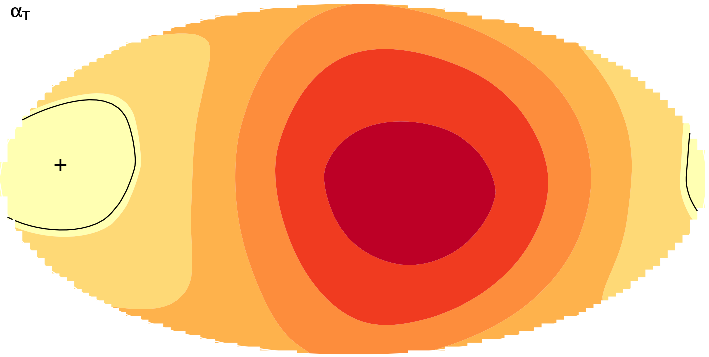

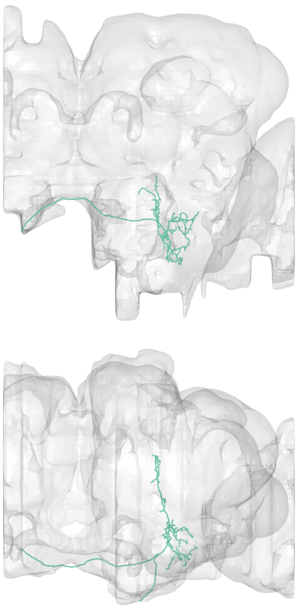

#45
bodyid =
1849974592

FOV = 55%
overlap of input LPTs' FOV = 20%
optimal rotation axis = [−52°,42°]
$\alpha_{R0} = 76°$
optimal translation axis = [2°,158°]
$\alpha_{T0} = 29°$
RTSA = 47°

input LPT neuons:
LPT21_R
LPT41_R_Nod3
LPT41_L_Nod3
LPT42_L_Nod4

PMPM

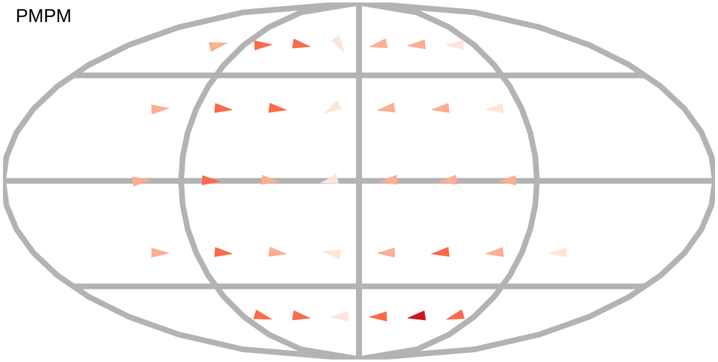

$\alpha_R$

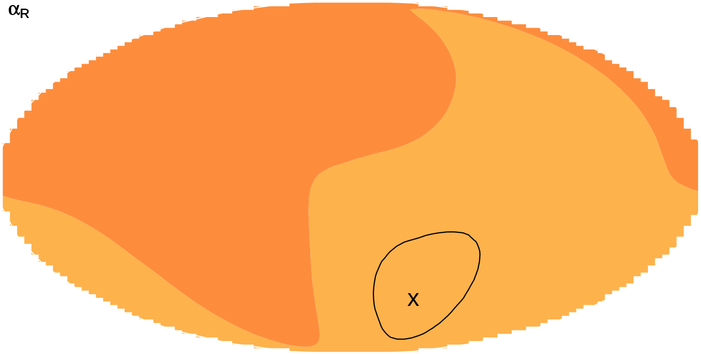

layer innervation

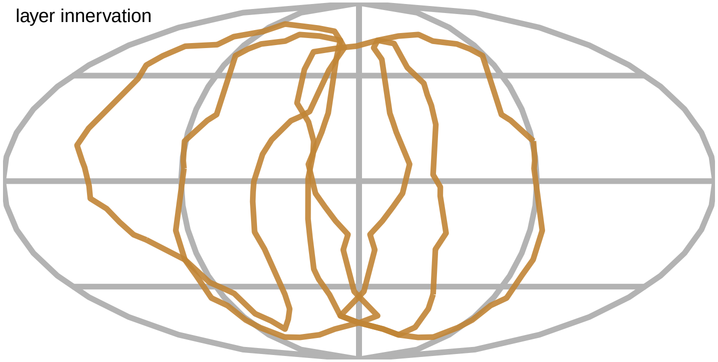

$\alpha_T$

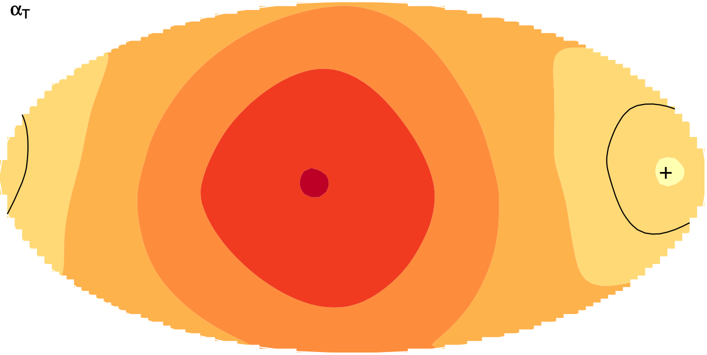

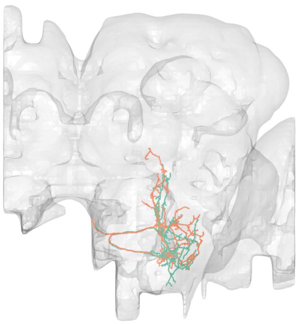

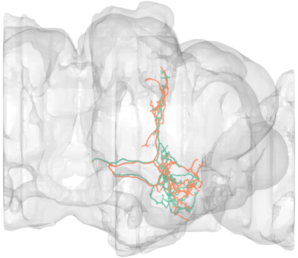

#46
bodyid =
2215550458
2153454486

FOV = 55%
overlap of input LPTs' FOV = 31%
optimal rotation axis = [−58°,−159°]
$\alpha_{R0}$ = 77°
optimal translation axis = [2°,166°]
$\alpha_{T0}$ = 17°
RTSA = 60°

input LPT neuons:
LPT41_R_Nod3
LPT42_L_Nod4
LPT40_R_Nod2
LPT40_L_Nod2

PMPM

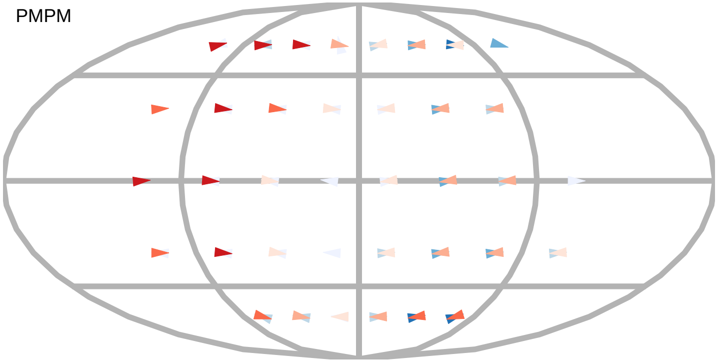

$\alpha_R$

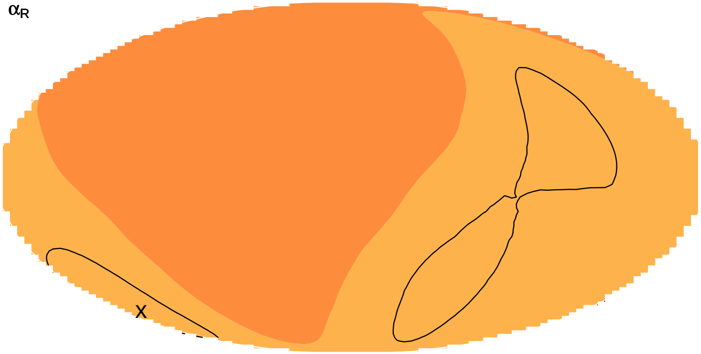

$\alpha_T$

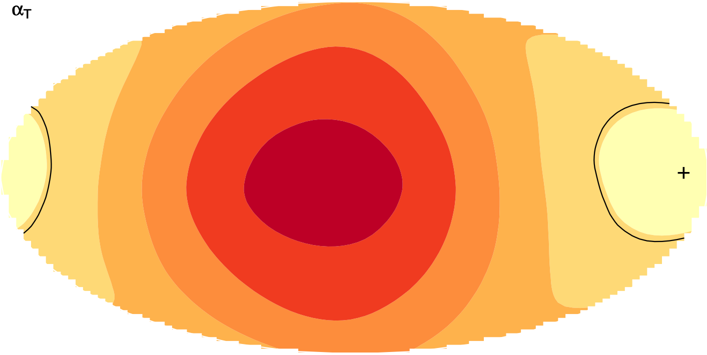

layer innervation

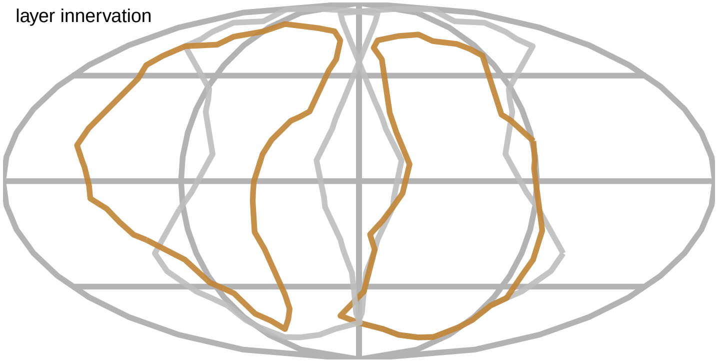

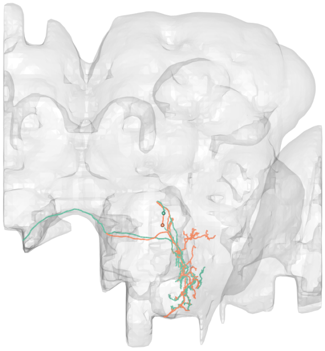

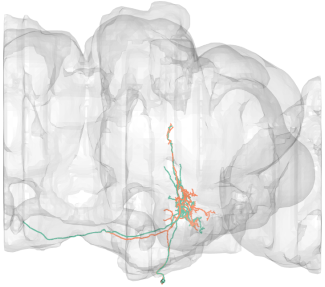

#47
bodyid =
2000028737
1937258605

FOV = 55%
overlap of input LPTs' FOV = 40%
optimal rotation axis = [58°,117°]
$\alpha_{R0} = 71°$
optimal translation axis = [4°,−180°]
$\alpha_{T0} = 22°$
RTSA = 49°

input LPT neuons:
LPT41_R_Nod3
LPT41_L_Nod3
LPT42_L_Nod4
LPT40_R_Nod2
LPT40_L_Nod2

layer innervation

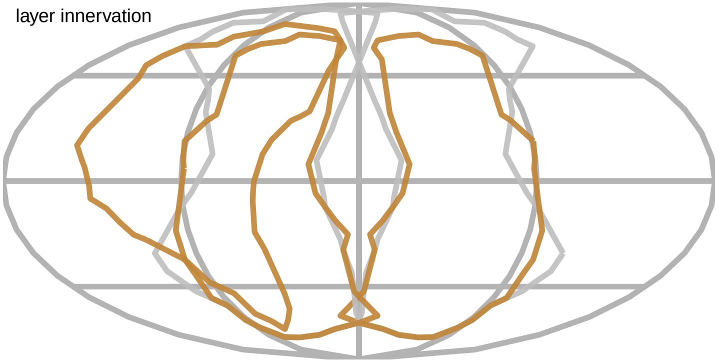

PMPM

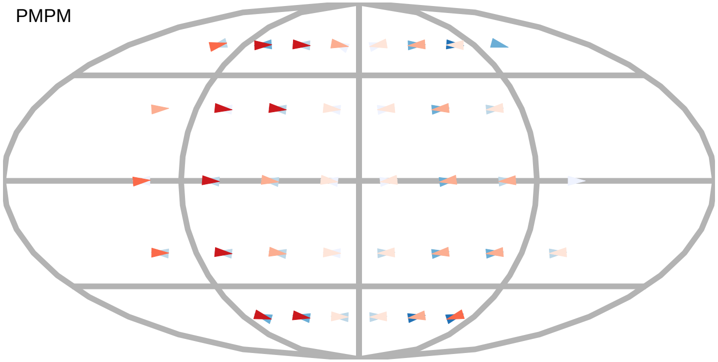

$\alpha_R$

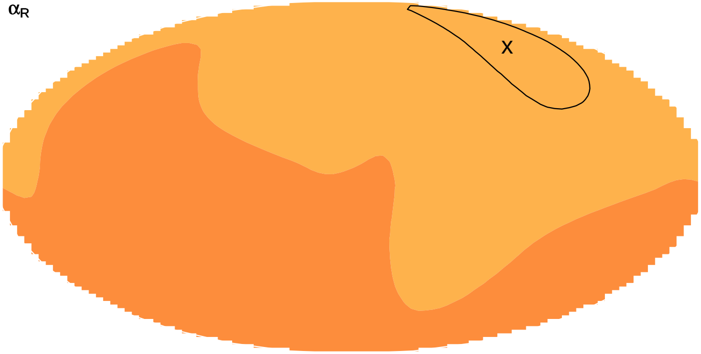


$\alpha_T$

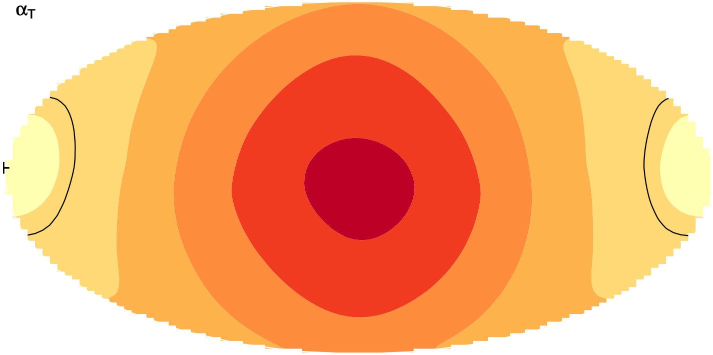

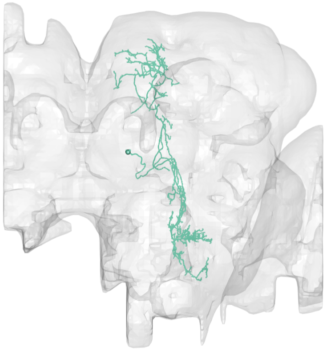

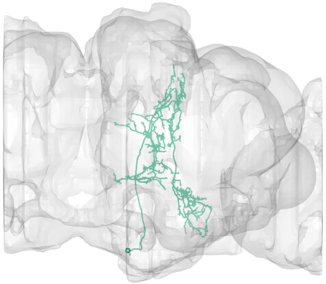

#48
bodyid =
482754144

FOV = 55%
overlap of input LPTs' FOV = 40%
optimal rotation axis = [58°,117°]
$\alpha_{R0} = 71°$
optimal translation axis = [4°,−180°]
$\alpha_{T0} = 22°$
RTSA = 49°

input LPT neuons:
LPT41_R_Nod3
LPT41_L_Nod3
LPT42_L_Nod4
LPT40_R_Nod2
LPT40_L_Nod2

layer innervation

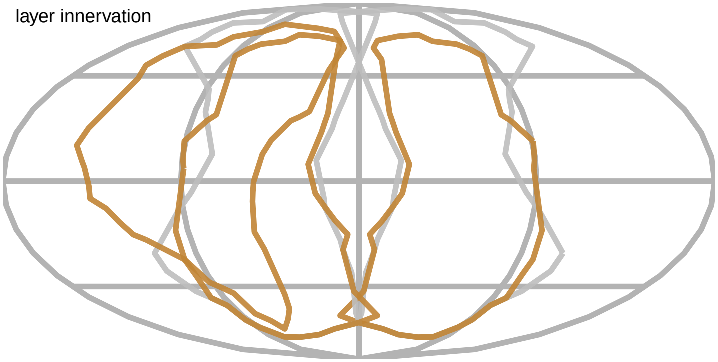

PMPM

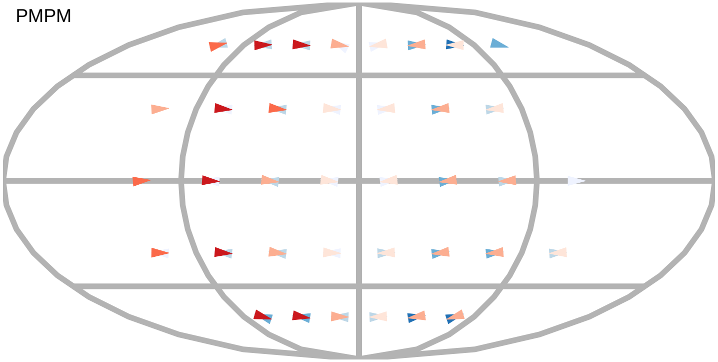

$\alpha_R$

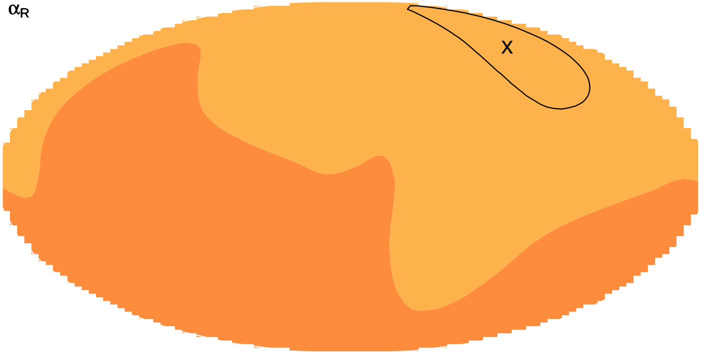


$\alpha_T$

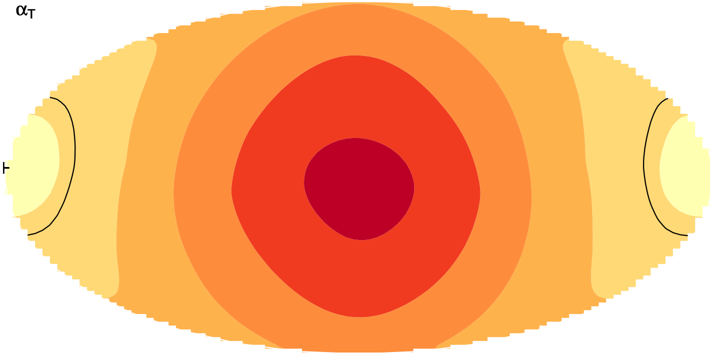

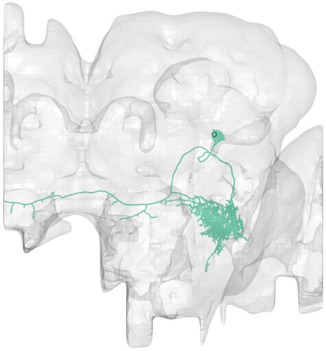

#49
bodyid =
5813022932

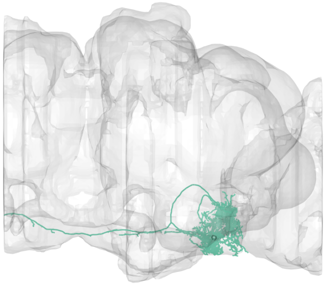

FOV = 55%
overlap of input LPTs' FOV = 33%
optimal rotation axis = [0°,−90°]
$\alpha_{R0} = 39°$
optimal translation axis = [−80°,0°]
$\alpha_{T0} = 14°$
RTSA = 25°

input LPT neuons:
LPT48_R_vCal3
LPT48_L_vCal3
LPT46_R_vCal1
LPT47_R_vCal2
LPT45_L_dCal1
LPT46_L_vCal1
LPT47_L_vCal2

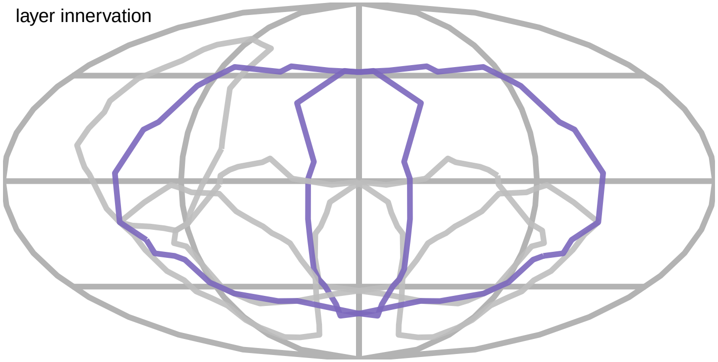


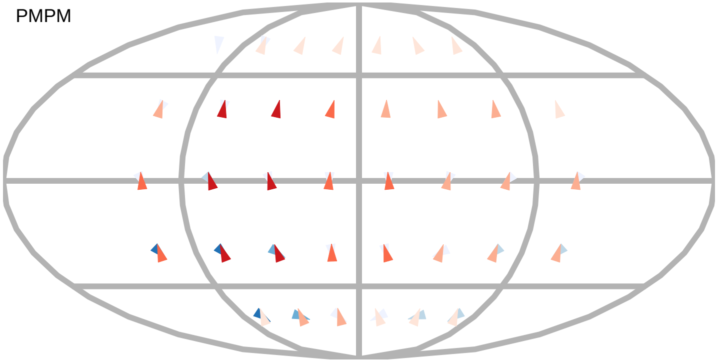


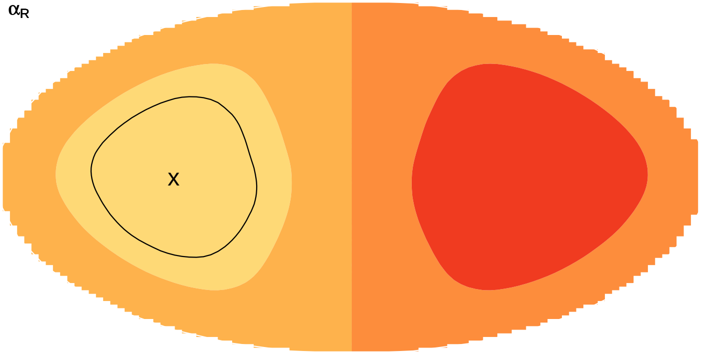


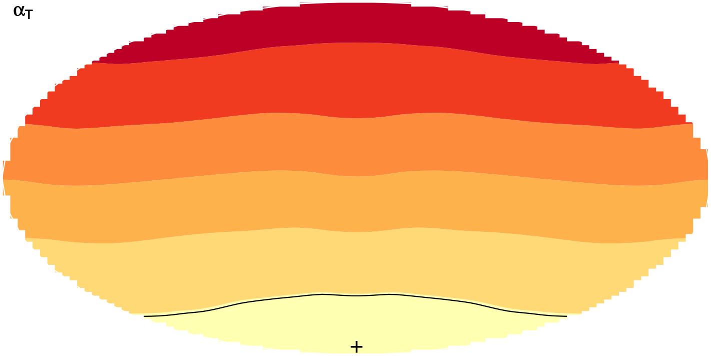

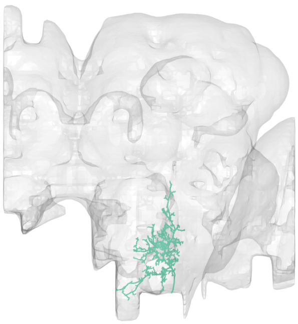

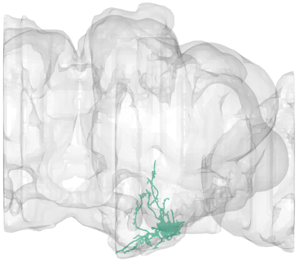

#50
bodyid =
5813024629

FOV = 56%
overlap of input LPTs' FOV = 26%
optimal rotation axis = [84°,0°]
$\alpha_{R0} = 8°$
optimal translation axis = [4°,−100°]
$\alpha_{T0} = 37°$
RTSA = −29°

input LPT neuons:
LPT03_R_HSS
LPT04_R_HST
LPT31_R
LPT43_L_H2
LPT22_R

PMPM

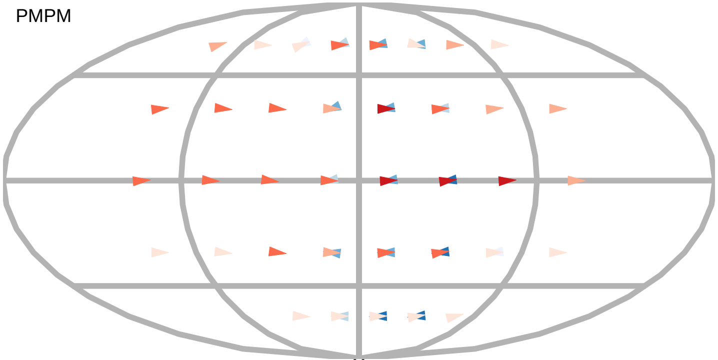

$\alpha_R$

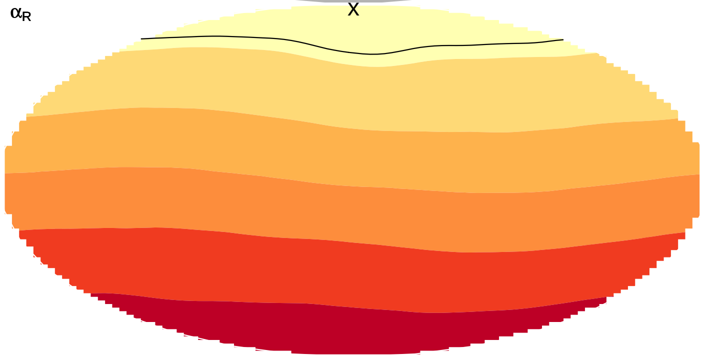


$\alpha_T$

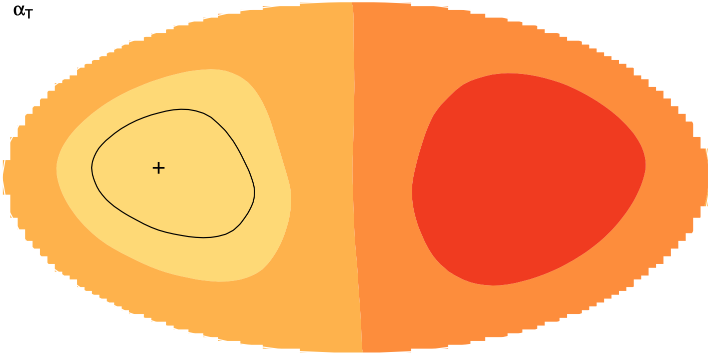

layer innervation

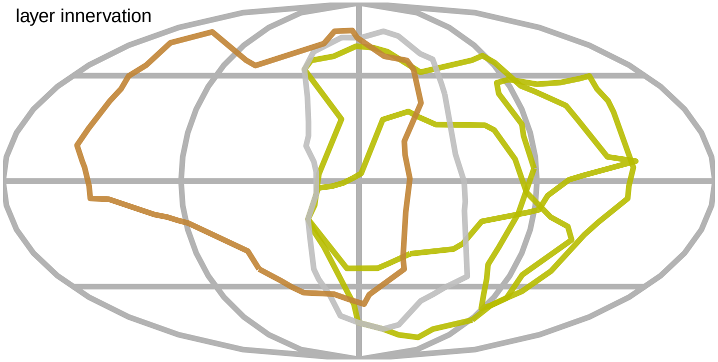

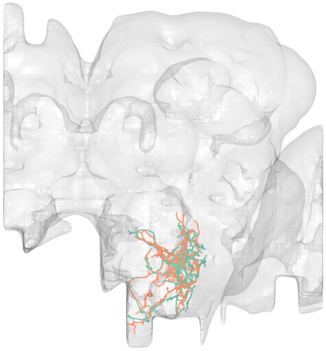

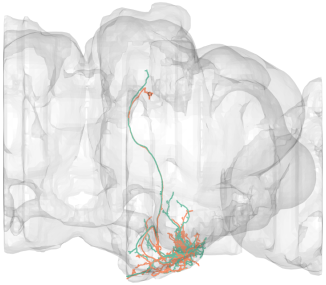

#51
bodyid =
1902144174
5813000953

FOV = 58%
overlap of input LPTs' FOV = 23%
optimal rotation axis = [86°,0°]
$\alpha_{R0} = 7°$
optimal translation axis = [0°,−86°]
$\alpha_{T0} = 38°$
RTSA = −31°

input LPT neuons:
LPT02_R_HSE
LPT03_R_HSS
LPT04_R_HST
LPT43_L_H2

layer innervation

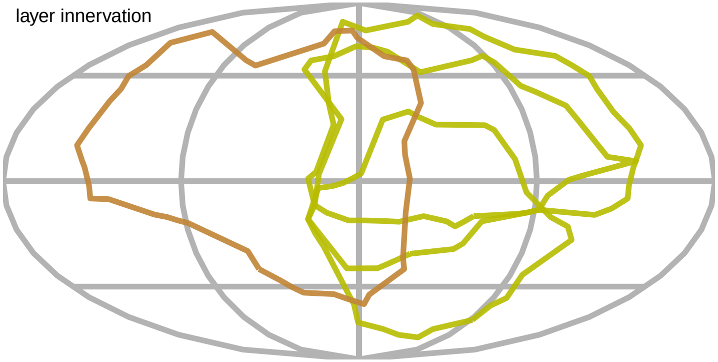

PMPM

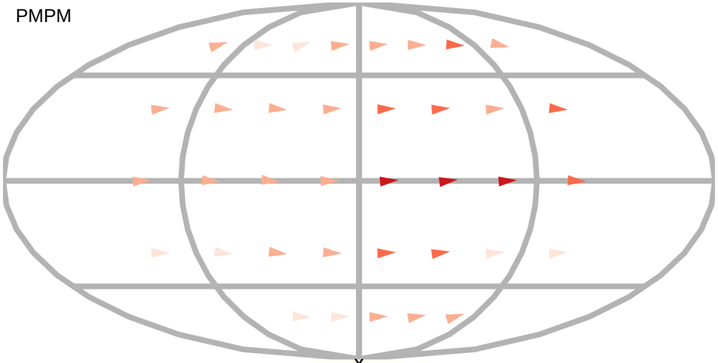

$\alpha_R$

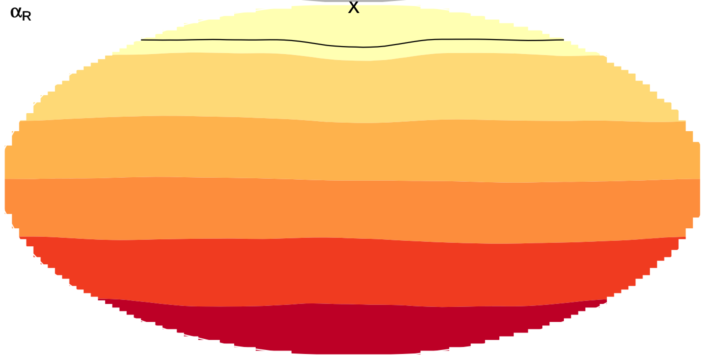

$\alpha_T$

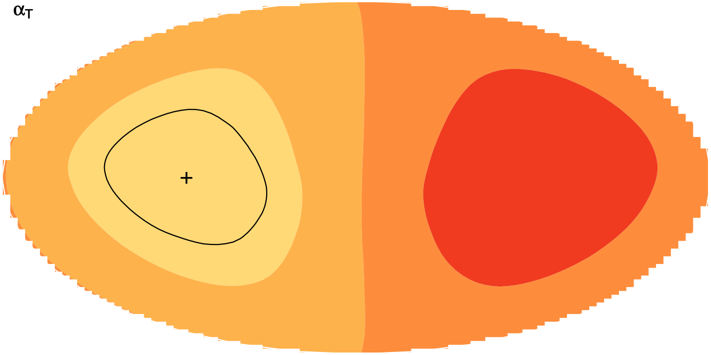

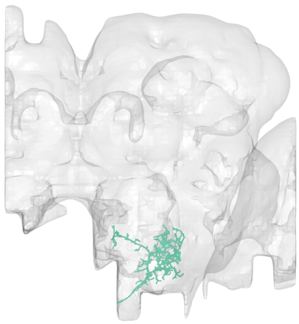

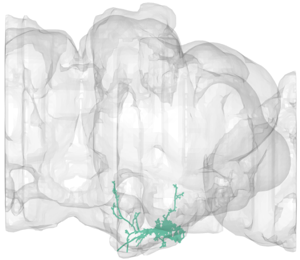

#52
bodyid =
5813078134

FOV = 58%
overlap of input LPTs' FOV = 27%
optimal rotation axis = [86°,0°]
$\alpha_{R0} = 7°$
optimal translation axis = [−4°,−80°]
$\alpha_{T0} = 37°$
RTSA = −30°

input LPT neuons:
LPT01_R_HSN
LPT02_R_HSE
LPT03_R_HSS
LPT04_R_HST
LPT43_L_H2

PMPM

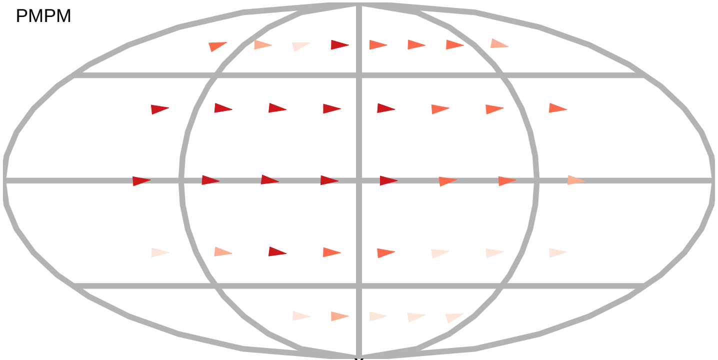

$\alpha_R$

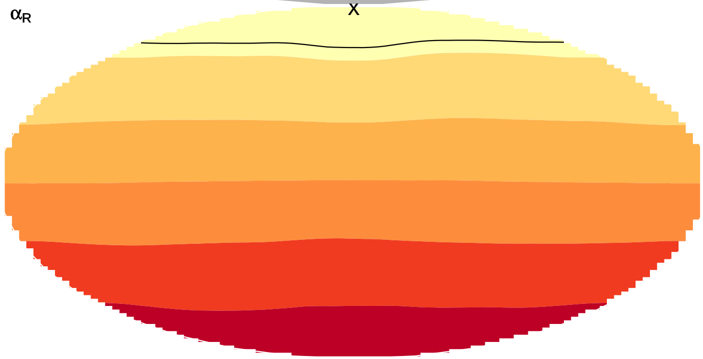

layer innervation

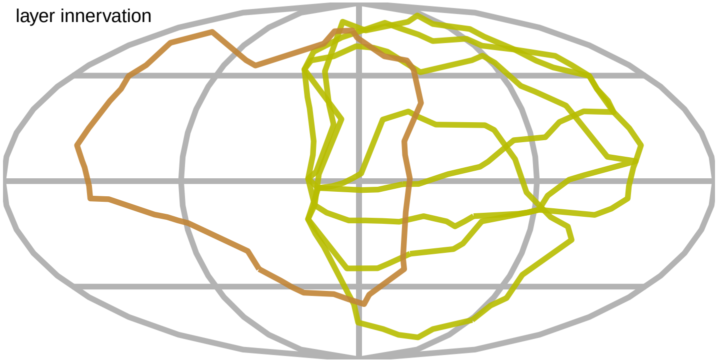

$\alpha_T$

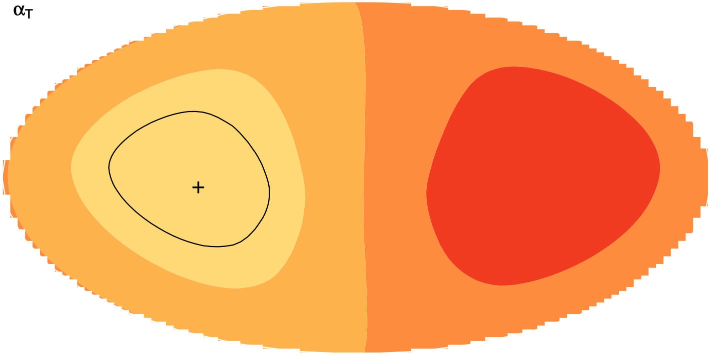

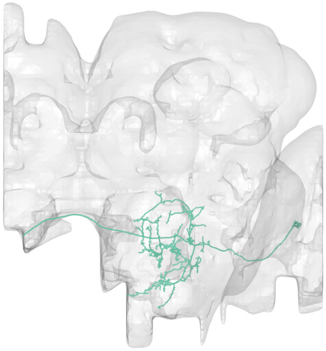

#53
bodyid =
5813013422

FOV = 58%
overlap of input LPTs' FOV = 14%
optimal rotation axis = [6°,−80°]
$\alpha_{R0} = 47°$
optimal translation axis = [−36°,17°]
$\alpha_{T0} = 39°$
RTSA = 8°

input LPT neuons:
LPT26_R
LPT27_R
LPT28_R
LPT50_L

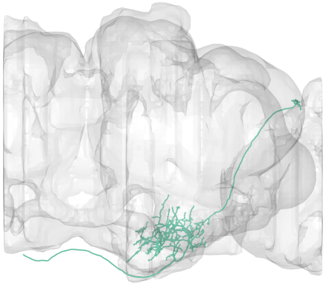

PMPM

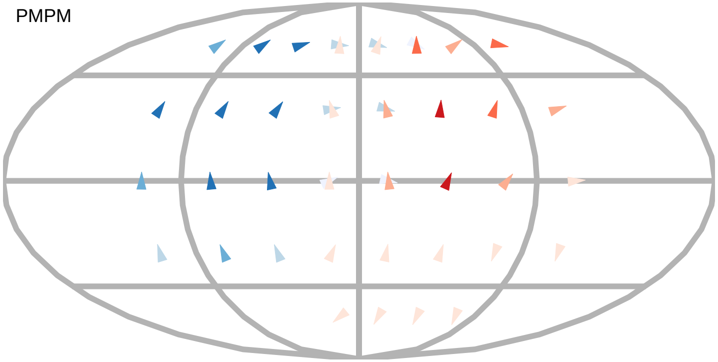

$\alpha_R$

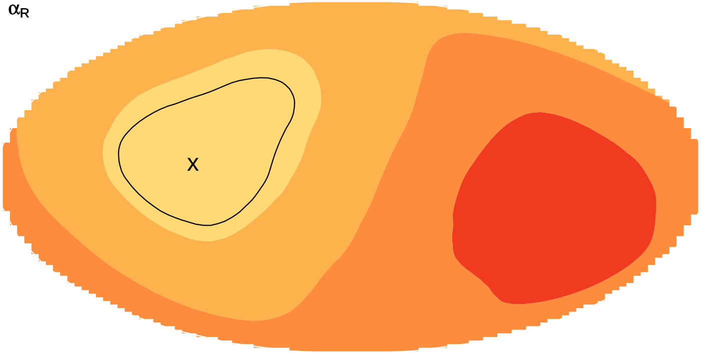

layer innervation

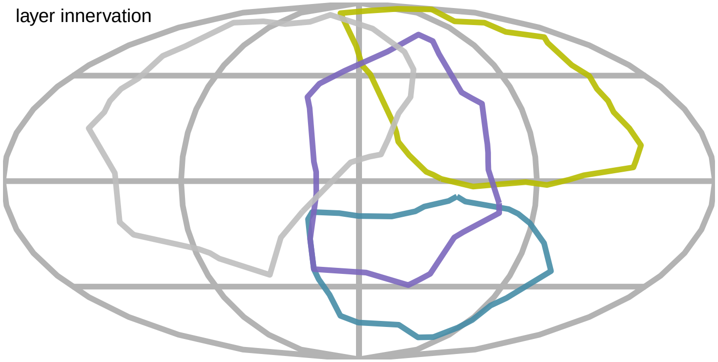

$\alpha_T$

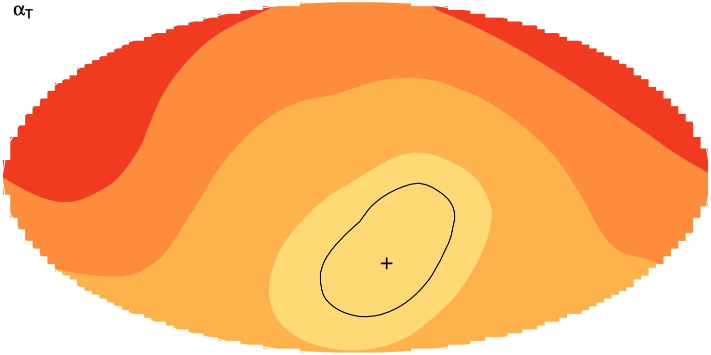

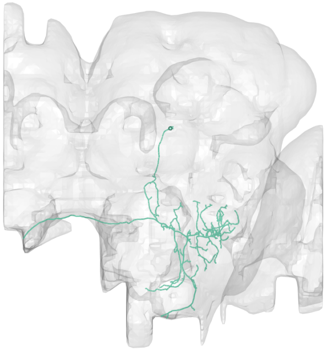

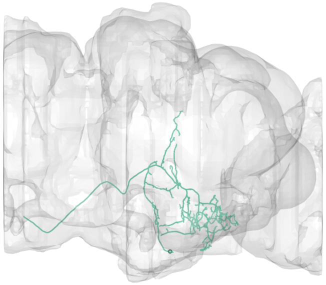

#54
bodyid =
1281812747

FOV = 58%
overlap of input LPTs' FOV = 38%
optimal rotation axis = [6°,−125°]
$\alpha_{R0} = 60°$
optimal translation axis = [−46°,141°]
$\alpha_{T0} = 45°$
RTSA = 15°

input LPT neuons:
LPT21_R
LPT30_R
LPT48_R_vCal3
LPT42_L_Nod4
LPT48_L_vCal3
LPT46_L_vCal1
LPT47_L_vCal2

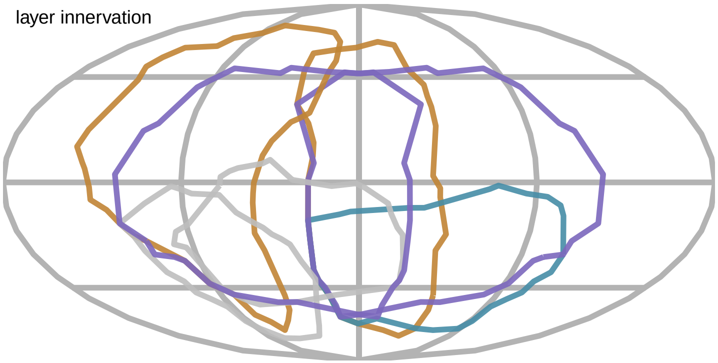


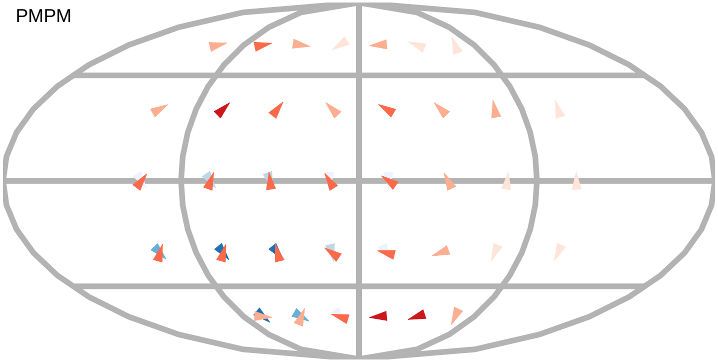


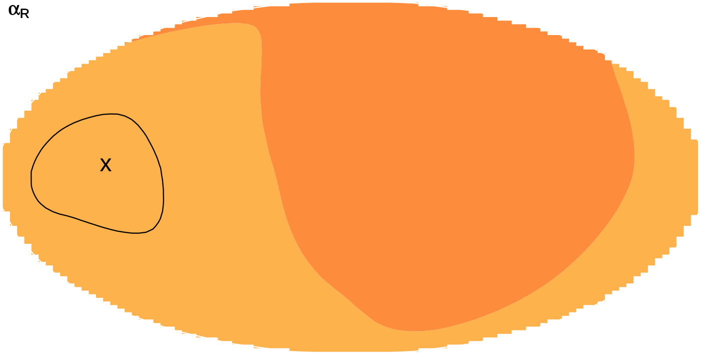


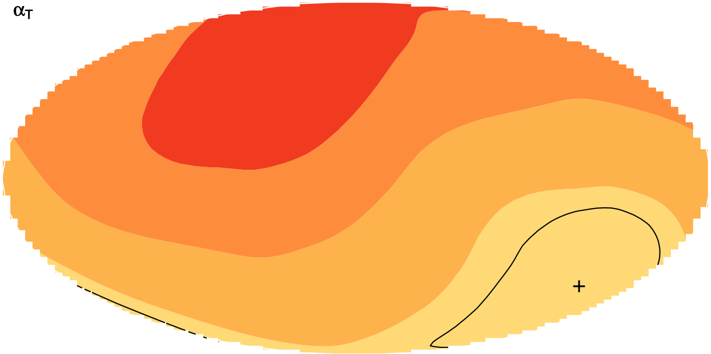

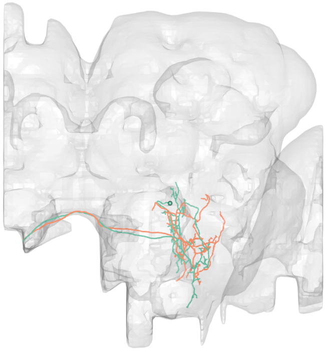

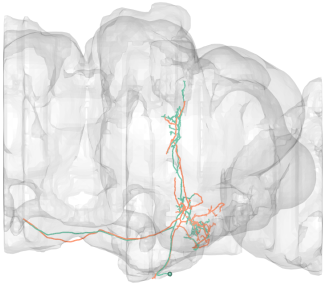

#55
bodyid =
1758224986
1713236401

FOV = 58%
overlap of input LPTs' FOV = 33%
optimal rotation axis = [−52°,42°]
$\alpha_{R0} = 76°$
optimal translation axis = [2°,158°]
$\alpha_{T0} = 29°$
RTSA = 47°

input LPT neuons:
LPT21_R
LPT41_R_Nod3
LPT41_L_Nod3
LPT42_L_Nod4
LPT40_R_Nod2

PMPM

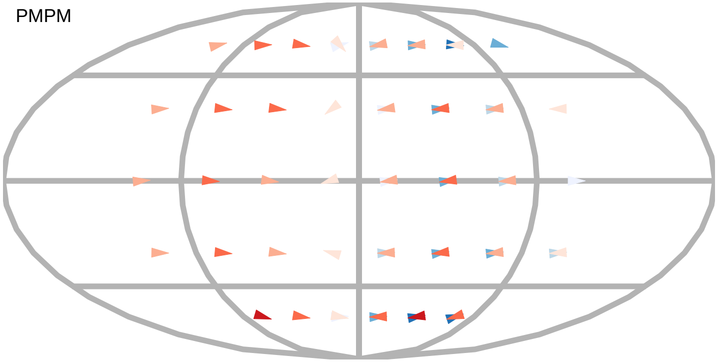

$\alpha_R$

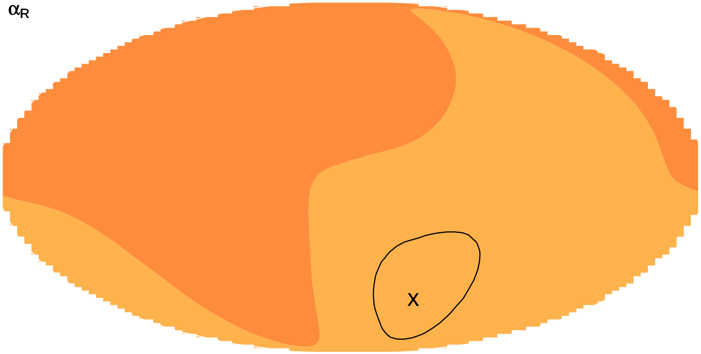


$\alpha_T$

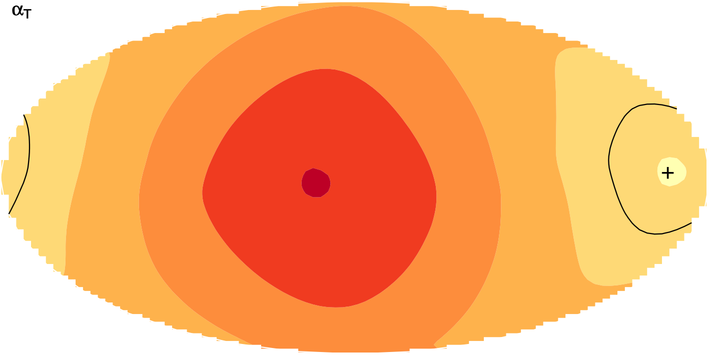


layer innervation

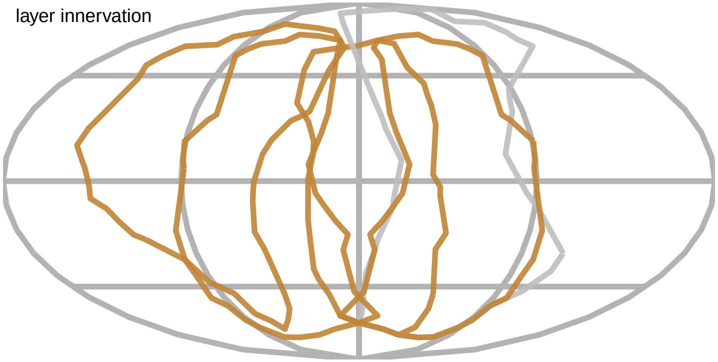

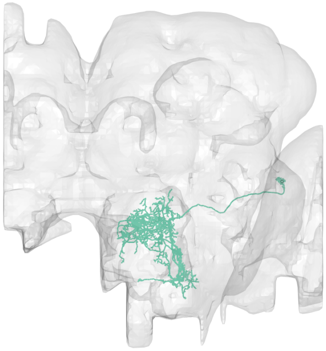

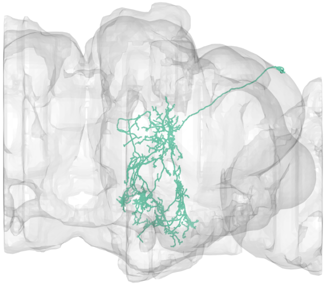

#56
bodyid =
1695145055

FOV = 60%
overlap of input LPTs' FOV = 42%
optimal rotation axis = [−52°,42°]
$\alpha_{R0} = 76°$
optimal translation axis = [2°,158°]
$\alpha_{T0} = 29°$
RTSA = 47°

input LPT neuons:
LPT21_R
LPT41_R_Nod3
LPT41_L_Nod3
LPT42_L_Nod4
LPT40_R_Nod2
LPT40_L_Nod2

layer innervation

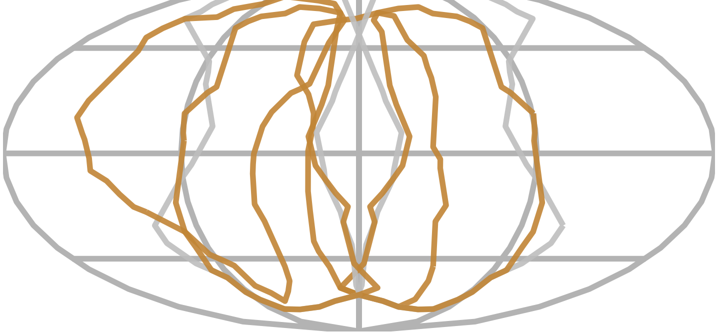

PMPM

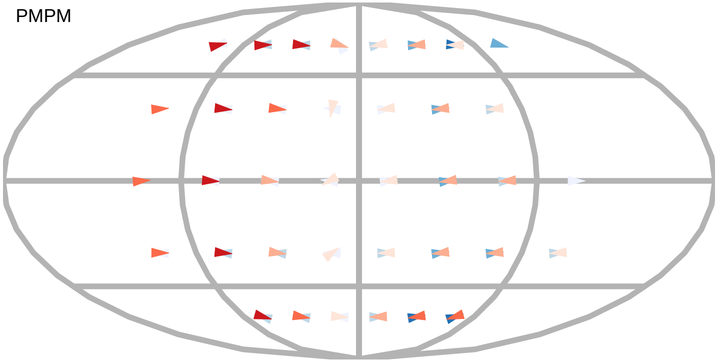

$\alpha_R$

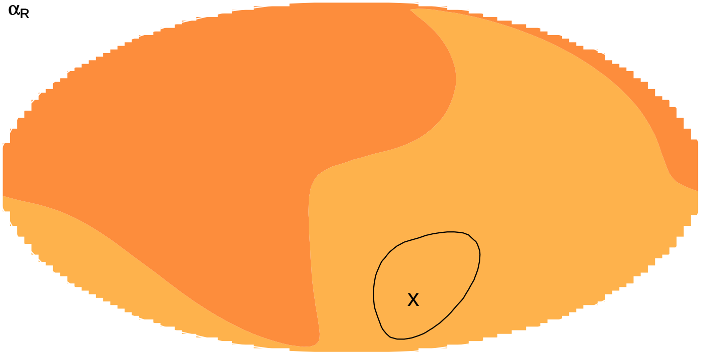


$\alpha_T$

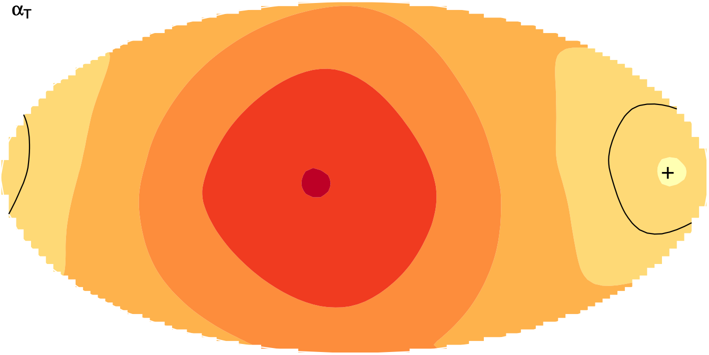

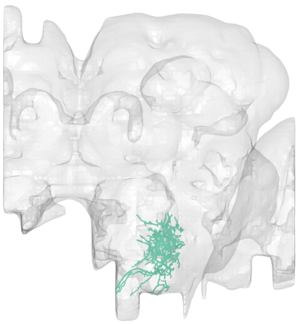

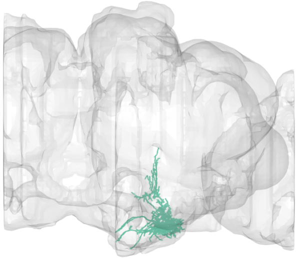

#57
bodyid =
1932497911

FOV = 62%
overlap of input LPTs' FOV = 50%
optimal rotation axis = [86°,−57°]
$\alpha_{R0} = 9°$
optimal translation axis = [−6°,−78°]
$\alpha_{T0} = 38°$
RTSA = −29°

input LPT neuons:
LPT01_R_HSN
LPT02_R_HSE
LPT03_R_HSS
LPT04_R_HST
LPT31_R
LPT43_L_H2
LPT44_L_Nod5
LPT50_L

layer innervation

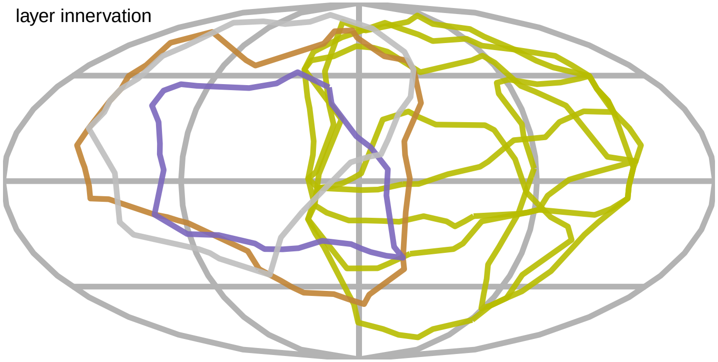

PMPM

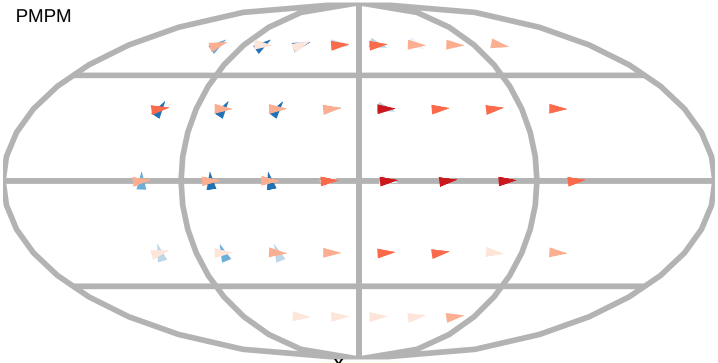

$\alpha_R$

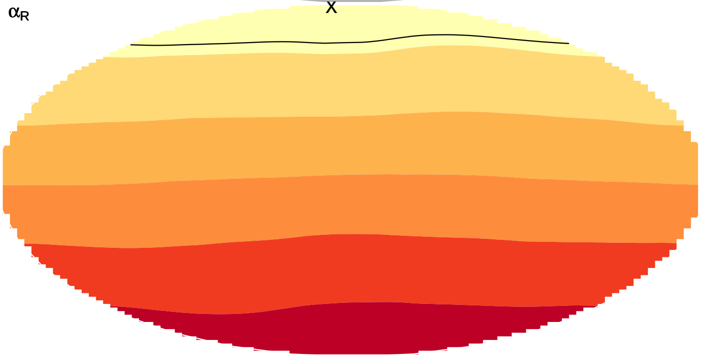


$\alpha_T$

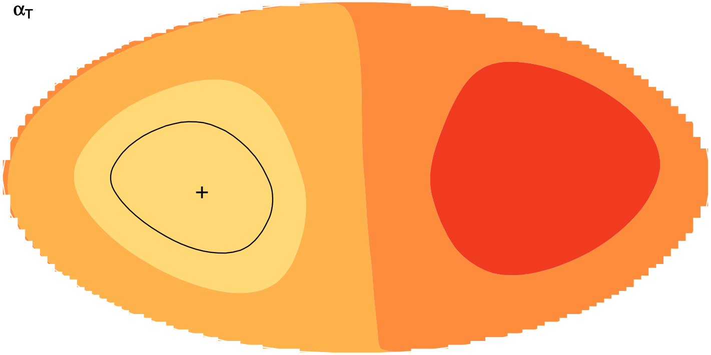

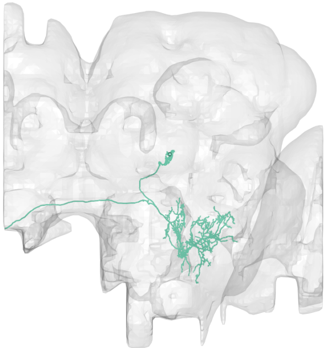

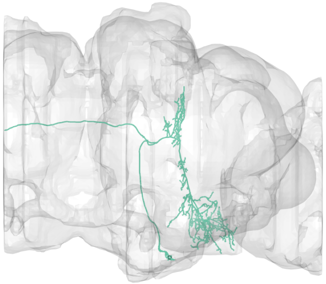

#58
bodyid =
1311125016

FOV = 62%
overlap of input LPTs' FOV = 27%
optimal rotation axis = [−18°,156°]
$\alpha_{R0} = 81°$
optimal translation axis = [4°,88°]
$\alpha_{T0} = 55°$
RTSA = 26°

input LPT neuons:
LPT21_R
LPT26_R
LPT27_R
LPT30_R
LPT31_R
LPT38_L_Nod1
LPT42_L_Nod4

layer innervation

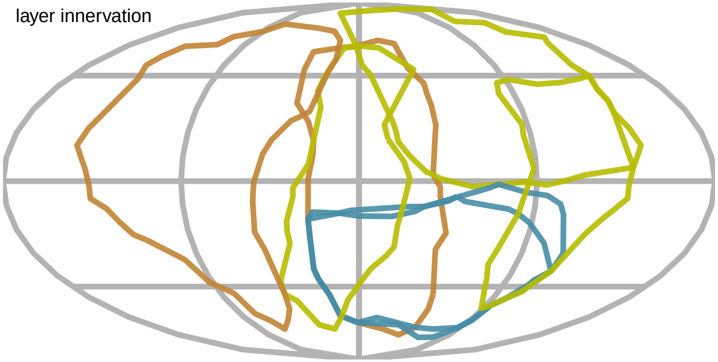

PMPM

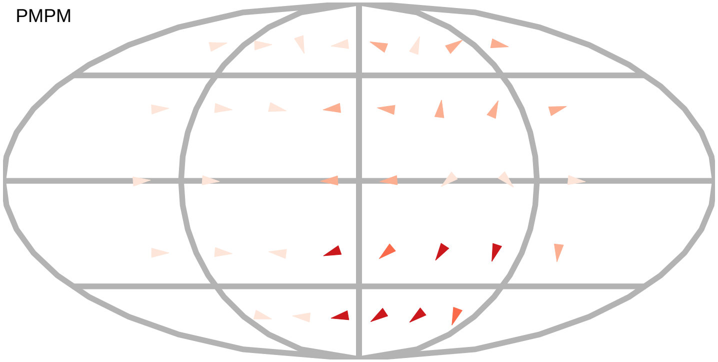

$\alpha_R$

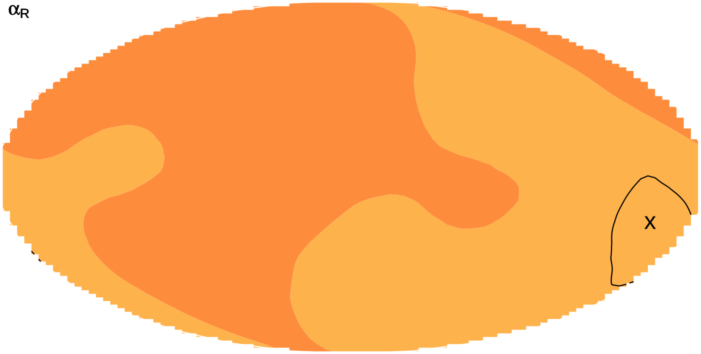


$\alpha_T$

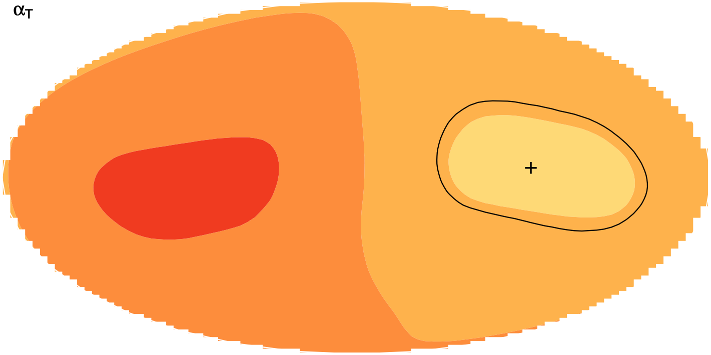

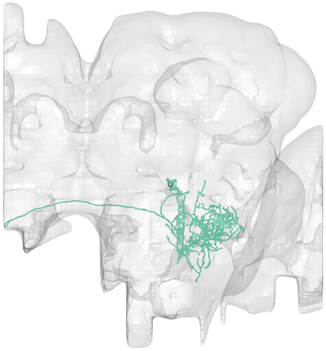

#59
bodyid =
1560734467

FOV = 66%
overlap of input LPTs' FOV = 36%
optimal rotation axis = [60°,100°]
$\alpha_{R0} = 81°$
optimal translation axis = [2°,70°]
$\alpha_{T0} = 61°$
RTSA = 20°

input LPT neuons:
LPT21_R
LPT26_R
LPT27_R
LPT30_R
LPT31_R
LPT38_L_Nod1
LPT42_L_Nod4
LPT46_R_vCal1
LPT46_L_vCal1
LPT47_L_vCal2

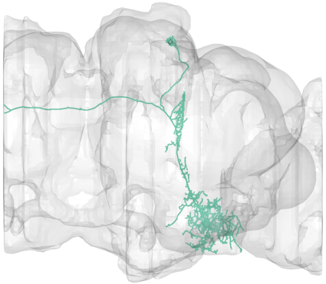

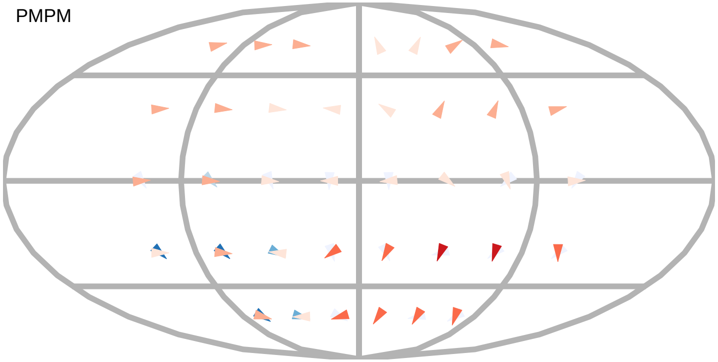


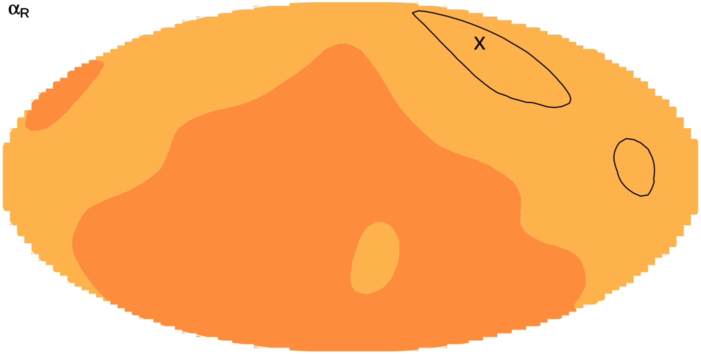


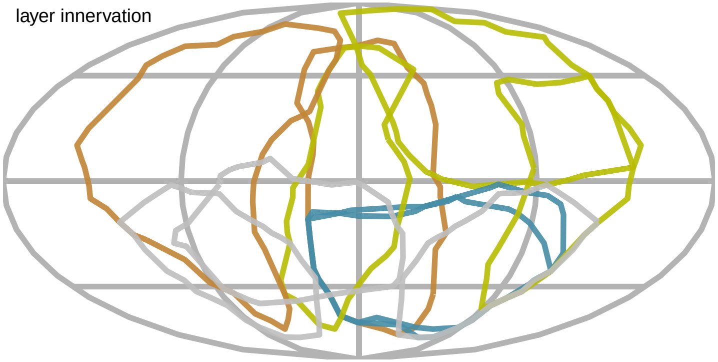


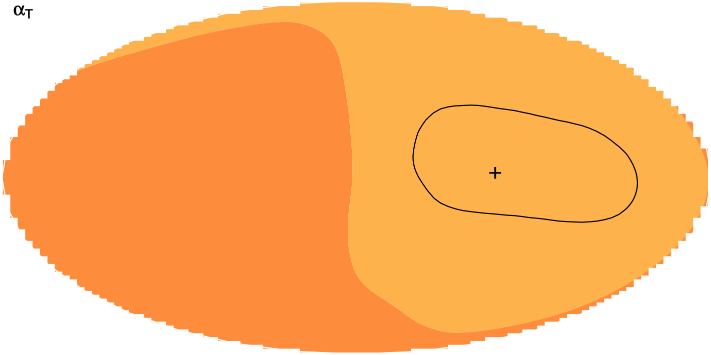

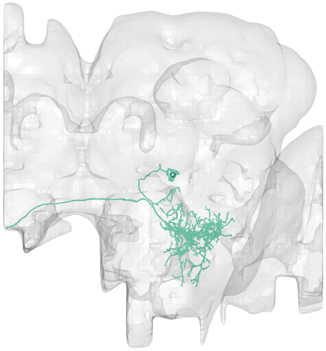

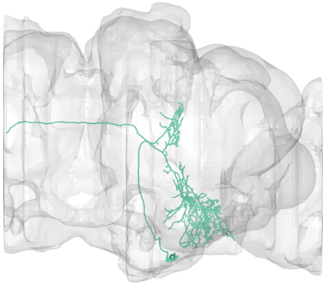

#60
bodyid =
1663767674

FOV = 67%
overlap of input LPTs' FOV = 46%
optimal rotation axis = [60°,100°]
$\alpha_{R0} = 84°$
optimal translation axis = [2°,136°]
$\alpha_{T0} = 59°$
RTSA = 25°

input LPT neuons:
LPT21_R
LPT26_R
LPT27_R
LPT30_R
LPT31_R
LPT41_R_Nod3
LPT38_L_Nod1
LPT41_L_Nod3
LPT42_L_Nod4

layer innervation

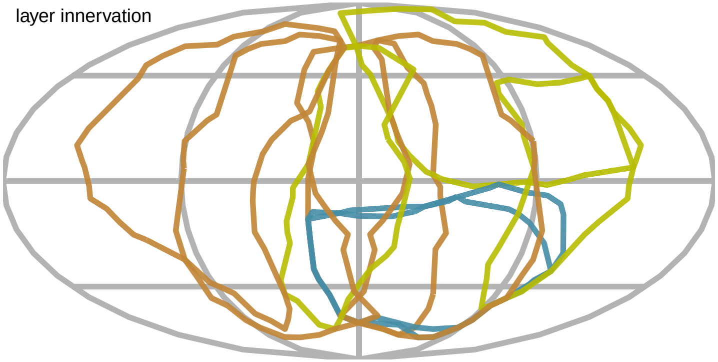

PMPM

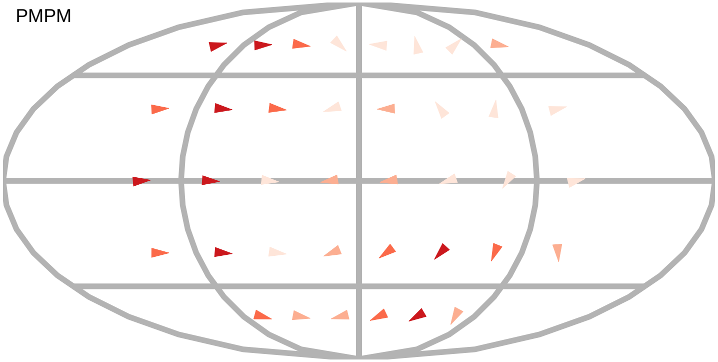

$\alpha_R$

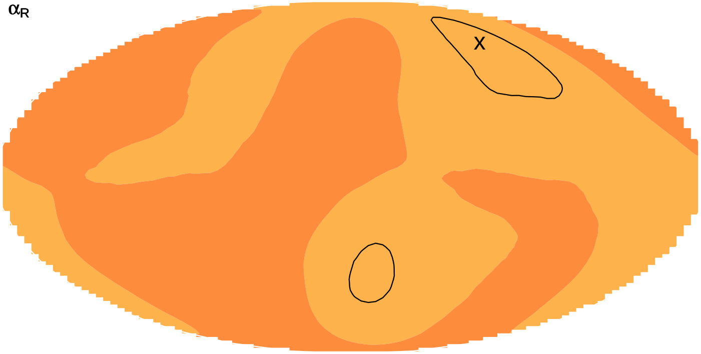


$\alpha_T$

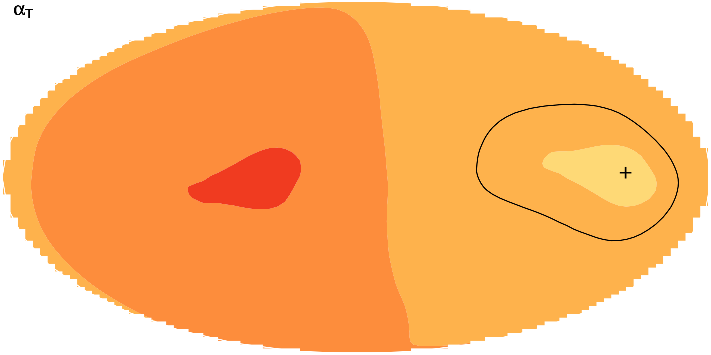

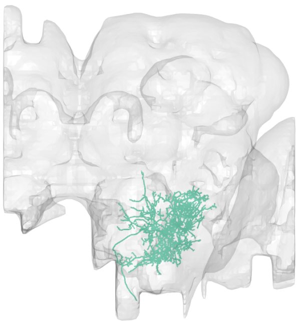

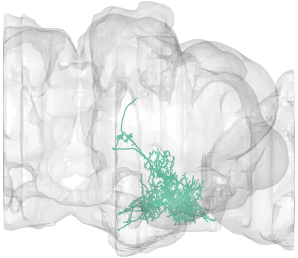

#61
bodyid =
1747625772

FOV = 67%
overlap of input LPTs' FOV = 40%
optimal rotation axis = [50°,96°]
$\alpha_{R0} = 81°$
optimal translation axis = [4°,74°]
$\alpha_{T0} = 61°$
RTSA = 20°

input LPT neuons:
LPT21_R
LPT26_R
LPT27_R
LPT30_R
LPT31_R
LPT38_L_Nod1
LPT42_L_Nod4
LPT49_L
LPT46_L_vCal1

PMPM

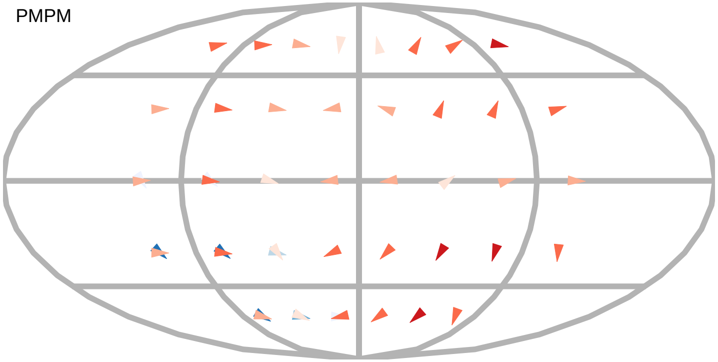

$\alpha_R$

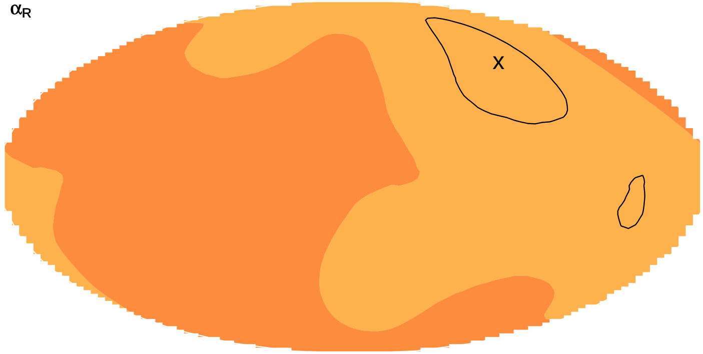

$\alpha_T$

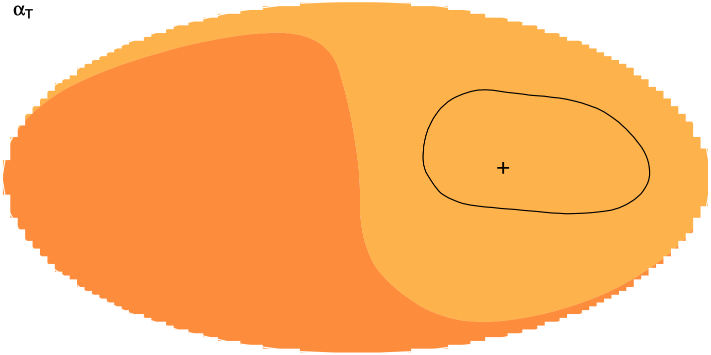

layer innervation

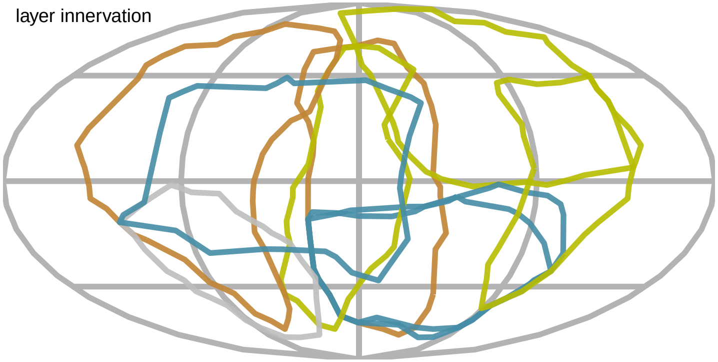

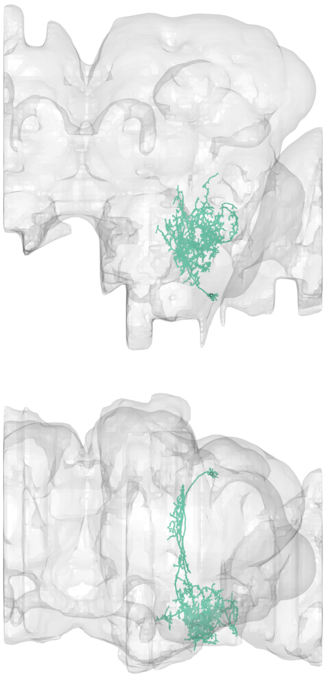

#62
bodyid =
5813067970

FOV = 67%
overlap of input LPTs' FOV = 51%
optimal rotation axis = [−14°,−159°]
$\alpha_{R0} = 69°$
optimal translation axis = [−12°,63°]
$\alpha_{T0} = 52°$
RTSA = 17°

input LPT neuons:
LPT21_R
LPT26_R
LPT27_R
LPT30_R
LPT31_R
LPT48_R_vCal3
LPT38_L_Nod1
LPT42_L_Nod4
LPT48_L_vCal3
LPT46_R_vCal1

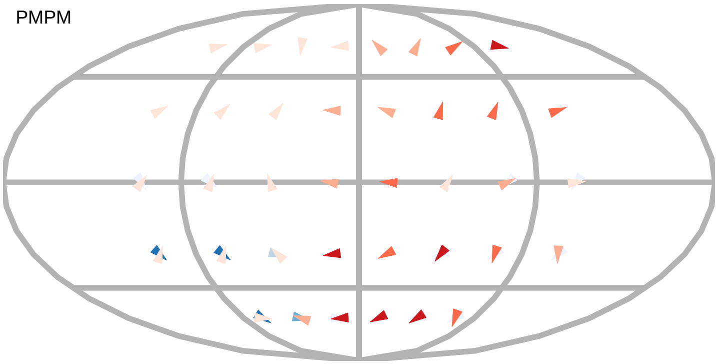


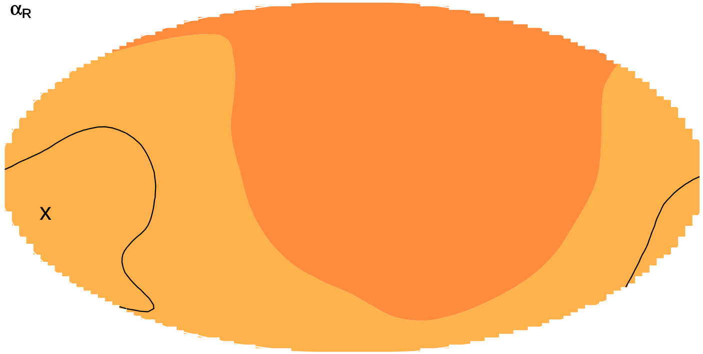


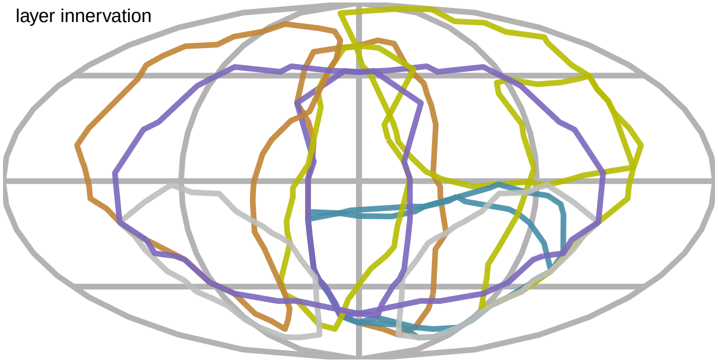


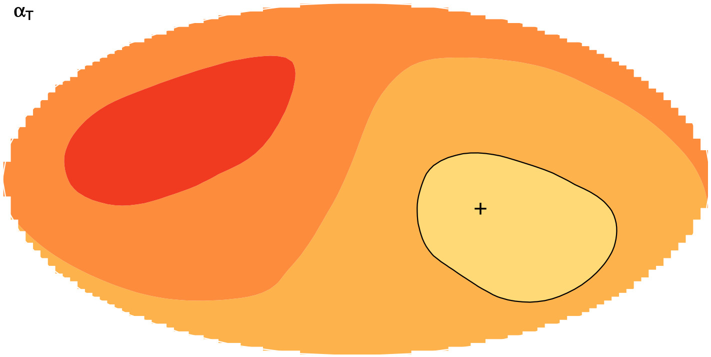
